# Proteomic Characterization of Isolated Arabidopsis Clathrin-Coated Vesicles Reveals Evolutionarily Conserved and Plant Specific Components

**DOI:** 10.1101/2021.09.16.460678

**Authors:** Dana A. Dahhan, Gregory D. Reynolds, Jessica J. Cárdenas, Dominique Eeckhout, Alexander Johnson, Klaas Yperman, Walter A. Kaufmann, Nou Vang, Xu Yan, Inhwan Hwang, Antje Heese, Geert De Jaeger, Jiri Friml, Daniel Van Damme, Jianwei Pan, Sebastian Y. Bednarek

## Abstract

In eukaryotes, clathrin-coated vesicles (CCVs) facilitate the internalization of material from the cell surface as well as the movement of cargo in post-Golgi trafficking pathways. This diversity of functions is partially provided by multiple monomeric and multimeric clathrin adaptor complexes that provide compartment and cargo selectivity. The adaptor-protein AP-1 complex operates as part of the secretory pathway at the *trans*-Golgi network, while the AP-2 complex and the TPLATE complex (TPC) jointly operate at the plasma membrane to execute clathrin-mediated endocytosis. Key to our further understanding of clathrin-mediated trafficking in plants will be the comprehensive identification and characterization of the network of evolutionarily conserved and plant-specific core and accessory machinery involved in the formation and targeting of CCVs. To facilitate these studies, we have analyzed the proteome of enriched *trans*-Golgi network/early endosome-derived and endocytic CCVs isolated from dividing and expanding suspension-cultured Arabidopsis cells. Tandem mass spectrometry analysis results were validated by differential chemical labeling experiments to identify proteins co-enriching with CCVs. Proteins enriched in CCVs included previously characterized CCV components and cargos such as the vacuolar sorting receptors in addition to conserved and plant-specific components whose function in clathrin-mediated trafficking has not been previously defined. Notably, in addition to AP-1 and AP-2, all subunits of the AP-4 complex, but not AP-3 or AP-5, were found to be in high abundance in the CCV proteome. The association of AP-4 with suspension-cultured Arabidopsis CCVs is further supported via additional biochemical data.

## Introduction

Vesicle trafficking is critical for the exchange of materials between the various biochemically and functionally distinct compartments of the biosynthetic/secretory and endocytic pathways. In particular, the trafficking of vacuolar proteins and the polar localization of plasma membrane proteins is critical for nutrient uptake, pathogen response, and organismal homeostasis.

Fundamental to the process of vesicle trafficking is the assembly of cytosolic coat protein complexes which cluster cargo and generate the membrane curvature necessary for a budding vesicle (Bonifacino and Glick, 2004; Robinson, 2015). The distinctive geometric lattice that surrounds clathrin-coated vesicles (CCVs) is composed of clathrin triskelia comprised of clathrin heavy chain (CHC) and clathrin light chain (CLC) subunits. Since their initial discovery in metazoans and plants (Gray, 1961; Roth and Porter, 1964; Bonnett and Newcomb, 1966), CCVs have been demonstrated to function in endocytosis (Robinson, 2015; Reynolds et al., 2018) and post-Golgi trafficking (Orci et al., 1985; Hirst et al., 2012). The highly choreographed multi-step process of clathrin-mediated endocytosis (CME) requires clathrin and a large number of endocytic accessory proteins (EAPs) that function at specific sites on the plasma membrane (PM) (i. e. clathrin-coated pits) to mediate cargo recruitment, membrane invagination, scission/severing of the nascent clathrin-coated vesicle from the PM and their uncoating prior to fusion with endosomes (Kaksonen and Roux, 2018; Wu and Wu, 2021; Taylor et al. 2011). Our current understanding of the complex network of proteins and lipids required for post-Golgi trafficking and endocytosis in plants largely comes from yeast and mammalian systems and the use of various biochemical, proteomic, genetic, and advanced quantitative live-cell imaging approaches. By comparison, mechanistic insight into clathrin-dependent membrane trafficking in plants remains limited.

CCVs facilitate multiple intracellular trafficking pathways via the action of adaptor protein complexes which are tasked with pathway-specific cargo recognition and clathrin recruitment at the PM and endomembrane compartments. The first clathrin adaptors to be identified were the assembly polypeptide (AP)-2 and AP-1 heterotetrameric complexes (Pearse and Robinson, 1984; Keen, 1987) that interact with protein cargo and specific phospholipids residing at the plasma membrane and the *trans*-Golgi Network (TGN), respectively (Robinson, 2015). Work has demonstrated that these functions are largely conserved in plants including the central role of AP-2 in endocytosis (Bashline et al., 2013; Di Rubbo et al., 2013; Fan et al., 2013; Kim et al., 2013) and of AP-1 in trafficking at TGN / early endosomes (TGN/EE) (Song et al., 2006; Park et al., 2013; Teh et al., 2013). The latter’s function is somewhat more complex as the plant TGN/EE appears to be comprised of numerous distinct sub-compartments with varying morphologies and functions in contrast to its metazoan counterpart (Dettmer et al., 2006; Viotti et al., 2010; Kang et al., 2011; Rosquete et al., 2018; Shimizu, et al., 2021; Heinze, et al. 2020). Moreover, an increasing number of studies have demonstrated that these core CCV trafficking proteins have evolved to accommodate processes critical for plant growth and development against the differing biophysical properties of plant cells including cell wall formation, cytokinesis, and pathogen response (Ekanayake et al., 2019; McMichael and Bednarek, 2013; Zhang et al., 2015; Gu et al., 2017).

In addition to AP-2 and AP-1, three additional AP complexes, AP-3 (Dell’Angelica et al., 1997; Simpson et al., 1997; Zwiewka et al., 2011), AP-4 (Dell’Angelica et al., 1999; Hirst et al., 1999), and AP-5 (Hirst et al., 2013) mediate post-Golgi endomembrane trafficking in multicellular eukaryotes. However, relative to AP-1 and AP-2, their interaction with the clathrin machinery in mammalian and other systems is less defined. Proteomic analyses of mammalian CCVs suggest that of the five evolutionarily conserved adaptor complexes, only AP-1 and AP-2 were found to be associated with the clathrin coat (Borner, et al. 2012, Blondeau, et al. PNAS 2004). Recent super-resolution microscopy of plant cells showed a lack of colocalization between AP4 and clathrin (Shimizu et al., 2021). Lastly, the association of AP-5 with clathrin metazoan and plants remains undetermined (Sanger et al., 2019).

Plants have shown evolutionary divergence in their key EAPs compared to other model systems including, for example, the TPLATE complex (TPC), which likely evolved in early eukaryotes but is notably absent from yeast and metazoans and which is essential for plant clathrin-mediated endocytosis (Gadeyne et al., 2014; Hirst, et al., 2014; Wang, et al., 2021; Bashline, et al., 2015). Similar to AP-2, TPC functions exclusively at the PM and acts as a central interaction hub required for the formation of endocytic CCVs (Gadeyne, et al., 2014; Wang, et al. 2016; Zhang, et al., 2015; Yperman, et al., 2021).

Additional conserved accessory proteins have been demonstrated to be required for CCV maturation including monomeric cargo adaptor families such as the ENTH/ANTH/VHS domain containing proteins which aid in cargo recognition and membrane deformation (Zouhar and Sauer, 2014; Fujimoto, 2020). In addition, CCV formation requires members of Arabidopsis dynamin related protein families DRP1s and DRP2s, which are necessary for scission of the budding vesicle from the plasma membrane (Bednarek and Backues, 2010). DRP1 and DRP2 protein assembly into clathrin-coated pits (CCPs) is critical for efficient CME and cell plate biogenesis but the precise mechanistic details of their function(s) in these processes are not fully established (Fujimoto, et al., 2010; Narasimhan, et al., 2020; Ekanayake, et al., 2021; Mravec, et al., 2011; Backues, et al. 2010; Konopka, et al., 2008). Various plant uncoating factors have been identified, and the mechanistic details are under investigation (Robinson, 2015; Adamowski et al., 2018). Trafficking factors including Rab and ARF-related small GTPases (Jurgens et al., 2015; Lipatova et al., 2015; Takemoto et al., 2018), vesicle tethering complexes (Jurgens et al., 2015; Takemoto et al., 2018), and soluble N-ethylmaleimide-sensitive factor adaptor protein receptors (SNARE) proteins (Yun and Kwon, 2017) add complexity to the regulation of clathrin-mediated trafficking pathways.

Defining the protein complement of post-Golgi and endocytic CCVs will further enhance our understanding of the evolutionarily conserved and plant-specific core and accessory machinery involved in the formation and targeting of plant clathrin-coated vesicles. Organelle proteomics is a useful tool for such studies as it can provide a global view of the protein content of a particular organelle thereby placing known and un- annotated gene products in a functional context. Several studies have employed mass spectrometry (MS) to answer questions of mammalian CCV content including analyses of rat brain CCVs to examine their role in synaptic vesicle recycling (Takamori et al., 2006; Blondeau et al., 2004) and of those isolated from HeLa cells (Borner et al., 2006; Borner et al., 2012). In addition to the identification of previously unknown CCV components, the relative abundance of proteins across CCVs isolated from rat brain and liver has been examined to interrogate pathway-specific proteins such as AP-1 and AP-2 subunits (functioning in secretion and endocytosis, respectively) and the proportion of CCVs involved in post-Golgi and endocytic trafficking in different cell types (Girard et al., 2005).

In plants, subcellular fractionation coupled with tandem MS has also been utilized to elucidate the protein content of endomembrane compartments including Arabidopsis Golgi cisternae (Parsons et al., 2012; Okekeogbu et al., 2019; Parsons et al., 2019), vacuoles (Carter et al., 2004), and syntaxin of plants 61 (SYP61)-positive TGN compartments (Drakakaki et al., 2012). Here, we report the proteomic assessment of CCVs isolated from undifferentiated Arabidopsis suspension cultured cells using tandem mass spectrometry and quantitative immunoblotting to elucidate the ensemble of proteins underlying clathrin-mediated trafficking in plants and to better understand the similarities and differences among trafficking protein pathways within eukaryotes.

## Results/Discussion

### CCV Isolation

In plants, CCVs mediate trafficking of secretory and endocytic cargo necessary for cell expansion, cytokinesis, nutrient uptake, and pathogen immunity pathways underlying growth, morphogenesis, and defense in a variety of developmentally-distinct cell types (Reynolds et al., 2018; Ekanayake, et al., 2019). Undifferentiated suspension-cultured plant cells offer, however, several benefits as a biological sample source for proteomic analysis of plant CCVs as they: 1) display high levels of division and expansion, processes that require a large flux of vesicular trafficking; 2) are easily scaled to provide large amounts of material; and 3) when grown under constant conditions, provide a population of cells uniform in cell type and development between biological replicates. With these features in mind, CCVs used for proteomic analysis in this study were isolated from 3-4 day-old T87 suspension-cultured Arabidopsis cells (Axelos et al., 1992). In addition, the availability of the Arabidopsis genome sequence and *in silico* proteome (Arabidopsis Genome, 2000; Berardini et al., 2015) as well as the T87 transcriptome datasets (Stolc et al., 2005) facilitate the assessment of the enrichment or depletion of proteins of interest in the CCV proteome.

For proteomic analyses, plant CCVs were isolated under pH conditions that inhibit clathrin cage disassembly using a fractionation scheme that includes differential, rate-zonal centrifugation, and a final equilibrium deuterium/Ficoll gradient as described previously (Reynolds et al., 2014). Prior to mass spectrometry analysis, the composition and quality of CCV preparations were assessed by morphological analysis using transmission electron microscopy (TEM) and immunoblotting for protein markers of the secretory and endocytic pathways. TEM analysis (Figure 1A) of the enriched clathrin-coated vesicle samples revealed that 65% and 48% of the vesicles in two independent replicates were coated and displayed the characteristic geometry of clathrin coats with a diameter of 70 nm. Scanning transmission electron micrographs of enriched CCVs at higher resolution show striking symmetry of the coat (Figures 1B and 1C).

**Figure 1.**
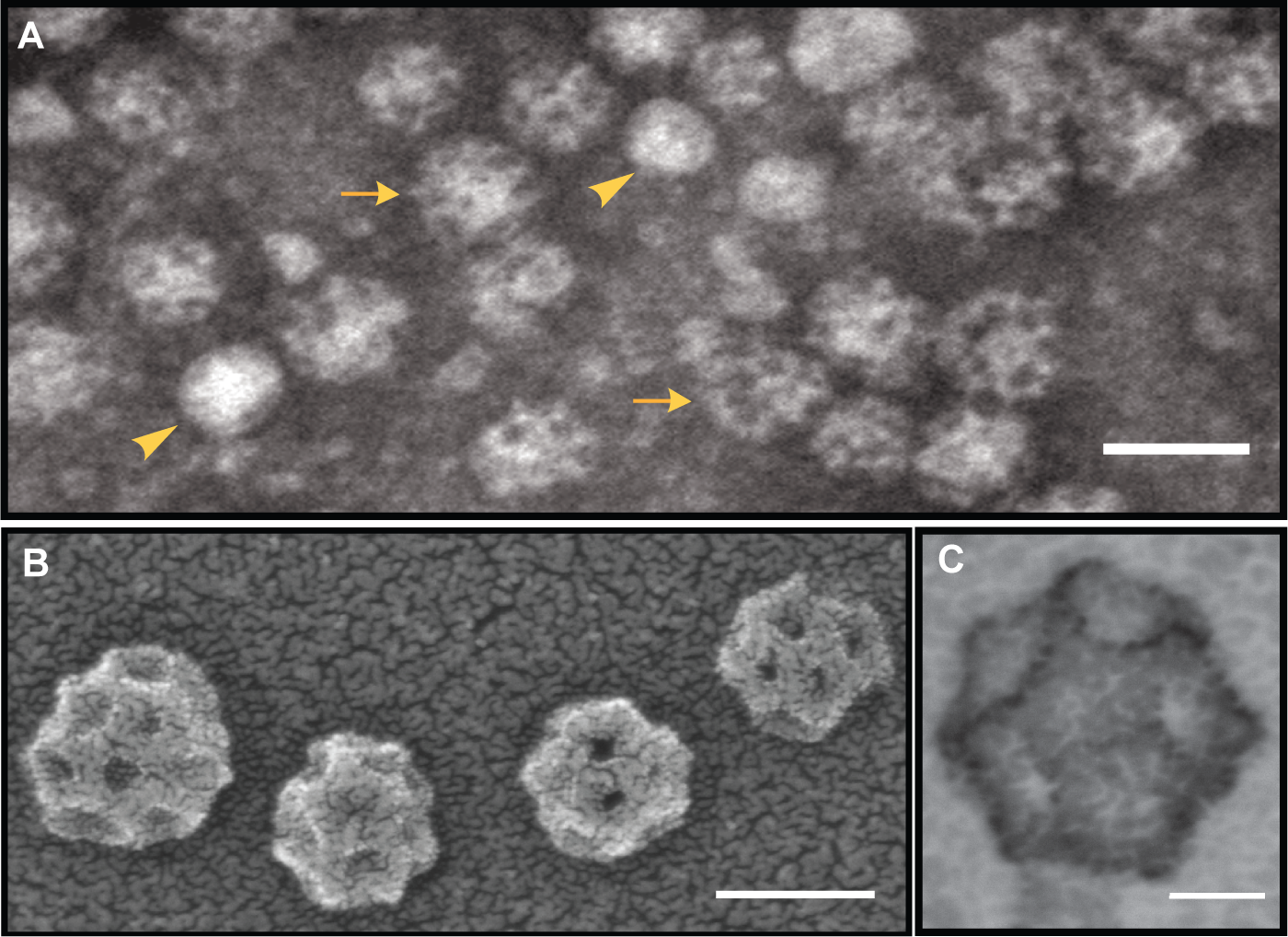
Electron microscopy of purified plant clathrin-coated vesicles. (A) Negative stain transmission electron micrographs of a typical CCV prepa- ration. (B-C) Positive stain scanning transmission electron micrographs of clathrin-coated vesicles. Scale bars (clockwise, starting top): 100 nm, 20 nm, and 50 nm. Uncoated vesicles are indicated by arrowheads and coated vesicles by arrows.

### CCV MS/MS Sample Preparation & Analysis

To establish a comprehensive understanding of the protein composition of plant CCVs, two independent, parallel proteomic workflows were performed on enriched CCV protein samples (Supplemental Figure 1). In the first methodology, CCVs isolated from four independent biological replicates were resolved via one-dimensional SDS-PAGE before in-gel digestion of proteins by trypsin and subjection to LC/MS-MS. To account for differences in protein sampling related to 1D SDS-PAGE sample fractionation and the use of detergents, we used a second methodology in which three independent CCV preparations were denatured in a urea buffer prior to trypsin digestion in-solution and separation of recovered peptides by LC/MS-MS.

Coomassie staining of a representative enriched CCV fraction separated by 1D SDS-PAGE is shown in Figure 2A. Gel slices 2, 6, and 8 contain bands of high protein abundance (intense Coomassie staining) which migrate at rates corresponding to the molecular weights of the clathrin coat proteins. To confirm the identity of the specific heavy or light chain clathrin isoforms in each polyacrylamide gel slice, we plotted the unique spectral counts as a percentage of unique spectral counts for that isoform across the entire SDS-PAGE gel and showed that the protein abundances in gel slices 2, 6, and 8 in Figure 2A corresponded to the presence of clathrin subunits (Figure 2B). Arabidopsis encodes two CHC isoforms, CHC1 and CHC2, of approximately the same mass (193kD) which share >90% sequence identity (Kitakura et al., 2011) and are transcribed at comparable levels in T87 suspension cultured cells (Stolc et al., 2005). In contrast to CHC, the three Arabidopsis CLC isoforms share only ∼55% sequence identity (Wang et al., 2013) and vary in predicted mass (37.2, 28.8, and 29.1 kD, CLC1-3, respectively). The spectral count and SDS-PAGE data confirmed that clathrin heavy chain contributes towards the high protein abundance of gel slice 2 (predicted molecular weight of CHC1 and CHC2, 193 kDa), and that clathrin light chain contribute towards the protein abundance of gel slices 6 and 8 (predicted molecular weight of CLC1, 37 kDa; of CLC2 and CLC3, 29 kDa). The identities of the other CCV-associated proteins in Figure 2A with abundances below the limits of detection by Coomassie staining were determined by proteomic and immunoblot analyses addressed below.

**Figure 2.**
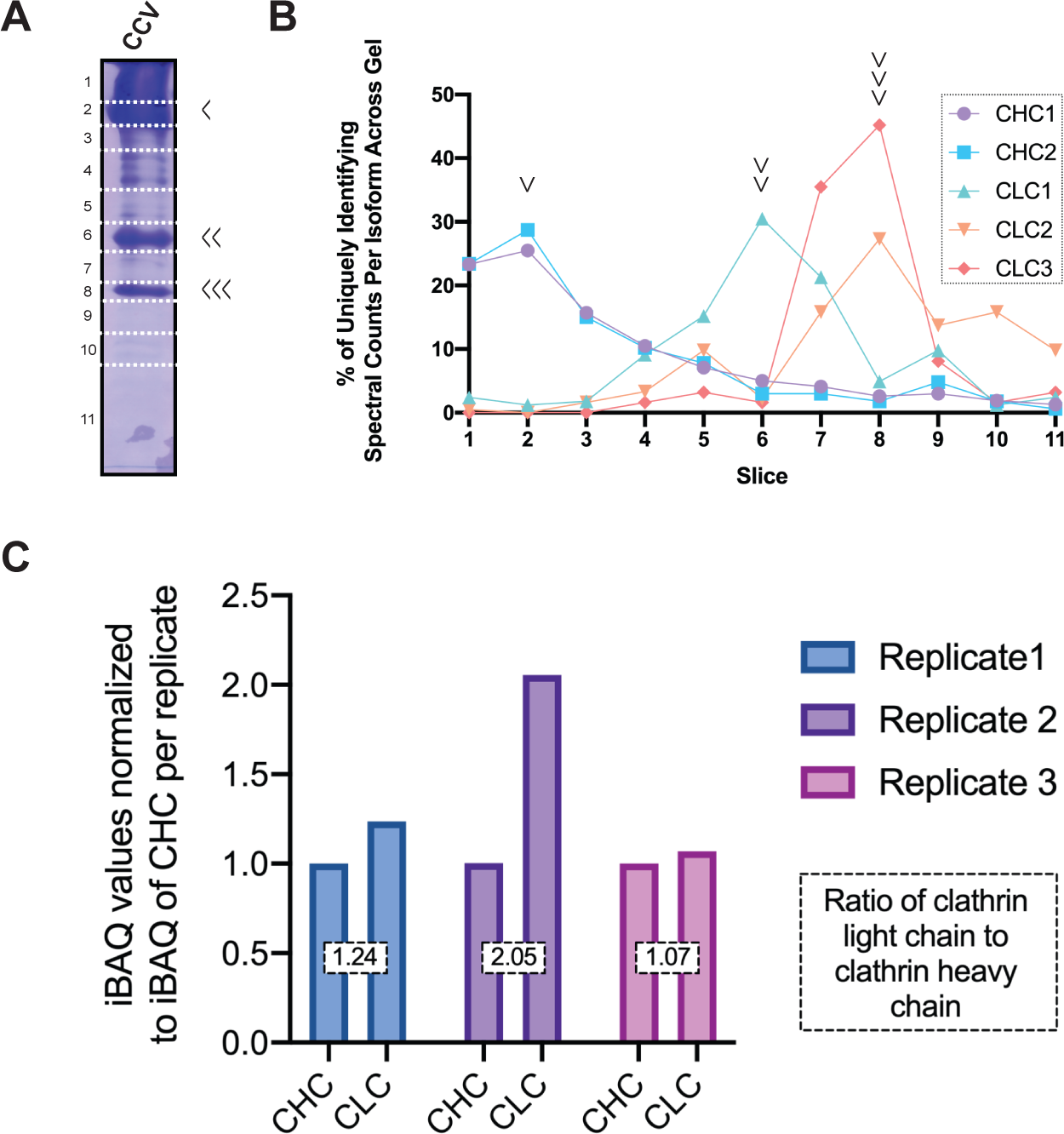
Distinction of clathrin isoforms by mass spectrometry and stoichiometry of clathrin subunits. (A) Representative Coomassie stained SDS-PAGE analysis of 1 mg of clathrin-coated vesicles purified by differential centrifugation from Arabidopsis T87 suspension cultured cells. After separation by SDS-PAGE, gels were sliced along the indicated dotted lines before in-gel trypsin digest and analysis of each fragment by LC/MS-MS. The identity of abundant CCV associated proteins marked by arrowheads to the right of the gel were found to be clathrin heavy (single arrow head) and light chain isoforms (CLC1, double arrowhead; CLC2 and CLC3, triple arrowheads) based on mass spectrometry and analysis in Figure 2B. (B) Spectral counts attributed to a specific clathrin heavy or light chain isoform from the unlabeled LC/MS-MS analysis of the CCV purifications separated by 1D SDS-PAGE and visualized by Coomassie staining in Figure 2A. Peaks in the line graphs were used to assign particular clathrin heavy and light chain isoforms to the indicated bands in Figure 2A. (C) Ratio of clathrin heavy chain to clathrin light chain subunits for three independent CCV purifications analyzed by LC/MS-MS without separation by one dimensional SDS-PAGE. The iBAQ value plotted on the y-axis is derived from the sums of the iBAQ intensity values assigned to clathrin light or heavy chain isoforms from the indicated replicate normalized to the sum of the iBAQ intensity value assigned to clathrin heavy chain for each replicate. The boxed number overlaid on the columns is the ratio of CLC:CHC for each replicate.

To define the CCV proteome, protein or protein group assignments were considered true if unique peptides denoting the protein or protein group were found in at least two biological replicates based on a 1% false discovery rate protein threshold and unique peptide assignments at 95% confidence (first methodology; Materials & Methods and Supplemental Figure 1B) or at a 1% false discovery rate protein and peptide threshold (second methodology; see Materials & Methods and Supplemental Figure 1C). Processed spectra from the first proteomic analysis were matched against the Arabidopsis protein sequence database using the Mascot searching algorithm (Perkins et al., 1999), while in the second approach, MS/MS spectra were analyzed with the MaxQuant software package.

Following these criteria, protein assignments from the four independent CCV preparations analyzed by Method 1 (in which CCVs were separated by 1D SDS-PAGE) comprised a list of 3,548 proteins (Supplemental Dataset 1), the vast majority of which (∼72%) were in relatively low abundance (< 50 spectral counts across four replicates). Total spectral counts have previously been used as an approximation of overall protein abundance in biological samples for those proteins with a reasonably high (>10-20) total number of counts (Lundgren et al., 2010). Accordingly, we compared total counts over four biological replicates without normalization as an approximation of overall protein abundance within CCVs and focused our analysis on those proteins with the highest associated spectral count totals. The proteomic data from CCVs fractionated by 1D SDS-PAGE and analyzed by tandem MS is presented in Supplemental Datasets 1 and 4 and Figures 2A, 2B, 3, and 5. Protein assignments obtained from the second CCV methodology resulted in a list of 1,981 protein groups (Supplemental Dataset 2). Intensity based absolute quantitation (iBAQ) values were used to sort protein groups within this dataset, as this method of label-free quantification method has been previously judged to be a metric for protein abundances within biological samples and enables comparison of these abundances (Arike, et al. 2012; Nagaraj, et al. 2011; Schwanhausser, et al. 2011). The data from this second methodology is presented in Supplemental Datasets 2 and 4 and Figures 2C and 3.

**Figure 3.**
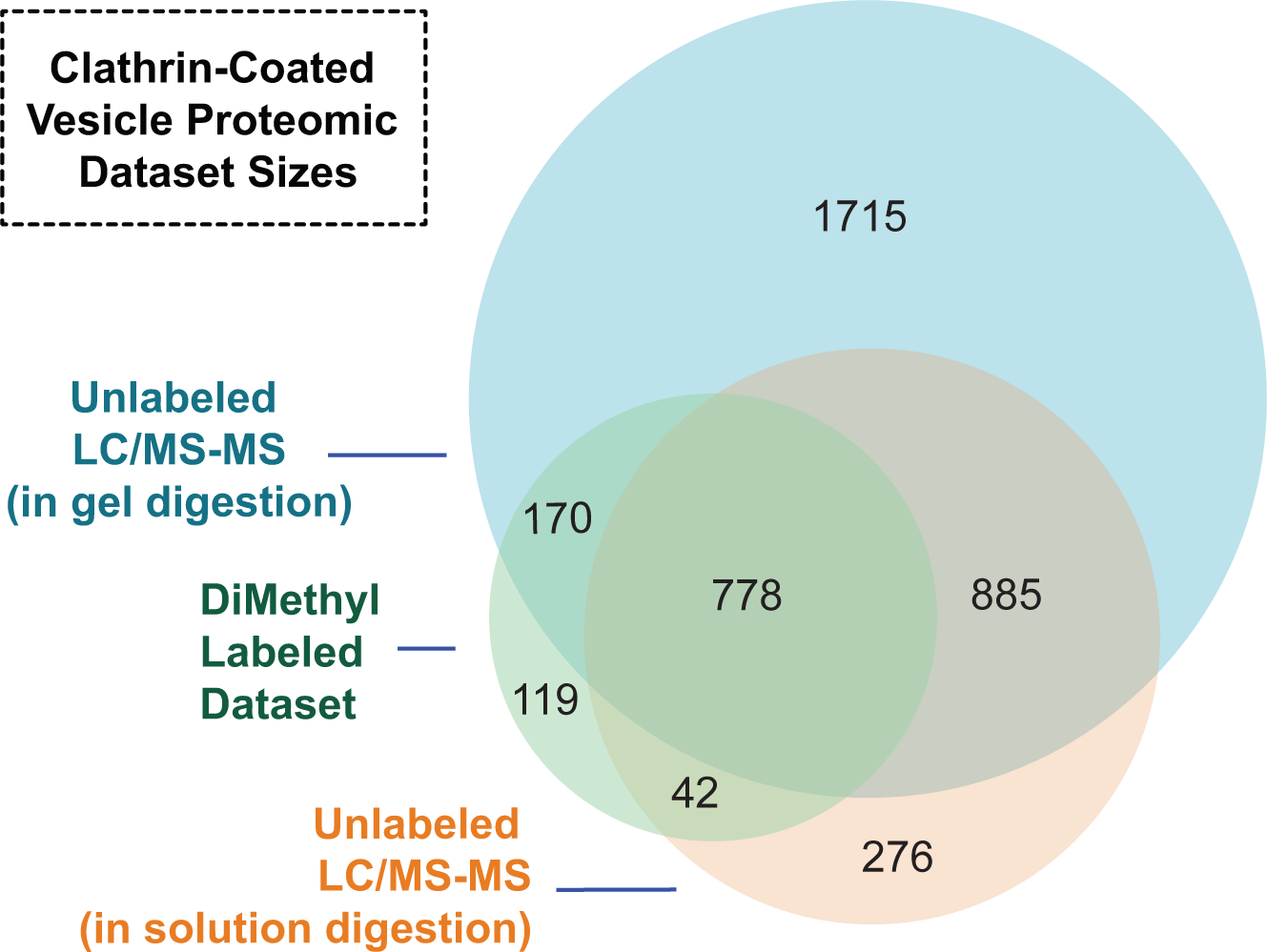
Sizes of and overlaps in proteomic datasets defining proteins associated with clathrin-coated vesicles purified from Arabidopsis cells. Four independent CCV preparations were separated by one-dimensional SDS-PAGE before LC/MS-MS. Proteins or proteins representative of protein groups were incorporated into the unla- beled LC/MS-MS dataset (blue) if total spectral counts for the protein/protein group were present in >/= 2 replicates (3,548 proteins). Three independent CCV preparations were not separated by SDS-PAGE but treated with urea before trypsin digest and subsequent LC/MS-MS. Protein groups were incorporated into in-solution digest, unlabeled LC/MS-MS dataset (orange) if they were identified in at least two replicates (1,981 protein groups); overlap with this dataset was determined using the first protein within the protein groups identified. Deuterium ficoll gradient load (DFGL) and CCV fractions from two independent preparations were reciprocally labeled with light and heavy formalde- hyde before separation with SDS-PAGE and subjection to LC/MS-MS. Proteins identified in at least one replicate as increased or decreased in abundance between CCV and DFGL fractions were included in the dimethyl labeling LC/MS-MS dataset (1,109 proteins; green). The sizes of circles and overlaps are proportional to the number of proteins or protein groups contained within.

Previous proteomic studies of CCVs purified from mammalian tissue have found that clathrin light and heavy chains exist in a stoichiometric 1:1 ratio, but also as non-stoichiometric ratios depending on the tissue and species source of the CCV sample (Borner, et al. 2012; Blondeau, et al. 2004; Girard, et al. 2005). We used iBAQ levels to analyze the abundances of clathrin heavy and light chain subunits and determine the ratio of subunits within the triskelion independently of immunoblotting based methods. We summed the iBAQ values for clathrin heavy chain subunits and for clathrin light chain subunits for each independent replicate and took the ratio between these values as the measure of CLC:CHC per replicate of 1.24, 2.05, and 1.07 (Figure 2C; Supplemental Dataset 2). These values generally support a 1:1 ratio but suggest that, in some CCV preparations from plant cells, clathrin light chain subunits are in excess.

The numbers of proteins comprising each proteomic dataset and shared between both sample preparations described above are illustrated in Figure 3. Cross-referencing the datasets derived from these independent, parallel workflows established an overlapping CCV proteome comprising 1,663 proteins.

### Assessment of Protein Enrichment and Depletion in CCV Fraction

To further refine the CCV proteome, we quantitatively compared the abundance of peptides in pre- and post- deuterium/Ficoll gradient samples, termed Deuterium Ficoll Gradient Load (DFGL) and CCV samples, respectively. To do so, we assessed the relative enrichment or depletion of proteins identified by mass spectrometry in an unbiased manner through a differential labeling strategy with stable isotope dimethyl moieties (Boersema et al., 2009). This methodology involves the reaction of peptide primary amines with either formaldehyde or deuterated formaldehyde to form methylated peptides, which results in identical peptides treated in this way differing by 4 Daltons. Accordingly, a quantitative ratio of an individual peptide’s abundance in the DFGL relative to the final enriched CCV preparation, as represented by spectral counts, can be derived. DFGL and CCV fractions from two independent biological replicates were separated by 1D SDS-PAGE prior to gel sectioning and in-gel digestion with trypsin. Peptides from both fractions were recovered and treated with dimethyl reagents as previously described (Boersema et al., 2009). In the second replicate, heavy (deuterated) and light labels applied to CCV and DFGL samples were swapped to control for potential discrepancies resulting in the identification of proteins which were enriched in (CCV:DFGL spectral count ratio ≥ 2.0) or depleted from (DFGL:CCV spectral count ratio ≥ 2.0) the purified clathrin-coated vesicles (Figure 3, Supplementary Dataset 3). Cross-referencing these datasets yielded a core set of 778 proteins in the enriched CCV purification sample as detected by all three proteomic workflows, 213 (27%) of which had a CCV:DFGL spectral count ratio ≥ 2.0, i.e. were enriched more than two-fold in the final preparatory step resulting in purified clathrin-coated vesicles (Figure 3, Supplemental Dataset 3).

In previous studies, immunoblotting has been used to validate differential centrifugation as a means to purify clathrin-coated vesicles from plant cells by confirming the depletion of markers of the endoplasmic reticulum (ER) and Golgi, as well as of other organelles including plastids, peroxisomes, and mitochondria, in addition to confirming the enrichment of CCV associated proteins (McMichael et al., 2013; Reynolds et al 2014). However, the fold enrichment or depletion of these proteins relative to their initial abundance in the lysate (S0.1) fraction has not been quantified. Here, we performed quantitative immunoblotting of equal amounts of protein from steps throughout the CCV purification process, including the DFGL and CCV fractions, to establish the fold enrichment and depletion of CCV mediated trafficking-associated and unassociated proteins, respectively, relative to the lysate (Figure 4). We then used the average fold enrichments between the DFGL and CCV fractions from three independent quantitative immunoblotting experiments (apart from immunoblotting for DRP2, where n = 2) to corroborate the values obtained by the dimethyl labeling proteomic workflow (Table 1).

**Figure 4.**
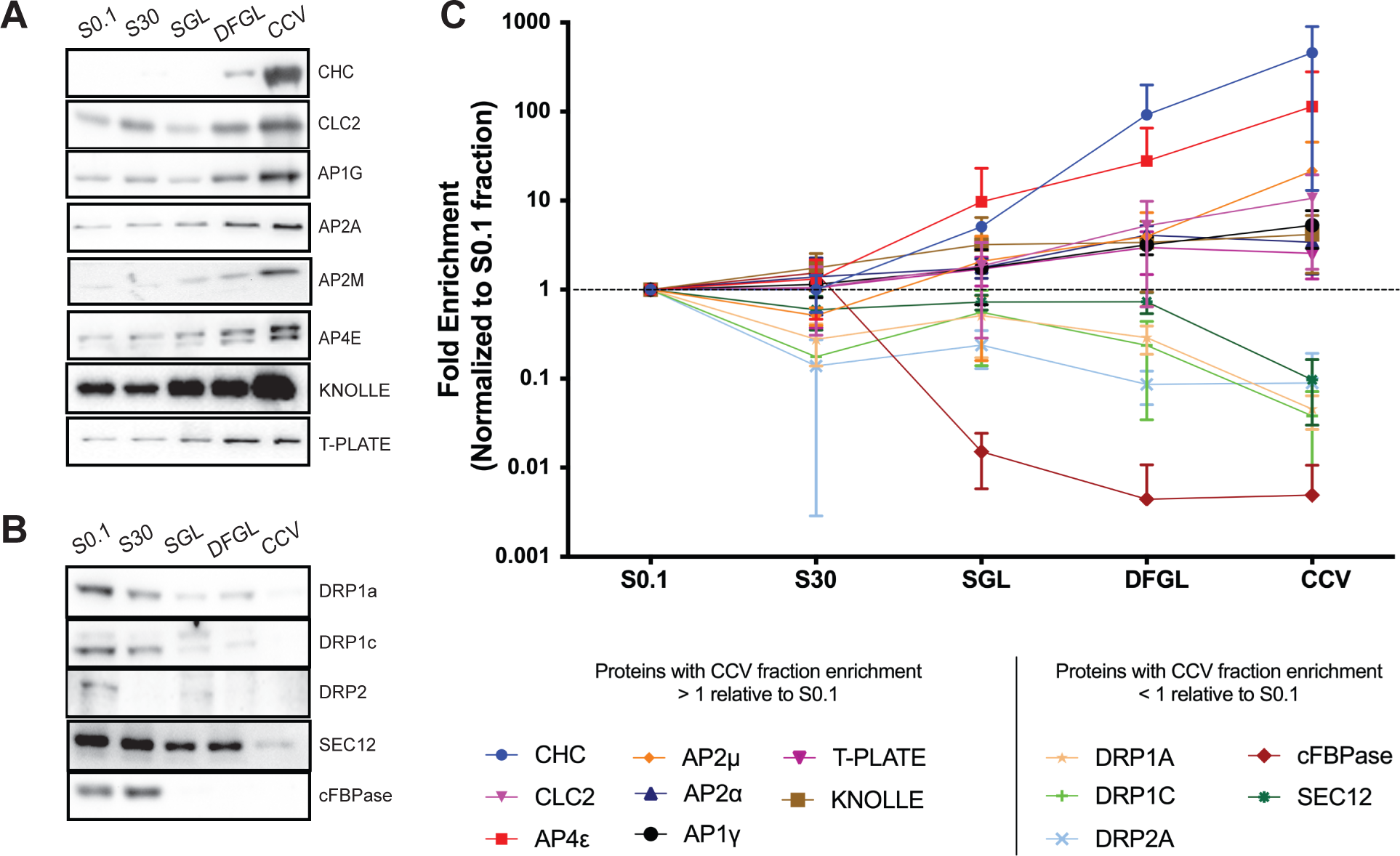
Stepwise enrichment and depletions of AP4 and trafficking and marker proteins throughout the CCV purification process. (A) Equal amounts of protein from S0.1 (lysate), S30, SGL, DFGL, and CCV fractions were immunoblotted with antibodies against known CCV associated proteins (CHC, CLC2, AP1G, AP2A, AP2M, and T-PLATE) as well as AP4E and the cell plate marker, KNOLLE. These proteins were found to be enriched in the final CCV fraction relative to the lysate (B) Equal amounts of protein from S0.1, S30, SGL, DFGL, and CCV fractions were immunoblotted with antibodies for proteins known to be transiently associated with the CCV formation process (DRP1c, DRP1a, and DRP2) as well as for organellar markers, cFBPase (cytosol) and SEC12 (endoplasmic reticulum). These proteins were found to be depleted in the final CCV fraction relative to the lysate. (C) Quantitation of the enrichment of proteins in A and B from three biological replicates (apart from DRP2, where n = 2). The mean signal intensity of each step relative to that of the mean signal intensity in the lysate for each protein was plotted on a logarithmic scale with error bars indicating standard deviation about the mean. Abbreviations used: S0.1, lysate; S30, 30,000 x g supernatant; SGL, sucrose step gradient load; DFGL, linear deuterium oxide/Ficoll gradient load; CCV, clathrin coated vesicle fraction.

**Table 1.** Enrichment and depletion profiles of CCV associated proteins and organellar markers. The ratio of protein levels in the CCV fraction relative to levels in the DFGL fraction (without normalizing to levels in the S0.1 fraction) are presented alongside standard deviation about the mean. For average ratios derived by immunoblotting, n = 3 biological replicates, except for DRP2 (n = 2). For dimethyl labeling values, a dash in the right-hand column indicates the protein was not detected in the labeling datasets. An absence of standard deviation about the mean in the right-hand column indicates the protein was present in only one of two replicates.

|  | Average Fold Enrichment of CCV Relative to DFGL |  |
| --- | --- | --- |
|  | Immunoblot | Dimethyl Labeling |
| **CHC** | 5.15 ± 2.37 | 5.68 ± 1.48 |
| **AP4E** | 3.14 ± 1.10 | 3.18 ± 0.60 |
| **CLC2** | 2.07 ± 0.14 | 18.17 ± 13.70 |
| **AP2M** | 4.33 ± 1.91 | 2.89 |
| **AP1G** | 1.69 ± 0.53 | 5.28 ± 0.72 |
| **KNOLLE** | 1.29 ± 0.15 | 6.28 ± 4.87 |
| **T-PLATE** | 0.88 ± 0.28 | 0.53 |
| **AP2A** | 0.84 ± 0.30 | 2.62 ± 1.06 |
| **cFBPase** | 0.77 ± 0.43 | - |
| **SEC12** | 0.13 ± 0.06 | - |
| **DRP1A** | 0.16 ± 0.02 | - |
| **DRP1C** | 0.32 ± 0.34 | - |
| **DRP2A** | 0.87 ± 0.59 | - |

The expected enrichment of CCV associated proteins, such as clathrin coat proteins CHC and CLC2 and subunits of AP-1 and AP-2 complexes, as well as the cell plate, TGN, and putative CCV cargo marker, KNOLLE (Boutte et al. 2010; Dhonukshe et al. 2006; Reichardt et al. 2007), is shown in Figure 4A. To demonstrate the removal of non-CCV trafficking associated proteins from the purified CCVs, the depletion of cFBPase and SEC12 proteins, markers for the cytosol and ER, respectively, between the S0.1 fraction and final CCV sample, is shown in Figure 4B. We also quantitated the intensity of the proteins in Figures 4A and 4B at the DFGL and CCV steps of the CCV purification scheme and compared these values to those obtained in the dimethyl labeling experiment (Table 1). The average fold enrichment values between the CCV and DFGL samples for the proteins in Figures 4A and 4B as determined by immunoblotting or dimethyl labeling are compared in Table 1, which shows that the general trends of enrichment of CCV associated proteins and depletion of markers of subcellular compartments not associated with CCV-mediated trafficking away from the final CCV sample are consistent across both methods. These quantitative immunoblotting data support the use of the dimethyl labeling proteomic dataset (Supplemental Dataset 3) as a tool for researchers investigating proteins of interest and the strength of potential connections to CCV mediated trafficking processes.

To assess the contributions of subcellular organelles to the complement of 539 proteins that were depleted at least 2-fold during the final CCV purification step, their predicted subcellular localizations were determined using the SUBAcon (<u>SUB</u>cellular Arabidopsis <u>con</u>sensus) bioinformatics tool, an algorithm which integrates experimental fluorescent and proteomic data, as well as computational prediction algorithms to identify a likely protein location (Hooper, et al. 2014) (Supplemental Dataset 5). The average fold depletion of each of the 539 proteins sorted by subcellular localization as identified by SUBAcon is shown in Figure 5A, and the contribution of each organelle to the abundance of spectral counts represented in the 539 protein depletion-dataset is depicted in Figure 5B. Approximately 54% of depleted spectra were attributed to proteins associated with the cytoplasm and other organelles likely not directly participating in clathrin-mediated trafficking, such as the endoplasmic reticulum (Fig. 5B). Demonstrating the effectiveness of the deuterium/Ficoll gradient, approximately 31% and 4% of all spectra corresponding to peptides more abundant in the DFGL relative to the final post centrifugation CCV fraction (i.e. peptides indicating the 539 depleted proteins) were attributed to components of the ribosome and 26S proteasome, respectively.

**Figure 5.**
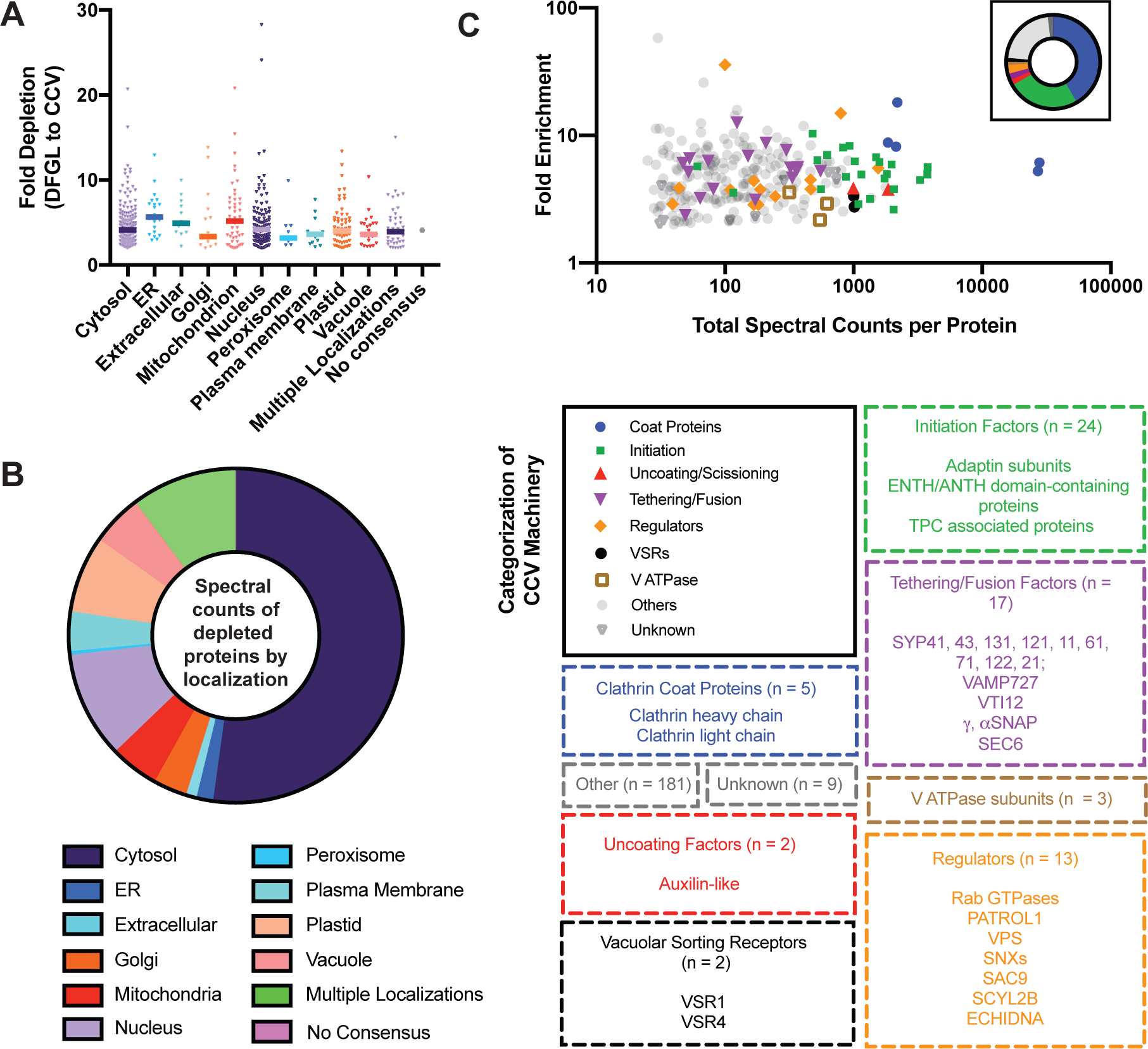
Annotations of proteins identified by shotgun CCV proteomics which were more than two-fold depleted or enriched in the last stage of the CCV purification process. (A) The average fold depletions of the 539 proteins that were more than two-fold depleted between the DFGL and CCV fractions and which overlapped with the CCV LC/MS-MS dataset deriving from 1D SDS-PAGE separation were plotted against the consensus subcellular localizations of the corresponding accession numbers predicted by the SUBA (SUBcellular Arabidopsis) algorithm based on experimental and computational data. (B) Proportion of total spectral counts across four biological replicates for each >2x depleted protein also present in the CCV LC/MS-MS dataset deriving from 1D SDS-PAGE separation as categorized by SUBA annotated subcellular localization. (C) The average fold enrichments of the 256 proteins that were more than two-fold enriched between the DFGL and CCV fractions and which overlapped with the CCV LC/MS-MS dataset deriving from 1D SDS-PAGE separation were plotted against the sum of the total spectral counts of each corresponding protein across four biological replicates. Functional categorization of these proteins were manually annotated; the size and composition of each functional category are indicated below.

Manual annotation of the 256 proteins that were enriched at least 2-fold in the final CCV fraction relative to the deuterium/Ficoll gradient load is depicted in Figure 5C. The fold enrichment and total spectral count are plotted for the enriched proteins sorted by manually assigned category relating to their roles in CCV trafficking. The fold enrichment and abundance of specific categories of CCV associated proteins will be discussed further below.

## VALIDATION OF CCV-ASSOCIATED PROTEINS

### Clathrin

Morphological and SDS-PAGE analyses demonstrated that clathrin heavy chain (CHC) and light chain (CLC) subunits were highly enriched in the CCV preparations (Figures 1A and 2A). Consistent with this, 42% of all spectra assigned to peptides corresponding to the 256 proteins that were enriched in the CCV fraction corresponded to the core clathrin coat components, namely CHC and CLC (Figure 5C). CHC1 and CHC2 were enriched 6- and 5-fold, respectively, in CCVs compared to the DFGL (Table 1, Supplemental Dataset 4), which is comparable to their 3-fold DFGL to CCV enrichment observed via quantitative immunoblotting (Figures 4A and 4C). CLC1-3 were 8-, 18-, and 9-fold enriched in CCVs relative to the DFGL in dimethyl labeling experiments (Supplemental Dataset 4) while quantitative immunoblotting with αCLC2 antibody supported an enrichment for the CLC2 isoform in the final CCV fraction (Table 1, Figures 4A and 4C).

### The AP-1, AP-2, and TPC Hetero-oligomeric Adaptor Protein Complexes

Subunits of the previously-characterized multimeric adaptor protein complexes AP-2, TPC, and AP-1 underlying endocytic and post-Golgi clathrin-dependent trafficking, respectively, were also identified in the suspension-cultured cell CCV proteome. All subunits, including large (A, B, G), medium (M), and small (S) proteins, of the canonical, conserved heterotetrameric adaptin AP-1 and AP-2 complexes were detected in our datasets in high abundance, accounting for 16% of all spectra assigned to peptides enriched in CCVs in dimethyl labeling experiments (Supplemental Dataset 4). AP1G and AP2A were 6- and 3-fold more abundant in CCVs relative to the DFGL as observed by dimethyl labeling (Table 1 and Supplemental Dataset 4). Consistent with this, AP1G was found to be enriched 2-fold and AP2A present in the final CCV fraction as found via quantitative immunoblotting (Table 1, Figures 4A and 4C), indicating that the enriched CCV preparations are a mixed population of post-Golgi and PM-derived CCVs. While both AP1G1 and AP1G2 paralogs were detected in the dimethyl labeling replicates at an enrichment of about 5-fold, only one of two AP2A paralogs, AP2A2, was detected in the labeling studies at an enrichment of 2.6-fold, suggesting that AP2A1 is less abundant in the suspension cultured cell CCV fraction. The medium and small subunits of the AP-1 and AP-2 complexes were enriched 3-fold or greater as were the AP-1/2 B1 and B2 large subunits (6- and 4- fold enrichment; Supplemental Dataset 4), which, similar to the case in Dictyostelium (Sosa et al., 2012) have been postulated to be interchangeably associated with AP-1 and AP-2 complexes in plants (Bassham et al., 2008). Efforts in recent years have established that the role of AP-1 in the trafficking of vacuolar cargo and clathrin recruitment in plant cells resembles that observed in yeast and mammalian systems. AP1M isoform mutants (*ap1m1* and *ap1m2*) both show defects in trafficking of the soluble vacuolar protease precursor proaleurain (Song et al., 2006; Park et al., 2013). In addition, the AP-1 complex is critical for proper targeting of membrane-bound cargo to the tonoplast, at least partially via cytoplasmic sorting signals such as the N-terminal dileucine motif found in VACUOLAR ION TRANSPORTER1 (VIT1), which is mislocalized to the PM in *ap1g* mutants (Wang et al., 2014). TGN/EE integrity is however comprised in *ap1* mutants, which manifests not only in defects in vacuolar protein transport but in exocytic trafficking to the plasma membrane and cell plate, as well as clathrin-mediated endocytosis (Park et al. 2013; Yan et al. 2021).

Recent characterization of the plant-specific TPC has revealed that the complex functions in endocytosis in concert with clathrin and AP-2 (Gadeyne et al., 2014; Bashline et al., 2015; Zhang et al., 2015; Wang et al., 2016). The two adapter complexes likely have overlapping as well as distinct functions (Gadeyne et al., 2014; Narasimhan et al., 2020; Johnson et al. bioRxiv 10.1101/2021.04.26.441441). Consistent with the former, both AP-2 and TPC bind clathrin and have been shown to interact with a common endocytic cargo protein, CESA6 (Bashline et al., 2013; Sanchez-Rodriguez et al., 2018), one of three CESAs identified in the CCV proteome (Supplemental Dataset 4). However, compared to *ap2* mutants, loss-of-function TPC subunit mutants display more severe biological phenotypic defects including pollen lethality (Van Damme et al., 2006; Gadeyne et al., 2014). Thus, given the essential nature of TPC in plant CME and the absence of homologs of most TPC subunits in yeast and metazoans, TPC is critical for functions unique to plants (Zhang et al., 2015). In our studies, mass spectrometry analysis identified all eight core TPC subunits in the proteome derived from 1D SDS-PAGE separated CCVs (Supplemental Dataset 4). However, unlike the subunits of the heterotetrameric AP-1 and AP-2 complexes, which all enriched in the CCV relative to DFGL fraction, the enrichment of the core TPC subunits were generally lower. Although the enrichment values of TPC subunits, TML and TASH3, were somewhat higher than those of the other subunits of TPC (e.g. TPLATE), no TPC subunit was strongly enriched in the last CCV purification step as detected by immunoblotting or dimethyl labeling (Table 1 and Supplemental Dataset 4). Consistent with recent data from Johnson et al. showing that TPC is structurally more external to endocytic CCVs than AP-2, localizing around clathrin and AP2, and that TPC is loosely associated with purified CCV (Johnson et al. bioRxiv), the EH domain-containing proteins EH1 and EH2 were present in the CCV proteome and similarly neither enriched alongside the vesicles as measured by the dimethyl labeling experiments (Supplemental Dataset 4). However, two ENTH-domain-containing TPC accessory components (AtECA4 and CAP1) were found in comparatively high abundance in CCVs (Supplemental Dataset 4). Given that ENTH proteins directly interact with membranes (Zouhar and Sauer, 2014), AtECA4 and CAP1 may be retained on CCVs to a larger extent than the core TPC. These data may explain why TPC components do not enrich to the degree of AP-2 subunits between the DFGL and CCV steps as measured by dimethyl labeling or immunoblotting, in that TPC subunits may have dissociated from the purified CCV. The abundance of TPC core and accessory proteins identified did not differ based on CCV sample preparation, including a similar enrichment of AtECA4 and CAP1 in CCVs relative to TPC core subunits (Supplemental Dataset 4).

### The AP-4 adapter complex, but not AP-3 and AP-5, is associated with CCVs

In addition to the known clathrin associated heterooligomeric adapter complexes, AP-1, AP-2 and TPC, the detection of all subunits of the less-studied AP-4 complex (AP4E, B, M, S) and the abundance thereof at levels comparable to that of AP-1 and AP-2 are notable in the CCV proteome (Supplemental Dataset 4). The enrichment of the AP4E large subunit in CCV preparations was similar to that of AP2A and AP1G subunits (4.6-, 2.6-, and 5.6-fold CCV:DFGL ratios, respectively) as was the medium AP4M subunit relative to those corresponding to AP-2 and AP-1 (3.2-, 2.9-, and 5.9-fold, respectively) as determined by differential labeling experiments (Supplemental Dataset 4). In mammals, AP4M was not detected by immunoblotting in purified CCVs and the AP-4 complex was revealed to associate with non-clathrin-coated vesicles near the TGN via immunogold electron microscopy (Hirst et al., 1999), suggesting AP-4 is not associated with clathrin in these organisms.

In plants, the four AP-4 subunits function together in a complex critical in trafficking to the protein storage vacuole (PSV). Similar to vacuolar sorting receptor (VSR) mutants *vsr1*, *vsr3*, and *vsr4* (Zouhar et al., 2010), GREEN FLUORESCENT SEED (GFS) loss-of-function AP-4 mutants *gfs4, gfs5, gfs6, ap4e-1* corresponding to AP4B, AP4M, AP4S, and AP4E, respectively mislocalize the PSV-targeted 12S globulin seed storage protein to the extracellular space (Fuji et al., 2016). Binding studies have demonstrated that the Arabidopsis AP4M subunit interacts with the cytoplasmic tail of VSR2 (Gershlick et al., 2014). Recently, evidence for interaction between AP4 subunits with DRPs and clathrin was shown via several independent approaches, including coimmunoprecipitations and yeast two-hybrid experiments (Fuji et al., 2016 and Shimizu, et al., 2021) To further investigate the association of AP-4 subunits with plant CCVs, we generated a recombinant antibody against the C-terminal 22 amino acids of the AP4-E large subunit. The specificity of this antibody was confirmed by immunoblotting of total protein extracts prepared from wild-type, *ap4e*, and complemented *ap4e* plants which showed the presence, absence, and presence, respectively, of a band corresponding to the molecular weight of AP4E (Supplemental Figure 2). AP4E co-enriched with *bona fide* CCV proteins and was approximately 90-fold more abundant in the CCV fraction compared with lysate and 3-fold more abundant in the CCV fraction compared to the DFGL (Table 1 and Figures 4A and 4C). As noted in Fuji et al., (2016), the expression levels of AP-4 subunits in Arabidopsis are comparable to those of AP-1 (Park et al., 2013), unlike in animals where AP-4 expression is an order of magnitude lower (Hirst et al., 2013), suggesting a more prominent role for AP-4 in plants.

Previous studies have indicated AP-4 functions at the TGN/EE (Shimizu, et al., 2021; Fuji, et al. 2016; Hirst, et al., 1999). Consistent with this, functional GFP-tagged AP4M colocalizes with the TGN-resident SNARE mRFP-SYP43 and the endocytic tracer FM4-64 in Arabidopsis root tip cells (Fuji et al., 2016). Recently, GFP-tagged AP4M was shown to localize to the TGN/EE at sites distinct from AP1M2 (Shimizu et al., 2021). To further investigate the subcellular distribution of AP-4 and whether it colocalizes with clathrin at the TGN/EE, we constructed lines that stably expressed N-terminal and C-terminal GFP- and RFP-tagged AP4E fusion proteins under control of the ubiquitin-10 promoter (Grefen et al., 2010). These constructs are functional *in vivo* as demonstrated by the rescue of the overall dwarf and abnormal growth phenotype observed in homozygous *ap4e-1* and *ap4e-2* plants (Supplemental Figure 3). Consistent with visualization of GFP-tagged AP4M (Fuji et al., 2016), *ap4e-2* lines that expressed GFP- AP4E primarily displayed cytosolic and endomembrane subcellular localization with signal notably absent from the plasma membrane and tonoplast (Supplemental Figure 4A).

We corroborated the subcellular localization of RFP-AP4E by pulse labeling, co- localization studies with the endocytic tracer dye, FM4-64. This dye has been used to distinguish plant endosomal compartments based on their spatial and temporal distribution of the dye upon internalization, for example, by labeling the TGN/EE within 2- 6 minutes of internalization (Dettmer et al., 2006; Viotti et al., 2010). We observed co- localization between RFP-AP4E and FM4-64 (Pearson’s correlation coefficient [PCC] = 0.705 utilizing a Costes’ automated threshold, Costes P = 1.00) on a similar timescale (Supplemental Figure 4B) which confirmed that AP4E, like AP4M and AP4S, localized to the TGN/EE (Shimizu et al., 2021; Fuji et al., 2016). Clathrin distribution within the cell is divided into soluble (cytosolic) and membrane associated pools, the latter of which includes the plasma membrane, TGN/EE and cell plate. Root epidermal cells expressing CLC2-GFP under control of the *clc2* native promoter (Konopka et al., 2008) and *proUB10::RFP-AP4E* showed occasional colocalization of these fluorophores (PCC = 0.37 utilizing a Costes’ automated threshold, Costes P = 1.00) in endosomal structures, but not at the PM nor at the cell plate (Supplemental Figure 4D). Taken together with the presence of AP-4 in the CCV proteome, these data suggest AP-4 is incorporated into CCVs, likely at the TGN as part of a trafficking pathway to the PSV.

In contrast to AP-1, AP-2, and AP-4, subunits of the AP-3 and AP-5 complexes were absent or were detected in only trace amounts in our CCV proteomes (Supplemental Dataset 4). Mammalian AP-3 is involved in trafficking between recycling and late endosomes (LE) with mixed evidence of a clathrin association (Hirst et al., 1999). In plants, AP-3 has been found to localize to compartments distinct from the TGN/EE, recycling endosome, and Golgi (Feraru et al., 2010). The Arabidopsis AP3B and AP3D mutants, *protein affected trafficking2* (*pat2*) and *pat4,* display defects in a vacuole biogenesis pathway apparently independent of the canonical PVC / MVB maturation sequence, though both mutants display overall normal growth and development, suggesting AP-3 mediates trafficking of some but not all tonoplast proteins (Feraru et al., 2010; Zwiewka et al., 2011; Wolfenstetter et al., 2012; Feng et al., 2017). Furthermore, while Zwiewka et al. observed that clathrin heavy chain was identified as a potential interactor of AP3 by immunoprecipitation (IP) of AP3B subunit and subsequent mass spectrometry (MS), this was not supported by IP of AP3D or in the CCV proteomic datasets presented herein (Supplemental Dataset 4, Zwiewka et al., 2011). Very little is known of the AP-5 complex, though recent studies in mammalian cells have suggested that it functions in late endosome to Golgi protein retrieval in a clathrin independent fashion (Hirst et al., 2011; Hirst et al., 2018). The AP-5 complex is essentially uncharacterized in plants, though its absence in the CCV proteome (Supplemental Dataset 4) suggests its function(s) are also clathrin independent.

### Clathrin Accessory Factors

In addition to hetero-oligomeric protein complexes, monomeric adapters, including members of the ENTH/ANTH/VHS domain-containing protein family and the Golgi-localized, gamma-ear-containing, ARF (ADP-ribosylation factor)-binding (GGA) family, facilitate cargo recognition and vesicle formation at various points in the late endomembrane system. GGA proteins, which function in CCV formation at the TGN in yeast and mammals (Bonifacino, 2004), are absent from plants (Zouhar and Sauer, 2014). Arabidopsis contains 35 ENTH/ANTH/VHS domain-containing genes, 24 of which are putatively expressed in T87 suspension-cultured cells (Stolc et al., 2005). In addition to the TPC accessory proteins AtECA4 and CAP1 (see above), 13 other ENTH/ANTH/VHS-domain containing proteins were identified in the suspension-cultured cell CCV proteome (Supplemental Dataset 4) including the ENTH domain-containing monomeric clathrin adaptors EPSIN1 (EPS1) and EPS2 (Collins et al., 2020; Song et al., 2006; Lee et al., 2007) and the TGN-localized MODIFIED TRANSPORT TO THE VACUOLE 1 (MTV1) (Sauer et al., 2013; Heinze et al., 2020).

EPS1 and MTV1 have previously been implicated in clathrin-mediated trafficking to the vacuole (Heinze et al., 2020; Sauer et al., 2013; Song et al., 2006). Consistent with additional roles in cargo trafficking to the PM, Epsin1 modulates the plasma membrane abundance of the immune receptor FLAGELLIN SENSING2 and its co-receptor, BRI1- ASSOCIATED KINASE (BAK1) for effective defense responses (Collins et al., 2020). In our study, EPS1, EPS2, and MTV1 enriched in CCVs relative to the DFGL in labeling experiments 5-, 5-, and 7-fold, respectively (Supplemental Dataset 4) consistent with data demonstrating their incorporation into CCVs (Sauer et al., 2013), biochemical interaction with clathrin and AP-1 (Song et al., 2006), and similar defects in the trafficking of the soluble vacuolar protease precursor proaleurain in *epsin1* and *ap1m* lines (Song et al., 2006; Park et al., 2013).

Mammalian and plant clathrin-mediated endocytosis are also regulated by Eps15 homology domain-containing proteins which function as membrane and/or protein adaptors (Bar, et al., 2008; Chen, et al., 1998; Yperman, et al., 2021; Schwihla, et al., 2020). The Arabidopsis Eps15 homology domain-containing proteins, EHD1 and EHD2, were also found in the CCV proteome derived from both methodologies (Supplemental Dataset 4).

Several other proteins putatively functioning as trafficking adaptors were identified in the CCV proteome datasets including the sorting nexin homologs SNX1 and SNX2b (Supplemental Dataset 4). Mammalian sorting nexins interact with subunits of the endosomal retromer coat protein complex in a clathrin-independent fashion (McGough and Cullen, 2013) to mediate protein export from the early endosome to the TGN and PM by retrieving sorting receptors (e.g. the mannose-6-phosphate receptor) from the lysosome (Burd and Cullen, 2014). However, an interaction between the Arabidopsis retromer (VPS26a, 26b, 29, 35a, 35b, 35c) and the three sorting nexin homologs (SNX1, 2a, 2b) has yet to be shown, and conflicting evidence in the literature regarding SNX and retromer localization in plant cells at the TGN and MVB (reviewed in (Heucken and Ivanov, 2018; Robinson, 2018)) suggest multiple, independent roles for these complexes. Intriguingly, 12S globulin trafficking to the PSV is disrupted in *snx* mutants while that of 2S globulin remains apparently normal (Pourcher et al., 2010), a phenotype also observed in *ap4* mutants (Fuji et al., 2016). Moreover, a YFP fusion with the *Pisum sativum* (pea) homolog of Arabidopsis VSR1 (BP80) colocalizes with AtSNX1 (Jaillais et al., 2008) and trafficking of a similar BP80 reporter depends on SNX1 and SNX2 function (Niemes et al., 2010). In contrast to SNX1 and SNX2b, components of the retromer core (VPS26A, VPS26B, VPS29, VPS35A, VPS35B, and VPS35C) were identified in the CCV proteome in low abundance (Supplementary Table 4) suggesting a possible role for SNX proteins in CCVs independent of retromer. Further studies are required to investigate the role of SNX proteins and a possible association with CCVs as an evolutionary divergence from the clathrin-independent nature of mammalian retromer / SNX function.

Several SH3Ps (SH3 domain-containing proteins) implicated in CCV trafficking were present in the suspension-cultured cell CCV proteome including the TASH3 subunit of the TPC, as well as SH3P1 and SH3P2, the latter of which is in high abundance and enriched in CCVs relative to DFGL fractions as measured by quantitative dimethyl labeling (8-fold CCV:DFGL, Supplemental Dataset 4) and by immunoblotting (Nagel et al., 2017). Ubiquitylation of PM proteins serves as a post-translational modification signaling for internalization and vacuolar degradation (Martins et al., 2015). SH3P2 localizes to PM and functions as a ubiquitin-binding protein that may facilitate ESCRT recognition of ubiquitylated cargo for subsequent degradation (Nagel et al., 2017). However, SH3P2 also localizes to the cell plate and to other clathrin-positive foci suggesting this and other ubiquitin adaptors may function at other membranes beyond the PM.

Another set of clathrin accessory factors identified in the CCV proteome are Arabidopsis homologs of animal SCY1-LIKE2 proteins (SCYL2A and SCYL2B); SCYL2B is in high abundance and was found to co-enrich with CCVs by quantitative dimethyl labeling MS/MS experiments (15-fold CCV:DFGL ratio, Supplemental Dataset 4). In animals, SCYL2 binds clathrin (Duwel and Ungewickell, 2006), localizes to the Golgi, TGN, and other endosomes, and is incorporated into CCVs (Conner and Schmid, 2005; Borner et al., 2007). Although less is known about their function in plants, SCYL2B was recently shown to localize to the TGN and interact with CHC as well as two related but functionally distinct TGN-associated SNAREs, VTI11 and VTI12, both of which were also identified in the CCV proteome (Supplemental Dataset 4; Jung et al., 2017). Taken together, these data suggest SCYL2A and SCYL2B function may reflect that of mammalian SCYL2 by mediating TGN CCV formation in Arabidopsis.

### Dynamin Related Proteins

Following cargo recognition and clathrin recruitment, the maturing CCV must separate from the plasma membrane prior to internalization. In metazoans, this is accomplished by the action of the dynamin GTPases which oligomerize around the vesicle neck and utilize GTP hydrolysis to exert a constricting, twisting force, achieving scission (Antonny et al., 2016; Cheng et al., 2021). In plants, members of the DYNAMIN RELATED PROTEIN 2 (DRP2) protein family, DRP2A and DRP2B, which are most closely related to mammalian dynamin (Backues et al., 2010; Smith et al., 2014), and the plant-specific DRP1 family members DRP1A and DRP1C (Collings et al., 2008; Smith et al., 2014; Ekanayake et al., 2021; Mravec et al., 2011) function together in cargo trafficking via CME (Ekanayake et al., 2021). DRP2 and DRP1 family members also biochemically interact with TPC and AP-2 (Gadeyne et al., 2014) and localize to PM CCPs (Konopka et al., 2008; Fujimoto et al., 2010; Wang et al., 2020).

Despite their clear association with PM CCPs (Konopka et al., 2008; Konopka and Bednarek, 2008; Fujimoto et al., 2010), DRP proteins were only detected at low levels in the total CCV proteome datasets (Table 1, Supplemental Dataset 4). Consistent with this, DRP1A and DRP1C show significant depletion in the CCV fraction relative to the DFGL as determined via quantitative immunoblotting (Figures 4B and 4C, Table 1). This apparent discrepancy may reflect a short residency of DRP proteins at the PM wherein DRPs are recruited, accomplish their function(s), and dissociate from the newly-formed CCV. The former hypothesis is supported by high-temporal-resolution imaging data showing recruitment of DRP2B, DRP1C, and DRP1A to endocytic foci at the PM simultaneously with or shortly after clathrin before their simultaneous disappearance, potentially reflecting a function in CCV maturation and/or scission (Konopka et al., 2008; Konopka and Bednarek, 2008; Fujimoto et al., 2010). The brief tenure of DRPs at nascent CCVs may also be reflected in the putative interaction between DRPs and the TASH3 subunits of the TPC (Gadeyne et al., 2014), which, along with other core TPC components, were likewise detected at relatively low abundance in the suspension-cultured cell CCV proteome (Supplemental Dataset 4). This parallel may reflect a functional coordination of recruitment and dissociation between DRPs and TPC in endocytic CCV formation. Alternatively, we cannot rule out the low abundance of DRP peptides in our dataset may be due to poor retention of the DRPs during the CCV isolation. In contrast, the DnaJ related AUXILIN-LIKE1 and AUXILIN-LIKE2 proteins, which are postulated to function in uncoating of CCVs (Lam et al., 2001; Adamowski et al., 2018), were readily identified in the suspension-cell cultured CCV proteome (Figure 5C; Supplemental Dataset 4), possibly due to the conditions used in vesicle preparation (pH 6.4) that block clathrin coat disassembly but which permit association of the AUXILIN-LIKE proteins with CCVs (Reynolds et al 2014). Furthermore, recent data suggests that uncoating of CCVs in plants may not occur immediately after scission but as the CCV is trafficked away from the plasma membrane (Narasimhan et al., 2021).

Our results appear consistent with those observed in mammalian and yeast systems including the apparent uncertainty of dynamin association with isolated mammalian CCVs with the protein being alternatively undetected (Blondeau et al., 2004) or observed with high confidence (Borner et al., 2006) in different preparations. However, the precise role(s) of members of the DRP2 and DRP1 protein families in CCV maturation and/or scission remains to be determined.

### Vacuolar Protein Trafficking

Plant Vacuolar Sorting Receptors (VSRs) bind soluble vacuolar proteins including hydrolytic enzymes and vacuolar storage proteins (e.g. 12S globulin and 2S albumin) in the lumen of the secretory pathway through recognition of sorting motifs of vacuolar ligands, sequestering these cargo from others which are secreted or targeted towards the plasma membrane (Shimada, et al. 2003). The primary mode of VSR function in plants had been postulated to reflect that of the mammalian mannose 6-phosphate receptor (MPR), which binds and traffics soluble lysosomal hydrolases via CCVs to the late endosome. The increasingly acidic nature of the maturing endosome triggers pH-dependent dissociation of its ligand, resulting in MPR recycling back to the TGN via retromer-mediated trafficking (Braulke and Bonifacino, 2009), at which the pH falls within the pH range for optimal MPR-ligand binding established by *in vitro* experiments (Tong et al. 1989). However, increasing evidence indicates that VSR-mediated vacuolar protein trafficking is distinct in plants (Robinson and Neuhaus, 2016). In one instance of divergence from the mammalian MPR sorting model, plant VSR-ligand interaction appears to be initiated in the ER and cis-Golgi rather than the TGN/EE (daSilva et al., 2005; Gershlick et al., 2014; Kunzl et al., 2016). Furthermore, the pH of the plant TGN (5.5) is lower than that of other compartments involved in VSR trafficking (Luo et al., 2015) ostensibly marking it as the site of ligand dissociation. Thus, the newly liberated VSRs must be recycled from the TGN back to the ER. In line with this hypothesis, a recent study has demonstrated the retrograde movement of nano-body tagged VSR from the TGN/EE to the early secretory pathway in tobacco protoplasts (Fruholz et al., 2018). Therefore, in addition to the AP/CCV-mediated anterograde trafficking of VSRs, alternative proposals have questioned whether VSR-laden CCVs carry soluble vacuolar cargo directly or are instead part of a recycling mechanism responsible for bringing VSRs back to the early secretory pathway after cargo dissociation in the TGN (Reviewed in (Kang and Hwang, 2014; Robinson and Neuhaus, 2016).

Previous analyses of isolated plant CCVs have identified abundant VSR proteins consistent with a role for CCVs as VSR carriers (Masclaux et al., 2005; De Marcos Lousa et al., 2012). Indeed, the first plant VSR to be discovered (BP80) was initially detected in CCVs isolated from pea (*P. sativum*) cotyledons (Kirsch et al., 1994), and VSRs have been shown to co-fractionate with clathrin and vacuolar cargos during CCV preparations (Jolliffe et al., 2004). Consistent with this, five VSR proteins were identified in the suspension-cultured cell CCV proteome, of which three (VSR1, 3, and 4) were in highest abundance (Figure 5C, Supplemental Dataset 4).

Adaptor protein 1 and 4 complexes mediate the interaction of clathrin with VSR- ligand complexes, facilitating clustering of vacuolar cargo and vesicular formation necessary for vacuolar protein trafficking. The function of the AP-1 complex as an adaptor of vacuolar trafficking in plants reflects that of its mammalian/yeast counterparts in lysosomal/vacuolar trafficking. Recent work in *S. cerevisiae* has demonstrated AP-1 is critical for retrograde recycling of cargo from mature to earlier Golgi cisternae (Papanikou et al., 2015; Day et al., 2018; Casler et al., 2019) suggesting that retrograde trafficking of VSRs and other cargo from the TGN and/or late Golgi to the early secretory pathway in plants may be mediated by AP-dependent CCVs. Moreover, binding of plant VSRs to AP1M is dependent on a cytoplasmic motif and is required for proper trafficking and maturation of the 12S globulin precursor seed storage protein (Gershlick et al., 2014; Fuji et al., 2016). Another member of the Arabidopsis VSR family, VSR4, interacts with AP1M2 partly through a complex cargo-recognition sequence. Mutations of this AP1M2 interaction motif results in VSR4 mislocalization to the PM and increased residency at the tonoplast, suggesting that this motif mediates both anterograde and retrograde VSR4 transport (Nishimura et al., 2016).

As noted above, mutants of AP-4 subunits mislocalize PSV cargo, a phenotype also observed in *vsr* and AP-1 subunit mutants (Fuji et al., 2016; Zouhar et al., 2010; Park et al., 2013), raising questions regarding the cargo and destination specificity of AP-1 and AP-4 dependent vacuolar trafficking. Pathway specific accessory proteins may aid in differentiating AP-1 and AP-4 function, such as the Arabidopsis homolog of the mammalian AP-4 accessory protein tepsin, MTV1, which has been shown to both bind clathrin and co-enrich with isolated CCVs as measured by immunoblotting and electron microscopy techniques (Heinze et al., 2020; Borner et al., 2012; Sauer et al., 2013). We have also found the AP4 interactor, MTV1, to be highly abundant in the Arabidopsis suspension-cultured cell CCV proteome and enriched in CCV pools relative to DFGL (7- fold enrichment, Supplementary Table 4). Recent studies present a genetic interaction between *AP4* and *MTV1* and demonstrate that MTV1 functions in vacuolar trafficking (Heinze, et al. 2020), and the presence of EPS1, AP-1, AP-4, VTI11, and VSR1, 3, and 4 in the CCV dataset (Figure 5C, Supplemental Dataset 4) considered alongside available data regarding the vacuolar trafficking defects of *ap1, ap4,* and *vsr* mutants suggest that AP-1 and AP-4 facilitate anterograde, and possibly retrograde, CCV-mediated trafficking of VSRs to some degree.

### Regulators of Clathrin-Mediated Trafficking

#### Phospholipid Metabolism

Intriguingly, one of the most abundant CCV proteome components identified was the phosphoinositide phosphatase SAC9, which is thought to regulate levels of phosphatidylinositol 4,5-bisphosphate [PI(4,5)P2] (Williams et al., 2005; Vollmer et al., 2011). In mammals, PI(4,5)P2 in the inner leaflet of the PM mediates CCP formation (Antonescu et al., 2011) through interactions with clathrin adaptor and accessory proteins including subunits of the AP-2 complex (AP2A, AP2B, AP2M) (Jackson et al., 2010), epsin (Itoh et al., 2001), and dynamin (Vallis et al., 1999). The importance of these interactions in plant CME remains to be demonstrated, though PI(4,5)P2 accumulates at sites of high membrane flux, like the growing pollen tube tip (Kost et al., 1999; Yao et al., 1999) and at the apex of expanding root hairs (Braun et al., 1999) and is incorporated into CCVs (Zhao et al., 2010). Moreover, mutants lacking the CCV-associated enzymes involved in the synthesis of PI(4,5)P2, phosphatidylinositol 4-phosphate 5-kinases PIP5K1 and PIP5K, show fewer CCPs at the PM of root epidermal cells (Ischebeck et al., 2013). Studies in plants have demonstrated that PI(4,5)P2 recruits the endocytic accessory factor AP2M to the plasma membrane and interacts with the EH1 subunit of the TPLATE complex, suggesting that this phospholipid plays a role in the regulation of clathrin mediated endocytosis (Doumane et al., 2021; Yperman et al., 2021). The presence of SAC9 in the CCV proteome (Figure 5C; Supplemental Dataset 4), taken together with the numerous defects in membrane morphology including vesicle accumulation observed in Arabidopsis *sac9* mutants (Vollmer et al., 2011), suggests that PI(4,5)P2 turnover in mature CCVs may play an important role in the regulation of CCV trafficking.

#### Small GTPases

Rabs and ARF GTPases regulate vesicle trafficking by modulating between GTP and GDP (active and inactive) bound states which govern interactions with downstream effector proteins. Arabidopsis maintains 57 Rabs and 27 ARFs, of which the former can be grouped into eight distinct clades corresponding to mammalian Rab groups mediating different trafficking pathways (Rutherford and Moore, 2002). Of the numerous Rabs identified in our study, members of the RabA1, D2, and E1 families were most abundant (Figure 5C; Supplemental Dataset 4). RabA proteins are most closely related to mammalian Rab11, which regulates endosomal trafficking and mediate TGN-PM trafficking in plants (Zhou, et al., 2020; Li, et al., 2017; Nielsen et al., 2008). In addition, members of the RabE family, which are orthologs of the Sec4p/Rab8 GTPase family, mediate exocytosis in yeast and mammalian cells (Rutherford and Moore, 2002) and regulate post-Golgi trafficking to the PM and to the cell plate during cytokinesis (Orr, et al., 2021; Speth et al., 2009; Ahn et al., 2013). RabD2 proteins are related to mammalian Rab1s which participate in the early secretory pathway. In plants however, RabD2 proteins localize to the Golgi / TGN (Pinheiro et al., 2009) and mediate post-Golgi trafficking at certain endosomes (Drakakaki et al., 2012).

Guanine nucleotide exchange factors (GEFs) are tasked with activating GTPases by catalyzing the exchange of GDP for GTP. Evidence has shown STOMATAL CYTOKINESIS DEFECTIVE1 (SCD1) and SCD2 proteins participate in a complex that interacts with RabE1s in a nucleotide-dependent matter as well as subunits of the exocyst (Mayers et al., 2017), suggesting the SCD complex may function as a RabGEF in exocytosis. Both SCD1 and SCD2 enrich with the purification of CCVs by immunoblot analysis (McMichael et al., 2013) and were detected, albeit at low levels, in both unlabeled CCV proteomic datasets (Supplemental Dataset 4). Despite the presence of SCD1 and SCD2 in enriched CCVs and impaired internalization and post-Golgi trafficking defects of *scd* mutants, it is not clear whether the SCD complex directly functions in endocytosis or functions in recycling endocytic machinery to the plasma membrane.

The Arabidopsis suspension-cultured cell CCV proteome contains several additional structural and regulatory components of the exocyst tethering complex including SEC6, SEC15b, SEC10, EXO84B, and EXO70A1 (Figure 5C; Supplemental Dataset 4). Mammalian SEC15, SEC10, and EXO84 interact with vesicle-bound small GTPases to facilitate secretory trafficking from the TGN while EXO70 localizes to target membranes (Wu and Guo, 2015). Exocyst function in Arabidopsis appears to be conserved, including putative subcellular localizations at sites with high rates of vesicle fusion, e.g. EXO70A1 and EXO84b at the CP (Fendrych et al., 2010).

In addition to GEFs, which initiate GTPase activation, GTPase activating proteins (GAPs) promoting Rab GTP hydrolysis to terminate GTPase signaling are likewise critical for controlling vesicle trafficking. The ARF-GAP NEVERSHED/AGD5/MTV4 which is required for vacuolar protein trafficking (Sauer et al., 2013), was previously demonstrated to co-enrich with CCVs using immunoblotting and immunoEM techniques. In agreement, this ARF-GAP is well represented in the CCV proteome (Supplemental Dataset 4).

#### SNAREs

In this study, we identified PM- and endosome-localized SNAREs in the CCV proteome (Eisenach et al., 2012; Ichikawa et al., 2014; Suwastika et al., 2008; Bassham et al., 2000; Ebine et al., 2011; da Silva Conceicao et al., 1997; Uemura et al., 2012). SNAREs localized to the PM, such as SYP121, VAMP722, SYP132, and SYP71, were enriched 7-, 5-, 8-, and 6-fold, respectively, in the CCV fraction. Endosomal SNAREs, SYP61, SYP41, VTI12, VAMP727, SYP21, and SYP43, were enriched 6-, 13-, 7-, 5-, 2-, and 9-fold, respectively (Figure 5C; Supplemental Dataset 4). The cell plate/cytokinesis-specific syntaxin and putative CCV cargo KNOLLE (KN) (Lauber et al., 1997; Boutte et al., 2010) was notably enriched in CCVs in dimethylabeling (6-fold) and modestly enriched in quantitative immunoblotting (1.3-fold) experiments (Figure 4, Table 1, Supplementary Table 4). In contrast, SNAREs localized to compartments likely not engaged in CCV trafficking, such as the tonoplast protein VAMP713 (Takemoto et al., 2018) and the Golgi-localized VAMP714 (Uemura et al., 2005), were depleted in CCVs 5- and 3-fold relative to the DFGL, respectively (Supplementary Table 4).

We also identified several homologs of mammalian trafficking regulators involved in vesicle fusion in the CCV proteome, including PROTON ATPASE TRANSLOCATION CONTROL 1 (PATROL1) which contains the Munc13 MUN domain (Figure 4C; Supplemental Dataset 4). While the mammalian Munc13 interacts with the SNARE syntaxin-1 to prime synaptic vesicles for fusion with the PM (Ma et al., 2011), less is known of MUN-domain containing protein function in plants. PATROL1 appears to modulate the delivery of the PM H+-ATPase AHA1 to the PM possibly through interactions with the exocyst complex and localizes to endosomes and dynamic foci at the cell cortex (Hashimoto-Sugimoto et al., 2013; Higaki et al., 2014; Zhu et al., 2018). The presence of both PATROL1 and AHA1 in the CCV proteome (Figure 5C; Supplemental Dataset 4) suggests PATROL may regulate the trafficking of CCVs containing AHA1 through interactions with its MUN domain. Another SNARE interacting protein, VPS45, was detected in abundance in the CCV proteome and found to enrich in CCVs in differential labeling experiments (5-fold CCV:DFGL enrichment, Supplemental Dataset 4). VPS45 is a Munc18 protein that binds and regulates the TGN SNARE complex comprised of SYP41, SYP61, YKT61, and VTI12, potentially to mediate fusion of PVC-derived vesicles at the TGN (Bassham et al., 2000; Zouhar et al., 2009). Given the presence of VPS45 and its cognate SNAREs (SYP41, SYP61, VTI12, YKT61) in the CCV proteome (Figure 5C; Supplemental Dataset 4), clathrin-mediated trafficking may play a role in the regulation / function of the complex, possibly by recycling individual SNAREs for subsequent fusion events.

## Conclusion

Space constraints prohibit a full discussion of all protein groups identified in the suspension-cultured cell CCV proteome; instead, we have reported on a subset of actively discussed protein groups and provided the complete CCV proteome as a rich data reference and resource for future investigations of proteins putatively involved in clathrin-mediated trafficking. An important caveat to interpreting the biological significance of the suspension cultured CCV proteome is that CCV composition reflects the tissue/cell type and developmental stage of the biological source material from which they are isolated. The data reported here reflects the content of CCVs isolated from undifferentiated, rapidly dividing and expanding Arabidopsis suspension cultured cells under conditions amenable to tissue culture. Future experiments probing CCV content in other cell and tissue types under varying conditions or in different genetic backgrounds might apply our isolation methodology (Reynolds et al., 2014) using seedlings as a sample source as recently described (Nagel et al., 2017; Mosesso et al., 2019).

Our proteomic data demonstrate that, in addition to the canonical clathrin coat adapters AP-1 and AP-2, the AP-4 complex is incorporated into plant CCVs in an apparent contrast to its function in metazoans (Mattera et al., 2015; Robinson, 2015). This result inspires numerous questions regarding the evolutionary divergence of plant trafficking proteins relative to other eukaryotes, as well as the identity of which pathway(s) might be mediated by AP-4 and clathrin. Current biochemical and genetic studies in the literature suggest AP-4 mediates the trafficking of specific cargos to the PSV (Gershlick et al., 2014; Fuji et al., 2016), though whether AP-4 function in CCVs directly facilitates the anterograde trafficking of soluble cargos and their cognate receptors (i.e. VSRs) or instead participates in the recycling thereof remains to be determined. Additional studies identifying the composition of AP-2, AP-1, and AP-4-positive CCVs are needed to better understand cargo specificity and identify the regulators governing formation, trafficking, and fusion of these vesicles. Nevertheless, the identification of AP-4 as a CCV-associated protein complex presented here, together with recent genetic and biochemical evidence, will contribute to future experiments focused on elucidating the mechanisms of AP-4- dependent post-Golgi trafficking to the PSV. The question remains: is the interaction between AP-4 and clathrin direct as is the case for AP-1 and AP-2? Coimmunoprecipitation experiments (Fuji et al., 2016; Shimizu, et al., 2021) and AP-4 abundance in the CCV proteome suggest that it might be, but a conclusive demonstration of a direct biochemical interaction remains to be determined. Determining the role of clathrin in VSR sorting, given the recent evidence of an ER/cis-Golgi to TGN/EE VSR- dependent, vacuolar trafficking pathway will be essential as plant endomembrane dynamics become better understood (Robinson and Pimpl, 2014; Robinson and Neuhaus 2016).

In contrast to the evolutionarily conserved endocytic adaptor AP-2, subunits of the TPLATE complex, which is essential for CME, were not significantly enriched in the final enrichment step towards purified CCVs suggesting that the TPLATE complex may dissociate more readily from CCVs following their budding from the plasma membrane relative to AP-2. This observation may be resolved by a recent study in which TPLATE dissociated from CCVs earlier than clathrin and which positioned the TPLATE complex on the periphery of a clathrin coated structure at the plasma membrane (Johnson et al. bioRxiv 2021).The positioning of the TPLATE complex to the periphery of the budding vesicle will be of significant biological interest.

As discussed above, the enriched CCV proteomic datasets contain numerous proteins that are unlikely to be associated with CCVs (e.g. ribosomal subunits). The presence of a protein in a shotgun MS/MS dataset, especially those of low abundance, must be taken as a preliminary indication only of its presence and therefore requires subsequent confirmation. Conversely, the absence or low abundance of proteins of interest might also be explained by low intracellular concentrations, such as in the case of regulatory proteins like the SCD1 and SCD2 subunits of the SCD complex which, while in low abundance in the MS/MS dataset, are shown to coenrich with isolated CCVs via immunoblotting and other techniques. Nevertheless, our CCV proteomics data expand our understanding of the plant endomembrane compartments for subsequent efforts to investigate the plant endomembrane network and their physiological function in the future.

## Supporting information

Supplemental Figures, Table, and References

## ACKNOWLEDGMENTS

The authors would like to acknowledge the VIB Proteomics Core Facility (VIB-UGent Center for Medical Biotechnology in Ghent, Belgium) and the Research Technology Support Facility Proteomics Core (Michigan State University in East Lansing, Michigan) for sample analysis, as well as the University of Wisconsin Biotechnology Center Mass Spectrometry Core Facility (Madison, WI) for help with data processing. Additionally, we are grateful to Sue Weintraub (UT Health San Antonio) and Sydney Thomas (UW-Madison) for assistance with data analysis. This research was supported by grants to S.Y.B. from the National Science Foundation (Nos. 1121998 and 1614915) and a Vilas Associate Award (University of Wisconsin, Madison, Graduate School); to J.P. from the National Natural Science Foundation of China (Nos. 91754104, 31820103008, and 31670283); to I.H. from the National Research Foundation of Korea (No. 2019R1A2B5B03099982). This research was also supported by the Scientific Service Units (SSU) of IST Austria through resources provided by the Electron microscopy Facility (EMF). A.J. is supported by funding from the Austrian Science Fund (FWF): I3630B25 to J.F. A.H. is supported by funding from the National Science Foundation (NSF IOS Nos. 1025837 and 1147032).

## AUTHOR CONTRIBUTIONS

D.A.D., G.D.R., S.Y.B., J.J.H., D.V, A.J., and J.P. conceived the study and designed the experiments. D.A.D., G.D.R., S.Y.B., J.J.H., D.V, A.J., J.P, K.Y., D.E., Y.X., W.K., and N.V. carried out experiments and conducted data analysis. G.D.R., D.A.D., and S.Y.B. wrote the manuscript. D.A.D., G.D.R., S.Y.B., J.J.H., A.H., D.V., A.J., K.Y., G.D., and D.E. edited the manuscript.

## MATERIALS AND METHODS

### Plant Materials and Growth Conditions

Seed stocks of Col-0 (CS70000) and Col-3 (CS708) were obtained from the Arabidopsis Biological Research Center (ABRC). Seed stocks of the *ap4e-1* and *ap4e-2* T-DNA insertion mutants (Fuji et al., 2016) were graciously provided by Tomoo Shimada of Kyoto University. CLC2-GFP transgenic plant lines were generated as previously described (Konopka et al., 2008). Seeds were sterilized in 70% (v/v) ethanol with 0.1% (v/v) Triton X-10 for 5 minutes and in 90% (v/v) ethanol for 1 minute prior to plating on ½ strength MS media (Murashige and Skoog, 1962) containing 0.6% (w/v) agar. Seeds were stratified without light at 4°C for 3 days prior to growing under continuous light at 22°C. Plants grown on soil were transferred from plates after 7 to 14 days to Metro-Mix 360 (SunGro Horticulture) and grown at 22°C long days (16 hours of light exposure). *ap4e-1* and *ap4e- 2* mutants were genotyped using primers GR1, 2, & 5 and GR3, 4, & 5, respectively.

Undifferentiated Arabidopsis T87 cells (Axelos et al., 1992) were maintained in MS media supplemented with 0.2 mg/L 2,4-Dichloropphenoxyacetic acid and 1.32 mM KH2PO4 under continuous light at 22°C on an orbital shaker at 140 RPM. Cells were passaged weekly at 1:10 dilution.

### Primers

GR1 – SAIL866C01_LP – 5’-CATGGGTATTGATGGTCTTGG-3’ GR2 – SAIL866C01_RP – AGACCAGAACAGCTAAGCACG GR3 – SAIL60E03_LP – ATAGGCTTCGAATCGAAGAGC GR4 – SAIL60E03_RP – ATGCAGGTGGAATCGTACTTG GR5 – SAIL_LB3 – TAGCATCTGAATTTCATAACCAATCTCGATA GR6 – SB1859STOPsense – CAAAGATCTCCTCGGCTGAGCACCTCTCTTCTTCA GR7 – SB1859STOPanti - TGAAGAAGAGAGGTGCTCAGCCGAGGAGATCTTTG GR8 – SB1859del673sense - CAAAAAAGCAGGCTTCGCAAGAAGGTCCATGGA GR9 – SB1859del673anti – CTCCATGGACCTTCTTGCGAAGCCTGCTTTTTTG

### Plasmid Construction and Plant Transformation

All oligonucleotide primers used in this study were synthesized by Integrated DNA Technologies. Transgenic plants were generated using the floral dip method (Clough and Bent, 1998) with the *Agrobacterium tumefaciens* strain EHA105. N-terminal and C-terminal GFP and RFP fusions with AP4E under control of the UBQ10 promoter were created in pUBN-Dest and pUBC-Dest vectors (Grefen et al., 2010) using Gateway™ cloning (Invitrogen). The AP4E CDS in pLIC6, obtained from ABRC clone DKLAT1G31730, was moved into pDONR221 (Invitrogen) using the BP Clonase kit. Stop codon insertion and frame correction for pUBN constructs was accomplished with primer pairs GR6&7 and GR8&9, respectively, before using the LR Clonase kit (Invitrogen) to move respective AP4E constructs into pUBN and pUBC destination vectors.

T1 seedlings from pUBN/pUBC transformed plants were sown on soil and selected by spray application of a 120 µg/ml Glufosinate (Liberty Herbicide, Bayer Crop Sciences) water solution containing 0.05% (v/v) Silwet-77 sufactant 7, 9, 12, and 14 days after germination.

### CCV Enrichment and Analysis

CCVs were isolated from undifferentiated Arabidopsis T87 cells as described (Reynolds et al., 2014). Total protein yield in the enriched CCV fraction was approximately 300-500 μg per biological replicate. Protein concentrations of individual fractions for subsequent immunoblotting analysis and total CCV yield were obtained using the Pierce® 660nm Protein Assay (Thermo Scientific).

Immunoblotting was performed as described (McMichael et al., 2013) Information about generation of antibodies and concentrations used is described in Supplemental Table 1.

### Anti-AP4E Antibody Generation

The c-terminal 22 amino acids of the Arabidopsis AP4 Epsilon subunit were cloned from the ABRC stock DKLAT1G31730 into the pAN4GST GST-expression vector and the resulting construct used for phage display as described (Blanc et al., 2014) by the Geneva Antibody Facility at the Université de Genève, Switzerland. Antibody specificity was tested by probing the total protein content of wild-type, *ap4e-1*, *ap4e-2*, and various transgenic plants expressing GRP and RFP fusions with AP4E (Fig. S2). Seedlings grown on plates as described for 7-14 days were flash frozen and homogenized via mortar and pestle under liquid nitrogen. Tissue was resuspended in 2X lamelli buffer, quantified using the Pierce® 660nm Protein Assay (Thermo Scientific), and diluted accordingly such that 23.5 µg of each sample in equal volumes could be loaded on an 11% Tris-HCl SDS-PAGE gel and analyzed via western blotting.

### Light Microscopy

All confocal imaging experiments were conducted on a Nikon A1R-Si+ microscope. Colocalization and localization observations were made in root tip epidermal cells of 5 to 7-day old seedlings grown on plates as described above. Seedlings were mounted for imaging in ½ MS media. FM4-64 treated samples were incubated in ½ MS containing 4µM FM4-64 for three minutes and mounted in the same solution before imaging after 6 minutes total incubation.

Colocalization analysis was performed using the JACoP plugin (Bolte and Cordelieres, 2006) in the Fiji (Schindelin et al., 2012) distribution of ImageJ2 (Rueden et al., 2017). Images were processed to remove background (Rolling ball 50-pixel diameter) and cropped to relevant ROIs before analysis with JACoP utilizing 1000 Costes randomizations with a point spread function of two pixels and Costes’ automated thresholding.

### STEM Imaging of Purified CCVs

4 µl of purified CCV preparation (0.33 mg/µl) were applied to carbon-coated and glow discharged (2 minutes at 7x10-1 mbar) 300-mesh copper EM grids (Electron Microscopy Sciences; CF300-CU) and incubated for four minutes at room temperature. Excess solution was removed with blotting paper and samples immediately fixed by a 20 minute incubation with 2% glutaraldehyde (v/v) in PEM buffer (100 mM PIPES, 1 mM MgCl2, 1 mM EGTA, pH 6.9). Samples were then washed with phosphate buffer (0.1 M, pH 7.4) and distilled water, followed by a 20 minute incubation with 0.1% tannic acid (w/v) in MilliQ water. After three washes in water, the samples were incubated with 0.2% aqueous uranyl acetate for 30 minutes at room temperature and then washed three times with water before being dehydrated in a series of graded ethanol (10%, 20%, 40%, 60%, 80%, 96% and 100% for 2 minutes each) and two washes with hexamethyldisilane (99%). Dried samples were then coated with 3 nm platinum and 4 nm carbon using an ACE600 coating device (Leica Microsystems). The sample grids were imaged using a JEOL JEM2800 scanning/transmission electron microscope at 200 kV.

### LC/MS/MS of CCVs separated by 1D SDS-PAGE

Enriched CCV fractions were resolved via 1D SDS-PAGE on a 4-15% Tris-HCl gradient gel (BioRad cat# 161-1158) at a constant 200V for ∼90 minutes. Gels were stained with Coomassie R250 and cut into ∼10 bands. Gel bands were digested in-gel as described (Shevchenko et al., 1996) with modifications. Briefly, gel bands were dehydrated using 100% acetonitrile and incubated with 10mM dithiothreitol in 100mM ammonium bicarbonate, pH∼8, at 56 ℃ for 45min, dehydrated again and incubated in the dark with 50mM iodoacetamide in 100mM ammonium bicarbonate for 20 minutes. Gel bands were then washed with ammonium bicarbonate and dehydrated again. Sequencing grade modified trypsin was prepared to 0.01ug/µL in 50mM ammonium bicarbonate and ∼50uL of this was added to each gel band so that the gel was completely submerged. Bands were then incubated at 37°C overnight. Peptides were extracted from the gel by water bath sonication in a solution of 60%ACN/1%TCA and vacuum dried to ∼2uL.

Enriched DFGL or CCV fractions for dimethyl labeling experiments were prepared in the same manner as the other CCV samples up to peptide extraction and vacuum drying except that a 12.5% Tris-HCl SDS polyacrylamide gel was used. Each peptide sample was then re-suspended in 100mM Triethylammonium and labeled in solution with dimethyl reagents (light label – C2H6, medium label – C2H2D4) according to (Boersema et al. 2009). After labeling, peptides were purified using solid phase extraction tips (OMIX, www.varian.com). Same slice samples from each condition were combined and dried to ∼2uL. Peptides were then re-suspended in 2% acetonitrile/0.1%TFA to 25uL. From this, 10uL were automatically injected by a Waters nanoAcquity Sample Manager (www.waters.com) and loaded for 5 minutes onto a Waters Symmetry C18 peptide trap (5um, 180um x 20mm) at 4uL/min in 5%ACN/0.1%Formic Acid. The bound peptides were then eluted onto a MICHROM Bioresources (www.michrom.com) 0.1 x 150mm column packed with 3u, 200A Magic C18AQ material over 90min with a gradient of 5% B to 35% B in 77min, ramping to 90%B at 79min, holding for 1min and returning to 5%B at 80.1min for the remainder of the analysis using a Waters nanoAcquity UPLC (Buffer A = 99.9% Water/0.1% Formic Acid, Buffer B = 99.9% Acetonitrile/0.1% Formic Acid) with an initial flow rate of 1uL/min.

Peptides were then re-suspended in 2% acetonitrile/0.1%TFA to 20uL. From this, 10uL were automatically injected by a Thermo (www.thermo.com) EASYnLC onto a Thermo Acclaim PepMap RSLC 0.075mm x 150mm C18 column and eluted at 250nL over 90min with a gradient of 5%B to 30%B in 79min, ramping to 100%B at 80min and held at 100%B for the duration of the run (Buffer A = 99.9% Water/0.1% Formic Acid, Buffer B = 99.9% Acetonitrile/0.1% Formic Acid). Eluted peptides were analyzed as follows: **CCV I**: peptides were sprayed into a ThermoFisher LTQ-FT Ultra mass spectrometer using a Michrom ADVANCE nanospray source with survey scans were taken in the FT (25000 resolution determined at m/z 400) and the top five ions in each survey scan then subjected to automatic low energy collision induced dissociation (CID) in the LTQ; **CCV II, Dimethyl Labeling Samples:** peptides were sprayed into a ThermoFisher LTQ Linear Ion trap mass spectrometer using a Michrom ADVANCE nanospray source with the top eight ions in each survey scan are then subjected to low energy CID in a data dependent manner; **CCV III & CCV IV**: peptides were sprayed into a ThermoFisher Q-Exactive mass spectrometer using a FlexSpray spray ion source with survey scans were taken in the Orbi trap (70,000 resolution, determined at m/z 200) and the top twelve ions in each survey scan are then subjected to automatic higher energy collision induced dissociation (HCD at 25%) with fragment spectra acquired at 17,500 resolution. The resulting MS/MS spectra from CCV replicates were converted to peak lists using Mascot Distiller, v2.4.3.3 (www.matrixscience.com) and searched against a custom database which included all entries in the TAIR10 protein sequence database (downloaded from www.arabidopsis.org), appended with common laboratory contaminants (downloaded from www.thegpm.org, cRAP project), using the Mascot searching algorithm, v2.4, with the following parameters: 2 allowed missed tryptic cleavages, fixed modification of carbamidomethyl cysteine, variable modification of oxidation of methionine, peptide tolerance ±5ppm, MS/MS tolerance 0.3 Da. False Discovery Rates (FDR) were calculated against a randomized database search. The Mascot output was then analyzed using Scaffold, v4.8.4 (Proteome Software Inc., Portland, OR) to probabilistically validate protein identifications. Assignments validated using the Scaffold 1% FDR protein threshold, containing 2 unique peptides, and meeting the 95% confidence peptide threshold were considered true. Assignments matching these criteria from four distinct biological replicates (CCV enrichments) were cross-referenced to eliminate duplicates and filtered to exclude proteins not present in at least two biological replicates, generating a list of 3,548 protein assignments.

Quantitation of labeled MS peaks from dimethyl labeling samples and processing of the resulting MS/MS spectra to peak lists was done using MaxQuant3, v1.2.2.5 (Cox and Mann, 2008). Peak lists were searched against the TAIR10 protein sequence database, downloaded from www.arabidopsis.org and appended with common laboratory contaminants using the Andromeda search algorithm within MaxQuant with the following parameters: two allowed missed tryptic cleavages, fixed modification of carbamidomethyl cysteine, variable modification of oxidation of methionine and acetylation of protein N-termini, DimethLys0, DimethNter0; DimethLys4, DimethNter4, peptide tolerance ±6ppm, MS/MS tolerance 20ppm, protein and peptide FDR filter set to 1%. A total of 1,109 unique proteins were identified between both replicates, 948 of which were identified in the LC/MS-MS dataset corresponding to CCVs digested in gel across two or more biological replicates.

Overlaps between proteomic datasets were identified by matching accession numbers between the 3,548 accession numbers in column 1 of Supplemental Dataset 1, the accession numbers of the 1,109 proteins identified in at least one of two dimethyl replicates (Supplemental Dataset 3), and the first accession number (corresponding to iBAQ values listed) of the majority accession numbers within each protein group (n = 1,981) in the alternative methodology CCV LC/MS-MS dataset (Supplemental Dataset 4). Overlaps between proteomic datasets were visualized using the application, BioVenn (Hulsen et al., 2008).

Protein localization of the 536 more than two-fold depleted proteins between the DFGL and CCV purification steps (Supplemental Dataset 3) were determined by submission of the corresponding accession numbers (Supplemental Dataset 5) to the SUBcellular Arabidopsis (SUBA) consensus algorithm (Hooper et al., 2014).

### LC/MS-MS of CCVs digested in solution

5µg of 1 µg/µl isolated CCVs were lysed in a urea lysis buffer containing 8 M urea, 20 mM HEPES pH 8.0, by repeatedly pipetting up and down. Proteins in each sample were reduced by adding 15 mM DTT and incubation for 30 minutes at 55°C. Alkylation of the proteins was done by addition of 30 mM iodoacetamide for 15 minutes at room temperature in the dark. The samples were diluted with 20 mM HEPES pH 8.0 to a urea concentration of 2 M and the proteins were digested with 4 µl Trypsin/LysC (Promega V5073: 20ug + 80uL 50mM acetic acid) for 4 hours at 37°C and boosted with an extra 2µl Trypsin/LysC (Promega V5073: 20ug + 80uL 50mM acetic acid) overnight at 37°C. Peptides were then purified on a OMIX C18 pipette tip (Agilent).

Purified peptides were re-dissolved in 25 µl loading solvent A (0.1% TFA in water/ACN (98:2, v/v)) and 5 µl was injected for LC-MS/MS analysis on an Ultimate 3000 RSLCnano ProFLow system in-line connected to a Q Exactive HF mass spectrometer (Thermo). Trapping was performed at 10 μl/min for 4 min in loading solvent A on a 20 mm trapping column (made in-house, 100 μm internal diameter (I.D.), 5 μm beads, C18 Reprosil-HD, Dr. Maisch, Germany) and the sample was loaded on a 200 mm analytical column (made in-house, 75 µm I.D., 1.9 µm beads C18 Reprosil-HD, Dr. Maisch). Peptides were eluted by a non-linear gradient from 2 to 55% MS solvent B (0.1% FA in water/acetonitrile (2:8, v/v)) over 175 minutes at a constant flow rate of 250 nl/min, reaching 99% MS solvent B after 200 minutes, followed by a 10 minute wash with 99% MS solvent B and re-equilibration with MS solvent A (0.1% FA in water). The column temperature was kept constant at 50°C in a column oven (Butterfly, Phoenix S&T). The mass spectrometer was operated in data-dependent mode, automatically switching between MS and MS/MS acquisition for the 16 most abundant ion peaks per MS spectrum. Full-scan MS spectra (375-1500 m/z) were acquired at a resolution of 60,000 in the orbitrap analyzer after accumulation to a target value of 3E6. The 16 most intense ions above a threshold value of 1.3E4 were isolated for fragmentation at a normalized collision energy of 28% after filling the trap at a target value of 1E5 for maximum 80 ms. MS/MS spectra (200-2000 m/z) were acquired at a resolution of 15,000 in the orbitrap analyzer.

Data analysis was performed with MaxQuant (version 1.6.10.43) using the built in Andromeda search engine with default settings, including a false discovery rate set at 1% on both the peptide and protein level. Spectra were searched against the Araport11plus database, consisting of the Araport11_genes.2016.06.pep.fasta downloaded from www.arabidopsis.org, extended with sequences of all types of possible contaminants in proteomics experiments in general. These contaminants include the cRAP protein sequences, a list of proteins commonly found in proteomics experiments, which are present either by accident or by unavoidable contamination of protein samples (The Global Proteome Machine, http://www.thegpm.org/crap/). In addition, commonly used tag sequences and typical contaminants, such as sequences from frequently used resins and proteases, were added. The Araport11plus database contains in total 49,057 sequence entries. The mass tolerance for precursor and fragment ions was set to 4.5 and 20 ppm, respectively, including matching between runs and false discovery rate set at 1% on PSM, peptide and protein level. Enzyme specificity was set as C-terminal to arginine and lysine (trypsin), also allowing cleavage at arginine/lysine-proline bonds with a maximum of two missed cleavages. Variable modifications were set to oxidation of methionine residues and acetylation of protein N-termini. Proteins were quantified by the MaxLFQ algorithm integrated in the MaxQuant software. Only protein groups with at least one unique peptide and with peptides identified in at least two of three biological replicates were retained, yielding a list of 1,981 protein groups (Supplemental Dataset 4).

## Plant Accession #s

Accession numbers corresponding to proteins identified in this study can be found in Supplemental Datasets 1-3.

## Proteomic Data Deposition

Files corresponding to raw proteomics data, as well as results, search, and peak list files can be accessed on the MassIVE data repository (MassIVE, University of California at San Diego) using the identifiers, PXD026180 or at doi:10.25345/C50F9D, or at the following URL: ftp://massive.ucsd.edu/MSV000087472/. A summary of the methods & protocols used, as well as gel images and supplementary files, can also be accessed here.

## Supplemental Figures & Table

Supplemental Figure 1: Schematics illustrating protocol for clathrin coated vesicle purification and workflows detailing the CCV proteome.

Supplemental Figure 2: AP4E antibody is specific for the AP4E subunit. Supplemental Figure 3: pUB10::GFP-AP4E is functional *in vivo*.

Supplemental Figure 4: AP4E colocalizes with FM4-64 and clathrin at the TGN. Supplemental Figure 5: Transmission electron microscopy of CCVs.

Supplemental Table 1: Antibodies used in this study.

<u>Supplemental Data files</u> One Excel file containing:

Supplemental Dataset 1: Merged CCV proteomic datasets

Supplemental Dataset 2 LC/MS-MS data corresponding to CCVs digested in solution Supplemental Dataset 3: CCV dimethyl labeling LC/MS-MS data

Supplemental Dataset 4: LC/MS-MS data corresponding to discussed proteins Supplemental Dataset 5: Predicted protein localization data for proteins more than two-fold depleted between DFGL and CCV fractions

## References

Adamowski, M., Narasimhan, M., Kania, U., Glanc, M., De Jaeger, G., and Friml, J. (2018). A Functional Study of AUXILIN-LIKE1 and 2, Two Putative Clathrin Uncoating Factors in Arabidopsis. Plant Cell 30, 700–716.

Ahn, C.S., Han, J.A., and Pai, H.S. (2013). Characterization of in vivo functions of Nicotiana benthamiana RabE1. Planta 237, 161–172.

Antonescu, C.N., Aguet, F., Danuser, G., and Schmid, S.L. (2011). Phosphatidylinositol-(4,5)- bisphosphate regulates clathrin-coated pit initiation, stabilization, and size. Mol Biol Cell 22, 2588–2600.

Antonny, B., Burd, C., De Camilli, P., Chen, E., Daumke, O., Faelber, K., Ford, M., Frolov, V.A., Frost, A., Hinshaw, J.E., Kirchhausen, T., Kozlov, M.M., Lenz, M., Low, H.H., McMahon, H., Merrifield, C., Pollard, T.D., Robinson, P.J., Roux, A., and Schmid, S. (2016). Membrane fission by dynamin: what we know and what we need to know. EMBO J 35, 2270–2284.

Arabidopsis Genome, I. (2000). Analysis of the genome sequence of the flowering plant Arabidopsis thaliana. Nature 408, 796–815.

Arike, L., Valgepea, K., Peil, L., Nahku, R., Adamberg, K., Vilu, R. (2012) Comparison and applications of label-free absolute proteome quantification methods on *Escherichia coli*. Journal of Proteomics. 75, 5437–5448.

Axelos, M., Curie, C., Mazzolini, L., Bardet, C., and Lescure, B. (1992). A Protocol for Transient Gene-Expression in Arabidopsis-Thaliana Protoplasts Isolated from Cell-Suspension Cultures. Plant Physiol Bioch 30, 123–128.

Backues, S.K., Korasick, D.A., Heese, A., and Bednarek, S.Y. (2010). The Arabidopsis dynamin-related protein2 family is essential for gametophyte development. Plant Cell 22, 3218–3231.

Bar, M., Aharon, M., Benjamin, S., Rotblat, B., Horowitz, M., Avni, A. (2008). AtEHDs, novel Arabidopsis EH-domain-containing proteins involved in endocytosis. Plant J 55, 1025–1038.

Bashline, L., Li, S., Zhu, X., and Gu, Y. (2015). The TWD40-2 protein and the AP2 complex cooperate in the clathrin-mediated endocytosis of cellulose synthase to regulate cellulose biosynthesis. Proc Natl Acad Sci U S A 112, 12870–12875.

Bashline, L., Li, S., Anderson, C.T., Lei, L., and Gu, Y. (2013). The endocytosis of cellulose synthase in Arabidopsis is dependent on mu2, a clathrin-mediated endocytosis adaptin. Plant Physiol 163, 150–160.

Bassham, D.C., Brandizzi, F., Otegui, M.S., and Sanderfoot, A.A. (2008). The secretory system of Arabidopsis. Arabidopsis Book 6, e0116.

Bassham, D.C., Sanderfoot, A.A., Kovaleva, V., Zheng, H., and Raikhel, N.V. (2000). AtVPS45 complex formation at the trans-Golgi network. Mol Biol Cell 11, 2251–2265.

Bednarek, S.Y., and Backues, S.K. (2010) Plant dynamin-related protein families DRP1 and DRP2 in plant development. Biochem Soc Trans 38, 797–806.

Berardini, T.Z., Reiser, L., Li, D., Mezheritsky, Y., Muller, R., Strait, E., and Huala, E. (2015). The Arabidopsis information resource: Making and mining the “gold standard” annotated reference plant genome. Genesis 53, 474–485.

Blanc, C., Zufferey, M., and Cosson, P. (2014). Use of in vivo biotinylated GST fusion proteins to select recombinant antibodies. ALTEX 31, 37–42.

Blondeau, F., Ritter, B., Allaire, P.D., Wasiak, S., Girard, M., Hussain, N.K., Angers, A., Legendre-Guillemin, V., Roy, L., Boismenu, D., Kearney, R.E., Bell, A.W., Bergeron, J.J., and McPherson, P.S. (2004). Tandem MS analysis of brain clathrin-coated vesicles reveals their critical involvement in synaptic vesicle recycling. Proc Natl Acad Sci U S A 101, 3833–3838.

Boersema, P.J., Raijmakers, R., Lemeer, S., Mohammed, S., and Heck, A.J. (2009). Multiplex peptide stable isotope dimethyl labeling for quantitative proteomics. Nat Protoc 4, 484–494.

Bolte, S., and Cordelieres, F.P. (2006). A guided tour into subcellular colocalization analysis in light microscopy. J Microsc 224, 213–232.

Bonifacino, J.S. (2004). The GGA proteins: adaptors on the move. Nat Rev Mol Cell Biol 5, 23–32.

Bonifacino, J.S. and Glick, B.S. (2004). The mechanisms of vesicle budding and fusion. Cell. 116, 153–166.

Bonnett, H.T., and Newcomb, E.H. (1966). Coated vesicles and other cytoplasmic components of growing root hairs of radish. Protoplasma 62, 16.

Borner, G.H., Harbour, M., Hester, S., Lilley, K.S., and Robinson, M.S. (2006). Comparative proteomics of clathrin-coated vesicles. J Cell Biol 175, 571–578.

Borner, G.H., Rana, A.A., Forster, R., Harbour, M., Smith, J.C., and Robinson, M.S. (2007). CVAK104 is a novel regulator of clathrin-mediated SNARE sorting. Traffic 8, 893–903.

Borner, G.H., Antrobus, R., Hirst, J., Bhumbra, G.S., Kozik, P., Jackson, L.P., Sahlender, D.A., and Robinson, M.S. (2012). Multivariate proteomic profiling identifies novel accessory proteins of coated vesicles. J Cell Biol 197, 141–160.

Boutte, Y., Frescatada-Rosa, M., Men, S., Chow, C.M., Ebine, K., Gustavsson, A., Johansson, L., Ueda, T., Moore, I., Jurgens, G., and Grebe, M. (2010). Endocytosis restricts Arabidopsis KNOLLE syntaxin to the cell division plane during late cytokinesis. EMBO J 29, 546–558.

Braulke, T. and Bonifacino, J. (2009) Sorting of lysosomal proteins. BBA Molecular Cell Research. 1793, 605–614.

Braun, M., Baluska, F., von Witsch, M., and Menzel, D. (1999). Redistribution of actin, profilin and phosphatidylinositol-4, 5-bisphosphate in growing and maturing root hairs. Planta 209, 435–443.

Burd, C., and Cullen, P.J. (2014). Retromer: a master conductor of endosome sorting. Cold Spring Harb Perspect Biol 6.

Carter, C., Pan, S., Zouhar, J., Avila, E.L., Girke, T., and Raikhel, N.V. (2004). The vegetative vacuole proteome of Arabidopsis thaliana reveals predicted and unexpected proteins. Plant Cell 16, 3285–3303.

Casler, J.C., Papanikou, E., Barrero, J.J., and Glick, B.S. (2019). Maturation-driven transport and AP-1- dependent recycling of a secretory cargo in the Golgi. J Cell Biol.

Chen, H., Fre, S., Slepnev, V., Capua, M., Takei, K., Butler, M., Di Fiore, P., De Camilli, P. (1998) Epsin is an EH-domain-binding protein implicated in clathrin-mediated endocytosis. Nature 394, 793–797.

Cheng, X., Chen, K., Dong, B., Yang, M., Filbrun, S., Myoung, Y., Huang, T., Gu, Y., Wang, G., Fang, N. (2021) Dynamin-dependent vesicle twist at the final stage of clathrin-mediated endocytosis. Nature Cell Biology 23, 859–869.

Clough, S.J., and Bent, A.F. (1998). Floral dip: a simplified method for Agrobacterium-mediated transformation of Arabidopsis thaliana. Plant J 16, 735–743.

Collings, D.A., Gebbie, L.K., Howles, P.A., Hurley, U.A., Birch, R.J., Cork, A.H., Hocart, C.H., Arioli, T., and Williamson, R.E. (2008). Arabidopsis dynamin-like protein DRP1A: a null mutant with widespread defects in endocytosis, cellulose synthesis, cytokinesis, and cell expansion. J Exp Bot 59, 361–376.

Collins, C.A., LaMontagne, E.D., Anderson, J.C., Ekanayake, G., Clarke, A.S., Bond, L.N., Salamango, D.J., Cornish, P.V., Peck, S.C., Heese, A. (2020) EPSIN1 modulates the plasma membrane abundance of FLAGELLIIN SENSING2 for effective immune responses. Plant Physiology. 182:1762–1775.

Conner, S.D., and Schmid, S.L. (2005). CVAK104 is a novel poly-L-lysine-stimulated kinase that targets the beta2-subunit of AP2. J Biol Chem 280, 21539–21544.

Cox, J., and Mann, M. (2008). MaxQuant enables high peptide identification rates, individualized p.p.b.- range mass accuracies and proteome-wide protein quantification. Nat Biotechnol 26, 1367–1372.

da Silva Conceicao, A., Marty-Mazars, D., Bassham, D.C., Sanderfoot, A.A., Marty, F., and Raikhel, N.V. (1997). The syntaxin homolog AtPEP12p resides on a late post-Golgi compartment in plants. Plant Cell 9, 571–582.

daSilva, L.L., Taylor, J.P., Hadlington, J.L., Hanton, S.L., Snowden, C.J., Fox, S.J., Foresti, O., Brandizzi, F., and Denecke, J. (2005). Receptor salvage from the prevacuolar compartment is essential for efficient vacuolar protein targeting. Plant Cell 17, 132–148.

Day, K.J., Casler, J.C., and Glick, B.S. (2018). Budding Yeast Has a Minimal Endomembrane System. Dev Cell 44, 56–72 e54.

De Marcos Lousa, C., Gershlick, D.C., and Denecke, J. (2012). Mechanisms and concepts paving the way towards a complete transport cycle of plant vacuolar sorting receptors. Plant Cell 24, 1714–1732.

Dell’Angelica, E.C., Ooi, C.E., and Bonifacino, J.S. (1997). Beta3A-adaptin, (Zwiewka et al., 2011). J Biol Chem 272, 15078–15084.

Dell’Angelica, E.C., Mullins, C., and Bonifacino, J.S. (1999). AP-4, a novel protein complex related to clathrin adaptors. J Biol Chem 274, 7278–7285.

Dettmer, J., Hong-Hermesdorf, A., Stierhof, Y.D., and Schumacher, K. (2006). Vacuolar H+-ATPase activity is required for endocytic and secretory trafficking in Arabidopsis. Plant Cell 18, 715–730.

Dhonukshe, P., Baluska, F., Schlicht, M., Hlavacka, A., Samaj, J., Friml, J., Gadella Jr., T. (2006) Endocytosis of cell surface material mediates cell plate formation during plant cytokinesis. Dev Cell 10: 137 – 150.

Di Rubbo, S., Irani, N.G., Kim, S.Y., Xu, Z.Y., Gadeyne, A., Dejonghe, W., Vanhoutte, I., Persiau, G., Eeckhout, D., Simon, S., Song, K., Kleine-Vehn, J., Friml, J., De Jaeger, G., Van Damme, D., Hwang, I., and Russinova, E. (2013). The clathrin adaptor complex AP-2 mediates endocytosis of brassinosteroid insensitive1 in Arabidopsis. Plant Cell 25, 2986–2997.

Doumane, M., Lebecq, A., Colin, L., Fangain, A., Stevens, F., Bareille, J., Hamant, O., Belkhadir, Y., Munnik, T., Jaillais, Y., Caillaud, M. (2021) Inducible depletion of PI(4,5)P2 by the synthetic iDePP system in *Arabidopsis*. Nature Plants 7, 587–597.

Drakakaki, G., van de Ven, W., Pan, S., Miao, Y., Wang, J., Keinath, N.F., Weatherly, B., Jiang, L., Schumacher, K., Hicks, G., and Raikhel, N. (2012). Isolation and proteomic analysis of the SYP61 compartment reveal its role in exocytic trafficking in Arabidopsis. Cell Res 22, 413–424.

Duwel, M., and Ungewickell, E.J. (2006). Clathrin-dependent association of CVAK104 with endosomes and the trans-Golgi network. Mol Biol Cell 17, 4513–4525.

Ebine, K., Fujimoto, M., Okatani, Y., Nishiyama, T., Goh, T., Ito, E., Dainobu, T., Nishitani, A., Uemura, T., Sato, M.H., Thordal-Christensen, H., Tsutsumi, N., Nakano, A., and Ueda, T. (2011). A membrane trafficking pathway regulated by the plant-specific RAB GTPase ARA6. Nat Cell Biol 13, 853–859.

Ekanayake, G., Smith, J., Jones, K., Stiers, H., Robinson, S., LaMontagne, E., Kostos, P., Cornish, P., Bednarek, S., Heese, A. (2021) DYNAMIN-RELATED PROTEIN DRP1A functions with DRP2B in plant growth, flg22-immune responses, and endocytosis. Plant Phys. 185:1986–2002.

Ekanayake, G., LaMontagne, E., Heese, A. (2019) Never walk alone: clathrin-coated vesicle (CCV) components in plant immunity. Annu Rev Phytopathol. 57: 307–409.

Eisenach, C., Chen, Z.H., Grefen, C., and Blatt, M.R. (2012). The trafficking protein SYP121 of Arabidopsis connects programmed stomatal closure and K(+) channel activity with vegetative growth. Plant J 69, 241–251.

Fan, L., Hao, H., Xue, Y., Zhang, L., Song, K., Ding, Z., Botella, M.A., Wang, H., and Lin, J. (2013). Dynamic analysis of Arabidopsis AP2 sigma subunit reveals a key role in clathrin-mediated endocytosis and plant development. Development 140, 3826–3837.

Fendrych, M., Synek, L., Pecenkova, T., Toupalova, H., Cole, R., Drdova, E., Nebesarova, J., Sedinova, M., Hala, M., Fowler, J.E., and Zarsky, V. (2010). The Arabidopsis Exocyst Complex Is Involved in Cytokinesis and Cell Plate Maturation. Plant Cell 22, 3053–3065.

Feng, Q., Song, S., Yu, S., Wang, J., Li, S., Zhang, Y. (2017). Adaptor protein-3-dependent vacuolar trafficking involves a subpopulation of COPII and HOPS tethering proteins. Plant Physiology 174, 1609–1620.

Feraru, E., Paciorek, T., Feraru, M.I., Zwiewka, M., De Groodt, R., De Rycke, R., Kleine-Vehn, J., and Friml, J. (2010). The AP-3 beta adaptin mediates the biogenesis and function of lytic vacuoles in Arabidopsis. Plant Cell 22, 2812–2824.

Fruholz, S., Fassler, F., Kolukisaoglu, U., and Pimpl, P. (2018). Nanobody-triggered lockdown of VSRs reveals ligand reloading in the Golgi. Nat Commun 9, 643.

Fuji, K., Shirakawa, M., Shimono, Y., Kunieda, T., Fukao, Y., Koumoto, Y., Takahashi, H., Hara-Nishimura, I., and Shimada, T. (2016). The Adaptor Complex AP-4 Regulates Vacuolar Protein Sorting at the trans-Golgi Network by Interacting with VACUOLAR SORTING RECEPTOR1. Plant Physiol 170, 211–219.

Fujimoto, M., Ebine, K., Nishimura, K., Tsutsumi, N., and Ueda, T. (2020) Longin R-SNARE is retrieved from the plasma membrane by ANTH domain-containing proteins in *Arabidopsis*. PNAS. 117, 25150–25158.

Fujimoto, M., Arimura, S., Ueda, T., Takanashi, H., Hayashi, Y., Nakano, A., and Tsutsumi, N. (2010). Arabidopsis dynamin-related proteins DRP2B and DRP1A participate together in clathrin-coated vesicle formation during endocytosis. Proc Natl Acad Sci U S A 107, 6094–6099.

Gadeyne, A., Sanchez-Rodriguez, C., Vanneste, S., Di Rubbo, S., Zauber, H., Vanneste, K., Van Leene, J., De Winne, N., Eeckhout, D., Persiau, G., Van De Slijke, E., Cannoot, B., Vercruysse, L., Mayers, J.R., Adamowski, M., Kania, U., Ehrlich, M., Schweighofer, A., Ketelaar, T., Maere, S., Bednarek, S.Y., Friml, J., Gevaert, K., Witters, E., Russinova, E., Persson, S., De Jaeger, G., and Van Damme, D. (2014). The TPLATE adaptor complex drives clathrin-mediated endocytosis in plants. Cell 156, 691–704.

Gershlick, D.C., Lousa Cde, M., Foresti, O., Lee, A.J., Pereira, E.A., daSilva, L.L., Bottanelli, F., and Denecke, J. (2014). Golgi-dependent transport of vacuolar sorting receptors is regulated by COPII, AP1, and AP4 protein complexes in tobacco. Plant Cell 26, 1308–1329.

Girard, M., Allaire, P.D., McPherson, P.S., and Blondeau, F. (2005). Non-stoichiometric relationship between clathrin heavy and light chains revealed by quantitative comparative proteomics of clathrin-coated vesicles from brain and liver. Mol Cell Proteomics 4, 1145–1154.

Gray, E.G. (1961). The granule cells, mossy synapses and Purkinje spine synapses of the cerebellum: light and electron microscope observations. J Anat 95, 345–356.

Grefen, C., Donald, N., Hashimoto, K., Kudla, J., Schumacher, K., and Blatt, M.R. (2010). A ubiquitin-10 promoter-based vector set for fluorescent protein tagging facilitates temporal stability and native protein distribution in transient and stable expression studies. Plant J 64, 355–365.

Gu, Y., Zavaliev, R., and Dong, X. (2017). Membrane Trafficking in Plant Immunity. Mol Plant 10, 1026–1034.

Hashimoto-Sugimoto, M., Higaki, T., Yaeno, T., Nagami, A., Irie, M., Fujimi, M., Miyamoto, M., Akita, K., Negi, J., Shirasu, K., Hasezawa, S., and Iba, K. (2013). A Munc13-like protein in Arabidopsis mediates H+-ATPase translocation that is essential for stomatal responses. Nat Commun 4, 2215.

Heinze, L., Freimuth, N., Robling, A., Hahnke, R., Riebschlager, S., Frohlich, A., Sampathkumar, A., McFarlane, H., Sauer, M. (2020) EPSIN1 and MTV1 define functionally overlapping but molecularly distinct *trans*-Golgi network subdomains in *Arabidopsis*. PNAS 117, 25880–25889.

Heucken, N., and Ivanov, R. (2018). The retromer, sorting nexins and the plant endomembrane protein trafficking. J Cell Sci 131.

Higaki, T., Hashimoto-Sugimoto, M., Akita, K., Iba, K., and Hasezawa, S. (2014). Dynamics and environmental responses of PATROL1 in Arabidopsis subsidiary cells. Plant Cell Physiol 55, 773–780.

Hirst, J., Irving, C., and Borner, G.H. (2013). Adaptor protein complexes AP-4 and AP-5: new players in endosomal trafficking and progressive spastic paraplegia. Traffic 14, 153–164.

Hirst, J., Bright, N.A., Rous, B., and Robinson, M.S. (1999). Characterization of a fourth adaptor-related protein complex. Mol Biol Cell 10, 2787–2802.

Hirst, J., Barlow, L.D., Francisco, G.C., Sahlender, D.A., Seaman, M.N., Dacks, J.B., and Robinson, M.S. (2011). The fifth adaptor protein complex. PLoS Biol 9, e1001170.

Hirst, J., Borner, G.H., Antrobus, R., Peden, A.A., Hodson, N.A., Sahlender, D.A., and Robinson, M.S. (2012). Distinct and overlapping roles for AP-1 and GGAs revealed by the “knocksideways” system. Curr Biol 22, 1711–1716.

Hirst, J., Schlacht, A., Norcott, J.P., Traynor, D., Bloomfield, G., Antrobus, R., Kay, R.R., Dacks, J.B., and Robinson, M.S. (2014). Characterization of TSET, an ancient and widespread membrane trafficking complex. Elife 3, e02866

Hirst, J., Itzhak, D., Antrobus, R., Borner, G., Robinson, M. (2018) Role of the AP-5 adaptor protein complex in late endosome-to-Golgi retrieval. PLoS Biology 16, e2004411.

Hooper, C., Tanz, S., Castleden, I., Vacher, M., Small, I., Millar, AH. (2014) SUBAcon: a consensus algorithm for unifying the subcellular localization data of the *Arabidopsis* proteome. Bioinformatics. 30, 3356–3364.

Hulsen, T., de Vlieg, J., Alkema, W. (2008) BioVenn - a web application for the comparison and visualization of biological lists using area-proportional Venn diagrams. BMC Genomics 9,1–6.

Ichikawa, M., Hirano, T., Enami, K., Fuselier, T., Kato, N., Kwon, C., Voigt, B., Schulze-Lefert, P., Baluska, F., and Sato, M.H. (2014). Syntaxin of plant proteins SYP123 and SYP132 mediate root hair tip growth in Arabidopsis thaliana. Plant Cell Physiol 55, 790–800.

Ischebeck, T., Werner, S., Krishnamoorthy, P., Lerche, J., Meijon, M., Stenzel, I., Lofke, C., Wiessner, T., Im, Y.J., Perera, I.Y., Iven, T., Feussner, I., Busch, W., Boss, W.F., Teichmann, T., Hause, B., Persson, S., and Heilmann, I. (2013). Phosphatidylinositol 4,5-bisphosphate influences PIN polarization by controlling clathrin-mediated membrane trafficking in Arabidopsis. Plant Cell 25, 4894–4911.

Itoh, T., Koshiba, S., Kigawa, T., Kikuchi, A., Yokoyama, S., and Takenawa, T. (2001). Role of the ENTH domain in phosphatidylinositol-4,5-bisphosphate binding and endocytosis. Science 291, 1047–1051.

Jackson, L.P., Kelly, B.T., McCoy, A.J., Gaffry, T., James, L.C., Collins, B.M., Honing, S., Evans, P.R., and Owen, D.J. (2010). A large-scale conformational change couples membrane recruitment to cargo binding in the AP2 clathrin adaptor complex. Cell 141, 1220–1229.

Jaillais, Y., Fobis-Loisy, I., Miege, C., and Gaude, T. (2008). Evidence for a sorting endosome in Arabidopsis root cells. Plant J 53, 237–247.

Johnson, A., Dahhan, D., Gnyliukh, N., Kaufmann, W., Zheden, V., Costanzo, T., Mahou, P., Hrtyan, M., Wie, J., Aguilera-Servin, J., van Damme, D., Beaurepaire E., Loose, M., Bednarek, S., Friml, J. (2021) The TPLATE complex mediates membrane bending during plant clathrin-mediated endocytosis. bioRxiv 10.1101/2021.04.26.441441.

Jolliffe, N.A., Brown, J.C., Neumann, U., Vicre, M., Bachi, A., Hawes, C., Ceriotti, A., Roberts, L.M., and Frigerio, L. (2004). Transport of ricin and 2S albumin precursors to the storage vacuoles of Ricinus communis endosperm involves the Golgi and VSR-like receptors. Plant J 39, 821–833.

Jung, J.Y., Lee, D.W., Ryu, S.B., Hwang, I., and Schachtman, D.P. (2017). SCYL2 Genes Are Involved in Clathrin-Mediated Vesicle Trafficking and Essential for Plant Growth. Plant Physiol 175, 194–209.

Jurgens, G., Park, M., Richter, S., Touihri, S., Krause, C., El Kasmi, F., and Mayer, U. (2015). Plant cytokinesis: a tale of membrane traffic and fusion. Biochem Soc Trans 43, 73–78.

Kaksonen, M., and Roux, A. (2018). Mechanisms of clathrin-mediated endocytosis. Nat Rev Mol Cell Biol. 19, 313–326.

Kang, B.H., Nielsen, E., Preuss, M.L., Mastronarde, D., and Staehelin, L.A. (2011). Electron tomography of RabA4b- and PI-4Kbeta1-labeled trans Golgi network compartments in Arabidopsis. Traffic 12, 313–329.

Kang, H., and Hwang, I. (2014). Vacuolar Sorting Receptor-Mediated Trafficking of Soluble Vacuolar Proteins in Plant Cells. Plants (Basel) 3, 392–408.

Keen, J.H. (1987). Clathrin assembly proteins: affinity purification and a model for coat assembly. J Cell Biol 105, 1989–1998.

Kim, S.Y., Xu, Z.Y., Song, K., Kim, D.H., Kang, H., Reichardt, I., Sohn, E.J., Friml, J., Juergens, G., and Hwang, I. (2013). Adaptor protein complex 2-mediated endocytosis is crucial for male reproductive organ development in Arabidopsis. Plant Cell 25, 2970–2985.

Kirsch, T., Paris, N., Butler, J.M., Beevers, L., and Rogers, J.C. (1994). Purification and initial characterization of a potential plant vacuolar targeting receptor. Proc Natl Acad Sci U S A 91, 3403–3407.

Kitakura, S., Vanneste, S., Robert, S., Lofke, C., Teichmann, T., Tanaka, H., and Friml, J. (2011). Clathrin mediates endocytosis and polar distribution of PIN auxin transporters in Arabidopsis. Plant Cell 23, 1920–1931.

Konopka, C.A., and Bednarek, S.Y. (2008). Comparison of the dynamics and functional redundancy of the Arabidopsis dynamin-related isoforms DRP1A and DRP1C during plant development. Plant Physiol 147, 1590–1602.

Konopka, C.A., Backues, S.K., and Bednarek, S.Y. (2008). Dynamics of Arabidopsis dynamin-related protein 1C and a clathrin light chain at the plasma membrane. Plant Cell 20, 1363–1380.

Kost, B., Lemichez, E., Spielhofer, P., Hong, Y., Tolias, K., Carpenter, C., and Chua, N.H. (1999). Rac homologues and compartmentalized phosphatidylinositol 4, 5-bisphosphate act in a common pathway to regulate polar pollen tube growth. J Cell Biol 145, 317–330.

Kunzl, F., Fruholz, S., Fassler, F., Li, B., and Pimpl, P. (2016). Receptor-mediated sorting of soluble vacuolar proteins ends at the trans-Golgi network/early endosome. Nat Plants 2, 16017.

Lam, B.C., Sage, T.L., Bianchi, F., and Blumwald, E. (2001). Role of SH3 domain-containing proteins in clathrin-mediated vesicle trafficking in Arabidopsis. Plant Cell 13, 2499–2512.

Lauber, M.H., Waizenegger, I., Steinmann, T., Schwarz, H., Mayer, U., Hwang, I., Lukowitz, W., and Jurgens, G. (1997). The Arabidopsis KNOLLE protein is a cytokinesis-specific syntaxin. Journal of Cell Biology 139, 1485–1493.

Lee, G.J., Kim, H., Kang, H., Jang, M., Lee, D.W., Lee, S., and Hwang, I. (2007). EpsinR2 interacts with clathrin, adaptor protein-3, AtVTI12, and phosphatidylinositol-3-phosphate. Implications for EpsinR2 function in protein trafficking in plant cells. Plant Physiol 143, 1561–1575.

Li, R., Rodriguez-Furlan, C., Wang, J., van de Ven, W., Gao, T., Raikhel, N., Hicks, G. (2017). Different endomembrane trafficking pathways establish apical and basal polarities. Plant Cell 29: 90–108.

Lipatova, Z., Hain, A.U., Nazarko, V.Y., and Segev, N. (2015). Ypt/Rab GTPases: principles learned from yeast. Crit Rev Biochem Mol Biol 50, 203–211.

Lundgren, D.H., Hwang, S.I., Wu, L., and Han, D.K. (2010). Role of spectral counting in quantitative proteomics. Expert Rev Proteomics 7, 39–53.

Luo, Y., Scholl, S., Doering, A., Zhang, Y., Irani, N.G., Rubbo, S.D., Neumetzler, L., Krishnamoorthy, P., Van Houtte, I., Mylle, E., Bischoff, V., Vernhettes, S., Winne, J., Friml, J., Stierhof, Y.D., Schumacher, K., Persson, S., and Russinova, E. (2015). V-ATPase activity in the TGN/EE is required for exocytosis and recycling in Arabidopsis. Nat Plants 1, 15094.

Ma, C., Li, W., Xu, Y., and Rizo, J. (2011). Munc13 mediates the transition from the closed syntaxin- Munc18 complex to the SNARE complex. Nat Struct Mol Biol 18, 542–549.

Martins, S., Dohmann, E.M., Cayrel, A., Johnson, A., Fischer, W., Pojer, F., Satiat-Jeunemaitre, B., Jaillais, Y., Chory, J., Geldner, N., and Vert, G. (2015). Internalization and vacuolar targeting of the brassinosteroid hormone receptor BRI1 are regulated by ubiquitination. Nat Commun 6, 6151.

Masclaux, F.G., Galaud, J.P., and Pont-Lezica, R. (2005). The riddle of the plant vacuolar sorting receptors. Protoplasma 226, 103–108.

Mattera, R., Guardia, C.M., Sidhu, S.S., and Bonifacino, J.S. (2015). Bivalent Motif-Ear Interactions Mediate the Association of the Accessory Protein Tepsin with the AP-4 Adaptor Complex. J Biol Chem 290, 30736–30749.

Mayers, J.R., Hu, T., Wang, C., Cardenas, J.J., Tan, Y., Pan, J., and Bednarek, S.Y. (2017). SCD1 and SCD2 Form a Complex That Functions with the Exocyst and RabE1 in Exocytosis and Cytokinesis. Plant Cell 29, 2610–2625.

McGough, I.J., and Cullen, P.J. (2013). Clathrin is not required for SNX-BAR-retromer-mediated carrier formation. J Cell Sci 126, 45–52.

McMichael, C.M., and Bednarek, S.Y. (2013). Cytoskeletal and membrane dynamics during higher plant cytokinesis. New Phytol 197, 1039–1057.

McMichael, C.M., Reynolds, G.D., Koch, L.M., Wang, C., Jiang, N., Nadeau, J., Sack, F.D., Gelderman, M.B., Pan, J., and Bednarek, S.Y. (2013). Mediation of clathrin-dependent trafficking during cytokinesis and cell expansion by Arabidopsis stomatal cytokinesis defective proteins. Plant Cell 25, 3910–3925.

Mosesso, N., Blaske, T., Nagel, M.K., Laumann, M., and Isono, E. (2019). Preparation of Clathrin-Coated Vesicles From Arabidopsis thaliana Seedlings. Front Plant Sci 9, 1972.

Mravec, J., Petrasek, J., Li N., Boeren, S., Karlova, R., Kitakura, S., Parezova, M., Naramoto, S., Nodzynski, T., Dhonukshe, P., Bednarek, S., Zazimalova, E., de Vries, S., Friml, J. (2011) Cell plate restricted association of DRP1A and PIN proteins is required for cell polarity establishment in Arabidopsis. Current Biology. 21, 1055–1060.

Murashige, T., and Skoog, F. (1962). A Revised Medium for Rapid Growth and Bio Assays with Tobacco Tissue Cultures. Physiol Plantarum 15, 473–497.

Nagaraj, N., Wisniewski, J., Geiger, T., Cox, J., Kircher, M., Kelso, J., Paabo, S., Mann, M. (2011) Deep proteome and transcriptome mapping of a human cancer cell line. Molecular Systems Biology 7: 1–8.

Nagel, M., Kalinowska, K., Vogel, K., Reynolds, G.D., Wu, Z., Anzenberger, F., Ichikawa, M., Tsutsumi, C., Sato, M.H., Kuster, B., Bednarek, S.Y., and Isono, E. (2017). Arabidopsis SH3P2 is an ubiquitin-binding protein that functions together with ESCRT-I and the deubiquitylating enzyme AMSH3. Proc Natl Acad Sci U S A.

Narasimhan, M., Johnson, A., Prizak, R., Kaufmann, W., Tan, S., Casillas-Perez, B., Friml, J. (2020). Evolutionarily unique mechanistic framework of clathrin-mediated endocytosis in plants. eLife. 9: 1–30.

Nielsen, E., Cheung, A.Y., and Ueda, T. (2008). The regulatory RAB and ARF GTPases for vesicular trafficking. Plant Physiol 147, 1516–1526.

Niemes, S., Langhans, M., Viotti, C., Scheuring, D., San Wan Yan, M., Jiang, L., Hillmer, S., Robinson, D.G., and Pimpl, P. (2010). Retromer recycles vacuolar sorting receptors from the trans-Golgi network. Plant J 61, 107–121.

Nishimura, K., Matsunami, E., Yoshida, S., Kohata, S., Yamauchi, J., Jisaka, M., Nagaya, T., Yokota, K., and Nakagawa, T. (2016). The tyrosine-sorting motif of the vacuolar sorting receptor VSR4 from Arabidopsis thaliana, which is involved in the interaction between VSR4 and AP1M2, mu1- adaptin type 2 of clathrin adaptor complex 1 subunits, participates in the post-Golgi sorting of VSR4. Biosci Biotechnol Biochem 80, 694–705.

Okekeogbu, I.O., Pattathil, S., Gonzalez Fernandez-Nino, S.M., Aryal, U.K., Penning, B.W., Lao, J., Heazlewood, J., Hahn, M.G., McCann, M.C., Carpita, N.C. (2019) Glycome and proteome components of Golgi membranes are common between two angiosperms with distinct cell-wall structures. Plant Cell 31,1094–1112.

Orci, L., Ravazzola, M., Amherdt, M., Louvard, D., and Perrelet, A. (1985). Clathrin-immunoreactive sites in the Golgi apparatus are concentrated at the trans pole in polypeptide hormone-secreting cells. Proc Natl Acad Sci U S A 82, 5385–5389.

Orr, R., Fury, F., Warner, E., Agar, E., Garbarino, J., Cabral, S., Dubuque, M., Butt, A., Munson, M., Vidali, L. (2021) Rab-E and its interaction with myosin XI are essential for polarised cell growth. New Phytologist 229: 1924–1936.

Papanikou, E., Day, K.J., Austin, J., and Glick, B.S. (2015). COPI selectively drives maturation of the early Golgi. Elife 4.

Park, M., Song, K., Reichardt, I., Kim, H., Mayer, U., Stierhof, Y.D., Hwang, I., and Jurgens, G. (2013). Arabidopsis mu-adaptin subunit AP1M of adaptor protein complex 1 mediates late secretory and vacuolar traffic and is required for growth. Proc Natl Acad Sci U S A 110, 10318–10323.

Parsons, H.T., Stevens, T.J., McFarlane, H.E., Vidal-Melgosa, S., Griss, J., Lawrence, N., Butler, R., Sousa, M.M.L., Salemi, M., Willats, W.G.T., Petzold, C.J., Heazlewood, J., Lilley, K. (2019) Separating Golgi Proteins from *Cis* to *Trans* Reveals Underlying Properties of Cisternal Localization. Plant Cell 31, 2010–2034.

Parsons, H.T., Christiansen, K., Knierim, B., Carroll, A., Ito, J., Batth, T.S., Smith-Moritz, A.M., Morrison, S., McInerney, P., Hadi, M.Z., Auer, M., Mukhopadhyay, A., Petzold, C.J., Scheller, H.V., Loque, D., and Heazlewood, J.L. (2012). Isolation and proteomic characterization of the Arabidopsis Golgi defines functional and novel components involved in plant cell wall biosynthesis. Plant Physiol 159, 12–26.

Pearse, B.M., and Robinson, M.S. (1984). Purification and properties of 100-kd proteins from coated vesicles and their reconstitution with clathrin. EMBO J 3, 1951–1957.

Perkins, D.N., Pappin, D.J., Creasy, D.M., and Cottrell, J.S. (1999). Probability-based protein identification by searching sequence databases using mass spectrometry data. Electrophoresis 20, 3551–3567.

Pinheiro, H., Samalova, M., Geldner, N., Chory, J., Martinez, A., and Moore, I. (2009). Genetic evidence that the higher plant Rab-D1 and Rab-D2 GTPases exhibit distinct but overlapping interactions in the early secretory pathway. J Cell Sci 122, 3749–3758.

Pourcher, M., Santambrogio, M., Thazar, N., Thierry, A.M., Fobis-Loisy, I., Miege, C., Jaillais, Y., and Gaude, T. (2010). Analyses of sorting nexins reveal distinct retromer-subcomplex functions in development and protein sorting in Arabidopsis thaliana. Plant Cell 22, 3980–3991.

Reichardt, I., Stierhof, Y., Mayer, U., Richter, S., Schwarz, H., Schumacher, K., Jurgens, G. (2007) Plant cytokinesis requires do novo secretory trafficking but not endocytosis. Current Biology 17, 2047–2053.

Reynolds, G.D., August, B., and Bednarek, S.Y. (2014). Preparation of enriched plant clathrin-coated vesicles by differential and density gradient centrifugation. Methods Mol Biol 1209, 163–177.

Reynolds, G.D., Wang, C., Pan, J., and Bednarek, S.Y. (2018). Inroads into Internalization: Five Years of Endocytic Exploration. Plant Physiol 176, 208–218.

Robinson, D.G. (2018). Retromer and VSR Recycling: A Red Herring? Plant Physiol 176, 483–484.

Robinson, D.G., and Neuhaus, J.M. (2016). Receptor-mediated sorting of soluble vacuolar proteins: myths, facts, and a new model. J Exp Bot 67, 4435–4449.

Robinson, D.G., and Pimpl, P. (2014) Receptor-mediated transport of vacuolar proteins: a critical analysis and a new model. Protoplasma 251, 247–264.

Robinson, M.S. (2015). Forty Years of Clathrin-coated Vesicles. Traffic 16, 1210–1238.

Rosquete, M.R., Davis, D.J., and Drakakaki, G. (2018). The Plant Trans-Golgi Network: Not Just a Matter of Distinction. Plant Physiol 176, 187–198.

Roth, T.F., and Porter, K.R. (1964). Yolk Protein Uptake in the Oocyte of the Mosquito Aedes Aegypti. L. J Cell Biol 20, 313–332.

Rueden, C.T., Schindelin, J., Hiner, M.C., DeZonia, B.E., Walter, A.E., Arena, E.T., and Eliceiri, K.W. (2017). ImageJ2: ImageJ for the next generation of scientific image data. BMC Bioinformatics 18, 529.

Rutherford, S., and Moore, I. (2002). The Arabidopsis Rab GTPase family: another enigma variation. Curr Opin Plant Biol 5, 518–528.

Sanchez-Rodriguez, C., Shi, Y., Kesten, C., Zhang, D., Sancho-Andres, G., Ivakov, A., Lampugnani, E.R., Sklodowski, K., Fujimoto, M., Nakano, A., Bacic, A., Wallace, I.S., Ueda, T., Van Damme, D., Zhou, Y., and Persson, S. (2018). The Cellulose Synthases Are Cargo of the TPLATE Adaptor Complex. Mol Plant 11, 346–349.

Sanger, A., Hirst, J., Davies, A.K., Robinson, M.S. (2019) Adaptor protein complexes and disease at a glance. Journal of Cell Science. 132, 1–8.

Sauer, M., Delgadillo, M.O., Zouhar, J., Reynolds, G.D., Pennington, J.G., Jiang, L., Liljegren, S.J., Stierhof, Y.D., De Jaeger, G., Otegui, M.S., Bednarek, S.Y., and Rojo, E. (2013). MTV1 and MTV4 encode plant-specific ENTH and ARF GAP proteins that mediate clathrin-dependent trafficking of vacuolar cargo from the trans-Golgi network. Plant Cell 25, 2217–2235.

Schindelin, J., Arganda-Carreras, I., Frise, E., Kaynig, V., Longair, M., Pietzsch, T., Preibisch, S., Rueden, C., Saalfeld, S., Schmid, B., Tinevez, J.Y., White, D.J., Hartenstein, V., Eliceiri, K., Tomancak, P., and Cardona, A. (2012). Fiji: an open-source platform for biological-image analysis. Nat Methods 9, 676–682.

Schwanhausser, B., Busse, D., Li, N., Dittmar, G., Schuchhardt, J., Wolf, J., Chen, W., Selbach, M. (2011) Global quantification of mammalian gene expression control. Nature. 473: 337–342.

Schwihla, M., and Korbei, B. (2020) The beginning of the end: initial steps in the degradation of plasma membrane proteins. Front Plant Sci. 11, 680.

Shevchenko, A., Wilm, M., Vorm, O., and Mann, M. (1996). Mass spectrometric sequencing of proteins silver-stained polyacrylamide gels. Anal Chem 68, 850–858.

Shimada, T., Fuji, K., Tamura, K., Kondo, M., Nishimura, M., Hara-Nishimura, I. (2003) Vacuolar sorting receptor for seed storage proteins in *Arabidopsis thaliana*. PNAS 100, 16095–16100.

Shimizu, Y., Takagi, J., Ito, E., Ito, Y., Ebine, K., Komatsu, Y., Goto Y., Sato, M., Toyooka, K., Ueda, T., Kurokawa, K., Uemura, T., Nakano, A. (2021). Cargo sorting zones in the *trans*-Golgi network visualized by super-resolution confocal live imaging microscopy in plants. Nature Communications. 12, 1–14.

Simpson, F., Peden, A.A., Christopoulou, L., and Robinson, M.S. (1997). Characterization of the adaptor-related protein complex, AP-3. J Cell Biol 137, 835–845.

Smith, J.M., Leslie, M.E., Robinson, S.J., Korasick, D.A., Zhang, T., Backues, S.K., Cornish, P.V., Koo, A.J., Bednarek, S.Y., and Heese, A. (2014). Loss of Arabidopsis thaliana Dynamin-Related Protein 2B reveals separation of innate immune signaling pathways. PLoS Pathog 10, e1004578.

Song, J., Lee, M.H., Lee, G.J., Yoo, C.M., and Hwang, I. (2006). Arabidopsis EPSIN1 plays an important role in vacuolar trafficking of soluble cargo proteins in plant cells via interactions with clathrin, AP-1, VTI11, and VSR1. Plant Cell 18, 2258–2274.

Sosa, R.T., Weber, M.M., Wen, Y., and O’Halloran, T.J. (2012). A single beta adaptin contributes to AP1 and AP2 complexes and clathrin function in Dictyostelium. Traffic 13, 305–316.

Speth, E.B., Imboden, L., Hauck, P., and He, S.Y. (2009). Subcellular localization and functional analysis of the Arabidopsis GTPase RabE. Plant Physiol 149, 1824–1837.

Stolc, V., Samanta, M.P., Tongprasit, W., Sethi, H., Liang, S., Nelson, D.C., Hegeman, A., Nelson, C., Rancour, D., Bednarek, S., Ulrich, E.L., Zhao, Q., Wrobel, R.L., Newman, C.S., Fox, B.G., Phillips, G.N., Jr., Markley, J.L., and Sussman, M.R. (2005). Identification of transcribed sequences in Arabidopsis thaliana by using high-resolution genome tiling arrays. Proc Natl Acad Sci U S A 102, 4453–4458.

Suwastika, I.N., Uemura, T., Shiina, T., Sato, M.H., and Takeyasu, K. (2008). SYP71, a plant-specific Qc-SNARE protein, reveals dual localization to the plasma membrane and the endoplasmic reticulum in Arabidopsis. Cell Struct Funct 33, 185–192.

Takamori, S., Holt, M., Stenius, K., Lemke, E., Gronborg, M., Riedel, D., Urlaub, H., Schenck, S., Brugger, B., Ringler, P., Muller, S., Rammer, B., Grater, F., Hub, J., De Groot, B., Mieskes, G., Moriyama, Y., Klingauf, J., Grubmuller, H., Heuser, J., Wieland, F., Jahn, R. (2006) Molecular anatomy of a trafficking organelle. Cell. 127: 831–846.

Takemoto, K., Ebine, K., Askani, J.C., Kruger, F., Gonzalez, Z.A., Ito, E., Goh, T., Schumacher, K., Nakano, A., and Ueda, T. (2018). Distinct sets of tethering complexes, SNARE complexes, and Rab GTPases mediate membrane fusion at the vacuole in Arabidopsis. Proc Natl Acad Sci U S A 115, E2457–E2466.

Taylor, M., Perraid, D., Merrifield, C. (2011). A high precision survey of the molecular dynamics of mammalian clathrin-mediated endocytosis. PLoS Biology 9: e1000604.

Teh, O.K., Shimono, Y., Shirakawa, M., Fukao, Y., Tamura, K., Shimada, T. (2013) The AP-1 mu adaptin is required for KNOLLE localization at the cell plate to mediate cytokinesis in Arabidopsis. Plant cell Physiology 43, 838–847.

Tong, P., Kornfeld, S. (1989) Ligand interactions of the cation-dependent mannose 6-phosphate receptor: comparison with the cation-independent mannose 6-phosphate receptor. J Biol Chem 264, 7970–7975.

Uemura, T., Sato, M.H., and Takeyasu, K. (2005). The longin domain regulates subcellular targeting of VAMP7 in Arabidopsis thaliana. FEBS Lett 579, 2842–2846.

Uemura, T., Kim, H., Saito, C., Ebine, K., Ueda, T., Schulze-Lefert, P., and Nakano, A. (2012). Qa-SNAREs localized to the trans-Golgi network regulate multiple transport pathways and extracellular disease resistance in plants. Proc Natl Acad Sci U S A 109, 1784–1789.

Vallis, Y., Wigge, P., Marks, B., Evans, P.R., and McMahon, H.T. (1999). Importance of the pleckstrin homology domain of dynamin in clathrin-mediated endocytosis. Curr Biol 9, 257–260.

Van Damme, D., Coutuer, S., De Rycke, R., Bouget, F., Inzé, D., Geelen, D. (2006) Somatic cytokinesis and pollen maturation in *Arabidopsis* depend on TPLATE, which has domains similar to coat proteins. Plant Cell, 18, 3502–3518.

Viotti, C., Bubeck, J., Stierhof, Y.D., Krebs, M., Langhans, M., van den Berg, W., van Dongen, W., Richter, S., Geldner, N., Takano, J., Jurgens, G., de Vries, S.C., Robinson, D.G., and Schumacher, K. (2010). Endocytic and secretory traffic in Arabidopsis merge in the trans-Golgi network/early endosome, an independent and highly dynamic organelle. Plant Cell 22, 1344–1357.

Vollmer, A.H., Youssef, N.N., and DeWald, D.B. (2011). Unique cell wall abnormalities in the putative phosphoinositide phosphatase mutant AtSAC9. Planta 234, 993–1005.

Wang, C., Yan, X., Chen, Q., Jiang, N., Fu, W., Ma, B., Liu, J., Li, C., Bednarek, S.Y., and Pan, J. (2013). Clathrin light chains regulate clathrin-mediated trafficking, auxin signaling, and development in Arabidopsis. Plant Cell 25, 499–516.

Wang, C., Hu, T., Yan, X., Meng, T., Wang, Y., Wang, Q., Zhang, X., Gu, Y., Sanchez-Rodriguez, C., Gadeyne, A., Lin, J., Persson, S., Van Damme, D., Li, C., Bednarek, S.Y., and Pan, J. (2016). Differential Regulation of Clathrin and Its Adaptor Proteins during Membrane Recruitment for Endocytosis. Plant Physiol 171, 215–229.

Wang, X., Cai, Y., Wang, H., Zeng, Y., Zhuang, X., Li, B., and Jiang, L. (2014). Trans-Golgi network-located AP1 gamma adaptins mediate dileucine motif-directed vacuolar targeting in Arabidopsis. Plant Cell 26, 4102–4118.

Wang, J., Mylle, E., Johnson, A., Besbrugge, N., De Jaeger G., Friml, J., Pleskot, R., Van Damme, D. (2020) High temporal resolution reveals simultaneous plasma membrane recruitment of TPLATE complex subunits. Plant Physiology 183, 986–997.

Wang, J., Yperman, K., Grones, P., Jiang, Q., Dragwidge, J., Mylle, E., Mor, E., Nolf, J., Eeckhout, D., De Jaeger, G., De Rybel, B., Pleskot, R., Van Damme, D. (2021) Conditional destabilization of the TPLATE complex impairs endocytic internalization. PNAS 118, 1-

Williams, M.E., Torabinejad, J., Cohick, E., Parker, K., Drake, E.J., Thompson, J.E., Hortter, M., and Dewald, D.B. (2005). Mutations in the Arabidopsis phosphoinositide phosphatase gene SAC9 lead to overaccumulation of PtdIns(4,5)P2 and constitutive expression of the stress-response pathway. Plant Physiol 138, 686–700.

Wolfenstetter, S., Wirsching, P., Dotzauer, D., Schneider, S., Sauer, N. (2012). Routes to the tonoplast: the sorting of tonoplast transporters in *Arabidopsis* mesophyll protoplasts. Plant Cell 24: 215–232.

Wu, B., and Guo, W. (2015). The Exocyst at a Glance. Journal of Cell Science 128, 2957–2964.

Wu, M., and Wu, X. (2021). A kinetic view of clathrin assembly and endocytic cargo sorting. Current Opinion in Cell Biology. 71, 130–138.

Yan, X., Wang, Y., Xu, M., Dahhan, D., Liu, C., Zhang, Y., Lin, J., Bednarek, S., Pan, J. (2021). Cross-talk between clathrin-dependent post-golgi trafficking and clathrin-mediated endocytosis in Arabidopsis root cells. Plant Cell. 10.1093/plcell/koab180.

Yao, L., Janmey, P., Frigeri, L.G., Han, W., Fujita, J., Kawakami, Y., Apgar, J.R., and Kawakami, T. (1999). Pleckstrin homology domains interact with filamentous actin. J Biol Chem 274, 19752–19761.

Yperman, K., Papageorgiou, A., Merceron, R., De Munck, S., Bloch, Y., Eeckhout, D., Jiang, Q., Tack, P., Grigoryan, R., Evangelidis, T., Van Leene, J., Vincze, L., Vandenabeele, P., Vanhaecke, F., Potocky, M., De Jaeger, G., Savvides, S., Tripsianes, K., Pleskot, R., Van Damme, D. (2021) Distinct EH domains of the endocytic TPLATE complex confer lipid and protein binding. Nature Communications 12, 1–11.

Yun, H.S., and Kwon, C. (2017). Vesicle trafficking in plant immunity. Curr Opin Plant Biol 40, 34–42.

Zhang, Y., Persson, S., Hirst, J., Robinson, M.S., van Damme, D., and Sanchez-Rodriguez, C. (2015). Change your TPLATE, change your fate: plant CME and beyond. Trends Plant Sci 20, 41–48.

Zhao, Y., Yan, A., Feijo, J.A., Furutani, M., Takenawa, T., Hwang, I., Fu, Y., and Yang, Z. (2010). Phosphoinositides regulate clathrin-dependent endocytosis at the tip of pollen tubes in Arabidopsis and tobacco. Plant Cell 22, 4031–4044.

Zhou, Y., Yang, Y., Niu, Y., Fan, T., Qian, D., Luo, C., Shi, Y., Li, S., An, L., Xiang, Y. (2020) The tip-localized phosphatidylserine established by Arabidopsis ALA3 is crucial for Rab GTPase-mediated vesicle trafficking and pollen tube growth. Plant Cell 32: 3170–3187.

Zhu, X., Li, S., Pan, S., Xin, X., and Gu, Y. (2018). CSI1, PATROL1, and exocyst complex cooperate in delivery of cellulose synthase complexes to the plasma membrane. Proc Natl Acad Sci U S A 115, E3578–E3587.

Zouhar, J., and Sauer, M. (2014). Helping hands for budding prospects: ENTH/ANTH/VHS accessory proteins in endocytosis, vacuolar transport, and secretion. Plant Cell 26, 4232–4244.

Zouhar, J., Rojo, E., and Bassham, D.C. (2009). AtVPS45 is a positive regulator of the SYP41/SYP61/VTI12 SNARE complex involved in trafficking of vacuolar cargo. Plant Physiol 149, 1668–1678.

Zouhar, J., Munoz, A., and Rojo, E. (2010). Functional specialization within the vacuolar sorting receptor family: VSR1, VSR3 and VSR4 sort vacuolar storage cargo in seeds and vegetative tissues. Plant J 64, 577–588.

Zwiewka, M., Feraru, E., Moller, B., Hwang, I., Feraru, M.I., Kleine-Vehn, J., Weijers, D., and Friml, J. (2011). The AP-3 adaptor complex is required for vacuolar function in Arabidopsis. Cell Res 21, 1711–1722.

