## Supplemental Figures, Table, and References for "Proteomic Characterization of Isolated Arabidopsis Clathrin-Coated Vesicles Reveals Evolutionarily Conserved and Plant Specific Components"

### (A) Overview of CCV purification process

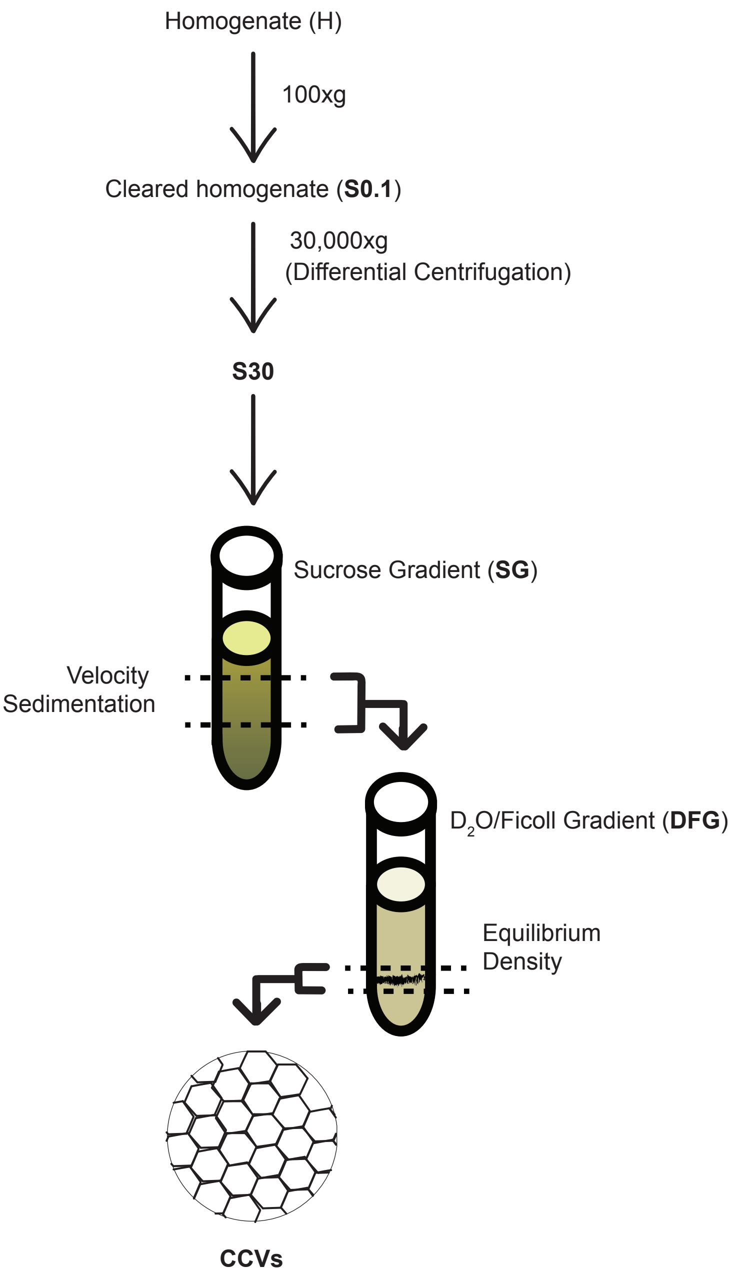

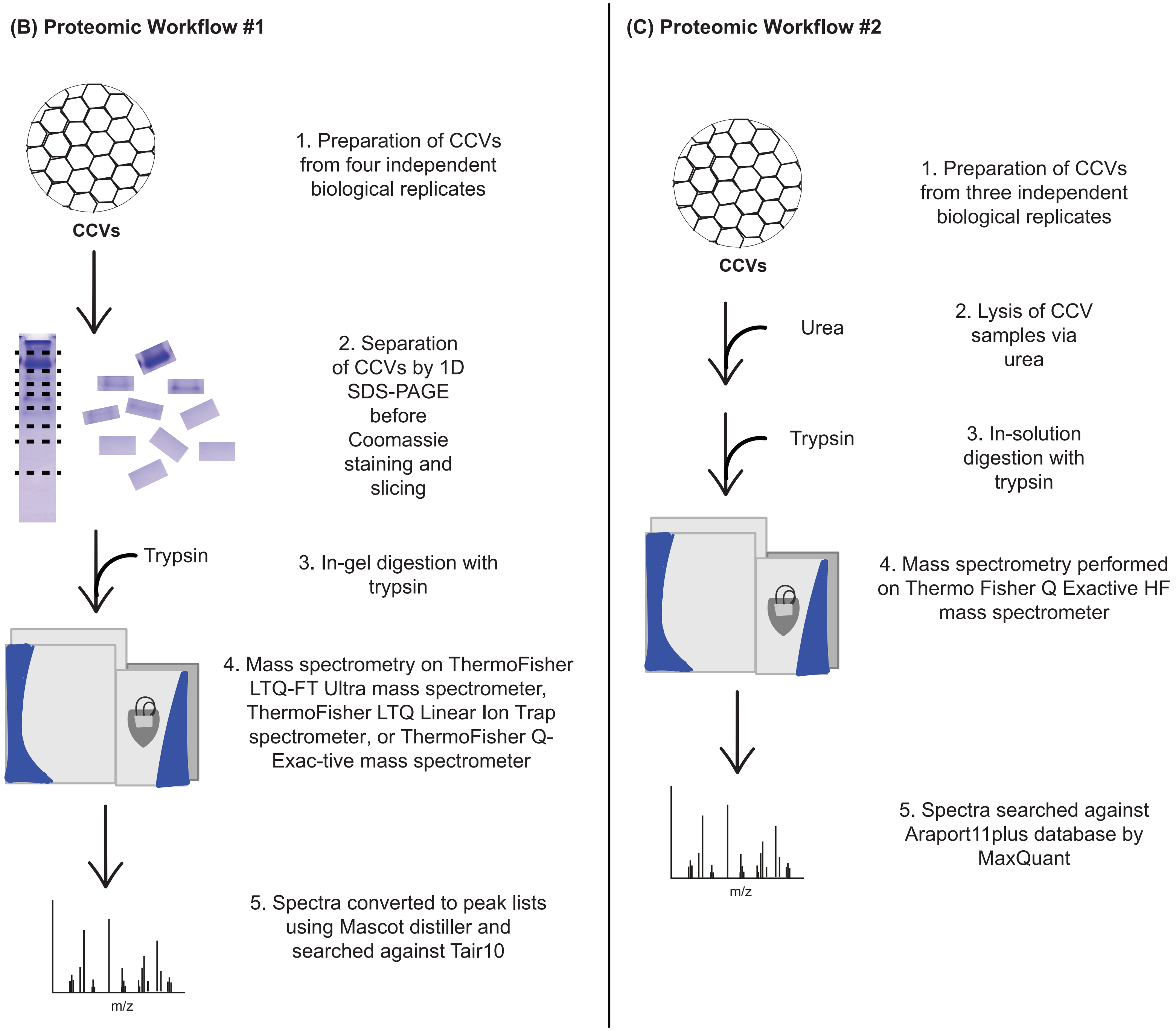
(B) Proteomic Workflow #1
CCVs
1. Preparation of CCVs from four independent biological replicates
2. Separation of CCVs by 1D SDS-PAGE before Coomassie staining and slicing
Trypsin
3. In-gel digestion with trypsin
4. Mass spectrometry on ThermoFisher LTQ-FT Ultra mass spectrometer, ThermoFisher LTQ Linear Ion Trap spectrometer, or ThermoFisher Q-Exac-tive mass spectrometer
m/z
5. Spectra converted to peak lists using Mascot distiller and searched against Tair10
(C) Proteomic Workflow #2
CCVs
1. Preparation of CCVs from three independent biological replicates
Urea
2. Lysis of CCV samples via urea
Trypsin
3. In-solution digestion with trypsin
4. Mass spectrometry performed on Thermo Fisher Q Exactive HF mass spectrometer
m/z
5. Spectra searched against Araport11plus database by MaxQuant

**(D) Proteomic Workflow #3**

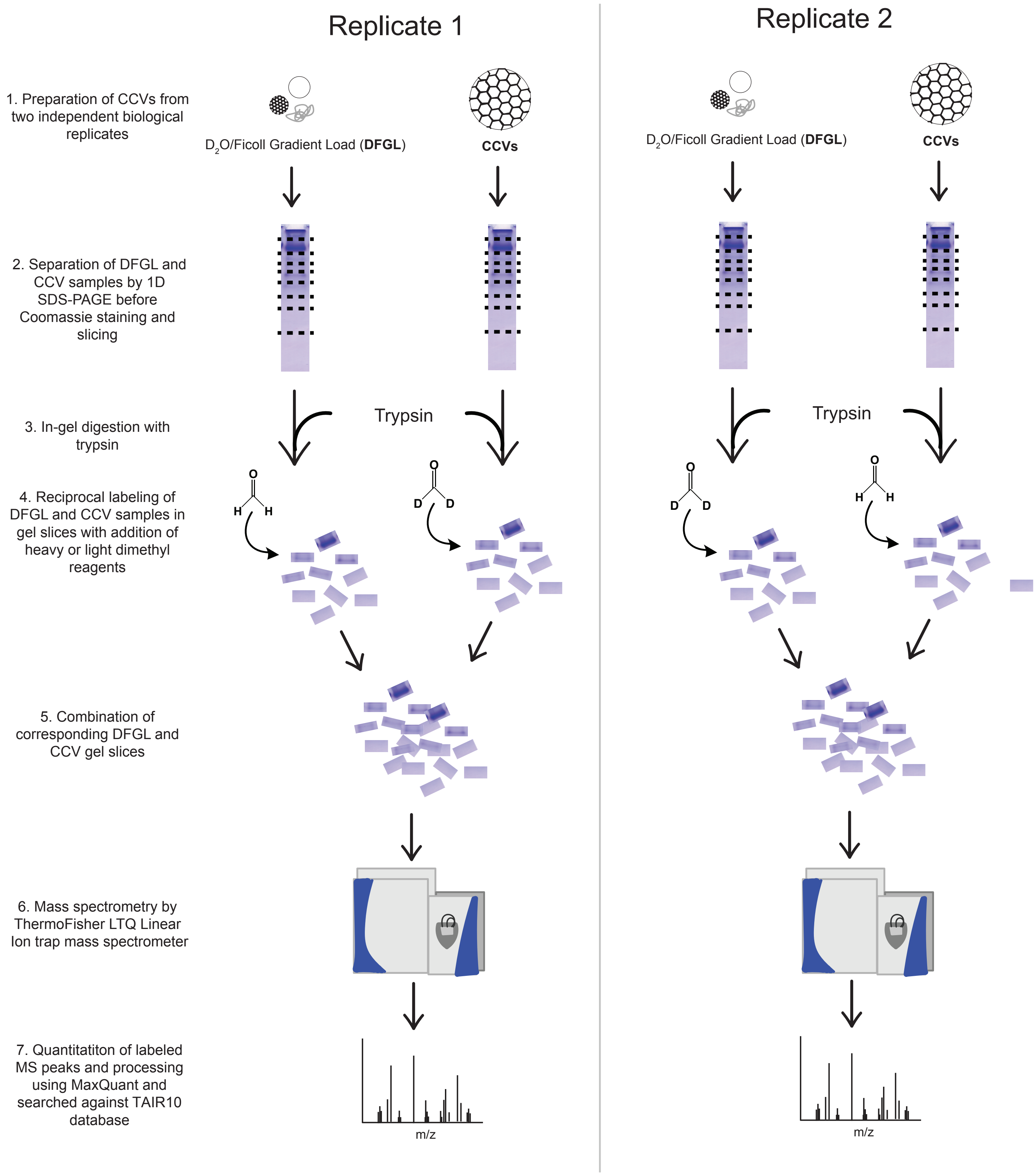

**Supplemental Figure 1. Schematics illustrating protocol for clathrin-coated vesicle purification and workflows detailing the CCV proteome.**

A. Abbreviated protocol for purification of clathrin-coated vesicles from Arabidopsis T87 suspension-cultured cells (adapted from Reynolds, et al. *MMB.* 2014). Steps in the purification process which were immunoblotted are highlighted in bold.

B. Proteomic workflow leading to collection of CCV proteomic dataset shown in Supplemental Dataset 1 and described in Materials & Methods.

C. Proteomic workflow leading to collection of CCV proteomic dataset shown in Supplemental Dataset 2 and described in Materials & Methods.

D. Proteomic workflow leading to collection of enrichment/depletion CCV proteomic dataset shown in Supplemental Dataset 3 and described in Materials & Methods.

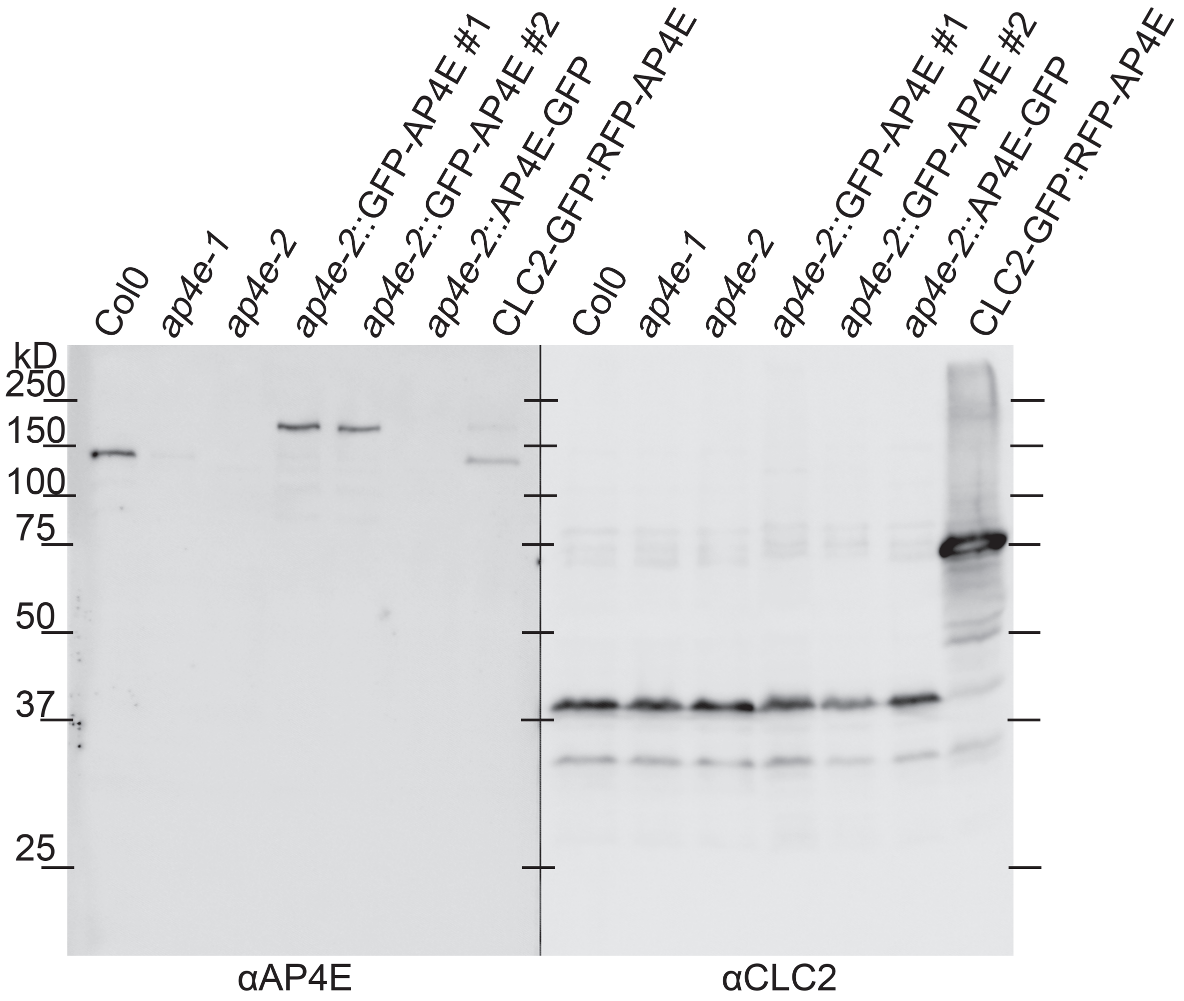


**Supplemental Figure 2: AP4E antibody is specific for the AP4E subunit**

Equal amounts of total protein extract from Arabidopsis seedlings were probed with αAP4E to demonstrate specificity and αCLC2 as a loading control. αAP4E recognizes the AP4E large subunit in wild type (Col0) and wild-type plants transformed with pUB10::RFP-AP4E and clc2pro::CLC2-GFP (CLC2-GFP:RFP-AP4E), but not in plants homozygous for either the ap4e-1 or ap4e-2 tDNA alleles (ap4e-1, ap4e-2, ap4e- 2::GFP-AP4E #1, ap4e-2::GFP-AP4E #2, ap4e-2:: AP4E-GFP). The larger GFP RFP fusion proteins were detected by αAP4E in plants transformed with pUBN::GFP-AP4E (two independent transformants, ap4e-2::GFP-AP4E #1, ap4e-2::GFP-AP4E #2) or pUB10::AP4e-GFP (ap4e-2:: AP4E-GFP).

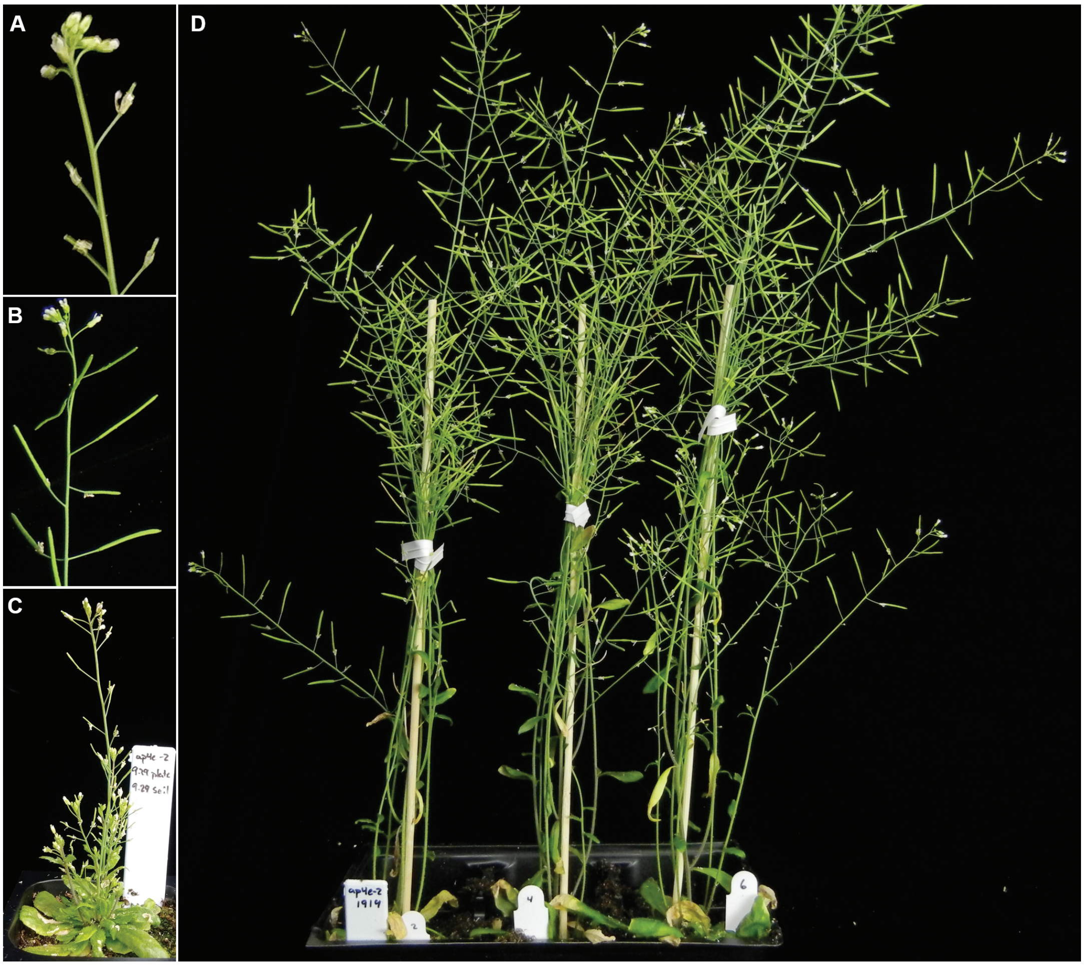


**Supplemental Figure 3: pUB10::GFP-AP4E is functional in vivo**

(A) ap4e-2 homozygotes display abnormal growth defects including stunted / aborted siliques.
(B) ap4e-2 homozygotes transformed with pUB10::GFP-AP4E display phenotypically wild-type inflorescences. (C) ap4e-2 homozygotes are overall dwarfed. (D) pUB10::GFP-AP4E rescues the dwarf phenotype of ap4e-2 homozygotes. Each plant in D derives from an independent transformant.

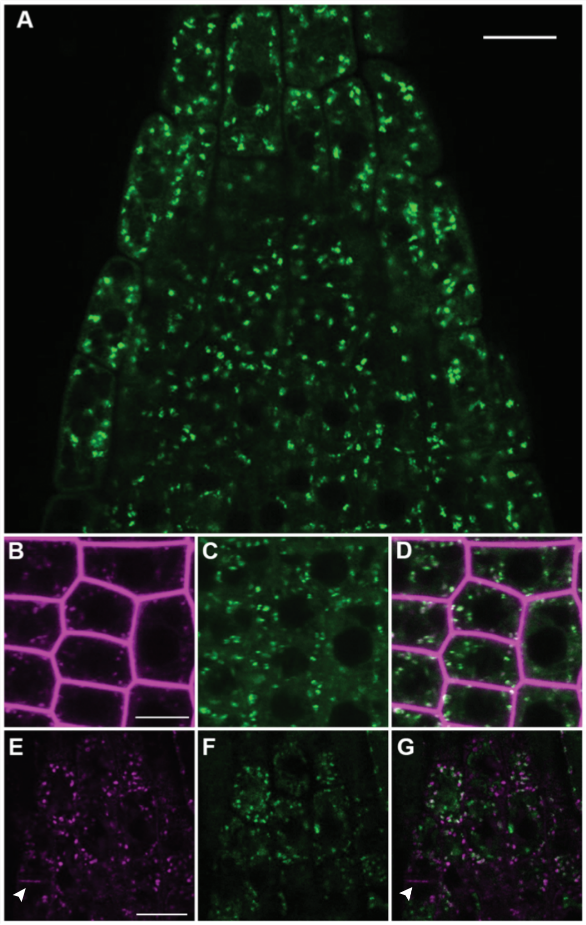


**Supplemental Figure 4. AP4E colocalizes with FM4-64 and clathrin at the TGN.**

(A) pUB10::GFP-AP4E expressed in homozygous ap4e-2 knockout lines has a primarily cytosolic and endomembrane localization in epidermal cells of seedling root tips.
(B) TGN/EE labeled with FM4-64 after 6 minute incubation colocalizes with pUB10::RFP-AP4E in homozygous ap4e-2 knockouts (C); (D) Merge.
(E) CLC2pro::CLC2-GFP and pUB10::RFP-AP4E (F) colocalize at the TGN but not at the PM nor cell plate; (G) Merge. Scale bars = 10 um. A white arrowhead indicates the cell plate.

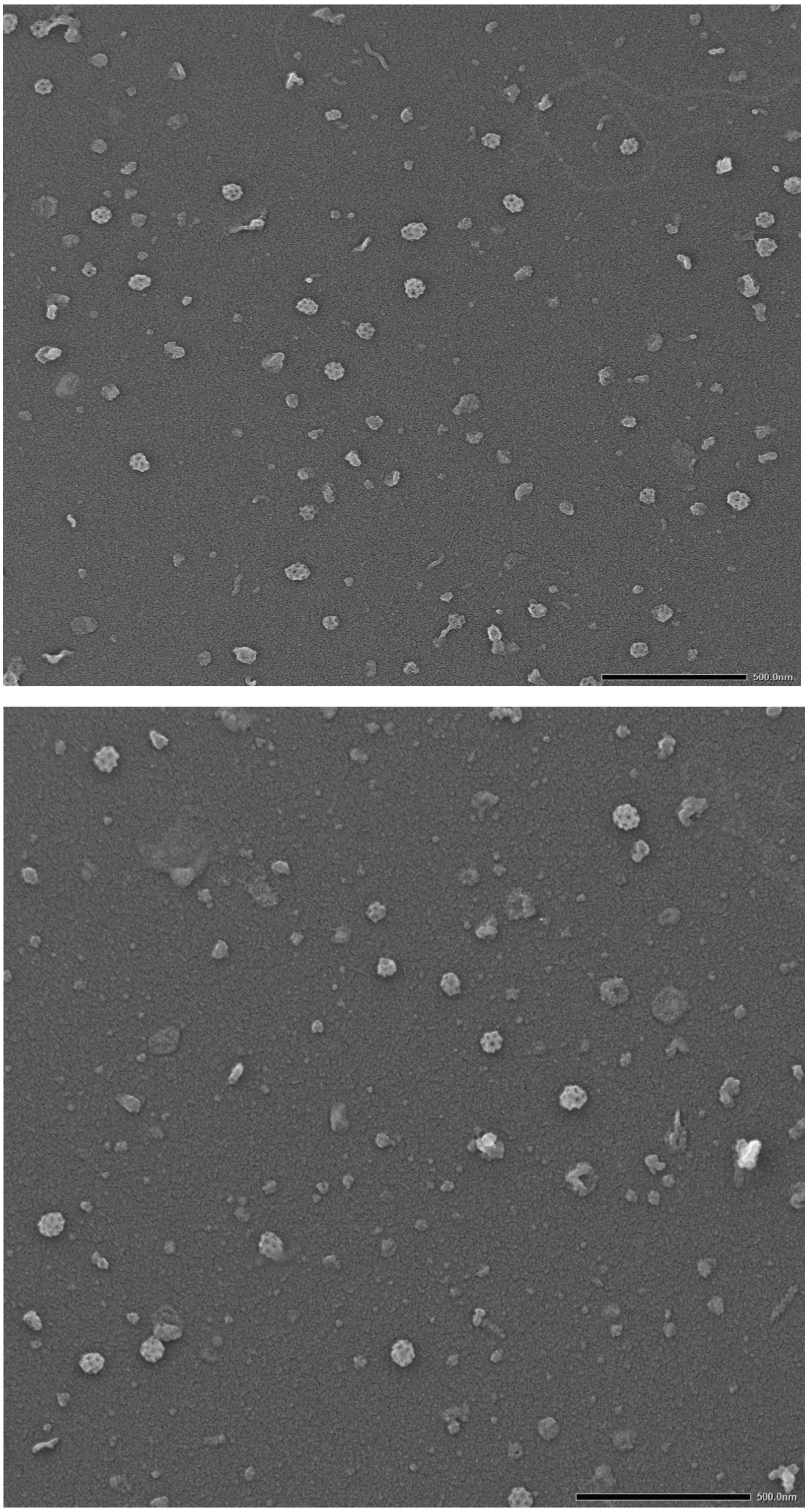


**Supplemental Figure 5: Transmission electron microscopy of CCVs.** Representative images collected by scanning transmission electron microscopy of CCVs purified by differential centrifugation from T87 suspension cultured Arabidopsis cells. Scale bar: 500 nm.

**Supplemental Table 1. Antibodies used in this study.** Information describing the antibodies used in this study are presented (clonality, dilution factor of the primary antibody, secondary antibody information, and citation).

| Antibodies | Clonality | Primary Dilution for Immunoblot | Secondary | Secondary Dilution | Citation in Manuscript |
|---|---|---|---|---|---|
| anti-CLC2 | polyclonal | 1:10000 | Rabbit | 1:5000 | Backues. Dissertation, University of Wisconsin-Madison. 2010. |
| anti-CHC | monoclonal | 1:1000 | Mouse | 1:5000 | Santa Cruz Biotechnology, sc-57684 |
| anti-AP2A | polyclonal | 1:500 | Rabbit | 1:5000 | Song et al. *Plant Physiology* 2012 |
| anti-AP1G | polyclonal | 1:2000 | Rabbit | 1:5000 | Park et al. *PNAS* 2013 |
| anti-AP2M | polyclonal | 1:250 | Rabbit | 1:5000 | Wang et al. *Plant Physiology.* 2016. |
| anti-KNOLLE | polyclonal | 1:1000 | Rabbit | 1:5000 | Rancour et al. *Plant Physiology.* 2002 |
| anti-TPLATE | polyclonal | 1:1000 | Rabbit | 1:5000 | DeJonghe, et al. *Nature Chem Biol.* 2019 |
| anti-SEC12 | polyclonal | 1:000 | Rabbit | 1:5000 | Bar-Peled and Raikhel. *Plant Physiology.* 1997. |
| anti-DRP1A | polyclonal | 1:500 | Rabbit | 1:5000 | Kang et al. *Plant Phys* 2001 |
| anti-DRP1C | polyclonal | 1:500 | Rabbit | 1:5000 | Kang et al. *Plant Journal.* 2003. |
| anti-DRP2 | polyclonal | 1:5000 | Rabbit | 1:5000 | Backues et al. *Plant Cell.* 2010. |
| anti-cFBPase | polyclonal | 1:5000 | Rabbit | 1:5000 | Agrisera, AS04 043 |
| anti-AP4E | Recombinant | 1:100 | Mouse | 1:5000 | This manuscript |

### Supplemental References:


**Axelos, M., Curie, C., Mazzolini, L., Bardet, C., and Lescure, B.** (1992). A Protocol for Transient Gene-Expression in Arabidopsis-Thaliana Protoplasts Isolated from Cell-Suspension Cultures. Plant Physiol Bioch **30,** 123-128.

**Backues, S.** (2010). Analysis of Arabidopsis DYNAMIN-RELATED PROTEIN 1 AND 2 (DRP1 AND DRP2) families. Doctoral dissertation, University of Wisconsin-Madison. ProQuest Dissertations Publishing.

**Bar-Peled, M., and Raikhel, N.V.** (1997). Characterization of AtSEC12 and AtSAR1. Proteins likely involved in endoplasmic reticulum and Golgi transport. Plant Physiol **114,** 315-324.

**Berardini, T.Z., Reiser, L., Li, D.H., Mezheritsky, Y., Muller, R., Strait, E., and Huala, E.** (2015). The Arabidopsis information resource: Making and mining the "gold standard" annotated reference plant genome. Genesis **53,** 474-485.

**Boersema, P.J., Raijmakers, R., Lemeer, S., Mohammed, S., and Heck, A.J.** (2009). Multiplex peptide stable isotope dimethyl labeling for quantitative proteomics. Nature protocols **4,** 484-494.

**Dejonghe, W., Sharma, I., Denoo, B., De Munck, S., Lu, Q., Mishev, K., Bulut, H., Mylle E., De Ryche, R., Vasileva, M., Savatin, D., Nerinckx, W., Staes, A., Drozdzecki, A., Audenaert, D., Yperman, K., Madder, A., Friml, J., Van Damme, D., Gevaert, K., Haucke, V., Savvides, S., Winne, J., Russinova, E.** (2019) Disruption of endocytosis through chemical inhibition of clathrin heavy chain function. Nature Chem Biology **15**, 641-649.

**Kang, B.H., Busse, J.S., Dickey, C., Rancour, D.M., Bednarek, S.Y.** (2001) The Arabidopsis cell plate-associated dynamin-like protein, ADL1Ap, is required for multiple stages of plant growth and development. Plant Physiology **126**, 47-68.

**Kang, B.H., Rancour, D.M., and Bednarek, S.Y.** (2003). The dynamin-like protein ADL1C is essential for plasma membrane maintenance during pollen maturation. Plant J **35,** 1-15.

**McMichael, C.M., Reynolds, G.D., Koch, L.M., Wang, C., Jiang, N., Nadeau, J., Sack, F.D., Gelderman, M.B., Pan, J., and Bednarek, S.Y.** (2013). Mediation of clathrin-dependent trafficking during cytokinesis and cell expansion by Arabidopsis stomatal cytokinesis defective proteins. The Plant cell **25,** 3910-3925.

**Park, M., Song, K., Reichardt, I., Kim, H., Mayer, U., Stierhof, Y.D., Hwang, I., and Jürgens, G.** (2013). Arabidopsis mu-adaptin subunit AP1M of adaptor protein complex 1 mediates late secretory and vacuolar traffic and is required for growth. Proceedings of the National Academy of Sciences of the United States of America **110,** 10318-10323.

**Rancour, D.M., Dickey, C.E., Park, S., and Bednarek, S.Y.** (2002). Characterization of AtCDC48. Evidence for multiple membrane fusion mechanisms at the plane of cell division in plants. Plant Physiol **130,** 1241-1253.

**Reynolds, G.D., August, B., and Bednarek, S.Y.** (2014). Preparation of enriched plant clathrin-coated vesicles by differential and density gradient centrifugation. Methods in molecular biology **1209,** 163-177.

**Song, K., Jang, M., Kim, S., Lee, G., Lee, G., Kim, D., Lee, Y., Cho, W., Hwang, I.** (2012) An A/ENTH domain-containing protein functions as an adaptor for clathrin-coated vesicles on the growing cell plate in Arabidopsis root cells. Plant Physiology **159**, 1013-1025.

**Wang, C., Hu, T., Yan, X., Meng, T., Wang, Y., Wang, Q., Zhang, X., Gu, Y., Sanchez-Rodriguez, C., Gadeyne, A., Lin, J., Persson, S., Van Damme, D., Li, C., Bednarek, S.Y., and Pan, J.** (2016). Differential Regulation of Clathrin and Its Adaptor Proteins during Membrane Recruitment for Endocytosis. Plant Physiol **171,** 215-229.
