## Supplemental Datasets for "Proteomic Characterization of Isolated Arabidopsis Clathrin-Coated Vesicles Reveals Evolutionarily Conserved and Plant Specific Components"

**Supplemental Dataset 1. Merged CCV proteomic datasets.** Merged data from CCV LC/MS-MS and dimethyl labeling experiments is presented using proteins identified in the LC/MS-MS of CCVs separated by SDS-PAGE before mass spectrometry as reference values. Columns: 1) TAIR accession numbers corresponding to the protein or protein representative of the group identified by LC/MS-MS of CCVs separated by SDS-PAGE (n = 3,548); 2) corresponding majority accession numbers identified in LC/MS-MS dataset of CCVs not separated by SDS-PAGE using column 1 accession numbers as reference; 3) TAIR annotation; 4) molecular weight of the gene product of the accession number in column 1 in kilodaltons; 5-8) abundance of protein as given by total spectral counts per independent biological replicate of the protein identified in column 1; 9) sum of total spectral counts across replicates; 10-11) ratio of protein abundance in CCV fraction relative to DFGL per replicate of the dimethyl labeling experiments; 12) average fold enrichment of protein in CCV fraction relative to DFGL purification step; 13-15) intensity based absolute quantitation (iBAQ) value of first protein listed in the majority accession numbers (Col. 2) of the corresponding protein groups in LC/MS-MS dataset of CCV proteins not separated by SDS-PAGE; 16) sum of iBAQ values across replicates.

| | | | | Total Spectral Counts (n = 3,548 proteins) | | | | | Fold Enrichment (CCV:DFGL) (n = 1,109 proteins in complete dataset) | | | iBAQ Values (n = 1,981 protein groups in complete dataset) | | | |
|---|---|---|---|---|---|---|---|---|---|---|---|---|---|---|---|
| Accession No. | Majority Accession No. | Annotation | MW | Rep 1 | Rep 2 | Rep 3 | Rep 4 | Sum | Rep 1 | Rep 2 | Average | Rep 1 | Rep 2 | Rep 3 | Sum |
| AT3G11130 | AT3G11130.1 | Clathrin, heavy chain | 193 kDa | 2144 | 3376 | 12637 | 9651 | 27808 | 7.34 | 4.88 | 6.11 | 6.7E+08 | 6.8E+08 | 2.6E+08 | 1.6E+09 |
| AT3G08530 | AT3G08530.1 | Clathrin, heavy chain | 193 kDa | 2101 | 3145 | 12447 | 9390 | 27083 | 6.82 | 3.67 | 5.25 | 6.7E+08 | 6.6E+08 | 2.4E+08 | 1.6E+09 |
| AT1G60070 | AT1G60070.1;AT1G60070.2 | Adaptor protein complex AP-1, gamma subunit | 95 kDa | 159 | 325 | 1935 | 1345 | 3764 | 5.06 | 6.20 | 5.63 | 9.5E+07 | 1.2E+08 | 2.9E+07 | 2.5E+08 |
| AT1G07920 | | GTP binding Elongation factor Tu family protein | 50 kDa | 112 | 231 | 1911 | 1351 | 3605 | 0.08 | 0.21 | 0.15 | | | | |
| AT1G23900 | AT1G23900.2;AT1G23900.1;AT1G23900.3 | gamma-adaptin 1 | 96 kDa | 201 | 372 | 1819 | 1353 | 3745 | 4.25 | 5.61 | 4.93 | 3.4E+07 | 4.5E+07 | 2.4E+07 | 1.0E+08 |
| AT4G11380 | AT4G11380.1;AT4G11380.2 | Adaptin family protein | 99 kDa | 214 | 204 | 1662 | 1692 | 3772 | 3.53 | 7.65 | 5.59 | 7.1E+07 | 5.9E+07 | 1.8E+07 | 1.5E+08 |
| AT4G23460 | AT4G23460.1;AT4G23460.2 | Adaptin family protein | 99 kDa | 201 | 186 | 1394 | 1506 | 3287 | 4.13 | 4.81 | 4.47 | 1.9E+07 | 1.7E+07 | 1.1E+07 | 4.7E+07 |
| AT2G40060 | AT2G40060.1 | Clathrin light chain protein | 29 kDa | 143 | 628 | 1011 | 408 | 2190 | 4.47 | 31.87 | 18.17 | 6.2E+08 | 1.0E+09 | 2.6E+08 | 1.9E+09 |
| AT2G47110 | | ubiquitin 6 | 18 kDa | 57 | 113 | 988 | 811 | 1969 | | 0.36 | 0.36 | | | | |
| AT1G31340 | | related to ubiquitin 1 | 17 kDa | 0 | 0 | 976 | 803 | 1779 | | 2.53 | 2.53 | | | | |
| AT1G78900 | AT1G78900.2;AT1G78900.1 | vacuolar ATP synthase subunit A | 69 kDa | 77 | 72 | 973 | 693 | 1815 | 0.12 | 0.46 | 0.29 | 1.8E+08 | 1.1E+08 | 2.0E+07 | 3.2E+08 |
| AT1G60780 | AT1G60780.1 | Clathrin adaptor complexes medium subunit family pr | 49 kDa | 96 | 258 | 908 | 793 | 2055 | 3.07 | 4.50 | 3.78 | 5.0E+07 | 6.6E+07 | 2.2E+07 | 1.4E+08 |
| AT2G20760 | AT2G20760.1 | Clathrin light chain protein | 37 kDa | 144 | 441 | 906 | 648 | 2139 | 3.57 | 12.77 | 8.17 | 8.9E+08 | 1.6E+09 | 2.5E+08 | 2.7E+09 |
| AT4G32285 | AT4G32285.2;AT4G32285.1 | ENTH/ANTH/VHS superfamily protein | 71 kDa | 157 | 248 | 863 | 705 | 1973 | 6.04 | 5.78 | 5.91 | 3.3E+07 | 4.9E+07 | 4.4E+06 | 8.6E+07 |
| AT5G22780 | AT5G22780.2;AT5G22780.1 | Adaptor protein complex AP-2, alpha subunit | 112 kDa | 155 | 220 | 862 | 814 | 2051 | 1.56 | 3.68 | 2.62 | 3.4E+06 | 2.7E+06 | 3.3E+05 | 6.4E+06 |
| AT2G18960 | AT2G18960.1;AT2G18960.3;AT2G18960.2 | H(+)-ATPase 1 | 104 kDa | 106 | 42 | 859 | 506 | 1513 | 1.61 | 0.44 | 1.03 | 4.5E+07 | 2.8E+07 | 3.9E+06 | 7.6E+07 |
| AT1G21660 | AT1G21660.1 | Chaperone DnaJ-domain superfamily protein | 57 kDa | 110 | 313 | 858 | 578 | 1859 | 2.56 | 4.98 | 3.77 | 7.2E+07 | 1.1E+08 | 2.4E+07 | 2.1E+08 |
| AT3G59770 | AT3G59770.1;AT3G59770.3;AT3G59770.2;AT3G59770.4 | sacI homology domain-containing protein / WW doma | 180 kDa | 55 | 117 | 840 | 540 | 1552 | 5.80 | 5.25 | 5.53 | 6.8E+06 | 8.0E+06 | 4.3E+05 | 1.5E+07 |
| AT4G20850 | AT4G20850.1 | tripeptidyl peptidase ii | 152 kDa | 16 | 7 | 832 | 594 | 1449 | 0.15 | 0.42 | 0.29 | 2.1E+08 | 3.5E+08 | 2.4E+05 | 5.6E+08 |
| AT5G22770 | AT5G22770.6;AT5G22770.5;AT5G22770.4;AT5G22770.3;AT5G22770.2;AT5G22770.1 | alpha-adaptin | 112 kDa | 155 | 216 | 823 | 779 | 1973 | | | | 1.5E+06 | 1.2E+06 | 5.4E+05 | 3.3E+06 |
| AT5G11710 | AT5G11710.3;AT5G11710.2;AT5G11710.1 | ENTH/VHS family protein | 61 kDa | 140 | 314 | 796 | 565 | 1815 | 3.31 | 6.66 | 4.98 | 1.1E+08 | 1.7E+08 | 3.3E+07 | 3.2E+08 |
| AT1G31730 | AT1G31730.1 | Adaptin family protein | 104 kDa | 175 | 260 | 779 | 528 | 1742 | 3.30 | 5.86 | 4.58 | 6.4E+07 | 8.7E+07 | 1.3E+07 | 1.6E+08 |
| AT4G30190 | AT4G30190.1;AT4G30190.2 | H(+)-ATPase 2 | 104 kDa | 94 | 22 | 743 | 458 | 1317 | | 0.16 | 0.16 | 2.5E+07 | 2.0E+07 | 1.6E+06 | 4.7E+07 |
| AT2G01820 | AT2G01820.1 | Leucine-rich repeat protein kinase family protein | 102 kDa | 73 | 61 | 690 | 546 | 1370 | 2.93 | 3.51 | 3.22 | 9.5E+07 | 7.4E+07 | 1.3E+07 | 1.8E+08 |
| AT2G25430 | AT2G25430.1 | epsin N-terminal homology (ENTH) domain-containing | 72 kDa | 100 | 147 | 672 | 668 | 1587 | 3.78 | 8.61 | 6.19 | 2.0E+07 | 3.0E+07 | 1.5E+06 | 5.2E+07 |
| AT3G16270 | AT3G16270.3;AT3G16270.2;AT3G16270.1 | ENTH/VHS family protein | 75 kDa | 96 | 244 | 666 | 488 | 1494 | | 6.74 | 6.74 | 1.7E+07 | 2.6E+07 | 9.6E+06 | 5.3E+07 |
| AT5G62690 | | tubulin beta chain 2 | 51 kDa | 117 | 429 | 616 | 623 | 1785 | | | | | | | |
| AT3G16760 | AT3G16760.1;AT3G16760.2 | Tetratricopeptide repeat (TPR)-like superfamily protein | 49 kDa | 93 | 451 | 590 | 644 | 1778 | 2.31 | 5.53 | 3.92 | 5.8E+07 | 1.1E+08 | 1.9E+07 | 1.8E+08 |
| AT3G51890 | AT3G51890.1 | Clathrin light chain protein | 29 kDa | 82 | 574 | 582 | 614 | 1852 | 11.07 | 6.49 | 8.78 | 1.4E+08 | 1.8E+08 | 2.0E+07 | 3.4E+08 |
| AT1G76030 | AT1G76030.1;AT1G20260.1 | ATPase, V1 complex, subunit B protein | 54 kDa | 56 | 97 | 579 | 602 | 1334 | | | | 3.4E+07 | 2.3E+07 | 3.0E+06 | 6.0E+07 |
| AT2G29550 | AT2G29550.1 | tubulin beta-7 chain | 51 kDa | 96 | 368 | 568 | 574 | 1606 | | 0.42 | 0.42 | 1.3E+07 | 2.1E+07 | 8.2E+06 | 4.3E+07 |
| AT5G23860 | | tubulin beta 8 | 51 kDa | 99 | 367 | 539 | 564 | 1569 | | | | | | | |
| AT5G44340 | AT5G44340.1 | tubulin beta chain 4 | 50 kDa | 73 | 296 | 513 | 485 | 1367 | 0.05 | 0.65 | 0.35 | 1.6E+07 | 2.2E+07 | 1.3E+06 | 3.9E+07 |
| AT4G20890 | | tubulin beta-9 chain | 50 kDa | 78 | 305 | 510 | 487 | 1380 | | | | | | | |
| AT4G38510 | AT4G38510.4;AT4G38510.3;AT4G38510.2;AT4G38510.1;AT4G38510.5 | ATPase, V1 complex, subunit B protein | 54 kDa | 49 | 79 | 509 | 510 | 1147 | 0.19 | 0.72 | 0.45 | 3.2E+06 | 2.6E+06 | 0.0E+00 | 5.8E+06 |
| AT3G13930 | AT3G13930.1 | Dihydrolipoamide acetyltransferase, long form protein | 58 kDa | 49 | 42 | 480 | 256 | 827 | 0.08 | 0.23 | 0.16 | 3.3E+08 | 6.9E+08 | 5.7E+07 | 1.1E+09 |
| AT4G24550 | AT4G24550.2;AT4G24550.1;AT4G24550.3 | Clathrin adaptor complexes medium subunit family pr | 51 kDa | 103 | 213 | 467 | 562 | 1345 | 3.47 | 2.88 | 3.17 | 2.9E+07 | 4.1E+07 | 6.6E+06 | 7.7E+07 |
| AT5G11490 | AT5G11490.2;AT5G11490.1;AT5G11490.3 | adaptin family protein | 93 kDa | 108 | 114 | 465 | 333 | 1020 | | 6.27 | 6.27 | 4.5E+07 | 6.4E+07 | 9.0E+06 | 1.2E+08 |
| AT5G02500 | AT5G02500.1;AT5G02500.2 | heat shock cognate protein 70-1 | 71 kDa | 66 | 57 | 464 | 364 | 951 | 0.21 | 0.33 | 0.27 | 6.7E+07 | 7.4E+07 | 7.6E+06 | 1.5E+08 |
| AT1G43170 | AT1G43170.9;AT1G43170.8;AT1G43170.7;AT1G43170.6;AT1G43170.5;AT1G43170.3;AT1G43170.2;AT1G43170.1;AT1G43170.4 | ribosomal protein 1 | 45 kDa | 31 | 84 | 463 | 305 | 883 | 0.14 | 0.33 | 0.23 | 4.3E+08 | 4.7E+08 | 1.2E+08 | 1.0E+09 |
| AT2G14720 | AT2G14720.2;AT2G14720.1 | vacuolar sorting receptor 4 | 70 kDa | 84 | 75 | 458 | 398 | 1015 | 1.47 | 4.03 | 2.75 | 6.0E+06 | 8.0E+06 | 2.2E+06 | 1.6E+07 |
| AT3G52850 | AT3G52850.1 | vacuolar sorting receptor homolog 1 | 69 kDa | 95 | 116 | 431 | 360 | 1002 | 3.68 | 2.99 | 3.33 | 4.2E+07 | 5.5E+07 | 7.4E+06 | 1.0E+08 |
| AT5G46630 | AT5G46630.1;AT5G46630.2 | Clathrin adaptor complexes medium subunit family pr | 49 kDa | 30 | 87 | 429 | 543 | 1089 | | 2.89 | 2.89 | 2.0E+07 | 2.1E+07 | 3.9E+06 | 4.4E+07 |
| AT2G14740 | AT2G14740.2;AT2G14740.1 | vaculolar sorting receptor 3 | 70 kDa | 79 | 67 | 429 | 376 | 951 | | | | 8.4E+05 | 8.2E+05 | 8.6E+05 | 2.5E+06 |
| AT5G57350 | AT5G57350.4;AT5G57350.2;AT5G57350.1;AT5G57350.3 | H(+)-ATPase 3 | 104 kDa | 37 | 14 | 426 | 280 | 757 | | | | 1.7E+06 | 4.9E+05 | 0.0E+00 | 2.2E+06 |
| AT4G12420 | AT4G12420.2;AT4G12420.1 | Cupredoxin superfamily protein | 66 kDa | 38 | 62 | 415 | 393 | 908 | 4.86 | 7.80 | 6.33 | 2.1E+08 | 1.7E+08 | 1.5E+07 | 4.0E+08 |
| AT1G30360 | AT1G30360.1 | Early-responsive to dehydration stress protein (ERD4) | 82 kDa | 18 | 0 | 407 | 295 | 720 | 4.71 | 1.28 | 2.99 | 8.0E+07 | 5.5E+07 | 1.0E+06 | 1.4E+08 |
| AT1G71410 | AT1G71410.2;AT1G71410.1 | ARM repeat superfamily protein | 99 kDa | 57 | 137 | 406 | 194 | 794 | | 14.94 | 14.94 | 1.6E+07 | 1.7E+07 | 3.8E+06 | 3.7E+07 |
| AT1G15690 | AT1G15690.1 | Inorganic H pyrophosphatase family protein | 81 kDa | 43 | 16 | 404 | 302 | 765 | 0.23 | 0.18 | 0.20 | 1.2E+08 | 7.1E+07 | 1.1E+07 | 2.0E+08 |
| AT1G04820 | | tubulin alpha-4 chain | 50 kDa | 98 | 186 | 403 | 395 | 1082 | 0.18 | 0.31 | 0.24 | | | | |
| AT2G43160 | AT2G43160.5;AT2G43160.3;AT2G43160.2;AT2G43160.1;AT2G43160.4 | ENTH/VHS family protein | 95 kDa | 93 | 270 | 401 | 386 | 1150 | 4.32 | 4.68 | 4.50 | 3.0E+07 | 4.0E+07 | 1.0E+07 | 8.1E+07 |
| AT3G09630 | AT3G09630.1;AT3G09630.2 | Ribosomal protein L4/L1 family | 45 kDa | 50 | 159 | 397 | 253 | 859 | 0.10 | 0.40 | 0.25 | 9.1E+07 | 1.2E+08 | 7.8E+07 | 2.9E+08 |
| AT4G12780 | AT4G12780.3;AT4G12780.2;AT4G12780.1;AT4G12780.6;AT4G12780.4;AT4G12780.5 | Chaperone DnaJ-domain superfamily protein | 100 kDa | 36 | 254 | 394 | 310 | 994 | 3.98 | 3.75 | 3.86 | 6.7E+05 | 1.9E+06 | 1.4E+06 | 4.0E+06 |
| AT5G12250 | AT5G12250.1 | beta-6 tubulin | 51 kDa | 72 | 210 | 390 | 390 | 1062 | 0.28 | 0.37 | 0.32 | 1.3E+06 | 1.6E+06 | 0.0E+00 | 2.9E+06 |
| AT1G75780 | | tubulin beta-1 chain | 50 kDa | 63 | 214 | 388 | 385 | 1050 | | | | | | | |
| AT4G04720 | AT4G04720.2;AT4G04720.1 | calcium-dependent protein kinase 21 | 60 kDa | 50 | 34 | 387 | 328 | 799 | 3.90 | 6.80 | 5.35 | 3.6E+07 | 2.1E+07 | 1.8E+06 | 5.9E+07 |
| AT5G02870 | AT5G02870.1;AT5G02870.2 | Ribosomal protein L4/L1 family | 45 kDa | 48 | 135 | 385 | 246 | 814 | 0.11 | 0.67 | 0.39 | 1.1E+08 | 1.1E+08 | 4.7E+07 | 2.7E+08 |
| AT2G01600 | AT2G01600.1 | ENTH/ANTH/VHS superfamily protein | 64 kDa | 51 | 80 | 383 | 361 | 875 | 2.66 | 6.79 | 4.72 | 1.9E+07 | 2.3E+07 | 2.3E+06 | 4.5E+07 |
| AT4G20830 | AT4G20830.2;AT4G20830.1 | FAD-binding Berberine family protein | 64 kDa | 2 | 17 | 382 | 337 | 738 | 2.43 | 6.00 | 4.22 | 2.1E+07 | 1.7E+07 | 3.7E+06 | 4.1E+07 |
| AT2G28520 | AT2G28520.1 | vacuolar proton ATPase A1 | 93 kDa | 17 | 26 | 378 | 199 | 620 | 3.92 | 1.91 | 2.91 | 7.0E+06 | 8.3E+06 | 4.0E+06 | 1.9E+07 |
| AT5G02490 | AT5G02490.1 | Heat shock protein 70 (Hsp 70) family protein | 71 kDa | 53 | 46 | 356 | 299 | 754 | | 0.37 | 0.37 | 4.1E+07 | 4.4E+07 | 4.1E+06 | 8.9E+07 |
| AT2G17380 | AT2G17380.1;AT4G35410.1 | associated protein 19 | 19 kDa | 37 | 89 | 353 | 144 | 623 | | 7.01 | 7.01 | 5.0E+07 | 7.9E+07 | 1.0E+07 | 1.4E+08 |
| AT1G10730 | AT1G10730.1 | Clathrin adaptor complexes medium subunit family pr | 49 kDa | 26 | 49 | 348 | 406 | 829 | | 8.02 | 8.02 | 3.8E+05 | 5.4E+05 | 6.2E+04 | 9.8E+05 |
| AT1G54920 | AT1G54920.2;AT1G54920.3;AT1G54920.4;AT1G54920.1 | unknown protein; FUNCTIONS IN: molecular_function | 98 kDa | 35 | 40 | 333 | 313 | 721 | 4.83 | 5.40 | 5.12 | 6.1E+06 | 6.7E+06 | 2.1E+06 | 1.5E+07 |

| | | | | | | | | | | | | | | | |
|---|---|---|---|---|---|---|---|---|---|---|---|---|---|---|---|
| AT3G49490 | AT3G49490.1 | unknown protein; Has 722 Blast hits to 186 proteins in | 105 kDa | 24 | 78 | 323 | 212 | 637 | | 5.08 | 5.08 | 2.5E+06 | 4.0E+06 | 5.6E+06 | 1.2E+07 |
| AT1G54920 | AT1G54920.2;AT1G54920.3;AT1G54920.4;AT1G54920.1 | unknown protein; FUNCTIONS IN: molecular_function | 99 kDa | 0 | 39 | 320 | 0 | 359 | 4.83 | 5.40 | 5.12 | 6.1E+06 | 6.7E+06 | 2.1E+06 | 1.5E+07 |
| AT3G25520 | AT3G25520.1;AT3G25520.3;AT3G25520.2 | ribosomal protein L5 | 34 kDa | 49 | 114 | 318 | 287 | 768 | 0.23 | 1.19 | 0.71 | 4.3E+07 | 4.1E+07 | 1.3E+07 | 9.7E+07 |
| AT4G12770 | AT4G12770.2;AT4G12770.1 | Chaperone DnaJ-domain superfamily protein | 98 kDa | 29 | 202 | 315 | 264 | 810 | | | | 1.2E+06 | 1.3E+06 | 1.3E+06 | 3.9E+06 |
| AT1G77140 | AT1G77140.1 | vacuolar protein sorting 45 | 65 kDa | 14 | 10 | 315 | 213 | 552 | | 5.25 | 5.25 | 5.9E+06 | 3.2E+06 | 2.1E+06 | 1.1E+07 |
| AT5G39740 | AT5G39740.2;AT5G39740.1 | ribosomal protein L5 B | 34 kDa | 47 | 104 | 310 | 281 | 742 | | 0.09 | 0.09 | 2.0E+07 | 1.8E+07 | 8.3E+06 | 4.7E+07 |
| AT4G12770 | AT4G12770.2;AT4G12770.1 | Chaperone DnaJ-domain superfamily protein | 98 kDa | 0 | 0 | 310 | 255 | 565 | | | | 1.2E+06 | 1.3E+06 | 1.3E+06 | 3.9E+06 |
| AT4G34660 | AT4G34660.1;AT4G34660.3;AT4G34660.2 | SH3 domain-containing protein | 41 kDa | 49 | 306 | 304 | 268 | 927 | 7.13 | 9.40 | 8.27 | 1.9E+07 | 2.9E+07 | 8.5E+06 | 5.6E+07 |
| AT5G51480 | AT5G51480.1 | SKU5 similar 2 | 66 kDa | 22 | 1 | 301 | 327 | 651 | 5.21 | 7.21 | 6.21 | 2.1E+07 | 2.1E+07 | 1.9E+06 | 4.4E+07 |
| AT1G54270 | AT1G54270.1;AT1G54270.2 | eif4a-2 | 47 kDa | 32 | 67 | 301 | 253 | 653 | | 0.31 | 0.31 | 5.0E+07 | 3.7E+07 | 3.9E+06 | 9.0E+07 |
| AT3G02880 | AT3G02880.1 | Leucine-rich repeat protein kinase family protein | 68 kDa | 61 | 24 | 301 | 221 | 607 | 2.49 | 5.94 | 4.22 | 2.5E+07 | 1.9E+07 | 6.2E+05 | 4.5E+07 |
| AT3G13920 | AT3G13920.1;AT3G13920.4;AT3G13920.3;AT3G13920.2;AT3G13920.5 | eukaryotic translation initiation factor 4A1 | 47 kDa | 34 | 66 | 298 | 247 | 645 | 0.09 | 0.20 | 0.14 | 2.2E+07 | 2.1E+07 | 0.0E+00 | 4.4E+07 |
| AT5G19770 | | tubulin alpha-3 | 50 kDa | 71 | 140 | 297 | 312 | 820 | 0.09 | 0.41 | 0.25 | | | | |
| AT2G20580 | AT2G20580.1 | 26S proteasome regulatory subunit S2 1A | ? | 43 | 17 | 297 | 190 | 547 | 0.19 | 0.27 | 0.23 | 2.0E+08 | 2.2E+08 | 2.9E+07 | 4.5E+08 |
| AT3G12580 | AT3G12580.1 | heat shock protein 70 | 71 kDa | 49 | 38 | 291 | 222 | 600 | | 0.46 | 0.46 | 1.6E+06 | 1.7E+06 | 4.4E+05 | 3.8E+06 |
| AT4G35410 | AT4G35410.2 | Clathrin adaptor complex small chain family protein | 19 kDa | 26 | 46 | 291 | 117 | 480 | 10.33 | | 10.33 | 2.8E+06 | 6.9E+06 | 1.3E+06 | 1.1E+07 |
| AT1G07770 | | ribosomal protein S15A | 15 kDa | 9 | 6 | 285 | 29 | 329 | 0.31 | 0.17 | 0.24 | | | | |
| AT4G16370 | AT4G16370.1 | oligopeptide transporter | 82 kDa | 7 | 1 | 284 | 326 | 618 | 6.70 | 5.61 | 6.16 | 1.4E+07 | 1.1E+07 | 5.5E+06 | 3.1E+07 |
| AT5G09810 | AT5G09810.1 | actin 7 | 42 kDa | 63 | 82 | 282 | 242 | 669 | 0.14 | 0.77 | 0.45 | 2.8E+07 | 2.8E+07 | 5.6E+06 | 6.2E+07 |
| AT3G09440 | AT3G09440.4;AT3G09440.3;AT3G09440.2;AT3G09440.1 | Heat shock protein 70 (Hsp 70) family protein | 71 kDa | 60 | 41 | 282 | 236 | 619 | | 0.27 | 0.27 | 1.8E+07 | 2.0E+07 | 1.2E+06 | 3.9E+07 |
| AT4G20720 | AT4G20720.1;AT4G20720.2 | dentin sialophosphoprotein-related | 80 kDa | 52 | 66 | 271 | 347 | 736 | 3.60 | 5.35 | 4.48 | 1.6E+07 | 2.0E+07 | 2.1E+06 | 3.8E+07 |
| AT4G08850 | AT4G08850.1;AT4G08850.2 | Leucine-rich repeat receptor-like protein kinase family | 115 kDa | 16 | 11 | 267 | 197 | 491 | 2.67 | 4.82 | 3.74 | 4.8E+07 | 4.2E+07 | 3.0E+06 | 9.3E+07 |
| AT4G16155 | AT4G16155.1 | dihydrolipoyl dehydrogenases | 67 kDa | 64 | 110 | 265 | 388 | 827 | 0.05 | 0.33 | 0.19 | 4.3E+07 | 2.5E+07 | 1.6E+06 | 6.9E+07 |
| AT4G32760 | AT4G32760.2;AT4G32760.1;AT4G32760.3 | ENTH/VHS/GAT family protein | 73 kDa | 33 | 80 | 263 | 147 | 523 | 8.88 | 3.86 | 6.37 | 1.4E+07 | 1.4E+07 | 8.7E+06 | 3.7E+07 |
| AT5G47480 | AT5G47480.1;AT5G47480.2 | RGPR-related | 146 kDa | 7 | 6 | 260 | 188 | 461 | | 5.18 | 5.18 | 3.5E+05 | 1.6E+06 | 7.9E+05 | 2.7E+06 |
| AT3G02260 | AT3G02260.2;AT3G02260.3;AT3G02260.4;AT3G02260.1 | auxin transport protein (BIG) | 568 kDa | 1 | 0 | 258 | 281 | 540 | | 0.17 | 0.17 | 1.7E+07 | 1.3E+07 | 2.9E+06 | 3.3E+07 |
| AT4G25240 | AT4G25240.2;AT4G25240.1 | SKU5 similar 1 | 66 kDa | 8 | 3 | 257 | 297 | 565 | 3.96 | 7.81 | 5.88 | 1.8E+07 | 1.3E+07 | 2.1E+06 | 3.3E+07 |
| AT5G62390 | AT5G62390.1 | BCL-2-associated athanogene 7 | 52 kDa | 26 | 51 | 256 | 210 | 543 | | 0.34 | 0.34 | 6.9E+06 | 1.1E+07 | 1.7E+06 | 2.0E+07 |
| AT3G28715 | AT3G28715.1;AT3G28715.2 | ATPase, V0/A0 complex, subunit C/D | 41 kDa | 28 | 71 | 255 | 193 | 547 | | 3.57 | 3.57 | 1.0E+07 | 3.3E+06 | 1.2E+05 | 1.4E+07 |
| AT3G28710 | AT3G28710.1 | ATPase, V0/A0 complex, subunit C/D | 41 kDa | 28 | 72 | 255 | 191 | 546 | 0.52 | 3.82 | 2.17 | 3.4E+06 | 3.3E+06 | 4.3E+05 | 7.1E+06 |
| AT3G16950 | AT3G16950.1;AT3G16950.2 | lipoamide dehydrogenase 1 | 67 kDa | 53 | 79 | 248 | 402 | 782 | | 0.48 | 0.48 | 3.0E+07 | 1.7E+07 | 1.2E+07 | 5.9E+07 |
| AT2G31610 | AT2G31610.1 | Ribosomal protein S3 family protein | 28 kDa | 32 | 94 | 248 | 127 | 501 | 0.11 | 0.74 | 0.42 | 8.8E+07 | 8.2E+07 | 2.3E+07 | 1.9E+08 |
| AT1G45000 | AT1G45000.1;AT1G45000.2 | AAA-type ATPase family protein | 45 kDa | 32 | 68 | 245 | 182 | 527 | 0.21 | 1.02 | 0.61 | 3.8E+07 | 4.5E+07 | 4.1E+06 | 8.8E+07 |
| AT4G34670 | AT4G34670.1 | Ribosomal protein S3Ae | 30 kDa | 27 | 71 | 245 | 110 | 453 | 0.12 | 0.29 | 0.20 | 1.1E+08 | 1.2E+08 | 2.0E+07 | 2.5E+08 |
| AT1G54220 | AT1G54220.2;AT1G54220.1 | Dihydrolipoamide acetyltransferase, long form protein | 58 kDa | 19 | 9 | 243 | 160 | 431 | 0.06 | 1.00 | 0.53 | 1.1E+08 | 2.4E+08 | 1.2E+07 | 3.7E+08 |
| AT5G15200 | AT5G15200.1;AT5G15200.2 | Ribosomal protein S4 | 23 kDa | 17 | 71 | 240 | 93 | 421 | 0.17 | 0.50 | 0.33 | 3.3E+08 | 3.7E+08 | 5.3E+07 | 7.5E+08 |
| AT1G34430 | AT1G34430.1 | 2-oxoacid dehydrogenases acyltransferase family prote | 48 kDa | 74 | 57 | 239 | 357 | 727 | 0.23 | 0.28 | 0.25 | 9.3E+07 | 5.4E+07 | 2.2E+07 | 1.7E+08 |
| AT5G58420 | AT5G58420.1 | Ribosomal protein S4 (RPS4A) family protein | 30 kDa | 0 | 97 | 238 | 0 | 335 | | | | 1.1E+07 | 2.5E+06 | 1.5E+06 | 1.5E+07 |
| AT2G17360 | | Ribosomal protein S4 (RPS4A) family protein | 30 kDa | 43 | 102 | 237 | 151 | 533 | | | | | | | |
| AT3G04840 | AT3G04840.1 | Ribosomal protein S3Ae | 30 kDa | 23 | 80 | 234 | 105 | 442 | 0.09 | 0.41 | 0.25 | 1.4E+08 | 1.6E+08 | 2.8E+07 | 3.2E+08 |
| AT4G33070 | AT4G33070.1 | Thiamine pyrophosphate dependent pyruvate decarbo | 66 kDa | 135 | 252 | 233 | 1042 | 1662 | 0.49 | 1.65 | 1.07 | 3.0E+08 | 8.2E+08 | 3.7E+07 | 1.2E+09 |
| AT3G05530 | AT3G05530.1 | regulatory particle triple-A ATPase 5A | 47 kDa | 41 | 70 | 233 | 195 | 539 | 0.16 | 0.65 | 0.41 | 3.9E+08 | 4.5E+08 | 4.0E+07 | 8.7E+08 |
| AT3G53870 | AT3G53870.1 | Ribosomal protein S3 family protein | 27 kDa | 0 | 82 | 232 | 123 | 437 | | 1.81 | 1.81 | 5.7E+07 | 4.5E+07 | 7.4E+06 | 1.1E+08 |
| AT5G35530 | AT5G35530.1 | Ribosomal protein S3 family protein | 27 kDa | 29 | 70 | 227 | 114 | 440 | | | | 4.6E+07 | 3.0E+07 | 3.5E+06 | 7.9E+07 |
| AT1G33120 | | Ribosomal protein L6 family | 22 kDa | 42 | 41 | 222 | 153 | 458 | 0.18 | 0.24 | 0.21 | | | | |
| AT2G38410 | AT2G38410.1 | ENTH/VHS/GAT family protein | 72 kDa | 35 | 24 | 219 | 279 | 557 | 3.52 | 4.03 | 3.78 | 3.7E+06 | 4.7E+06 | 7.4E+05 | 9.1E+06 |
| AT2G34480 | AT2G34480.1;AT2G34480.2 | Ribosomal protein L18ae/LX family protein | 21 kDa | 22 | 24 | 217 | 121 | 384 | 0.15 | 0.21 | 0.18 | 3.0E+08 | 3.3E+08 | 9.2E+07 | 7.2E+08 |
| AT5G57110 | AT5G57110.3;AT5G57110.2;AT5G57110.1 | autoinhibited Ca2+-ATPase, isoform 8 | 116 kDa | 7 | 0 | 215 | 192 | 414 | | 0.47 | 0.47 | 6.1E+07 | 5.9E+07 | 3.0E+06 | 1.2E+08 |
| AT4G12730 | AT4G12730.1 | FASCICLIN-like arabinogalactan 2 | 43 kDa | 7 | 12 | 215 | 176 | 410 | | 2.84 | 2.84 | 5.4E+07 | 5.6E+07 | 2.2E+06 | 1.1E+08 |
| AT5G43010 | AT5G43010.1 | regulatory particle triple-A ATPase 4A | 45 kDa | 30 | 56 | 215 | 162 | 463 | | | | 1.1E+07 | 1.7E+07 | 2.9E+06 | 3.0E+07 |
| AT3G56150 | AT3G56150.1;AT3G56150.2 | eukaryotic translation initiation factor 3C | 103 kDa | 18 | 18 | 215 | 122 | 373 | 0.14 | 0.35 | 0.25 | 3.1E+07 | 3.7E+07 | 1.5E+07 | 8.3E+07 |
| AT4G12650 | AT4G12650.1 | Endomembrane protein 70 protein family | 74 kDa | 17 | 29 | 214 | 204 | 464 | | 0.30 | 0.30 | 1.1E+07 | 7.8E+06 | 4.3E+06 | 2.3E+07 |
| AT5G58290 | AT5G58290.1 | regulatory particle triple-A ATPase 3 | 46 kDa | 27 | 48 | 210 | 112 | 397 | 0.14 | 0.81 | 0.47 | 5.2E+08 | 6.5E+08 | 5.2E+07 | 1.2E+09 |
| AT1G24180 | AT1G24180.1 | Thiamin diphosphate-binding fold (THDP-binding) supe | 43 kDa | 41 | 43 | 208 | 131 | 423 | 0.12 | 0.71 | 0.41 | 2.2E+08 | 4.8E+08 | 3.7E+07 | 7.4E+08 |
| AT4G11420 | AT4G11420.1 | eukaryotic translation initiation factor 3A | ? | 9 | 10 | 208 | 119 | 346 | | 0.21 | 0.21 | 4.2E+07 | 4.1E+07 | 1.6E+07 | 9.8E+07 |
| AT3G62870 | AT3G62870.1 | Ribosomal protein L7Ae/L30e/S12e/Gadd45 family pro | 29 kDa | 34 | 101 | 208 | 103 | 446 | 0.09 | 0.66 | 0.37 | 1.4E+08 | 1.5E+08 | 2.3E+07 | 3.2E+08 |
| AT5G19990 | AT5G19990.1;AT5G19990.3;AT5G19990.2 | regulatory particle triple-A ATPase 6A | 47 kDa | 34 | 30 | 207 | 158 | 429 | | 1.30 | 1.30 | 6.7E+07 | 1.0E+08 | 1.6E+06 | 1.7E+08 |
| AT3G09200 | AT3G09200.1;AT3G09200.2 | Ribosomal protein L10 family protein | 34 kDa | 17 | 61 | 206 | 172 | 456 | 0.07 | 0.24 | 0.15 | 1.6E+08 | 1.9E+08 | 7.5E+07 | 4.2E+08 |
| AT3G59290 | AT3G59290.1 | ENTH/VHS family protein | 110 kDa | 31 | 90 | 206 | 163 | 490 | | | | 6.0E+05 | 9.8E+05 | 3.1E+05 | 1.9E+06 |
| AT5G20000 | AT5G20000.1 | AAA-type ATPase family protein | 47 kDa | 34 | 30 | 206 | 157 | 427 | | | | 3.7E+07 | 4.6E+07 | 1.5E+06 | 8.4E+07 |
| AT2G01690 | AT2G01690.1;AT2G01690.2 | ARM repeat superfamily protein | 84 kDa | 18 | 4 | 203 | 179 | 404 | 0.28 | 0.55 | 0.42 | 3.0E+07 | 1.7E+07 | 2.2E+07 | 7.0E+07 |
| AT2G36530 | AT2G36530.1;AT2G36530.2 | Enolase | 48 kDa | 51 | 119 | 197 | 258 | 625 | 0.49 | 2.54 | 1.52 | 3.2E+07 | 2.6E+07 | 3.3E+06 | 6.1E+07 |
| AT5G20230 | | blue-copper-binding protein | 20 kDa | 2 | 92 | 197 | 173 | 464 | 4.68 | 12.00 | 8.34 | | | | |
| AT3G24550 | AT3G24550.1 | proline extensin-like receptor kinase 1 | 69 kDa | 10 | 31 | 197 | 123 | 361 | | 5.50 | 5.50 | 4.8E+07 | 5.6E+07 | 2.0E+06 | 1.1E+08 |
| AT3G49010 | AT3G49010.7;AT3G49010.6;AT3G49010.3;AT3G49010.2;AT3G49010.1;AT3G49010.4;AT3G49010.5 | breast basic conserved 1 | 24 kDa | 19 | 64 | 192 | 111 | 386 | 0.11 | 0.35 | 0.23 | 5.0E+08 | 5.7E+08 | 1.6E+08 | 1.2E+09 |
| AT1G06220 | AT1G06220.3;AT1G06220.2;AT1G06220.1 | Ribosomal protein S5/Elongation factor G/III/V family p | 110 kDa | 4 | 5 | 192 | 106 | 307 | 0.17 | 0.21 | 0.19 | 1.4E+07 | 1.9E+07 | 1.5E+06 | 3.4E+07 |
| AT3G47950 | | H(+)-ATPase 4 | 106 kDa | 0 | 14 | 189 | 131 | 334 | | | | | | | |
| AT2G01250 | AT2G01250.1;AT2G01250.2 | Ribosomal protein L30/L7 family protein | 28 kDa | 21 | 63 | 188 | 122 | 394 | 0.06 | 0.58 | 0.32 | 1.0E+08 | 8.4E+07 | 2.7E+08 | 4.6E+08 |
| AT1G30450 | AT1G30450.3;AT1G30450.2;AT1G30450.1;AT1G30450.4 | cation-chloride co-transporter 1 | 107 kDa | 4 | 0 | 188 | 65 | 257 | | | | 2.3E+06 | 3.0E+06 | 1.0E+06 | 6.3E+06 |
| AT1G58380 | | Ribosomal protein S5 family protein | 31 kDa | 23 | 60 | 187 | 101 | 371 | 0.10 | 0.44 | 0.27 | | | | |
| AT4G27090 | AT4G27090.1 | Ribosomal protein L14 | 16 kDa | 21 | 3 | 187 | 85 | 296 | 0.19 | 0.13 | 0.16 | 5.8E+08 | 6.0E+08 | 1.1E+08 | 1.3E+09 |
| AT5G24010 | AT5G24010.1 | Protein kinase superfamily protein | 92 kDa | 13 | 13 | 186 | 164 | 376 | 7.60 | 4.49 | 6.05 | 2.1E+07 | 1.3E+07 | 2.2E+05 | 3.3E+07 |
| AT1G74050 | AT1G74050.1;AT1G74060.1 | Ribosomal protein L6 family protein | 26 kDa | 29 | 83 | 186 | 147 | 445 | 0.12 | 0.51 | 0.32 | 3.3E+08 | 4.2E+08 | 1.3E+08 | 8.9E+08 |
| AT5G50850 | AT5G50850.1 | Transketolase family protein | 39 kDa | 39 | 28 | 186 | 113 | 366 | 0.14 | 0.14 | 0.14 | 5.8E+08 | 1.1E+09 | 8.7E+07 | 1.7E+09 |
| AT1G53210 | AT1G53210.1 | sodium/calcium exchanger family protein / calcium-bi | 63 kDa | 6 | 0 | 184 | 149 | 339 | 0.20 | 0.31 | 0.26 | 4.6E+07 | 3.0E+07 | 4.8E+05 | 7.7E+07 |
| AT1G53840 | AT1G53840.1 | pectin methylesterase 1 | 64 kDa | 25 | 23 | 184 | 136 | 368 | | 6.94 | 6.94 | 1.3E+07 | 1.1E+07 | 6.8E+04 | 2.4E+07 |
| AT2G41840 | AT2G41840.1 | Ribosomal protein S5 family protein | 31 kDa | 26 | 69 | 184 | 103 | 382 | 0.12 | 0.28 | 0.20 | 1.4E+08 | 1.5E+08 | 2.3E+07 | 3.2E+08 |
| AT1G53750 | AT1G53750.1 | regulatory particle triple-A 1A | 48 kDa | 25 | 31 | 183 | 115 | 354 | | 0.49 | 0.49 | 2.6E+08 | 3.6E+08 | 2.8E+07 | 6.6E+08 |
| AT4G39080 | AT4G39080.1 | vacuolar proton ATPase A3 | 93 kDa | 10 | 5 | 181 | 143 | 339 | 0.39 | 0.21 | 0.30 | 1.8E+07 | 1.0E+07 | 8.9E+06 | 3.7E+07 |
| AT5G51570 | AT5G51570.1 | SPFH/Band 7/PHB domain-containing membrane-asso | 32 kDa | 33 | 54 | 180 | 135 | 402 | 0.47 | 4.42 | 2.45 | 5.9E+06 | 1.9E+06 | 6.2E+06 | 1.4E+07 |
| AT3G52990 | AT3G52990.1;AT3G52990.2 | Pyruvate kinase family protein | 57 kDa | 15 | 38 | 180 | 114 | 347 | | 0.23 | 0.23 | 2.7E+07 | 2.0E+07 | 1.8E+06 | 4.9E+07 |

| | | | | | | | | | | | | | | | |
|---|---|---|---|---|---|---|---|---|---|---|---|---|---|---|---|
| AT5G07120 | AT5G07120.1;AT5G07120.2;AT5G07120.3 | sorting nexin 2B | 64 kDa | 35 | 62 | 179 | 186 | 462 | | 4.48 | 4.48 | 1.6E+07 | 2.0E+07 | 3.2E+06 | 3.9E+07 |
| AT5G06140 | AT5G06140.1 | sorting nexin 1 | 47 kDa | 52 | 73 | 177 | 162 | 464 | | 3.79 | 3.79 | 2.4E+07 | 2.6E+07 | 2.5E+06 | 5.3E+07 |
| AT2G44120 | AT2G44120.1;AT2G44120.2 | Ribosomal protein L30/L7 family protein | 28 kDa | 23 | 75 | 177 | 120 | 395 | 0.08 | 0.27 | 0.17 | 1.3E+08 | 1.3E+08 | 1.7E+08 | 4.2E+08 |
| AT2G47610 | AT2G47610.1 | Ribosomal protein L7Ae/L30e/S12e/Gadd45 family prc | 29 kDa | 31 | 82 | 174 | 97 | 384 | | 0.65 | 0.65 | 8.8E+07 | 9.8E+07 | 2.3E+07 | 2.1E+08 |
| AT1G12840 | AT1G12840.1 | vacuolar ATP synthase subunit C (VATC) / V-ATPase C sul | 43 kDa | 18 | 36 | 173 | 131 | 358 | 0.22 | 2.00 | 1.11 | 5.2E+07 | 2.9E+07 | 4.3E+06 | 8.6E+07 |
| AT3G52200 | AT3G52200.1;AT3G52200.2 | Dihydrolipoamide acetyltransferase, long form protein | 69 kDa | 9 | 3 | 173 | 111 | 296 | | 0.28 | 0.28 | 9.8E+07 | 1.9E+08 | 2.1E+07 | 3.1E+08 |
| AT1G29150 | AT1G29150.2;AT1G29150.1 | non-ATPase subunit 9 | 47 kDa | 23 | 12 | 170 | 124 | 329 | 0.19 | 1.12 | 0.65 | 3.1E+08 | 3.5E+08 | 4.0E+07 | 7.1E+08 |
| AT2G05920 | AT2G05920.1 | Subtilase family protein | 80 kDa | 4 | 13 | 168 | 118 | 303 | 0.18 | 0.25 | 0.22 | 1.7E+07 | 1.9E+07 | 1.0E+06 | 3.8E+07 |
| AT3G25860 | AT3G25860.1 | 2-oxoacid dehydrogenases acyltransferase family prote | 50 kDa | 58 | 89 | 167 | 217 | 531 | 0.09 | 0.30 | 0.20 | 1.1E+08 | 5.5E+07 | 2.2E+07 | 1.9E+08 |
| AT5G17920 | AT5G17920.2;AT5G17920.1 | Cobalamin-independent synthase family protein | 84 kDa | 40 | 36 | 167 | 167 | 410 | 0.45 | 0.90 | 0.67 | 4.8E+07 | 4.5E+07 | 1.5E+06 | 9.5E+07 |
| AT5G55480 | AT5G55480.1 | SHV3-like 1 | 84 kDa | 10 | 30 | 167 | 150 | 357 | 2.82 | 5.80 | 4.31 | 7.1E+07 | 6.2E+07 | 8.0E+06 | 1.4E+08 |
| AT1G64790 | AT1G64790.1;AT1G64790.3;AT1G64790.2 | ILITYHIA | 285 kDa | 0 | 1 | 167 | 114 | 282 | | 0.15 | 0.15 | 1.1E+07 | 9.9E+06 | 4.2E+06 | 2.5E+07 |
| AT4G31140 | AT4G31140.1 | O-Glycosyl hydrolases family 17 protein | 53 kDa | 29 | 30 | 166 | 183 | 408 | | 11.82 | 11.82 | 3.1E+07 | 2.3E+07 | 1.7E+06 | 5.5E+07 |
| AT2G05220 | AT2G05220.2;AT2G05220.1 | Ribosomal S17 family protein | 16 kDa | 38 | 5 | 166 | 109 | 318 | | | | 1.0E+07 | 4.5E+07 | 3.7E+06 | 5.8E+07 |
| AT3G19930 | AT3G19930.2;AT3G19930.1 | sugar transporter 4 | 57 kDa | 7 | 42 | 165 | 126 | 340 | 4.75 | 2.89 | 3.82 | 2.2E+07 | 1.8E+07 | 3.6E+05 | 4.0E+07 |
| AT4G29040 | AT4G29040.1 | regulatory particle AAA-ATPase 2A | 49 kDa | 23 | 31 | 164 | 155 | 373 | | | | 5.9E+06 | 7.7E+06 | 2.0E+05 | 1.4E+07 |
| AT5G20290 | AT5G20290.1 | Ribosomal protein S8e family protein | 25 kDa | 36 | 105 | 163 | 82 | 386 | 0.17 | 0.48 | 0.33 | 1.3E+09 | 1.6E+09 | 2.6E+08 | 3.2E+09 |
| AT2G27710 | AT2G27710.3;AT2G27710.2;AT2G27710.1;AT2G27710.4 | 60S acidic ribosomal protein family | 11 kDa | 15 | 0 | 163 | 40 | 218 | 0.28 | 0.22 | 0.25 | 2.8E+08 | 3.2E+08 | 1.8E+08 | 7.9E+08 |
| AT1G67140 | AT1G67140.1;AT1G67140.3;AT1G67140.2 | HEAT repeat-containing protein | 244 kDa | 24 | 4 | 162 | 359 | 549 | | | | 1.6E+06 | 1.1E+06 | 0.0E+00 | 2.7E+06 |
| AT3G12110 | | actin-11 | 42 kDa | 48 | 45 | 162 | 136 | 391 | | | | | | | |
| AT4G28470 | AT4G28470.1 | 26S proteasome regulatory subunit S2 1B | 98 kDa | 0 | 13 | 162 | 99 | 274 | | 0.07 | 0.07 | 3.2E+07 | 3.8E+07 | 5.0E+06 | 7.5E+07 |
| AT1G56070 | AT1G56070.3;AT1G56070.2;AT1G56070.1 | Ribosomal protein S5/Elongation factor G/III/V family | 94 kDa | 29 | 9 | 161 | 107 | 306 | 0.11 | 0.17 | 0.14 | 7.1E+07 | 6.5E+07 | 2.2E+07 | 1.6E+08 |
| AT3G42050 | AT3G42050.1 | vacuolar ATP synthase subunit H family protein | 50 kDa | 25 | 25 | 159 | 136 | 345 | 0.12 | 0.50 | 0.31 | 4.6E+07 | 2.0E+07 | 5.6E+06 | 7.2E+07 |
| AT1G80070 | AT1G80070.1;AT4G38780.1 | Pre-mRNA-processing-splicing factor | 275 kDa | 3 | 1 | 158 | 161 | 323 | 0.21 | 0.12 | 0.17 | 2.6E+07 | 3.1E+07 | 6.2E+06 | 6.3E+07 |
| AT2G36580 | AT2G36580.1 | Pyruvate kinase family protein | 58 kDa | 19 | 35 | 158 | 99 | 311 | | | | 1.3E+07 | 1.1E+07 | 9.1E+05 | 2.4E+07 |
| AT3G52140 | AT3G52140.1;AT3G52140.2;AT3G52140.4;AT3G52140.3 | tetratricopeptide repeat (TPR)-containing protein | 153 kDa | 1 | 9 | 155 | 94 | 259 | | 0.19 | 0.19 | 1.7E+08 | 1.6E+08 | 4.8E+07 | 3.7E+08 |
| AT2G37620 | | actin 1 | 42 kDa | 45 | 45 | 154 | 130 | 374 | 0.32 | 1.27 | 0.80 | | | | |
| AT1G20200 | AT1G20200.1 | PAM domain (PCI/PINT associated module) protein | 56 kDa | 26 | 21 | 154 | 112 | 313 | 0.18 | 0.53 | 0.36 | 1.0E+08 | 9.8E+07 | 1.9E+07 | 2.2E+08 |
| AT1G28490 | AT1G28490.1;AT1G28490.3;AT1G28490.2 | syntaxin of plants 61 | 28 kDa | 17 | 95 | 154 | 101 | 367 | 3.65 | 7.54 | 5.60 | 5.3E+06 | 8.8E+06 | 2.5E+06 | 1.7E+07 |
| AT1G50920 | AT1G50920.1 | Nucleolar GTP-binding protein | 77 kDa | 1 | 5 | 154 | 89 | 249 | | 0.36 | 0.36 | 5.3E+07 | 3.6E+07 | 1.5E+07 | 1.0E+08 |
| AT1G18540 | AT1G18540.1 | Ribosomal protein L6 family protein | 26 kDa | 21 | 59 | 153 | 110 | 343 | 0.12 | 0.54 | 0.33 | 9.1E+07 | 1.3E+08 | 4.6E+07 | 2.7E+08 |
| AT1G59870 | AT1G59870.1 | ABC-2 and Plant PDR ABC-type transporter family prote | 165 kDa | 11 | 0 | 152 | 131 | 294 | | 0.34 | 0.34 | 1.1E+07 | 8.4E+06 | 1.0E+05 | 2.0E+07 |
| AT1G75680 | AT1G75680.1 | glycosyl hydrolase 9B7 | 58 kDa | 1 | 5 | 152 | 89 | 247 | | 5.78 | 5.78 | 1.9E+07 | 1.1E+07 | 2.9E+06 | 3.3E+07 |
| AT5G25230 | AT5G25230.2;AT5G25230.1 | Ribosomal protein S5/Elongation factor G/III/V family | 109 kDa | 0 | 5 | 152 | 88 | 245 | | | | 4.9E+05 | 1.1E+06 | 9.9E+04 | 1.7E+06 |
| AT3G62290 | AT3G62290.3;AT3G62290.2;AT3G62290.1 | ADP-ribosylation factor A1E | 21 kDa | 51 | 22 | 151 | 137 | 361 | | | | 7.7E+06 | 6.6E+06 | 0.0E+00 | 1.4E+07 |
| AT1G04270 | AT1G04270.2;AT1G04270.1 | cytosolic ribosomal protein S15 | 17 kDa | 8 | 1 | 150 | 53 | 212 | 0.17 | 0.58 | 0.37 | 2.0E+07 | 1.1E+07 | 3.5E+07 | 6.5E+07 |
| AT3G53020 | AT3G53020.1 | Ribosomal protein L24e family protein | 19 kDa | 10 | 9 | 149 | 65 | 233 | | | | 3.7E+07 | 9.9E+07 | 6.3E+06 | 1.4E+08 |
| AT1G14320 | AT1G14320.1;AT1G14320.2 | Ribosomal protein L16p/L10e family protein | 25 kDa | 10 | 34 | 148 | 55 | 247 | 0.05 | 0.17 | 0.11 | 6.2E+07 | 7.5E+07 | 1.9E+07 | 1.6E+08 |
| AT4G22750 | AT4G22750.1 | DHHC-type zinc finger family protein | 34 kDa | 4 | 7 | 147 | 129 | 287 | | | | 2.0E+07 | 2.4E+07 | 1.5E+07 | 5.9E+07 |
| AT3G13580 | AT3G13580.9;AT3G13580.8;AT3G13580.7;AT3G13580.6;AT3G13580.5;AT3G13580.4;AT3G13580.3;AT3G13580.2;AT3G13580.1 | Ribosomal protein L30/L7 family protein | 28 kDa | 19 | 56 | 146 | 97 | 318 | | 0.23 | 0.23 | 2.0E+07 | 2.0E+07 | 3.7E+06 | 4.4E+07 |
| AT1G66580 | AT1G66580.1 | senescence associated gene 24 | 25 kDa | 9 | 31 | 146 | 53 | 239 | 0.12 | 0.20 | 0.16 | 7.2E+06 | 3.9E+06 | 9.2E+06 | 2.0E+07 |
| AT2G20450 | AT2G20450.1 | Ribosomal protein L14 | 16 kDa | 15 | 0 | 145 | 71 | 231 | 0.48 | 0.15 | 0.32 | 9.9E+07 | 1.6E+08 | 2.9E+07 | 2.8E+08 |
| AT4G23650 | AT4G23650.1 | calcium-dependent protein kinase 6 | 59 kDa | 38 | 12 | 144 | 148 | 342 | | 2.64 | 2.64 | 2.2E+07 | 8.2E+06 | 1.1E+05 | 3.0E+07 |
| AT4G11150 | AT4G11150.1 | vacuolar ATP synthase subunit E1 | 26 kDa | 26 | 64 | 144 | 134 | 368 | 0.33 | 1.67 | 1.00 | 9.7E+07 | 6.0E+07 | 3.3E+06 | 1.6E+08 |
| AT3G14990 | AT3G14990.1;AT3G14990.3;AT3G14990.2 | Class I glutamine amidotransferase-like superfamily prc | 42 kDa | 10 | 31 | 144 | 129 | 314 | 0.15 | 0.50 | 0.32 | 3.3E+07 | 1.5E+07 | 9.6E+05 | 4.9E+07 |
| AT3G11940 | AT3G11940.2;AT3G11940.1 | ribosomal protein 5A | 23 kDa | 25 | 70 | 143 | 90 | 328 | | | | 7.2E+07 | 5.9E+07 | 6.7E+06 | 1.4E+08 |
| AT5G58090 | AT5G58090.1 | O-Glycosyl hydrolases family 17 protein | 52 kDa | 9 | 20 | 142 | 118 | 289 | | 4.84 | 4.84 | 3.9E+07 | 2.3E+07 | 1.4E+06 | 6.3E+07 |
| AT3G18780 | AT3G18780.3;AT3G18780.2;AT1G49240.1;AT3G18780.1 | actin 2 | 42 kDa | 29 | 44 | 142 | 111 | 326 | | | | 5.3E+06 | 5.1E+06 | 6.0E+05 | 1.1E+07 |
| AT3G58510 | AT3G58510.3;AT3G58510.2;AT3G58510.1 | DEA(D/H)-box RNA helicase family protein | 66 kDa | 10 | 25 | 142 | 84 | 261 | 0.10 | 0.29 | 0.20 | 8.8E+06 | 7.4E+06 | 3.6E+06 | 2.0E+07 |
| AT1G22780 | | Ribosomal protein S13/S18 family | 18 kDa | 20 | 1 | 142 | 59 | 222 | 0.20 | 0.22 | 0.21 | | | | |
| AT3G51550 | AT3G51550.1 | Malectin/receptor-like protein kinase family protein | 98 kDa | 9 | 10 | 141 | 97 | 257 | 3.29 | 5.63 | 4.46 | 9.1E+06 | 8.5E+06 | 8.4E+04 | 1.8E+07 |
| AT1G80410 | AT1G80410.1;AT1G80410.2 | tetratricopeptide repeat (TPR)-containing protein | 102 kDa | 6 | 10 | 141 | 90 | 247 | 0.11 | 0.45 | 0.28 | 4.2E+07 | 4.7E+07 | 9.7E+06 | 9.9E+07 |
| AT3G24830 | AT3G24830.1 | Ribosomal protein L13 family protein | 23 kDa | 12 | 28 | 141 | 83 | 264 | | 0.38 | 0.38 | 7.5E+07 | 1.0E+08 | 2.1E+07 | 2.0E+08 |
| AT5G09900 | AT5G09900.2;AT5G09900.1;AT5G09900.3 | 26S proteasome regulatory subunit, putative (RPN5) | 51 kDa | 22 | 22 | 140 | 103 | 287 | 0.13 | 0.98 | 0.55 | 2.0E+08 | 1.8E+08 | 1.8E+07 | 4.0E+08 |
| AT1G07660 | | Histone superfamily protein | 11 kDa | 16 | 0 | 140 | 61 | 217 | 0.23 | 0.36 | 0.30 | | | | |
| AT1G20960 | AT1G20960.2;AT1G20960.1 | U5 small nuclear ribonucleoprotein helicase, putative | 247 kDa | 0 | 0 | 138 | 148 | 286 | | 0.15 | 0.15 | 1.7E+07 | 2.0E+07 | 6.3E+06 | 4.4E+07 |
| AT1G64760 | AT1G64760.2;AT1G64760.1 | O-Glycosyl hydrolases family 17 protein | 52 kDa | 14 | 31 | 138 | 119 | 302 | | 8.87 | 8.87 | 9.2E+07 | 7.0E+07 | 6.6E+06 | 1.7E+08 |
| AT3G17840 | AT3G17840.1 | receptor-like kinase 902 | 70 kDa | 17 | 0 | 138 | 109 | 264 | | 3.95 | 3.95 | 4.2E+06 | 4.2E+06 | 1.5E+05 | 8.6E+06 |
| AT3G20410 | AT3G20410.2;AT3G20410.1 | calmodulin-domain protein kinase 9 | 60 kDa | 14 | 5 | 137 | 131 | 287 | | | | 1.9E+06 | 4.0E+05 | 0.0E+00 | 2.3E+06 |
| AT5G16590 | AT5G16590.1 | Leucine-rich repeat protein kinase family protein | 67 kDa | 32 | 6 | 137 | 111 | 286 | | 2.98 | 2.98 | 3.4E+06 | 3.2E+06 | 0.0E+00 | 6.6E+06 |
| AT2G18020 | AT2G18020.1 | Ribosomal protein L2 family | 28 kDa | 26 | 85 | 137 | 101 | 349 | 0.15 | 0.26 | 0.21 | 2.5E+08 | 2.8E+08 | 2.2E+08 | 7.5E+08 |
| AT1G72730 | AT1G72730.1 | DEA(D/H)-box RNA helicase family protein | 47 kDa | 24 | 53 | 136 | 105 | 318 | | | | 6.3E+05 | 0.0E+00 | 2.4E+05 | 8.7E+05 |
| AT5G47200 | AT5G47200.1;AT4G17530.1 | RAB GTPase homolog 1A | 22 kDa | 29 | 103 | 136 | 86 | 354 | | | | 4.8E+07 | 3.0E+07 | 1.8E+06 | 8.0E+07 |
| AT3G46040 | AT3G46040.1 | ribosomal protein S15A D | 15 kDa | 0 | 4 | 134 | 28 | 166 | | | | 4.7E+06 | 5.8E+06 | 5.2E+05 | 1.1E+07 |
| AT5G47490 | AT5G47490.1;AT5G47490.2;AT5G47490.3 | RGPR-related | 148 kDa | 3 | 0 | 133 | 98 | 234 | | | | 0.0E+00 | 0.0E+00 | 1.7E+05 | 1.7E+05 |
| AT2G03450 | AT2G03450.1 | purple acid phosphatase 9 | 74 kDa | 9 | 4 | 133 | 75 | 221 | | 3.88 | 3.88 | | | | |
| AT2G33120 | AT2G33120.1;AT2G33120.2 | synaptobrevin-related protein 1 | 25 kDa | 26 | 81 | 132 | 94 | 333 | 2.06 | 7.20 | 4.63 | 2.7E+06 | 2.3E+06 | 3.6E+05 | 5.4E+06 |
| AT3G17390 | AT3G17390.1 | S-adenosylmethionine synthetase family protein | 43 kDa | 24 | 19 | 131 | 106 | 280 | | 0.45 | 0.45 | 7.0E+07 | 5.2E+07 | 6.9E+06 | 1.3E+08 |
| AT2G42740 | | ribosomal protein large subunit 16A | 21 kDa | 35 | 19 | 131 | 90 | 275 | 0.22 | 0.25 | 0.24 | | | | |
| AT1G22870 | AT1G22870.4;AT1G22870.1;AT1G22870.2;AT1G22870.3 | Protein kinase family protein with ARM repeat domain | 100 kDa | 31 | 49 | 131 | 72 | 283 | | | | 4.9E+05 | 1.3E+05 | 0.0E+00 | 6.2E+05 |
| AT1G71880 | AT1G71880.1 | sucrose-proton symporter 1 | 55 kDa | 3 | 0 | 130 | 133 | 266 | | 0.44 | 0.44 | 5.1E+06 | 4.5E+06 | 1.8E+05 | 9.8E+06 |
| AT4G21940 | | calcium-dependent protein kinase 15 | 63 kDa | 0 | 11 | 129 | 134 | 274 | | | | | | | |
| AT3G05590 | AT3G05590.1;AT3G05590.3;AT3G05590.2 | ribosomal protein L18 | 21 kDa | 18 | 56 | 129 | 101 | 304 | | | | 2.6E+07 | 3.0E+07 | 1.9E+07 | 7.5E+07 |
| AT4G17530 | | RAB GTPase homolog 1C | 22 kDa | 28 | 103 | 129 | 85 | 345 | 0.32 | 0.55 | 0.43 | | | | |
| AT5G44790 | AT5G44790.1 | copper-exporting ATPase / responsive-to-antagonist 1 / | 107 kDa | 1 | 1 | 128 | 70 | 200 | | | | 1.6E+07 | 2.3E+07 | 1.0E+06 | 3.9E+07 |
| AT4G24820 | AT4G24820.2;AT4G24820.1 | 26S proteasome, regulatory subunit Rpn7;Proteasome | 44 kDa | 5 | 3 | 127 | 87 | 222 | 0.13 | 0.18 | 0.15 | 3.1E+08 | 3.9E+08 | 4.6E+07 | 7.5E+08 |
| AT5G24710 | AT5G24710.1 | Transducin/WD40 repeat-like superfamily protein | 148 kDa | 1 | 3 | 126 | 125 | 255 | | 0.25 | 0.25 | 8.2E+06 | 9.0E+06 | 2.7E+06 | 2.0E+07 |
| AT5G48450 | AT5G48450.2;AT5G48450.1 | SKU5 similar 3 | 69 kDa | 1 | 3 | 125 | 102 | 231 | 1.96 | 1.86 | 1.91 | 8.9E+06 | 7.4E+06 | 2.4E+05 | 1.7E+07 |

| | | | | | | | | | | | | | | |
|---|---|---|---|---|---|---|---|---|---|---|---|---|---|---|
| AT5G08080 | AT5G08080.4;AT5G08080.1;AT5G08080.3;AT5G08080.5;AT5G08080.2 | syntaxin of plants 132 | 34 kDa | 29 | 51 | 125 | 95 | 300 | | 7.63 | 7.63 | 1.4E+08 | 1.1E+08 | 3.7E+06 | 2.6E+08 |
| AT3G02650 | AT3G02650.1 | Tetratricopeptide repeat (TPR)-like superfamily protein | 64 kDa | 10 | 14 | 125 | 52 | 201 | | 2.56 | 2.56 | 9.0E+07 | 1.2E+08 | 1.8E+07 | 2.3E+08 |
| AT2G42520 | AT2G42520.3;AT2G42520.2;AT2G42520.1 | P-loop containing nucleoside triphosphate hydrolases | 68 kDa | 10 | 29 | 124 | 72 | 235 | | 0.34 | 0.34 | 6.8E+06 | 6.0E+06 | 2.7E+06 | 1.5E+07 |
| AT3G09740 | AT3G09740.1 | syntaxin of plants 71 | 30 kDa | 34 | 42 | 123 | 123 | 322 | 2.66 | 8.43 | 5.55 | 4.2E+07 | 4.0E+07 | 1.6E+06 | 8.3E+07 |
| AT3G07390 | AT3G07390.1 | auxin-responsive family protein | 28 kDa | 16 | 16 | 122 | 71 | 225 | 2.92 | 3.72 | 3.32 | 2.1E+08 | 1.5E+08 | 9.4E+06 | 3.6E+08 |
| AT3G14600 | AT3G14600.1 | Ribosomal protein L18ae/LX family protein | 21 kDa | 0 | 11 | 122 | 57 | 190 | | | | 2.8E+07 | 3.1E+07 | 2.9E+06 | 6.2E+07 |
| AT3G47730 | AT3G47730.1 | ATP-binding cassette A2 | 109 kDa | 36 | 15 | 121 | 86 | 258 | | 7.15 | 7.15 | 7.7E+06 | 8.0E+06 | 8.8E+05 | 1.7E+07 |
| AT3G57490 | AT3G57490.1 | Ribosomal protein S5 family protein | 30 kDa | 0 | 0 | 121 | 61 | 182 | | | | 1.1E+06 | 6.5E+05 | 8.6E+05 | 2.6E+06 |
| AT1G06700 | AT1G06700.3;AT1G06700.2;AT1G06700.1 | Protein kinase superfamily protein | 40 kDa | 15 | 26 | 120 | 122 | 283 | | 6.76 | 6.76 | 6.1E+06 | 3.3E+06 | 1.3E+05 | 9.5E+06 |
| AT5G25100 | AT5G25100.1;AT5G25100.2;AT5G10840.1 | Endomembrane protein 70 protein family | 74 kDa | 6 | 0 | 118 | 99 | 223 | | 0.12 | 0.12 | 4.3E+06 | 3.2E+06 | 3.2E+06 | 1.1E+07 |
| AT2G27530 | AT2G27530.2;AT2G27530.1 | Ribosomal protein L1p/L10e family | 24 kDa | 8 | 36 | 118 | 65 | 227 | 0.09 | 0.37 | 0.23 | 1.7E+08 | 1.5E+08 | 8.6E+07 | 4.1E+08 |
| AT1G30400 | AT1G30400.2;AT1G30400.1 | multidrug resistance-associated protein 1 | 182 kDa | 2 | 0 | 117 | 136 | 255 | 0.27 | 0.17 | 0.22 | 1.1E+07 | 5.4E+06 | 1.0E+06 | 1.7E+07 |
| AT3G57290 | AT3G57290.1 | eukaryotic translation initiation factor 3E | 52 kDa | 25 | 37 | 117 | 102 | 281 | | 0.79 | 0.79 | 2.3E+07 | 3.2E+07 | 1.8E+07 | 7.2E+07 |
| AT3G25290 | AT3G25290.2;AT3G25290.1 | Auxin-responsive family protein | 43 kDa | 12 | 2 | 116 | 120 | 250 | | | | 5.1E+07 | 4.1E+07 | 4.5E+06 | 9.7E+07 |
| AT2G29560 | AT2G29560.1 | cytosolic enolase | 52 kDa | 9 | 27 | 115 | 77 | 228 | | | | 1.3E+06 | 3.2E+06 | 5.8E+05 | 5.1E+06 |
| AT5G06320 | AT5G06320.1 | NDR1/HIN1-like 3 | 26 kDa | 25 | 43 | 114 | 94 | 276 | 2.85 | 7.01 | 4.93 | 1.3E+08 | 9.2E+07 | 7.4E+06 | 2.3E+08 |
| AT4G36130 | AT4G36130.1 | Ribosomal protein L2 family | 28 kDa | 20 | 61 | 114 | 80 | 275 | 0.01 | 0.49 | 0.25 | 3.5E+07 | 3.3E+07 | 5.8E+07 | 1.3E+08 |
| AT4G19006 | AT4G19006.1;AT4G19006.2 | Proteasome component (PCI) domain protein | 44 kDa | 12 | 38 | 114 | 67 | 231 | | 1.96 | 1.96 | 1.3E+08 | 1.2E+08 | 1.4E+07 | 2.6E+08 |
| AT5G15980 | AT5G15980.1 | Pentatricopeptide repeat (PPR) superfamily protein | 75 kDa | 1 | 8 | 114 | 63 | 186 | | 0.99 | 0.99 | 6.8E+07 | 9.7E+07 | 1.3E+07 | 1.8E+08 |
| AT1G04810 | AT1G04810.1 | 26S proteasome regulatory complex, non-ATPase subc | 109 kDa | 0 | 1 | 114 | 62 | 177 | | 0.25 | 0.25 | 4.1E+07 | 3.0E+07 | 3.4E+06 | 7.5E+07 |
| AT2G07360 | AT2G07360.1;AT2G07360.2 | SH3 domain-containing protein | 130 kDa | 1 | 12 | 113 | 93 | 219 | | 1.29 | 1.29 | 6.6E+06 | 8.1E+06 | 2.2E+06 | 1.7E+07 |
| AT4G31700 | AT4G31700.1;AT4G31700.2 | ribosomal protein S6 | 28 kDa | 22 | 60 | 113 | 76 | 271 | 0.10 | 0.49 | 0.29 | 7.8E+07 | 1.0E+08 | 2.1E+07 | 2.0E+08 |
| AT4G02930 | AT4G02930.1 | GTP binding Elongation factor Tu family protein | 49 kDa | 15 | 46 | 113 | 73 | 247 | 0.14 | 0.68 | 0.41 | 1.1E+08 | 1.4E+08 | 2.3E+07 | 2.7E+08 |
| AT1G48480 | AT1G48480.1 | receptor-like kinase 1 | 71 kDa | 10 | 1 | 112 | 102 | 225 | | 5.57 | 5.57 | 4.7E+05 | 5.1E+05 | 2.4E+05 | 1.2E+06 |
| AT5G61820 | AT5G61820.2;AT5G61820.1 | FUNCTIONS IN: molecular_function unknown; INVOLVE | 53 kDa | 27 | 57 | 112 | 92 | 288 | | 4.41 | 4.41 | 6.5E+05 | 6.0E+05 | 8.4E+05 | 2.1E+06 |
| AT1G64740 | | alpha-1 tubulin | 50 kDa | 0 | 47 | 111 | 167 | 325 | | | | | | | |
| AT1G14910 | AT1G14910.2;AT1G14910.1 | ENTH/ANTH/VHS superfamily protein | 76 kDa | 17 | 13 | 111 | 101 | 242 | | | | 1.3E+06 | 1.4E+06 | 2.9E+05 | 2.9E+06 |
| AT3G58570 | AT3G58570.1 | P-loop containing nucleoside triphosphate hydrolases | 69 kDa | 0 | 24 | 111 | 70 | 205 | | 0.47 | 0.47 | 2.4E+06 | 2.5E+06 | 3.9E+05 | 5.3E+06 |
| AT2G32730 | AT2G32730.1 | 26S proteasome regulatory complex, non-ATPase subc | 109 kDa | 3 | 1 | 111 | 68 | 183 | 0.12 | 0.24 | 0.18 | 1.5E+08 | 1.4E+08 | 1.2E+07 | 2.9E+08 |
| AT3G07110 | AT3G07110.1;AT3G07110.2 | Ribosomal protein L13 family protein | 23 kDa | 9 | 42 | 109 | 69 | 229 | 0.10 | 0.27 | 0.19 | 1.6E+08 | 1.3E+08 | 4.8E+07 | 3.3E+08 |
| AT1G67430 | AT1G67430.1;AT1G67430.2 | Ribosomal protein L22p/L17e family protein | 20 kDa | 12 | 12 | 108 | 48 | 180 | | | | 0.0E+00 | 0.0E+00 | 2.9E+06 | 2.9E+06 |
| AT5G15090 | AT5G15090.2;AT5G15090.1 | voltage dependent anion channel 3 | 29 kDa | 25 | 1 | 107 | 131 | 264 | 0.04 | 0.13 | 0.09 | 3.9E+07 | 6.8E+07 | 1.6E+07 | 1.2E+08 |
| AT3G58460 | AT3G58460.1;AT3G58460.2 | RHOMBOID-like protein 15 | 47 kDa | 23 | 23 | 106 | 115 | 267 | 10.15 | 1.95 | 6.05 | 6.6E+06 | 1.6E+07 | 6.3E+06 | 2.9E+07 |
| AT5G10360 | AT5G10360.1;AT5G10360.2 | Ribosomal protein S6e | 28 kDa | 25 | 57 | 106 | 86 | 274 | 0.07 | 0.39 | 0.23 | 1.8E+08 | 3.4E+08 | 5.4E+07 | 5.7E+08 |
| AT3G17420 | AT3G17420.1 | glyoxysomal protein kinase 1 | 53 kDa | 15 | 8 | 106 | 84 | 213 | | 10.87 | 10.87 | 7.2E+06 | 4.3E+06 | 0.0E+00 | 1.1E+07 |
| AT4G18100 | AT4G18100.1 | Ribosomal protein L32e | 16 kDa | 21 | 0 | 106 | 48 | 175 | 0.08 | 0.13 | 0.11 | 8.2E+07 | 1.0E+08 | 4.1E+07 | 2.3E+08 |
| AT1G27400 | AT1G27400.1 | Ribosomal protein L22p/L17e family protein | 20 kDa | 18 | 0 | 106 | 41 | 165 | 0.18 | 0.34 | 0.26 | 8.3E+06 | 9.6E+06 | 1.9E+06 | 2.0E+07 |
| AT3G57330 | AT3G57330.1;AT3G57330.2 | autoinhibited Ca2+-ATPase 11 | 112 kDa | 0 | 0 | 104 | 78 | 182 | | 0.25 | 0.25 | 2.6E+06 | 9.9E+05 | 4.0E+05 | 4.0E+06 |
| AT5G42020 | AT5G42020.3;AT5G42020.1;AT5G42020.2 | Heat shock protein 70 (Hsp 70) family protein | 74 kDa | 11 | 3 | 104 | 73 | 191 | 0.09 | 0.18 | 0.14 | 3.1E+06 | 2.4E+06 | 1.3E+06 | 6.8E+06 |
| AT5G26710 | AT5G26710.1 | Glutamyl/glutaminyl-tRNA synthetase, class Ic | 81 kDa | 3 | 2 | 104 | 66 | 175 | 0.17 | 0.36 | 0.26 | 1.1E+08 | 9.8E+07 | 2.5E+06 | 2.2E+08 |
| AT5G45620 | AT5G45620.1;AT5G45620.2 | Proteasome component (PCI) domain protein | 44 kDa | 13 | 23 | 104 | 59 | 199 | | 1.73 | 1.73 | 9.8E+07 | 1.1E+08 | 8.7E+06 | 2.2E+08 |
| AT5G22140 | AT5G22140.1;AT5G22140.2 | FAD/NAD(P)-binding oxidoreductase family protein | 40 kDa | 5 | 17 | 103 | 128 | 253 | | | | 1.9E+07 | 7.4E+06 | 5.6E+05 | 2.7E+07 |
| AT5G27850 | AT5G27850.1;AT5G27850.2 | Ribosomal protein L18e/L15 superfamily protein | 21 kDa | 22 | 53 | 103 | 106 | 284 | 0.14 | 0.36 | 0.25 | 4.5E+07 | 6.1E+07 | 1.7E+07 | 1.2E+08 |
| AT4G30260 | AT4G30260.3;AT4G30260.1;AT4G30260.2 | Integral membrane Yip1 family protein | 30 kDa | 8 | 7 | 103 | 78 | 196 | | | | 8.0E+06 | 1.1E+07 | 2.5E+06 | 2.2E+07 |
| AT1G32050 | AT1G32050.1 | SCAMP family protein | 30 kDa | 13 | 24 | 103 | 65 | 205 | | | | 9.7E+07 | 7.2E+07 | 2.4E+06 | 1.7E+08 |
| AT4G13930 | AT4G13930.1 | serine hydroxymethyltransferase 4 | 52 kDa | 30 | 47 | 102 | 82 | 261 | 0.18 | 0.24 | 0.21 | 2.6E+08 | 3.8E+08 | 1.2E+07 | 6.6E+08 |
| AT5G61740 | AT5G61740.3;AT5G61740.2;AT5G61740.1 | ABC2 homolog 14 | 94 kDa | 15 | 1 | 102 | 77 | 195 | | | | 3.3E+06 | 2.4E+06 | 0.0E+00 | 5.8E+06 |
| AT2G39450 | AT2G39450.1 | Cation efflux family protein | 45 kDa | 4 | 1 | 102 | 75 | 182 | | 7.11 | 7.11 | 2.7E+06 | 2.7E+06 | 2.3E+05 | 5.6E+06 |
| AT3G01780 | AT3G01780.1;AT3G01780.2 | ARM repeat superfamily protein | 131 kDa | 1 | 0 | 101 | 107 | 209 | | 0.53 | 0.53 | 6.1E+06 | 5.3E+06 | 1.3E+06 | 1.3E+07 |
| AT5G67130 | AT5G67130.1 | PLC-like phosphodiesterases superfamily protein | 47 kDa | 2 | 11 | 101 | 99 | 213 | | 7.68 | 7.68 | 2.2E+07 | 1.5E+07 | 0.0E+00 | 3.7E+07 |
| AT3G46060 | AT3G46060.3;AT3G46060.2;AT3G46060.1 | RAB GTPase homolog 8A | 24 kDa | 51 | 91 | 101 | 82 | 325 | 0.58 | | 0.58 | 9.4E+06 | 7.8E+06 | 0.0E+00 | 1.7E+07 |
| AT4G05520 | AT4G05520.1;AT4G05520.2 | EPS15 homology domain 2 | 61 kDa | 117 | 52 | 101 | 81 | 351 | | | | 2.8E+07 | 3.4E+07 | 4.6E+04 | 6.2E+07 |
| AT5G62630 | AT5G62630.1 | hipl2 protein precursor | 76 kDa | 8 | 17 | 101 | 78 | 204 | | 3.56 | 3.56 | 2.3E+07 | 1.7E+07 | 1.2E+06 | 4.1E+07 |
| AT5G28540 | AT5G28540.1 | heat shock protein 70 (Hsp 70) family protein | 74 kDa | 12 | 3 | 100 | 72 | 187 | 0.10 | 0.29 | 0.20 | 3.1E+05 | 5.1E+05 | 0.0E+00 | 8.2E+05 |
| AT1G05150 | | Calcium-binding tetratricopeptide family protein | 90 kDa | 4 | 0 | 100 | 66 | 170 | | 5.58 | 5.58 | | | | |
| AT5G62740 | AT5G62740.1 | SPFH/Band 7/PHB domain-containing membrane-assoc | 31 kDa | 58 | 112 | 99 | 146 | 415 | 1.02 | 5.27 | 3.15 | 3.5E+07 | 2.8E+07 | 3.4E+06 | 6.6E+07 |
| AT2G17930 | AT2G17930.1 | Phosphatidylinositol 3- and 4-kinase family protein wit | 436 kDa | 0 | 0 | 99 | 123 | 222 | | 0.36 | 0.36 | 4.3E+04 | 4.9E+04 | 1.8E+04 | 1.1E+05 |
| AT2G25800 | AT2G25800.1 | Protein of unknown function (DUF810) | 110 kDa | 57 | 18 | 99 | 72 | 246 | 3.22 | 3.42 | 3.32 | 2.1E+06 | 3.1E+06 | 0.0E+00 | 5.3E+06 |
| AT1G53500 | AT1G53500.1 | NAD-dependent epimerase/dehydratase family protein | 75 kDa | 0 | 4 | 99 | 40 | 143 | | 0.32 | 0.32 | 3.4E+06 | 3.0E+06 | 3.1E+05 | 6.7E+06 |
| AT5G55730 | AT5G55730.2;AT5G55730.1 | FASCICLIN-like arabinogalactan 1 | 45 kDa | 4 | 10 | 98 | 85 | 197 | | 6.31 | 6.31 | 7.5E+07 | 7.7E+07 | 5.4E+06 | 1.6E+08 |
| AT4G34450 | AT4G34450.1 | coatomer gamma-2 subunit, putative / gamma-2 coat p | 98 kDa | 18 | 10 | 98 | 84 | 210 | 0.17 | 0.33 | 0.25 | 8.2E+06 | 1.0E+07 | 1.2E+06 | 2.0E+07 |
| AT1G66250 | AT1G66250.1 | O-Glycosyl hydrolases family 17 protein | 54 kDa | 8 | 2 | 98 | 56 | 164 | | 4.70 | 4.70 | 1.1E+07 | 7.8E+06 | 3.8E+05 | 1.9E+07 |
| AT5G54960 | AT5G54960.1 | pyruvate decarboxylase-2 | 66 kDa | 62 | 146 | 97 | 512 | 817 | 0.10 | 1.38 | 0.74 | 2.6E+08 | 7.0E+08 | 1.0E+07 | 9.8E+08 |
| AT5G54310 | AT5G54310.1 | ARF-GAP domain 5 | 53 kDa | 13 | 12 | 97 | 74 | 196 | | | | 2.9E+06 | 2.5E+06 | 4.6E+06 | 9.9E+06 |
| AT2G27810 | AT2G27810.1;AT2G27810.4;AT2G27810.3;AT2G27810.2 | nucleobase-ascorbate transporter 12 | 77 kDa | 0 | 0 | 97 | 53 | 150 | | | | 1.0E+07 | 1.0E+07 | 4.7E+04 | 2.0E+07 |
| AT1G72370 | AT1G72370.2;AT1G72370.1 | 40s ribosomal protein SA | 32 kDa | 26 | 69 | 96 | 82 | 273 | 0.21 | 0.45 | 0.33 | 3.2E+08 | 2.8E+08 | 7.8E+07 | 6.8E+08 |
| AT5G10840 | | Endomembrane protein 70 protein family | 74 kDa | 6 | 0 | 96 | 77 | 179 | | 0.07 | 0.07 | | | | |
| AT1G52780 | AT1G52780.1 | Protein of unknown function (DUF2921) | 120 kDa | 0 | 4 | 96 | 73 | 173 | | | | 0.0E+00 | 1.7E+05 | 2.9E+05 | 4.6E+05 |
| AT4G00100 | AT4G00100.1 | ribosomal protein S13A | 17 kDa | 15 | 0 | 96 | 60 | 171 | | 0.12 | 0.12 | 9.1E+06 | 1.3E+07 | 2.5E+06 | 2.5E+07 |
| AT3G45600 | AT3G45600.1;AT3G45600.2 | tetraspanin3 | 32 kDa | 36 | 97 | 96 | 59 | 288 | 3.91 | 12.42 | 8.16 | 6.5E+07 | 3.9E+07 | 2.9E+06 | 1.1E+08 |
| AT1G52980 | AT1G52980.1 | GTP-binding family protein | 65 kDa | 0 | 0 | 96 | 58 | 154 | | | | 1.3E+07 | 1.5E+07 | 5.3E+06 | 3.3E+07 |
| AT1G70600 | AT1G70600.1 | Ribosomal protein L18e/L15 superfamily protein | 16 kDa | 26 | 6 | 96 | 50 | 178 | 0.10 | | 0.10 | 6.8E+07 | 1.3E+08 | 5.8E+06 | 2.0E+08 |
| AT5G46430 | AT5G46430.2;AT5G46430.1 | Ribosomal protein L32e | 15 kDa | 19 | 0 | 96 | 41 | 156 | | 0.41 | 0.41 | 1.0E+08 | 1.4E+08 | 2.3E+07 | 2.6E+08 |
| AT5G45750 | AT5G45750.1 | RAB GTPase homolog A1C | 24 kDa | 38 | 48 | 95 | 104 | 285 | | | | 5.0E+07 | 2.3E+07 | 2.2E+06 | 7.5E+07 |
| AT1G69840 | AT1G69840.7;AT1G69840.6;AT1G69840.5;AT1G69840.4;AT1G69840.3;AT1G69840.2;AT1G69840.1 | SPFH/Band 7/PHB domain-containing membrane-assoc | 31 kDa | 43 | 41 | 95 | 79 | 258 | 1.36 | 4.20 | 2.78 | 2.3E+07 | 1.2E+07 | 9.2E+05 | 3.6E+07 |
| AT2G38040 | AT2G38040.2;AT2G38040.1 | acetyl Co-enzyme a carboxylase carboxyltransferase alp | 85 kDa | 0 | 3 | 95 | 69 | 167 | | 0.49 | 0.49 | 2.2E+07 | 2.1E+07 | 4.5E+06 | 4.7E+07 |
| AT3G03780 | AT3G03780.3;AT3G03780.2;AT3G03780.1 | methionine synthase 2 | 85 kDa | 0 | 18 | 94 | 94 | 206 | | | | 9.9E+05 | 7.7E+05 | 0.0E+00 | 1.8E+06 |
| AT1G02780 | AT1G02780.1 | Ribosomal protein L19e family protein | 25 kDa | 12 | 37 | 94 | 47 | 190 | 0.11 | 0.49 | 0.30 | 2.5E+08 | 4.2E+08 | 7.1E+07 | 7.4E+08 |
| AT1G75990 | AT1G75990.1 | PAM domain (PCI/PINT associated module) protein | 56 kDa | 14 | 14 | 93 | 76 | 197 | | 0.88 | 0.88 | 5.3E+06 | 7.3E+06 | 3.8E+05 | 1.3E+07 |
| AT5G59840 | | Ras-related small GTP-binding family protein | 24 kDa | 52 | 87 | 93 | 73 | 305 | | 0.69 | 0.69 | | | | |
| AT4G13350 | AT4G13350.2;AT4G13350.1 | NSP (nuclear shuttle protein)-interacting GTPase | 65 kDa | 21 | 51 | 93 | 61 | 226 | | 6.62 | 6.62 | 3.3E+06 | 9.0E+06 | 2.0E+06 | 1.4E+07 |
| AT3G05280 | AT3G05280.1 | Integral membrane Yip1 family protein | 31 kDa | 11 | 7 | 93 | 56 | 167 | | 4.40 | 4.40 | 3.3E+06 | 4.7E+06 | 2.2E+06 | 1.0E+07 |
| AT3G55280 | | ribosomal protein L23AB | 17 kDa | 15 | 11 | 93 | 49 | 168 | | | | | | | |

| | | | | | | | | | | | | | | | |
|---|---|---|---|---|---|---|---|---|---|---|---|---|---|---|---|
| AT1G13440 | | glyceraldehyde-3-phosphate dehydrogenase C2 | 37 kDa | 32 | 58 | 92 | 182 | 364 | 0.60 | 3.82 | 2.21 | | | | |
| AT2G21410 | AT2G21410.1 | vacuolar proton ATPase A2 | 93 kDa | 0 | 4 | 92 | 85 | 181 | 0.67 | 0.21 | 0.44 | 9.0E+05 | 4.8E+05 | 0.0E+00 | 1.4E+06 |
| AT5G42090 | AT5G42090.1 | Lung seven transmembrane receptor family protein | 50 kDa | 13 | 10 | 92 | 70 | 185 | | | | 1.7E+06 | 4.6E+05 | 4.3E+06 | 6.4E+06 |
| AT3G23750 | AT3G23750.1 | Leucine-rich repeat protein kinase family protein | 100 kDa | 14 | 4 | 91 | 103 | 212 | | | | 7.0E+06 | 3.0E+06 | 4.4E+04 | 1.0E+07 |
| AT2G47000 | AT2G47000.1;AT2G47000.5;AT2G47000.6;AT2G47000.2;AT2G47000.3;AT2G47000.7 | ATP binding cassette subfamily B4 | 139 kDa | 5 | 0 | 91 | 80 | 176 | | | | 5.6E+05 | 4.6E+05 | 0.0E+00 | 1.0E+06 |
| AT4G26510 | | uridine kinase-like 4 | 53 kDa | 0 | 4 | 91 | 71 | 166 | | | | | | | |
| AT3G58730 | AT3G58730.1 | vacuolar ATP synthase subunit D (VATD) / V-ATPase D su | 29 kDa | 21 | 36 | 91 | 58 | 206 | 0.31 | 2.26 | 1.29 | 3.6E+07 | 2.0E+07 | 3.9E+06 | 6.0E+07 |
| AT2G03270 | AT2G03270.1 | DNA-binding protein, putative | 71 kDa | 1 | 0 | 91 | 56 | 148 | 0.11 | 0.31 | 0.21 | 7.1E+06 | 3.6E+06 | 2.7E+06 | 1.3E+07 |
| AT1G09590 | | Translation protein SH3-like family protein | 19 kDa | 9 | 1 | 91 | 44 | 145 | 0.12 | 0.39 | 0.26 | | | | |
| AT3G04120 | AT3G04120.1;AT1G13440.1;AT1G13440.2 | glyceraldehyde-3-phosphate dehydrogenase C subunit | 37 kDa | 0 | 58 | 90 | 186 | 334 | | | | 4.8E+07 | 3.5E+07 | 4.2E+05 | 8.2E+07 |
| AT5G58480 | AT5G58480.1 | O-Glycosyl hydrolases family 17 protein | 52 kDa | 2 | 0 | 90 | 80 | 172 | | | | 7.7E+06 | 6.2E+06 | 6.8E+05 | 1.5E+07 |
| AT4G01850 | AT4G01850.2;AT4G01850.1 | S-adenosylmethionine synthetase 2 | 43 kDa | 10 | 13 | 90 | 63 | 176 | | | | 4.7E+07 | 2.1E+07 | 1.8E+06 | 7.0E+07 |
| AT3G54030 | AT3G54030.1 | Protein kinase protein with tetratricopeptide repeat d | 55 kDa | 15 | 26 | 89 | 85 | 215 | 1.53 | 2.90 | 2.22 | 1.7E+07 | 9.2E+06 | 8.1E+04 | 2.7E+07 |
| AT2G01660 | AT2G01660.2;AT2G01660.3;AT2G01660.1 | plasmodesmata-located protein 6 | 30 kDa | 3 | 14 | 89 | 73 | 179 | | 6.50 | 6.50 | 2.3E+07 | 1.9E+07 | 2.2E+05 | 4.2E+07 |
| AT1G11260 | AT1G11260.1 | sugar transporter 1 | 58 kDa | 1 | 8 | 89 | 71 | 169 | | | | 1.2E+07 | 5.2E+06 | 2.8E+05 | 1.8E+07 |
| AT3G07020 | AT3G07020.1;AT3G07020.2;AT3G07020.3 | UDP-Glycosyltransferase superfamily protein | 69 kDa | 10 | 12 | 89 | 61 | 172 | | 2.23 | 2.23 | 9.6E+06 | 5.3E+06 | 7.6E+05 | 1.6E+07 |
| AT3G50590 | AT3G50590.2;AT3G50590.1 | Transducin/WD40 repeat-like superfamily protein | 176 kDa | 1 | 3 | 88 | 91 | 183 | | 0.20 | 0.20 | 4.4E+06 | 4.2E+06 | 7.3E+05 | 9.4E+06 |
| AT5G27640 | AT5G27640.1;AT5G27640.3;AT5G27640.2 | translation initiation factor 3B1 | 82 kDa | 4 | 0 | 88 | 57 | 149 | 0.08 | 0.32 | 0.20 | 1.2E+07 | 1.1E+07 | 1.4E+06 | 2.5E+07 |
| AT5G28060 | AT5G28060.1 | Ribosomal protein S24e family protein | 15 kDa | 9 | 1 | 88 | 42 | 140 | 0.34 | 0.57 | 0.45 | 5.4E+08 | 5.8E+08 | 1.6E+08 | 1.3E+09 |
| AT1G61900 | AT1G61900.2;AT1G61900.3;AT1G61900.1 | unknown protein; FUNCTIONS IN: molecular_function | 45 kDa | 13 | 1 | 87 | 52 | 153 | | | | 5.0E+07 | 4.5E+07 | 1.8E+06 | 9.7E+07 |
| AT1G48830 | AT1G48830.2;AT1G48830.1 | Ribosomal protein S7e family protein | 22 kDa | 5 | 14 | 87 | 47 | 153 | 0.21 | 0.27 | 0.24 | 1.5E+08 | 1.6E+08 | 3.5E+07 | 3.4E+08 |
| AT1G04760 | | vesicle-associated membrane protein 726 | 25 kDa | 15 | 36 | 87 | 35 | 173 | | | | | | | |
| AT3G54300 | AT3G54300.3;AT3G54300.2;AT3G54300.1 | vesicle-associated membrane protein 727 | 27 kDa | 31 | 103 | 86 | 141 | 361 | 2.54 | 8.06 | 5.30 | 5.4E+07 | 7.4E+07 | 6.3E+06 | 1.3E+08 |
| AT5G58440 | AT5G58440.1 | sorting nexin 2A | ? | 0 | 12 | 86 | 83 | 181 | | | | 1.7E+06 | 2.4E+06 | 4.7E+05 | 4.6E+06 |
| AT1G11130 | AT1G11130.2;AT1G11130.1 | Leucine-rich repeat protein kinase family protein | 84 kDa | 11 | 6 | 86 | 74 | 177 | | 5.32 | 5.32 | 1.2E+07 | 8.9E+06 | 7.6E+04 | 2.1E+07 |
| AT3G03140 | | Tudor/PWWP/MBT superfamily protein | 87 kDa | 0 | 1 | 86 | 62 | 149 | | | | | | | |
| AT5G57460 | AT5G57460.1 | unknown protein; FUNCTIONS IN: molecular_function | 71 kDa | 1 | 2 | 86 | 59 | 148 | | 1.38 | 1.38 | 3.4E+06 | 4.3E+06 | 9.5E+05 | 8.7E+06 |
| AT1G27190 | AT1G27190.1 | Leucine-rich repeat protein kinase family protein | 65 kDa | 4 | 8 | 86 | 51 | 149 | | 0.74 | 0.74 | 3.1E+06 | 3.8E+05 | 0.0E+00 | 3.4E+06 |
| AT4G19210 | AT4G19210.1 | RNAse l inhibitor protein 2 | 68 kDa | 0 | 0 | 86 | 47 | 133 | 0.15 | 0.17 | 0.16 | 1.4E+07 | 1.5E+07 | 1.6E+07 | 4.4E+07 |
| ATMG00090 | ATMG00090.1 | structural constituent of ribosome;protein binding | 65 kDa | 7 | 0 | 86 | 37 | 130 | | 1.40 | 1.40 | 5.5E+07 | 6.0E+07 | 9.6E+06 | 1.2E+08 |
| AT3G11250 | AT3G11250.1 | Ribosomal protein L10 family protein | 34 kDa | 10 | 40 | 85 | 84 | 219 | 0.17 | | 0.17 | 4.5E+07 | 5.6E+07 | 2.9E+07 | 1.3E+08 |
| AT5G39850 | AT5G39850.1 | Ribosomal protein S4 | 23 kDa | 0 | 35 | 84 | 61 | 180 | 0.44 | | 0.44 | 4.1E+07 | 3.6E+07 | 1.4E+07 | 9.1E+07 |
| AT1G32090 | AT1G32090.1 | early-responsive to dehydration stress protein (ERD4) | 93 kDa | 2 | 0 | 84 | 53 | 139 | | | | 1.2E+07 | 8.2E+06 | 7.0E+04 | 2.1E+07 |
| AT2G09990 | AT2G09990.1 | Ribosomal protein S5 domain 2-like superfamily protei | 17 kDa | 12 | 2 | 83 | 51 | 148 | 0.09 | 0.09 | 0.09 | 4.4E+07 | 2.6E+07 | 8.1E+06 | 7.8E+07 |
| AT5G25780 | AT5G25780.1 | eukaryotic translation initiation factor 3B-2 | 82 kDa | 0 | 0 | 83 | 46 | 129 | | | | 0.0E+00 | 0.0E+00 | 2.9E+05 | 2.9E+05 |
| AT1G59900 | AT1G59900.1;AT1G59900.2 | pyruvate dehydrogenase complex E1 alpha subunit | 43 kDa | 21 | 12 | 83 | 45 | 161 | 0.18 | 0.36 | 0.27 | 1.5E+08 | 2.7E+08 | 2.8E+07 | 4.4E+08 |
| AT3G22230 | AT3G22230.1 | Ribosomal L27e protein family | 16 kDa | 12 | 1 | 83 | 31 | 127 | | 0.16 | 0.16 | 1.1E+08 | 9.5E+07 | 2.4E+07 | 2.3E+08 |
| AT2G42910 | AT2G42910.1 | Phosphoribosyltransferase family protein | 38 kDa | 17 | 2 | 83 | 22 | 124 | | 6.85 | 6.85 | 2.1E+06 | 3.9E+06 | 0.0E+00 | 6.0E+06 |
| AT1G04480 | | Ribosomal protein L14p/L23e family protein | 15 kDa | 10 | 0 | 83 | 20 | 113 | | 0.26 | 0.26 | | | | |
| AT1G56110 | AT1G56110.1 | homolog of nucleolar protein NOP56 | 59 kDa | 0 | 6 | 82 | 61 | 149 | | 0.28 | 0.28 | 1.9E+07 | 1.9E+07 | 1.2E+07 | 5.1E+07 |
| AT5G02610 | AT5G02610.1;AT5G02610.2 | Ribosomal L29 family protein | 17 kDa | 6 | 0 | 82 | 58 | 146 | 0.60 | | 0.60 | 3.0E+07 | 5.1E+07 | 1.6E+07 | 9.7E+07 |
| AT3G09500 | AT3G09500.1 | Ribosomal L29 family protein | 14 kDa | 7 | 0 | 82 | 44 | 133 | 0.20 | 0.18 | 0.19 | 1.4E+07 | 1.9E+07 | 3.9E+06 | 3.7E+07 |
| AT4G15000 | AT4G15000.1;AT4G15000.2 | Ribosomal L27e protein family | 16 kDa | 16 | 1 | 82 | 44 | 143 | 0.14 | 0.34 | 0.24 | 1.6E+08 | 2.1E+08 | 2.2E+07 | 3.9E+08 |
| AT3G60245 | AT3G60245.1 | Zinc-binding ribosomal protein family protein | 10 kDa | 6 | 2 | 82 | 14 | 104 | | 0.24 | 0.24 | 2.7E+08 | 3.0E+08 | 5.0E+07 | 6.2E+08 |
| AT3G46290 | AT3G46290.1 | hercules receptor kinase 1 | 91 kDa | 12 | 6 | 81 | 91 | 190 | | 3.16 | 3.16 | 9.4E+06 | 6.1E+06 | 0.0E+00 | 1.5E+07 |
| AT4G18800 | AT4G18800.1 | RAB GTPase homolog A1D | 24 kDa | 23 | 44 | 81 | 90 | 238 | | 2.86 | 2.86 | 4.4E+06 | 1.7E+06 | 0.0E+00 | 6.0E+06 |
| AT5G18500 | AT5G18500.6;AT5G18500.5;AT5G18500.4;AT5G18500.3;AT5G18500.2;AT5G18500.1 | Protein kinase superfamily protein | 54 kDa | 3 | 5 | 81 | 76 | 165 | | | | 1.7E+06 | 9.9E+05 | 0.0E+00 | 2.7E+06 |
| AT5G23540 | AT5G23540.1;AT5G23540.2 | Mov34/MPN/PAD-1 family protein | 34 kDa | 14 | 33 | 81 | 64 | 192 | 0.15 | 0.82 | 0.48 | 4.6E+08 | 6.5E+08 | 7.4E+07 | 1.2E+09 |
| AT2G07698 | AT2G07698.1;ATMG01190.1 | ATPase, F1 complex, alpha subunit protein | 86 kDa | 2 | 2 | 81 | 61 | 146 | 0.17 | 0.19 | 0.18 | 1.1E+07 | 1.5E+07 | 8.7E+05 | 2.7E+07 |
| AT5G55920 | AT5G55920.1 | S-adenosyl-L-methionine-dependent methyltransferase | 77 kDa | 2 | 0 | 81 | 60 | 143 | | 0.39 | 0.39 | 6.5E+07 | 5.4E+07 | 1.2E+07 | 1.3E+08 |
| AT1G16920 | AT1G16920.1 | RAB GTPase homolog A1B | 24 kDa | 31 | 39 | 80 | 97 | 247 | | | | 6.4E+06 | 4.3E+06 | 2.1E+05 | 1.1E+07 |
| AT5G22330 | AT5G22330.1 | P-loop containing nucleoside triphosphate hydrolases | 50 kDa | 4 | 16 | 80 | 81 | 181 | 0.26 | 0.59 | 0.42 | 1.6E+07 | 1.1E+07 | 3.7E+06 | 3.1E+07 |
| AT3G63260 | AT3G63260.1;AT3G63260.2 | Protein kinase superfamily protein | 43 kDa | 5 | 16 | 80 | 50 | 151 | | | | 2.9E+06 | 1.4E+06 | 6.9E+04 | 4.4E+06 |
| AT2G37990 | AT2G37990.1 | ribosome biogenesis regulatory protein (RRS1) family p | 36 kDa | 2 | 1 | 80 | 49 | 132 | 0.16 | 2.20 | 1.18 | 6.4E+07 | 7.5E+07 | 1.3E+07 | 1.5E+08 |
| AT3G04920 | AT3G04920.1;AT3G04920.2 | Ribosomal protein S24e family protein | 15 kDa | 11 | 0 | 80 | 44 | 135 | 0.29 | 0.29 | 0.29 | 4.7E+08 | 3.8E+08 | 1.0E+08 | 9.5E+08 |
| AT5G18380 | | Ribosomal protein S5 domain 2-like superfamily protei | 17 kDa | 13 | 4 | 79 | 56 | 152 | 0.38 | 0.32 | 0.35 | | | | |
| AT1G64520 | AT1G64520.1 | regulatory particle non-ATPase 12A | 31 kDa | 20 | 48 | 79 | 54 | 201 | 0.22 | 2.15 | 1.18 | 4.8E+08 | 4.9E+08 | 5.3E+07 | 1.0E+09 |
| AT1G78570 | AT1G78570.1 | rhamnose biosynthesis 1 | 75 kDa | 0 | 4 | 79 | 34 | 117 | 0.11 | | 0.11 | 3.4E+06 | 4.0E+06 | 1.4E+05 | 7.5E+06 |
| AT1G64200 | AT1G64200.1;AT1G64200.2;AT1G64200.3 | vacuolar H+-ATPase subunit E isoform 3 | 27 kDa | 0 | 34 | 78 | 116 | 228 | | 1.48 | 1.48 | 4.4E+06 | 7.4E+05 | 2.1E+06 | 7.2E+06 |
| AT1G63500 | AT1G63500.1 | Protein kinase protein with tetratricopeptide repeat d | 55 kDa | 19 | 31 | 78 | 83 | 211 | | | | 1.3E+07 | 8.6E+06 | 2.8E+05 | 2.2E+07 |
| AT3G49910 | AT3G49910.1 | Translation protein SH3-like family protein | 17 kDa | 24 | 3 | 78 | 63 | 168 | 0.12 | 0.48 | 0.30 | 5.2E+08 | 7.2E+08 | 2.5E+08 | 1.5E+09 |
| AT2G24360 | AT2G24360.1 | Protein kinase superfamily protein | 46 kDa | 13 | 8 | 78 | 61 | 160 | | | | 2.6E+06 | 9.5E+05 | 1.8E+05 | 3.8E+06 |
| AT5G49720 | | glycosyl hydrolase 9A1 | 69 kDa | 6 | 8 | 78 | 59 | 151 | | 4.53 | 4.53 | | | | |
| AT1G02520 | | P-glycoprotein 11 | 138 kDa | 0 | 0 | 78 | 50 | 128 | | | | | | | |
| AT2G30050 | AT2G30050.1 | transducin family protein / WD-40 repeat family protei | 33 kDa | 13 | 24 | 78 | 46 | 161 | | 3.58 | 3.58 | 5.8E+06 | 7.7E+06 | 9.8E+05 | 1.5E+07 |
| AT3G14790 | AT3G14790.1;AT3G14790.2 | rhamnose biosynthesis 3 | 75 kDa | 0 | 4 | 78 | 32 | 114 | | | | 3.4E+06 | 3.2E+06 | 2.2E+06 | 8.7E+06 |
| AT2G38770 | AT2G38770.1 | P-loop containing nucleoside triphosphate hydrolases | 173 kDa | 0 | 0 | 77 | 52 | 129 | | 0.16 | 0.16 | 2.7E+07 | 3.0E+07 | 5.8E+06 | 6.3E+07 |
| AT3G02080 | AT3G02080.1 | Ribosomal protein S19e family protein | 16 kDa | 11 | 0 | 77 | 25 | 113 | 0.27 | 0.26 | 0.26 | 6.9E+07 | 6.1E+07 | 2.1E+07 | 1.5E+08 |
| AT1G07870 | AT1G07870.1;AT1G07870.2 | Protein kinase superfamily protein | ? | 2 | 6 | 76 | 88 | 172 | | 2.31 | 2.31 | 1.8E+06 | 9.6E+05 | 0.0E+00 | 2.8E+06 |
| AT2G22125 | AT2G22125.1 | binding | 231 kDa | 0 | 1 | 76 | 82 | 159 | | 0.25 | 0.25 | 6.7E+05 | 4.7E+05 | 8.5E+03 | 1.1E+06 |
| AT5G08690 | AT5G08690.1;AT5G08670.1;AT5G08680.1 | ATP synthase alpha/beta family protein | 60 kDa | 17 | 17 | 76 | 80 | 190 | | | | 1.3E+07 | 2.1E+07 | 3.1E+06 | 3.7E+07 |
| AT5G03520 | AT5G03520.1;AT5G03520.2 | RAB GTPase homolog 8C | 24 kDa | 38 | 62 | 76 | 54 | 230 | | | | 3.2E+06 | 2.9E+06 | 0.0E+00 | 6.1E+06 |
| AT5G46280 | AT5G46280.1;AT5G46280.2;AT5G46280.3;AT5G46280.4 | Minichromosome maintenance (MCM2/3/5) family pr | 86 kDa | 0 | 0 | 76 | 51 | 127 | | 0.29 | 0.29 | 1.7E+07 | 4.3E+07 | 3.8E+06 | 6.4E+07 |
| AT1G19520 | AT1G19520.1 | pentatricopeptide (PPR) repeat-containing protein | 83 kDa | 5 | 0 | 76 | 39 | 120 | | 0.61 | 0.61 | 7.6E+07 | 7.8E+07 | 7.5E+06 | 1.6E+08 |
| AT1G01510 | | NAD(P)-binding Rossmann-fold superfamily protein | 70 kDa | 0 | 3 | 75 | 66 | 144 | | | | | | | |
| AT5G25040 | AT5G25040.1;AT5G25040.4;AT5G25040.2;AT5G25040.5;AT5G25040.3 | Major facilitator superfamily protein | 58 kDa | 15 | 3 | 75 | 60 | 153 | | 1.85 | 1.85 | 1.2E+06 | 1.1E+06 | 4.6E+04 | 2.3E+06 |
| AT2G40360 | AT2G40360.2;AT2G40360.1 | Transducin/WD40 repeat-like superfamily protein | 85 kDa | 0 | 0 | 75 | 55 | 130 | | 0.24 | 0.24 | 1.1E+07 | 1.2E+07 | 4.7E+06 | 2.7E+07 |
| AT5G14520 | AT5G14520.1 | pescadillo-related | 68 kDa | 0 | 0 | 75 | 50 | 125 | | 0.23 | 0.23 | 1.1E+07 | 2.1E+07 | 8.3E+06 | 4.1E+07 |
| AT1G41880 | AT1G41880.2;AT1G41880.1 | Ribosomal protein L35Ae family protein | 13 kDa | 8 | 1 | 75 | 32 | 116 | | 1.82 | 1.82 | 2.3E+08 | 2.7E+08 | 8.8E+06 | 5.1E+08 |

| | | | | | | | | | | | | | | | |
|---|---|---|---|---|---|---|---|---|---|---|---|---|---|---|---|
| AT2G30740 | AT2G30740.9;AT2G30740.8;AT2G30740.7;AT2G30740.6;AT2G30740.5;AT2G30740.4;AT2G30740.3;AT2G30740.2;AT2G30740.15;AT2G30740.14;AT2G30740.13;AT2G30740.12;AT2G30740.11;AT2G30740.10;AT2G30740.1 | Protein kinase superfamily protein | 41 kDa | 17 | 20 | 74 | 59 | 170 | | | | 2.5E+06 | 9.9E+05 | 0.0E+00 | 3.5E+06 |
| AT3G55410 | AT3G55410.2;AT3G55410.1 | 2-oxoglutarate dehydrogenase, E1 component | 115 kDa | 0 | 0 | 74 | 49 | 123 | | 0.23 | 0.23 | 1.6E+07 | 3.3E+07 | 3.7E+06 | 5.2E+07 |
| AT1G16870 | AT1G16870.1 | mitochondrial 28S ribosomal protein S29-related | 54 kDa | 13 | 10 | 74 | 44 | 141 | | 1.76 | 1.76 | 6.8E+07 | 9.8E+07 | 7.8E+06 | 1.7E+08 |
| AT3G08510 | AT3G08510.4;AT3G08510.3;AT3G08510.2;AT3G08510.1 | phospholipase C 2 | 66 kDa | 0 | 2 | 74 | 31 | 107 | | | | 2.1E+06 | 3.4E+05 | 0.0E+00 | 2.4E+06 |
| AT5G18520 | | Lung seven transmembrane receptor family protein | 50 kDa | 10 | 0 | 73 | 139 | 222 | | 0.26 | 0.26 | | | | |
| AT3G15060 | AT3G15060.1 | RAB GTPase homolog A1G | 24 kDa | 22 | 38 | 73 | 83 | 216 | | 2.86 | 2.86 | 3.5E+06 | 1.5E+06 | 0.0E+00 | 5.0E+06 |
| AT1G02500 | AT1G02500.2;AT1G02500.1 | S-adenosylmethionine synthetase 1 | 43 kDa | 0 | 0 | 73 | 61 | 134 | 0.18 | | 0.18 | 3.0E+06 | 4.5E+06 | 0.0E+00 | 7.5E+06 |
| AT4G04740 | | calcium-dependent protein kinase 23 | 59 kDa | 8 | 12 | 73 | 60 | 153 | | | | | | | |
| AT1G70520 | AT1G70520.1 | cysteine-rich RLK (RECEPTOR-like protein kinase) 2 | 72 kDa | 0 | 2 | 73 | 42 | 117 | | 10.35 | 10.35 | 9.9E+05 | 5.9E+05 | 1.8E+04 | 1.6E+06 |
| AT1G12360 | AT1G12360.1 | Sec1/munc18-like (SM) proteins superfamily | 75 kDa | 23 | 9 | 73 | 37 | 142 | | | | 9.5E+05 | 1.0E+06 | 0.0E+00 | 2.0E+06 |
| AT2G34730 | | myosin heavy chain-related | 95 kDa | 0 | 0 | 73 | 17 | 90 | | | | | | | |
| AT3G02740 | AT3G02740.2;AT3G02740.1 | Eukaryotic aspartyl protease family protein | 53 kDa | 12 | 6 | 72 | 163 | 253 | | 9.63 | 9.63 | 5.5E+07 | 2.7E+07 | 1.4E+06 | 8.4E+07 |
| AT4G16120 | AT4G16120.1 | COBRA-like protein-7 precursor | 73 kDa | 5 | 11 | 72 | 82 | 170 | | | | 6.7E+05 | 1.2E+06 | 8.3E+04 | 1.9E+06 |
| AT1G31970 | AT1G31970.1 | DEA(D/H)-box RNA helicase family protein | 60 kDa | 0 | 3 | 72 | 53 | 128 | 0.06 | 0.21 | 0.14 | 3.1E+07 | 2.7E+07 | 2.4E+06 | 6.0E+07 |
| AT2G36880 | AT2G36880.2;AT2G36880.1 | methionine adenosyltransferase 3 | 42 kDa | 7 | 19 | 72 | 52 | 150 | 0.12 | 0.59 | 0.35 | 4.6E+07 | 4.2E+07 | 3.7E+06 | 9.2E+07 |
| AT5G05780 | AT5G05780.1;AT5G05780.2 | RP non-ATPase subunit 8A | 35 kDa | 10 | 27 | 72 | 43 | 152 | 0.13 | 2.14 | 1.13 | 1.7E+08 | 1.8E+08 | 2.9E+07 | 3.8E+08 |
| AT3G53430 | AT3G53430.1 | Ribosomal protein L11 family protein | 18 kDa | 13 | 8 | 72 | 42 | 135 | | 0.36 | 0.36 | 1.6E+07 | 2.3E+07 | 5.8E+06 | 4.5E+07 |
| AT3G09900 | | RAB GTPase homolog E1E | 24 kDa | 31 | 55 | 72 | 40 | 198 | | | | | | | |
| AT5G16130 | AT5G16130.1 | Ribosomal protein S7e family protein | 22 kDa | 10 | 13 | 72 | 39 | 134 | 0.14 | 0.36 | 0.25 | 1.8E+08 | 2.1E+08 | 2.1E+07 | 4.1E+08 |
| AT1G42440 | AT1G42440.1 | FUNCTIONS IN: molecular_function unknown; INVOLVE | 89 kDa | 0 | 0 | 72 | 30 | 102 | | 0.42 | 0.42 | 1.5E+07 | 1.9E+07 | 8.2E+06 | 4.2E+07 |
| AT1G79550 | AT1G79550.2;AT1G79550.1 | phosphoglycerate kinase | 42 kDa | 12 | 17 | 71 | 106 | 206 | 2.06 | 7.85 | 4.96 | 2.4E+07 | 2.5E+07 | 6.0E+04 | 4.9E+07 |
| AT3G44110 | AT3G44110.1;AT3G44110.2 | DNAJ homologue 3 | 46 kDa | 20 | 28 | 71 | 65 | 184 | 0.11 | | 0.11 | 1.0E+08 | 9.8E+07 | 1.6E+07 | 2.2E+08 |
| AT5G11890 | AT5G11890.1 | FUNCTIONS IN: molecular_function unknown; LOCATE | 31 kDa | 4 | 8 | 71 | 41 | 124 | | 15.78 | 15.78 | 3.1E+07 | 3.1E+07 | 9.8E+05 | 6.3E+07 |
| AT3G05710 | AT3G05710.1;AT3G05710.2 | syntaxin of plants 43 | 37 kDa | 5 | 6 | 70 | 127 | 208 | | 8.72 | 8.72 | 0.0E+00 | 9.5E+05 | 5.3E+04 | 1.0E+06 |
| AT4G29900 | AT4G29900.2;AT4G29900.1 | autoinhibited Ca(2+)-ATPase 10 | 117 kDa | 0 | 0 | 70 | 67 | 137 | | | | 2.2E+06 | 1.5E+06 | 0.0E+00 | 3.7E+06 |
| AT5G64420 | AT5G64420.1 | DNA polymerase V family | 146 kDa | 0 | 5 | 70 | 64 | 139 | 0.12 | | 0.12 | 9.1E+06 | 7.8E+06 | 2.6E+06 | 1.9E+07 |
| AT2G47800 | AT2G47800.1 | multidrug resistance-associated protein 4 | 169 kDa | 3 | 0 | 70 | 61 | 134 | 0.34 | 0.20 | 0.27 | 4.8E+06 | 2.0E+06 | 1.5E+05 | 7.0E+06 |
| AT2G25270 | AT2G25270.1 | unknown protein; LOCATED IN: plasma membrane; EXP | ? | 9 | 1 | 70 | 60 | 140 | | 8.32 | 8.32 | 1.5E+07 | 9.4E+06 | 9.8E+05 | 2.5E+07 |
| AT1G11330 | AT1G11330.2;AT1G11330.1;AT1G11330.3 | S-locus lectin protein kinase family protein | 94 kDa | 17 | 11 | 70 | 54 | 152 | | | | 1.6E+06 | 9.9E+05 | 0.0E+00 | 2.6E+06 |
| AT4G22690 | AT4G22690.1;AT4G22710.1 | cytochrome P450, family 706, subfamily A, polypeptid | 63 kDa | 0 | 0 | 70 | 51 | 121 | 0.19 | 0.47 | 0.33 | 8.9E+05 | 7.4E+05 | 2.1E+05 | 1.8E+06 |
| AT4G38630 | AT4G38630.1 | regulatory particle non-ATPase 10 | 41 kDa | 8 | 14 | 70 | 51 | 143 | | 1.98 | 1.98 | 2.8E+08 | 3.7E+08 | 3.1E+07 | 6.8E+08 |
| AT4G16720 | AT4G16720.1;AT4G17390.1 | Ribosomal protein L23/L15e family protein | 24 kDa | 15 | 26 | 70 | 50 | 161 | 0.07 | 0.32 | 0.19 | 8.8E+08 | 9.8E+08 | 1.8E+08 | 2.0E+09 |
| AT5G20010 | AT5G20010.1;AT5G55190.1;AT5G20020.1 | RAS-related nuclear protein-1 | 25 kDa | 14 | 16 | 70 | 49 | 149 | 0.14 | 0.31 | 0.23 | 5.0E+08 | 5.0E+08 | 1.7E+07 | 1.0E+09 |
| AT2G19790 | AT2G19790.1 | SNARE-like superfamily protein | 17 kDa | 22 | 40 | 70 | 45 | 177 | | | | 5.6E+07 | 8.7E+07 | 2.6E+06 | 1.5E+08 |
| AT5G06970 | AT5G06970.1 | Protein of unknown function (DUF810) | 125 kDa | 21 | 6 | 70 | 44 | 141 | | | | 6.3E+05 | 1.4E+06 | 0.0E+00 | 2.1E+06 |
| AT4G04700 | | calcium-dependent protein kinase 27 | 55 kDa | 3 | 3 | 70 | 36 | 112 | | 3.20 | 3.20 | | | | |
| AT5G48760 | AT5G48760.2;AT5G48760.1 | Ribosomal protein L13 family protein | 24 kDa | 4 | 16 | 70 | 27 | 117 | | 0.22 | 0.22 | 2.6E+07 | 2.9E+07 | 5.6E+06 | 6.1E+07 |
| AT4G35100 | AT4G35100.2;AT4G35100.1 | plasma membrane intrinsic protein 3 | 30 kDa | 9 | 0 | 69 | 72 | 150 | | 3.30 | 3.30 | 1.6E+07 | 1.0E+07 | 6.1E+06 | 3.2E+07 |
| AT5G61730 | AT5G61730.1;AT5G61730.2 | ABC2 homolog 11 | 104 kDa | 14 | 0 | 69 | 66 | 149 | | | | 7.8E+05 | 7.3E+05 | 9.8E+04 | 1.6E+06 |
| AT5G56590 | AT5G56590.1 | O-Glycosyl hydrolases family 17 protein | 56 kDa | 2 | 1 | 69 | 55 | 127 | | 3.36 | 3.36 | 3.8E+06 | 3.3E+06 | 0.0E+00 | 7.1E+06 |
| AT3G02200 | AT3G02200.2;AT3G02200.1 | Proteasome component (PCI) domain protein | 47 kDa | 11 | 45 | 69 | 52 | 177 | 0.11 | 0.84 | 0.48 | 1.4E+07 | 1.4E+07 | 5.4E+06 | 3.3E+07 |
| AT5G28740 | AT5G28740.1 | Tetratricopeptide repeat (TPR)-like superfamily protein | 107 kDa | 0 | 0 | 69 | 50 | 119 | | 0.56 | 0.56 | 8.2E+06 | 1.1E+07 | 4.6E+05 | 2.0E+07 |
| AT2G32450 | | Calcium-binding tetratricopeptide family protein | 90 kDa | 0 | 0 | 69 | 46 | 115 | | | | | | | |
| AT5G03700 | AT5G03700.1 | D-mannose binding lectin protein with Apple-like carb | 54 kDa | 1 | 7 | 69 | 46 | 123 | | 3.27 | 3.27 | 3.2E+06 | 2.4E+06 | 0.0E+00 | 5.7E+06 |
| AT5G47910 | AT5G47910.1 | respiratory burst oxidase homologue D | 104 kDa | 0 | 0 | 69 | 32 | 101 | | | | 1.1E+06 | 1.2E+06 | 0.0E+00 | 2.3E+06 |
| AT1G59740 | | Major facilitator superfamily protein | 66 kDa | 0 | 0 | 69 | 29 | 98 | | | | | | | |
| AT4G02195 | AT4G02195.1 | syntaxin of plants 42 | 36 kDa | 10 | 8 | 68 | 116 | 202 | | | | 9.3E+05 | 6.1E+05 | 1.4E+05 | 1.7E+06 |
| AT4G04910 | AT4G04910.1 | AAA-type ATPase family protein | 81 kDa | 1 | 1 | 68 | 76 | 146 | | | | 5.0E+06 | 3.1E+06 | 5.8E+04 | 8.2E+06 |
| AT1G48030 | AT1G48030.5;AT1G48030.4;AT1G48030.3;AT1G48030.2;AT1G48030.1 | mitochondrial lipoamide dehydrogenase 1 | 54 kDa | 9 | 2 | 68 | 60 | 139 | | | | 3.7E+07 | 6.7E+07 | 1.1E+07 | 1.1E+08 |
| AT2G37230 | AT2G37230.1 | Tetratricopeptide repeat (TPR)-like superfamily protein | 86 kDa | 1 | 0 | 68 | 50 | 119 | | 1.00 | 1.00 | 2.2E+07 | 3.3E+07 | 6.1E+06 | 6.0E+07 |
| AT5G15080 | AT5G15080.1;AT3G01300.1 | Protein kinase superfamily protein | 55 kDa | 2 | 1 | 68 | 47 | 118 | | | | 4.2E+06 | 2.4E+06 | 2.7E+05 | 6.8E+06 |
| AT4G39200 | AT4G39200.2;AT4G39200.1 | Ribosomal protein S25 family protein | 12 kDa | 4 | 0 | 68 | 19 | 91 | | 0.15 | 0.15 | 2.4E+06 | 5.0E+06 | 7.7E+06 | 1.5E+07 |
| AT2G34660 | AT2G34660.1;AT2G34660.2;AT2G34660.3 | multidrug resistance-associated protein 2 | 182 kDa | 0 | 0 | 67 | 72 | 139 | | 0.28 | 0.28 | 7.6E+05 | 4.2E+05 | 0.0E+00 | 1.2E+06 |
| AT4G12980 | AT4G12980.1 | Auxin-responsive family protein | 42 kDa | 0 | 0 | 67 | 69 | 136 | | | | 5.0E+07 | 4.1E+07 | 5.2E+05 | 9.1E+07 |
| AT4G31840 | AT4G31840.1 | early nodulin-like protein 15 | 19 kDa | 0 | 0 | 67 | 68 | 135 | 1.14 | | 1.14 | 2.6E+08 | 2.2E+08 | 1.3E+07 | 4.9E+08 |
| AT3G63460 | AT3G63460.1;AT3G63460.3;AT3G63460.2 | transducin family protein / WD-40 repeat family protei | 120 kDa | 15 | 11 | 67 | 45 | 138 | | 0.63 | 0.63 | 1.9E+06 | 2.7E+06 | 0.0E+00 | 4.6E+06 |
| AT5G47010 | AT5G47010.1 | RNA helicase, putative | 137 kDa | 1 | 0 | 67 | 39 | 107 | | | | 3.1E+06 | 4.6E+06 | 7.0E+05 | 8.4E+06 |
| AT3G52640 | AT3G52640.3;AT3G52640.1;AT3G52640.2 | Zn-dependent exopeptidases superfamily protein | 73 kDa | 2 | 1 | 67 | 38 | 108 | | | | 5.8E+06 | 2.7E+06 | 0.0E+00 | 8.5E+06 |
| AT2G43460 | | Ribosomal L38e protein family | 8 kDa | 7 | 0 | 67 | 25 | 99 | 0.54 | 0.16 | 0.35 | | | | |
| AT2G37050 | AT2G37050.3;AT2G37050.1;AT2G37050.4 | Leucine-rich repeat protein kinase family protein | 104 kDa | 0 | 0 | 66 | 56 | 122 | | | | 4.6E+06 | 3.7E+06 | 0.0E+00 | 8.3E+06 |
| AT4G20360 | AT4G20360.2;AT4G20360.1 | RAB GTPase homolog E1B | 52 kDa | 8 | 18 | 66 | 52 | 144 | 0.15 | 0.40 | 0.27 | 2.1E+07 | 1.5E+07 | 1.1E+07 | 4.7E+07 |
| AT3G18740 | AT3G18740.1 | Ribosomal protein L7Ae/L30e/S12e/Gadd45 family prc | 12 kDa | 14 | 15 | 66 | 46 | 141 | | 0.17 | 0.17 | 6.9E+07 | 7.1E+07 | 5.9E+06 | 1.4E+08 |
| AT4G25630 | AT4G25630.1 | fibrillarin 2 | 34 kDa | 3 | 11 | 66 | 31 | 111 | 0.07 | | 0.07 | 7.3E+06 | 6.6E+06 | 5.8E+06 | 2.0E+07 |
| AT5G19550 | AT5G19550.1 | aspartate aminotransferase 2 | 44 kDa | 23 | 15 | 65 | 80 | 183 | 0.36 | 1.86 | 1.11 | 3.6E+07 | 4.0E+07 | 0.0E+00 | 7.7E+07 |
| AT3G62700 | AT3G62700.1 | multidrug resistance-associated protein 10 | 172 kDa | 3 | 0 | 65 | 72 | 140 | | 0.20 | 0.20 | 1.7E+06 | 6.1E+05 | 0.0E+00 | 2.3E+06 |
| AT3G59110 | AT3G59110.1 | Protein kinase superfamily protein | 57 kDa | 2 | 1 | 65 | 61 | 129 | | | | 5.7E+06 | 4.5E+06 | 0.0E+00 | 1.0E+07 |
| AT3G53610 | AT3G53610.3;AT3G53610.2;AT3G53610.1 | RAB GTPase homolog 8 | 24 kDa | 39 | 58 | 65 | 49 | 211 | | | | 1.1E+06 | 1.4E+06 | 5.4E+04 | 2.5E+06 |
| AT5G22060 | AT5G22060.1 | DNAJ homologue 2 | 46 kDa | 15 | 27 | 65 | 48 | 155 | 0.10 | 0.38 | 0.24 | 8.6E+07 | 8.6E+07 | 1.8E+07 | 1.9E+08 |
| AT5G61170 | AT5G61170.1 | Ribosomal protein S19e family protein | 16 kDa | 15 | 0 | 65 | 22 | 102 | | 0.17 | 0.17 | 2.0E+07 | 1.5E+07 | 1.4E+07 | 4.9E+07 |
| AT3G54200 | AT3G54200.1 | Late embryogenesis abundant (LEA) hydroxyproline-ric | 26 kDa | 11 | 17 | 64 | 70 | 162 | 3.04 | 13.08 | 8.06 | 2.4E+07 | 1.7E+07 | 9.5E+05 | 4.2E+07 |
| AT3G52930 | AT3G52930.1 | Aldolase superfamily protein | 39 kDa | 9 | 18 | 64 | 55 | 146 | | | | 1.5E+07 | 6.9E+06 | 0.0E+00 | 2.2E+07 |
| AT3G47780 | AT3G47780.1;AT3G47740.1 | ABC2 homolog 6 | 105 kDa | 21 | 1 | 64 | 53 | 139 | | | | 7.6E+05 | 7.9E+05 | 0.0E+00 | 1.6E+06 |
| AT3G57530 | AT3G57530.2;AT3G57530.1 | calcium-dependent protein kinase 32 | 61 kDa | 2 | 4 | 64 | 53 | 123 | | | | 2.7E+06 | 2.0E+06 | 0.0E+00 | 4.6E+06 |
| AT3G51800 | AT3G51800.3;AT3G51800.1;AT3G51800.2 | metallopeptidase M24 family protein | 43 kDa | 2 | 12 | 64 | 50 | 128 | | 0.50 | 0.50 | 3.9E+08 | 4.1E+08 | 1.4E+08 | 9.3E+08 |
| AT3G42660 | AT3G42660.1 | transducin family protein / WD-40 repeat family protei | 105 kDa | 0 | 0 | 64 | 40 | 104 | | 0.34 | 0.34 | 1.4E+06 | 6.2E+05 | 8.2E+05 | 2.9E+06 |
| AT4G26690 | AT4G26690.1 | PLC-like phosphodiesterase family protein | 83 kDa | 4 | 10 | 63 | 81 | 158 | | 3.23 | 3.23 | 4.2E+07 | 2.7E+07 | 4.0E+05 | 7.0E+07 |
| AT3G17410 | AT3G17410.2;AT3G17410.1 | Protein kinase superfamily protein | 40 kDa | 8 | 26 | 63 | 51 | 148 | | 8.03 | 8.03 | 8.0E+05 | 6.0E+05 | 0.0E+00 | 1.4E+06 |
| AT3G29310 | | calmodulin-binding protein-related | 61 kDa | 1 | 1 | 63 | 47 | 112 | | | | | | | |
| AT1G04170 | AT1G04170.1;AT1G04170.2 | eukaryotic translation initiation factor 2 gamma subun | 51 kDa | 2 | 2 | 63 | 43 | 110 | | 0.19 | 0.19 | 6.5E+07 | 6.0E+07 | 1.9E+07 | 1.4E+08 |

| | | | | | | | | | | | | | | | |
|---|---|---|---|---|---|---|---|---|---|---|---|---|---|---|---|
| AT5G60790 | AT5G60790.1 | ABC transporter family protein | 67 kDa | 0 | 0 | 63 | 34 | 97 | | 0.81 | 0.81 | 1.2E+07 | 1.0E+07 | 1.2E+07 | 3.5E+07 |
| AT3G11270 | AT3G11270.1;AT3G11270.2 | Mov34/MPN/PAD-1 family protein | 35 kDa | 9 | 21 | 63 | 31 | 124 | 0.20 | 1.96 | 1.08 | 4.7E+07 | 5.1E+07 | 8.4E+06 | 1.1E+08 |
| AT5G52470 | AT5G52470.1;AT5G52470.2 | fibrillarin 1 | 33 kDa | 0 | 8 | 63 | 28 | 99 | | 0.20 | 0.20 | 3.5E+06 | 2.9E+06 | 7.0E+05 | 7.1E+06 |
| AT3G28500 | AT3G28500.1 | 60S acidic ribosomal protein family | 12 kDa | 6 | 0 | 63 | 11 | 80 | 0.38 | 0.16 | 0.27 | 2.6E+08 | 3.0E+08 | 1.9E+08 | 7.5E+08 |
| AT5G43100 | AT5G43100.2;AT5G43100.1 | Eukaryotic aspartyl protease family protein | 70 kDa | 5 | 6 | 62 | 53 | 126 | | 3.31 | 3.31 | 9.2E+05 | 2.5E+06 | 0.0E+00 | 3.4E+06 |
| AT1G66150 | AT1G66150.1 | transmembrane kinase 1 | 102 kDa | 2 | 2 | 62 | 50 | 116 | | | | 5.3E+06 | 2.8E+06 | 0.0E+00 | 8.1E+06 |
| AT4G21710 | AT4G21710.1 | DNA-directed RNA polymerase family protein | 135 kDa | 1 | 0 | 62 | 44 | 107 | | 0.19 | 0.19 | 1.4E+07 | 1.1E+07 | 3.3E+06 | 2.8E+07 |
| AT2G20010 | AT2G20010.2;AT2G20010.1 | Protein of unknown function (DUF810) | 106 kDa | 61 | 28 | 62 | 37 | 188 | | 3.78 | 3.78 | 6.3E+06 | 7.9E+06 | 0.0E+00 | 1.4E+07 |
| AT5G23900 | AT5G23900.1 | Ribosomal protein L13e family protein | 23 kDa | 0 | 6 | 62 | 32 | 100 | 0.25 | 0.49 | 0.37 | 1.6E+07 | 1.9E+07 | 2.3E+07 | 5.9E+07 |
| AT1G77940 | AT1G77940.1 | Ribosomal protein L7Ae/L30e/S12e/Gadd45 family prc | 12 kDa | 10 | 11 | 62 | 29 | 112 | | 0.13 | 0.13 | 1.3E+08 | 1.7E+08 | 3.6E+07 | 3.4E+08 |
| AT1G48210 | AT1G48210.7;AT1G48210.3;AT1G48210.2;AT1G48210.1;AT1G48210.6;AT1G48210.5;AT1G48210.4 | Protein kinase superfamily protein | 40 kDa | 9 | 28 | 61 | 43 | 141 | | 9.67 | 9.67 | 1.9E+06 | 1.1E+06 | 0.0E+00 | 2.9E+06 |
| AT2G07690 | AT2G07690.1;AT2G07690.2 | Minichromosome maintenance (MCM2/3/5) family prc | 81 kDa | 0 | 0 | 61 | 42 | 103 | | 0.43 | 0.43 | 4.8E+06 | 5.8E+06 | 3.0E+05 | 1.1E+07 |
| AT3G18130 | AT3G18130.1 | receptor for activated C kinase 1C | 36 kDa | 8 | 16 | 61 | 41 | 126 | 0.16 | | 0.16 | 2.2E+07 | 1.9E+07 | 3.3E+06 | 4.5E+07 |
| AT3G55200 | | Cleavage and polyadenylation specificity factor (CPSF) / | 135 kDa | 2 | 0 | 61 | 40 | 103 | 0.17 | 0.15 | 0.16 | | | | |
| AT3G24660 | AT3G24660.1 | transmembrane kinase-like 1 | 73 kDa | 0 | 0 | 61 | 38 | 99 | | 3.75 | 3.75 | 8.8E+06 | 7.6E+06 | 1.2E+05 | 1.7E+07 |
| AT3G57150 | AT3G57150.1 | homologue of NAP57 | 63 kDa | 1 | 0 | 61 | 37 | 99 | | 0.15 | 0.15 | 2.5E+07 | 3.6E+07 | 9.5E+06 | 7.0E+07 |
| AT1G49670 | AT1G49670.1;AT1G49670.2 | ARP protein (REF) | 68 kDa | 0 | 0 | 61 | 31 | 92 | | 0.42 | 0.42 | 4.3E+07 | 3.7E+07 | 1.4E+07 | 9.4E+07 |
| AT1G64550 | AT1G64550.1 | general control non-repressible 3 | 80 kDa | 0 | 1 | 61 | 28 | 90 | 0.15 | 0.26 | 0.21 | 1.8E+07 | 2.1E+07 | 4.0E+06 | 4.2E+07 |
| AT1G33590 | AT1G33590.1;AT1G33590.3;AT1G33590.2 | Leucine-rich repeat (LRR) family protein | 52 kDa | 0 | 3 | 60 | 98 | 161 | 0.08 | 0.24 | 0.16 | 6.4E+07 | 3.7E+07 | 7.0E+06 | 1.1E+08 |
| AT1G76040 | | calcium-dependent protein kinase 29 | 64 kDa | 0 | 0 | 60 | 77 | 137 | | | | | | | |
| AT5G03340 | | ATPase, AAA-type, CDC48 protein | 90 kDa | 62 | 28 | 60 | 64 | 214 | | | | | | | |
| AT1G10490 | AT1G10490.2;AT1G10490.1 | Domain of unknown function (DUF1726) ;Putative ATP | 116 kDa | 0 | 1 | 60 | 35 | 96 | | 0.23 | 0.23 | 2.0E+06 | 1.6E+06 | 2.1E+06 | 5.8E+06 |
| AT3G48930 | AT3G48930.1 | Nucleic acid-binding, OB-fold-like protein | 18 kDa | 23 | 10 | 60 | 35 | 128 | 0.21 | 0.33 | 0.27 | 3.5E+08 | 3.9E+08 | 5.9E+07 | 8.0E+08 |
| AT5G41520 | AT5G41520.1;AT5G41520.2 | RNA binding Plectin/S10 domain-containing protein | 20 kDa | 22 | 0 | 59 | 41 | 122 | 0.35 | 0.29 | 0.32 | 2.0E+08 | 1.8E+08 | 3.2E+07 | 4.1E+08 |
| AT1G21630 | AT1G21630.3;AT1G21630.1;AT1G21630.4;AT1G21630.2 | Calcium-binding EF hand family protein | 132 kDa | 0 | 2 | 58 | 60 | 120 | | | | 2.7E+06 | 3.0E+06 | 9.3E+05 | 6.6E+06 |
| AT2G33340 | AT2G33340.1;AT2G33340.2;AT2G33340.3 | MOS4-associated complex 3B | 57 kDa | 0 | 0 | 58 | 51 | 109 | | 0.28 | 0.28 | 2.0E+07 | 2.3E+07 | 2.6E+06 | 4.6E+07 |
| AT2G17120 | AT2G17120.1 | lysm domain GPI-anchored protein 2 precursor | 38 kDa | 11 | 14 | 58 | 46 | 129 | | | | 2.2E+07 | 2.0E+07 | 5.1E+05 | 4.2E+07 |
| AT2G16440 | AT2G16440.1 | Minichromosome maintenance (MCM2/3/5) family prc | 94 kDa | 0 | 0 | 58 | 16 | 74 | | 0.71 | 0.71 | 1.4E+07 | 1.2E+07 | 3.1E+06 | 2.9E+07 |
| AT5G01320 | | Thiamine pyrophosphate dependent pyruvate decarbo | 65 kDa | 0 | 50 | 57 | 227 | 334 | | | | | | | |
| AT1G61670 | AT1G61670.2;AT1G61670.1 | Lung seven transmembrane receptor family protein | 58 kDa | 3 | 2 | 57 | 59 | 121 | | | | 1.9E+06 | 1.2E+06 | 5.6E+04 | 3.2E+06 |
| AT1G42470 | AT1G42470.1 | Patched family protein | 140 kDa | 4 | 0 | 57 | 46 | 107 | | | | 2.1E+07 | 1.6E+07 | 1.5E+05 | 3.7E+07 |
| AT1G09330 | AT1G09330.1;AT1G09330.2 | unknown protein; FUNCTIONS IN: molecular_function | 21 kDa | 9 | 0 | 57 | 44 | 110 | 3.01 | 4.47 | 3.74 | 1.6E+06 | 2.9E+06 | 1.3E+06 | 5.8E+06 |
| AT3G02090 | AT3G02090.1;AT3G02090.2 | Insulinase (Peptidase family M16) protein | 59 kDa | 7 | 1 | 57 | 39 | 104 | 0.06 | 0.18 | 0.12 | 1.4E+07 | 2.3E+07 | 4.7E+05 | 3.8E+07 |
| AT4G02230 | AT4G02230.1 | Ribosomal protein L19e family protein | 24 kDa | 6 | 25 | 57 | 31 | 119 | | | | 7.5E+07 | 8.6E+07 | 9.2E+06 | 1.7E+08 |
| AT3G20290 | AT3G20290.3;AT3G20290.2;AT3G20290.1 | EPS15 homology domain 1 | 61 kDa | 92 | 41 | 57 | 27 | 217 | | | | 2.2E+06 | 2.5E+06 | 0.0E+00 | 4.7E+06 |
| AT5G57290 | AT5G57290.3;AT5G57290.1;AT5G57290.2 | 60S acidic ribosomal protein family | 12 kDa | 4 | 0 | 57 | 11 | 72 | 0.21 | 0.17 | 0.19 | 2.4E+08 | 2.0E+08 | 4.6E+07 | 4.9E+08 |
| AT5G62680 | | Major facilitator superfamily protein | 68 kDa | 1 | 4 | 56 | 61 | 122 | | | | | | | |
| AT1G61250 | AT1G61250.2;AT1G61250.1 | secretory carrier 3 | 33 kDa | 7 | 23 | 56 | 53 | 139 | | 4.00 | 4.00 | 3.9E+07 | 2.1E+07 | 0.0E+00 | 6.1E+07 |
| AT3G46830 | AT3G46830.2;AT3G46830.1 | RAB GTPase homolog A2C | 24 kDa | 26 | 33 | 56 | 45 | 160 | 0.15 | | 0.15 | 5.1E+06 | 5.5E+06 | 0.0E+00 | 1.1E+07 |
| AT1G63660 | AT1G63660.1;AT1G63660.2 | GMP synthase (glutamine-hydrolyzing), putative / gluta | 59 kDa | 3 | 7 | 56 | 39 | 105 | | 0.28 | 0.28 | 3.9E+07 | 3.4E+07 | 2.3E+06 | 7.5E+07 |
| AT3G51740 | AT3G51740.1 | inflorescence meristem receptor-like kinase 2 | 90 kDa | 8 | 1 | 55 | 69 | 133 | | | | 4.4E+06 | 7.2E+06 | 0.0E+00 | 1.2E+07 |
| AT5G56010 | | heat shock protein 81-3 | 80 kDa | 0 | 0 | 55 | 60 | 115 | | | | | | | |
| AT2G26300 | AT2G26300.2;AT2G26300.1 | G protein alpha subunit 1 | 45 kDa | 2 | 4 | 55 | 47 | 108 | | 2.12 | 2.12 | 3.5E+06 | 1.9E+06 | 0.0E+00 | 5.4E+06 |
| AT3G48870 | AT3G48870.2;AT3G48870.4;AT3G48870.3;AT3G48870.1 | Clp ATPase | 106 kDa | 0 | 1 | 55 | 42 | 98 | 0.08 | 0.31 | 0.19 | 1.9E+05 | 0.0E+00 | 0.0E+00 | 1.9E+05 |
| AT1G48630 | AT1G48630.1 | receptor for activated C kinase 1B | 36 kDa | 0 | 17 | 55 | 37 | 109 | | 0.21 | 0.21 | 1.0E+07 | 1.3E+07 | 5.0E+06 | 2.8E+07 |
| AT2G27100 | AT2G27100.1 | C2H2 zinc-finger protein SERRATE (SE) | 81 kDa | 0 | 1 | 55 | 35 | 91 | 0.29 | 0.51 | 0.40 | 3.5E+06 | 5.1E+06 | 8.4E+05 | 9.5E+06 |
| AT1G01100 | AT1G01100.4;AT1G01100.2;AT1G01100.1;AT1G01100.3 | 60S acidic ribosomal protein family | 11 kDa | 8 | 0 | 55 | 10 | 73 | 0.28 | 0.12 | 0.20 | 2.5E+08 | 2.7E+08 | 1.9E+08 | 7.1E+08 |
| AT3G16240 | | delta tonoplast integral protein | 25 kDa | 4 | 9 | 54 | 166 | 233 | 0.22 | 0.65 | 0.44 | | | | |
| AT1G03230 | AT1G03230.1 | Eukaryotic aspartyl protease family protein | 46 kDa | 1 | 1 | 54 | 82 | 138 | 0.02 | 0.24 | 0.13 | 4.4E+07 | 3.7E+07 | 4.2E+06 | 8.5E+07 |
| AT5G56030 | AT5G56030.1;AT5G56010.1;AT5G56030.2;AT5G56000.1 | heat shock protein 81-2 | 80 kDa | 12 | 0 | 54 | 60 | 126 | | 1.27 | 1.27 | 1.0E+07 | 1.1E+07 | 0.0E+00 | 2.2E+07 |
| AT1G29940 | AT1G29940.1 | nuclear RNA polymerase A2 | 132 kDa | 0 | 0 | 54 | 57 | 111 | | 0.68 | 0.68 | 3.4E+06 | 3.5E+06 | 3.2E+06 | 1.0E+07 |
| AT1G56140 | AT1G56140.1 | Leucine-rich repeat transmembrane protein kinase | 114 kDa | 0 | 1 | 54 | 50 | 105 | | | | 1.2E+06 | 9.7E+05 | 0.0E+00 | 2.1E+06 |
| AT4G08350 | AT4G08350.1 | global transcription factor group A2 | 115 kDa | 0 | 0 | 54 | 47 | 101 | | | | 8.1E+05 | 8.7E+05 | 1.1E+05 | 1.8E+06 |
| AT3G27300 | AT3G27300.3;AT3G27300.2;AT3G27300.1;AT3G27300.5;AT3G27300.4 | glucose-6-phosphate dehydrogenase 5 | 59 kDa | 0 | 0 | 54 | 32 | 86 | 0.09 | 0.34 | 0.22 | 3.8E+07 | 3.2E+07 | 1.1E+06 | 7.1E+07 |
| AT5G13110 | AT5G13110.1 | glucose-6-phosphate dehydrogenase 2 | 67 kDa | 0 | 6 | 54 | 31 | 91 | | | | 3.9E+06 | 2.3E+06 | 2.2E+05 | 6.4E+06 |
| AT5G02450 | AT5G02450.1 | Ribosomal protein L36e family protein | 12 kDa | 8 | 0 | 54 | 29 | 91 | 0.20 | 0.57 | 0.38 | 2.2E+08 | 2.4E+08 | 4.9E+07 | 5.1E+08 |
| AT2G17760 | AT2G17760.1 | Eukaryotic aspartyl protease family protein | 56 kDa | 7 | 4 | 54 | 27 | 92 | | | | 9.7E+06 | 6.8E+06 | 0.0E+00 | 1.7E+07 |
| AT1G18080 | AT1G18080.1 | Transducin/WD40 repeat-like superfamily protein | 36 kDa | 0 | 2 | 54 | 5 | 61 | | 0.12 | 0.12 | 8.3E+07 | 1.1E+08 | 3.8E+07 | 2.3E+08 |
| AT5G42100 | AT5G42100.2;AT5G42100.1 | beta-1,3-glucanase_putative | 45 kDa | 11 | 11 | 53 | 70 | 145 | 3.48 | 4.93 | 4.21 | 7.0E+07 | 5.6E+07 | 2.4E+07 | 1.5E+08 |
| AT4G36080 | | phosphotransferases, alcohol group as acceptor;bindin | 434 kDa | 0 | 0 | 53 | 56 | 109 | | | | | | | |
| AT5G40870 | AT5G40870.1 | uridine kinase/uracil phosphoribosyltransferase 1 | 54 kDa | 0 | 0 | 53 | 49 | 102 | | 0.71 | 0.71 | 4.7E+06 | 4.2E+06 | 1.3E+06 | 1.0E+07 |
| ATCG00500 | ATCG00500.1 | acetyl-CoA carboxylase carboxyl transferase subunit be | 56 kDa | 0 | 2 | 53 | 47 | 102 | | 0.56 | 0.56 | 2.1E+07 | 1.5E+07 | 5.3E+06 | 4.2E+07 |
| AT5G11150 | AT5G11150.2;AT5G11150.1 | vesicle-associated membrane protein 713 | 25 kDa | 2 | 0 | 53 | 46 | 101 | 0.23 | 0.18 | 0.21 | 1.6E+07 | 5.8E+06 | 2.0E+06 | 2.4E+07 |
| AT2G36850 | | glucan synthase-like 8 | 218 kDa | 0 | 0 | 53 | 43 | 96 | 0.48 | 0.63 | 0.56 | | | | |
| AT2G46280 | AT2G46280.2;AT2G46280.1;AT2G46280.3 | TGF-beta receptor interacting protein 1 | 36 kDa | 7 | 1 | 53 | 43 | 104 | 0.16 | 0.44 | 0.30 | 7.0E+07 | 8.8E+07 | 2.4E+07 | 1.8E+08 |
| AT1G25570 | | Di-glucose binding protein with Leucine-rich repeat do | 69 kDa | 9 | 3 | 53 | 37 | 102 | | | | | | | |
| AT1G31130 | AT1G31130.1 | unknown protein; FUNCTIONS IN: molecular_function | ? | 20 | 12 | 53 | 24 | 109 | 2.88 | 1.43 | 2.16 | | | | |
| AT5G18620 | AT5G18620.1;AT5G18620.2 | chromatin remodeling factor17 | 124 kDa | 0 | 0 | 53 | 21 | 74 | | 0.36 | 0.36 | 8.3E+05 | 8.1E+05 | 0.0E+00 | 1.6E+06 |
| AT3G06400 | | chromatin-remodeling protein 11 | 122 kDa | 0 | 0 | 53 | 18 | 71 | | | | | | | |
| AT2G21580 | AT2G21580.2;AT2G21580.1 | Ribosomal protein S25 family protein | 12 kDa | 3 | 0 | 53 | 16 | 72 | | 0.50 | 0.50 | 5.5E+07 | 3.8E+07 | 2.5E+06 | 9.5E+07 |
| AT3G53230 | | ATPase, AAA-type, CDC48 protein | 90 kDa | 57 | 26 | 52 | 57 | 192 | | | | | | | |
| AT3G11820 | AT3G11820.2;AT3G11820.1 | syntaxin of plants 121 | 38 kDa | 20 | 24 | 52 | 55 | 151 | | 6.86 | 6.86 | 2.4E+06 | 2.2E+06 | 8.1E+04 | 4.8E+06 |
| AT2G36380 | AT2G36380.1;AT1G66950.1 | pleiotropic drug resistance 6 | 164 kDa | 4 | 1 | 52 | 45 | 102 | | | | 1.8E+06 | 1.6E+06 | 0.0E+00 | 3.4E+06 |
| AT1G64880 | AT1G64880.1 | Ribosomal protein S5 family protein | 60 kDa | 5 | 0 | 52 | 44 | 101 | | 3.46 | 3.46 | 3.7E+07 | 5.8E+07 | 3.9E+06 | 9.8E+07 |
| AT3G57660 | AT3G57660.2;AT3G57660.1 | nuclear RNA polymerase A1 | 188 kDa | 0 | 0 | 52 | 40 | 92 | | 0.16 | 0.16 | 1.8E+06 | 1.9E+06 | 1.5E+06 | 5.2E+06 |
| AT2G36160 | AT2G36160.1 | Ribosomal protein S11 family protein | 16 kDa | 16 | 1 | 52 | 33 | 102 | 0.07 | 0.14 | 0.11 | 5.3E+07 | 4.8E+07 | 1.5E+07 | 1.2E+08 |
| AT1G70530 | AT1G70530.3;AT1G70530.2;AT1G70530.1 | cysteine-rich RLK (RECEPTOR-like protein kinase) 3 | 72 kDa | 1 | 3 | 52 | 30 | 86 | | | | 1.5E+06 | 1.5E+06 | 0.0E+00 | 3.0E+06 |
| AT2G18840 | | Integral membrane Yip1 family protein | 30 kDa | 0 | 0 | 52 | 30 | 82 | | | | | | | |
| AT3G11510 | AT3G11510.1 | Ribosomal protein S11 family protein | 16 kDa | 16 | 1 | 52 | 30 | 99 | 0.14 | 0.13 | 0.14 | 6.1E+07 | 6.0E+07 | 1.1E+07 | 1.3E+08 |
| AT1G61870 | AT1G61870.1 | pentatricopeptide repeat 336 | 46 kDa | 2 | 9 | 52 | 12 | 75 | | 4.00 | 4.00 | 6.1E+07 | 9.4E+07 | 1.2E+07 | 1.7E+08 |
| AT5G47700 | AT5G47700.2;AT5G47700.1 | 60S acidic ribosomal protein family | 11 kDa | 7 | 0 | 52 | 9 | 68 | 0.25 | 0.12 | 0.18 | 3.3E+08 | 5.1E+08 | 2.2E+08 | 1.1E+09 |

| | | | | | | | | | | | | | | | |
|---|---|---|---|---|---|---|---|---|---|---|---|---|---|---|---|
| AT3G45980 | AT3G45980.1;AT3G46030.1;AT1G07790.1;AT2G28720.1;AT5G02570.1;AT2G37470.1;AT3G09480.1 | Histone superfamily protein | 16 kDa | 0 | 6 | 52 | 0 | 58 | | | | 2.4E+06 | 2.2E+06 | 5.0E+05 | 5.1E+06 |
| AT1G07890 | AT1G07890.6;AT1G07890.8;AT1G07890.7;AT1G07890.5;AT1G07890.4;AT1G07890.3;AT1G07890.2;AT1G07890.1 | ascorbate peroxidase 1 | 28 kDa | 15 | 25 | 51 | 75 | 166 | 0.87 | 9.98 | 5.42 | 3.2E+07 | 1.9E+07 | 2.4E+06 | 5.3E+07 |
| AT5G05160 | AT5G05160.1 | Leucine-rich repeat protein kinase family protein | 71 kDa | 0 | 0 | 51 | 64 | 115 | | | | 2.4E+06 | 1.6E+06 | 0.0E+00 | 3.9E+06 |
| AT3G07160 | AT3G07160.1;AT3G07160.3;AT3G07160.2;AT2G36850.1 | glucan synthase-like 10 | 217 kDa | 1 | 0 | 51 | 55 | 107 | | | | | | | |
| AT3G22520 | | unknown protein; INVOLVED IN: biological_process un | ? | 4 | 2 | 51 | 55 | 112 | | 0.30 | 0.30 | | | | |
| AT3G54140 | AT3G54140.2;AT3G54140.1 | peptide transporter 1 | 64 kDa | 0 | 0 | 51 | 52 | 103 | | | | 7.0E+06 | 4.2E+06 | 4.5E+05 | 1.2E+07 |
| AT1G20760 | AT1G20760.1 | Calcium-binding EF hand family protein | 110 kDa | 1 | 3 | 51 | 49 | 104 | | | | 3.4E+05 | 7.9E+05 | 1.2E+05 | 1.3E+06 |
| AT2G36460 | AT2G36460.1;AT2G36460.3;AT2G36460.2 | Aldolase superfamily protein | 38 kDa | 7 | 17 | 51 | 44 | 119 | | 3.02 | 3.02 | 2.8E+06 | 1.5E+06 | 0.0E+00 | 4.3E+06 |
| AT1G65930 | AT1G65930.1 | cytosolic NADP+-dependent isocitrate dehydrogenase | 46 kDa | 17 | 22 | 51 | 37 | 127 | | | | 3.0E+07 | 1.7E+07 | 0.0E+00 | 4.7E+07 |
| AT1G62020 | AT1G62020.1 | Coatomer, alpha subunit | 137 kDa | 0 | 0 | 51 | 32 | 83 | | 0.29 | 0.29 | 2.9E+06 | 3.9E+06 | 1.4E+05 | 6.9E+06 |
| AT5G24430 | | Calcium-dependent protein kinase (CDPK) family prote | 67 kDa | 0 | 0 | 51 | 31 | 82 | | 3.13 | 3.13 | | | | |
| AT4G10320 | AT4G10320.1 | tRNA synthetase class I (I, L, M and V) family protein | 135 kDa | 0 | 0 | 51 | 29 | 80 | 0.12 | 0.26 | 0.19 | 9.1E+07 | 8.5E+07 | 1.1E+06 | 1.8E+08 |
| AT5G54500 | AT5G54500.1;AT5G54500.2 | flavodoxin-like quinone reductase 1 | 22 kDa | 12 | 26 | 51 | 28 | 117 | 1.93 | 3.09 | 2.51 | 3.5E+07 | 1.4E+07 | 6.7E+05 | 5.0E+07 |
| AT3G02560 | AT3G02560.3;AT3G02560.2;AT3G02560.1 | Ribosomal protein S7e family protein | 22 kDa | 7 | 0 | 51 | 23 | 81 | | | | 6.1E+07 | 6.7E+07 | 2.5E+07 | 1.5E+08 |
| AT2G27720 | AT2G27720.1;AT2G27720.3;AT2G27720.2;AT2G27720.4 | 60S acidic ribosomal protein family | 11 kDa | 10 | 0 | 51 | 13 | 74 | 0.25 | 0.23 | 0.24 | 8.1E+07 | 1.3E+08 | 3.5E+07 | 2.5E+08 |
| AT5G41260 | AT5G41260.1 | Protein kinase protein with tetratricopeptide repeat d | 55 kDa | 7 | 13 | 50 | 60 | 130 | | | | 2.1E+05 | 3.5E+04 | 6.3E+05 | 8.8E+05 |
| AT5G59010 | AT5G59010.2;AT5G59010.1 | Protein kinase protein with tetratricopeptide repeat d | 55 kDa | 0 | 12 | 50 | 54 | 116 | | | | 3.7E+05 | 2.5E+05 | 5.4E+05 | 1.2E+06 |
| AT1G52540 | AT1G52540.2;AT1G52540.1 | Protein kinase superfamily protein | 40 kDa | 1 | 2 | 50 | 49 | 102 | | | | 4.7E+05 | 2.5E+05 | 0.0E+00 | 7.2E+05 |
| AT2G26730 | AT2G26730.1 | Leucine-rich repeat protein kinase family protein | 72 kDa | 11 | 0 | 50 | 42 | 103 | | | | 1.2E+06 | 4.6E+05 | 0.0E+00 | 1.6E+06 |
| AT1G50200 | | Alanyl-tRNA synthetase | 110 kDa | 6 | 0 | 50 | 41 | 97 | | 0.18 | 0.18 | | | | |
| AT1G28340 | AT1G28340.1 | receptor like protein 4 | 68 kDa | 13 | 10 | 50 | 35 | 108 | | 2.14 | 2.14 | 2.9E+05 | 1.6E+05 | 0.0E+00 | 4.5E+05 |
| AT5G65750 | AT5G65750.1 | 2-oxoglutarate dehydrogenase, E1 component | 116 kDa | 0 | 0 | 50 | 27 | 77 | 0.08 | | 0.08 | 4.3E+06 | 1.1E+07 | 8.6E+05 | 1.6E+07 |
| AT3G07050 | AT3G07050.1 | GTP-binding family protein | 66 kDa | 0 | 2 | 50 | 26 | 78 | | 0.23 | 0.23 | 1.5E+07 | 1.6E+07 | 1.0E+07 | 4.1E+07 |
| AT5G22880 | AT5G22880.1 | histone B2 | 16 kDa | 10 | 6 | 50 | 26 | 92 | | | | 9.2E+06 | 6.3E+06 | 7.9E+05 | 1.6E+07 |
| AT1G71830 | | somatic embryogenesis receptor-like kinase 1 | 69 kDa | 1 | 3 | 50 | 19 | 73 | | | | | | | |
| AT2G13820 | AT2G13820.2;AT2G13820.1;AT2G13820.3 | Bifunctional inhibitor/lipid-transfer protein/seed stora | 13 kDa | 3 | 36 | 49 | 80 | 168 | | 4.48 | 4.48 | 5.1E+08 | 4.5E+08 | 1.4E+07 | 9.7E+08 |
| AT2G43030 | AT2G43030.1 | Ribosomal protein L3 family protein | 29 kDa | 12 | 28 | 49 | 50 | 139 | | 4.16 | 4.16 | 4.9E+06 | 4.4E+06 | 8.4E+05 | 1.0E+07 |
| AT4G18910 | AT4G18910.1;AT4G18910.2 | NOD26-like intrinsic protein 1;2 | 31 kDa | 3 | 5 | 49 | 47 | 104 | | | | 4.1E+06 | 3.0E+06 | 2.1E+05 | 7.3E+06 |
| AT3G22330 | AT3G22330.1 | putative mitochondrial RNA helicase 2 | 65 kDa | 2 | 9 | 49 | 45 | 105 | | 0.29 | 0.29 | 1.3E+07 | 1.2E+07 | 3.4E+06 | 2.9E+07 |
| AT1G03870 | AT1G03870.1;AT5G44130.1 | FASCICLIN-like arabinoogalactan 9 | 26 kDa | 1 | 2 | 49 | 44 | 96 | | 6.07 | 6.07 | 3.5E+06 | 1.9E+06 | 4.7E+05 | 5.9E+06 |
| AT5G11240 | AT5G11240.1 | transducin family protein / WD-40 repeat family protei | 67 kDa | 0 | 1 | 49 | 37 | 87 | | 0.31 | 0.31 | 1.3E+07 | 1.4E+07 | 5.8E+06 | 3.2E+07 |
| AT5G10290 | AT5G10290.2;AT5G10290.1 | leucine-rich repeat transmembrane protein kinase fami | 69 kDa | 0 | 0 | 49 | 33 | 82 | | | | 6.1E+05 | 4.0E+05 | 0.0E+00 | 1.0E+06 |
| AT3G61260 | AT3G61260.1 | Remorin family protein | 23 kDa | 11 | 26 | 49 | 32 | 118 | | 6.45 | 6.45 | 1.9E+07 | 9.4E+06 | 3.3E+05 | 2.8E+07 |
| AT3G15730 | | phospholipase D alpha 1 | 92 kDa | 1 | 0 | 49 | 24 | 74 | | | | | | | |
| AT4G33430 | AT4G33430.1;AT4G33430.2;AT2G13800.4;AT2G13800.2;AT2G13800.3;AT2G13800.1;AT2G13790.1;AT1G71830.1 | BRI1-associated receptor kinase | 74 kDa | 1 | 3 | 49 | 21 | 74 | | | | 4.0E+06 | 2.6E+06 | 0.0E+00 | 6.6E+06 |
| AT3G58660 | AT3G58660.1 | Ribosomal protein L1p/L10e family | 50 kDa | 0 | 1 | 49 | 17 | 67 | | | | 4.4E+07 | 3.5E+07 | 1.7E+07 | 9.6E+07 |
| AT3G15380 | | Plasma-membrane choline transporter family protein | 79 kDa | 7 | 3 | 48 | 51 | 109 | | | | | | | |
| AT1G64330 | AT1G64330.1 | myosin heavy chain-related | 64 kDa | 3 | 2 | 48 | 43 | 96 | 0.10 | | 0.10 | 5.4E+05 | 1.6E+05 | 0.0E+00 | 7.0E+05 |
| AT1G11910 | AT1G11910.1;AT1G11910.2 | aspartic proteinase A1 | 55 kDa | 13 | 12 | 48 | 42 | 115 | 0.19 | 0.11 | 0.15 | 5.1E+06 | 8.0E+06 | 4.4E+06 | 1.8E+07 |
| AT3G56190 | AT3G56190.1;AT3G56190.2 | alpha-soluble NSF attachment protein 2 | 33 kDa | 27 | 60 | 48 | 37 | 172 | | 3.11 | 3.11 | 1.2E+07 | 8.6E+06 | 1.8E+06 | 2.2E+07 |
| AT1G24560 | | unknown protein; BEST Arabidopsis thaliana protein m | ? | 9 | 17 | 47 | 72 | 145 | | | | | | | |
| AT5G26742 | AT5G26742.2;AT5G26742.1;AT5G26742.3 | DEAD box RNA helicase (RH3) | 81 kDa | 0 | 4 | 47 | 40 | 91 | 0.09 | 0.29 | 0.19 | 1.8E+07 | 1.5E+07 | 1.7E+06 | 3.5E+07 |
| AT1G24510 | AT1G24510.1;AT1G24510.2;AT1G24510.3 | TCP-1/cpn60 chaperonin family protein | 51 kDa | 1 | 3 | 47 | 38 | 89 | | | | 1.6E+07 | 1.8E+07 | 3.4E+06 | 3.8E+07 |
| AT5G04885 | AT5G04885.3;AT5G04885.1;AT5G04885.2 | Glycosyl hydrolase family protein | 72 kDa | 0 | 0 | 47 | 33 | 80 | | | | 1.5E+06 | 1.8E+06 | 3.1E+05 | 3.6E+06 |
| AT1G22280 | AT1G22280.3;AT1G22280.1 | phytochrome-associated protein phosphatase type 2C | 31 kDa | 9 | 31 | 47 | 32 | 119 | | 14.25 | 14.25 | 7.5E+06 | 4.1E+06 | 0.0E+00 | 1.2E+07 |
| AT4G35300 | AT4G35300.5;AT4G35300.9;AT4G35300.3;AT4G35300.2;AT4G35300.8;AT4G35300.7;AT4G35300.6;AT4G35300.4;AT4G35300.11;AT4G35300.10;AT4G35300.1 | tonoplast monosaccharide transporter2 | 80 kDa | 0 | 0 | 47 | 31 | 78 | | 0.28 | 0.28 | 3.8E+06 | 5.5E+06 | 2.1E+05 | 9.5E+06 |
| AT5G27490 | | Integral membrane Yip1 family protein | 31 kDa | 3 | 1 | 47 | 31 | 82 | | | | | | | |
| AT5G03540 | | exocyst subunit exo70 family protein A1 | 72 kDa | 17 | 2 | 47 | 29 | 95 | | | | | | | |
| AT3G53740 | AT3G53740.1;AT3G53740.4;AT3G53740.3;AT3G53740.2 | Ribosomal protein L36e family protein | 12 kDa | 9 | 0 | 47 | 25 | 81 | | | | 5.6E+07 | 3.5E+07 | 6.9E+06 | 9.8E+07 |
| AT3G53990 | AT3G53990.1;AT3G53990.2 | Adenine nucleotide alpha hydrolases-like superfamily p | 18 kDa | 28 | 0 | 47 | 23 | 98 | 8.89 | 2.07 | 5.48 | 1.4E+08 | 9.8E+07 | 1.6E+06 | 2.4E+08 |
| AT4G25530 | AT4G25530.1 | FLOWERING WAGENINGEN | 76 kDa | 0 | 0 | 47 | 21 | 68 | | 0.51 | 0.51 | 7.2E+05 | 0.0E+00 | 4.3E+05 | 1.2E+06 |
| AT1G19910 | AT1G19910.2;AT1G19910.1;AT4G38920.2;AT4G34720.2 | ATPase, F0/V0 complex, subunit C protein | 17 kDa | 12 | 3 | 47 | 10 | 72 | | | | 1.1E+08 | 2.5E+07 | 3.6E+07 | 1.7E+08 |
| AT2G20850 | AT2G20850.1 | STRUBBELIG-receptor family 1 | 85 kDa | 2 | 4 | 46 | 52 | 104 | | | | 3.1E+06 | 3.7E+06 | 8.5E+04 | 6.9E+06 |
| AT1G09770 | AT1G09770.1 | cell division cycle 5 | 96 kDa | 0 | 0 | 46 | 49 | 95 | 0.14 | 0.30 | 0.22 | 5.6E+06 | 9.7E+06 | 1.0E+06 | 1.6E+07 |
| AT4G20260 | AT4G20260.9;AT4G20260.8;AT4G20260.7;AT4G20260.3;AT4G20260.2;AT4G20260.10;AT4G20260.1;AT4G20260.4;AT4G20260.5 | plasma-membrane associated cation-binding protein 1 | 25 kDa | 2 | 9 | 46 | 45 | 102 | 0.77 | 5.63 | 3.20 | 2.4E+07 | 1.1E+07 | 2.5E+06 | 3.7E+07 |
| AT1G09630 | | RAB GTPase 11C | 24 kDa | 17 | 25 | 46 | 41 | 129 | | | | | | | |
| AT5G19450 | AT5G19450.2;AT5G19450.1 | calcium-dependent protein kinase 19 | 60 kDa | 1 | 7 | 46 | 39 | 93 | | | | 2.4E+06 | 1.8E+06 | 0.0E+00 | 4.2E+06 |
| AT3G46740 | AT3G46740.1 | translocon at the outer envelope membrane of chlorop | 89 kDa | 1 | 0 | 46 | 32 | 79 | 0.07 | 0.24 | 0.15 | 4.0E+06 | 8.0E+06 | 5.5E+05 | 1.3E+07 |
| AT5G27120 | AT5G27120.1 | NOP56-like pre RNA processing ribonucleoprotein | 59 kDa | 0 | 4 | 46 | 32 | 82 | | 0.34 | 0.34 | 5.8E+06 | 3.6E+06 | 4.3E+06 | 1.4E+07 |
| AT2G21390 | AT2G21390.1 | Coatomer, alpha subunit | 136 kDa | 0 | 1 | 46 | 29 | 76 | | 0.19 | 0.19 | 5.5E+05 | 1.2E+06 | 1.1E+05 | 1.8E+06 |
| AT1G72230 | AT1G72230.1 | Cupredoxin superfamily protein | 18 kDa | 10 | 0 | 46 | 15 | 71 | | 7.70 | 7.70 | 4.0E+07 | 4.2E+07 | 0.0E+00 | 8.2E+07 |
| AT2G30970 | AT2G30970.2;AT2G30970.1 | aspartate aminotransferase 1 | 48 kDa | 16 | 39 | 45 | 49 | 149 | 0.09 | 0.52 | 0.30 | 1.3E+07 | 1.8E+07 | 3.2E+05 | 3.1E+07 |
| AT2G19440 | | O-Glycosyl hydrolases family 17 protein | 52 kDa | 2 | 9 | 45 | 47 | 103 | | 5.35 | 5.35 | | | | |
| AT5G62270 | AT5G62270.2;AT5G62270.1 | FUNCTIONS IN: molecular_function unknown; INVOLVI | 48 kDa | 5 | 2 | 45 | 44 | 96 | | 1.29 | 1.29 | 4.0E+07 | 5.9E+07 | 3.3E+06 | 1.0E+08 |
| AT5G48380 | AT5G48380.1 | BAK1-interacting receptor-like kinase 1 | 69 kDa | 1 | 4 | 45 | 38 | 88 | | | | 1.2E+07 | 7.5E+06 | 0.0E+00 | 1.9E+07 |
| AT3G23620 | AT3G23620.1 | Ribosomal RNA processing Brix domain protein | 36 kDa | 2 | 4 | 45 | 36 | 87 | | 1.08 | 1.08 | 5.6E+07 | 5.1E+07 | 1.4E+07 | 1.2E+08 |
| AT4G02350 | | exocyst complex component sec15B | 88 kDa | 15 | 2 | 45 | 35 | 97 | | | | | | | |
| AT3G61240 | AT3G61240.2;AT3G61240.1 | DEA(D/H)-box RNA helicase family protein | 57 kDa | 1 | 0 | 45 | 34 | 80 | | | | 8.2E+06 | 8.0E+06 | 2.1E+06 | 1.8E+07 |
| AT2G26780 | AT2G26780.2;AT2G26780.1 | ARM repeat superfamily protein | 201 kDa | 2 | 0 | 45 | 33 | 80 | | 0.36 | 0.36 | 2.2E+07 | 2.2E+07 | 2.4E+06 | 4.7E+07 |

| | | | | | | | | | | | | | | | |
|---|---|---|---|---|---|---|---|---|---|---|---|---|---|---|---|
| AT3G20050 | AT3G20050.1 | T-complex protein 1 alpha subunit | 59 kDa | 4 | 8 | 45 | 33 | 90 | | 0.19 | 0.19 | 2.5E+06 | 2.7E+06 | 2.2E+05 | 5.5E+06 |
| AT5G12370 | AT5G12370.3;AT5G12370.2;AT5G12370.1 | exocyst complex component sec10 | 90 kDa | 19 | 3 | 45 | 32 | 99 | | | | 2.9E+05 | 5.7E+05 | 0.0E+00 | 8.6E+05 |
| AT1G77630 | AT1G77630.1 | Peptidoglycan-binding LysM domain-containing protei | 44 kDa | 6 | 5 | 45 | 30 | 86 | | | | 8.0E+07 | 6.5E+07 | 1.4E+06 | 1.5E+08 |
| AT1G44910 | AT1G44910.2;AT1G44910.1 | pre-mRNA-processing protein 40A | 109 kDa | 0 | 0 | 45 | 26 | 71 | | 0.66 | 0.66 | 2.4E+07 | 2.4E+07 | 4.3E+06 | 5.2E+07 |
| AT4G37870 | AT4G37870.1 | phosphoenolpyruvate carboxykinase 1 | 73 kDa | 0 | 0 | 45 | 23 | 68 | | 0.36 | 0.36 | 1.3E+07 | 1.8E+07 | 0.0E+00 | 3.1E+07 |
| AT5G02960 | AT5G02960.1 | Ribosomal protein S12/S23 family protein | 16 kDa | 9 | 9 | 45 | 21 | 84 | | | | 4.5E+07 | 6.7E+07 | 1.8E+07 | 1.3E+08 |
| AT3G01390 | AT3G01390.4;AT3G01390.3;AT3G01390.2;AT3G01390.1 | vacuolar membrane ATPase 10 | 12 kDa | 15 | 0 | 45 | 20 | 80 | 1.69 | 1.19 | 1.44 | 3.2E+08 | 2.2E+08 | 2.5E+07 | 5.6E+08 |
| AT3G62220 | AT3G62220.3;AT3G62220.1;AT3G62220.2 | Protein kinase superfamily protein | 39 kDa | 10 | 36 | 44 | 39 | 129 | | 8.95 | 8.95 | 3.1E+06 | 1.7E+06 | 1.1E+06 | 5.8E+06 |
| AT1G43890 | AT1G43890.3;AT1G43890.2;AT1G43890.1 | RAB GTPASE HOMOLOG B18 | 24 kDa | 13 | 18 | 44 | 35 | 110 | 0.15 | 0.66 | 0.41 | 1.1E+07 | 6.8E+06 | 5.5E+05 | 1.9E+07 |
| AT3G55450 | | PBS1-like 1 | 43 kDa | 0 | 3 | 44 | 35 | 82 | | | | | | | |
| AT5G64760 | AT5G64760.3;AT5G64760.2;AT5G64760.1 | regulatory particle non-ATPase subunit 5B | 51 kDa | 0 | 9 | 44 | 35 | 88 | | | | 2.2E+06 | 1.7E+06 | 5.2E+05 | 4.5E+06 |
| AT5G42720 | AT5G42720.1 | Glycosyl hydrolase family 17 protein | 47 kDa | 0 | 1 | 44 | 30 | 75 | | 5.24 | 5.24 | 8.5E+06 | 9.8E+06 | 3.3E+04 | 1.8E+07 |
| AT1G24280 | AT1G24280.1 | glucose-6-phosphate dehydrogenase 3 | 67 kDa | 1 | 5 | 44 | 28 | 78 | | 0.43 | 0.43 | 1.9E+06 | 1.6E+06 | 0.0E+00 | 3.5E+06 |
| AT5G35200 | AT5G35200.2;AT5G35200.1 | ENTH/ANTH/VHS superfamily protein | 61 kDa | 0 | 2 | 44 | 28 | 74 | | | | 2.1E+05 | 6.3E+05 | 0.0E+00 | 8.4E+05 |
| AT3G05560 | AT3G05560.3;AT3G05560.2;AT3G05560.1 | Ribosomal L22e protein family | 14 kDa | 13 | 0 | 44 | 23 | 80 | 0.34 | 0.13 | 0.24 | 7.2E+07 | 6.8E+07 | 1.5E+07 | 1.6E+08 |
| AT3G57940 | AT3G57940.2;AT3G57940.1 | Domain of unknown function (DUF1726) ;Putative ATP | 115 kDa | 0 | 1 | 44 | 23 | 68 | | | | 1.3E+05 | 6.4E+04 | 3.1E+05 | 5.0E+05 |
| AT3G56340 | AT3G56340.1 | Ribosomal protein S26e family protein | 15 kDa | 8 | 0 | 44 | 13 | 65 | | | | 8.2E+07 | 5.1E+07 | 3.8E+07 | 1.7E+08 |
| AT3G10380 | AT3G10380.2;AT3G10380.1 | subunit of exocyst complex 8 | 117 kDa | 10 | 0 | 43 | 41 | 94 | | | | 1.3E+05 | 1.4E+05 | 0.0E+00 | 2.7E+05 |
| AT3G62560 | | Ras-related small GTP-binding family protein | 22 kDa | 0 | 20 | 43 | 37 | 100 | | | | | | | |
| AT4G29360 | AT4G29360.1;AT4G29360.2 | O-Glycosyl hydrolases family 17 protein | 58 kDa | 1 | 0 | 43 | 37 | 81 | 3.41 | 8.07 | 5.74 | 1.4E+06 | 1.2E+06 | 0.0E+00 | 2.6E+06 |
| AT4G02600 | AT4G02600.2;AT4G02600.1 | Seven transmembrane MLO family protein | 59 kDa | 0 | 0 | 43 | 36 | 79 | | | | 2.8E+06 | 1.4E+06 | 0.0E+00 | 4.1E+06 |
| AT2G40660 | AT2G40660.1 | Nucleic acid-binding, OB-fold-like protein | 42 kDa | 4 | 11 | 43 | 35 | 93 | | 1.63 | 1.63 | 1.7E+08 | 1.4E+08 | 4.4E+06 | 3.2E+08 |
| AT5G50920 | | CLPC homologue 1 | ? | 0 | 1 | 43 | 35 | 79 | | | | | | | |
| AT3G27190 | | uridine kinase-like 2 | 54 kDa | 0 | 0 | 43 | 34 | 77 | | | | | | | |
| AT1G55620 | | chloride channel F | 84 kDa | 4 | 2 | 43 | 32 | 81 | | | | | | | |
| AT3G18190 | AT3G18190.1 | TCP-1/cpn60 chaperonin family protein | 58 kDa | 0 | 7 | 43 | 32 | 82 | | 0.47 | 0.47 | 1.0E+07 | 9.3E+06 | 4.6E+06 | 2.4E+07 |
| AT4G10450 | | Ribosomal protein L6 family | 22 kDa | 0 | 5 | 43 | 31 | 79 | | 0.35 | 0.35 | | | | |
| AT5G64270 | AT5G64270.1 | splicing factor, putative | 141 kDa | 0 | 0 | 43 | 30 | 73 | | 0.12 | 0.12 | 4.0E+06 | 5.0E+06 | 2.0E+06 | 1.1E+07 |
| AT5G62890 | AT5G62890.3;AT5G62890.2;AT5G62890.1 | Xanthine/uracil permease family protein | 58 kDa | 16 | 2 | 43 | 28 | 89 | | 1.56 | 1.56 | 1.1E+07 | 3.1E+06 | 0.0E+00 | 1.4E+07 |
| AT1G55890 | AT1G55890.1 | Tetratricopeptide repeat (TPR)-like superfamily protein | 45 kDa | 2 | 7 | 43 | 27 | 79 | | 1.75 | 1.75 | 4.1E+07 | 4.1E+07 | 8.7E+06 | 9.0E+07 |
| AT1G06840 | AT1G06840.1 | Leucine-rich repeat protein kinase family protein | 106 kDa | 0 | 1 | 43 | 26 | 70 | | 1.81 | 1.81 | 8.9E+05 | 3.3E+05 | 2.7E+04 | 1.2E+06 |
| AT5G27770 | AT5G27770.1 | Ribosomal L22e protein family | 14 kDa | 13 | 0 | 43 | 21 | 77 | 0.43 | 0.16 | 0.29 | 1.0E+08 | 1.1E+08 | 5.4E+06 | 2.2E+08 |
| AT1G74270 | AT1G74270.1;AT1G07070.1 | Ribosomal protein L35Ae family protein | 13 kDa | 0 | 0 | 43 | 19 | 62 | | | | 0.0E+00 | 6.5E+06 | 0.0E+00 | 6.5E+06 |
| AT5G57870 | AT5G57870.2;AT5G57870.1 | MIF4G domain-containing protein / MA3 domain-cont | 86 kDa | 0 | 0 | 43 | 19 | 62 | 0.26 | 0.20 | 0.23 | 1.0E+07 | 9.4E+06 | 5.0E+06 | 2.4E+07 |
| AT5G60250 | AT5G60250.1 | zinc finger (C3HC4-type RING finger) family protein | 73 kDa | 0 | 0 | 43 | 15 | 58 | | | | 6.8E+05 | 8.3E+05 | 2.5E+05 | 1.7E+06 |
| AT5G51280 | AT5G51280.1 | DEAD-box protein abstrakt, putative | 66 kDa | 0 | 1 | 42 | 47 | 90 | | | | 2.3E+06 | 2.8E+06 | 1.6E+06 | 6.7E+06 |
| AT5G26240 | | chloride channel D | 87 kDa | 0 | 2 | 42 | 45 | 89 | | | | | | | |
| AT3G05060 | AT3G05060.1 | NOP56-like pre RNA processing ribonucleoprotein | 59 kDa | 0 | 4 | 42 | 43 | 89 | | 0.22 | 0.22 | 5.3E+06 | 4.9E+06 | 2.0E+06 | 1.2E+07 |
| AT1G56330 | AT1G56330.1;AT3G62560.1 | secretion-associated RAS 1B | 22 kDa | 17 | 21 | 42 | 41 | 121 | 0.24 | 0.41 | 0.32 | 1.6E+07 | 1.1E+07 | 2.1E+06 | 2.9E+07 |
| AT3G22310 | AT3G22310.1;AT3G22310.2 | putative mitochondrial RNA helicase 1 | 64 kDa | 0 | 5 | 42 | 40 | 87 | | | | 7.4E+06 | 7.9E+06 | 3.7E+06 | 1.9E+07 |
| AT3G62310 | AT3G62310.1;AT3G62310.3;AT3G62310.2 | RNA helicase family protein | 83 kDa | 2 | 0 | 42 | 38 | 82 | | | | 1.8E+06 | 1.6E+06 | 2.3E+05 | 3.6E+06 |
| AT2G47060 | AT2G47060.2;AT2G47060.1;AT2G47060.4;AT2G47060.5 | Protein kinase superfamily protein | 44 kDa | 6 | 21 | 42 | 36 | 105 | | | | 7.7E+05 | 0.0E+00 | 0.0E+00 | 7.7E+05 |
| AT2G39660 | AT2G39660.1;AT5G02290.1;AT5G02290.2 | botrytis-induced kinase1 | 44 kDa | 0 | 6 | 42 | 34 | 82 | | | | 3.7E+06 | 2.6E+06 | 0.0E+00 | 6.3E+06 |
| AT4G38600 | AT4G38600.1;AT4G38600.3;AT4G38600.2 | HEAT repeat ;HECT-domain (ubiquitin-transferase) | 203 kDa | 1 | 0 | 42 | 32 | 75 | | | | 5.1E+06 | 4.5E+06 | 6.3E+05 | 1.0E+07 |
| AT4G18060 | AT4G18060.1 | SH3 domain-containing protein | 40 kDa | 28 | 18 | 42 | 28 | 116 | | 3.55 | 3.55 | 1.1E+06 | 1.5E+06 | 0.0E+00 | 2.6E+06 |
| AT1G13900 | | Purple acid phosphatases superfamily protein | 74 kDa | 12 | 6 | 42 | 26 | 86 | | | | | | | |
| AT1G48240 | AT1G48240.1 | novel plant snare 12 | 30 kDa | 5 | 3 | 42 | 23 | 73 | | | | 8.4E+06 | 1.2E+07 | 0.0E+00 | 2.0E+07 |
| AT4G13510 | | ammonium transporter 1;1 | 54 kDa | 4 | 0 | 42 | 23 | 69 | | | | | | | |
| AT5G06110 | AT5G06110.2;AT5G06110.1 | DnaJ domain ;Myb-like DNA-binding domain | 76 kDa | 0 | 0 | 42 | 19 | 61 | | 0.20 | 0.20 | 1.4E+07 | 1.2E+07 | 4.0E+06 | 3.0E+07 |
| AT3G17440 | AT3G17440.2;AT3G17440.1 | novel plant snare 13 | 25 kDa | 4 | 5 | 42 | 14 | 65 | | 6.05 | 6.05 | 1.2E+07 | 1.4E+07 | 0.0E+00 | 2.6E+07 |
| AT3G16785 | | phospholipase D P1 | 125 kDa | 0 | 1 | 42 | 11 | 54 | | | | | | | |
| AT5G43330 | AT5G43330.1 | Lactate/malate dehydrogenase family protein | 36 kDa | 12 | 36 | 41 | 58 | 147 | | 5.26 | 5.26 | 3.2E+05 | 1.4E+06 | 0.0E+00 | 1.7E+06 |
| AT1G08680 | | ARF GAP-like zinc finger-containing protein ZIGA4 | 71 kDa | 0 | 7 | 41 | 55 | 103 | | | | | | | |
| AT4G02080 | AT4G02080.1 | secretion-associated RAS super family 2 | 22 kDa | 14 | 21 | 41 | 37 | 113 | | | | 3.7E+06 | 3.1E+06 | 0.0E+00 | 6.8E+06 |
| AT1G10580 | AT1G10580.1 | Transducin/WD40 repeat-like superfamily protein | 65 kDa | 0 | 0 | 41 | 34 | 75 | | | | 1.6E+07 | 2.5E+07 | 1.5E+06 | 4.3E+07 |
| AT4G39520 | AT4G39520.1 | GTP-binding protein-related | 41 kDa | 2 | 7 | 41 | 29 | 79 | | 0.70 | 0.70 | 2.3E+07 | 1.4E+07 | 1.2E+06 | 3.8E+07 |
| AT3G52580 | AT3G52580.1 | Ribosomal protein S11 family protein | 16 kDa | 14 | 0 | 41 | 27 | 82 | 0.22 | 0.47 | 0.35 | 3.3E+07 | 4.3E+07 | 1.1E+07 | 8.7E+07 |
| AT1G22410 | AT1G22410.1 | Class-II DAHP synthetase family protein | 58 kDa | 0 | 7 | 41 | 25 | 73 | | 0.27 | 0.27 | 3.2E+06 | 3.6E+06 | 3.9E+06 | 1.1E+07 |
| AT5G62190 | AT5G62190.1 | DEAD box RNA helicase (PRH75) | 73 kDa | 0 | 1 | 41 | 23 | 65 | | 0.22 | 0.22 | 2.7E+06 | 1.9E+06 | 2.6E+06 | 7.1E+06 |
| AT5G50420 | | O-fucosyltransferase family protein | 64 kDa | 0 | 0 | 40 | 62 | 102 | | | | | | | |
| AT1G18450 | AT1G18450.1 | actin-related protein 4 | 49 kDa | 0 | 2 | 40 | 47 | 89 | | | | 1.0E+07 | 1.0E+07 | 1.5E+06 | 2.2E+07 |
| AT3G44690 | | unknown protein; Has 30201 Blast hits to 17322 prote | 138 kDa | 0 | 0 | 40 | 44 | 84 | | | | | | | |
| AT3G60800 | AT3G60800.1 | DHHC-type zinc finger family protein | 35 kDa | 0 | 1 | 40 | 42 | 83 | | | | | | | |
| AT1G76680 | AT1G76680.1;AT1G76680.2;AT1G76690.1 | 12-oxophytodienoate reductase 1 | 41 kDa | 0 | 0 | 40 | 41 | 81 | | 6.83 | 6.83 | 2.6E+06 | 8.6E+05 | 0.0E+00 | 3.5E+06 |
| AT3G08590 | AT3G08590.1;AT3G08590.2 | Phosphoglycerate mutase, 2,3-bisphosphoglycerate-in | 61 kDa | 2 | 2 | 40 | 39 | 83 | | 0.58 | 0.58 | 2.3E+06 | 1.8E+06 | 0.0E+00 | 4.1E+06 |
| AT1G01620 | | plasma membrane intrinsic protein 1C | 31 kDa | 2 | 1 | 40 | 36 | 79 | | | | | | | |
| AT3G28450 | AT3G28450.1 | Leucine-rich repeat protein kinase family protein | 67 kDa | 1 | 1 | 40 | 31 | 73 | | 2.23 | 2.23 | 9.1E+06 | 4.7E+06 | 0.0E+00 | 1.4E+07 |
| AT1G71820 | AT1G71820.1;AT1G71820.2 | SEC6 | 86 kDa | 11 | 0 | 40 | 30 | 81 | | 3.78 | 3.78 | 3.3E+05 | 5.0E+05 | 0.0E+00 | 8.3E+05 |
| AT3G04230 | AT3G04230.1 | Ribosomal protein S5 domain 2-like superfamily protei | 17 kDa | 5 | 2 | 40 | 30 | 77 | | | | 6.5E+06 | 6.9E+06 | 1.1E+06 | 1.4E+07 |
| AT2G45810 | | DEA(D/H)-box RNA helicase family protein | 60 kDa | 0 | 0 | 40 | 29 | 69 | | 0.25 | 0.25 | | | | |
| AT5G22840 | AT5G22840.1 | Protein kinase superfamily protein | 61 kDa | 0 | 0 | 40 | 25 | 65 | | | | 6.5E+06 | 4.5E+06 | 9.1E+05 | 1.2E+07 |
| AT1G14000 | AT1G14000.1 | VH1-interacting kinase | 49 kDa | 1 | 6 | 40 | 24 | 71 | | | | 6.7E+06 | 6.4E+06 | 8.2E+05 | 1.4E+07 |
| AT3G16810 | AT3G16810.1 | pumilio 24 | 72 kDa | 0 | 0 | 40 | 22 | 62 | | | | 3.5E+06 | 1.9E+06 | 9.1E+05 | 6.3E+06 |
| AT5G40760 | AT5G40760.2;AT5G40760.1 | glucose-6-phosphate dehydrogenase 6 | 59 kDa | 0 | 3 | 40 | 20 | 63 | | | | 1.3E+07 | 1.1E+07 | 2.2E+05 | 2.4E+07 |
| AT1G08700 | | Presenilin-1 | 49 kDa | 0 | 0 | 40 | 19 | 59 | | | | | | | |
| AT4G15640 | AT4G15640.2;AT4G15640.1 | unknown protein; BEST Arabidopsis thaliana protein m | ? | 3 | 12 | 40 | 16 | 71 | | 0.57 | 0.57 | 2.1E+07 | 2.8E+07 | 2.9E+06 | 5.2E+07 |
| AT5G23740 | AT5G23740.1 | ribosomal protein S11-beta | 18 kDa | 12 | 7 | 40 | 16 | 75 | | 0.39 | 0.39 | 9.1E+07 | 1.2E+08 | 9.8E+06 | 2.2E+08 |
| AT5G10240 | AT5G10240.1;AT5G10240.2 | asparagine synthetase 3 | 65 kDa | 0 | 0 | 40 | 15 | 55 | | | | 3.3E+06 | 4.6E+06 | 5.9E+05 | 8.5E+06 |
| AT3G48500 | | Nucleic acid-binding, OB-fold-like protein | 79 kDa | 0 | 0 | 40 | 14 | 54 | | | | | | | |
| AT1G42550 | AT1G42550.1 | plastid movement impaired1 | 94 kDa | 0 | 0 | 39 | 57 | 96 | | | | 1.0E+06 | 9.4E+05 | 0.0E+00 | 2.0E+06 |
| AT1G08360 | AT1G08360.1 | Ribosomal protein L1p/L10e family | 24 kDa | 5 | 24 | 39 | 39 | 107 | | 0.15 | 0.15 | 2.4E+07 | 2.7E+07 | 7.3E+06 | 5.8E+07 |
| AT3G51670 | AT3G51670.1 | SEC14 cytosolic factor family protein / phosphoglycer | ? | 5 | 0 | 39 | 36 | 80 | | | | 7.8E+06 | 6.7E+06 | 8.5E+04 | 1.5E+07 |
| AT2G02800 | | protein kinase 2B | 46 kDa | 0 | 1 | 39 | 32 | 72 | | | | | | | |
| AT3G17240 | AT3G17240.3;AT3G17240.1 | lipoamide dehydrogenase 2 | 54 kDa | 7 | 2 | 39 | 32 | 80 | | | | 5.1E+06 | 1.4E+07 | 7.9E+05 | 2.0E+07 |
| AT2G40290 | AT2G40290.1;AT2G40290.3;AT2G40290.2 | Eukaryotic translation initiation factor 2 subunit 1 | 39 kDa | 4 | 6 | 39 | 29 | 78 | 0.14 | 0.21 | 0.18 | 2.8E+07 | 3.4E+07 | 1.2E+07 | 7.5E+07 |

| | | | | | | | | | | | | | | | |
|---|---|---|---|---|---|---|---|---|---|---|---|---|---|---|---|
| AT4G23850 | AT4G23850.1 | AMP-dependent synthetase and ligase family protein | 75 kDa | 0 | 0 | 39 | 27 | 66 | 0.11 | 0.49 | 0.30 | 8.2E+05 | 7.5E+05 | 1.5E+06 | 3.1E+06 |
| AT4G32520 | AT4G32520.2;AT4G32520.1 | serine hydroxymethyltransferase 3 | 58 kDa | 1 | 2 | 39 | 25 | 67 | | 0.25 | 0.25 | 4.0E+07 | 6.4E+07 | 2.2E+06 | 1.1E+08 |
| AT1G27090 | AT1G27090.1 | glycine-rich protein | ? | 0 | 0 | 39 | 24 | 63 | | 0.47 | 0.47 | 2.4E+07 | 2.1E+07 | 2.6E+06 | 4.8E+07 |
| AT4G25210 | AT4G25210.1 | DNA-binding storekeeper protein-related transcription | 40 kDa | 0 | 0 | 39 | 18 | 57 | | | | 1.4E+06 | 2.6E+06 | 0.0E+00 | 4.0E+06 |
| AT5G46700 | AT5G46700.1 | Tetraspanin family protein | 30 kDa | 11 | 18 | 39 | 18 | 86 | | 2.91 | 2.91 | 1.7E+08 | 1.5E+08 | 1.3E+07 | 3.3E+08 |
| AT3G55940 | | Phosphoinositide-specific phospholipase C family prot | 66 kDa | 0 | 1 | 39 | 15 | 55 | | | | | | | |
| AT1G04410 | AT1G04410.1 | Lactate/malate dehydrogenase family protein | 36 kDa | 12 | 24 | 38 | 67 | 141 | 0.83 | 5.65 | 3.24 | 4.2E+06 | 3.5E+06 | 0.0E+00 | 7.8E+06 |
| AT3G61650 | | gamma-tubulin | 53 kDa | 1 | 9 | 38 | 63 | 111 | 0.22 | 0.44 | 0.33 | | | | |
| AT2G44610 | AT2G44610.1;AT5G10260.1 | Ras-related small GTP-binding family protein | 23 kDa | 22 | 33 | 38 | 48 | 141 | | 0.50 | 0.50 | 8.3E+06 | 6.8E+06 | 2.7E+05 | 1.5E+07 |
| AT3G54820 | AT3G54820.1 | plasma membrane intrinsic protein 2;5 | 31 kDa | 3 | 1 | 38 | 44 | 86 | | | | 9.4E+06 | 6.2E+06 | 1.4E+07 | 2.9E+07 |
| AT1G07410 | | RAB GTPase homolog A2B | 24 kDa | 15 | 0 | 38 | 41 | 94 | | | | | | | |
| AT5G16050 | AT5G16050.2;AT5G16050.1 | general regulatory factor 5 | 30 kDa | 28 | 20 | 38 | 39 | 125 | 0.60 | 3.71 | 2.15 | 2.0E+06 | 1.2E+06 | 0.0E+00 | 3.2E+06 |
| AT4G15900 | AT4G15900.1 | pleiotropic regulatory locus 1 | 54 kDa | 0 | 0 | 38 | 34 | 72 | | 0.65 | 0.65 | 6.5E+06 | 1.0E+07 | 3.6E+05 | 1.7E+07 |
| AT5G52650 | AT5G52650.1 | RNA binding Plectin/S10 domain-containing protein | 20 kDa | 11 | 1 | 38 | 34 | 84 | 0.22 | 0.24 | 0.23 | 2.8E+07 | 2.3E+07 | 1.6E+06 | 5.2E+07 |
| AT1G34300 | AT1G34300.1 | lectin protein kinase family protein | 91 kDa | 0 | 0 | 38 | 31 | 69 | | | | 2.3E+06 | 1.8E+06 | 0.0E+00 | 4.1E+06 |
| AT3G28690 | | Protein kinase superfamily protein | 47 kDa | 0 | 3 | 38 | 31 | 72 | | | | | | | |
| AT5G35430 | AT5G35430.1 | Tetratricopeptide repeat (TPR)-like superfamily protein | 87 kDa | 0 | 0 | 38 | 27 | 65 | | | | 8.8E+06 | 1.0E+07 | 4.5E+05 | 1.9E+07 |
| AT3G62370 | | heme binding | 40 kDa | 4 | 6 | 38 | 26 | 74 | | 3.20 | 3.20 | | | | |
| AT2G17790 | AT2G17790.1 | VPS35 homolog A | 89 kDa | 0 | 4 | 38 | 25 | 67 | | 0.88 | 0.88 | 7.5E+05 | 1.1E+05 | 1.2E+05 | 9.8E+05 |
| AT5G49830 | | exocyst complex component 84B | 90 kDa | 7 | 2 | 38 | 25 | 72 | | | | | | | |
| AT2G06210 | AT2G06210.1 | binding | 124 kDa | 0 | 0 | 38 | 21 | 59 | | 0.50 | 0.50 | 5.0E+06 | 4.4E+06 | 5.1E+05 | 9.9E+06 |
| AT5G43980 | AT5G43980.1 | plasmodesmata-located protein 1 | 33 kDa | 30 | 16 | 38 | 21 | 105 | | 8.53 | 8.53 | 6.9E+07 | 6.5E+07 | 8.0E+04 | 1.3E+08 |
| AT1G69620 | AT1G69620.1 | ribosomal protein L34 | 14 kDa | 8 | 3 | 38 | 20 | 69 | 0.21 | | 0.21 | 2.3E+08 | 2.4E+08 | 6.1E+07 | 5.3E+08 |
| AT2G34040 | AT2G34040.1;AT2G34040.3;AT2G34040.2 | Apoptosis inhibitory protein 5 (API5) | 62 kDa | 0 | 4 | 38 | 12 | 54 | 0.11 | 0.25 | 0.18 | 1.3E+07 | 9.4E+06 | 2.9E+06 | 2.5E+07 |
| AT5G46570 | AT5G46570.1 | BR-signaling kinase 2 | 55 kDa | 2 | 6 | 37 | 53 | 98 | | | | 9.8E+05 | 1.1E+05 | 0.0E+00 | 1.1E+06 |
| AT2G02040 | AT2G02040.1 | peptide transporter 2 | 64 kDa | 0 | 0 | 37 | 42 | 79 | | 0.29 | 0.29 | 1.9E+07 | 5.8E+06 | 0.0E+00 | 2.4E+07 |
| AT2G25060 | AT2G25060.1 | early nodulin-like protein 14 | 19 kDa | 0 | 0 | 37 | 41 | 78 | | 1.97 | 1.97 | 1.7E+08 | 1.5E+08 | 2.7E+06 | 3.3E+08 |
| AT5G67630 | AT5G67630.1 | P-loop containing nucleoside triphosphate hydrolases | 52 kDa | 1 | 7 | 37 | 41 | 86 | | 0.45 | 0.45 | 9.1E+06 | 6.2E+06 | 1.8E+06 | 1.7E+07 |
| AT1G50700 | | calcium-dependent protein kinase 33 | 59 kDa | 0 | 2 | 37 | 39 | 78 | | | | | | | |
| AT2G20190 | AT2G20190.1 | CLIP-associated protein | 159 kDa | 0 | 17 | 37 | 35 | 89 | 0.12 | 1.12 | 0.62 | 6.6E+05 | 7.2E+05 | 2.3E+05 | 1.6E+06 |
| AT2G37970 | AT2G37970.1 | SOUL heme-binding family protein | 25 kDa | 10 | 26 | 37 | 35 | 108 | 0.44 | 2.78 | 1.61 | 6.0E+06 | 5.2E+06 | 1.3E+05 | 1.1E+07 |
| AT3G61820 | AT3G61820.1 | Eukaryotic aspartyl protease family protein | 51 kDa | 0 | 1 | 37 | 35 | 73 | | 0.45 | 0.45 | 8.0E+06 | 9.0E+06 | 4.3E+06 | 2.1E+07 |
| AT2G27600 | AT2G27600.1 | AAA-type ATPase family protein | ? | 3 | 1 | 37 | 33 | 74 | | | | 7.9E+05 | 6.5E+05 | 0.0E+00 | 1.4E+06 |
| AT1G30570 | AT1G30570.1 | hercules receptor kinase 2 | 94 kDa | 0 | 3 | 37 | 32 | 72 | | | | 3.9E+05 | 4.3E+05 | 0.0E+00 | 8.2E+05 |
| AT1G55150 | AT1G55150.1;AT1G55150.2 | DEA(D/H)-box RNA helicase family protein | 56 kDa | 0 | 8 | 37 | 31 | 76 | | 0.20 | 0.20 | 5.5E+06 | 4.2E+06 | 2.3E+06 | 1.2E+07 |
| AT4G02630 | AT4G02630.1 | Protein kinase superfamily protein | 55 kDa | 0 | 2 | 37 | 31 | 70 | | | | 1.1E+06 | 1.1E+06 | 0.0E+00 | 2.2E+06 |
| AT3G49870 | AT3G49870.2;AT3G49870.1;AT5G67560.1 | ADP-ribosylation factor-like A1C | 20 kDa | 8 | 3 | 37 | 30 | 78 | 0.52 | | 0.52 | 3.0E+07 | 1.5E+07 | 2.2E+06 | 4.8E+07 |
| AT3G01340 | | Transducin/WD40 repeat-like superfamily protein | 33 kDa | 6 | 16 | 37 | 26 | 85 | | | | | | | |
| AT4G02060 | AT4G02060.2;AT4G02060.1 | Minichromosome maintenance (MCM2/3/5) family pro | 80 kDa | 0 | 1 | 37 | 25 | 63 | 0.12 | 0.10 | 0.11 | 5.9E+06 | 3.9E+06 | 1.5E+06 | 1.1E+07 |
| AT1G72440 | AT1G72440.1 | CCAAT-binding factor | 117 kDa | 0 | 1 | 37 | 24 | 62 | | 0.33 | 0.33 | 4.9E+06 | 3.4E+06 | 1.3E+06 | 9.6E+06 |
| AT3G53110 | AT3G53110.1 | P-loop containing nucleoside triphosphate hydrolases | 55 kDa | 2 | 0 | 37 | 24 | 63 | | 0.62 | 0.62 | 6.3E+07 | 9.3E+07 | 3.1E+06 | 1.6E+08 |
| AT2G38360 | AT2G38360.1 | prenylated RAB acceptor 1.B4 | 24 kDa | 4 | 11 | 37 | 23 | 75 | 0.07 | 1.56 | 0.82 | 9.6E+06 | 1.3E+07 | 0.0E+00 | 2.3E+07 |
| AT3G06720 | AT3G06720.2;AT3G06720.1 | importin alpha isoform 1 | 59 kDa | 3 | 3 | 37 | 23 | 66 | | 0.43 | 0.43 | 3.6E+06 | 8.3E+06 | 5.7E+06 | 1.8E+07 |
| AT3G52470 | AT3G52470.1 | Late embryogenesis abundant (LEA) hydroxyproline-ric | 23 kDa | 11 | 6 | 37 | 20 | 74 | | 8.10 | 8.10 | 3.5E+07 | 2.1E+07 | 0.0E+00 | 5.6E+07 |
| AT1G43620 | AT1G43620.8;AT1G43620.7;AT1G43620.6;AT1G43620.5;AT1G43620.4;AT1G43620.3;AT1G43620.2;AT1G43620.1 | UDP-Glycosyltransferase superfamily protein | 68 kDa | 0 | 0 | 37 | 19 | 56 | | | | 1.5E+06 | 5.5E+05 | 0.0E+00 | 2.1E+06 |
| AT3G16780 | AT3G16780.1 | Ribosomal protein L19e family protein | 24 kDa | 0 | 10 | 37 | 17 | 64 | | | | 3.8E+07 | 5.2E+07 | 1.0E+07 | 1.0E+08 |
| AT5G55070 | AT5G55070.2;AT5G55070.1 | Dihydrolipoamide succinyltransferase | 50 kDa | 0 | 3 | 36 | 51 | 90 | | | | 4.8E+06 | 1.4E+07 | 4.8E+05 | 2.0E+07 |
| AT3G03050 | | cellulose synthase-like D3 | 129 kDa | 0 | 0 | 36 | 43 | 79 | | | | | | | |
| AT4G00660 | AT4G00660.2;AT4G00660.1 | RNAhelicase-like 8 | 58 kDa | 0 | 0 | 36 | 38 | 74 | | | | 1.8E+06 | 1.9E+06 | 1.2E+06 | 4.9E+06 |
| AT3G17750 | | Protein kinase superfamily protein | 127 kDa | 0 | 0 | 36 | 35 | 71 | | | | | | | |
| AT3G11730 | AT3G11730.1 | Ras-related small GTP-binding family protein | 23 kDa | 12 | 26 | 36 | 30 | 104 | | | | | | | |
| AT3G53780 | AT3G53780.2;AT3G53780.3 | RHOMBOID-like protein 4 | 44 kDa | 0 | 0 | 36 | 28 | 64 | | 3.49 | 3.49 | 4.4E+07 | 3.5E+07 | 4.4E+04 | 8.0E+07 |
| AT5G26360 | AT5G26360.1 | TCP-1/cpn60 chaperonin family protein | 60 kDa | 0 | 1 | 36 | 28 | 65 | | | | 1.2E+07 | 9.9E+06 | 1.8E+06 | 2.4E+07 |
| AT5G60960 | AT5G60960.1 | Pentatricopeptide repeat (PPR) superfamily protein | 59 kDa | 1 | 1 | 36 | 28 | 66 | | 0.80 | 0.80 | 8.3E+06 | 1.2E+07 | 1.4E+06 | 2.2E+07 |
| AT2G20840 | | Secretory carrier membrane protein (SCAMP) family pro | 32 kDa | 1 | 0 | 36 | 27 | 64 | | | | | | | |
| AT3G19760 | AT3G19760.1 | eukaryotic initiation factor 4A-III | ? | 0 | 3 | 36 | 27 | 66 | | 0.49 | 0.49 | 5.5E+06 | 2.8E+06 | 7.7E+05 | 9.1E+06 |
| AT5G65240 | | Leucine-rich repeat protein kinase family protein | 68 kDa | 1 | 0 | 36 | 27 | 64 | | | | | | | |
| AT3G58750 | AT3G58750.1 | citrate synthase 2 | 57 kDa | 0 | 11 | 36 | 23 | 70 | | 0.22 | 0.22 | 1.7E+07 | 1.8E+07 | 1.6E+06 | 3.7E+07 |
| AT5G24650 | AT5G24650.1 | Mitochondrial import inner membrane translocase sub | 28 kDa | 19 | 5 | 36 | 22 | 82 | 0.09 | 0.28 | 0.19 | 2.1E+07 | 3.4E+07 | 1.2E+07 | 6.7E+07 |
| AT5G35735 | AT5G35735.1 | Auxin-responsive family protein | 44 kDa | 1 | 0 | 36 | 18 | 55 | | | | 4.5E+06 | 3.6E+06 | 0.0E+00 | 8.1E+06 |
| AT1G29320 | AT1G29320.1;AT1G29320.2 | Transducin/WD40 repeat-like superfamily protein | 52 kDa | 0 | 0 | 36 | 16 | 52 | | 0.37 | 0.37 | 6.4E+06 | 1.1E+07 | 1.5E+06 | 1.9E+07 |
| AT2G33150 | AT2G33150.1 | peroxisomal 3-ketoacyl-CoA thiolase 3 | 49 kDa | 2 | 8 | 36 | 16 | 62 | | | | 7.6E+07 | 6.3E+07 | 5.0E+06 | 1.4E+08 |
| AT2G47980 | AT2G47980.1 | sister-chromatid cohesion protein 3 | 126 kDa | 0 | 0 | 36 | 16 | 52 | | | | 1.7E+05 | 4.0E+05 | 0.0E+00 | 5.7E+05 |
| AT1G04510 | AT1G04510.2;AT1G04510.1 | MOS4-associated complex 3A | 57 kDa | 0 | 0 | 35 | 47 | 82 | | 0.13 | 0.13 | 1.1E+07 | 1.2E+07 | 3.2E+06 | 2.7E+07 |
| AT4G09000 | AT4G09000.1;AT4G09000.2;AT1G35160.1;AT1G35160.2 | general regulatory factor 1 | 30 kDa | 27 | 30 | 35 | 47 | 139 | | 5.08 | 5.08 | 2.6E+06 | 2.4E+06 | 0.0E+00 | 5.0E+06 |
| AT2G33580 | AT2G33580.1 | Protein kinase superfamily protein | 73 kDa | 4 | 4 | 35 | 40 | 83 | | | | 2.3E+06 | 2.1E+06 | 0.0E+00 | 4.5E+06 |
| AT3G02530 | AT3G02530.1 | TCP-1/cpn60 chaperonin family protein | 59 kDa | 2 | 0 | 35 | 27 | 64 | | 0.21 | 0.21 | 1.8E+06 | 1.2E+06 | 3.1E+05 | 3.3E+06 |
| AT4G18465 | AT4G18465.1 | RNA helicase family protein | 78 kDa | 0 | 1 | 35 | 26 | 62 | | 0.18 | 0.18 | 1.6E+06 | 1.8E+06 | 1.1E+06 | 4.5E+06 |
| AT4G26600 | AT4G26600.1;AT4G26600.5;AT4G26600.3;AT4G26600.7;AT4G26600.6;AT4G26600.8;AT4G26600.4;AT4G26600.2 | S-adenosyl-L-methionine-dependent methyltransferase | ? | 0 | 0 | 35 | 26 | 61 | | 0.48 | 0.48 | 3.9E+06 | 2.4E+06 | 1.5E+06 | 7.8E+06 |
| AT2G18720 | AT2G18720.1;AT2G18720.2;AT2G18720.3 | Translation elongation factor EF1A/initiation factor IF2 | 51 kDa | 0 | 0 | 35 | 24 | 59 | | 0.27 | 0.27 | 1.6E+07 | 2.1E+07 | 2.6E+06 | 3.9E+07 |
| AT5G12480 | AT5G12480.1;AT5G12480.2 | calmodulin-domain protein kinase 7 | 60 kDa | 0 | 5 | 35 | 23 | 63 | | | | 2.1E+06 | 1.1E+06 | 0.0E+00 | 3.2E+06 |
| AT1G74790 | AT1G74790.1 | catalytics | 75 kDa | 4 | 0 | 35 | 22 | 61 | | 4.93 | 4.93 | 5.6E+05 | 1.0E+06 | 0.0E+00 | 1.6E+06 |
| AT3G09570 | AT3G09570.1 | Lung seven transmembrane receptor family protein | 50 kDa | 1 | 0 | 35 | 19 | 55 | | | | | | | |
| AT3G23540 | | alpha/beta-Hydrolases superfamily protein | 63 kDa | 0 | 3 | 35 | 19 | 57 | | | | | | | |
| AT5G63420 | AT5G63420.1 | RNA-metabolising metallo-beta-lactamase family prote | 101 kDa | 0 | 0 | 35 | 19 | 54 | | | | 2.0E+06 | 1.8E+06 | 4.7E+05 | 4.3E+06 |
| AT5G42080 | AT5G42080.2;AT5G42080.3;AT5G42080.4;AT5G42080.1 | dynamin-like protein | 68 kDa | 0 | 1 | 35 | 18 | 54 | | | | 7.2E+05 | 7.5E+05 | 0.0E+00 | 1.5E+06 |
| AT4G15770 | AT4G15770.1 | RNA binding | 21 kDa | 0 | 1 | 35 | 16 | 52 | | 0.24 | 0.24 | 9.6E+07 | 1.0E+08 | 3.3E+07 | 2.3E+08 |
| AT1G04260 | AT1G04260.1 | CAMV movement protein interacting protein 7 | 19 kDa | 9 | 0 | 35 | 11 | 55 | | 3.07 | 3.07 | 5.9E+06 | 4.6E+06 | 0.0E+00 | 1.0E+07 |
| AT5G47520 | | RAB GTPase homolog A5A | 24 kDa | 0 | 11 | 34 | 64 | 109 | | | | | | | |
| AT1G71890 | AT1G71890.1 | Major facilitator superfamily protein | 55 kDa | 0 | 0 | 34 | 54 | 88 | | | | 8.1E+05 | 6.7E+05 | 0.0E+00 | 1.5E+06 |
| ATCG00480 | ATCG00480.1 | ATP synthase subunit beta | 54 kDa | 7 | 6 | 34 | 50 | 97 | 0.12 | 0.45 | 0.28 | 9.2E+06 | 7.0E+06 | 1.3E+06 | 1.8E+07 |
| AT1G32860 | AT1G32860.1 | Glycosyl hydrolase superfamily protein | 45 kDa | 0 | 1 | 34 | 45 | 80 | | | | 1.9E+06 | 4.7E+06 | 0.0E+00 | 6.6E+06 |
| AT3G57990 | AT3G57990.1 | unknown protein; Has 1497 Blast hits to 1323 proteins | 40 kDa | 0 | 8 | 34 | 44 | 86 | | 1.76 | 1.76 | 5.3E+05 | 2.1E+06 | 1.1E+06 | 3.7E+06 |
| AT4G00710 | | BR-signaling kinase 3 | 55 kDa | 6 | 10 | 34 | 44 | 94 | | | | | | | |
| AT3G03960 | AT3G03960.1 | TCP-1/cpn60 chaperonin family protein | 59 kDa | 1 | 7 | 34 | 41 | 83 | | 0.76 | 0.76 | 2.4E+06 | 2.5E+06 | 1.7E+05 | 5.1E+06 |
| AT1G51630 | | O-fucosyltransferase family protein | 47 kDa | 3 | 10 | 34 | 40 | 87 | | | | | | | |

| | | | | | | | | | | | | | | | |
|---|---|---|---|---|---|---|---|---|---|---|---|---|---|---|---|
| AT4G35230 | AT4G35230.1 | BR-signaling kinase 1 | 57 kDa | 7 | 15 | 34 | 36 | 92 | | | | 3.9E+06 | 2.1E+06 | 0.0E+00 | 6.0E+06 |
| AT1G73590 | | Auxin efflux carrier family protein | 67 kDa | 5 | 2 | 34 | 32 | 73 | | | | | | | |
| AT1G72150 | AT1G72150.1 | PATELLIN 1 | 64 kDa | 0 | 0 | 34 | 27 | 61 | | | | 5.7E+05 | 9.7E+05 | 5.7E+04 | 1.6E+06 |
| AT1G69295 | AT1G69295.1;AT1G69295.2 | plasmodesmata callose-binding protein 4 | 22 kDa | 6 | 0 | 34 | 25 | 65 | | 12.43 | 12.43 | 6.1E+07 | 4.9E+07 | 2.9E+06 | 1.1E+08 |
| AT5G15610 | AT5G15610.2;AT5G15610.1 | Proteasome component (PCI) domain protein | 46 kDa | 2 | 24 | 34 | 25 | 85 | | 1.08 | 1.08 | 0.0E+00 | 1.2E+06 | 4.0E+05 | 1.6E+06 |
| AT1G43710 | AT1G43710.1 | Pyridoxal phosphate (PLP)-dependent transferases supe | 54 kDa | 0 | 0 | 34 | 23 | 57 | | 0.41 | 0.41 | 2.5E+06 | 1.8E+06 | 0.0E+00 | 4.2E+06 |
| AT4G35260 | AT4G35260.1;AT4G35650.1;AT2G17130.2;AT2G17130.1 | isocitrate dehydrogenase 1 | 40 kDa | 4 | 8 | 34 | 21 | 67 | | | | 3.0E+06 | 4.8E+06 | 2.2E+06 | 1.0E+07 |
| AT1G45688 | | unknown protein; FUNCTIONS IN: molecular_function | ? | 0 | 0 | 34 | 20 | 54 | | | | | | | |
| AT2G01630 | AT2G01630.1;AT2G01630.3;AT2G01630.2 | O-Glycosyl hydrolases family 17 protein | 54 kDa | 4 | 0 | 34 | 20 | 58 | | 7.95 | 7.95 | 1.0E+07 | 6.0E+06 | 2.8E+05 | 1.6E+07 |
| AT4G26910 | AT4G26910.3;AT4G26910.2;AT4G26910.1 | Dihydrolipoamide succinyltransferase | 40 kDa | 1 | 5 | 33 | 52 | 91 | | 0.86 | 0.86 | 1.2E+06 | 3.8E+06 | 6.7E+05 | 5.6E+06 |
| AT1G02130 | | RAS 5 | 23 kDa | 0 | 33 | 33 | 35 | 101 | | | | | | | |
| AT1G08560 |  | syntaxin of plants 111 | 35 kDa | 4 | 3 | 33 | 34 | 74 | 1.41 | 11.15 | 6.28 |  |  |  |  |
| AT1G49340 | | Phosphatidylinositol 3- and 4-kinase family protein | 224 kDa | 0 | 0 | 33 | 34 | 67 | | | | | | | |
| AT2G19580 | AT2G19580.1 | tetraspanin2 | 30 kDa | 8 | 15 | 33 | 29 | 85 | | 4.72 | 4.72 | 1.3E+07 | 9.7E+06 | 4.1E+05 | 2.3E+07 |
| AT3G57000 | AT3G57000.1 | nucleolar essential protein-related | 34 kDa | 0 | 0 | 33 | 28 | 61 | | | | 1.4E+07 | 1.3E+07 | 5.1E+06 | 3.2E+07 |
| AT4G16130 | AT4G16130.1 | arabinose kinase | 114 kDa | 0 | 0 | 33 | 28 | 61 | | 0.21 | 0.21 | 1.6E+06 | 1.2E+06 | 0.0E+00 | 2.8E+06 |
| AT4G14230 | | CBS domain-containing protein with a domain of unkn | 53 kDa | 0 | 1 | 33 | 27 | 61 | | | | | | | |
| AT5G07300 | AT5G07300.1 | Calcium-dependent phospholipid-binding Copine fami | 64 kDa | 1 | 0 | 33 | 27 | 61 | | | | 8.1E+05 | 7.4E+05 | 0.0E+00 | 1.5E+06 |
| AT5G60670 | AT5G60670.1 | Ribosomal protein L11 family protein | 18 kDa | 9 | 0 | 33 | 27 | 69 | | 0.83 | 0.83 | 8.7E+06 | 1.0E+07 | 9.1E+06 | 2.8E+07 |
| AT4G38350 | AT4G38350.1;AT4G38350.2 | Patched family protein | 140 kDa | 0 | 0 | 33 | 25 | 58 | | | | 5.9E+06 | 2.0E+06 | 1.3E+05 | 8.0E+06 |
| AT5G20350 | | Ankyrin repeat family protein with DHHC zinc finger do | 68 kDa | 1 | 3 | 33 | 25 | 62 | | | | | | | |
| AT5G67560 | | ADP-ribosylation factor-like A1D | 20 kDa | 0 | 0 | 33 | 25 | 58 | | | | | | | |
| AT3G59350 | | Protein kinase superfamily protein | 46 kDa | 9 | 9 | 33 | 24 | 75 | | | | | | | |
| AT5G15550 | AT5G15550.1;AT5G15550.3;AT5G15550.2 | Transducin/WD40 repeat-like superfamily protein | 47 kDa | 1 | 2 | 33 | 24 | 60 | | 0.19 | 0.19 | 2.5E+07 | 2.5E+07 | 8.7E+06 | 5.8E+07 |
| AT4G16630 | AT4G16630.1 | DEA(D/H)-box RNA helicase family protein | 89 kDa | 0 | 0 | 33 | 23 | 56 | | 0.35 | 0.35 | 4.5E+05 | 3.6E+05 | 9.5E+04 | 9.1E+05 |
| AT3G62120 | AT3G62120.3;AT3G62120.2;AT3G62120.1 | Class II aaRS and biotin synthetases superfamily protein | 61 kDa | 0 | 0 | 33 | 21 | 54 | | | | 7.0E+06 | 5.2E+06 | 1.4E+05 | 1.2E+07 |
| AT2G45960 | AT2G45960.2;AT2G45960.1;AT2G45960.3;AT1G01620.2;AT3G61430.2;AT3G61430.1;AT1G01620.1;AT4G23400.1;AT4G00430.1 | plasma membrane intrinsic protein 1B | 31 kDa | 2 | 2 | 33 | 20 | 57 | | | | 4.2E+07 | 1.6E+07 | 9.1E+06 | 6.7E+07 |
| AT4G03560 | AT4G03560.1 | two-pore channel 1 | 85 kDa | 0 | 0 | 33 | 19 | 52 |  | 0.18 | 0.18 | 1.6E+07 | 8.2E+06 | 0.0E+00 | 2.4E+07 |
| AT3G29810 | | COBRA-like protein 2 precursor | 50 kDa | 0 | 0 | 33 | 17 | 50 | | | | | | | |
| AT2G39420 | AT2G39420.1 | alpha/beta-Hydrolases superfamily protein | 36 kDa | 0 | 1 | 33 | 16 | 50 | | | | | | | |
| AT3G13670 | | Protein kinase family protein | 79 kDa | 0 | 1 | 33 | 16 | 50 | | | | | | | |
| AT2G07734 | | Alpha-L RNA-binding motif/Ribosomal protein S4 famil | 43 kDa | 0 | 0 | 33 | 12 | 45 | | 0.10 | 0.10 | | | | |
| AT1G53770 | | O-fucosyltransferase family protein | 64 kDa | 0 | 0 | 32 | 46 | 78 | | | | | | | |
| AT1G78300 | AT1G78300.1 | general regulatory factor 2 | 29 kDa | 16 | 12 | 32 | 42 | 102 | | 4.64 | 4.64 | 1.7E+06 | 3.8E+05 | 0.0E+00 | 2.1E+06 |
| AT5G62560 | AT5G62560.1;AT3G47820.1 | RING/U-box superfamily protein with ARM repeat dom | 60 kDa | 0 | 1 | 32 | 32 | 65 | | | | 9.9E+05 | 6.8E+05 | 0.0E+00 | 1.7E+06 |
| AT2G47990 | AT2G47990.1 | transducin family protein / WD-40 repeat family protei | 59 kDa | 0 | 0 | 32 | 31 | 63 | | | | 1.1E+07 | 9.6E+06 | 3.9E+06 | 2.5E+07 |
| AT1G80930 | AT1G80930.1 | MIF4G domain-containing protein / MA3 domain-cont | 104 kDa | 0 | 0 | 32 | 30 | 62 | | 0.28 | 0.28 | 1.2E+05 | 2.1E+05 | 6.1E+04 | 3.9E+05 |
| AT4G02620 | AT4G02620.1 | vacuolar ATPase subunit F family protein | 14 kDa | 4 | 0 | 32 | 28 | 64 | 0.51 | 0.61 | 0.56 | 2.1E+07 | 1.1E+07 | 3.6E+06 | 3.6E+07 |
| AT5G64370 | AT5G64370.1 | beta-ureidopropionase | 46 kDa | 0 | 0 | 32 | 28 | 60 | | | | 5.7E+07 | 5.4E+07 | 0.0E+00 | 1.1E+08 |
| AT2G42960 | | Protein kinase superfamily protein | 55 kDa | 0 | 1 | 32 | 27 | 60 | | | | | | | |
| ATCG00120 | ATCG00120.1 | ATP synthase subunit alpha | 55 kDa | 3 | 4 | 32 | 24 | 63 | 0.05 | 0.17 | 0.11 | 5.6E+06 | 3.1E+06 | 4.8E+05 | 9.2E+06 |
| AT3G07100 | AT3G07100.1 | Sec23/Sec24 protein transport family protein | 114 kDa | 1 | 0 | 32 | 23 | 56 | | | | 1.9E+06 | 1.2E+06 | 1.5E+05 | 3.3E+06 |
| AT5G50000 | | Protein kinase superfamily protein | 43 kDa | 0 | 1 | 32 | 23 | 56 | | | | | | | |
| AT2G47730 | AT2G47730.2;AT2G47730.1 | glutathione S-transferase phi 8 | 29 kDa | 1 | 2 | 32 | 22 | 57 | | | | 9.2E+06 | 8.6E+06 | 0.0E+00 | 1.8E+07 |
| AT4G36750 | AT4G36750.1 | Quinone reductase family protein | 29 kDa | 11 | 8 | 32 | 20 | 71 | | 5.85 | 5.85 | 2.6E+06 | 1.1E+06 | 0.0E+00 | 3.7E+06 |
| AT5G14170 | AT5G14170.1 | SWIB/MDM2 domain superfamily protein | 59 kDa | 0 | 1 | 32 | 20 | 53 | | | | 7.3E+06 | 6.9E+06 | 5.9E+05 | 1.5E+07 |
| AT5G22100 | AT5G22100.1 | RNA cyclase family protein | 41 kDa | 0 | 0 | 32 | 19 | 51 | | 0.41 | 0.41 | 5.0E+05 | 6.5E+05 | 9.6E+05 | 2.1E+06 |
| AT5G10190 | | Major facilitator superfamily protein | 53 kDa | 2 | 0 | 32 | 16 | 50 | | | | | | | |
| AT5G22360 | AT5G22360.1 | vesicle-associated membrane protein 714 | 25 kDa | 1 | 3 | 32 | 16 | 52 | | 0.31 | 0.31 | 2.1E+06 | 0.0E+00 | 2.3E+05 | 2.4E+06 |
| AT3G43980 | | Ribosomal protein S14p/S29e family protein | 6 kDa | 0 | 0 | 32 | 13 | 45 | | | | | | | |
| AT4G31810 | AT4G31810.1 | ATP-dependent caseinolytic (Clp) protease/crotonase fa | 46 kDa | 3 | 4 | 32 | 12 | 51 | | 2.14 | 2.14 | 3.4E+07 | 4.9E+07 | 5.7E+06 | 8.8E+07 |
| AT3G43810 | | calmodulin 7 | 17 kDa | 16 | 0 | 32 | 9 | 57 | | | | | | | |
| AT1G26670 | | Vesicle transport v-SNARE family protein | 25 kDa | 1 | 12 | 32 | 8 | 53 | 6.59 | | 6.59 | | | | |
| AT5G26980 | AT5G26980.3;AT5G26980.2;AT5G26980.1 | syntaxin of plants 41 | 36 kDa | 6 | 11 | 31 | 76 | 124 | | 12.55 | 12.55 | 0.0E+00 | 9.6E+05 | 0.0E+00 | 9.6E+05 |
| AT5G22300 | AT5G22300.1;AT5G22300.2 | nitrilase 4 | 39 kDa | 0 | 1 | 31 | 64 | 96 | | | | 1.9E+06 | 3.4E+06 | 9.5E+06 | 1.5E+07 |
| AT4G16480 | AT4G16480.3;AT4G16480.2;AT4G16480.1 | inositol transporter 4 | 63 kDa | 0 | 0 | 31 | 60 | 91 | | | | 7.2E+06 | 5.1E+06 | 0.0E+00 | 1.2E+07 |
| AT5G49630 | | amino acid permease 6 | 53 kDa | 0 | 0 | 31 | 53 | 84 | | | | | | | |
| AT3G63520 | AT3G63520.1 | carotenoid cleavage dioxygenase 1 | 61 kDa | 3 | 3 | 31 | 33 | 70 | | 1.32 | 1.32 | 1.3E+06 | 7.1E+05 | 0.0E+00 | 2.0E+06 |
| AT5G19280 | AT5G19280.1;AT5G19280.2 | kinase associated protein phosphatase | 65 kDa | 1 | 4 | 31 | 33 | 69 | | | | 1.0E+06 | 6.4E+05 | 1.1E+05 | 1.8E+06 |
| AT4G31490 | | Coatomer, beta subunit | 106 kDa | 0 | 0 | 31 | 28 | 59 | | | | | | | |
| AT1G07570 | | Protein kinase superfamily protein | 46 kDa | 0 | 1 | 31 | 26 | 58 | | | | | | | |
| AT2G32920 | AT2G32920.1 | PDI-like 2-3 | 48 kDa | 6 | 4 | 31 | 26 | 67 | | | | 5.5E+07 | 6.9E+07 | 5.9E+06 | 1.3E+08 |
| AT3G54540 | AT3G54540.2;AT3G54540.1 | general control non-repressible 4 | 80 kDa | 0 | 0 | 31 | 26 | 57 | | 0.21 | 0.21 | 6.4E+06 | 7.5E+06 | 2.2E+06 | 1.6E+07 |
| AT1G73450 | | Protein kinase superfamily protein | 129 kDa | 0 | 0 | 31 | 25 | 56 | | | | | | | |
| AT5G59520 | AT5G59520.1 | ZRT/IRT-like protein 2 | 38 kDa | 13 | 1 | 31 | 25 | 70 | | 3.27 | 3.27 | 4.7E+06 | 4.1E+06 | 0.0E+00 | 8.8E+06 |
| AT3G13050 | | Major facilitator superfamily protein | 55 kDa | 0 | 0 | 31 | 24 | 55 | | | | | | | |
| AT3G19300 | | Protein kinase superfamily protein | 73 kDa | 0 | 0 | 31 | 23 | 54 | | | | | | | |
| AT4G13010 | AT4G13010.1 | Oxidoreductase, zinc-binding dehydrogenase family pr | 34 kDa | 6 | 40 | 31 | 22 | 99 | | 1.12 | 1.12 | 4.2E+06 | 3.4E+06 | 0.0E+00 | 7.6E+06 |
| AT5G16070 | | TCP-1/cpn60 chaperonin family protein | 59 kDa | 0 | 0 | 31 | 20 | 51 | | | | | | | |
| AT1G47550 | AT1G47550.2;AT1G47550.1;AT1G47560.1 | exocyst complex component sec3A | 100 kDa | 18 | 1 | 31 | 19 | 69 | | | | 1.9E+05 | 2.1E+05 | 0.0E+00 | 4.0E+05 |
| AT3G11964 | AT3G11964.2;AT3G11964.1 | RNA binding;RNA binding | 212 kDa | 0 | 1 | 31 | 17 | 49 | | | | 8.3E+05 | 2.8E+05 | 6.5E+05 | 1.8E+06 |
| AT4G22120 | | ERD (early-responsive to dehydration stress) family prot | 88 kDa | 0 | 0 | 31 | 12 | 43 | | | | | | | |
| AT3G18600 | AT3G18600.1 | P-loop containing nucleoside triphosphate hydrolases | 64 kDa | 0 | 0 | 31 | 10 | 41 | | 0.25 | 0.25 | 5.9E+06 | 4.6E+06 | 3.0E+06 | 1.4E+07 |
| AT1G76850 | | exocyst complex component sec5 | 122 kDa | 10 | 1 | 31 | 9 | 51 | | | | | | | |
| AT5G65110 | AT5G65110.1;AT5G65110.2 | acyl-CoA oxidase 2 | 77 kDa | 0 | 0 | 31 | 7 | 38 | | | | 1.4E+06 | 1.1E+06 | 0.0E+00 | 2.5E+06 |
| AT3G13620 | | Amino acid permease family protein | 53 kDa | 0 | 0 | 30 | 50 | 80 | | | | | | | |
| AT3G01290 | | SPFH/Band 7/PHB domain-containing membrane-assoc | 31 kDa | 26 | 16 | 30 | 48 | 120 | | 5.83 | 5.83 | | | | |
| AT5G36230 | AT5G36230.1;AT5G36230.2 | ARM repeat superfamily protein | 47 kDa | 1 | 4 | 30 | 35 | 70 | | 0.41 | 0.41 | 3.4E+06 | 2.3E+06 | 2.4E+06 | 8.2E+06 |
| AT2G45470 | AT2G45470.1 | FASCICLIN-like arabinogalactan protein 8 | 43 kDa | 0 | 1 | 30 | 32 | 63 | | | | 4.3E+07 | 2.5E+07 | 6.0E+06 | 7.4E+07 |
| AT5G56460 | AT5G56460.1 | Protein kinase superfamily protein | 46 kDa | 7 | 10 | 30 | 32 | 79 | | | | 4.2E+05 | 2.8E+05 | 0.0E+00 | 7.0E+05 |
| AT4G18760 | AT4G18760.1 | receptor like protein 51 | 46 kDa | 3 | 0 | 30 | 30 | 63 | | | | 9.3E+06 | 3.7E+06 | 1.1E+06 | 1.4E+07 |
| AT1G54320 | AT1G54320.1 | LEM3 (ligand-effect modulator 3) family protein / CDC5 | 39 kDa | 1 | 2 | 30 | 28 | 61 | | | | 2.7E+06 | 2.2E+06 | 0.0E+00 | 4.9E+06 |
| AT2G40650 | AT2G40650.1 | PRP38 family protein | 43 kDa | 0 | 1 | 30 | 28 | 59 | | | | 1.7E+06 | 8.4E+05 | 2.9E+05 | 2.9E+06 |
| AT1G47240 | | NRAMP metal ion transporter 2 | 58 kDa | 0 | 0 | 30 | 27 | 57 | | | | | | | |
| AT3G11650 | | NDR1/HIN1-like 2 | 27 kDa | 1 | 4 | 30 | 27 | 62 | | 6.39 | 6.39 | | | | |
| AT2G11520 | AT2G11520.1 | calmodulin-binding receptor-like cytoplasmic kinase 3 | 57 kDa | 0 | 3 | 30 | 26 | 59 | | | | 7.3E+05 | 6.8E+05 | 0.0E+00 | 1.4E+06 |

| | | | | | | | | | | | | | | | |
|---|---|---|---|---|---|---|---|---|---|---|---|---|---|---|---|
| AT4G21700 | | Protein of unknown function (DUF2921) | 109 kDa | 0 | 0 | 30 | 26 | 56 | | | | | | | |
| AT5G41770 | AT5G41770.1 | crooked neck protein, putative / cell cycle protein, put | 85 kDa | 0 | 0 | 30 | 26 | 56 | | | | 7.3E+06 | 8.2E+06 | 3.8E+05 | 1.6E+07 |
| AT5G54380 | AT5G54380.1;AT2G21480.1 | protein kinase family protein | 93 kDa | 2 | 3 | 30 | 26 | 61 | | | | 9.3E+05 | 8.8E+05 | 0.0E+00 | 1.8E+06 |
| AT1G47830 | AT1G47830.1 | SNARE-like superfamily protein | 17 kDa | 2 | 4 | 30 | 25 | 61 | | 5.71 | 5.71 | 2.8E+07 | 2.7E+07 | 2.9E+06 | 5.7E+07 |
| AT2G47250 | AT2G47250.1 | RNA helicase family protein | 83 kDa | 0 | 0 | 30 | 25 | 55 | | 0.14 | 0.14 | 3.1E+05 | 0.0E+00 | 0.0E+00 | 3.1E+05 |
| AT3G58690 | AT3G58690.1 | Protein kinase superfamily protein | 44 kDa | 1 | 1 | 30 | 25 | 57 | | | | 1.0E+06 | 5.7E+05 | 0.0E+00 | 1.6E+06 |
| AT5G65700 | AT5G65700.2;AT5G65700.1 | Leucine-rich receptor-like protein kinase family proteir | 109 kDa | 0 | 0 | 30 | 24 | 54 | | | | 1.2E+06 | 4.1E+05 | 5.4E+04 | 1.6E+06 |
| AT2G40300 | AT2G40300.1 | ferritin 4 | 29 kDa | 9 | 0 | 30 | 23 | 62 | 1.50 | 3.04 | 2.27 | 1.8E+08 | 1.0E+08 | 0.0E+00 | 2.8E+08 |
| AT2G37340 | AT2G37340.1;AT2G37340.5;AT2G37340.3;AT2G37340.6;AT2G37340.4;AT2G37340.2 | arginine/serine-rich zinc knuckle-containing protein 3 | 33 kDa | 0 | 4 | 30 | 20 | 54 | 0.37 | 3.75 | 2.06 | 5.4E+07 | 5.6E+07 | 3.7E+06 | 1.1E+08 |
| AT3G44850 | AT3G44850.1 | Protein kinase superfamily protein | 61 kDa | 0 | 1 | 30 | 20 | 51 | | | | 2.5E+06 | 1.9E+06 | 1.5E+06 | 5.9E+06 |
| AT3G50530 | | CDPK-related kinase | 67 kDa | 0 | 0 | 30 | 18 | 48 | | | | | | | |
| AT4G28700 | | ammonium transporter 1;4 | 54 kDa | 0 | 0 | 30 | 17 | 47 | | | | | | | |
| AT5G55660 | AT5G55660.1 | DEK domain-containing chromatin associated protein | 87 kDa | 0 | 0 | 30 | 15 | 45 | | 0.91 | 0.91 | 5.5E+06 | 6.6E+06 | 1.7E+06 | 1.4E+07 |
| AT1G51850 | | Leucine-rich repeat protein kinase family protein | 96 kDa | 0 | 0 | 30 | 12 | 42 | | | | | | | |
| AT2G23810 | AT2G23810.1 | tetraspanin8 | 31 kDa | 20 | 37 | 30 | 12 | 99 | | 9.41 | 9.41 | 1.3E+08 | 7.8E+07 | 4.7E+06 | 2.1E+08 |
| AT4G36440 | AT4G36440.1 | unknown protein; FUNCTIONS IN: molecular_function | 43 kDa | 1 | 0 | 30 | 11 | 42 | | | | 0.0E+00 | 4.0E+04 | 6.4E+05 | 6.8E+05 |
| AT3G06530 | AT3G06530.3;AT3G06530.1;AT3G06530.2;AT3G06530.5;AT3G06530.4 | ARM repeat superfamily protein | 246 kDa | 0 | 1 | 29 | 45 | 75 | | 0.13 | 0.13 | 1.3E+06 | 1.4E+06 | 5.1E+05 | 3.3E+06 |
| AT4G24400 | AT4G24400.2;AT4G24400.1 | CBL-interacting protein kinase 8 | 50 kDa | 2 | 5 | 29 | 42 | 78 | | | | 3.8E+06 | 1.6E+06 | 0.0E+00 | 5.4E+06 |
| AT1G65730 | AT1G65730.1 | YELLOW STRIPE like 7 | 76 kDa | 0 | 0 | 29 | 41 | 70 | | | | 2.6E+06 | 1.2E+06 | 0.0E+00 | 3.7E+06 |
| AT5G15650 | | reversibly glycosylated polypeptide 2 | 41 kDa | 0 | 2 | 29 | 39 | 70 | | | | | | | |
| AT3G02520 | | general regulatory factor 7 | 30 kDa | 22 | 18 | 29 | 38 | 107 | | 4.36 | 4.36 | | | | |
| AT1G74470 | AT1G74470.1 | Pyridine nucleotide-disulphide oxidoreductase family | 52 kDa | 0 | 1 | 29 | 36 | 66 | | 0.36 | 0.36 | 9.2E+05 | 8.9E+05 | 0.0E+00 | 1.8E+06 |
| AT5G14120 | AT5G14120.1 | Major facilitator superfamily protein | 63 kDa | 0 | 0 | 29 | 34 | 63 | | 0.20 | 0.20 | 1.6E+06 | 4.4E+05 | 2.3E+05 | 2.2E+06 |
| AT4G21540 | AT4G21540.1;AT4G21540.3 | sphingosine kinase 1 | 54 kDa | 0 | 0 | 29 | 32 | 61 | | | | 5.6E+05 | 2.3E+05 | 0.0E+00 | 7.9E+05 |
| AT4G25940 | | ENTH/ANTH/VHS superfamily protein | 68 kDa | 4 | 2 | 29 | 32 | 67 | | | | | | | |
| AT1G61730 | AT1G61730.1 | DNA-binding storekeeper protein-related transcription | 41 kDa | 0 | 2 | 29 | 29 | 60 | | | | 5.9E+06 | 6.0E+06 | 0.0E+00 | 1.2E+07 |
| AT2G19470 | AT2G19470.1 | casein kinase I-like 5 | 49 kDa | 1 | 0 | 29 | 28 | 58 | | | | 7.8E+05 | 1.1E+06 | 0.0E+00 | 1.9E+06 |
| AT1G30580 | AT1G30580.1 | GTP binding | 44 kDa | 0 | 5 | 29 | 27 | 61 | 0.30 | 0.52 | 0.41 | 1.3E+07 | 7.9E+06 | 6.0E+05 | 2.1E+07 |
| AT5G59700 | | Protein kinase superfamily protein | 92 kDa | 0 | 3 | 29 | 26 | 58 | | | | | | | |
| AT1G26850 | | S-adenosyl-L-methionine-dependent methyltransferase | 70 kDa | 0 | 0 | 29 | 25 | 54 | | | | | | | |
| AT2G02560 | AT2G02560.2;AT2G02560.1 | cullin-associated and neddylation dissociated | 135 kDa | 1 | 0 | 29 | 25 | 55 | | | | 8.4E+06 | 6.0E+06 | 2.0E+06 | 1.6E+07 |
| AT4G16143 | AT4G16143.2;AT4G16143.1 | importin alpha isoform 2 | 59 kDa | 2 | 6 | 29 | 24 | 61 | 0.05 | 0.27 | 0.16 | 1.6E+06 | 1.1E+06 | 4.1E+06 | 6.8E+06 |
| AT3G12280 | AT3G12280.2;AT3G12280.1 | retinoblastoma-related 1 | 112 kDa | 0 | 2 | 29 | 22 | 53 | | | | 1.3E+06 | 2.0E+06 | 3.8E+05 | 3.7E+06 |
| AT2G23070 | AT2G23070.1;AT5G67380.2;AT3G50000.1;AT5G67380.1;AT2G23080.2;AT2G23080.1 | Protein kinase superfamily protein | 50 kDa | 0 | 1 | 29 | 21 | 51 | | | | 5.8E+06 | 5.5E+06 | 7.5E+05 | 1.2E+07 |
| AT4G18950 | AT4G18950.2;AT4G18950.1 | Integrin-linked protein kinase family | 53 kDa | 2 | 0 | 29 | 20 | 51 | | | | 4.5E+06 | 2.7E+06 | 4.2E+06 | 1.1E+07 |
| AT5G01600 | AT5G01600.1 | ferretin 1 | 28 kDa | 8 | 0 | 29 | 20 | 57 | 2.43 | | 2.43 | 1.8E+08 | 8.4E+07 | 5.2E+05 | 2.7E+08 |
| AT4G23270 | AT4G23270.1;AT4G23270.2;AT4G23270.3 | cysteine-rich RLK (RECEPTOR-like protein kinase) 19 | 72 kDa | 0 | 0 | 29 | 19 | 48 | | | | 3.4E+05 | 3.4E+05 | 0.0E+00 | 6.7E+05 |
| AT5G47320 | AT5G47320.1 | ribosomal protein S19 | 24 kDa | 4 | 10 | 29 | 17 | 60 | | 1.15 | 1.15 | 1.4E+08 | 2.6E+08 | 2.0E+07 | 4.2E+08 |
| AT1G50480 | AT1G50480.1 | 10-formyltetrahydrofolate synthetase | 68 kDa | 0 | 0 | 29 | 16 | 45 | | 0.50 | 0.50 | 5.3E+06 | 4.9E+06 | 1.1E+06 | 1.1E+07 |
| AT2G18900 | AT2G18900.1 | Transducin/WD40 repeat-like superfamily protein | 88 kDa | 0 | 0 | 29 | 15 | 44 | | 0.25 | 0.25 | 4.2E+06 | 5.1E+06 | 1.9E+06 | 1.1E+07 |
| AT5G65010 | AT5G65010.1;AT5G65010.2 | asparagine synthetase 2 | 65 kDa | 0 | 0 | 29 | 13 | 42 | | 0.18 | 0.18 | 4.2E+06 | 5.6E+06 | 0.0E+00 | 9.8E+06 |
| AT5G52820 | AT5G52820.1 | WD-40 repeat family protein / notchless protein, putat | 53 kDa | 0 | 1 | 29 | 12 | 42 | | 0.27 | 0.27 | 5.0E+06 | 4.8E+06 | 1.5E+06 | 1.1E+07 |
| AT5G40780 | | lysine histidine transporter 1 | 50 kDa | 0 | 0 | 29 | 10 | 39 | | | | | | | |
| AT3G59920 | AT3G59920.1 | RAB GDP dissociation inhibitor 2 | 50 kDa | 19 | 3 | 28 | 50 | 100 | | 35.80 | 35.80 | 9.0E+06 | 5.6E+06 | 0.0E+00 | 1.5E+07 |
| AT3G51840 | AT3G51840.1 | acyl-CoA oxidase 4 | 48 kDa | 28 | 28 | 28 | 45 | 129 | | | | 1.7E+07 | 1.9E+07 | 1.1E+05 | 3.6E+07 |
| AT1G35160 | | GF14 protein phi chain | 30 kDa | 19 | 18 | 28 | 36 | 101 | | | | | | | |
| AT5G16120 | AT5G16120.3;AT5G16120.1;AT5G16120.2;AT5G16120.4 | alpha/beta-Hydrolases superfamily protein | 39 kDa | 3 | 5 | 28 | 36 | 72 | 0.18 | | 0.18 | 1.1E+06 | 5.8E+05 | 0.0E+00 | 1.7E+06 |
| AT5G13000 | | glucan synthase-like 12 | 226 kDa | 0 | 0 | 28 | 29 | 57 | | | | | | | |
| AT1G60850 | AT1G60850.4;AT1G60850.2;AT1G60850.1;AT1G60850.3 | DNA-directed RNA polymerase family protein | 42 kDa | 0 | 13 | 28 | 28 | 69 | | 0.83 | 0.83 | 5.4E+06 | 4.8E+06 | 3.4E+06 | 1.4E+07 |
| AT4G14240 | | CBS domain-containing protein with a domain of unkn | 54 kDa | 0 | 0 | 28 | 28 | 56 | | | | | | | |
| AT5G58430 | | exocyst subunit exo70 family protein B1 | 71 kDa | 0 | 0 | 28 | 28 | 56 | | | | | | | |
| AT1G32380 | AT1G32380.1;AT1G32380.2 | phosphoribosyl pyrophosphate (PRPP) synthase 2 | 43 kDa | 0 | 8 | 28 | 27 | 63 | | | | 3.7E+06 | 3.4E+06 | 9.6E+05 | 8.0E+06 |
| AT2G42710 | AT2G42710.1 | Ribosomal protein L1p/L10e family | 46 kDa | 0 | 6 | 28 | 26 | 60 | | | | 2.2E+07 | 3.6E+07 | 4.9E+06 | 6.3E+07 |
| AT3G47960 | | Major facilitator superfamily protein | 71 kDa | 0 | 1 | 28 | 26 | 55 | | | | | | | |
| AT1G31850 | | S-adenosyl-L-methionine-dependent methyltransferase | 69 kDa | 0 | 1 | 28 | 23 | 52 | | | | | | | |
| AT1G78200 | AT1G78200.5;AT1G78200.4;AT1G78200.3;AT1G78200.2;AT1G78200.1 | Protein phosphatase 2C family protein | 31 kDa | 3 | 11 | 28 | 23 | 65 | | 10.57 | 10.57 | 6.3E+06 | 3.5E+06 | 0.0E+00 | 9.9E+06 |
| AT3G18790 | AT3G18790.1 | FUNCTIONS IN: molecular_function unknown; INVOLVE | 35 kDa | 1 | 0 | 28 | 22 | 51 | | 1.49 | 1.49 | 1.5E+07 | 2.2E+07 | 2.0E+06 | 3.9E+07 |
| AT5G22440 | AT5G22440.2;AT5G22440.1 | Ribosomal protein L1p/L10e family | 25 kDa | 0 | 29 | 28 | 22 | 79 | | | | 1.7E+07 | 2.0E+07 | 8.8E+06 | 4.6E+07 |
| AT1G73650 | AT1G73650.3;AT1G73650.1;AT1G73650.4;AT1G73650.5;AT1G73650.2 | Protein of unknown function (DUF1295) | 33 kDa | 1 | 5 | 28 | 21 | 55 | | 0.69 | 0.69 | 1.2E+07 | 1.6E+07 | 3.0E+05 | 2.8E+07 |
| AT1G77470 | | replication factor C subunit 3 | 41 kDa | 0 | 0 | 28 | 21 | 49 | | | | | | | |
| AT5G26830 | AT5G26830.1 | Threonyl-tRNA synthetase | 81 kDa | 0 | 1 | 28 | 21 | 50 | | 0.17 | 0.17 | 1.3E+07 | 1.4E+07 | 7.9E+05 | 2.8E+07 |
| AT1G07360 | AT1G07360.1 | CCCH-type zinc fingerfamily protein with RNA-binding | 54 kDa | 0 | 0 | 28 | 20 | 48 | | | | 5.6E+06 | 1.2E+07 | 4.2E+05 | 1.8E+07 |
| AT3G12780 | AT3G12780.1 | phosphoglycerate kinase 1 | 50 kDa | 3 | 2 | 28 | 20 | 53 | | | | 4.1E+06 | 1.8E+06 | 0.0E+00 | 5.9E+06 |
| AT1G20950 | AT1G20950.1 | Phosphofructokinase family protein | 67 kDa | 0 | 0 | 28 | 17 | 45 | | | | 6.5E+06 | 3.9E+06 | 5.3E+05 | 1.1E+07 |
| AT1G42960 | AT1G42960.1 | expressed protein localized to the inner membrane of t | 18 kDa | 8 | 0 | 28 | 17 | 53 | 0.15 | 0.18 | 0.16 | 1.2E+08 | 8.1E+07 | 2.3E+07 | 2.2E+08 |
| AT2G39990 | AT2G39990.1 | eukaryotic translation initiation factor 2 | 32 kDa | 11 | 37 | 28 | 17 | 93 | 0.10 | 0.62 | 0.36 | 2.5E+07 | 2.8E+07 | 1.3E+07 | 6.6E+07 |
| AT3G55620 | AT3G55620.1 | Translation initiation factor IF6 | 26 kDa | 2 | 2 | 28 | 17 | 49 | 0.23 | 0.47 | 0.35 | 1.5E+08 | 1.2E+08 | 4.5E+07 | 3.2E+08 |
| AT3G21630 | AT3G21630.1 | chitin elicitor receptor kinase 1 | 67 kDa | 0 | 0 | 28 | 16 | 44 | | | | 6.2E+05 | 4.4E+05 | 0.0E+00 | 1.1E+06 |
| AT5G50380 | AT5G50380.1 | exocyst subunit exo70 family protein F1 | 77 kDa | 0 | 0 | 28 | 16 | 44 | | | | 1.2E+05 | 8.1E+05 | 0.0E+00 | 9.3E+05 |
| AT1G26880 | AT1G26880.1;AT1G26880.2 | Ribosomal protein L34e superfamily protein | 14 kDa | 6 | 0 | 28 | 15 | 49 | | 0.49 | 0.49 | 9.3E+07 | 6.9E+07 | 4.2E+07 | 2.0E+08 |
| AT1G51710 | AT1G51710.2;AT1G51710.1 | ubiquitin-specific protease 6 | 54 kDa | 0 | 2 | 28 | 15 | 45 | | | | 5.6E+07 | 6.1E+07 | 2.7E+06 | 1.2E+08 |
| AT5G18420 | AT5G18420.3;AT5G18420.1;AT5G18420.2;AT5G18420.4 | unknown protein; FUNCTIONS IN: molecular_function | 50 kDa | 0 | 0 | 28 | 15 | 43 | | | | 4.4E+06 | 8.1E+06 | 2.0E+06 | 1.5E+07 |
| AT2G13790 | | somatic embryogenesis receptor-like kinase 4 | 69 kDa | 0 | 0 | 28 | 14 | 42 | | | | | | | |
| AT1G62390 | AT1G62390.1 | Octicosapeptide/Phox/Bem1p (PB1) domain-containin | 83 kDa | 0 | 0 | 28 | 13 | 41 | | 0.18 | 0.18 | 1.7E+07 | 2.3E+07 | 6.7E+06 | 4.7E+07 |
| AT5G04830 | AT5G04830.2;AT5G04830.1 | Nuclear transport factor 2 (NTF2) family protein | 20 kDa | 1 | 10 | 28 | 13 | 52 | 0.16 | 0.22 | 0.19 | 6.2E+06 | 8.7E+06 | 0.0E+00 | 1.5E+07 |
| AT5G15810 | AT5G15810.1 | N2,N2-dimethylguanosine tRNA methyltransferase | 77 kDa | 0 | 0 | 28 | 13 | 41 | | 0.48 | 0.48 | 1.4E+07 | 1.3E+07 | 2.3E+05 | 2.7E+07 |
| AT4G26100 | AT4G26100.1;AT5G57015.1;AT1G72710.1 | casein kinase 1 | 51 kDa | 0 | 0 | 28 | 12 | 40 | | | | 1.1E+05 | 8.8E+04 | 1.9E+05 | 3.9E+05 |
| AT3G11830 | AT3G11830.1;AT3G11830.2 | TCP-1/cpn60 chaperonin family protein | 60 kDa | 0 | 0 | 28 | 11 | 39 | | | | 2.6E+06 | 2.8E+06 | 8.4E+04 | 5.4E+06 |
| AT5G50320 | AT5G50320.1 | radical SAM domain-containing protein / GCN5-related | 64 kDa | 0 | 0 | 28 | 8 | 36 | | | | 2.3E+06 | 2.1E+06 | 5.1E+05 | 4.9E+06 |
| AT3G18280 | AT3G18280.1;AT3G18280.2 | Bifunctional inhibitor/lipid-transfer protein/seed stora | 10 kDa | 0 | 0 | 28 | 5 | 33 | | | | 7.0E+08 | 9.6E+08 | 9.7E+07 | 1.8E+09 |
| AT5G03630 | AT5G03630.1 | Pyridine nucleotide-disulphide oxidoreductase family | 47 kDa | 0 | 1 | 27 | 50 | 78 | | | | 4.8E+06 | 1.1E+06 | 0.0E+00 | 5.9E+06 |
| AT1G17620 | AT1G17620.1 | Late embryogenesis abundant (LEA) hydroxyproline-ric | ? | 4 | 4 | 27 | 38 | 73 | | 4.79 | 4.79 | 7.9E+06 | 7.0E+06 | 0.0E+00 | 1.5E+07 |

| | | | | | | | | | | | | | | | |
|---|---|---|---|---|---|---|---|---|---|---|---|---|---|---|---|
| AT1G22300 | AT1G22300.3;AT1G22300.2;AT1G22300.1 | general regulatory factor 10 | 29 kDa | 12 | 8 | 27 | 37 | 84 | | 6.36 | 6.36 | 6.9E+06 | 3.3E+06 | 0.0E+00 | 1.0E+07 |
| AT3G14840 | | Leucine-rich repeat transmembrane protein kinase | 112 kDa | 0 | 1 | 27 | 35 | 63 | | | | | | | |
| AT1G21880 | AT1G21880.1;AT1G21880.2 | lysm domain GPI-anchored protein 1 precursor | 43 kDa | 8 | 1 | 27 | 32 | 68 | | | | 2.6E+08 | 1.2E+08 | 8.2E+06 | 4.0E+08 |
| AT2G36070 | AT2G36070.1 | translocase inner membrane subunit 44-2 | 54 kDa | 3 | 8 | 27 | 29 | 67 | | | | | | | |
| AT5G10450 | AT5G10450.2;AT5G10450.1;AT5G10450.3;AT5G10450.4 | G-box regulating factor 6 | 28 kDa | 9 | 8 | 27 | 29 | 73 | | 1.84 | 1.84 | 2.0E+06 | 1.9E+06 | 0.0E+00 | 3.9E+06 |
| AT2G16250 | AT2G16250.1;AT2G16250.2 | Leucine-rich repeat protein kinase family protein | 100 kDa | 0 | 0 | 27 | 24 | 51 | | | | 1.8E+06 | 1.2E+06 | 0.0E+00 | 3.0E+06 |
| AT1G54610 | AT1G54610.3;AT1G54610.2;AT1G54610.1;AT5G50860.2;AT5G44290.2;AT1G09600.1;AT1G03740.2;AT1G53050.1;AT1G53050.2;AT5G44290.5;AT5G44290.1;AT5G44290.4;AT5G44290.3;AT5G44290.7;AT5G44290.6;AT3G05050.2;AT3G05050.1;AT3G05050.3;AT5G50860.1;AT1G03740.1 | Protein kinase superfamily protein | 63 kDa | 0 | 0 | 27 | 23 | 50 | | | | 2.3E+06 | 1.4E+06 | 0.0E+00 | 3.7E+06 |
| AT1G36180 | AT1G36180.1 | acetyl-CoA carboxylase 2 | 263 kDa | 0 | 0 | 27 | 22 | 49 | | 0.18 | 0.18 | 3.1E+06 | 2.1E+06 | 2.2E+05 | 5.4E+06 |
| AT2G45820 | AT2G45820.1 | Remorin family protein | 21 kDa | 17 | 54 | 27 | 22 | 120 | | | | 2.3E+07 | 1.7E+07 | 1.4E+06 | 4.2E+07 |
| AT2G15430 | AT2G15430.1;AT2G15430.3;AT2G15430.2 | DNA-directed RNA polymerase family protein | 35 kDa | 0 | 3 | 27 | 21 | 51 | | 1.48 | 1.48 | 3.0E+07 | 2.9E+07 | 2.0E+06 | 6.1E+07 |
| AT4G36945 | AT4G36945.1 | PLC-like phosphodiesterases superfamily protein | 44 kDa | 2 | 1 | 27 | 20 | 50 | | | | 2.0E+06 | 1.7E+06 | 0.0E+00 | 3.7E+06 |
| AT4G32410 | | cellulose synthase 1 | 122 kDa | 0 | 0 | 27 | 18 | 45 | | | | | | | |
| AT1G12920 | AT1G12920.1 | eukaryotic release factor 1-2 | 49 kDa | 1 | 4 | 27 | 17 | 49 | | | | 1.3E+06 | 2.4E+06 | 2.1E+05 | 3.9E+06 |
| AT2G01650 | AT2G01650.1;AT2G01650.2;AT2G01650.3 | plant UBX domain-containing protein 2 | 51 kDa | 33 | 8 | 27 | 17 | 85 | | 1.17 | 1.17 | 2.7E+06 | 3.8E+06 | 1.6E+05 | 6.7E+06 |
| AT3G49240 | AT3G49240.1 | Pentatricopeptide repeat (PPR) superfamily protein | 71 kDa | 0 | 0 | 27 | 16 | 43 | | 0.31 | 0.31 | 6.3E+06 | 7.3E+06 | 1.2E+06 | 1.5E+07 |
| AT1G06720 | AT1G06720.2;AT1G06720.1;AT1G06720.3 | P-loop containing nucleoside triphosphate hydrolases | ? | 0 | 0 | 27 | 15 | 42 | | | | 1.8E+06 | 2.3E+06 | 1.0E+06 | 5.1E+06 |
| AT1G53645 | AT1G53645.1 | hydroxyproline-rich glycoprotein family protein | 57 kDa | 3 | 0 | 27 | 14 | 44 | | | | 3.4E+07 | 3.5E+07 | 4.0E+06 | 7.3E+07 |
| AT1G76550 | AT1G76550.1 | Phosphofructokinase family protein | 68 kDa | 0 | 0 | 27 | 13 | 40 | | | | 3.3E+06 | 3.8E+06 | 0.0E+00 | 7.1E+06 |
| AT3G17465 | AT3G17465.1 | ribosomal protein L3 plastid | 35 kDa | 2 | 9 | 27 | 13 | 51 | | 1.26 | 1.26 | 2.3E+07 | 4.4E+07 | 4.3E+06 | 7.1E+07 |
| AT1G66340 | | Signal transduction histidine kinase, hybrid-type, ethyl | 83 kDa | 0 | 0 | 27 | 12 | 39 | | | | | | | |
| AT3G13560 | | O-Glycosyl hydrolases family 17 protein | 54 kDa | 0 | 0 | 27 | 11 | 38 | | | | | | | |
| AT4G32150 | | vesicle-associated membrane protein 711 | 25 kDa | 0 | 1 | 27 | 11 | 39 | | | | | | | |
| AT2G19740 | AT2G19740.1;AT4G26230.1 | Ribosomal protein L31e family protein | 14 kDa | 3 | 0 | 27 | 9 | 39 | | 0.28 | 0.28 | 2.6E+07 | 2.6E+07 | 0.0E+00 | 5.2E+07 |
| AT1G14830 | | DYNAMIN-like 1C | 69 kDa | 0 | 1 | 27 | 8 | 36 | | | | | | | |
| AT4G31985 | | Ribosomal protein L39 family protein | 6 kDa | 0 | 0 | 27 | 8 | 35 | | 0.12 | 0.12 | | | | |
| AT2G18740 | AT2G18740.1;AT2G18740.2;AT4G30330.1 | Small nuclear ribonucleoprotein family protein | 10 kDa | 2 | 0 | 27 | 4 | 33 | | 0.12 | 0.12 | 2.9E+07 | 4.3E+07 | 8.8E+06 | 8.1E+07 |
| AT5G61480 | AT5G61480.1 | Leucine-rich repeat protein kinase family protein | 115 kDa | 1 | 0 | 26 | 33 | 60 | | | | 8.2E+05 | 6.1E+05 | 0.0E+00 | 1.4E+06 |
| AT3G01540 | AT3G01540.1;AT3G01540.4;AT3G01540.3;AT3G01540.2 | DEAD box RNA helicase 1 | 68 kDa | 0 | 1 | 26 | 32 | 59 | | | | 3.1E+06 | 3.4E+06 | 4.1E+05 | 7.0E+06 |
| AT5G16910 | AT5G16910.1 | cellulose-synthase like D2 | 128 kDa | 0 | 0 | 26 | 31 | 57 | | | | | | | |
| AT4G02510 | AT4G02510.4;AT4G02510.3;AT4G02510.2;AT4G02510.1 | translocon at the outer envelope membrane of chlorop | 161 kDa | 0 | 0 | 26 | 27 | 53 | | 0.22 | 0.22 | 5.2E+05 | 4.6E+05 | 2.6E+04 | 1.0E+06 |
| AT4G22010 | AT4G22010.1 | SKU5 similar 4 | 60 kDa | 0 | 0 | 26 | 27 | 53 | | 0.26 | 0.26 | | | | |
| AT5G65270 | AT5G65270.1 | RAB GTPase homolog A4A | 25 kDa | 7 | 16 | 26 | 27 | 76 | | 0.46 | 0.46 | 6.6E+05 | 0.0E+00 | 0.0E+00 | 6.6E+05 |
| AT1G78880 | AT1G78880.1 | Ubiquitin-specific protease family C19-related protein | 50 kDa | 0 | 0 | 26 | 26 | 52 | | | | 3.5E+06 | 1.3E+06 | 0.0E+00 | 4.8E+06 |
| AT2G22660 | AT2G22660.2;AT2G22660.3;AT2G22660.1 | Protein of unknown function (duplicated DUF1399) | 89 kDa | 1 | 1 | 26 | 26 | 54 | | | | 4.7E+05 | 3.1E+05 | 0.0E+00 | 7.8E+05 |
| AT2G44530 | AT2G44530.2;AT2G44530.1 | Phosphoribosyltransferase family protein | 43 kDa | 0 | 5 | 26 | 26 | 57 | | | | 3.6E+06 | 9.1E+05 | 2.4E+06 | 6.9E+06 |
| AT4G03550 | | glucan synthase-like 5 | 207 kDa | 0 | 0 | 26 | 24 | 50 | | | | | | | |
| AT4G23250 | | kinases;protein kinases | 116 kDa | 0 | 0 | 26 | 24 | 50 | | | | | | | |
| AT2G35350 | | poltergeist like 1 | 87 kDa | 0 | 0 | 26 | 23 | 49 | | | | | | | |
| AT4G02010 | AT4G02010.1 | Protein kinase superfamily protein | 79 kDa | 0 | 1 | 26 | 22 | 49 | | | | 9.7E+05 | 7.6E+05 | 0.0E+00 | 1.7E+06 |
| AT3G55430 | AT3G55430.1 | O-Glycosyl hydrolases family 17 protein | 48 kDa | 0 | 1 | 26 | 21 | 48 | | 0.57 | 0.57 | 3.7E+06 | 2.8E+06 | 9.1E+05 | 7.4E+06 |
| AT5G49930 | AT5G49930.1 | zinc knuckle (CCHC-type) family protein | 121 kDa | 0 | 0 | 26 | 21 | 47 | | 0.29 | 0.29 | 5.4E+06 | 4.8E+06 | 2.7E+06 | 1.3E+07 |
| AT1G59820 | | aminophospholipid ATPase 3 | 138 kDa | 0 | 1 | 26 | 17 | 44 | | | | | | | |
| AT1G72710 | | casein kinase 1-like protein 2 | 52 kDa | 0 | 0 | 26 | 17 | 43 | | | | | | | |
| AT3G51850 | AT3G51850.3;AT3G51850.2;AT3G51850.1 | calcium-dependent protein kinase 13 | 59 kDa | 0 | 0 | 26 | 17 | 43 | | | | 6.1E+06 | 3.4E+06 | 0.0E+00 | 9.5E+06 |
| AT3G60830 | AT3G60830.1 | actin-related protein 7 | 40 kDa | 0 | 0 | 26 | 16 | 42 | | 1.58 | 1.58 | 1.1E+07 | 8.6E+06 | 2.6E+06 | 2.2E+07 |
| AT1G69850 | | nitrate transporter 1:2 | 64 kDa | 1 | 0 | 26 | 15 | 42 | | | | | | | |
| AT3G06650 | AT3G06650.2;AT3G06650.1 | ATP-citrate lyase B-1 | 66 kDa | 0 | 4 | 26 | 15 | 45 | 0.10 | 0.28 | 0.19 | 7.1E+05 | 4.2E+06 | 0.0E+00 | 4.9E+06 |
| AT5G23880 | AT5G23880.1 | cleavage and polyadenylation specificity factor 100 | 82 kDa | 0 | 1 | 26 | 15 | 42 | | | | 1.4E+06 | 1.4E+06 | 2.7E+05 | 3.2E+06 |
| AT5G43500 | AT5G43500.2;AT5G43500.1;AT5G43500.3;AT5G43500.4 | actin-related protein 9 | 66 kDa | 0 | 1 | 26 | 15 | 42 | | | | 0.0E+00 | 0.0E+00 | 0.0E+00 | 0.0E+00 |
| AT5G20920 | AT5G20920.2;AT5G20920.3;AT5G20920.1 | eukaryotic translation initiation factor 2 beta subunit | 31 kDa | 4 | 9 | 26 | 13 | 52 | 0.18 | | 0.18 | 5.0E+07 | 5.1E+07 | 1.3E+07 | 1.1E+08 |
| AT5G13560 | | unknown protein; BEST Arabidopsis thaliana protein m | ? | 0 | 0 | 26 | 12 | 38 | | | | | | | |
| AT5G18700 | AT5G18700.1 | Protein kinase family protein with ARM repeat domain | ? | 0 | 1 | 26 | 11 | 38 | | | | 3.7E+05 | 2.4E+05 | 8.8E+04 | 6.9E+05 |
| AT2G40840 | AT2G40840.1 | disproportionating enzyme 2 | 110 kDa | 0 | 0 | 26 | 1 | 27 | | | | 6.0E+06 | 9.1E+06 | 0.0E+00 | 1.5E+07 |
| AT1G01090 | AT1G01090.1 | pyruvate dehydrogenase E1 alpha | 47 kDa | 10 | 7 | 25 | 69 | 111 | | | | 2.8E+06 | 5.6E+05 | 7.5E+05 | 4.1E+06 |
| AT1G01540 | AT1G01540.2;AT1G01540.1;AT4G01330.3;AT4G01330.1;AT4G01330.2 | Protein kinase superfamily protein | 52 kDa | 0 | 0 | 25 | 31 | 56 | | | | 1.6E+07 | 5.5E+06 | 0.0E+00 | 2.2E+07 |
| AT1G77180 | AT1G77180.3;AT1G77180.2;AT1G77180.1 | chromatin protein family | 69 kDa | 0 | 0 | 25 | 31 | 56 | | 0.31 | 0.31 | 8.1E+06 | 9.1E+06 | 1.0E+06 | 1.8E+07 |
| AT3G25610 | | ATPase E1-E2 type family protein / haloacid dehalogena | 136 kDa | 0 | 0 | 25 | 28 | 53 | | | | | | | |
| AT1G06400 | AT1G06400.1 | Ras-related small GTP-binding family protein | 24 kDa | 15 | 12 | 25 | 27 | 79 | | | | 4.8E+06 | 1.9E+06 | 0.0E+00 | 6.7E+06 |
| AT5G13160 | | Protein kinase superfamily protein | 50 kDa | 2 | 2 | 25 | 27 | 56 | | | | | | | |
| AT5G11670 | | NADP-malic enzyme 2 | 64 kDa | 1 | 3 | 25 | 26 | 55 | 0.09 | | 0.09 | | | | |
| AT1G06470 | | Nucleotide/sugar transporter family protein | 46 kDa | 0 | 0 | 25 | 25 | 50 | | | | | | | |
| AT2G43770 | AT2G43770.1 | Transducin/WD40 repeat-like superfamily protein | 38 kDa | 0 | 0 | 25 | 25 | 50 | | 0.23 | 0.23 | 2.8E+07 | 3.7E+07 | 4.2E+06 | 7.0E+07 |
| AT5G64080 | AT5G64080.2;AT5G64080.1 | Bifunctional inhibitor/lipid-transfer protein/seed stora | 18 kDa | 2 | 6 | 25 | 24 | 57 | | 1.41 | 1.41 | 6.1E+08 | 6.3E+08 | 1.3E+07 | 1.2E+09 |
| AT1G47670 | AT1G47670.1 | Transmembrane amino acid transporter family protein | 57 kDa | 0 | 0 | 25 | 23 | 48 | | | | 2.6E+07 | 2.3E+07 | 0.0E+00 | 4.9E+07 |
| AT1G62810 | | Copper amine oxidase family protein | 80 kDa | 0 | 0 | 25 | 22 | 47 | | | | | | | |
| AT1G71380 | | cellulase 3 | 53 kDa | 0 | 0 | 25 | 20 | 45 | | 0.26 | 0.26 | | | | |
| AT2G39210 | | Major facilitator superfamily protein | 66 kDa | 0 | 0 | 25 | 20 | 45 | | | | | | | |
| AT4G23220 | | cysteine-rich RLK (RECEPTOR-like protein kinase) 14 | 82 kDa | 0 | 0 | 25 | 19 | 44 | | | | | | | |
| AT1G02690 | AT1G02690.1;AT1G02690.2 | importin alpha isoform 6 | 59 kDa | 0 | 2 | 25 | 18 | 45 | | 0.61 | 0.31 | 5.6E+05 | 7.6E+05 | 4.2E+05 | 1.7E+06 |
| AT1G75500 | AT1G75500.2;AT1G75500.1 | Walls Are Thin 1 | 43 kDa | 10 | 2 | 25 | 18 | 55 | 0.40 | 0.37 | 0.39 | 8.6E+06 | 8.0E+06 | 0.0E+00 | 1.7E+07 |
| AT5G05170 | | Cellulose synthase family protein | 120 kDa | 0 | 0 | 25 | 18 | 43 | | | | | | | |
| AT4G04695 | | calcium-dependent protein kinase 31 | 55 kDa | 0 | 0 | 25 | 17 | 42 | | | | | | | |
| AT4G33510 | AT4G33510.1;AT4G33510.2 | 3-deoxy-d-arabino-heptulosonate 7-phosphate synthas | 56 kDa | 0 | 2 | 25 | 17 | 44 | | 0.34 | 0.34 | 1.7E+06 | 8.5E+05 | 3.8E+05 | 2.9E+06 |

| | | | | | | | | | | | | | | | |
|---|---|---|---|---|---|---|---|---|---|---|---|---|---|---|---|
| AT5G14220 | AT5G14220.1;AT5G14220.4;AT5G14220.5;AT5G14220.3;AT5G14220.2 | Flavin containing amine oxidoreductase family | 56 kDa | 0 | 0 | 25 | 17 | 42 | 0.08 | 0.29 | 0.18 | 2.8E+07 | 4.4E+07 | 1.4E+06 | 7.3E+07 |
| AT4G34500 | | Protein kinase superfamily protein | 48 kDa | 0 | 0 | 25 | 16 | 41 | | | | | | | |
| AT3G45030 | | Ribosomal protein S10p/S20e family protein | 14 kDa | 5 | 0 | 25 | 15 | 45 | 0.06 | 0.46 | 0.26 | | | | |
| AT5G61770 | AT5G61770.1;AT5G61770.3;AT5G61770.2 | PETER PAN-like protein | 39 kDa | 0 | 0 | 25 | 15 | 40 | | | | 3.9E+06 | 3.5E+06 | 9.2E+05 | 8.3E+06 |
| AT2G24050 | AT2G24050.1 | MIF4G domain-containing protein / MA3 domain-cont | 83 kDa | 0 | 6 | 25 | 14 | 45 | | 0.20 | 0.20 | 8.1E+05 | 5.4E+05 | 3.3E+06 | 4.6E+06 |
| AT3G56110 | AT3G56110.4;AT3G56110.3;AT3G56110.2;AT3G56110.1 | prenylated RAB acceptor 1.B1 | 23 kDa | 10 | 3 | 25 | 14 | 52 | | | | 7.9E+06 | 1.1E+07 | 0.0E+00 | 1.9E+07 |
| AT1G53730 | AT1G53730.1;AT1G53730.2 | STRUBBELIG-receptor family 6 | 78 kDa | 0 | 0 | 25 | 13 | 38 | | | | 3.0E+06 | 1.3E+06 | 0.0E+00 | 4.2E+06 |
| AT2G28000 | AT2G28000.1 | chaperonin-60alpha | 62 kDa | 0 | 0 | 25 | 13 | 38 | 0.17 | 0.17 | 0.17 | 1.2E+06 | 8.5E+05 | 0.0E+00 | 2.0E+06 |
| AT4G10710 | AT4G10710.1;AT4G10710.2;AT4G10710.3 | global transcription factor C | 121 kDa | 0 | 0 | 25 | 12 | 37 | | | | 2.3E+06 | 1.8E+06 | 6.0E+05 | 4.7E+06 |
| AT5G25460 | AT5G25460.1 | Protein of unknown function, DUF642 | 40 kDa | 3 | 0 | 25 | 12 | 40 | | 0.14 | 0.14 | 4.5E+07 | 2.7E+07 | 0.0E+00 | 7.1E+07 |
| AT5G05470 | AT5G05470.1 | eukaryotic translation initiation factor 2 alpha subunit | 39 kDa | 0 | 2 | 25 | 9 | 36 | 0.22 | 0.65 | 0.44 | 1.7E+07 | 1.8E+07 | 8.2E+06 | 4.2E+07 |
| AT1G24706 | AT1G24706.3;AT1G24706.6;AT1G24706.5;AT1G24706.4;AT1G24706.1;AT1G24706.7;AT1G24706.2;AT1G24706.8;AT1G24706.9 | THO2 | 207 kDa | 0 | 0 | 25 | 8 | 33 | | | | 1.1E+06 | 1.2E+06 | 6.9E+05 | 3.0E+06 |
| AT3G09980 | | Family of unknown function (DUF662) | 21 kDa | 2 | 2 | 25 | 8 | 37 | | | | | | | |
| AT5G06870 | | polygalacturonase inhibiting protein 2 | 37 kDa | 0 | 0 | 25 | 8 | 33 | | | | | | | |
| AT1G51060 | | histone H2A 10 | 14 kDa | 10 | 0 | 25 | 7 | 42 | 0.26 | 0.12 | 0.19 | | | | |
| AT5G38480 | AT5G38480.3;AT5G38480.2;AT5G38480.1 | general regulatory factor 3 | 29 kDa | 24 | 18 | 24 | 38 | 104 | | 7.02 | 7.02 | 3.2E+06 | 2.3E+06 | 0.0E+00 | 5.5E+06 |
| AT2G01810 | AT2G01810.1 | RING/FYVE/PHD zinc finger superfamily protein | 79 kDa | 0 | 0 | 24 | 37 | 61 | | | | 3.5E+05 | 1.6E+05 | 0.0E+00 | 5.0E+05 |
| AT5G26780 | AT5G26780.4;AT5G26780.1;AT5G26780.3;AT5G26780.2 | serine hydroxymethyltransferase 2 | 57 kDa | 1 | 2 | 24 | 30 | 57 | | | | 1.8E+07 | 3.3E+07 | 3.4E+05 | 5.1E+07 |
| AT1G62380 | AT1G62380.1 | ACC oxidase 2 | 36 kDa | 8 | 17 | 24 | 28 | 77 | 2.25 | 9.64 | 5.95 | 2.3E+07 | 1.3E+07 | 0.0E+00 | 3.7E+07 |
| AT5G43780 | AT5G43780.1 | Pseudouridine synthase/archaeosine transglycosylase-l | 52 kDa | 0 | 10 | 24 | 28 | 62 | | | | 7.4E+06 | 4.0E+06 | 2.0E+06 | 1.3E+07 |
| AT1G11820 | AT1G11820.2;AT1G11820.3;AT1G11820.1 | O-Glycosyl hydrolases family 17 protein | 56 kDa | 0 | 1 | 24 | 27 | 52 | | | | 3.6E+06 | 2.1E+06 | 0.0E+00 | 5.7E+06 |
| AT1G15530 | AT1G15530.1;AT3G20860.1;AT3G59730.1;AT3G44200.2;AT4G29050.2;AT4G29050.1;AT1G70110.1;AT3G59700.1;AT3G59740.1;AT2G43700.1;AT1G70130.1;AT2G43700.4;AT2G43700.5;AT2G43700.2;AT2G43700.3;AT1G54510.4;AT1G54510.3;AT1G54510.2;AT1G54510.1;AT3G63280.4;AT3G04810.3;AT3G04810.2;AT3G04810.1;AT5G28290.2;AT5G28290.1;AT3G63280.1;AT3G63280.2;AT3G63280.3;AT5G28290.3;AT3G44200.1 | Concanavalin A-like lectin protein kinase family proteir | 73 kDa | 0 | 0 | 24 | 27 | 51 | | | | 1.5E+05 | 1.1E+05 | 0.0E+00 | 2.6E+05 |
| AT5G10020 | AT5G10020.1;AT5G10020.2 | Leucine-rich receptor-like protein kinase family proteir | 115 kDa | 0 | 0 | 24 | 26 | 50 | | | | 1.1E+06 | 8.0E+05 | 0.0E+00 | 1.9E+06 |
| AT3G13380 | | BRI1-like 3 | 127 kDa | 0 | 0 | 24 | 25 | 49 | | | | | | | |
| AT3G50000 | | casein kinase II, alpha chain 2 | 47 kDa | 2 | 5 | 24 | 25 | 56 | | | | | | | |
| AT2G31260 | | autophagy 9 (APG9) | 99 kDa | 0 | 0 | 24 | 24 | 48 | | | | | | | |
| AT1G56145 | | Leucine-rich repeat transmembrane protein kinase | 112 kDa | 0 | 2 | 24 | 23 | 49 | | | | | | | |
| AT1G55810 | | uridine kinase-like 3 | 52 kDa | 0 | 0 | 24 | 22 | 46 | | | | | | | |
| AT2G01830 | | CHASE domain containing histidine kinase protein | 118 kDa | 0 | 0 | 24 | 22 | 46 | | | | | | | |
| AT2G41560 | | autoinhibited Ca(2+)-ATPase, isoform 4 | 113 kDa | 0 | 0 | 24 | 22 | 46 | | | | | | | |
| AT1G74740 | | calcium-dependent protein kinase 30 | 61 kDa | 0 | 2 | 24 | 21 | 47 | | | | | | | |
| AT2G31810 | AT2G31810.1;AT2G31810.2;AT2G31810.3 | ACT domain-containing small subunit of acetolactate s | 54 kDa | 0 | 8 | 24 | 21 | 53 | | | | 1.6E+07 | 1.3E+07 | 6.7E+05 | 3.0E+07 |
| AT3G18240 | AT3G18240.2;AT3G18240.1;AT4G21460.1;AT4G21460.2 | Ribosomal protein S24/S35, mitochondrial | 48 kDa | 1 | 5 | 24 | 21 | 51 | | 1.89 | 1.89 | 6.3E+07 | 9.2E+07 | 1.0E+07 | 1.7E+08 |
| AT4G20410 | AT4G20410.1;AT4G20410.2 | gamma-soluble NSF attachment protein | 32 kDa | 7 | 12 | 24 | 21 | 64 | | 3.21 | 3.21 | 4.2E+06 | 2.0E+06 | 0.0E+00 | 6.3E+06 |
| AT5G67380 | | casein kinase alpha 1 | 48 kDa | 0 | 4 | 24 | 20 | 48 | | 2.11 | 2.11 | | | | |
| AT1G30680 | AT1G30680.1 | toprim domain-containing protein | 80 kDa | 0 | 0 | 24 | 19 | 43 | | | | 8.8E+06 | 5.5E+06 | 7.6E+05 | 1.5E+07 |
| AT1G76810 | AT1G76810.1 | eukaryotic translation initiation factor 2 (eIF-2) family | | 142 kDa | 0 | 0 | 24 | 19 | 43 | | 0.61 | 0.61 | 1.4E+07 | 1.1E+07 | 3.1E+06 | 2.8E+07 |
| AT2G18730 | AT2G18730.1 | diacylglycerol kinase 3 | 54 kDa | 0 | 0 | 24 | 18 | 42 | | | | 3.7E+06 | 2.4E+06 | 6.4E+04 | 6.2E+06 |
| AT1G32400 | AT1G32400.3;AT1G32400.2;AT1G32400.1 | tobamovirus multiplication 2A | 32 kDa | 6 | 8 | 24 | 17 | 55 | 0.48 | 0.97 | 0.72 | 9.9E+06 | 1.0E+07 | 0.0E+00 | 2.0E+07 |
| AT2G02860 | | sucrose transporter 2 | 64 kDa | 0 | 0 | 24 | 17 | 41 | | | | | | | |
| AT3G49080 | AT3G49080.1 | Ribosomal protein S5 domain 2-like superfamily protei | 48 kDa | 6 | 3 | 24 | 17 | 50 | | | | 5.7E+07 | 8.7E+07 | 1.1E+07 | 1.6E+08 |
| AT3G49370 | | Calcium-dependent protein kinase (CDPK) family prote | 66 kDa | 0 | 0 | 24 | 17 | 41 | | | | | | | |
| AT1G11050 | | Protein kinase superfamily protein | 69 kDa | 3 | 1 | 24 | 16 | 44 | | | | | | | |
| AT3G05090 | AT3G05090.3;AT3G05090.2;AT3G05090.1 | Transducin/WD40 repeat-like superfamily protein | 82 kDa | 0 | 0 | 24 | 16 | 40 | | | | 2.4E+06 | 2.0E+06 | 0.0E+00 | 4.4E+06 |
| AT3G25800 | AT3G25800.2;AT3G25800.3;AT3G25800.1;AT1G13320.4;AT1G13320.2;AT1G13320.3;AT1G13320.1;AT1G25490.1 | protein phosphatase 2A subunit A2 | 66 kDa | 0 | 1 | 24 | 16 | 41 | | | | 9.3E+05 | 7.0E+05 | 4.6E+04 | 1.7E+06 |
| AT4G35600 | | Protein kinase superfamily protein | 46 kDa | 0 | 1 | 24 | 16 | 41 | | | | | | | |
| AT5G63950 | AT5G63950.1;AT5G63950.2 | chromatin remodeling 24 | 123 kDa | 0 | 0 | 24 | 15 | 39 | | | | 8.4E+05 | 8.4E+05 | 1.1E+05 | 1.8E+06 |
| AT2G19730 | AT2G19730.3;AT2G19730.2;AT2G19730.1 | Ribosomal L28e protein family | 16 kDa | 4 | 0 | 24 | 13 | 41 | | 0.15 | 0.15 | 4.6E+07 | 6.9E+07 | 1.8E+07 | 1.3E+08 |
| AT2G22475 | | GRAM domain family protein | 32 kDa | 0 | 21 | 24 | 13 | 58 | | | | | | | |
| AT4G34870 | AT4G34870.1 | rotamase cyclophilin 5 | 18 kDa | 9 | 0 | 24 | 13 | 46 | 3.99 | | 3.99 | 1.1E+07 | 1.0E+07 | 0.0E+00 | 2.1E+07 |
| AT5G57015 | | casein kinase I-like 12 | 49 kDa | 0 | 0 | 24 | 13 | 37 | | | | | | | |
| AT4G30800 | AT4G30800.1 | Nucleic acid-binding, OB-fold-like protein | 18 kDa | 7 | 0 | 24 | 11 | 42 | | 0.19 | 0.19 | 1.1E+07 | 1.1E+07 | 6.0E+05 | 2.3E+07 |
| AT1G77590 | AT1G77590.1;AT1G77590.2 | long chain acyl-CoA synthetase 9 | 76 kDa | 0 | 0 | 24 | 10 | 34 | | | | 7.4E+05 | 9.5E+05 | 2.6E+05 | 1.9E+06 |
| AT1G79470 | AT1G79470.1 | Aldolase-type TIM barrel family protein | 54 kDa | 0 | 0 | 24 | 10 | 34 | | | | 3.0E+07 | 2.1E+07 | 2.5E+05 | 5.0E+07 |
| AT3G61110 | | ribosomal protein S27 | 10 kDa | 7 | 0 | 24 | 10 | 41 | | 0.20 | 0.20 | | | | |
| AT4G31580 | AT4G31580.2;AT4G31580.1 | serine/arginine-rich 22 | 22 kDa | 0 | 5 | 24 | 10 | 39 | 0.18 | 0.47 | 0.33 | 6.2E+07 | 6.7E+07 | 8.6E+06 | 1.4E+08 |
| AT1G15930 | AT1G15930.2;AT1G15930.1 | Ribosomal protein L7Ae/L30e/S12e/Gadd45 family prc | 15 kDa | 2 | 0 | 24 | 9 | 35 | | 0.12 | 0.12 | 1.2E+08 | 8.5E+07 | 2.8E+07 | 2.3E+08 |
| AT4G03415 | AT4G03415.3;AT4G03415.2;AT4G03415.1;AT1G03590.1 | Protein phosphatase 2C family protein | 52 kDa | 0 | 0 | 24 | 8 | 32 | | | | 6.9E+05 | 1.1E+06 | 0.0E+00 | 1.8E+06 |
| AT1G25490 | | ARM repeat superfamily protein | 65 kDa | 0 | 0 | 24 | 7 | 31 | | | | | | | |
| AT5G47210 | AT5G47210.1;AT5G47210.3;AT5G47210.2 | Hyaluronan / mRNA binding family | 38 kDa | 0 | 0 | 24 | 7 | 31 | | | | 6.3E+06 | 6.1E+06 | 8.9E+06 | 2.1E+07 |
| AT3G46280 | | protein kinase-related | ? | 0 | 5 | 24 | 6 | 35 | | | | | | | |
| AT3G02230 | | reversibly glycosylated polypeptide 1 | 41 kDa | 0 | 2 | 23 | 34 | 59 | | 1.73 | 1.73 | | | | |
| AT3G44150 | AT3G44150.1 | unknown protein; FUNCTIONS IN: molecular_function | ? | 14 | 16 | 23 | 32 | 85 | | 3.16 | 3.16 | 3.2E+07 | 3.0E+07 | 1.0E+06 | 6.4E+07 |

| | | | | | | | | | | | | | | | |
|---|---|---|---|---|---|---|---|---|---|---|---|---|---|---|---|
| AT3G22890 | AT3G22890.1 | ATP sulfurylase 1 | 51 kDa | 2 | 5 | 23 | 30 | 60 | | | | 5.9E+06 | 5.8E+06 | 1.4E+06 | 1.3E+07 |
| AT3G49750 | AT3G49750.2;AT3G49750.1;AT5G65830.1 | receptor like protein 44 | 30 kDa | 3 | 12 | 23 | 30 | 68 | | 25.94 | 25.94 | 1.8E+07 | 1.7E+07 | 0.0E+00 | 3.5E+07 |
| AT5G67470 | | formin homolog 6 | 99 kDa | 0 | 0 | 23 | 28 | 51 | | | | | | | |
| AT2G26890 | AT2G26890.1 | DNAJ heat shock N-terminal domain-containing protei | 279 kDa | 0 | 0 | 23 | 26 | 49 | | | | 2.1E+05 | 1.3E+05 | 0.0E+00 | 3.5E+05 |
| AT2G35390 | AT2G35390.3 | Phosphoribosyltransferase family protein | 38 kDa | 0 | 8 | 23 | 22 | 53 | | 0.71 | 0.71 | 0.0E+00 | 0.0E+00 | 7.3E+05 | 7.3E+05 |
| AT5G02290 | | Protein kinase superfamily protein | 44 kDa | 0 | 2 | 23 | 22 | 47 | | | | | | | |
| AT2G41820 | AT2G41820.1 | Leucine-rich repeat protein kinase family protein | 97 kDa | 0 | 0 | 23 | 21 | 44 | | | | 2.1E+06 | 1.9E+07 | 0.0E+00 | 2.1E+07 |
| AT5G47100 | AT5G47100.1;AT5G47100.2 | calcineurin B-like protein 9 | 25 kDa | 3 | 18 | 23 | 21 | 65 | | | | 6.7E+06 | 1.3E+06 | 0.0E+00 | 7.9E+06 |
| AT3G50050 | | Eukaryotic aspartyl protease family protein | 70 kDa | 0 | 0 | 23 | 20 | 43 | | | | | | | |
| AT4G39990 | AT4G39990.1 | RAB GTPase homolog A4B | 24 kDa | 5 | 14 | 23 | 20 | 62 | | | | 4.2E+06 | 1.8E+06 | 3.6E+05 | 6.4E+06 |
| AT5G39510 | AT5G39510.1 | Vesicle transport v-SNARE family protein | 25 kDa | 4 | 6 | 23 | 20 | 53 | 0.39 | 1.60 | 1.00 | 4.0E+06 | 2.8E+06 | 9.9E+05 | 7.7E+06 |
| AT1G18850 | AT1G18850.1 | unknown protein; FUNCTIONS IN: molecular_function | 44 kDa | 0 | 2 | 23 | 19 | 44 | | | | 2.5E+06 | 2.9E+06 | 2.0E+06 | 7.5E+06 |
| AT1G15500 | AT1G15500.1;AT1G80300.1 | TLC ATP/ADP transporter | 68 kDa | 0 | 0 | 23 | 18 | 41 | 0.37 | 0.09 | 0.23 | 3.0E+06 | 3.9E+06 | 5.6E+04 | 7.0E+06 |
| AT3G01280 | AT3G01280.1 | voltage dependent anion channel 1 | 29 kDa | 6 | 0 | 23 | 18 | 47 | 0.02 | 0.18 | 0.10 | 4.6E+06 | 1.3E+07 | 4.9E+06 | 2.2E+07 |
| AT5G64440 | AT5G64440.1 | fatty acid amide hydrolase | 66 kDa | 0 | 2 | 23 | 18 | 43 | | | | 9.4E+05 | 3.0E+05 | 0.0E+00 | 1.2E+06 |
| AT1G53165 | | Protein kinase superfamily protein | 75 kDa | 5 | 0 | 23 | 17 | 45 | | | | | | | |
| AT3G17020 | AT3G17020.1 | Adenine nucleotide alpha hydrolases-like superfamily p | 18 kDa | 20 | 0 | 23 | 17 | 60 | 4.16 | 2.39 | 3.28 | 6.6E+06 | 6.5E+06 | 2.4E+06 | 1.6E+07 |
| AT4G09720 | AT4G09720.6;AT4G09720.5;AT4G09720.2;AT4G09720.1;AT4G09720.4;AT4G09720.3 | RAB GTPase homolog G3A | 23 kDa | 1 | 9 | 23 | 17 | 50 | | 0.99 | 0.99 | 4.6E+06 | 3.3E+06 | 2.3E+05 | 8.1E+06 |
| AT1G48620 | AT1G48620.1 | high mobility group A5 | 51 kDa | 0 | 0 | 23 | 16 | 39 | | | | 1.4E+06 | 2.9E+06 | 9.6E+05 | 5.2E+06 |
| AT1G52320 | AT1G52320.5;AT1G52320.2;AT1G52320.1;AT1G52320.4;AT1G52320.3 | unknown protein; FUNCTIONS IN: molecular_function | 90 kDa | 0 | 0 | 23 | 16 | 39 | | | | 1.5E+06 | 1.7E+06 | 0.0E+00 | 3.1E+06 |
| AT1G60200 | AT1G60200.4;AT1G60200.3;AT1G60200.2;AT1G60200.1 | splicing factor PWI domain-containing protein / RNA re | 101 kDa | 0 | 0 | 23 | 16 | 39 | | 0.29 | 0.29 | 7.3E+06 | 7.6E+06 | 1.2E+06 | 1.6E+07 |
| AT1G06660 | | unknown protein; BEST Arabidopsis thaliana protein m | ? | 0 | 0 | 23 | 15 | 38 | | | | | | | |
| AT4G23180 | | cysteine-rich RLK (RECEPTOR-like protein kinase) 10 | 74 kDa | 0 | 0 | 23 | 15 | 38 | | | | | | | |
| AT5G49460 | AT5G49460.2;AT5G49460.1 | ATP citrate lyase subunit B 2 | 66 kDa | 0 | 4 | 23 | 13 | 40 | | | | 3.7E+05 | 3.4E+06 | 0.0E+00 | 3.7E+06 |
| AT3G47370 | AT3G47370.3;AT3G47370.2;AT3G47370.1 | Ribosomal protein S10p/S20e family protein | 14 kDa | 0 | 0 | 23 | 12 | 35 | 0.05 | 0.12 | 0.09 | 6.2E+07 | 6.0E+07 | 1.3E+07 | 1.4E+08 |
| AT3G44190 | AT3G44190.2;AT3G44190.1 | FAD/NAD(P)-binding oxidoreductase family protein | 40 kDa | 3 | 3 | 23 | 11 | 40 | | | | 2.9E+06 | 2.7E+06 | 4.0E+05 | 6.0E+06 |
| AT5G09860 | AT5G09860.2;AT5G09860.1 | nuclear matrix protein-related | 68 kDa | 0 | 0 | 23 | 11 | 34 | | | | 1.5E+06 | 1.9E+06 | 5.8E+05 | 4.0E+06 |
| AT5G17910 | AT5G17910.3;AT5G17910.2;AT5G17910.1 | unknown protein; FUNCTIONS IN: molecular_function | 150 kDa | 0 | 0 | 22 | 66 | 88 | | | | 7.9E+04 | 5.0E+04 | 0.0E+00 | 1.3E+05 |
| AT4G30990 | AT4G30990.1;AT4G30990.2;AT4G30990.3 | ARM repeat superfamily protein | 295 kDa | 0 | 1 | 22 | 44 | 67 | | 0.15 | 0.15 | 1.5E+05 | 1.1E+05 | 8.0E+05 | 1.1E+06 |
| AT1G29470 | | S-adenosyl-L-methionine-dependent methyltransferase | 87 kDa | 0 | 1 | 22 | 40 | 63 | | | | | | | |
| AT2G36570 | AT2G36570.1 | Leucine-rich repeat protein kinase family protein | 74 kDa | 2 | 2 | 22 | 37 | 63 | | | | 6.0E+06 | 2.7E+06 | 0.0E+00 | 8.7E+06 |
| AT1G17270 | | O-fucosyltransferase family protein | 64 kDa | 0 | 1 | 22 | 35 | 58 | | | | | | | |
| AT2G36910 | AT2G36910.1 | ATP binding cassette subfamily B1 | 141 kDa | 1 | 0 | 22 | 31 | 54 | | | | 2.3E+06 | 6.3E+05 | 0.0E+00 | 2.9E+06 |
| AT1G36160 | AT1G36160.2;AT1G36160.1 | acetyl-CoA carboxylase 1 | 251 kDa | 0 | 0 | 22 | 26 | 48 | | | | 2.1E+05 | 7.0E+04 | 0.0E+00 | 2.8E+05 |
| AT2G39805 | AT2G39805.1;AT2G39805.2 | Integral membrane Yip1 family protein | 31 kDa | 1 | 0 | 22 | 24 | 47 | | | | 1.5E+07 | 2.5E+07 | 6.2E+05 | 4.0E+07 |
| AT1G42970 | AT1G42970.1 | glyceraldehyde-3-phosphate dehydrogenase B subunit | 48 kDa | 0 | 1 | 22 | 22 | 45 | | | | 1.2E+07 | 2.3E+07 | 0.0E+00 | 3.5E+07 |
| AT1G53430 | | Leucine-rich repeat transmembrane protein kinase | 114 kDa | 0 | 1 | 22 | 22 | 45 | | | | | | | |
| AT3G26935 | AT3G26935.1 | DHHC-type zinc finger family protein | 50 kDa | 0 | 0 | 22 | 22 | 44 | | | | 6.7E+05 | 3.1E+05 | 0.0E+00 | 9.8E+05 |
| AT3G15890 | | Protein kinase superfamily protein | 41 kDa | 0 | 2 | 22 | 20 | 44 | | 6.26 | 6.26 | | | | |
| AT5G18610 | AT5G18610.3;AT5G18610.2;AT5G18610.1 | Protein kinase superfamily protein | 56 kDa | 0 | 1 | 22 | 19 | 42 | | | | 7.3E+05 | 4.2E+05 | 0.0E+00 | 1.2E+06 |
| AT1G68220 | | Protein of unknown function (DUF1218) | 21 kDa | 0 | 0 | 22 | 17 | 39 | | | | | | | |
| AT3G14350 | AT3G14350.2;AT3G14350.3;AT3G14350.1 | STRUBBELIG-receptor family 7 | 74 kDa | 3 | 1 | 22 | 17 | 43 | | | | 1.5E+06 | 1.6E+06 | 0.0E+00 | 3.0E+06 |
| AT4G35790 | AT4G35790.3;AT4G35790.2;AT4G35790.1 | phospholipase D delta | 99 kDa | 0 | 0 | 22 | 16 | 38 | | | | 3.1E+05 | 1.7E+05 | 0.0E+00 | 4.9E+05 |
| AT1G48050 | AT1G48050.1;AT1G48050.2;AT1G48050.3 | Ku80 family protein | 77 kDa | 0 | 0 | 22 | 15 | 37 | | | | 1.5E+06 | 2.3E+06 | 9.0E+04 | 3.8E+06 |
| AT1G78380 | AT1G78380.1 | glutathione S-transferase TAU 19 | 26 kDa | 0 | 8 | 22 | 15 | 45 | | | | 4.7E+06 | 4.1E+06 | 0.0E+00 | 8.9E+06 |
| AT5G17410 | AT5G17410.1;AT5G17410.2 | Spc97 / Spc98 family of spindle pole body (SBP) compo | 77 kDa | 0 | 0 | 22 | 14 | 36 | | | | 0.0E+00 | 0.0E+00 | 6.5E+04 | 6.5E+04 |
| AT5G20830 | | sucrose synthase 1 | 93 kDa | 5 | 1 | 22 | 14 | 42 | | | | | | | |
| AT1G18180 | AT1G18180.2;AT1G18180.1 | Protein of unknown function (DUF1295) | 35 kDa | 0 | 0 | 22 | 13 | 35 | | | | 2.3E+06 | 4.9E+05 | 0.0E+00 | 2.8E+06 |
| AT2G20990 | | synaptotagmin A | 62 kDa | 0 | 0 | 22 | 13 | 35 | | | | | | | |
| AT3G18035 | AT3G18035.1;AT3G18035.2 | winged-helix DNA-binding transcription factor family p | 52 kDa | 0 | 0 | 22 | 13 | 35 | | | | 1.4E+07 | 1.3E+07 | 4.9E+06 | 3.2E+07 |
| AT3G19770 | AT3G19770.1;AT3G19770.2 | Vacuolar sorting protein 9 (VPS9) domain | 58 kDa | 0 | 1 | 22 | 13 | 36 | | | | 4.4E+05 | 6.3E+05 | 5.5E+04 | 1.1E+06 |
| AT4G14160 | AT4G14160.2;AT4G14160.1;AT4G14160.3 | Sec23/Sec24 protein transport family protein | 85 kDa | 2 | 1 | 22 | 13 | 38 | | | | 1.2E+06 | 1.0E+06 | 8.6E+05 | 3.1E+06 |
| AT1G80750 | AT1G80750.1 | Ribosomal protein L30/L7 family protein | 28 kDa | 0 | 2 | 22 | 11 | 35 | 0.07 | 0.45 | 0.26 | 7.2E+06 | 1.1E+07 | 2.6E+06 | 2.0E+07 |
| AT4G21380 | | receptor kinase 3 | 96 kDa | 0 | 0 | 22 | 10 | 32 | | | | | | | |
| AT5G66210 | | calcium-dependent protein kinase 28 | 59 kDa | 0 | 0 | 22 | 10 | 32 | | | | | | | |
| AT4G35800 | AT4G35800.2;AT4G35800.1 | RNA polymerase II large subunit | 205 kDa | 0 | 0 | 22 | 9 | 31 | | | | 4.7E+05 | 9.4E+05 | 3.1E+04 | 1.4E+06 |
| AT5G58070 | AT5G58070.1 | temperature-induced lipocalin | 21 kDa | 10 | 11 | 22 | 6 | 49 | 0.39 | 0.19 | 0.29 | 3.8E+07 | 3.4E+07 | 1.4E+06 | 7.4E+07 |
| AT5G16970 | AT5G16970.1;AT5G16990.1 | alkenal reductase | 38 kDa | 2 | 0 | 21 | 49 | 72 | | 3.95 | 3.95 | 6.8E+06 | 5.2E+06 | 0.0E+00 | 1.2E+07 |
| AT3G10410 | AT3G10410.1 | SERINE CARBOXYPEPTIDASE-LIKE 49 | 57 kDa | 0 | 9 | 21 | 37 | 67 | | | | | | | |
| AT5G35160 | | Endomembrane protein 70 protein family | 71 kDa | 4 | 0 | 21 | 33 | 58 | | | | | | | |
| AT1G60770 | AT1G60770.1 | Tetratricopeptide repeat (TPR)-like superfamily protein | 56 kDa | 0 | 1 | 21 | 32 | 54 | | | | 4.4E+06 | 9.0E+06 | 7.5E+05 | 1.4E+07 |
| AT2G34680 | | Outer arm dynein light chain 1 protein | 187 kDa | 0 | 0 | 21 | 26 | 47 | | | | | | | |
| AT1G77380 | AT1G77380.1 | amino acid permease 3 | 52 kDa | 0 | 0 | 21 | 25 | 46 | | | | 2.9E+06 | 3.0E+06 | 0.0E+00 | 5.9E+06 |
| AT1G55610 | AT1G55610.2;AT1G55610.1 | BRI1 like | 127 kDa | 0 | 0 | 21 | 23 | 44 | | | | 2.6E+06 | 2.3E+06 | 0.0E+00 | 4.9E+06 |
| AT4G27060 | | ARM repeat superfamily protein | 94 kDa | 0 | 0 | 21 | 23 | 44 | | | | | | | |
| AT1G49300 | AT1G49300.2;AT1G49300.1 | RAB GTPase homolog G3E | 23 kDa | 3 | 9 | 21 | 21 | 54 | 0.27 | | 0.27 | 4.1E+06 | 1.2E+06 | 0.0E+00 | 5.4E+06 |
| AT4G23260 | | cysteine-rich RLK (RECEPTOR-like protein kinase) 18 | 73 kDa | 0 | 0 | 21 | 21 | 42 | | | | | | | |
| AT3G46550 | | Fasciclin-like arabinogalactan family protein | 44 kDa | 0 | 0 | 21 | 20 | 41 | | | | | | | |
| AT5G01020 | | Protein kinase superfamily protein | 46 kDa | 0 | 1 | 21 | 20 | 42 | | | | | | | |
| AT1G53440 | | Leucine-rich repeat transmembrane protein kinase | 115 kDa | 0 | 1 | 21 | 19 | 41 | | | | | | | |
| AT3G18820 | AT3G18820.1 | RAB GTPase homolog G3F | 23 kDa | 4 | 10 | 21 | 18 | 53 | | | | 6.3E+06 | 3.0E+06 | 1.4E+05 | 9.4E+06 |
| AT5G10010 | AT5G10010.1 | unknown protein; FUNCTIONS IN: molecular_function | ? | 0 | 0 | 21 | 15 | 36 | | | | 1.4E+07 | 1.7E+07 | 6.8E+05 | 3.2E+07 |
| AT1G71860 | AT1G71860.2;AT1G71860.3;AT1G71860.1 | protein tyrosine phosphatase 1 | 38 kDa | 0 | 7 | 21 | 14 | 42 | | 5.42 | 5.42 | 3.3E+06 | 2.1E+06 | 0.0E+00 | 5.4E+06 |
| AT3G24160 | | putative type 1 membrane protein | 39 kDa | 2 | 2 | 21 | 14 | 39 | 1.66 | 0.79 | 1.22 | | | | |
| AT3G42170 | AT3G42170.2;AT3G42170.1 | BED zinc finger ;hAT family dimerisation domain | 79 kDa | 0 | 0 | 21 | 14 | 35 | | | | 3.2E+06 | 3.5E+06 | 1.8E+05 | 6.8E+06 |
| AT1G18890 | | calcium-dependent protein kinase 1 | 61 kDa | 0 | 0 | 21 | 13 | 34 | | | | | | | |
| AT2G43070 | AT2G43070.1;AT2G43070.2;AT2G43070.3;AT2G43070.4 | SIGNAL PEPTIDE PEPTIDASE-LIKE 3 | 59 kDa | 0 | 0 | 21 | 13 | 34 | | 0.46 | 0.46 | 1.3E+06 | 1.0E+06 | 0.0E+00 | 2.4E+06 |
| AT3G13300 | AT3G13300.2;AT3G13300.1;AT3G13300.3 | Transducin/WD40 repeat-like superfamily protein | 146 kDa | 0 | 0 | 21 | 13 | 34 | | 0.13 | 0.13 | 1.4E+06 | 1.6E+06 | 3.5E+05 | 3.4E+06 |
| AT3G54180 | AT3G54180.1 | cyclin-dependent kinase B1;1 | 35 kDa | 1 | 9 | 21 | 13 | 44 | | | | 4.3E+06 | 1.6E+06 | 2.5E+05 | 6.2E+06 |
| AT4G28200 | AT4G28200.1 | FUNCTIONS IN: molecular_function unknown; INVOLVI | 75 kDa | 0 | 0 | 21 | 13 | 34 | | | | 1.2E+06 | 1.8E+06 | 1.2E+05 | 3.1E+06 |
| AT5G04550 | | Protein of unknown function (DUF668) | 67 kDa | 0 | 0 | 21 | 13 | 34 | | | | | | | |

| | | | | | | | | | | | | | | | |
|---|---|---|---|---|---|---|---|---|---|---|---|---|---|---|---|
| AT1G52360 | AT1G52360.3;AT1G52360.4;AT1G52360.1;AT1G52360.2;AT3G15980.5 | Coatomer, beta' subunit | 110 kDa | 0 | 0 | 21 | 12 | 33 | | 0.25 | 0.25 | 7.7E+05 | 8.6E+05 | 4.0E+04 | 1.7E+06 |
| AT3G53500 | AT3G53500.2;AT3G53500.3;AT3G53500.1 | RNA-binding (RRM/RBD/RNP motifs) family protein wit | 32 kDa | 0 | 4 | 21 | 12 | 37 | | 2.01 | 2.01 | 1.6E+07 | 1.3E+07 | 1.2E+06 | 3.0E+07 |
| AT1G05910 | AT1G05910.1 | cell division cycle protein 48-related / CDC48-related | 134 kDa | 0 | 0 | 21 | 11 | 32 | | 0.63 | 0.63 | 8.6E+04 | 0.0E+00 | 2.7E+05 | 3.5E+05 |
| AT5G05450 | AT5G05450.1 | P-loop containing nucleoside triphosphate hydrolases | 67 kDa | 0 | 0 | 21 | 10 | 31 | | | | 1.3E+06 | 1.2E+06 | 4.4E+05 | 2.9E+06 |
| AT5G06360 | AT5G06360.1 | Ribosomal protein S8e family protein | 30 kDa | 0 | 1 | 21 | 10 | 32 | | 0.23 | 0.23 | 3.7E+07 | 3.5E+07 | 2.8E+06 | 7.5E+07 |
| AT1G30240 | | FUNCTIONS IN: binding; INVOLVED IN: biological_proce | 90 kDa | 0 | 0 | 21 | 9 | 30 | | | | | | | |
| AT5G02310 | AT5G02310.2;AT5G02310.3;AT5G02310.1 | proteolysis 6 | 224 kDa | 0 | 1 | 21 | 9 | 31 | | | | 4.7E+05 | 3.7E+05 | 4.0E+04 | 8.8E+05 |
| AT3G18760 | AT3G18760.2;AT3G18760.1 | Translation elongation factor EF1B/ribosomal protein | 16 kDa | 7 | 0 | 21 | 8 | 36 | | 2.34 | 2.34 | 3.5E+07 | 6.5E+07 | 7.7E+06 | 1.1E+08 |
| AT3G48560 | AT3G48560.1 | chlorsulfuron/imidazolinone resistant 1 | 73 kDa | 0 | 0 | 21 | 8 | 29 | | | | 1.6E+07 | 2.0E+07 | 4.1E+04 | 3.6E+07 |
| AT4G23190 | | cysteine-rich RLK (RECEPTOR-like protein kinase) 11 | 74 kDa | 0 | 0 | 21 | 8 | 29 | | | | | | | |
| AT3G13160 | AT3G13160.1 | Tetratricopeptide repeat (TPR)-like superfamily protein | 45 kDa | 1 | 18 | 21 | 6 | 46 | | | | 3.0E+07 | 3.4E+07 | 5.7E+06 | 7.0E+07 |
| AT5G46860 | AT5G46860.1 | Syntaxin/t-SNARE family protein | 29 kDa | 6 | 13 | 20 | 56 | 95 | | | | 1.9E+07 | 1.5E+07 | 9.7E+05 | 3.5E+07 |
| AT3G09820 | AT3G09820.1;AT3G09820.2 | adenosine kinase 1 | 38 kDa | 4 | 2 | 20 | 41 | 67 | | | | 9.6E+05 | 7.7E+05 | 0.0E+00 | 1.7E+06 |
| AT2G45010 | AT2G45010.1 | PLAC8 family protein | 27 kDa | 5 | 19 | 20 | 39 | 83 | | 6.16 | 6.16 | 1.5E+07 | 8.5E+06 | 4.3E+06 | 2.7E+07 |
| AT3G56090 | AT3G56090.1 | ferritin 3 | 29 kDa | 8 | 0 | 20 | 32 | 60 | 0.51 | 1.29 | 0.90 | 2.0E+08 | 1.1E+08 | 1.7E+06 | 3.2E+08 |
| AT1G06950 | AT1G06950.1 | translocon at the inner envelope membrane of chlorop | 112 kDa | 1 | 0 | 20 | 30 | 51 | 0.06 | 0.16 | 0.11 | 2.5E+06 | 3.0E+06 | 4.1E+04 | 5.5E+06 |
| AT5G65430 | AT5G65430.2;AT5G65430.1;AT5G65430.3 | general regulatory factor 8 | 28 kDa | 0 | 8 | 20 | 25 | 53 | | | | 6.1E+05 | 5.4E+05 | 0.0E+00 | 1.1E+06 |
| AT2G03510 | AT2G03510.1 | SPFH/Band 7/PHB domain-containing membrane-assoc | 41 kDa | 2 | 1 | 20 | 24 | 47 | 0.06 | 0.22 | 0.14 | | | | |
| AT4G31170 | | Protein kinase superfamily protein | 46 kDa | 0 | 0 | 20 | 23 | 43 | | | | | | | |
| AT3G03610 | | ELMO/CED-12 family protein | 37 kDa | 7 | 15 | 20 | 21 | 63 | | 7.26 | 7.26 | | | | |
| AT4G30440 | | UDP-D-glucuronate 4-epimerase 1 | 47 kDa | 0 | 0 | 20 | 21 | 41 | | | | | | | |
| AT4G01330 | | Protein kinase superfamily protein | 54 kDa | 0 | 0 | 20 | 20 | 40 | | | | | | | |
| AT5G25754 | | RNA polymerase I-associated factor PAF67 | 60 kDa | 1 | 2 | 20 | 20 | 43 | | 1.10 | 1.10 | | | | |
| AT3G23820 | | UDP-D-glucuronate 4-epimerase 6 | 51 kDa | 0 | 2 | 20 | 19 | 41 | | | | | | | |
| AT4G15780 | AT4G15780.1 | vesicle-associated membrane protein 724 | 25 kDa | 5 | 8 | 20 | 19 | 52 | | | | | | | |
| AT4G17170 | | RAB GTPase homolog B1C | 23 kDa | 8 | 14 | 20 | 19 | 61 | | | | | | | |
| AT3G25140 | | Nucleotide-diphospho-sugar transferases superfamily p | 64 kDa | 0 | 0 | 20 | 18 | 38 | | | | | | | |
| AT4G02840 | AT4G02840.1;AT4G02840.2 | Small nuclear ribonucleoprotein family protein | 13 kDa | 5 | 2 | 20 | 18 | 45 | | 0.30 | 0.30 | 6.6E+07 | 7.1E+07 | 2.7E+06 | 1.4E+08 |
| AT5G63120 | AT5G63120.2;AT5G63120.5;AT5G63120.4;AT5G63120.3;AT5G63120.1 | P-loop containing nucleoside triphosphate hydrolases | 65 kDa | 1 | 3 | 20 | 18 | 42 | | 0.34 | 0.34 | 1.6E+06 | 2.0E+06 | 4.2E+05 | 4.1E+06 |
| AT1G10840 | AT1G10840.1;AT1G10840.2 | translation initiation factor 3 subunit H1 | 38 kDa | 0 | 10 | 20 | 17 | 47 | | 0.85 | 0.85 | 4.9E+06 | 9.3E+06 | 3.8E+06 | 1.8E+07 |
| AT2G24230 | | Leucine-rich repeat protein kinase family protein | 94 kDa | 0 | 1 | 20 | 17 | 38 | | | | | | | |
| AT2G38020 | | vacuoleless1 (VCL1) | 97 kDa | 0 | 0 | 20 | 17 | 37 | | | | | | | |
| AT5G61530 | | small G protein family protein / RhoGAP family protein | 42 kDa | 0 | 15 | 20 | 17 | 52 | | | | | | | |
| AT1G16860 | | Ubiquitin-specific protease family C19-related protein | 51 kDa | 0 | 0 | 20 | 16 | 36 | | | | | | | |
| AT3G11710 | AT3G11710.1 | lysyl-tRNA synthetase 1 | 71 kDa | 0 | 0 | 20 | 16 | 36 | | | | 3.6E+06 | 7.3E+06 | 7.9E+04 | 1.1E+07 |
| AT4G32300 | | S-domain-2 5 | 90 kDa | 0 | 0 | 20 | 16 | 36 | | | | | | | |
| AT5G66200 | | armadillo repeat only 2 | 71 kDa | 1 | 1 | 20 | 16 | 38 | | | | | | | |
| AT1G18390 | AT1G18390.1;AT1G18390.2 | Protein kinase superfamily protein | ? | 0 | 1 | 20 | 15 | 36 | | | | 0.0E+00 | 5.2E+04 | 0.0E+00 | 5.2E+04 |
| AT1G72160 | | Sec14p-like phosphatidylinositol transfer family protei | 56 kDa | 0 | 0 | 20 | 15 | 35 | | | | | | | |
| AT4G03390 | AT4G03390.2;AT4G03390.1 | STRUBBELIG-receptor family 3 | 85 kDa | 0 | 0 | 20 | 15 | 35 | | | | 5.4E+05 | 6.3E+05 | 2.4E+04 | 1.2E+06 |
| AT5G10730 | AT5G10730.1 | NAD(P)-binding Rossmann-fold superfamily protein | 31 kDa | 5 | 9 | 20 | 15 | 49 | | 0.89 | 0.89 | 1.1E+07 | 1.7E+07 | 7.1E+05 | 2.9E+07 |
| AT5G61910 | | DCD (Development and Cell Death) domain protein | 147 kDa | 1 | 0 | 20 | 15 | 36 | | | | | | | |
| AT5G64070 | | phosphatidylinositol 4-OH kinase beta1 | 126 kDa | 0 | 0 | 20 | 15 | 35 | | | | | | | |
| AT1G04690 | AT1G04690.1 | potassium channel beta subunit 1 | 37 kDa | 0 | 2 | 20 | 14 | 36 | | 0.18 | 0.18 | 6.1E+06 | 2.2E+06 | 1.5E+06 | 9.8E+06 |
| AT3G12670 | AT3G12670.1 | CTP synthase family protein | 64 kDa | 0 | 0 | 20 | 14 | 34 | | | | 3.9E+06 | 3.8E+06 | 5.8E+05 | 8.3E+06 |
| AT5G41600 | | VIRB2-interacting protein 3 | 29 kDa | 7 | 0 | 20 | 14 | 41 | | | | | | | |
| AT5G52040 | AT5G52040.1;AT5G52040.2;AT5G52040.7;AT5G52040.5;AT5G52040.6;AT5G52040.4;AT5G52040.3;AT4G25500.4;AT4G25500.1 | RNA-binding (RRM/RBD/RNP motifs) family protein | 41 kDa | 0 | 0 | 20 | 14 | 34 | | 1.26 | 1.26 | 1.5E+07 | 2.2E+07 | 1.4E+06 | 3.8E+07 |
| AT4G34220 | AT4G34220.1 | Leucine-rich repeat protein kinase family protein | 83 kDa | 0 | 0 | 20 | 13 | 33 | | | | | | | |
| AT5G06220 | AT5G06220.2;AT5G06220.3;AT5G06220.1 | LETM1-like protein | 102 kDa | 0 | 0 | 20 | 13 | 33 | | | | | | | |
| AT1G26120 | | alpha/beta-Hydrolases superfamily protein | 53 kDa | 2 | 1 | 20 | 12 | 35 | | | | | | | |
| AT1G34750 | | Protein phosphatase 2C family protein | 31 kDa | 4 | 4 | 20 | 12 | 40 | | 3.64 | 3.64 | | | | |
| AT1G65030 | AT1G65030.1 | Transducin/WD40 repeat-like superfamily protein | 37 kDa | 0 | 2 | 20 | 12 | 34 | | 0.21 | 0.21 | 5.7E+06 | 6.3E+06 | 1.5E+06 | 1.3E+07 |
| AT4G26870 | AT4G26870.1 | Class II aminoacyl-tRNA and biotin synthetases superfar | 60 kDa | 0 | 0 | 20 | 12 | 32 | | | | 2.1E+06 | 1.2E+06 | 5.1E+04 | 3.3E+06 |
| AT3G63130 | AT3G63130.2;AT3G63130.1 | RAN GTPase activating protein 1 | 59 kDa | 0 | 0 | 20 | 11 | 31 | | | | | | | |
| AT4G29410 | AT4G29410.2;AT4G29410.1 | Ribosomal L28e protein family | 16 kDa | 2 | 0 | 20 | 11 | 33 | | 0.71 | 0.71 | 2.6E+07 | 3.5E+07 | 3.0E+06 | 6.5E+07 |
| AT5G35790 | AT5G35790.1 | glucose-6-phosphate dehydrogenase 1 | 65 kDa | 1 | 0 | 20 | 11 | 32 | | | | 7.1E+05 | 5.0E+05 | 0.0E+00 | 1.2E+06 |
| AT3G19100 | | Protein kinase superfamily protein | 67 kDa | 0 | 0 | 20 | 10 | 30 | | | | | | | |
| AT1G15420 | AT1G15420.2;AT1G15420.1 | CONTAINS InterPro DOMAIN/s: Small-subunit processo | 30 kDa | 2 | 0 | 20 | 9 | 31 | 0.06 | 0.65 | 0.35 | 3.4E+06 | 3.5E+06 | 6.1E+06 | 1.3E+07 |
| AT3G26360 | AT3G26360.2;AT3G26360.1 | Ribosomal protein S21 family protein | 12 kDa | 0 | 0 | 20 | 9 | 29 | | | | 2.5E+07 | 2.8E+07 | 2.8E+06 | 5.6E+07 |
| AT2G25760 | | Protein kinase family protein | 76 kDa | 0 | 0 | 20 | 8 | 28 | | | | | | | |
| AT1G08130 | AT1G08130.1;AT1G49250.1 | DNA ligase 1 | 88 kDa | 1 | 0 | 20 | 7 | 28 | | 0.38 | 0.38 | 1.2E+06 | 9.1E+05 | 1.3E+05 | 2.2E+06 |
| AT3G02320 | AT3G02320.1 | N2,N2-dimethylguanosine tRNA methyltransferase | 66 kDa | 0 | 0 | 20 | 7 | 27 | | | | 3.6E+06 | 3.8E+06 | 6.8E+04 | 7.5E+06 |
| AT3G03600 | AT3G03600.1 | ribosomal protein S2 | 25 kDa | 0 | 6 | 20 | 5 | 31 | | | | 1.3E+07 | 1.7E+07 | 3.2E+06 | 3.3E+07 |
| AT3G08930 | AT3G08930.1 | LMBR1-like membrane protein | 57 kDa | 4 | 0 | 20 | 5 | 29 | | | | 2.2E+06 | 2.0E+06 | 0.0E+00 | 4.2E+06 |
| AT3G56760 | | Protein kinase superfamily protein | 65 kDa | 0 | 0 | 20 | 5 | 25 | | | | | | | |
| AT4G13940 | AT4G13940.2;AT4G13940.3;AT4G13940.1;AT4G13940.4;AT3G23810.1 | S-adenosyl-L-homocysteine hydrolase | 53 kDa | 6 | 2 | 19 | 45 | 72 | | | | 7.5E+06 | 2.7E+06 | 0.0E+00 | 1.0E+07 |
| AT1G35720 | AT1G35720.1 | annexin 1 | 36 kDa | 0 | 5 | 19 | 28 | 52 | | 5.49 | 5.49 | 6.8E+06 | 2.0E+06 | 0.0E+00 | 8.8E+06 |
| AT5G47940 | | unknown protein; Has 1807 Blast hits to 1807 proteins | 82 kDa | 8 | 6 | 19 | 21 | 54 | | | | | | | |
| AT1G52930 | AT1G52930.1 | Ribosomal RNA processing Brix domain protein | 37 kDa | 0 | 0 | 19 | 20 | 39 | | 0.18 | 0.18 | 2.1E+06 | 2.2E+06 | 1.9E+06 | 6.2E+06 |
| AT3G07080 | | EamA-like transporter family | 48 kDa | 0 | 0 | 19 | 20 | 39 | | | | | | | |
| AT1G80510 | | Transmembrane amino acid transporter family protein | 53 kDa | 0 | 0 | 19 | 19 | 38 | | | | | | | |
| AT2G17220 | | Protein kinase superfamily protein | 46 kDa | 0 | 1 | 19 | 19 | 39 | | | | | | | |
| AT4G02570 | AT4G02570.4;AT4G02570.3;AT4G02570.2;AT4G02570.1 | cullin 1 | 86 kDa | 0 | 0 | 19 | 19 | 38 | | | | 2.4E+06 | 9.9E+05 | 4.3E+04 | 3.4E+06 |
| AT5G14610 | | DEAD box RNA helicase family protein | 76 kDa | 0 | 1 | 19 | 19 | 39 | | 0.19 | 0.19 | | | | |
| AT5G47880 | AT5G47880.2;AT5G47880.1 | eukaryotic release factor 1-1 | 49 kDa | 0 | 3 | 19 | 18 | 40 | | | | 2.6E+06 | 3.3E+06 | 4.1E+05 | 6.3E+06 |
| AT5G59980 | AT5G59980.1;AT5G59980.3;AT5G59980.4;AT5G59980.2 | Polymerase/histidinol phosphatase-like | 78 kDa | 0 | 0 | 19 | 17 | 36 | | | | 3.2E+05 | 3.1E+05 | 2.0E+05 | 8.3E+05 |
| AT1G67320 | AT1G67320.1;AT1G67320.2;AT1G67320.3 | DNA primase, large subunit family | 52 kDa | 0 | 0 | 19 | 16 | 35 | | | | 2.1E+05 | 2.9E+05 | 3.2E+04 | 5.4E+05 |
| AT1G16670 | | Protein kinase superfamily protein | 43 kDa | 0 | 2 | 19 | 15 | 36 | | | | | | | |
| AT2G06990 | AT2G06990.1 | RNA helicase, ATP-dependent, SK12/DOB1 protein | 112 kDa | 0 | 0 | 19 | 15 | 34 | 0.76 | | 0.76 | 2.2E+05 | 1.3E+05 | 5.4E+04 | 4.0E+05 |
| AT3G05940 | | Protein of unknown function (DUF300) | 47 kDa | 0 | 0 | 19 | 15 | 34 | | | | | | | |
| AT4G01037 | AT4G01037.1 | Ubiquitin carboxyl-terminal hydrolase family protein | 61 kDa | 0 | 0 | 19 | 15 | 34 | | | | 3.5E+05 | 4.4E+05 | 0.0E+00 | 7.9E+05 |

| | | | | | | | | | | | | | | | |
|---|---|---|---|---|---|---|---|---|---|---|---|---|---|---|---|
| AT5G04930 | | aminophospholipid ATPase 1 | 130 kDa | 0 | 0 | 19 | 15 | 34 | | | | | | | |
| AT5G40490 | AT5G40490.1;AT5G40490.2 | RNA-binding (RRM/RBD/RNP motifs) family protein | 42 kDa | 0 | 0 | 19 | 15 | 34 | | | | 4.2E+07 | 2.9E+07 | 7.3E+06 | 7.8E+07 |
| AT5G62710 | | Leucine-rich repeat protein kinase family protein | 67 kDa | 0 | 1 | 19 | 15 | 35 | | | | | | | |
| AT2G28890 | | poltergeist like 4 | 72 kDa | 0 | 0 | 19 | 14 | 33 | | | | | | | |
| AT4G12120 | AT4G12120.1 | Sec1/munc18-like (SM) proteins superfamily | 75 kDa | 5 | 3 | 19 | 13 | 40 | | | | 4.8E+05 | 7.1E+05 | 0.0E+00 | 1.2E+06 |
| AT4G39980 | AT4G39980.1 | 3-deoxy-D-arabino-heptulosonate 7-phosphate syntha | 58 kDa | 0 | 3 | 19 | 13 | 35 | | 0.25 | 0.25 | 7.6E+06 | 5.7E+06 | 0.0E+00 | 1.3E+07 |
| AT5G20890 | AT5G20890.1 | TCP-1/cpn60 chaperonin family protein | 57 kDa | 1 | 1 | 19 | 13 | 34 | | | | 9.9E+05 | 1.1E+06 | 5.6E+04 | 2.1E+06 |
| AT5G48800 | | Phototropic-responsive NPH3 family protein | 70 kDa | 0 | 0 | 19 | 13 | 32 | | | | | | | |
| AT1G65220 | AT1G65220.1 | ARM repeat superfamily protein | 47 kDa | 0 | 2 | 19 | 12 | 33 | | | | 3.3E+06 | 3.8E+06 | 1.7E+05 | 7.2E+06 |
| AT4G21450 | AT4G21450.1;AT4G21450.2;AT4G21450.3 | PapD-like superfamily protein | 33 kDa | 2 | 3 | 19 | 12 | 36 | | | | 3.9E+06 | 1.8E+06 | 0.0E+00 | 5.7E+06 |
| AT4G20110 | | VACUOLAR SORTING RECEPTOR 7 | 70 kDa | 0 | 0 | 19 | 11 | 30 | | | | | | | |
| AT4G35470 | | plant intracellular ras group-related LRR 4 | 61 kDa | 0 | 0 | 19 | 11 | 30 | | | | | | | |
| AT4G38580 | AT4G38580.2;AT4G38580.1 | farnesylated protein 6 | 17 kDa | 13 | 0 | 19 | 11 | 43 | | 5.29 | 5.29 | 4.6E+07 | 3.4E+06 | 2.6E+05 | 5.0E+07 |
| AT5G66420 | AT5G66420.3;AT5G66420.1;AT5G66420.2 | LOCATED IN: cellular_component unknown; EXPRESSEI | 81 kDa | 0 | 0 | 19 | 11 | 30 | | | | 9.2E+05 | 3.1E+05 | 0.0E+00 | 1.2E+06 |
| AT1G04710 | AT1G04710.1 | peroxisomal 3-ketoacyl-CoA thiolase 4 | 47 kDa | 0 | 0 | 19 | 10 | 29 | | | | 1.4E+06 | 1.2E+06 | 2.2E+05 | 2.8E+06 |
| AT1G13320 | | protein phosphatase 2A subunit A3 | 66 kDa | 1 | 1 | 19 | 10 | 31 | | | | | | | |
| AT1G16350 | AT1G16350.1 | Aldolase-type TIM barrel family protein | 54 kDa | 1 | 0 | 19 | 10 | 30 | | 0.36 | 0.36 | 4.3E+07 | 2.5E+07 | 0.0E+00 | 6.8E+07 |
| AT1G48650 | | DEA(D/H)-box RNA helicase family protein | 134 kDa | 0 | 0 | 19 | 10 | 29 | | | | | | | |
| AT2G31060 | AT2G31060.1;AT2G31060.2;AT2G31060.3 | elongation factor family protein | 73 kDa | 0 | 0 | 19 | 10 | 29 | | 0.29 | 0.29 | 1.4E+06 | 1.8E+06 | 1.2E+06 | 4.3E+06 |
| AT1G49580 | | Calcium-dependent protein kinase (CDPK) family prote | 68 kDa | 0 | 0 | 19 | 9 | 28 | | | | | | | |
| AT3G10530 | AT3G10530.1 | Transducin/WD40 repeat-like superfamily protein | 60 kDa | 0 | 0 | 19 | 9 | 28 | | | | 1.8E+05 | 2.6E+06 | 2.9E+05 | 3.1E+06 |
| AT4G28600 | | no pollen germination related 2 | 83 kDa | 0 | 0 | 19 | 9 | 28 | | | | | | | |
| AT5G03660 | AT5G03660.2;AT5G03660.1;AT5G03660.3 | Family of unknown function (DUF662) | 20 kDa | 4 | 2 | 19 | 9 | 34 | | | | 4.1E+06 | 2.8E+06 | 0.0E+00 | 6.9E+06 |
| AT4G05410 | AT4G05410.1 | Transducin/WD40 repeat-like superfamily protein | 57 kDa | 0 | 0 | 19 | 8 | 27 | | 0.32 | 0.32 | 6.0E+06 | 9.6E+06 | 1.3E+06 | 1.7E+07 |
| AT4G11600 | AT4G11600.1 | glutathione peroxidase 6 | 26 kDa | 3 | 6 | 19 | 8 | 36 | | 0.62 | 0.62 | 6.1E+06 | 1.3E+06 | 0.0E+00 | 7.4E+06 |
| AT1G59610 | | dynamin-like 3 | 100 kDa | 0 | 1 | 19 | 7 | 27 | | | | | | | |
| AT4G29430 | AT4G29430.1 | ribosomal protein S15A E | 15 kDa | 0 | 0 | 19 | 3 | 22 | | 0.24 | 0.24 | 5.2E+07 | 1.0E+08 | 5.6E+06 | 1.6E+08 |
| AT3G23390 | AT3G23390.1;AT4G14320.1;AT4G14320.2 | Zinc-binding ribosomal protein family protein | 12 kDa | 0 | 0 | 19 | 1 | 20 | | | | 6.0E+07 | 2.3E+07 | 4.9E+07 | 1.3E+08 |
| AT5G53870 | AT5G53870.2;AT5G53870.1 | early nodulin-like protein 1 | 38 kDa | 0 | 0 | 18 | 77 | 95 | | | | 6.1E+06 | 3.8E+06 | 2.0E+05 | 1.0E+07 |
| AT1G65720 | | unknown protein; FUNCTIONS IN: molecular_function | 20 kDa | 4 | 0 | 18 | 38 | 60 | | | | | | | |
| AT3G51580 | AT3G51580.1;AT3G51580.2 | unknown protein; FUNCTIONS IN: molecular_function | 42 kDa | 5 | 8 | 18 | 33 | 64 | | | | 3.2E+05 | 1.9E+05 | 0.0E+00 | 5.1E+05 |
| AT1G23190 | AT1G23190.1 | Phosphoglucomutase/phosphomannomutase family p | 63 kDa | 0 | 0 | 18 | 27 | 45 | | | | 3.4E+06 | 3.1E+06 | 0.0E+00 | 6.5E+06 |
| AT1G65240 | AT1G65240.1 | Eukaryotic aspartyl protease family protein | 52 kDa | 2 | 6 | 18 | 25 | 51 | | 4.67 | 4.67 | 3.3E+06 | 2.3E+06 | 0.0E+00 | 5.5E+06 |
| AT5G10260 | | RAB GTPase homolog H1E | 23 kDa | 0 | 0 | 18 | 24 | 42 | | | | | | | |
| AT1G11580 | | methylesterase PCR A | 62 kDa | 0 | 0 | 18 | 23 | 41 | | | | | | | |
| AT3G16100 | | RAB GTPase homolog G3C | 23 kDa | 0 | 13 | 18 | 23 | 54 | | | | | | | |
| AT3G21540 | AT3G21540.1 | transducin family protein / WD-40 repeat family protei | 106 kDa | 0 | 0 | 18 | 22 | 40 | | | | 2.0E+06 | 2.6E+06 | 3.5E+05 | 4.9E+06 |
| AT4G25730 | AT4G25730.1 | FtsJ-like methyltransferase family protein | 92 kDa | 0 | 0 | 18 | 22 | 40 | | 0.04 | 0.04 | 0.0E+00 | 6.7E+04 | 3.5E+05 | 4.2E+05 |
| AT2G34560 | AT2G34560.1;AT2G34560.2 | P-loop containing nucleoside triphosphate hydrolases | 44 kDa | 0 | 0 | 18 | 21 | 39 | | | | 0.0E+00 | 0.0E+00 | 4.7E+05 | 4.7E+05 |
| AT5G11680 | AT5G11680.1 | FUNCTIONS IN: molecular_function unknown; INVOLVI | 23 kDa | 6 | 19 | 18 | 21 | 64 | | 5.77 | 5.77 | 7.9E+06 | 4.8E+06 | 7.0E+04 | 1.3E+07 |
| AT3G53420 | AT3G53420.2;AT3G53420.1;AT2G37180.1;AT2G37170.1;AT2G37170.2 | plasma membrane intrinsic protein 2A | 30 kDa | 0 | 1 | 18 | 20 | 39 | | | | 1.5E+06 | 9.2E+05 | 0.0E+00 | 2.4E+06 |
| AT5G11520 | AT5G11520.1 | aspartate aminotransferase 3 | 49 kDa | 5 | 4 | 18 | 20 | 47 | | | | 1.8E+06 | 3.0E+06 | 1.5E+05 | 5.0E+06 |
| AT1G63000 | AT1G63000.1 | nucleotide-rhamnose synthase/epimerase-reductase | 34 kDa | 0 | 3 | 18 | 19 | 40 | | | | 1.3E+06 | 2.8E+06 | 0.0E+00 | 4.1E+06 |
| AT3G13470 | AT3G13470.1;AT1G55490.5;AT1G55490.4;AT1G55490.3;AT1G55490.2;AT1G55490.1 | TCP-1/cpn60 chaperonin family protein | 63 kDa | 0 | 1 | 18 | 19 | 38 | | | | 2.8E+05 | 6.5E+05 | 2.8E+05 | 1.2E+06 |
| AT3G55740 | | proline transporter 2 | 48 kDa | 1 | 0 | 18 | 19 | 38 | | | | | | | |
| AT2G37760 | AT2G37760.7;AT2G37760.6;AT2G37760.4;AT2G37760.3;AT2G37760.5;AT2G37760.2;AT2G37760.1 | NAD(P)-linked oxidoreductase superfamily protein | 35 kDa | 2 | 9 | 18 | 18 | 47 | | | | 1.5E+06 | 1.6E+05 | 0.0E+00 | 1.7E+06 |
| AT2G33730 | AT2G33730.1 | P-loop containing nucleoside triphosphate hydrolases | 85 kDa | 0 | 0 | 18 | 17 | 35 | | | | 1.6E+06 | 1.9E+06 | 3.2E+05 | 3.8E+06 |
| AT4G35650 | | isocitrate dehydrogenase III | 40 kDa | 0 | 7 | 18 | 17 | 42 | | 0.93 | 0.93 | | | | |
| AT5G42980 | AT5G42980.1 | thioredoxin 3 | 13 kDa | 3 | 0 | 18 | 17 | 38 | | 2.09 | 2.09 | 5.1E+07 | 2.0E+07 | 5.1E+06 | 7.6E+07 |
| AT5G56900 | AT5G56900.2;AT5G56900.3 | CwfJ-like family protein / zinc finger (CCCH-type) family | 66 kDa | 0 | 0 | 18 | 17 | 35 | | | | 1.2E+06 | 9.0E+05 | 1.7E+05 | 2.2E+06 |
| AT3G01810 | AT3G01810.3;AT3G01810.2;AT3G01810.1 | FUNCTIONS IN: molecular_function unknown; INVOLVI | 102 kDa | 1 | 0 | 18 | 16 | 35 | | | | 1.5E+06 | 8.1E+05 | 0.0E+00 | 2.3E+06 |
| AT3G47670 | | Plant invertase/pectin methylesterase inhibitor superfa | 30 kDa | 5 | 15 | 18 | 15 | 53 | 1.21 | 8.30 | 4.76 | | | | |
| AT4G02480 | AT4G02480.1;AT1G02890.1;AT1G02890.2 | AAA-type ATPase family protein | 137 kDa | 0 | 0 | 18 | 15 | 33 | | | | 4.8E+05 | 8.0E+05 | 0.0E+00 | 1.3E+06 |
| AT5G47070 | | Protein kinase superfamily protein | 47 kDa | 0 | 2 | 18 | 15 | 35 | | | | | | | |
| AT3G17970 | | translocon at the outer membrane of chloroplasts 64-II | 64 kDa | 0 | 0 | 18 | 13 | 31 | | | | | | | |
| AT3G47570 | | Leucine-rich repeat protein kinase family protein | 111 kDa | 1 | 1 | 18 | 13 | 33 | | | | | | | |
| AT4G21120 | | amino acid transporter 1 | 65 kDa | 2 | 0 | 18 | 13 | 33 | | | | | | | |
| AT2G37710 | | receptor lectin kinase | 76 kDa | 0 | 0 | 18 | 12 | 30 | | | | | | | |
| AT3G01560 | AT3G01560.1 | Protein of unknown function (DUF1421) | 55 kDa | 0 | 4 | 18 | 12 | 34 | | | | 9.8E+05 | 8.9E+05 | 0.0E+00 | 1.9E+06 |
| AT3G47520 | AT3G47520.1 | malate dehydrogenase | 42 kDa | 4 | 1 | 18 | 12 | 35 | 0.09 | 0.16 | 0.13 | 9.5E+06 | 1.3E+07 | 4.6E+05 | 2.3E+07 |
| AT3G48860 | AT3G48860.3;AT3G48860.2;AT3G48860.1 | AT3G48860.2 | ? | 4 | 1 | 18 | 12 | 35 | | | | 1.0E+06 | 8.3E+05 | 0.0E+00 | 1.8E+06 |
| AT2G29580 | | CCCH-type zinc fingerfamily protein with RNA-binding | 54 kDa | 0 | 0 | 18 | 11 | 29 | | 0.35 | 0.35 | | | | |
| AT5G25590 | | Protein of unknown function (DUF630 and DUF632) | ? | 0 | 0 | 18 | 11 | 29 | | | | | | | |
| AT1G03050 | | ENTH/ANTH/VHS superfamily protein | 67 kDa | 0 | 0 | 18 | 10 | 28 | | | | | | | |
| AT4G24290 | | MAC/Perforin domain-containing protein | 68 kDa | 0 | 0 | 18 | 10 | 28 | | | | | | | |
| AT5G08180 | AT5G08180.2;AT5G08180.1 | Ribosomal protein L7Ae/L30e/S12e/Gadd45 family prc | 17 kDa | 0 | 0 | 18 | 10 | 28 | | 0.16 | 0.16 | 9.8E+07 | 7.5E+07 | 3.6E+07 | 2.1E+08 |
| AT5G47540 | | Mo25 family protein | 39 kDa | 0 | 0 | 18 | 10 | 28 | | | | | | | |
| AT1G20780 | | senescence-associated E3 ubiquitin ligase 1 | 88 kDa | 0 | 2 | 18 | 9 | 29 | | | | | | | |
| AT4G20980 | AT4G20980.4;AT4G20980.3;AT4G20980.2;AT4G20980.1 | Eukaryotic translation initiation factor 3 subunit 7 (eIF | 67 kDa | 0 | 0 | 18 | 9 | 27 | 0.10 | | 0.10 | 2.9E+06 | 1.3E+06 | 2.8E+06 | 7.0E+06 |
| AT4G23630 | AT4G23630.1;AT4G23630.2 | VIRB2-interacting protein 1 | 31 kDa | 3 | 6 | 18 | 9 | 36 | 0.28 | | 0.28 | 0.0E+00 | 2.5E+06 | 0.0E+00 | 2.5E+06 |
| AT5G09400 | | K+ uptake permease 7 | 95 kDa | 0 | 0 | 18 | 9 | 27 | | | | | | | |
| AT5G13650 | AT5G13650.1;AT5G13650.2 | elongation factor family protein | 74 kDa | 0 | 0 | 18 | 9 | 27 | | 0.08 | 0.08 | 2.4E+06 | 1.1E+06 | 2.9E+05 | 3.7E+06 |
| AT5G49990 | | Xanthine/uracil permease family protein | 57 kDa | 0 | 0 | 18 | 9 | 27 | | | | | | | |
| AT4G03430 | AT4G03430.3;AT4G03430.2;AT4G03430.1 | pre-mRNA splicing factor-related | 116 kDa | 0 | 0 | 18 | 8 | 26 | 0.25 | | 0.25 | 2.7E+06 | 2.5E+06 | 2.9E+05 | 5.4E+06 |
| AT1G10290 | | dynamin-like protein 6 | 99 kDa | 0 | 0 | 18 | 7 | 25 | | | | | | | |
| AT2G16600 | AT2G16600.1;AT2G16600.2 | rotamase CYP 3 | 18 kDa | 4 | 0 | 18 | 7 | 29 | | | | 4.4E+07 | 4.0E+07 | 0.0E+00 | 8.4E+07 |
| AT3G61130 | | galacturonosyltransferase 1 | 77 kDa | 0 | 0 | 18 | 7 | 25 | | | | | | | |
| AT2G44860 | AT2G44860.2;AT2G44860.1 | Ribosomal protein L24e family protein | 19 kDa | 0 | 0 | 18 | 6 | 24 | | | | 4.9E+07 | 5.5E+07 | 2.4E+07 | 1.3E+08 |
| AT3G03940 | | Protein kinase family protein | 78 kDa | 0 | 0 | 18 | 6 | 24 | | | | | | | |
| AT3G58100 | AT3G58100.1;AT3G58100.2 | plasmodesmata callose-binding protein 5 | 19 kDa | 6 | 0 | 18 | 6 | 30 | 13.01 | 3.42 | 8.21 | 1.7E+07 | 1.9E+07 | 2.0E+06 | 3.8E+07 |
| AT5G64670 | AT5G64670.1 | Ribosomal protein L18e/L15 superfamily protein | 31 kDa | 0 | 4 | 18 | 6 | 28 | | | | 1.7E+07 | 3.6E+07 | 4.2E+06 | 5.8E+07 |
| AT5G44710 | AT5G44710.1 | CONTAINS InterPro DOMAIN/s: Ribosomal protein S27/ | 12 kDa | 3 | 0 | 18 | 4 | 25 | | | | 9.6E+07 | 1.3E+08 | 4.6E+06 | 2.3E+08 |

| | | | | | | | | | | | | | | | |
|---|---|---|---|---|---|---|---|---|---|---|---|---|---|---|---|
| AT5G56940 | AT5G56940.1 | Ribosomal protein S16 family protein | 15 kDa | 6 | 0 | 18 | 4 | 28 | | | | 3.5E+07 | 5.0E+07 | 1.3E+07 | 9.7E+07 |
| AT3G52880 | AT3G52880.1;AT3G52880.2 | monodehydroascorbate reductase 1 | 46 kDa | 1 | 0 | 17 | 47 | 65 | | | | 5.7E+06 | 2.9E+06 | 0.0E+00 | 8.6E+06 |
| AT1G65260 | AT1G65260.2;AT1G65260.1 | plastid transcriptionally active 4 | 36 kDa | 3 | 4 | 17 | 42 | 66 | 0.28 | 0.96 | 0.62 | 6.7E+07 | 5.8E+07 | 1.1E+06 | 1.3E+08 |
| AT1G25260 | AT1G25260.1 | Ribosomal protein L10 family protein | 27 kDa | 0 | 0 | 17 | 29 | 46 | | 0.89 | 0.89 | 3.3E+07 | 2.9E+07 | 9.1E+06 | 7.1E+07 |
| AT2G15690 | AT2G15690.1 | Tetratricopeptide repeat (TPR)-like superfamily protein | 66 kDa | 0 | 0 | 17 | 28 | 45 | | | | 1.0E+07 | 1.3E+07 | 2.2E+06 | 2.5E+07 |
| AT2G44100 | AT2G44100.2;AT2G44100.1 | guanosine nucleotide diphosphate dissociation inhibit | 50 kDa | 7 | 1 | 17 | 28 | 53 | | | | 2.8E+06 | 1.9E+06 | 0.0E+00 | 4.7E+06 |
| AT1G47128 | | Granulin repeat cysteine protease family protein | 51 kDa | 0 | 0 | 17 | 24 | 41 | | | | | | | |
| AT1G07750 | AT1G07750.1 | RmlC-like cupins superfamily protein | 38 kDa | 2 | 5 | 17 | 22 | 46 | | 5.71 | 5.71 | 1.5E+06 | 1.3E+06 | 0.0E+00 | 2.8E+06 |
| AT2G07180 | | Protein kinase superfamily protein | 49 kDa | 0 | 1 | 17 | 22 | 40 | | | | | | | |
| AT3G55440 | AT3G55440.1 | triosephosphate isomerase | 27 kDa | 14 | 20 | 17 | 22 | 73 | 3.24 | 13.48 | 8.36 | 2.5E+07 | 1.6E+07 | 0.0E+00 | 4.1E+07 |
| AT2G42790 | AT2G42790.1 | citrate synthase 3 | 56 kDa | 0 | 13 | 17 | 21 | 51 | | 0.42 | 0.42 | 1.4E+07 | 1.2E+07 | 3.1E+06 | 2.9E+07 |
| AT3G07410 | AT3G07410.1;AT5G47520.1 | RAB GTPase homolog A5B | 24 kDa | 10 | 13 | 17 | 19 | 59 | | | | 1.3E+06 | 1.0E+06 | 0.0E+00 | 2.4E+06 |
| AT4G14350 | | AGC (cAMP-dependent, cGMP-dependent and protein k | 63 kDa | 0 | 0 | 17 | 19 | 36 | | | | | | | |
| AT2G01750 | AT2G01750.1;AT2G01750.2;AT1G14840.2;AT1G14840.1 | microtubule-associated proteins 70-3 | 70 kDa | 0 | 0 | 17 | 17 | 34 | | 0.25 | 0.25 | | | | |
| AT3G49670 | AT3G49670.1 | Leucine-rich receptor-like protein kinase family protein | 109 kDa | 0 | 0 | 17 | 17 | 34 | | | | 1.4E+06 | 3.8E+05 | 0.0E+00 | 1.8E+06 |
| AT3G51310 | AT3G51310.1 | VPS35 homolog C | 89 kDa | 0 | 4 | 17 | 17 | 38 | | | | 6.3E+05 | 5.2E+05 | 9.8E+04 | 1.3E+06 |
| AT5G11700 | AT5G11700.1;AT5G11700.2 | AT5G11700.2 | ? | 0 | 0 | 17 | 17 | 34 | | | | 2.8E+05 | 1.5E+05 | 0.0E+00 | 4.3E+05 |
| AT5G65290 | | LMBR1-like membrane protein | 83 kDa | 0 | 1 | 17 | 16 | 34 | | | | | | | |
| AT4G34230 | AT4G34230.1;AT4G34230.2 | cinnamyl alcohol dehydrogenase 5 | 39 kDa | 1 | 0 | 17 | 15 | 33 | | | | 3.2E+06 | 6.6E+06 | 0.0E+00 | 9.8E+06 |
| AT5G16000 | AT5G16000.1 | NSP-interacting kinase 1 | 71 kDa | 0 | 0 | 17 | 15 | 32 | | | | 3.0E+06 | 1.5E+05 | 0.0E+00 | 3.1E+06 |
| AT5G38830 | AT5G38830.1 | Cysteinyl-tRNA synthetase, class Ia family protein | 58 kDa | 3 | 0 | 17 | 14 | 34 | | | | 1.3E+07 | 1.0E+07 | 4.3E+04 | 2.3E+07 |
| AT1G63430 | | Leucine-rich repeat protein kinase family protein | 74 kDa | 0 | 0 | 17 | 13 | 30 | | | | | | | |
| AT3G08600 | AT3G08600.1 | Protein of unknown function (DUF1191) | 35 kDa | 2 | 0 | 17 | 13 | 32 | | | | 7.1E+05 | 8.6E+05 | 0.0E+00 | 1.6E+06 |
| AT3G47550 | | RING/FYVE/PHD zinc finger superfamily protein | 32 kDa | 0 | 7 | 17 | 13 | 37 | | | | | | | |
| AT1G29030 | | Apoptosis inhibitory protein 5 (API5) | 62 kDa | 0 | 0 | 17 | 12 | 29 | | | | | | | |
| AT2G03430 | AT2G03430.1 | Ankyrin repeat family protein | 26 kDa | 4 | 17 | 17 | 12 | 50 | | | | 1.4E+06 | 1.6E+06 | 4.7E+05 | 3.5E+06 |
| AT3G54920 | | Pectin lyase-like superfamily protein | 54 kDa | 0 | 0 | 17 | 12 | 29 | | | | | | | |
| AT4G24260 | | glycosyl hydrolase 9A3 | 70 kDa | 0 | 3 | 17 | 12 | 32 | | | | | | | |
| AT1G21170 | | Exocyst complex component SEC5 | 123 kDa | 2 | 0 | 17 | 11 | 30 | | | | | | | |
| AT5G07910 | | Leucine-rich repeat (LRR) family protein | ? | 1 | 6 | 17 | 11 | 35 | | 6.76 | 6.76 | | | | |
| AT5G64500 | | Major facilitator superfamily protein | 52 kDa | 0 | 0 | 17 | 11 | 28 | | | | | | | |
| AT1G03590 | | Protein phosphatase 2C family protein | 51 kDa | 0 | 0 | 17 | 10 | 27 | | | | | | | |
| AT1G20580 | AT1G20580.1 | Small nuclear ribonucleoprotein family protein | 14 kDa | 5 | 0 | 17 | 9 | 31 | | | | 1.9E+07 | 2.8E+07 | 6.7E+06 | 5.4E+07 |
| AT2G33450 | AT2G33450.1 | Ribosomal L28 family | 16 kDa | 0 | 0 | 17 | 9 | 26 | | | | 0.0E+00 | 6.0E+05 | 0.0E+00 | 6.0E+05 |
| AT4G27270 | | Quinone reductase family protein | 22 kDa | 0 | 0 | 17 | 9 | 26 | | | | | | | |
| AT5G04770 | AT5G04770.2;AT5G04770.1 | cationic amino acid transporter 6 | 63 kDa | 0 | 0 | 17 | 9 | 26 | | | | 1.0E+06 | 6.9E+05 | 0.0E+00 | 1.7E+06 |
| AT1G44900 | AT1G44900.2;AT1G44900.1 | minichromosome maintenance (MCM2/3/5) family prc | 106 kDa | 0 | 0 | 17 | 8 | 25 | | | | 7.8E+06 | 8.1E+06 | 2.4E+05 | 1.6E+07 |
| AT5G63570 | AT5G63570.2;AT5G63570.1 | glutamate-1-semialdehyde-2,1-aminomutase | 50 kDa | 4 | 3 | 17 | 8 | 32 | | | | 3.2E+06 | 1.9E+06 | 0.0E+00 | 5.2E+06 |
| AT5G65960 | AT5G65960.1 | GTP binding | 44 kDa | 11 | 5 | 17 | 8 | 41 | | | | 1.7E+06 | 3.0E+06 | 0.0E+00 | 4.7E+06 |
| AT1G15290 | AT1G15290.2;AT1G15290.1 | Tetratricopeptide repeat (TPR)-like superfamily protein | 178 kDa | 0 | 0 | 17 | 7 | 24 | | | | 0.0E+00 | 0.0E+00 | 9.1E+04 | 9.1E+04 |
| AT5G42920 | AT5G42920.2;AT5G42920.1 | THO complex, subunit 5 | 93 kDa | 0 | 0 | 17 | 7 | 24 | | | | 3.0E+05 | 2.2E+05 | 3.3E+05 | 8.5E+05 |
| AT3G48730 | AT3G48730.1 | glutamate-1-semialdehyde 2,1-aminomutase 2 | 50 kDa | 0 | 3 | 17 | 6 | 26 | | | | 1.1E+06 | 8.7E+05 | 0.0E+00 | 1.9E+06 |
| AT5G39040 | | transporter associated with antigen processing protein | 69 kDa | 0 | 0 | 17 | 6 | 23 | | | | | | | |
| AT2G24170 | | Endomembrane protein 70 protein family | 73 kDa | 0 | 0 | 17 | 3 | 20 | | | | | | | |
| AT5G03300 | AT5G03300.3;AT5G03300.2;AT5G03300.1 | adenosine kinase 2 | 38 kDa | 0 | 1 | 16 | 36 | 53 | | | | 1.4E+06 | 0.0E+00 | 0.0E+00 | 1.4E+06 |
| AT4G22730 | AT4G22730.1;AT4G22730.2 | Leucine-rich repeat protein kinase family protein | 76 kDa | 0 | 0 | 16 | 27 | 43 | | | | 9.7E+05 | 4.0E+05 | 0.0E+00 | 1.4E+06 |
| AT1G77120 | AT1G77120.1 | alcohol dehydrogenase 1 | 41 kDa | 3 | 0 | 16 | 25 | 44 | | | | 3.0E+06 | 1.9E+06 | 0.0E+00 | 4.9E+06 |
| AT2G23140 | | RING/U-box superfamily protein with ARM repeat dom | 90 kDa | 0 | 0 | 16 | 23 | 39 | | | | | | | |
| AT1G03220 | AT1G03220.1 | Eukaryotic aspartyl protease family protein | 46 kDa | 0 | 0 | 16 | 20 | 36 | | | | 6.2E+05 | 4.8E+05 | 0.0E+00 | 1.1E+06 |
| AT3G52400 | AT3G52400.1 | syntaxin of plants 122 | 38 kDa | 4 | 5 | 16 | 19 | 44 | | 5.15 | 5.15 | 2.4E+07 | 1.5E+07 | 2.9E+04 | 3.9E+07 |
| AT5G15490 | AT5G15490.1 | UDP-glucose 6-dehydrogenase family protein | 53 kDa | 5 | 6 | 16 | 19 | 46 | | 1.16 | 1.16 | 3.5E+06 | 4.1E+06 | 9.7E+04 | 7.8E+06 |
| AT5G56500 | AT5G56500.2;AT5G56500.1 | TCP-1/cpn60 chaperonin family protein | 63 kDa | 0 | 1 | 16 | 18 | 35 | | | | 8.3E+05 | 9.0E+05 | 0.0E+00 | 1.7E+06 |
| AT5G19440 | AT5G19440.1 | NAD(P)-binding Rossmann-fold superfamily protein | 36 kDa | 0 | 8 | 16 | 17 | 41 | | 3.17 | 3.17 | 7.1E+06 | 3.8E+06 | 0.0E+00 | 1.1E+07 |
| AT5G67200 | | Leucine-rich repeat protein kinase family protein | 74 kDa | 0 | 0 | 16 | 17 | 33 | | | | | | | |
| AT1G07670 | | endomembrane-type CA-ATPase 4 | 116 kDa | 0 | 2 | 16 | 16 | 34 | | 0.13 | 0.13 | | | | |
| AT1G21080 | | DNAJ heat shock N-terminal domain-containing protei | 44 kDa | 2 | 10 | 16 | 16 | 44 | | | | | | | |
| AT4G39820 | | Tetratricopeptide repeat (TPR)-like superfamily protein | 46 kDa | 0 | 0 | 16 | 16 | 32 | | | | | | | |
| AT5G43320 | | casein kinase I-like 8 | 54 kDa | 0 | 0 | 16 | 16 | 32 | | | | | | | |
| AT5G44130 | | FASCICLIN-like arabinogalactan protein 13 precursor | 26 kDa | 0 | 1 | 16 | 16 | 33 | | 3.98 | 3.98 | | | | |
| AT2G23080 | | Protein kinase superfamily protein | 39 kDa | 0 | 0 | 16 | 15 | 31 | | | | | | | |
| AT2G43130 | | P-loop containing nucleoside triphosphate hydrolases | 24 kDa | 0 | 12 | 16 | 15 | 43 | | | | | | | |
| AT3G17250 | | Protein phosphatase 2C family protein | 48 kDa | 0 | 1 | 16 | 15 | 32 | | | | | | | |
| AT4G29130 | | hexokinase 1 | 54 kDa | 0 | 0 | 16 | 15 | 31 | | 0.28 | 0.28 | | | | |
| AT4G04940 | AT4G04940.1;AT4G04940.2 | transducin family protein / WD-40 repeat family protei | 102 kDa | 0 | 1 | 16 | 14 | 31 | | | | 9.6E+05 | 9.7E+05 | 4.8E+04 | 2.0E+06 |
| AT5G08610 | AT5G08610.1 | P-loop containing nucleoside triphosphate hydrolases | 94 kDa | 0 | 0 | 16 | 14 | 30 | | | | 1.1E+06 | 7.7E+05 | 3.7E+05 | 2.2E+06 |
| AT1G76390 | | ARM repeat superfamily protein | 89 kDa | 0 | 0 | 16 | 13 | 29 | | | | | | | |
| AT2G20230 | AT2G20230.1 | Tetraspanin family protein | 30 kDa | 0 | 2 | 16 | 13 | 31 | | 0.30 | 0.30 | 2.0E+06 | 1.4E+06 | 0.0E+00 | 3.3E+06 |
| AT4G17615 | | calcineurin B-like protein 1 | 25 kDa | 0 | 0 | 16 | 13 | 29 | | 10.69 | 10.69 | | | | |
| AT4G23720 | | Protein of unknown function (DUF1191) | 35 kDa | 0 | 0 | 16 | 13 | 29 | | | | | | | |
| AT2G47620 | AT2G47620.1;AT2G47620.2 | SWITCH/sucrose nonfermenting 3A | 57 kDa | 0 | 0 | 16 | 12 | 28 | | | | 5.8E+05 | 5.5E+05 | 7.6E+04 | 1.2E+06 |
| AT3G11530 | | Vacuolar protein sorting 55 (VPS55) family protein | 12 kDa | 2 | 1 | 16 | 12 | 31 | 3.37 | 2.42 | 2.90 | | | | |
| AT2G20060 | AT2G20060.1 | Ribosomal protein L4/L1 family | 33 kDa | 1 | 0 | 16 | 11 | 28 | | | | 3.0E+07 | 6.3E+07 | 7.0E+06 | 1.0E+08 |
| AT2G43320 | | S-adenosyl-L-methionine-dependent methyltransferase | 39 kDa | 0 | 2 | 16 | 11 | 29 | | | | | | | |
| AT3G26650 | AT3G26650.1;AT1G12900.2;AT1G12900.4;AT1G12900.3;AT1G12900.1;AT1G12900.5 | glyceraldehyde 3-phosphate dehydrogenase A subunit | 42 kDa | 0 | 4 | 16 | 11 | 31 | | | | 7.5E+06 | 1.1E+07 | 0.0E+00 | 1.9E+07 |
| AT5G39590 | AT5G39590.1 | TLD-domain containing nucleolar protein | 60 kDa | 3 | 0 | 16 | 11 | 30 | | | | 2.6E+06 | 1.3E+06 | 0.0E+00 | 3.8E+06 |
| AT5G44635 | AT5G44635.1;AT5G44635.2 | minichromosome maintenance (MCM2/3/5) family prc | 93 kDa | 0 | 0 | 16 | 11 | 27 | | | | 8.6E+06 | 9.8E+06 | 3.4E+05 | 1.9E+07 |
| AT3G49430 | AT3G49430.2;AT3G49430.7;AT3G49430.6;AT3G49430.5;AT3G49430.4;AT3G49430.3;AT3G49430.1 | SER/ARG-rich protein 34A | 34 kDa | 0 | 1 | 16 | 10 | 27 | | | | 2.0E+07 | 2.0E+07 | 5.2E+06 | 4.5E+07 |
| AT1G05890 | | RING/U-box superfamily protein | 62 kDa | 0 | 0 | 16 | 9 | 25 | | | | | | | |
| AT1G32490 | AT1G32490.2;AT1G32490.1;AT2G35340.2;AT2G35340.1 | RNA helicase family protein | 118 kDa | 0 | 0 | 16 | 9 | 25 | | | | 4.8E+05 | 4.9E+05 | 3.0E+04 | 9.9E+05 |
| AT1G15440 | AT1G15440.1;AT1G15440.2 | periodic tryptophan protein 2 | 100 kDa | 0 | 0 | 16 | 8 | 24 | | 0.23 | 0.23 | 1.0E+06 | 4.6E+05 | 4.9E+05 | 2.0E+06 |
| AT1G45145 | AT1G45145.1 | thioredoxin H-type 5 | 13 kDa | 0 | 0 | 16 | 7 | 23 | | 2.77 | 2.77 | 5.9E+06 | 5.1E+06 | 0.0E+00 | 1.1E+07 |
| AT2G18220 | AT2G18220.1 | Noc2p family | 87 kDa | 0 | 0 | 16 | 7 | 23 | | | | 1.8E+06 | 1.7E+06 | 4.1E+05 | 3.9E+06 |
| AT2G34357 | AT2G34357.1 | ARM repeat superfamily protein | 142 kDa | 0 | 0 | 16 | 7 | 23 | | | | 1.9E+05 | 9.3E+04 | 0.0E+00 | 2.8E+05 |
| AT3G08710 | AT3G08710.3;AT3G08710.2;AT3G08710.1 | thioredoxin H-type 9 | 15 kDa | 4 | 0 | 16 | 7 | 27 | | | | 6.3E+06 | 2.9E+06 | 0.0E+00 | 9.2E+06 |
| AT5G13680 | AT5G13680.1 | IKI3 family protein | 147 kDa | 0 | 0 | 16 | 7 | 23 | | 0.18 | 0.18 | 4.3E+06 | 4.3E+06 | 2.6E+05 | 8.8E+06 |
| AT3G63080 | | glutathione peroxidase 5 | 19 kDa | 4 | 0 | 16 | 6 | 26 | | | | | | | |
| AT5G47930 | AT5G47930.1 | Zinc-binding ribosomal protein family protein | 9 kDa | 4 | 0 | 16 | 6 | 26 | | 0.13 | 0.13 | 2.6E+07 | 3.2E+07 | 2.8E+07 | 8.6E+07 |
| AT1G69670 | | cullin 3B | 86 kDa | 0 | 0 | 16 | 5 | 21 | | | | | | | |
| AT2G18040 | AT2G18040.1 | peptidylprolyl cis/trans isomerase, NIMA-interacting 1 | 13 kDa | 0 | 0 | 16 | 4 | 20 | | 0.82 | 0.82 | 2.6E+06 | 1.2E+06 | 0.0E+00 | 3.8E+06 |

| | | | | | | | | | | | | | | | |
|---|---|---|---|---|---|---|---|---|---|---|---|---|---|---|---|
| AT2G32060 | AT2G32060.3;AT2G32060.2;AT2G32060.1 | Ribosomal protein L7Ae/L30e/S12e/Gadd45 family prc | 15 kDa | 0 | 0 | 16 | 4 | 20 | | 0.26 | 0.26 | 6.1E+07 | 3.8E+07 | 1.6E+07 | 1.1E+08 |
| AT5G40080 | AT5G40080.1;AT5G39800.1 | Mitochondrial ribosomal protein L27 | 10 kDa | 0 | 0 | 16 | 1 | 17 | | | | 4.0E+07 | 4.9E+07 | 3.9E+06 | 9.3E+07 |
| AT4G33360 | | NAD(P)-binding Rossmann-fold superfamily protein | 38 kDa | 0 | 0 | 15 | 29 | 44 | | 1.09 | 1.09 | | | | |
| AT1G70940 | | Auxin efflux carrier family protein | 69 kDa | 0 | 0 | 15 | 27 | 42 | | | | | | | |
| AT2G26260 | | 3beta-hydroxysteroid-dehydrogenase/decarboxylase is | 62 kDa | 0 | 0 | 15 | 27 | 42 | | | | | | | |
| AT4G17140 | AT4G17140.1;AT4G17140.2;AT4G17140.3 | pleckstrin homology (PH) domain-containing protein | 471 kDa | 0 | 0 | 15 | 27 | 42 | | 0.56 | 0.56 | 5.6E+05 | 6.8E+05 | 0.0E+00 | 1.2E+06 |
| AT2G34510 | | Protein of unknown function, DUF642 | 44 kDa | 0 | 0 | 15 | 26 | 41 | | | | | | | |
| AT1G12230 | | Aldolase superfamily protein | 47 kDa | 1 | 13 | 15 | 24 | 53 | | | | | | | |
| AT1G79610 | AT1G79610.1 | Na+/H+ antiporter 6 | 59 kDa | 0 | 0 | 15 | 23 | 38 | | | | 4.9E+05 | 7.8E+05 | 0.0E+00 | 1.3E+06 |
| AT5G65687 | | Major facilitator superfamily protein | 54 kDa | 1 | 0 | 15 | 23 | 39 | | | | | | | |
| AT1G52280 | | RAB GTPase homolog G3D | 23 kDa | 0 | 10 | 15 | 20 | 45 | | 0.22 | 0.22 | | | | |
| AT4G20940 | | Leucine-rich receptor-like protein kinase family proteir | 105 kDa | 0 | 0 | 15 | 19 | 34 | | | | | | | |
| AT1G23890 | AT1G23890.1;AT1G23890.2 | NHL domain-containing protein | 44 kDa | 0 | 5 | 15 | 18 | 38 | | | | 1.2E+06 | 1.5E+06 | 0.0E+00 | 2.6E+06 |
| AT1G57720 | AT1G57720.2;AT1G57720.1 | Translation elongation factor EF1B, gamma chain | 46 kDa | 0 | 0 | 15 | 18 | 33 | | | | 5.2E+06 | 1.4E+07 | 0.0E+00 | 1.9E+07 |
| AT2G03480 | | QUASIMODO2 LIKE 2 | 68 kDa | 0 | 0 | 15 | 18 | 33 | | | | | | | |
| AT5G22850 | AT5G22850.1 | Eukaryotic aspartyl protease family protein | 52 kDa | 2 | 0 | 15 | 18 | 35 | | | | 8.7E+06 | 5.6E+06 | 0.0E+00 | 1.4E+07 |
| AT2G31880 | | Leucine-rich repeat protein kinase family protein | 71 kDa | 0 | 1 | 15 | 17 | 33 | | | | | | | |
| AT5G23060 | AT5G23060.2;AT5G23060.1 | calcium sensing receptor | 41 kDa | 6 | 0 | 15 | 17 | 38 | | 2.31 | 2.31 | 1.0E+07 | 5.0E+06 | 1.6E+05 | 1.5E+07 |
| AT5G49890 | AT5G49890.1 | chloride channel C | 85 kDa | 0 | 0 | 15 | 17 | 32 | | 0.19 | 0.19 | 1.4E+06 | 0.0E+00 | 2.5E+05 | 1.7E+06 |
| AT1G51570 | | Calcium-dependent lipid-binding (CaLB domain) plant | 89 kDa | 6 | 0 | 15 | 16 | 37 | | | | | | | |
| AT5G41880 | | DNA primases;DNA primases | 52 kDa | 0 | 0 | 15 | 16 | 31 | | | | | | | |
| AT4G17270 | | Mo25 family protein | 40 kDa | 1 | 2 | 15 | 15 | 33 | | | | | | | |
| AT5G63680 | AT5G63680.3;AT5G63680.2;AT5G63680.1 | Pyruvate kinase family protein | 55 kDa | 0 | 2 | 15 | 15 | 32 | | | | 5.1E+06 | 6.8E+06 | 2.7E+04 | 1.2E+07 |
| AT1G18390 | AT1G18390.1;AT1G18390.2 | Protein kinase superfamily protein | 73 kDa | 0 | 1 | 15 | 14 | 30 | | | | 0.0E+00 | 5.2E+04 | 0.0E+00 | 5.2E+04 |
| AT5G61790 | AT5G61790.1 | calnexin 1 | 60 kDa | 0 | 0 | 15 | 14 | 29 | 0.06 | 0.18 | 0.12 | 7.9E+05 | 1.1E+06 | 0.0E+00 | 1.9E+06 |
| AT4G03080 | | BRI1 suppressor 1 (BSU1)-like 1 | 96 kDa | 0 | 0 | 15 | 13 | 28 | | | | | | | |
| AT1G50240 | AT1G50240.3;AT1G50240.2 | Protein kinase family protein with ARM repeat domain | 145 kDa | 0 | 0 | 15 | 12 | 27 | | | | 1.8E+05 | 8.9E+04 | 0.0E+00 | 2.7E+05 |
| AT3G59140 | AT3G59140.1 | multidrug resistance-associated protein 14 | 163 kDa | 1 | 0 | 15 | 12 | 28 | | 0.40 | 0.40 | 6.1E+05 | 1.2E+05 | 5.4E+03 | 7.3E+05 |
| AT5G08620 | AT5G08620.1 | DEA(D/H)-box RNA helicase family protein | 62 kDa | 0 | 0 | 15 | 12 | 27 | | | | 6.3E+05 | 0.0E+00 | 3.8E+05 | 1.0E+06 |
| AT5G13890 | | Family of unknown function (DUF716) | 34 kDa | 4 | 20 | 15 | 12 | 51 | | | | | | | |
| AT1G64090 | AT1G64090.1;AT1G64090.2 | Reticulan like protein B3 | 29 kDa | 8 | 11 | 15 | 11 | 45 | 0.17 | | 0.17 | 9.4E+05 | 1.1E+06 | 6.0E+05 | 2.6E+06 |
| AT1G75850 | | VPS35 homolog B | 89 kDa | 0 | 0 | 15 | 11 | 26 | | | | | | | |
| AT2G38550 | AT2G38550.1 | Transmembrane proteins 14C | 37 kDa | 1 | 2 | 15 | 11 | 29 | 0.07 | | 0.07 | 6.5E+05 | 1.9E+06 | 0.0E+00 | 2.5E+06 |
| AT3G48750 | AT3G48750.1 | cell division control 2 | 34 kDa | 1 | 13 | 15 | 11 | 40 | | | | 1.9E+06 | 0.0E+00 | 3.8E+05 | 2.2E+06 |
| AT4G39090 | | Papain family cysteine protease | 40 kDa | 0 | 4 | 15 | 11 | 30 | | | | | | | |
| AT5G42540 | AT5G42540.1;AT5G42540.2 | exoribonuclease 2 | 116 kDa | 0 | 0 | 15 | 11 | 26 | | | | 1.6E+07 | 1.2E+07 | 1.0E+06 | 2.9E+07 |
| AT1G73110 | | P-loop containing nucleoside triphosphate hydrolases | 48 kDa | 0 | 2 | 15 | 10 | 27 | | | | | | | |
| AT3G12740 | | ALA-interacting subunit 1 | 39 kDa | 0 | 0 | 15 | 10 | 25 | | | | | | | |
| AT5G06260 | | TLD-domain containing nucleolar protein | 47 kDa | 0 | 0 | 15 | 10 | 25 | | | | | | | |
| AT5G20600 | AT5G20600.1 | FUNCTIONS IN: molecular_function unknown; INVOLVE | 60 kDa | 0 | 0 | 15 | 10 | 25 | | | | 3.7E+05 | 0.0E+00 | 4.2E+05 | 7.9E+05 |
| AT1G80670 | | Transducin/WD40 repeat-like superfamily protein | 38 kDa | 0 | 10 | 15 | 9 | 34 | | | | | | | |
| AT2G16485 | AT2G16485.2;AT2G16485.1 | nucleic acid binding;zinc ion binding;DNA binding | 195 kDa | 0 | 0 | 15 | 9 | 24 | | | | 4.2E+06 | 3.4E+06 | 1.0E+05 | 7.6E+06 |
| AT2G46290 | | Transducin/WD40 repeat-like superfamily protein | 39 kDa | 1 | 0 | 15 | 9 | 25 | | | | | | | |
| AT4G29010 | AT4G29010.1 | Enoyl-CoA hydratase/isomerase family | 78 kDa | 0 | 0 | 15 | 9 | 24 | | | | 7.1E+06 | 1.2E+07 | 9.6E+05 | 2.0E+07 |
| AT4G40050 | | Protein of unknown function (DUF3550/UPF0682) | 68 kDa | 2 | 0 | 15 | 9 | 26 | | | | | | | |
| AT5G54110 | | membrane-associated mannitol-induced | 30 kDa | 1 | 9 | 15 | 9 | 34 | | | | | | | |
| AT1G76300 | AT1G76300.1 | snRNP core protein SMD3 | 14 kDa | 0 | 0 | 15 | 8 | 23 | | 0.20 | 0.20 | 7.9E+06 | 1.2E+07 | 2.8E+06 | 2.2E+07 |
| AT2G17265 | AT2G17265.1 | homoserine kinase | 39 kDa | 0 | 0 | 15 | 8 | 23 | | | | 1.1E+07 | 1.0E+07 | 2.1E+06 | 2.3E+07 |
| AT5G15450 | AT5G15450.1 | casein lytic proteinase B3 | 109 kDa | 0 | 0 | 15 | 8 | 23 | | | | 4.3E+06 | 5.2E+06 | 0.0E+00 | 9.5E+06 |
| AT3G07300 | | NagB/RpiA/CoA transferase-like superfamily protein | 44 kDa | 0 | 2 | 15 | 7 | 24 | | | | | | | |
| AT5G06680 | | spindle pole body component 98 | ? | 0 | 0 | 15 | 7 | 22 | | | | | | | |
| AT5G27740 | AT5G27740.1 | ATPase family associated with various cellular activities | 41 kDa | 0 | 0 | 15 | 7 | 22 | | | | 5.1E+05 | 8.5E+05 | 3.7E+05 | 1.7E+06 |
| AT1G04950 | | TATA BOX ASSOCIATED FACTOR II 59 | 61 kDa | 0 | 0 | 15 | 6 | 21 | | | | | | | |
| AT1G05460 | | P-loop containing nucleoside triphosphate hydrolases | 113 kDa | 0 | 0 | 15 | 6 | 21 | | | | | | | |
| AT1G65430 | | IBR domain-containing protein | 65 kDa | 0 | 0 | 15 | 6 | 21 | | | | | | | |
| AT3G09720 | AT3G09720.2;AT3G09720.1 | P-loop containing nucleoside triphosphate hydrolases | 61 kDa | 0 | 0 | 15 | 6 | 21 | | | | 1.5E+06 | 9.1E+05 | 1.4E+05 | 2.6E+06 |
| AT1G31812 | AT1G31812.1 | acyl-CoA-binding protein 6 | 10 kDa | 0 | 0 | 15 | 5 | 20 | | | | 1.0E+08 | 3.5E+07 | 3.6E+05 | 1.4E+08 |
| AT2G07696 | | Ribosomal protein S7p/S5e family protein | 17 kDa | 8 | 0 | 15 | 5 | 28 | 0.49 | 0.46 | 0.47 | | | | |
| AT3G59650 | AT3G59650.1;AT3G59650.2 | mitochondrial ribosomal protein L51/S25/CI-B8 family | 14 kDa | 0 | 0 | 15 | 5 | 20 | | 0.50 | 0.50 | 4.3E+07 | 8.4E+07 | 3.7E+06 | 1.3E+08 |
| AT4G30930 | AT4G30930.1 | Ribosomal protein L21 | 31 kDa | 0 | 1 | 15 | 5 | 21 | | | | 4.2E+07 | 7.3E+07 | 4.8E+06 | 1.2E+08 |
| AT5G14050 | AT5G14050.1 | Transducin/WD40 repeat-like superfamily protein | 61 kDa | 0 | 0 | 15 | 5 | 20 | | | | 1.7E+06 | 1.9E+06 | 1.4E+05 | 3.7E+06 |
| AT5G26800 | AT5G26800.1 | FUNCTIONS IN: molecular_function unknown; INVOLVE | ? | 2 | 0 | 15 | 5 | 22 | 0.48 | 0.63 | 0.56 | 1.2E+08 | 1.3E+08 | 9.5E+06 | 2.6E+08 |
| AT2G06530 | AT2G06530.1;AT2G06530.2 | SNF7 family protein | 25 kDa | 0 | 0 | 15 | 4 | 19 | | | | 1.1E+06 | 2.0E+06 | 0.0E+00 | 3.0E+06 |
| AT2G41140 | | CDPK-related kinase 1 | 64 kDa | 0 | 0 | 15 | 4 | 19 | | | | | | | |
| AT3G18165 | AT3G18165.1 | modifier of snc1,4 | 30 kDa | 0 | 2 | 15 | 4 | 21 | | | | 1.1E+07 | 1.3E+07 | 1.4E+05 | 2.4E+07 |
| AT4G00810 | AT4G00810.2;AT4G00810.1 | 60S acidic ribosomal protein family | 11 kDa | 0 | 0 | 15 | 2 | 17 | | 0.39 | 0.39 | 2.0E+08 | 2.8E+08 | 5.3E+07 | 5.3E+08 |
| AT2G24200 | AT2G24200.3;AT2G24200.2;AT2G24200.1;AT4G30920.1 | Cytosol aminopeptidase family protein | 55 kDa | 8 | 1 | 14 | 35 | 58 | | | | 2.7E+06 | 2.6E+06 | 0.0E+00 | 5.3E+06 |
| AT3G54840 | | Ras-related small GTP-binding family protein | 22 kDa | 0 | 3 | 14 | 23 | 40 | | | | | | | |
| AT4G27690 | AT4G27690.2;AT4G27690.1 | vacuolar protein sorting 26B | 35 kDa | 0 | 1 | 14 | 21 | 36 | | | | 0.0E+00 | 0.0E+00 | 2.1E+05 | 2.1E+05 |
| AT4G37930 | AT4G37930.1 | serine transhydroxymethyltransferase 1 | 57 kDa | 0 | 0 | 14 | 21 | 35 | | | | 8.1E+06 | 7.1E+06 | 6.7E+04 | 1.5E+07 |
| AT1G70730 | | Phosphoglucomutase/phosphomannomutase family p | 63 kDa | 0 | 0 | 14 | 20 | 34 | | | | | | | |
| AT3G03250 | AT3G03250.3;AT3G03250.1;AT3G03250.2;AT5G17310.3;AT5G17310.5;AT5G17310.1 | UDP-GLUCOSE PYROPHOSPHORYLASE 1 | 52 kDa | 0 | 2 | 14 | 20 | 36 | | | | 5.0E+05 | 2.0E+05 | 0.0E+00 | 7.0E+05 |
| AT3G29360 | AT3G29360.2;AT3G29360.1;AT5G39320.1 | UDP-glucose 6-dehydrogenase family protein | 53 kDa | 6 | 5 | 14 | 20 | 45 | | | | 1.2E+06 | 6.8E+05 | 0.0E+00 | 1.9E+06 |
| AT4G35860 | AT4G35860.1;AT4G17170.1;AT4G35860.2;AT4G17160.1 | GTP-binding 2 | 23 kDa | 6 | 8 | 14 | 20 | 48 | | | | 1.3E+06 | 9.4E+05 | 0.0E+00 | 2.2E+06 |
| AT1G60420 | AT1G60420.1 | DC1 domain-containing protein | 65 kDa | 0 | 0 | 14 | 19 | 33 | | | | | | | |
| AT4G10840 | | Tetratricopeptide repeat (TPR)-like superfamily protein | 66 kDa | 0 | 0 | 14 | 19 | 33 | | | | | | | |
| AT3G15460 | AT3G15460.1 | Ribosomal RNA processing Brix domain protein | 36 kDa | 0 | 0 | 14 | 18 | 32 | | | | 0.0E+00 | 0.0E+00 | 8.0E+05 | 8.0E+05 |
| AT3G22590 | AT3G22590.1 | PLANT HOMOLOGOUS TO PARAFIBROMIN | 48 kDa | 0 | 0 | 14 | 18 | 32 | | | | 5.0E+06 | 6.1E+06 | 7.0E+05 | 1.2E+07 |
| AT1G04120 | | multidrug resistance-associated protein 5 | 169 kDa | 0 | 0 | 14 | 17 | 31 | | | | | | | |
| AT1G31660 | | CONTAINS InterPro DOMAIN/s: Bystin (InterPro:IPR007 | 51 kDa | 0 | 0 | 14 | 17 | 31 | | 0.22 | 0.22 | | | | |
| AT2G25730 | | unknown protein; Has 157 Blast hits to 144 proteins in | 275 kDa | 0 | 0 | 14 | 17 | 31 | | | | | | | |
| AT3G23310 | | AGC (cAMP-dependent, cGMP-dependent and protein k | 65 kDa | 0 | 0 | 14 | 17 | 31 | | | | | | | |
| AT4G17180 | AT4G17180.1 | O-Glycosyl hydrolases family 17 protein | 52 kDa | 0 | 0 | 14 | 17 | 31 | | | | 3.9E+05 | 5.2E+05 | 0.0E+00 | 9.1E+05 |
| AT5G11170 | | DEAD/DEAH box RNA helicase family protein | 48 kDa | 0 | 2 | 14 | 17 | 33 | | | | | | | |
| AT1G17750 | AT1G17750.1 | PEP1 receptor 2 | 119 kDa | 0 | 1 | 14 | 16 | 31 | | | | | | | |
| AT5G25090 | AT5G25090.1 | early nodulin-like protein 13 | 20 kDa | 0 | 0 | 14 | 16 | 30 | | | | 3.6E+06 | 2.1E+06 | 0.0E+00 | 5.8E+06 |
| AT1G06530 | | Tropomyosin-related | 36 kDa | 0 | 0 | 14 | 15 | 29 | 0.14 | | 0.14 | | | | |
| AT2G19760 | AT2G19760.1 | profilin 1 | 14 kDa | 7 | 0 | 14 | 15 | 36 | | 3.09 | 3.09 | 3.6E+07 | 1.7E+07 | 0.0E+00 | 5.3E+07 |
| AT5G17990 | AT5G17990.1 | tryptophan biosynthesis 1 | 47 kDa | 0 | 18 | 14 | 15 | 47 | | | | 2.5E+06 | 1.8E+06 | 0.0E+00 | 4.3E+06 |
| AT1G72680 | AT1G72680.1 | cinnamyl-alcohol dehydrogenase | 39 kDa | 1 | 0 | 14 | 14 | 29 | | | | 2.6E+07 | 6.1E+06 | 0.0E+00 | 3.2E+07 |
| AT2G39050 | AT2G39050.1 | hydroxyproline-rich glycoprotein family protein | 36 kDa | 0 | 3 | 14 | 13 | 30 | | | | 2.0E+06 | 4.1E+05 | 0.0E+00 | 2.4E+06 |

| | | | | | | | | | | | | | | | |
|---|---|---|---|---|---|---|---|---|---|---|---|---|---|---|---|
| AT1G25390 | | Protein kinase superfamily protein | 71 kDa | 0 | 1 | 14 | 12 | 27 | | | | | | | |
| AT3G22910 | | ATPase E1-E2 type family protein / haloacid dehalogena | 113 kDa | 0 | 0 | 14 | 12 | 26 | | | | | | | |
| AT3G57030 | | Calcium-dependent phosphotriesterase superfamily pr | 41 kDa | 0 | 0 | 14 | 12 | 26 | | | | | | | |
| AT1G09760 | AT1G09760.1 | U2 small nuclear ribonucleoprotein A | 28 kDa | 4 | 6 | 14 | 11 | 35 | 0.11 | 0.20 | 0.15 | 7.1E+06 | 1.3E+07 | 4.1E+06 | 2.5E+07 |
| AT1G31420 | | Leucine-rich repeat protein kinase family protein | 65 kDa | 0 | 0 | 14 | 11 | 25 | | | | | | | |
| AT2G48010 | AT2G48010.1 | receptor-like kinase in in flowers 3 | 67 kDa | 0 | 1 | 14 | 11 | 26 | | | | 0.0E+00 | 0.0E+00 | 0.0E+00 | 0.0E+00 |
| AT1G02890 | | AAA-type ATPase family protein | 136 kDa | 0 | 0 | 14 | 10 | 24 | | | | | | | |
| AT1G08880 | | Histone superfamily protein | 15 kDa | 9 | 0 | 14 | 10 | 33 | 0.11 | | 0.11 | | | | |
| AT1G09020 | | homolog of yeast sucrose nonfermenting 4 | 53 kDa | 0 | 2 | 14 | 10 | 26 | | | | | | | |
| AT2G35190 | AT2G35190.1 | novel plant snare 11 | 30 kDa | 0 | 2 | 14 | 10 | 26 | | | | 1.8E+06 | 7.2E+05 | 0.0E+00 | 2.5E+06 |
| AT2G35960 | AT2G35960.1 | NDR1/HIN1-like 12 | 23 kDa | 0 | 0 | 14 | 10 | 24 | | | | 8.8E+05 | 4.5E+05 | 0.0E+00 | 1.3E+06 |
| AT4G21660 | AT4G21660.1;AT4G21660.2;AT4G21660.3 | proline-rich spliceosome-associated (PSP) family protei | 69 kDa | 0 | 0 | 14 | 10 | 24 | | | | 1.2E+07 | 1.0E+07 | 2.7E+06 | 2.5E+07 |
| AT5G16830 | | syntaxin of plants 21 | 31 kDa | 8 | 17 | 14 | 10 | 49 | | 2.36 | 2.36 | | | | |
| AT2G35620 | | Leucine-rich repeat protein kinase family protein | 64 kDa | 0 | 0 | 14 | 9 | 23 | | | | | | | |
| AT3G10050 | AT3G10050.1 | L-O-methylthreonine resistant 1 | 65 kDa | 0 | 0 | 14 | 9 | 23 | | | | 1.9E+05 | 6.3E+05 | 1.5E+05 | 9.8E+05 |
| AT1G75840 | | RAC-like GTP binding protein 5 | 22 kDa | 5 | 2 | 14 | 8 | 29 | | | | | | | |
| AT3G22320 | AT3G22320.1 | Eukaryotic rpb5 RNA polymerase subunit family protei | 24 kDa | 0 | 1 | 14 | 8 | 23 | | 0.43 | 0.43 | 5.8E+06 | 5.6E+06 | 3.4E+05 | 1.2E+07 |
| AT5G18280 | | apyrase 2 | 69 kDa | 0 | 2 | 14 | 8 | 24 | | | | | | | |
| AT2G21790 | AT2G21790.1 | ribonucleotide reductase 1 | 92 kDa | 0 | 0 | 14 | 7 | 21 | | | | 6.2E+05 | 1.3E+06 | 2.3E+04 | 1.9E+06 |
| AT2G27130 | AT2G27130.1 | Bifunctional inhibitor/lipid-transfer protein/seed stora | 18 kDa | 1 | 1 | 14 | 7 | 23 | | | | 8.2E+06 | 4.8E+06 | 0.0E+00 | 1.3E+07 |
| AT5G58670 | | phospholipase C1 | 64 kDa | 0 | 1 | 14 | 7 | 22 | | | | | | | |
| AT5G64740 | | cellulose synthase 6 | 123 kDa | 0 | 0 | 14 | 7 | 21 | | | | | | | |
| AT1G05500 | | Calcium-dependent lipid-binding (CaLB domain) family | 63 kDa | 0 | 0 | 14 | 6 | 20 | | | | | | | |
| AT1G20090 | | RHO-related protein from plants 2 | 22 kDa | 4 | 1 | 14 | 6 | 25 | | | | | | | |
| AT1G54570 | AT1G54570.1 | Esterase/lipase/thioesterase family protein | 78 kDa | 0 | 0 | 14 | 6 | 20 | | | | | | | |
| AT2G47680 | AT2G47680.1 | zinc finger (CCCH type) helicase family protein | 115 kDa | 0 | 0 | 14 | 6 | 20 | | | | 3.1E+05 | 2.4E+05 | 2.3E+05 | 7.9E+05 |
| AT4G34460 | AT4G34460.2;AT4G34460.3;AT4G34460.4;AT4G34460.1 | GTP binding protein beta 1 | 41 kDa | 0 | 2 | 14 | 6 | 22 | | | | 2.2E+06 | 2.0E+06 | 0.0E+00 | 4.2E+06 |
| AT1G69340 | | appr-1-p processing enzyme family protein | 63 kDa | 0 | 0 | 14 | 5 | 19 | | | | | | | |
| AT2G46700 | | CDPK-related kinase 3 | 67 kDa | 0 | 0 | 14 | 5 | 19 | | | | | | | |
| AT3G20670 | AT3G20670.1 | histone H2A 13 | 14 kDa | 6 | 0 | 14 | 5 | 25 | | | | 2.4E+07 | 2.8E+07 | 3.8E+06 | 5.6E+07 |
| AT5G27540 | | MIRO-related GTP-ase 1 | 72 kDa | 0 | 0 | 14 | 5 | 19 | | 0.15 | 0.15 | | | | |
| AT1G49400 | AT1G49400.1 | Nucleic acid-binding, OB-fold-like protein | 13 kDa | 4 | 0 | 14 | 4 | 22 | | | | 9.4E+07 | 1.3E+08 | 5.6E+06 | 2.3E+08 |
| AT2G44290 | | Bifunctional inhibitor/lipid-transfer protein/seed stora | 22 kDa | 2 | 6 | 14 | 4 | 26 | | | | | | | |
| AT5G46210 | AT5G46210.1 | cullin4 | 91 kDa | 0 | 1 | 14 | 4 | 19 | | | | 3.2E+05 | 2.0E+05 | 2.2E+05 | 7.4E+05 |
| AT5G60920 | | COBRA-like extracellular glycosyl-phosphatidyl inositol | 51 kDa | 0 | 0 | 14 | 3 | 17 | | | | | | | |
| ATCG00650 | | ribosomal protein S18 | 12 kDa | 2 | 0 | 14 | 3 | 19 | 0.57 | 1.09 | 0.83 | | | | |
| AT4G17520 | AT4G17520.1 | Hyaluronan / mRNA binding family | 39 kDa | 0 | 0 | 14 | 2 | 16 | | | | 7.9E+05 | 8.5E+05 | 2.6E+06 | 4.3E+06 |
| AT1G26630 | AT1G26630.1;AT1G26630.2 | Eukaryotic translation initiation factor 5A-1 (eIF-5A 1) | 17 kDa | 4 | 0 | 14 | 1 | 19 | | | | 2.7E+07 | 2.8E+07 | 1.4E+07 | 6.9E+07 |
| AT5G07250 | | RHOMBOID-like protein 3 | 38 kDa | 0 | 0 | 13 | 25 | 38 | | | | | | | |
| AT4G17550 | AT4G17550.1 | Major facilitator superfamily protein | 59 kDa | 0 | 0 | 13 | 18 | 31 | | | | 2.4E+06 | 1.0E+06 | 0.0E+00 | 3.4E+06 |
| AT1G09780 | | Phosphoglycerate mutase, 2,3-bisphosphoglycerate-in | 61 kDa | 0 | 1 | 13 | 17 | 31 | | | | | | | |
| AT1G26460 | | Tetratricopeptide repeat (TPR)-like superfamily protein | 70 kDa | 0 | 4 | 13 | 17 | 34 | | | | | | | |
| AT4G26570 | AT4G26570.1;AT4G26570.2 | calcineurin B-like 3 | 26 kDa | 0 | 2 | 13 | 16 | 31 | 0.30 | 2.08 | 1.19 | 8.8E+05 | 4.0E+05 | 0.0E+00 | 1.3E+06 |
| AT1G76690 | | 12-oxophytodienoate reductase 2 | 42 kDa | 0 | 0 | 13 | 15 | 28 | | | | | | | |
| AT1G07810 | | ER-type Ca2+-ATPase 1 | 116 kDa | 0 | 2 | 13 | 14 | 29 | | 0.26 | 0.26 | | | | |
| AT1G30900 | | VACUOLAR SORTING RECEPTOR 6 | 70 kDa | 2 | 4 | 13 | 14 | 33 | | | | | | | |
| AT1G31280 | AT1G31280.1 | Argonaute family protein | 113 kDa | 0 | 0 | 13 | 13 | 26 | | 0.39 | 0.39 | 3.5E+05 | 8.3E+05 | 0.0E+00 | 1.2E+06 |
| AT2G23150 | | natural resistance-associated macrophage protein 3 | 56 kDa | 0 | 0 | 13 | 13 | 26 | | | | | | | |
| AT2G27260 | AT2G27260.1 | Late embryogenesis abundant (LEA) hydroxyproline-ric | 28 kDa | 0 | 6 | 13 | 13 | 32 | | | | 9.0E+07 | 1.9E+06 | 0.0E+00 | 9.2E+07 |
| AT3G08680 | | Leucine-rich repeat protein kinase family protein | 69 kDa | 0 | 0 | 13 | 12 | 25 | | | | | | | |
| AT4G19640 | AT4G19640.2;AT4G19640.3;AT5G45130.1;AT4G19640.1;AT3G54840.3;AT3G54840.2;AT3G54840.1;AT5G45130.2 | Ras-related small GTP-binding family protein | 22 kDa | 0 | 3 | 13 | 12 | 28 | | | | 1.6E+06 | 1.2E+06 | 1.4E+06 | 4.2E+06 |
| AT5G50160 | AT5G50160.2;AT5G50160.1 | ferric reduction oxidase 8 | 83 kDa | 1 | 0 | 13 | 12 | 26 | | | | 1.6E+06 | 7.5E+05 | 0.0E+00 | 2.3E+06 |
| AT1G50730 | | unknown protein; Has 218 Blast hits to 209 proteins in | 104 kDa | 0 | 0 | 13 | 11 | 24 | | | | | | | |
| AT3G01090 | | SNF1 kinase homolog 10 | 58 kDa | 0 | 0 | 13 | 11 | 24 | | | | | | | |
| AT3G19820 | AT3G19820.3;AT3G19820.2;AT3G19820.1 | cell elongation protein / DWARF1 / DIMINUTO (DIM) | 65 kDa | 0 | 0 | 13 | 11 | 24 | | | | | | | |
| AT1G14810 | AT1G14810.1;AT1G14810.2 | semialdehyde dehydrogenase family protein | 41 kDa | 2 | 5 | 13 | 10 | 30 | | 0.86 | 0.86 | 1.3E+07 | 2.2E+07 | 0.0E+00 | 3.5E+07 |
| AT1G51980 | AT1G51980.1;AT1G51980.2 | Insulinase (Peptidase family M16) protein | 54 kDa | 0 | 0 | 13 | 10 | 23 | | 0.20 | 0.20 | 4.4E+06 | 9.4E+06 | 0.0E+00 | 1.4E+07 |
| AT1G79730 | AT1G79730.1 | hydroxyproline-rich glycoprotein family protein | 67 kDa | 0 | 0 | 13 | 10 | 23 | | | | 1.6E+06 | 2.1E+06 | 4.5E+05 | 4.2E+06 |
| AT3G15010 | AT3G15010.2;AT3G15010.1 | RNA-binding (RRM/RBD/RNP motifs) family protein | 42 kDa | 0 | 2 | 13 | 10 | 25 | | 0.34 | 0.34 | 8.3E+06 | 1.0E+07 | 3.3E+06 | 2.2E+07 |
| AT2G17510 | | ribonuclease II family protein | 114 kDa | 0 | 0 | 13 | 9 | 22 | | 0.38 | 0.38 | | | | |
| AT2G19480 | | nucleosome assembly protein 1;2 | 44 kDa | 0 | 0 | 13 | 9 | 22 | | | | | | | |
| AT2G26510 | | Xanthine/uracil permease family protein | 60 kDa | 0 | 0 | 13 | 9 | 22 | | | | | | | |
| AT2G31740 | AT2G31740.1 | S-adenosyl-L-methionine-dependent methyltransferase | 85 kDa | 0 | 0 | 13 | 9 | 22 | | | | 1.0E+06 | 1.1E+06 | 3.7E+05 | 2.5E+06 |
| AT2G37860 | AT2G37860.4;AT2G37860.3;AT2G37860.2;AT2G37860.1 | Protein of unknown function (DUF3411) | 47 kDa | 0 | 0 | 13 | 9 | 22 | | 0.28 | 0.28 | 5.7E+05 | 7.2E+05 | 0.0E+00 | 1.3E+06 |
| AT3G07030 | | Alba DNA/RNA-binding protein | 43 kDa | 0 | 1 | 13 | 9 | 23 | | | | | | | |
| AT5G13280 | | aspartate kinase 1 | 62 kDa | 0 | 0 | 13 | 9 | 22 | | | | | | | |
| AT1G16240 | AT1G16240.3;AT1G16240.4;AT1G16240.2;AT1G16240.1 | syntaxin of plants 51 | 26 kDa | 0 | 2 | 13 | 8 | 23 | | | | | | | |
| AT3G03340 | AT3G03340.3;AT3G03340.1;AT3G03340.2 | LUC7 related protein | 47 kDa | 0 | 0 | 13 | 8 | 21 | | | | 1.6E+06 | 1.2E+06 | 6.8E+05 | 3.4E+06 |
| AT3G16060 | | ATP binding microtubule motor family protein | 77 kDa | 0 | 0 | 13 | 8 | 21 | | | | | | | |
| AT3G25680 | | FUNCTIONS IN: molecular_function unknown; INVOLVE | 63 kDa | 0 | 0 | 13 | 8 | 21 | | | | | | | |
| AT5G64200 | AT5G64200.2;AT5G64200.1 | ortholog of human splicing factor SC35 | 35 kDa | 0 | 3 | 13 | 8 | 24 | | 1.11 | 1.11 | 1.6E+07 | 1.2E+07 | 2.2E+06 | 3.0E+07 |
| AT2G17250 | | CCAAT-binding factor | 65 kDa | 0 | 0 | 13 | 7 | 20 | | | | | | | |
| AT2G20780 | | Major facilitator superfamily protein | 57 kDa | 0 | 0 | 13 | 7 | 20 | | | | | | | |
| AT3G05500 | | Rubber elongation factor protein (REF) | 27 kDa | 0 | 0 | 13 | 7 | 20 | | | | | | | |
| AT3G17350 | AT3G17350.2;AT3G17350.1 | unknown protein; FUNCTIONS IN: molecular_function | ? | 0 | 0 | 13 | 7 | 20 | | | | 0.0E+00 | 1.6E+06 | 0.0E+00 | 1.6E+06 |
| AT3G43240 | AT3G43240.1 | ARID/BRIGHT DNA-binding domain-containing protein | 83 kDa | 0 | 0 | 13 | 7 | 20 | | | | 6.3E+05 | 6.2E+05 | 2.3E+05 | 1.5E+06 |
| AT5G26860 | AT5G26860.1 | lon protease 1 | ? | 0 | 0 | 13 | 7 | 20 | | | | 5.8E+06 | 6.0E+06 | 9.3E+04 | 1.2E+07 |
| AT5G54830 | | DOMON domain-containing protein / dopamine beta-m | 101 kDa | 0 | 0 | 13 | 7 | 20 | | | | | | | |
| AT5G60300 | | Concanavalin A-like lectin protein kinase family protei | 79 kDa | 0 | 0 | 13 | 7 | 20 | | | | | | | |
| AT5G64640 | | Plant invertase/pectin methylesterase inhibitor superfa | 65 kDa | 0 | 0 | 13 | 7 | 20 | | | | | | | |
| AT1G07830 | AT1G07830.1 | ribosomal protein L29 family protein | 17 kDa | 2 | 0 | 13 | 6 | 21 | | 0.49 | 0.49 | 2.3E+07 | 3.1E+07 | 5.6E+06 | 6.0E+07 |
| AT1G17290 | | alanine aminotransferas | 60 kDa | 5 | 0 | 13 | 6 | 24 | | | | | | | |
| AT4G29830 | AT4G29830.1 | Transducin/WD40 repeat-like superfamily protein | 34 kDa | 0 | 1 | 13 | 6 | 20 | | 0.36 | 0.36 | 2.2E+07 | 3.3E+07 | 3.1E+06 | 5.8E+07 |
| AT5G18570 | | GTP1/OBG family protein | 76 kDa | 0 | 0 | 13 | 6 | 19 | | | | | | | |
| AT5G48030 | AT5G48030.1 | gametophytic factor 2 | 49 kDa | 0 | 1 | 13 | 6 | 20 | | | | 5.7E+06 | 7.5E+06 | 0.0E+00 | 1.3E+07 |
| AT5G53070 | AT5G53070.1 | Ribosomal protein L9/RNase H1 | 25 kDa | 0 | 1 | 13 | 6 | 20 | | | | 2.9E+07 | 4.0E+07 | 6.7E+06 | 7.6E+07 |
| AT5G60990 | AT5G60990.1 | DEA(D/H)-box RNA helicase family protein | 51 kDa | 0 | 0 | 13 | 6 | 19 | | | | 7.4E+06 | 1.7E+06 | 1.1E+06 | 1.0E+07 |
| ATCG00190 | | RNA polymerase subunit beta | 121 kDa | 0 | 0 | 13 | 6 | 19 | | | | | | | |
| AT2G03820 | AT2G03820.1 | nonsense-mediated mRNA decay NMD3 family protein | 59 kDa | 0 | 0 | 13 | 5 | 18 | | | | 1.1E+06 | 5.0E+05 | 0.0E+00 | 1.6E+06 |
| AT2G30620 | AT2G30620.1;AT2G30620.2 | winged-helix DNA-binding transcription factor family p | 28 kDa | 0 | 0 | 13 | 5 | 18 | | | | 8.8E+06 | 8.4E+06 | 7.5E+05 | 1.8E+07 |

| | | | | | | | | | | | | | | | |
|---|---|---|---|---|---|---|---|---|---|---|---|---|---|---|---|
| AT5G24850 | AT5G24850.1 | cryptochrome 3 | 60 kDa | 0 | 0 | 13 | 5 | 18 | | | | 1.2E+05 | 9.9E+04 | 0.0E+00 | 2.2E+05 |
| AT1G75660 | AT1G75660.1 | 5'-3' exoribonuclease 3 | 117 kDa | 0 | 0 | 13 | 4 | 17 | | | | 3.3E+05 | 2.1E+05 | 0.0E+00 | 5.4E+05 |
| AT2G24590 | AT2G24590.1 | RNA recognition motif and CCHC-type zinc finger doma | 22 kDa | 2 | 6 | 13 | 4 | 25 | | | | 1.9E+06 | 1.6E+06 | 0.0E+00 | 3.5E+06 |
| AT4G07410 | AT4G07410.1;AT4G07410.2 | Transducin family protein / WD-40 repeat family protei | 90 kDa | 0 | 0 | 13 | 4 | 17 | | | | 9.0E+06 | 6.4E+06 | 1.3E+06 | 1.7E+07 |
| AT4G23710 | AT4G23710.1 | vacuolar ATP synthase subunit G2 | 12 kDa | 2 | 0 | 13 | 4 | 19 | 0.37 | 0.69 | 0.53 | 7.7E+07 | 4.3E+07 | 4.1E+06 | 1.2E+08 |
| AT3G52560 | AT3G52560.1;AT2G36060.3;AT2G36060.1;AT3G52560.2;AT3G52560.4 | ubiquitin E2 variant 1D-4 | 17 kDa | 2 | 0 | 13 | 2 | 17 | | | | 1.8E+06 | 1.4E+06 | 0.0E+00 | 3.2E+06 |
| AT4G11010 | AT4G11010.1 | nucleoside diphosphate kinase 3 | 26 kDa | 3 | 0 | 13 | 2 | 18 | | 0.19 | 0.19 | 5.8E+07 | 7.8E+07 | 9.1E+06 | 1.5E+08 |
| AT5G12190 | AT5G12190.1 | RNA-binding (RRM/RBD/RNP motifs) family protein | 14 kDa | 2 | 0 | 13 | 2 | 17 | 0.64 | 0.21 | 0.42 | 6.3E+06 | 7.5E+06 | 1.4E+06 | 1.5E+07 |
| AT4G01560 | | Ribosomal RNA processing Brix domain protein | 40 kDa | 0 | 0 | 13 | 1 | 14 | | | | | | | |
| AT3G54720 | | Peptidase M28 family protein | 77 kDa | 0 | 0 | 12 | 31 | 43 | | | | | | | |
| AT3G63250 | AT3G63250.2;AT3G63250.3;AT3G63250.1 | homocysteine methyltransferase 2 | 36 kDa | 4 | 1 | 12 | 20 | 37 | | 2.69 | 2.69 | 7.6E+05 | 8.7E+05 | 1.6E+05 | 1.8E+06 |
| AT2G34300 | AT2G34300.3;AT2G34300.2;AT2G34300.1 | S-adenosyl-L-methionine-dependent methyltransferase | 87 kDa | 2 | 1 | 12 | 19 | 34 | | | | 0.0E+00 | 0.0E+00 | 3.9E+04 | 3.9E+04 |
| AT4G20440 | AT4G20440.5;AT4G20440.4;AT4G20440.3;AT4G20440.2;AT4G20440.1;AT5G44500.2;AT5G44500.1 | small nuclear ribonucleoprotein associated protein B | 27 kDa | 1 | 6 | 12 | 19 | 38 | | 0.87 | 0.87 | 1.7E+06 | 1.4E+07 | 4.1E+06 | 1.9E+07 |
| AT5G15350 | AT5G15350.1 | early nodulin-like protein 17 | 19 kDa | 2 | 7 | 12 | 18 | 39 | | | | 5.8E+06 | 6.3E+06 | 1.5E+06 | 1.4E+07 |
| AT1G66880 | | Protein kinase superfamily protein | 144 kDa | 0 | 1 | 12 | 16 | 29 | | | | | | | |
| AT3G03520 | | non-specific phospholipase C3 | 59 kDa | 0 | 1 | 12 | 16 | 29 | | | | | | | |
| AT5G17010 | AT5G17010.2;AT5G17010.4;AT5G17010.3;AT5G17010.1 | Major facilitator superfamily protein | 54 kDa | 0 | 0 | 12 | 16 | 28 | | | | 1.1E+06 | 0.0E+00 | 0.0E+00 | 1.1E+06 |
| AT1G61380 | | S-domain-1 29 | 90 kDa | 0 | 2 | 12 | 15 | 29 | | | | | | | |
| AT3G13080 | AT3G13080.2;AT3G13080.5;AT3G13080.1;AT3G13080.4;AT3G13080.3;AT3G13090.1 | multidrug resistance-associated protein 3 | 169 kDa | 1 | 2 | 12 | 15 | 30 | | | | 1.4E+05 | 7.8E+04 | 0.0E+00 | 2.1E+05 |
| AT5G08540 | AT5G08540.1 | unknown protein; FUNCTIONS IN: molecular_function | 39 kDa | 1 | 2 | 12 | 15 | 30 | | 0.25 | 0.25 | 9.0E+06 | 1.5E+07 | 2.5E+05 | 2.4E+07 |
| AT5G53530 | | vacuolar protein sorting 26A | 35 kDa | 0 | 1 | 12 | 15 | 28 | | | | | | | |
| AT1G47530 | | MATE efflux family protein | 52 kDa | 0 | 0 | 12 | 14 | 26 | | 0.21 | 0.21 | | | | |
| AT1G51940 | | protein kinase family protein / peptidoglycan-binding I | 71 kDa | 0 | 1 | 12 | 14 | 27 | | | | | | | |
| AT1G75270 | AT1G75270.1 | dehydroascorbate reductase 2 | 23 kDa | 2 | 5 | 12 | 13 | 32 | 3.78 | | 3.78 | 1.6E+06 | 3.5E+05 | 0.0E+00 | 2.0E+06 |
| AT2G39730 | AT2G39730.3;AT2G39730.2;AT2G39730.1 | rubisco activase | 52 kDa | 0 | 0 | 12 | 13 | 25 | | | | 6.8E+06 | 7.0E+06 | 0.0E+00 | 1.4E+07 |
| AT5G06700 | | Plant protein of unknown function (DUF828) | 68 kDa | 0 | 0 | 12 | 13 | 25 | | | | | | | |
| AT5G40370 | AT5G40370.1;AT5G40370.2 | Glutaredoxin family protein | 12 kDa | 1 | 0 | 12 | 13 | 26 | | 2.98 | 2.98 | 1.7E+07 | 8.7E+06 | 0.0E+00 | 2.6E+07 |
| AT1G09160 | AT1G09160.1;AT1G09160.2;AT1G47380.1 | Protein phosphatase 2C family protein | 46 kDa | 0 | 0 | 12 | 12 | 24 | | | | 4.0E+05 | 3.2E+05 | 0.0E+00 | 7.3E+05 |
| AT1G74850 | | plastid transcriptionally active 2 | 96 kDa | 0 | 0 | 12 | 12 | 24 | | | | | | | |
| AT2G33040 | AT2G33040.1 | gamma subunit of Mt ATP synthase | 35 kDa | 0 | 3 | 12 | 12 | 27 | | 0.46 | 0.46 | 3.8E+06 | 6.0E+06 | 1.9E+06 | 1.2E+07 |
| AT3G17940 | | Galactose mutarotase-like superfamily protein | 37 kDa | 1 | 0 | 12 | 12 | 25 | | | | | | | |
| AT4G02650 | | ENTH/ANTH/VHS superfamily protein | 68 kDa | 0 | 0 | 12 | 12 | 24 | | | | | | | |
| AT4G27680 | | P-loop containing nucleoside triphosphate hydrolases | 44 kDa | 0 | 10 | 12 | 12 | 34 | | | | | | | |
| AT5G26340 | AT5G26340.1 | Major facilitator superfamily protein | 57 kDa | 2 | 1 | 12 | 12 | 27 | | | | 2.0E+07 | 1.7E+07 | 0.0E+00 | 3.7E+07 |
| AT1G08450 | AT1G08450.2;AT1G08450.3;AT1G08450.1 | calreticulin 3 | 50 kDa | 0 | 0 | 12 | 11 | 23 | | | | 2.1E+06 | 2.0E+06 | 3.0E+05 | 4.5E+06 |
| AT2G17130 | | isocitrate dehydrogenase subunit 2 | 40 kDa | 0 | 0 | 12 | 11 | 23 | | | | | | | |
| AT2G40316 | | FUNCTIONS IN: molecular_function unknown; LOCATEI | 30 kDa | 0 | 2 | 12 | 11 | 25 | | | | | | | |
| AT4G23640 | | Potassium transporter family protein | 87 kDa | 0 | 0 | 12 | 11 | 23 | | | | | | | |
| AT5G40260 | AT5G40260.1;AT5G40260.2 | Nodulin MtN3 family protein | 27 kDa | 4 | 2 | 12 | 11 | 29 | | | | 2.8E+07 | 2.0E+07 | 0.0E+00 | 4.8E+07 |
| AT5G56890 | | Protein kinase superfamily protein | 119 kDa | 0 | 0 | 12 | 11 | 23 | | | | | | | |
| AT1G31440 | AT1G31440.1 | SH3 domain-containing protein | 49 kDa | 0 | 3 | 12 | 10 | 25 | | | | 8.8E+04 | 7.4E+04 | 0.0E+00 | 1.6E+05 |
| AT3G15980 | AT3G15980.1;AT3G15980.7;AT3G15980.6;AT3G15980.4;AT3G15980.3;AT3G15980.2 | Coatomer, beta' subunit | 102 kDa | 0 | 0 | 12 | 10 | 22 | | | | 2.2E+05 | 2.3E+05 | 0.0E+00 | 4.5E+05 |
| AT5G34850 | | purple acid phosphatase 26 | 55 kDa | 0 | 0 | 12 | 10 | 22 | | | | | | | |
| AT5G39410 | | Saccharopine dehydrogenase | 50 kDa | 0 | 0 | 12 | 10 | 22 | | | | | | | |
| AT1G28440 | AT1G28440.1 | HAESA-like 1 | 109 kDa | 0 | 0 | 12 | 9 | 21 | | | | 2.4E+05 | 2.5E+05 | 0.0E+00 | 5.0E+05 |
| AT1G64040 | AT1G64040.1 | type one serine/threonine protein phosphatase 3 | 36 kDa | 0 | 0 | 12 | 9 | 21 | | | | 4.8E+06 | 4.3E+06 | 5.3E+05 | 9.6E+06 |
| AT1G10390 | | Nucleoporin autopeptidase | 106 kDa | 0 | 0 | 12 | 8 | 20 | | | | | | | |
| AT4G28400 | | Protein phosphatase 2C family protein | 31 kDa | 0 | 9 | 12 | 8 | 29 | | 2.73 | 2.73 | | | | |
| AT4G33700 | | CBS domain-containing protein with a domain of unkn | 47 kDa | 0 | 0 | 12 | 8 | 20 | | | | | | | |
| AT5G66860 | AT5G66860.1 | Ribosomal protein L25/Gln-tRNA synthetase, anti-codo | 27 kDa | 0 | 0 | 12 | 8 | 20 | | | | 1.2E+07 | 1.7E+07 | 2.5E+06 | 3.2E+07 |
| ATCG00180 | | DNA-directed RNA polymerase family protein | 79 kDa | 0 | 0 | 12 | 8 | 20 | | | | | | | |
| AT4G09320 | AT4G09320.1 | Nucleoside diphosphate kinase family protein | 19 kDa | 6 | 0 | 12 | 7 | 25 | | 3.93 | 3.93 | 1.2E+07 | 6.3E+06 | 0.0E+00 | 1.8E+07 |
| AT1G07630 | | pol-like 5 | 74 kDa | 0 | 0 | 12 | 6 | 18 | | | | | | | |
| AT1G77750 | AT1G77750.1 | Ribosomal protein S13/S18 family | 17 kDa | 6 | 0 | 12 | 6 | 24 | | 1.26 | 1.26 | 5.4E+07 | 8.6E+07 | 1.3E+07 | 1.5E+08 |
| AT2G34970 | | Trimeric LpxA-like enzyme | 82 kDa | 0 | 0 | 12 | 6 | 18 | | | | | | | |
| AT2G20800 | | NAD(P)H dehydrogenase B4 | 65 kDa | 0 | 0 | 12 | 5 | 17 | | | | | | | |
| AT2G41800 | AT2G41800.1 | Protein of unknown function, DUF642 | 40 kDa | 0 | 0 | 12 | 5 | 17 | | | | | | | |
| AT4G37880 | AT4G37880.1 | LisH/CRA/RING-U-box domains-containing protein | 44 kDa | 0 | 0 | 12 | 5 | 17 | | | | 4.1E+06 | 4.9E+06 | 1.8E+05 | 9.2E+06 |
| AT4G38130 | AT4G38130.1;AT4G38130.2 | histone deacetylase 1 | 56 kDa | 0 | 1 | 12 | 5 | 18 | | 0.35 | 0.35 | 6.6E+05 | 1.7E+06 | 6.3E+05 | 2.9E+06 |
| AT5G09770 | AT5G09770.1;AT5G64650.1 | Ribosomal protein L17 family protein | 18 kDa | 0 | 0 | 12 | 5 | 17 | | | | 1.9E+07 | 2.1E+07 | 1.8E+06 | 4.2E+07 |
| AT5G19620 | | outer envelope protein of 80 kDa | 80 kDa | 0 | 0 | 12 | 5 | 17 | | | | | | | |
| AT5G23535 | AT5G23535.2;AT5G23535.1 | KOW domain-containing protein | 17 kDa | 0 | 0 | 12 | 5 | 17 | | | | 5.8E+07 | 7.8E+07 | 1.5E+07 | 1.5E+08 |
| AT5G56020 | | Got1/Sft2-like vescicle transport protein family | 25 kDa | 0 | 0 | 12 | 5 | 17 | | | | | | | |
| AT1G11960 | | ERD (early-responsive to dehydration stress) family prot | 88 kDa | 0 | 0 | 12 | 4 | 16 | | | | | | | |
| AT2G07050 | | cycloartenol synthase 1 | 86 kDa | 0 | 0 | 12 | 4 | 16 | | | | | | | |
| AT3G11450 | AT3G11450.1 | DnaJ domain ;Myb-like DNA-binding domain | 74 kDa | 0 | 0 | 12 | 4 | 16 | | | | 5.9E+06 | 5.5E+06 | 5.2E+04 | 1.1E+07 |
| AT5G05680 | | nuclear pore complex protein-related | 90 kDa | 0 | 0 | 12 | 4 | 16 | | | | | | | |
| AT5G15020 | | SIN3-like 2 | 155 kDa | 0 | 0 | 12 | 4 | 16 | | | | | | | |
| AT5G39950 | AT5G39950.1 | thioredoxin 2 | 15 kDa | 0 | 0 | 12 | 4 | 16 | | 0.31 | 0.31 | 1.4E+07 | 9.5E+06 | 6.6E+05 | 2.4E+07 |
| AT5G49210 | AT5G49210.1;AT5G49210.2 | unknown protein; Has 3675 Blast hits to 2315 proteins | 24 kDa | 0 | 1 | 12 | 4 | 17 | | | | 6.4E+07 | 8.0E+07 | 6.0E+06 | 1.5E+08 |
| AT2G24500 | AT2G24500.1 | Zinc finger protein 622 | 45 kDa | 0 | 0 | 12 | 3 | 15 | | | | 0.0E+00 | 0.0E+00 | 1.2E+06 | 1.2E+06 |
| AT2G25170 | AT2G25170.4;AT2G25170.1;AT2G25170.2;AT2G25170.3 | chromatin remodeling factor CHD3 (PICKLE) | 158 kDa | 0 | 0 | 12 | 3 | 15 | | | | 4.7E+04 | 7.9E+05 | 0.0E+00 | 8.4E+05 |
| AT5G16660 | AT5G16660.2;AT5G16660.1 | unknown protein; FUNCTIONS IN: molecular_function | 18 kDa | 2 | 0 | 12 | 3 | 17 | 0.12 | 0.13 | 0.12 | 1.0E+07 | 7.0E+06 | 0.0E+00 | 1.7E+07 |
| AT2G45640 | AT2G45640.1;AT2G45640.2 | SIN3 associated polypeptide P18 | 17 kDa | 0 | 0 | 12 | 2 | 14 | | | | 1.8E+06 | 1.3E+06 | 9.0E+04 | 3.1E+06 |
| AT4G29520 | AT4G29520.1 | LOCATED IN: endoplasmic reticulum, plasma membran | 34 kDa | 0 | 0 | 12 | 2 | 14 | | | | 1.6E+06 | 1.6E+06 | 0.0E+00 | 3.2E+06 |
| AT5G15680 | | ARM repeat superfamily protein | 243 kDa | 0 | 0 | 11 | 23 | 34 | | | | | | | |
| AT4G35850 | AT4G35850.1 | Pentatricopeptide repeat (PPR) superfamily protein | 50 kDa | 1 | 6 | 11 | 18 | 36 | | | | 8.9E+05 | 8.1E+05 | 5.1E+04 | 1.7E+06 |
| AT3G21190 | | O-fucosyltransferase family protein | 47 kDa | 0 | 3 | 11 | 17 | 31 | | | | | | | |
| AT3G54470 | AT3G54470.1 | uridine 5'-monophosphate synthase / UMP synthase (P' | 52 kDa | 0 | 0 | 11 | 17 | 28 | | | | 1.5E+07 | 1.4E+07 | 4.6E+05 | 3.0E+07 |
| AT4G26900 | AT4G26900.1 | HIS HF | 64 kDa | 0 | 3 | 11 | 17 | 31 | | 0.91 | 0.91 | 1.0E+06 | 3.2E+05 | 0.0E+00 | 1.4E+06 |
| AT2G31960 | | glucan synthase-like 3 | 226 kDa | 0 | 0 | 11 | 16 | 27 | | | | | | | |
| AT4G27730 | | oligopeptide transporter 1 | 82 kDa | 0 | 0 | 11 | 16 | 27 | | | | | | | |
| AT1G60710 | | NAD(P)-linked oxidoreductase superfamily protein | 38 kDa | 0 | 2 | 11 | 15 | 28 | | | | | | | |

| | | | | | | | | | | | | | | | |
|---|---|---|---|---|---|---|---|---|---|---|---|---|---|---|---|
| AT5G02800 | | Protein kinase superfamily protein | 42 kDa | 0 | 3 | 11 | 15 | 29 | | | | | | | |
| AT5G13760 | | Plasma-membrane choline transporter family protein | 64 kDa | 0 | 0 | 11 | 15 | 26 | | | | | | | |
| AT1G06670 | AT1G06670.2;AT1G06670.1 | nuclear DEIH-boxhelicase | 175 kDa | 0 | 0 | 11 | 14 | 25 | | | | 7.8E+05 | 7.1E+05 | 4.5E+05 | 1.9E+06 |
| AT3G11560 | AT3G11560.5;AT3G11560.1;AT3G11560.4;AT3G11560.3;AT3G11560.2 | LETM1-like protein | 98 kDa | 0 | 0 | 11 | 14 | 25 | | 0.43 | 0.43 | 6.4E+05 | 0.0E+00 | 0.0E+00 | 6.4E+05 |
| AT3G52500 | AT3G52500.1 | Eukaryotic aspartyl protease family protein | 51 kDa | 0 | 4 | 11 | 14 | 29 | | | | 2.2E+07 | 1.4E+07 | 1.3E+06 | 3.7E+07 |
| AT5G05210 | AT5G05210.2;AT5G05210.1 | Surfeit locus protein 6 | 45 kDa | 0 | 0 | 11 | 14 | 25 | | | | 3.5E+06 | 5.1E+06 | 7.7E+05 | 9.3E+06 |
| AT5G49760 | AT5G49760.1 | Leucine-rich repeat protein kinase family protein | 105 kDa | 0 | 0 | 11 | 14 | 25 | | | | 1.9E+05 | 1.3E+05 | 0.0E+00 | 3.3E+05 |
| AT1G65320 | | Cystathionine beta-synthase (CBS) family protein | 47 kDa | 0 | 0 | 11 | 13 | 24 | | | | | | | |
| AT3G06480 | AT3G06480.2;AT3G06480.1 | DEAD box RNA helicase family protein | 120 kDa | 1 | 0 | 11 | 13 | 25 | | | | 0.0E+00 | 0.0E+00 | 1.6E+05 | 1.6E+05 |
| AT3G62770 | | Transducin/WD40 repeat-like superfamily protein | 43 kDa | 0 | 0 | 11 | 13 | 24 | | | | | | | |
| AT5G10540 | | Zincin-like metalloproteases family protein | 79 kDa | 1 | 0 | 11 | 13 | 25 | | | | | | | |
| AT1G35220 | | unknown protein; FUNCTIONS IN: molecular_function | 111 kDa | 0 | 0 | 11 | 12 | 23 | | | | | | | |
| AT1G47380 | | Protein phosphatase 2C family protein | 46 kDa | 1 | 1 | 11 | 12 | 25 | | | | | | | |
| AT3G62830 | | NAD(P)-binding Rossmann-fold superfamily protein | 50 kDa | 0 | 0 | 11 | 12 | 23 | | | | | | | |
| AT5G24380 | | YELLOW STRIPE like 2 | 73 kDa | 1 | 0 | 11 | 12 | 24 | | | | | | | |
| AT1G05940 | | cationic amino acid transporter 9 | 60 kDa | 0 | 0 | 11 | 11 | 22 | | | | | | | |
| AT1G11300 | | protein serine/threonine kinases;protein kinases;ATP b | 184 kDa | 2 | 2 | 11 | 11 | 26 | | | | | | | |
| AT1G21690 | AT1G21690.2;AT1G21690.4;AT1G21690.1;AT1G21690.3 | ATPase family associated with various cellular activities | 38 kDa | 0 | 0 | 11 | 11 | 22 | | | | 7.8E+05 | 1.2E+06 | 4.3E+05 | 2.4E+06 |
| AT2G23820 | AT2G23820.2;AT2G23820.1 | Metal-dependent phosphohydrolase | 28 kDa | 0 | 11 | 11 | 11 | 33 | | | | 3.0E+07 | 1.9E+07 | 0.0E+00 | 4.9E+07 |
| AT3G10350 | | P-loop containing nucleoside triphosphate hydrolases | 45 kDa | 0 | 3 | 11 | 10 | 24 | | | | | | | |
| AT3G22960 | | Pyruvate kinase family protein | 65 kDa | 0 | 0 | 11 | 10 | 21 | | | | | | | |
| AT5G51660 | AT5G51660.1;AT5G51660.4;AT5G51660.2;AT5G51660.3 | cleavage and polyadenylation specificity factor 160 | 158 kDa | 0 | 0 | 11 | 10 | 21 | | | | 1.5E+06 | 6.5E+05 | 3.1E+05 | 2.4E+06 |
| AT1G17470 | AT1G17470.2;AT1G17470.1 | developmentally regulated G-protein 1 | 45 kDa | 0 | 0 | 11 | 9 | 20 | | | | 6.8E+05 | 8.4E+05 | 2.8E+05 | 1.8E+06 |
| AT1G22610 | | C2 calcium/lipid-binding plant phosphoribosyltransfer | 118 kDa | 3 | 0 | 11 | 9 | 23 | | | | | | | |
| AT1G52370 | AT1G52370.4;AT1G52370.3;AT1G52370.2;AT1G52370.1;AT4G28360.1 | Ribosomal protein L22p/L17e family protein | 30 kDa | 0 | 2 | 11 | 9 | 22 | | | | 9.1E+06 | 1.7E+07 | 5.7E+05 | 2.7E+07 |
| AT1G73390 | | Endosomal targeting BRO1-like domain-containing pro | 47 kDa | 0 | 0 | 11 | 9 | 20 | | | | | | | |
| AT1G73640 | | RAB GTPase homolog A6A | 26 kDa | 0 | 11 | 11 | 9 | 31 | | | | | | | |
| AT2G18510 | AT2G18510.1 | RNA-binding (RRM/RBD/RNP motifs) family protein | 40 kDa | 0 | 1 | 11 | 9 | 21 | | 0.66 | 0.66 | 7.5E+06 | 7.2E+06 | 1.1E+06 | 1.6E+07 |
| AT3G54290 | | FUNCTIONS IN: molecular_function unknown; INVOLVE | 39 kDa | 1 | 0 | 11 | 9 | 21 | | | | | | | |
| AT3G54560 | AT3G54560.2;AT3G54560.1;AT2G38810.4;AT2G38810.3;AT2G38810.2;AT2G38810.1 | histone H2A 11 | 15 kDa | 3 | 0 | 11 | 9 | 23 | | | | 4.9E+07 | 3.6E+07 | 1.6E+07 | 1.0E+08 |
| AT1G02180 | | ferredoxin-related | 25 kDa | 2 | 4 | 11 | 8 | 25 | 2.92 | 5.02 | 3.97 | | | | |
| AT1G10670 | AT1G10670.4;AT1G10670.2;AT1G10670.1;AT1G10670.3;AT1G60810.2;AT1G60810.1 | ATP-citrate lyase A-1 | 47 kDa | 0 | 0 | 11 | 8 | 19 | | | | | | | |
| AT1G63810 | AT1G63810.2;AT1G63810.1 | CONTAINS InterPro DOMAIN/s: Nrap protein (InterPro:I | 120 kDa | 0 | 0 | 11 | 8 | 19 | | 0.25 | 0.25 | 0.0E+00 | 0.0E+00 | 3.0E+05 | 3.0E+05 |
| AT1G66980 | | suppressor of npr1-1 constitutive 4 | 124 kDa | 0 | 2 | 11 | 8 | 21 | | | | | | | |
| AT2G19130 | | S-locus lectin protein kinase family protein | 92 kDa | 0 | 1 | 11 | 8 | 20 | | | | | | | |
| AT2G34260 | AT2G34260.2;AT2G34260.1 | transducin family protein / WD-40 repeat family protei | 38 kDa | 0 | 0 | 11 | 8 | 19 | | | | 4.6E+06 | 5.1E+06 | 0.0E+00 | 9.7E+06 |
| AT3G06510 | | Glycosyl hydrolase superfamily protein | 71 kDa | 0 | 0 | 11 | 8 | 19 | | | | | | | |
| AT4G13780 | AT4G13780.1 | methionine--tRNA ligase, putative / methionyl-tRNA syı | ? | 0 | 0 | 11 | 8 | 19 | | | | 9.9E+05 | 6.4E+05 | 0.0E+00 | 1.6E+06 |
| AT5G14420 | AT5G14420.4;AT5G14420.3;AT5G14420.2;AT5G14420.1 | RING domain ligase2 | 52 kDa | 0 | 0 | 11 | 8 | 19 | | | | 1.2E+06 | 1.1E+06 | 0.0E+00 | 2.3E+06 |
| AT5G48370 | AT5G48370.1;AT5G48370.2;AT5G48370.3 | Thioesterase/thiol ester dehydrase-isomerase superfam | 49 kDa | 0 | 0 | 11 | 8 | 19 | | 0.32 | 0.32 | 8.8E+05 | 7.8E+05 | 0.0E+00 | 1.7E+06 |
| AT1G07910 | AT1G07910.1;AT1G07910.2 | RNAligase | 123 kDa | 0 | 0 | 11 | 7 | 18 | | | | 1.3E+05 | 3.2E+05 | 0.0E+00 | 4.5E+05 |
| AT3G08580 | AT3G08580.2;AT3G08580.1 | ADP/ATP carrier 1 | 41 kDa | 5 | 0 | 11 | 7 | 23 | | 0.13 | 0.13 | 2.3E+06 | 3.9E+06 | 0.0E+00 | 6.2E+06 |
| AT3G17450 | | hAT dimerisation domain-containing protein | 101 kDa | 0 | 0 | 11 | 7 | 18 | | | | | | | |
| AT4G31880 | | LOCATED IN: cytosol, chloroplast; EXPRESSED IN: 24 pla | 94 kDa | 0 | 0 | 11 | 7 | 18 | | 0.10 | 0.10 | | | | |
| AT5G09260 | AT5G09260.1 | vacuolar protein sorting-associated protein 20.2 | 25 kDa | 4 | 1 | 11 | 7 | 23 | | | | 3.4E+06 | 0.0E+00 | 0.0E+00 | 3.4E+06 |
| AT5G16290 | AT5G16290.2;AT5G16290.1 | VALINE-TOLERANT 1 | 52 kDa | 0 | 0 | 11 | 7 | 18 | | | | 3.7E+06 | 2.3E+06 | 1.6E+04 | 6.0E+06 |
| AT5G22050 | | Protein kinase superfamily protein | 34 kDa | 1 | 2 | 11 | 7 | 21 | | | | | | | |
| AT1G18150 | AT1G18150.3;AT1G18150.2;AT1G18150.1 | Protein kinase superfamily protein | 66 kDa | 0 | 0 | 11 | 6 | 17 | | | | 9.9E+04 | 0.0E+00 | 0.0E+00 | 9.9E+04 |
| AT1G49040 | | stomatal cytokinesis defective / SCD1 protein (SCD1) | 132 kDa | 2 | 0 | 11 | 6 | 19 | | | | | | | |
| AT2G17480 | | Seven transmembrane MLO family protein | 67 kDa | 0 | 0 | 11 | 6 | 17 | | | | | | | |
| AT3G61860 | AT3G61860.1 | RNA-binding (RRM/RBD/RNP motifs) family protein | 31 kDa | 1 | 10 | 11 | 6 | 28 | 0.13 | 3.10 | 1.62 | 1.5E+07 | 2.0E+07 | 4.2E+05 | 3.6E+07 |
| AT5G17440 | AT5G17440.1;AT5G17440.2 | LUC7 related protein | 47 kDa | 0 | 0 | 11 | 6 | 17 | | | | 1.9E+06 | 3.5E+06 | 0.0E+00 | 5.4E+06 |
| AT5G50200 | | nitrate transmembrane transporters | 23 kDa | 0 | 0 | 11 | 6 | 17 | | | | | | | |
| AT5G65720 | AT5G65720.1;AT5G65720.3;AT5G65720.2 | nitrogen fixation S (NIFS)-like 1 | 50 kDa | 0 | 0 | 11 | 6 | 17 | | | | 6.5E+06 | 7.4E+06 | 3.3E+04 | 1.4E+07 |
| AT1G01300 | AT1G01300.1 | Eukaryotic aspartyl protease family protein | 52 kDa | 0 | 0 | 11 | 5 | 16 | | | | | | | |
| AT1G27500 | | Tetratricopeptide repeat (TPR)-like superfamily protein | 71 kDa | 0 | 0 | 11 | 5 | 16 | | | | | | | |
| AT3G63400 | AT3G63400.2;AT3G63400.4;AT3G63400.3;AT3G63400.1 | Cyclophilin-like peptidyl-prolyl cis-trans isomerase fam | 64 kDa | 0 | 0 | 11 | 5 | 16 | | | | 7.2E+06 | 9.6E+06 | 4.9E+05 | 1.7E+07 |
| AT4G23540 | AT4G23540.1 | ARM repeat superfamily protein | 123 kDa | 0 | 0 | 11 | 5 | 16 | | | | 1.4E+05 | 7.6E+04 | 4.4E+05 | 6.6E+05 |
| AT5G42150 | AT5G42150.1 | Glutathione S-transferase family protein | 36 kDa | 2 | 4 | 11 | 5 | 22 | 0.06 | | 0.06 | 2.0E+06 | 2.1E+06 | 1.7E+05 | 4.3E+06 |
| AT5G58060 | | SNARE-like superfamily protein | 23 kDa | 3 | 2 | 11 | 5 | 21 | | | | | | | |
| ATCG00790 | ATCG00790.1 | ribosomal protein L16 | 15 kDa | 0 | 0 | 11 | 5 | 16 | | 0.37 | 0.37 | 8.8E+06 | 2.8E+06 | 4.4E+06 | 1.6E+07 |
| AT1G15810 | AT1G15810.1 | S15/NS1, RNA-binding protein | 47 kDa | 0 | 0 | 11 | 4 | 15 | | | | 1.7E+07 | 2.5E+07 | 1.6E+06 | 4.3E+07 |
| AT3G04580 | | Signal transduction histidine kinase, hybrid-type, ethyl | 86 kDa | 0 | 0 | 11 | 4 | 15 | | | | | | | |
| AT3G26560 | AT3G26560.1 | ATP-dependent RNA helicase, putative | 134 kDa | 0 | 0 | 11 | 4 | 15 | | | | 2.0E+05 | 1.9E+05 | 4.7E+04 | 4.4E+05 |
| AT3G53760 | | GAMMA-TUBULIN COMPLEX PROTEIN 4 | 86 kDa | 0 | 0 | 11 | 4 | 15 | | | | | | | |
| AT4G28050 | AT4G28050.1 | tetraspanin7 | 30 kDa | 2 | 16 | 11 | 4 | 33 | | 9.10 | 9.10 | 4.6E+06 | 3.6E+06 | 0.0E+00 | 8.2E+06 |
| AT4G28450 | AT4G28450.1 | nucleotide binding;protein binding | 51 kDa | 0 | 0 | 11 | 4 | 15 | | | | 2.0E+06 | 2.2E+06 | 5.2E+05 | 4.7E+06 |
| AT4G29680 | | Alkaline-phosphatase-like family protein | 55 kDa | 0 | 0 | 11 | 4 | 15 | | | | | | | |
| AT5G13780 | AT5G13780.1 | Acyl-CoA N-acyltransferases (NAT) superfamily protein | 22 kDa | 0 | 0 | 11 | 4 | 15 | | 0.06 | 0.06 | 2.6E+07 | 1.3E+07 | 3.5E+06 | 4.3E+07 |
| AT2G23420 | AT2G23420.2;AT2G23420.1 | nicotinate phosphoribosyltransferase 2 | 62 kDa | 0 | 0 | 11 | 3 | 14 | | | | 4.6E+06 | 2.0E+06 | 0.0E+00 | 6.6E+06 |
| AT2G47640 | | Small nuclear ribonucleoprotein family protein | 13 kDa | 3 | 0 | 11 | 3 | 17 | | | | | | | |
| AT3G01790 | AT3G01790.1;AT3G01790.2;AT3G01790.3 | Ribosomal protein L13 family protein | 23 kDa | 0 | 3 | 11 | 3 | 17 | | | | 1.0E+07 | 1.3E+07 | 1.2E+06 | 2.4E+07 |
| AT3G51250 | AT3G51250.2;AT3G51250.1 | Senescence/dehydration-associated protein-related | 50 kDa | 0 | 1 | 11 | 3 | 15 | | | | 9.0E+05 | 1.7E+06 | 0.0E+00 | 2.6E+06 |
| AT5G11110 | AT5G11110.1 | sucrose phosphate synthase 2F | 117 kDa | 0 | 0 | 11 | 3 | 14 | | | | 2.8E+06 | 3.5E+06 | 0.0E+00 | 6.3E+06 |
| AT2G20490 | AT2G20490.1;AT2G20490.3;AT2G20490.2 | nucleolar RNA-binding Nop10p family protein | 7 kDa | 0 | 0 | 11 | 2 | 13 | 0.07 | 0.15 | 0.11 | 2.4E+07 | 1.0E+08 | 2.4E+07 | 1.5E+08 |
| AT3G47810 | AT3G47810.3;AT3G47810.2;AT3G47810.1 | Calcineurin-like metallo-phosphoesterase superfamily ( | 21 kDa | 0 | 7 | 11 | 2 | 20 | | | | 1.2E+07 | 1.0E+07 | 9.8E+05 | 2.3E+07 |
| ATCG00820 | | ribosomal protein S19 | 11 kDa | 0 | 0 | 11 | 2 | 13 | | 0.45 | 0.45 | | | | |
| AT1G16890 | | ubiquitin-conjugating enzyme 36 | 17 kDa | 2 | 0 | 11 | 1 | 14 | | | | | | | |
| AT3G21465 | AT3G21465.1;AT3G21465.2 | unknown protein; BEST Arabidopsis thaliana protein m | ? | 0 | 2 | 11 | 1 | 14 | | | | 1.2E+06 | 2.3E+06 | 1.7E+05 | 3.7E+06 |
| AT4G27640 | AT4G27640.1 | ARM repeat superfamily protein | 115 kDa | 0 | 1 | 11 | 1 | 13 | | | | | | | |
| AT5G03730 | | Protein kinase superfamily protein | 90 kDa | 0 | 0 | 11 | 1 | 12 | | | | | | | |

| | | | | | | | | | | | | | | | |
|---|---|---|---|---|---|---|---|---|---|---|---|---|---|---|---|
| AT2G31390 | | pfkB-like carbohydrate kinase family protein | 35 kDa | 0 | 0 | 10 | 24 | 34 | | | | | | | |
| AT4G39420 | | unknown protein; INVOLVED IN: biological_process un | 355 kDa | 0 | 0 | 10 | 22 | 32 | | | | | | | |
| AT1G73080 | AT1G73080.1 | PEP1 receptor 1 | 123 kDa | 0 | 0 | 10 | 21 | 31 | | | | 1.3E+06 | 5.8E+05 | 0.0E+00 | 1.9E+06 |
| AT4G27520 | AT4G27520.1 | early nodulin-like protein 2 | 35 kDa | 0 | 0 | 10 | 20 | 30 | | | | 1.8E+07 | 1.2E+07 | 0.0E+00 | 3.0E+07 |
| AT5G05010 | AT5G05010.2;AT5G05010.1 | clathrin adaptor complexes medium subunit family prc | 58 kDa | 0 | 1 | 10 | 20 | 31 | | | | 1.4E+06 | 1.6E+06 | 2.3E+05 | 3.2E+06 |
| AT3G15220 | | Protein kinase superfamily protein | 76 kDa | 0 | 0 | 10 | 19 | 29 | | | | | | | |
| AT2G47960 | AT2G47960.1 | unknown protein; CONTAINS InterPro DOMAIN/s: Prot | 49 kDa | 0 | 0 | 10 | 18 | 28 | | | | 1.0E+06 | 1.5E+06 | 2.4E+05 | 2.8E+06 |
| AT1G67120 | AT1G67120.1;AT1G67120.2 | ATPases;nucleotide binding;ATP binding;nucleoside-tri | 611 kDa | 0 | 0 | 10 | 16 | 26 | | 0.18 | 0.18 | 5.0E+04 | 0.0E+00 | 9.3E+02 | 5.1E+04 |
| AT1G60620 | AT1G60620.2;AT1G60620.1 | RNA polymerase I subunit 43 | 43 kDa | 0 | 0 | 10 | 15 | 25 | | | | 6.3E+05 | 6.4E+05 | 1.7E+05 | 1.4E+06 |
| AT4G22130 | | STRUBBELIG-receptor family 8 | 76 kDa | 0 | 0 | 10 | 15 | 25 | | | | | | | |
| AT5G03290 | AT5G03290.1 | isocitrate dehydrogenase V | 41 kDa | 0 | 3 | 10 | 15 | 28 | | | | 3.2E+06 | 6.6E+06 | 0.0E+00 | 9.8E+06 |
| AT5G65950 | AT5G65950.1 | FUNCTIONS IN: molecular_function unknown; INVOLVE | 132 kDa | 0 | 0 | 10 | 15 | 25 | | | | 2.2E+05 | 2.5E+05 | 0.0E+00 | 4.7E+05 |
| AT2G42010 | | phospholipase D beta 1 | 121 kDa | 0 | 0 | 10 | 14 | 24 | | | | | | | |
| AT4G22570 | AT4G22570.1;AT4G22570.2 | adenine phosphoribosyl transferase 3 | 20 kDa | 1 | 6 | 10 | 14 | 31 | | 3.72 | 3.72 | 1.5E+07 | 9.1E+06 | 2.6E+06 | 2.7E+07 |
| AT5G14250 | | Proteasome component (PCI) domain protein | 48 kDa | 0 | 0 | 10 | 14 | 24 | | | | | | | |
| AT5G16990 | | Zinc-binding dehydrogenase family protein | 38 kDa | 0 | 0 | 10 | 14 | 24 | | | | | | | |
| AT5G57020 | AT5G57020.1 | myristoyl-CoA:protein N-myristoyltransferase | 50 kDa | 0 | 0 | 10 | 14 | 24 | | | | 1.4E+07 | 1.5E+07 | 1.3E+06 | 3.1E+07 |
| AT1G54115 | | cation calcium exchanger 4 | 71 kDa | 2 | 5 | 10 | 12 | 29 | | | | | | | |
| AT1G58030 | | cationic amino acid transporter 2 | 67 kDa | 0 | 0 | 10 | 12 | 22 | | | | | | | |
| AT2G17800 | | Arabidopsis RAC-like 1 | 22 kDa | 3 | 4 | 10 | 12 | 29 | | | | | | | |
| AT3G10690 | AT3G10690.1 | DNA GYRASE A | 105 kDa | 0 | 0 | 10 | 12 | 22 | | | | 1.2E+05 | 0.0E+00 | 8.4E+04 | 2.0E+05 |
| AT4G09750 | | NAD(P)-binding Rossmann-fold superfamily protein | 35 kDa | 0 | 0 | 10 | 12 | 22 | | | | | | | |
| AT4G37910 | AT4G37910.2;AT4G37910.1 | mitochondrial heat shock protein 70-1 | 73 kDa | 0 | 2 | 10 | 12 | 24 | | | | 1.6E+06 | 1.9E+06 | 4.1E+05 | 3.9E+06 |
| AT1G30690 | | Sec14p-like phosphatidylinositol transfer family protei | 61 kDa | 0 | 0 | 10 | 11 | 21 | | | | | | | |
| AT1G55310 | AT1G55310.1;AT1G55310.3;AT1G55310.5;AT1G55310.2 | SC35-like splicing factor 33 | 33 kDa | 0 | 1 | 10 | 11 | 22 | | | | 7.9E+06 | 8.5E+06 | 2.0E+06 | 1.8E+07 |
| AT5G23200 | | unknown protein; BEST Arabidopsis thaliana protein m | ? | 0 | 0 | 10 | 11 | 21 | | | | | | | |
| AT5G26740 | | Protein of unknown function (DUF300) | 47 kDa | 0 | 1 | 10 | 11 | 22 | | | | | | | |
| AT5G64960 | | cyclin dependent kinase group C2 | 57 kDa | 0 | 2 | 10 | 11 | 23 | | | | | | | |
| AT1G31830 | AT1G31830.2;AT1G31830.1 | Amino acid permease family protein | 55 kDa | 0 | 0 | 10 | 10 | 20 | | | | 2.4E+06 | 1.5E+06 | 0.0E+00 | 4.0E+06 |
| AT1G77030 | AT1G77030.1 | hydrolases, acting on acid anhydrides, in phosphorus-c | 94 kDa | 0 | 0 | 10 | 10 | 20 | | | | 5.9E+04 | 3.7E+04 | 2.1E+05 | 3.0E+05 |
| AT2G24420 | | DNA repair ATPase-related | 50 kDa | 0 | 0 | 10 | 10 | 20 | | | | | | | |
| AT3G22990 | AT3G22990.1 | ARM repeat superfamily protein | 50 kDa | 0 | 0 | 10 | 10 | 20 | | | | 2.5E+06 | 5.6E+05 | 2.2E+05 | 3.3E+06 |
| AT4G38050 | | Xanthine/uracil permease family protein | 78 kDa | 0 | 0 | 10 | 10 | 20 | | | | | | | |
| AT1G30270 | | CBL-interacting protein kinase 23 | 54 kDa | 0 | 0 | 10 | 9 | 19 | | | | | | | |
| AT3G04260 | | plastid transcriptionally active 3 | 103 kDa | 0 | 0 | 10 | 9 | 19 | | | | | | | |
| AT3G07040 | | NB-ARC domain-containing disease resistance protein | 107 kDa | 0 | 0 | 10 | 9 | 19 | 7.68 | | 7.68 | | | | |
| AT3G12590 | AT3G12590.2;AT3G12590.1 | unknown protein; INVOLVED IN: biological_process un | 131 kDa | 1 | 0 | 10 | 9 | 20 | | | | 5.3E+05 | 2.4E+05 | 0.0E+00 | 7.6E+05 |
| AT3G54860 | | Sec1/munc18-like (SM) proteins superfamily | 69 kDa | 0 | 0 | 10 | 9 | 19 | | | | | | | |
| AT4G22290 | | Ubiquitin-specific protease family C19-related protein | 48 kDa | 0 | 0 | 10 | 9 | 19 | | | | | | | |
| AT4G27020 | | unknown protein; FUNCTIONS IN: molecular_function | ? | 0 | 0 | 10 | 9 | 19 | | | | | | | |
| AT4G33530 | | K+ uptake permease 5 | 95 kDa | 0 | 0 | 10 | 9 | 19 | | | | | | | |
| AT5G20650 | | copper transporter 5 | 16 kDa | 0 | 0 | 10 | 9 | 19 | 0.52 | 0.16 | 0.34 | | | | |
| AT5G60360 | | aleurain-like protease | 40 kDa | 2 | 9 | 10 | 9 | 30 | | | | | | | |
| AT1G63120 | | RHOMBOID-like 2 | 35 kDa | 2 | 3 | 10 | 8 | 23 | | | | | | | |
| AT2G05760 | | Xanthine/uracil permease family protein | 57 kDa | 0 | 0 | 10 | 8 | 18 | | | | | | | |
| AT2G21620 | | Adenine nucleotide alpha hydrolases-like superfamily p | 21 kDa | 2 | 2 | 10 | 8 | 22 | | | | | | | |
| AT2G40380 | | prenylated RAB acceptor 1.B2 | 23 kDa | 4 | 8 | 10 | 8 | 30 | | 3.86 | 3.86 | | | | |
| AT5G13010 | AT5G13010.1 | RNA helicase family protein | 142 kDa | 0 | 0 | 10 | 8 | 18 | | | | | | | |
| AT5G15270 | AT5G15270.3;AT5G15270.2;AT5G15270.1 | RNA-binding KH domain-containing protein | 60 kDa | 0 | 0 | 10 | 8 | 18 | | | | 1.2E+06 | 4.0E+05 | 0.0E+00 | 1.6E+06 |
| AT5G43940 | AT5G43940.1;AT5G43940.2 | GroES-like zinc-binding dehydrogenase family protein | 41 kDa | 1 | 0 | 10 | 8 | 19 | | | | 2.5E+06 | 2.2E+06 | 0.0E+00 | 4.7E+06 |
| AT5G64090 | | FUNCTIONS IN: molecular_function unknown; INVOLVE | ? | 0 | 2 | 10 | 8 | 20 | | | | | | | |
| AT1G11020 | | RING/FYVE/PHD zinc finger superfamily protein | 36 kDa | 0 | 0 | 10 | 7 | 17 | | | | | | | |
| AT2G27080 | | Late embryogenesis abundant (LEA) hydroxyproline-ric | 29 kDa | 0 | 0 | 10 | 7 | 17 | | | | | | | |
| AT2G37280 | | pleiotropic drug resistance 5 | 160 kDa | 0 | 0 | 10 | 7 | 17 | | | | | | | |
| AT2G38620 | | cyclin-dependent kinase B1;2 | 36 kDa | 0 | 2 | 10 | 7 | 19 | | 1.15 | 1.15 | | | | |
| AT3G60190 | | DYNAMIN-like 1E | 70 kDa | 0 | 0 | 10 | 7 | 17 | | | | | | | |
| AT4G17486 | | PPPDE putative thiol peptidase family protein | 25 kDa | 1 | 9 | 10 | 7 | 27 | | | | | | | |
| AT5G01460 | AT5G01460.1 | LMBR1-like membrane protein | 57 kDa | 6 | 0 | 10 | 7 | 23 | | | | 2.9E+06 | 1.9E+06 | 0.0E+00 | 4.8E+06 |
| AT5G44100 | | casein kinase I-like 7 | 54 kDa | 0 | 0 | 10 | 7 | 17 | | | | | | | |
| AT2G01850 | AT2G01850.1 | endoxyloglucan transferase A3 | 38 kDa | 1 | 3 | 10 | 6 | 20 | | | | | | | |
| AT2G44640 | AT2G44640.1 | FUNCTIONS IN: molecular_function unknown; INVOLVE | 50 kDa | 0 | 0 | 10 | 6 | 16 | | | | 5.3E+06 | 6.4E+06 | 0.0E+00 | 1.2E+07 |
| AT4G05420 | AT4G05420.1;AT4G05420.2 | damaged DNA binding protein 1A | 121 kDa | 0 | 0 | 10 | 6 | 16 | | | | 1.1E+06 | 4.7E+05 | 2.4E+05 | 1.8E+06 |
| AT4G13180 | | NAD(P)-binding Rossmann-fold superfamily protein | 28 kDa | 0 | 5 | 10 | 6 | 21 | | | | | | | |
| AT5G01410 | AT5G01410.2;AT5G01410.1 | Aldolase-type TIM barrel family protein | 33 kDa | 0 | 0 | 10 | 6 | 16 | | | | | | | |
| AT5G18190 | | Protein kinase family protein | 77 kDa | 0 | 0 | 10 | 6 | 16 | | | | | | | |
| AT5G25060 | | RNA recognition motif (RRM)-containing protein | 108 kDa | 0 | 0 | 10 | 6 | 16 | 58.14 | | 58.14 | | | | |
| AT5G26850 | AT5G26850.4;AT5G26850.3;AT5G26850.2;AT5G26850.1 | Uncharacterized protein | 110 kDa | 0 | 0 | 10 | 6 | 16 | | | | | | | |
| AT5G28840 | | GDP-D-mannose 3',5'-epimerase | 43 kDa | 0 | 0 | 10 | 6 | 16 | | | | | | | |
| AT1G09200 | | Histone superfamily protein | 15 kDa | 1 | 0 | 10 | 5 | 16 | | | | | | | |
| AT1G72500 | | LOCATED IN: plasma membrane; EXPRESSED IN: 23 plan | 85 kDa | 0 | 0 | 10 | 5 | 15 | | | | | | | |
| AT2G19540 | AT2G19540.1 | Transducin family protein / WD-40 repeat family protei | 51 kDa | 0 | 0 | 10 | 5 | 15 | | | | 1.0E+07 | 1.2E+07 | 5.3E+06 | 2.7E+07 |
| AT4G21580 | | oxidoreductase, zinc-binding dehydrogenase family prc | 34 kDa | 5 | 0 | 10 | 5 | 20 | | | | | | | |
| AT4G37510 | | Ribonuclease III family protein | ? | 0 | 0 | 10 | 5 | 15 | | | | | | | |
| AT5G13480 | AT5G13480.3;AT5G13480.4;AT5G13480.1;AT5G13480.2 | Transducin/WD40 repeat-like superfamily protein | 72 kDa | 0 | 0 | 10 | 5 | 15 | | | | 3.7E+05 | 2.0E+05 | 2.5E+04 | 6.0E+05 |
| AT5G40950 | AT5G40950.1 | ribosomal protein large subunit 27 | 22 kDa | 0 | 0 | 10 | 5 | 15 | | | | | | | |
| AT5G53170 | | FTSH protease 11 | 89 kDa | 0 | 0 | 10 | 5 | 15 | | | | | | | |
| ATMG00560 | ATMG00560.1;AT2G07715.1 | Nucleic acid-binding, OB-fold-like protein | 38 kDa | 0 | 0 | 10 | 5 | 15 | | | | 2.6E+07 | 3.6E+07 | 2.2E+06 | 6.4E+07 |
| AT1G52300 | | Zinc-binding ribosomal protein family protein | 11 kDa | 0 | 2 | 10 | 4 | 16 | | | | | | | |
| AT1G69410 | AT1G69410.1;AT1G13950.1 | eukaryotic elongation factor 5A-3 | 17 kDa | 2 | 0 | 10 | 4 | 16 | | | | 6.7E+06 | 5.3E+06 | 3.5E+06 | 1.6E+07 |
| AT2G36130 | AT2G36130.1 | Cyclophilin-like peptidyl-prolyl cis-trans isomerase fam | 18 kDa | 2 | 0 | 10 | 4 | 16 | | 0.47 | 0.47 | 1.5E+07 | 2.6E+07 | 9.3E+05 | 4.1E+07 |
| AT3G59990 | AT3G59990.4;AT3G59990.3;AT3G59990.2;AT3G59990.1 | methionine aminopeptidase 2B | 49 kDa | 0 | 0 | 10 | 4 | 14 | | | | 1.1E+07 | 7.3E+06 | 8.9E+06 | 2.7E+07 |
| AT4G36940 | AT4G36940.1 | nicotinate phosphoribosyltransferase 1 | 62 kDa | 0 | 1 | 10 | 4 | 15 | | | | 2.8E+06 | 1.4E+06 | 1.5E+05 | 4.4E+06 |
| AT5G04280 | AT5G04280.1;AT5G04280.5;AT5G04280.4;AT5G04280.3;AT5G04280.2 | RNA-binding (RRM/RBD/RNP motifs) family protein wit | 33 kDa | 0 | 1 | 10 | 4 | 15 | | | | 7.3E+05 | 1.3E+06 | 0.0E+00 | 2.0E+06 |
| AT5G65900 | AT5G65900.1 | DEA(D/H)-box RNA helicase family protein | 72 kDa | 0 | 0 | 10 | 4 | 14 | | | | 2.8E+05 | 2.2E+05 | 2.4E+05 | 7.4E+05 |
| AT1G73180 | AT1G73180.2;AT1G73180.1 | Eukaryotic translation initiation factor eIF2A family prc | 56 kDa | 0 | 0 | 10 | 3 | 13 | | 0.34 | 0.34 | 2.4E+06 | 1.8E+06 | 1.3E+05 | 4.3E+06 |
| AT2G47300 | AT2G47300.4;AT2G47300.2 | ribonuclease Ps | 93 kDa | 0 | 0 | 10 | 3 | 13 | | | | | | | |
| AT5G55850 | AT5G55850.3;AT5G55850.1;AT5G55850.2 | RPM1-interacting protein 4 (RIN4) family protein | 10 kDa | 4 | 0 | 10 | 3 | 17 | | | | 7.7E+07 | 4.0E+07 | 7.0E+05 | 1.2E+08 |
| AT1G23280 | AT1G23280.1 | MAK16 protein-related | 36 kDa | 0 | 0 | 10 | 2 | 12 | | | | 7.8E+06 | 5.4E+06 | 1.3E+06 | 1.4E+07 |
| AT3G58840 | | Tropomyosin-related | 37 kDa | 0 | 0 | 10 | 2 | 12 | | | | | | | |
| AT5G19530 | | S-adenosyl-L-methionine-dependent methyltransferase | 39 kDa | 0 | 1 | 10 | 2 | 13 | | | | | | | |
| AT5G20160 | | Ribosomal protein L7Ae/L30e/S12e/Gadd45 family prc | 18 kDa | 1 | 0 | 10 | 2 | 13 | | | | | | | |

| | | | | | | | | | | | | | | | |
|---|---|---|---|---|---|---|---|---|---|---|---|---|---|---|---|
| AT5G40200 | AT5G40200.1 | DegP protease 9 | 65 kDa | 0 | 0 | 10 | 2 | 12 | | 0.19 | 0.19 | 2.3E+06 | 3.6E+06 | 1.2E+07 | 1.8E+07 |
| AT3G06610 | AT3G06610.1 | DNA-binding enhancer protein-related | 13 kDa | 0 | 0 | 10 | 1 | 11 | | 0.41 | 0.41 | 2.4E+07 | 1.2E+07 | 9.0E+06 | 4.6E+07 |
| AT5G53320 | | Leucine-rich repeat protein kinase family protein | 67 kDa | 0 | 0 | 10 | 1 | 11 | | | | | | | |
| AT5G17310 | | UDP-glucose pyrophosphorylase 2 | 52 kDa | 1 | 2 | 9 | 24 | 36 | | 3.35 | 3.35 | | | | |
| AT4G30920 | | Cytosol aminopeptidase family protein | 61 kDa | 0 | 1 | 9 | 18 | 28 | | | | | | | |
| AT5G43060 | AT5G43060.1 | Granulin repeat cysteine protease family protein | 51 kDa | 2 | 0 | 9 | 17 | 28 | | | | 2.3E+05 | 4.7E+05 | 0.0E+00 | 7.0E+05 |
| AT5G64030 | AT5G64030.1 | S-adenosyl-L-methionine-dependent methyltransferase | 93 kDa | 0 | 4 | 9 | 17 | 30 | | | | 0.0E+00 | 5.7E+05 | 0.0E+00 | 5.7E+05 |
| AT1G73720 | | transducin family protein / WD-40 repeat family protei | 58 kDa | 0 | 0 | 9 | 15 | 24 | | | | | | | |
| AT3G51270 | AT3G51270.2;AT3G51270.4;AT3G51270.3;AT3G51270.1 | protein serine/threonine kinases;ATP binding;catalytic | 53 kDa | 0 | 0 | 9 | 15 | 24 | | | | 1.0E+06 | 1.0E+06 | 0.0E+00 | 2.1E+06 |
| AT5G17330 | | glutamate decarboxylase | 57 kDa | 0 | 0 | 9 | 15 | 24 | | | | | | | |
| AT1G27450 | AT1G27450.4;AT1G27450.2;AT1G27450.1;AT1G27450.3 | adenine phosphoribosyl transferase 1 | 20 kDa | 2 | 4 | 9 | 14 | 29 | | | | 1.5E+07 | 9.7E+06 | 0.0E+00 | 2.4E+07 |
| AT3G02760 | AT3G02760.2;AT3G02760.1 | Class II aaRS and biotin synthetases superfamily protein | 98 kDa | 0 | 0 | 9 | 14 | 23 | | | | 1.0E+06 | 1.2E+06 | 0.0E+00 | 2.2E+06 |
| AT1G09640 | AT1G09640.2;AT1G09640.1 | Translation elongation factor EF1B, gamma chain | 47 kDa | 0 | 0 | 9 | 13 | 22 | | | | 1.9E+06 | 2.0E+06 | 0.0E+00 | 3.9E+06 |
| AT3G11800 | | unknown protein; FUNCTIONS IN: molecular_function | ? | 4 | 9 | 9 | 13 | 35 | | | | | | | |
| AT3G13990 | AT3G13990.2;AT3G13990.1 | Kinase-related protein of unknown function (DUF1296 | 92 kDa | 0 | 0 | 9 | 13 | 22 | | | | 3.4E+05 | 4.2E+05 | 2.0E+05 | 9.6E+05 |
| AT3G23990 | AT3G23990.1 | heat shock protein 60 | 61 kDa | 1 | 0 | 9 | 13 | 23 | 0.10 | 0.21 | 0.16 | 1.2E+06 | 2.5E+06 | 0.0E+00 | 3.7E+06 |
| AT3G33530 | | Transducin family protein / WD-40 repeat family protei | 147 kDa | 0 | 0 | 9 | 13 | 22 | | | | | | | |
| AT3G47990 | AT3G47990.2 | SUGAR-INSENSITIVE 3 | 41 kDa | 0 | 0 | 9 | 13 | 22 | | | | | | | |
| AT4G23400 | | plasma membrane intrinsic protein 1;5 | 31 kDa | 0 | 0 | 9 | 12 | 21 | | | | | | | |
| AT5G47690 | AT5G47690.3;AT5G47690.1;AT5G47690.2;AT5G47690.4 | binding | 181 kDa | 0 | 0 | 9 | 12 | 21 | | | | 1.3E+06 | 1.0E+06 | 1.7E+05 | 2.5E+06 |
| AT1G11280 | | S-locus lectin protein kinase family protein | 93 kDa | 2 | 2 | 9 | 10 | 23 | | | | | | | |
| AT1G48760 | | delta-adaptin | 97 kDa | 0 | 0 | 9 | 10 | 19 | | | | | | | |
| AT3G13570 | AT3G13570.1 | SC35-like splicing factor 30A | 30 kDa | 0 | 7 | 9 | 10 | 26 | | 0.81 | 0.81 | 3.1E+06 | 2.4E+06 | 7.7E+05 | 6.3E+06 |
| AT3G59970 | | methylenetetrahydrofolate reductase 1 | 66 kDa | 0 | 0 | 9 | 10 | 19 | | | | | | | |
| AT5G03320 | | Protein kinase superfamily protein | 47 kDa | 0 | 2 | 9 | 10 | 21 | | | | | | | |
| AT5G08570 | AT5G08570.3;AT5G08570.2;AT5G08570.1 | Pyruvate kinase family protein | 55 kDa | 0 | 0 | 9 | 10 | 19 | | | | 4.9E+06 | 5.8E+06 | 4.4E+05 | 1.1E+07 |
| AT5G10080 | AT5G10080.1 | Eukaryotic aspartyl protease family protein | 58 kDa | 0 | 0 | 9 | 10 | 19 | | | | | | | |
| AT5G35360 | AT5G35360.3;AT5G35360.1;AT5G35360.2 | acetyl Co-enzyme a carboxylase biotin carboxylase subu | 58 kDa | 0 | 0 | 9 | 10 | 19 | | | | 6.8E+06 | 8.0E+06 | 0.0E+00 | 1.5E+07 |
| AT1G08860 | AT1G08860.1 | Calcium-dependent phospholipid-binding Copine fami | 64 kDa | 0 | 0 | 9 | 9 | 18 | | | | 9.6E+04 | 6.5E+04 | 0.0E+00 | 1.6E+05 |
| AT1G28110 | AT1G28110.2;AT1G28110.1 | serine carboxypeptidase-like 45 | 52 kDa | 0 | 3 | 9 | 9 | 21 | 0.23 | 0.19 | 0.21 | | | | |
| AT3G02750 | | Protein phosphatase 2C family protein | 54 kDa | 0 | 0 | 9 | 9 | 18 | | | | | | | |
| AT3G49725 | AT3G49725.1 | GTP-binding protein, HflX | 70 kDa | 0 | 0 | 9 | 9 | 18 | | | | 2.7E+05 | 8.3E+05 | 1.7E+05 | 1.3E+06 |
| AT4G01370 | | MAP kinase 4 | 43 kDa | 1 | 0 | 9 | 9 | 19 | | | | | | | |
| AT4G36630 | AT4G36630.1 | Vacuolar sorting protein 39 | 111 kDa | 0 | 0 | 9 | 9 | 18 | | | | | | | |
| AT5G01950 | | Leucine-rich repeat protein kinase family protein | 106 kDa | 0 | 1 | 9 | 9 | 19 | | | | | | | |
| AT1G02305 | AT1G02305.1;AT1G02300.1 | Cysteine proteinases superfamily protein | 40 kDa | 0 | 0 | 9 | 8 | 17 | | | | 3.4E+06 | 4.0E+06 | 0.0E+00 | 7.3E+06 |
| AT1G18880 | | Major facilitator superfamily protein | 65 kDa | 0 | 0 | 9 | 8 | 17 | | | | | | | |
| AT1G29880 | | glycyl-tRNA synthetase / glycine--tRNA ligase | 82 kDa | 0 | 0 | 9 | 8 | 17 | | | | | | | |
| AT1G53050 | | Protein kinase superfamily protein | 78 kDa | 0 | 0 | 9 | 8 | 17 | | | | | | | |
| AT1G78770 | | anaphase promoting complex 6 | 62 kDa | 0 | 0 | 9 | 8 | 17 | | | | | | | |
| AT1G79590 | | syntaxin of plants 52 | 26 kDa | 0 | 0 | 9 | 8 | 17 | | | | | | | |
| AT2G20900 | | diacylglycerol kinase 5 | 57 kDa | 0 | 1 | 9 | 8 | 18 | | | | | | | |
| AT2G30860 | AT2G30860.1;AT2G30860.2 | glutathione S-transferase PHI 9 | 24 kDa | 0 | 4 | 9 | 8 | 21 | 0.25 | | 0.25 | 1.6E+07 | 4.5E+06 | 0.0E+00 | 2.0E+07 |
| AT2G38120 | AT2G38120.2;AT2G38120.1 | Transmembrane amino acid transporter family protein | 54 kDa | 0 | 0 | 9 | 8 | 17 | | | | 1.5E+07 | 5.3E+06 | 0.0E+00 | 2.0E+07 |
| AT3G11400 | AT3G11400.1;AT3G11400.2 | eukaryotic translation initiation factor 3G1 | 36 kDa | 4 | 0 | 9 | 8 | 21 | | 0.17 | 0.17 | 6.4E+06 | 5.1E+06 | 2.9E+06 | 1.4E+07 |
| AT3G57880 | | Calcium-dependent lipid-binding (CaLB domain) plant | 89 kDa | 8 | 0 | 9 | 8 | 25 | | | | | | | |
| AT4G01700 | AT4G01700.1 | Chitinase family protein | 31 kDa | 0 | 2 | 9 | 8 | 19 | | | | | | | |
| AT4G08810 | | calcium ion binding | 64 kDa | 0 | 0 | 9 | 8 | 17 | | | | | | | |
| AT4G16340 | | guanyl-nucleotide exchange factors;GTPase binding;GT | 207 kDa | 0 | 0 | 9 | 8 | 17 | | | | | | | |
| AT4G21800 | AT4G21800.2;AT4G21800.1 | P-loop containing nucleoside triphosphate hydrolases | 43 kDa | 0 | 1 | 9 | 8 | 18 | | | | | | | |
| AT5G04290 | | kow domain-containing transcription factor 1 | 158 kDa | 0 | 0 | 9 | 8 | 17 | | | | | | | |
| AT5G55830 | | Concanavalin A-like lectin protein kinase family protein | 76 kDa | 0 | 0 | 9 | 8 | 17 | | | | | | | |
| AT5G60730 | | Anion-transporting ATPase | 43 kDa | 0 | 0 | 9 | 8 | 17 | | | | | | | |
| AT1G78120 | | Tetratricopeptide repeat (TPR)-like superfamily protein | 60 kDa | 0 | 1 | 9 | 7 | 17 | | | | | | | |
| AT2G31410 | AT2G31410.1 | unknown protein; Has 1719 Blast hits to 1091 proteins | 23 kDa | 0 | 2 | 9 | 7 | 18 | | | | 1.5E+07 | 1.6E+07 | 4.1E+06 | 3.5E+07 |
| AT3G48760 | | DHHC-type zinc finger family protein | 54 kDa | 0 | 0 | 9 | 7 | 16 | | | | | | | |
| AT3G56100 | | meristematic receptor-like kinase | 77 kDa | 0 | 0 | 9 | 7 | 16 | | | | | | | |
| AT4G26110 | | nucleosome assembly protein1;1 | 43 kDa | 0 | 0 | 9 | 7 | 16 | | | | | | | |
| AT5G16240 | AT5G16240.1 | Plant stearoyl-acyl-carrier-protein desaturase family pr | 45 kDa | 0 | 1 | 9 | 7 | 17 | | | | 3.1E+06 | 2.8E+06 | 7.4E+05 | 6.6E+06 |
| AT5G22790 | AT5G22790.1 | reticulata-related 1 | 47 kDa | 0 | 0 | 9 | 7 | 16 | | | | 4.1E+05 | 4.7E+05 | 0.0E+00 | 8.9E+05 |
| AT5G25150 | AT5G25150.2;AT5G25150.1 | TBP-associated factor 5 | 74 kDa | 0 | 0 | 9 | 7 | 16 | | | | 0.0E+00 | 0.0E+00 | 3.2E+04 | 3.2E+04 |
| AT5G50860 | | Protein kinase superfamily protein | 64 kDa | 0 | 0 | 9 | 7 | 16 | | | | | | | |
| ATCG01120 | | chloroplast ribosomal protein S15 | 11 kDa | 0 | 0 | 9 | 7 | 16 | | 0.37 | 0.37 | | | | |
| AT1G27900 | | RNA helicase family protein | 78 kDa | 0 | 0 | 9 | 6 | 15 | | | | | | | |
| AT1G71697 | | choline kinase 1 | 40 kDa | 0 | 0 | 9 | 6 | 15 | | | | | | | |
| AT2G32850 | | Protein kinase superfamily protein | 75 kDa | 0 | 3 | 9 | 6 | 18 | | | | | | | |
| AT2G43950 | AT2G43950.2;AT2G43950.1 | chloroplast outer envelope protein 37 | 39 kDa | 0 | 0 | 9 | 6 | 15 | | | | 1.6E+06 | 1.6E+06 | 0.0E+00 | 3.2E+06 |
| AT2G44790 | | uclacyanin 2 | 20 kDa | 8 | 5 | 9 | 6 | 28 | | | | | | | |
| AT3G18520 | | histone deacetylase 15 | 60 kDa | 0 | 0 | 9 | 6 | 15 | | | | | | | |
| AT3G26700 | | Protein kinase superfamily protein | 42 kDa | 0 | 7 | 9 | 6 | 22 | | | | | | | |
| AT4G17720 | | RNA-binding (RRM/RBD/RNP motifs) family protein | 34 kDa | 0 | 0 | 9 | 6 | 15 | | | | | | | |
| AT5G04130 | AT5G04130.3;AT5G04130.2;AT5G04130.1 | DNA GYRASE B2 | 81 kDa | 0 | 0 | 9 | 6 | 15 | | | | 8.1E+05 | 7.1E+05 | 0.0E+00 | 1.5E+06 |
| AT1G05020 | | ENTH/ANTH/VHS superfamily protein | 72 kDa | 1 | 1 | 9 | 5 | 16 | | | | | | | |
| AT1G05960 | | ARM repeat superfamily protein | 109 kDa | 0 | 0 | 9 | 5 | 14 | | | | | | | |
| AT1G24450 | AT1G24450.1 | Ribonuclease III family protein | 21 kDa | 1 | 0 | 9 | 5 | 15 | | | | 3.1E+06 | 2.5E+06 | 1.4E+06 | 7.0E+06 |
| AT1G72340 | | NagB/RpiA/CoA transferase-like superfamily protein | 42 kDa | 0 | 0 | 9 | 5 | 14 | | | | | | | |
| AT2G40940 | | ethylene response sensor 1 | 68 kDa | 0 | 0 | 9 | 5 | 14 | | | | | | | |
| AT3G25230 | | rotamase FKBP 1 | 61 kDa | 0 | 0 | 9 | 5 | 14 | | | | | | | |
| AT3G54260 | | TRICHOME BIREFRINGENCE-LIKE 36 | 44 kDa | 0 | 0 | 9 | 5 | 14 | | | | | | | |
| AT4G04640 | AT4G04640.1 | ATPase, F1 complex, gamma subunit protein | 41 kDa | 0 | 2 | 9 | 5 | 16 | | 0.50 | 0.50 | 4.1E+06 | 1.3E+06 | 6.3E+04 | 5.5E+06 |
| AT4G31210 | | DNA topoisomerase, type IA, core | 142 kDa | 0 | 0 | 9 | 5 | 14 | | | | | | | |
| AT5G22110 | AT5G22110.2;AT5G22110.1 | DNA polymerase epsilon subunit B2 | 60 kDa | 0 | 0 | 9 | 5 | 14 | | | | | | | |
| AT5G40540 | | Protein kinase superfamily protein | 40 kDa | 0 | 2 | 9 | 5 | 16 | | | | | | | |
| ATCG00900 | | Ribosomal protein S7p/S5e family protein | 17 kDa | 0 | 0 | 9 | 5 | 14 | | 0.57 | 0.57 | | | | |
| AT2G23980 | | cyclic nucleotide-gated channel 6 | 85 kDa | 0 | 0 | 9 | 4 | 13 | | | | | | | |
| AT3G52090 | AT3G52090.1;AT3G52090.2 | DNA-directed RNA polymerase, RBP11-like | 14 kDa | 1 | 0 | 9 | 4 | 14 | | 0.36 | 0.36 | 3.4E+07 | 3.4E+07 | 1.0E+07 | 7.8E+07 |
| AT3G55070 | AT3G55070.1 | LisH/CRA/RING-U-box domains-containing protein | 48 kDa | 0 | 0 | 9 | 4 | 13 | | | | 2.3E+06 | 1.8E+06 | 2.2E+05 | 4.3E+06 |
| AT5G04600 | AT5G04600.1 | RNA-binding (RRM/RBD/RNP motifs) family protein | 25 kDa | 0 | 0 | 9 | 4 | 13 | | | | 2.5E+07 | 1.9E+07 | 1.4E+07 | 5.8E+07 |
| AT5G19290 | AT5G19290.1 | alpha/beta-Hydrolases superfamily protein | 37 kDa | 0 | 0 | 9 | 4 | 13 | | | | 3.9E+06 | 3.2E+06 | 1.6E+05 | 7.2E+06 |
| AT5G43470 | | Disease resistance protein (CC-NBS-LRR class) family | 105 kDa | 0 | 0 | 9 | 4 | 13 | | | | | | | |
| ATCG00380 | ATCG00380.1 | chloroplast ribosomal protein S4 | 23 kDa | 0 | 0 | 9 | 4 | 13 | | 0.27 | 0.27 | 2.9E+06 | 2.2E+06 | 2.8E+05 | 5.3E+06 |
| AT1G67720 | | Leucine-rich repeat protein kinase family protein | 104 kDa | 0 | 0 | 9 | 3 | 12 | | | | | | | |
| AT1G77840 | AT1G77840.1 | Translation initiation factor IF2/IF5 | 49 kDa | 0 | 0 | 9 | 3 | 12 | | | | 7.3E+04 | 5.2E+04 | 0.0E+00 | 1.3E+05 |

| | | | | | | | | | | | | | | | |
|---|---|---|---|---|---|---|---|---|---|---|---|---|---|---|---|
| AT2G40540 | | potassium transporter 2 | 89 kDa | 0 | 0 | 9 | 3 | 12 | | | | | | | |
| AT4G08950 | AT4G08950.1 | Phosphate-responsive 1 family protein | 34 kDa | 2 | 1 | 9 | 3 | 15 | 0.20 | | 0.20 | 3.1E+05 | 6.4E+05 | 0.0E+00 | 9.5E+05 |
| AT4G24160 | | alpha/beta-Hydrolases superfamily protein | 47 kDa | 0 | 0 | 9 | 3 | 12 | | | | | | | |
| AT4G33760 | | tRNA synthetase class II (D, K and N) family protein | 75 kDa | 0 | 0 | 9 | 3 | 12 | | | | | | | |
| AT5G07140 | | Protein kinase superfamily protein | 64 kDa | 0 | 0 | 9 | 3 | 12 | | | | | | | |
| AT1G29250 | AT1G29250.1 | Alba DNA/RNA-binding protein | 15 kDa | 0 | 0 | 9 | 2 | 11 | | | | 0.0E+00 | 2.6E+06 | 3.1E+06 | 5.7E+06 |
| AT1G79190 | | ARM repeat superfamily protein | 148 kDa | 0 | 0 | 9 | 2 | 11 | | | | | | | |
| AT2G03890 | | phosphoinositide 4-kinase gamma 7 | 73 kDa | 0 | 0 | 9 | 2 | 11 | | | | | | | |
| AT3G54210 | | Ribosomal protein L17 family protein | 23 kDa | 0 | 0 | 9 | 2 | 11 | | | | | | | |
| AT5G44860 | | unknown protein; BEST Arabidopsis thaliana protein m | ? | 0 | 0 | 9 | 2 | 11 | | | | | | | |
| AT3G06810 | AT3G06810.1 | acyl-CoA dehydrogenase-related | 92 kDa | 0 | 0 | 9 | 1 | 10 | | | | 1.2E+06 | 2.3E+06 | 0.0E+00 | 3.5E+06 |
| AT3G13640 | | RNAse l inhibitor protein 1 | 68 kDa | 0 | 0 | 9 | 1 | 10 | | | | | | | |
| AT5G53560 | AT5G53560.1 | cytochrome B5 isoform E | 15 kDa | 3 | 0 | 9 | 0 | 12 | 0.19 | 0.11 | 0.15 | 3.0E+06 | 3.6E+06 | 5.0E+05 | 7.1E+06 |
| AT1G30120 | | pyruvate dehydrogenase E1 beta | 44 kDa | 1 | 12 | 8 | 19 | 40 | | 0.46 | 0.46 | | | | |
| AT4G02990 | | Mitochondrial transcription termination factor family | 62 kDa | 0 | 0 | 8 | 18 | 26 | | | | | | | |
| AT3G43190 | | sucrose synthase 4 | 93 kDa | 0 | 0 | 8 | 17 | 25 | | | | | | | |
| AT4G32750 | | unknown protein; Has 30201 Blast hits to 17322 prote | 33 kDa | 0 | 0 | 8 | 17 | 25 | | | | | | | |
| AT1G04980 | AT1G04980.1 | PDI-like 2-2 | 48 kDa | 0 | 2 | 8 | 15 | 25 | | | | 4.6E+06 | 6.7E+06 | 2.6E+04 | 1.1E+07 |
| AT1G63780 | AT1G63780.1 | Ribosomal RNA processing Brix domain protein | 34 kDa | 0 | 0 | 8 | 15 | 23 | | 1.66 | 1.66 | 8.5E+06 | 8.3E+06 | 2.0E+06 | 1.9E+07 |
| AT1G60790 | | Plant protein of unknown function (DUF828) | 61 kDa | 0 | 0 | 8 | 14 | 22 | | | | | | | |
| AT3G61050 | | Calcium-dependent lipid-binding (CaLB domain) family | 55 kDa | 3 | 3 | 8 | 14 | 28 | | | | | | | |
| AT5G46840 | AT5G46840.1;AT5G46840.2 | RNA-binding (RRM/RBD/RNP motifs) family protein | 56 kDa | 0 | 0 | 8 | 13 | 21 | | | | 1.2E+06 | 1.4E+06 | 9.8E+05 | 3.6E+06 |
| AT1G80300 | | nucleotide transporter 1 | 68 kDa | 0 | 0 | 8 | 11 | 19 | | | | | | | |
| AT2G24490 | | replicon protein A2 | 31 kDa | 0 | 1 | 8 | 11 | 20 | | | | | | | |
| AT5G43420 | | RING/U-box superfamily protein | 42 kDa | 0 | 0 | 8 | 11 | 19 | | | | | | | |
| AT1G17120 | | cationic amino acid transporter 8 | 65 kDa | 0 | 0 | 8 | 10 | 18 | | | | | | | |
| AT2G34460 | AT2G34460.1 | NAD(P)-binding Rossmann-fold superfamily protein | 30 kDa | 2 | 11 | 8 | 10 | 31 | | | | | | | |
| AT2G39910 | | ARM repeat superfamily protein | 50 kDa | 0 | 0 | 8 | 10 | 18 | | | | | | | |
| AT2G40700 | AT2G40700.1 | P-loop containing nucleoside triphosphate hydrolases | 68 kDa | 0 | 0 | 8 | 10 | 18 | | | | 9.5E+06 | 0.0E+00 | 1.3E+05 | 9.6E+06 |
| AT3G26020 | | Protein phosphatase 2A regulatory B subunit family pro | 59 kDa | 0 | 0 | 8 | 10 | 18 | 0.10 | | 0.10 | | | | |
| AT3G50670 | AT3G50670.1;AT3G50670.2 | U1 small nuclear ribonucleoprotein-70K | 50 kDa | 0 | 0 | 8 | 10 | 18 | | | | 5.5E+06 | 8.8E+06 | 2.9E+05 | 1.5E+07 |
| AT5G13530 | | protein kinases;ubiquitin-protein ligases | 178 kDa | 0 | 3 | 8 | 10 | 21 | | | | | | | |
| AT5G56970 | | cytokinin oxidase 3 | 59 kDa | 0 | 0 | 8 | 10 | 18 | | | | | | | |
| AT1G12780 | | UDP-D-glucose/UDP-D-galactose 4-epimerase 1 | 39 kDa | 0 | 1 | 8 | 9 | 18 | | | | | | | |
| AT1G20330 | | sterol methyltransferase 2 | 40 kDa | 0 | 0 | 8 | 9 | 17 | | | | | | | |
| AT1G45233 | | THO complex, subunit 5 | 52 kDa | 0 | 0 | 8 | 9 | 17 | | | | | | | |
| AT1G57700 | | Protein kinase superfamily protein | 78 kDa | 0 | 0 | 8 | 9 | 17 | | | | | | | |
| AT1G76660 | | FUNCTIONS IN: molecular_function unknown; INVOLVE | 46 kDa | 0 | 0 | 8 | 9 | 17 | | | | | | | |
| AT3G56990 | AT3G56990.2;AT3G56990.3;AT3G56990.1 | embryo sac development arrest 7 | 80 kDa | 0 | 0 | 8 | 9 | 17 | | | | | | | |
| AT2G21130 | | Cyclophilin-like peptidyl-prolyl cis-trans isomerase fam | 18 kDa | 0 | 0 | 8 | 8 | 16 | | | | | | | |
| AT2G30110 | | ubiquitin-activating enzyme 1 | 120 kDa | 0 | 0 | 8 | 8 | 16 | | | | | | | |
| AT2G32240 | AT2G32240.1 | FUNCTIONS IN: molecular_function unknown; INVOLVE | ? | 0 | 0 | 8 | 8 | 16 | | | | 1.9E+06 | 2.6E+06 | 0.0E+00 | 4.5E+06 |
| AT3G17820 | AT3G17820.1 | glutamine synthetase 1.3 | 39 kDa | 0 | 0 | 8 | 8 | 16 | | | | 1.0E+06 | 1.1E+06 | 0.0E+00 | 2.1E+06 |
| AT3G33520 | | actin-related protein 6 | 47 kDa | 0 | 0 | 8 | 8 | 16 | | | | | | | |
| AT3G55960 | | Haloacid dehalogenase-like hydrolase (HAD) superfamil | 35 kDa | 0 | 0 | 8 | 8 | 16 | | | | | | | |
| AT3G58760 | | Integrin-linked protein kinase family | 53 kDa | 0 | 0 | 8 | 8 | 16 | | | | | | | |
| AT4G21150 | | ribophorin II (RPN2) family protein | 75 kDa | 0 | 0 | 8 | 8 | 16 | 0.11 | 0.49 | 0.30 | | | | |
| AT5G01890 | | Leucine-rich receptor-like protein kinase family protein | 104 kDa | 0 | 0 | 8 | 8 | 16 | | | | | | | |
| AT5G03040 | AT5G03040.3;AT5G03040.2;AT5G03040.1 | IQ-domain 2 | 50 kDa | 0 | 0 | 8 | 8 | 16 | | | | 6.1E+05 | 7.2E+05 | 0.0E+00 | 1.3E+06 |
| AT1G16970 | AT1G16970.1 | KU70 homolog | 70 kDa | 0 | 0 | 8 | 7 | 15 | | | | 4.1E+05 | 5.3E+05 | 0.0E+00 | 9.4E+05 |
| AT1G18040 | | cyclin-dependent kinase D1;3 | 45 kDa | 0 | 2 | 8 | 7 | 17 | | | | | | | |
| AT1G21380 | | Target of Myb protein 1 | 56 kDa | 0 | 0 | 8 | 7 | 15 | | | | | | | |
| AT1G50910 | AT1G50910.1 | unknown protein; Has 1105 Blast hits to 802 proteins i | 56 kDa | 0 | 0 | 8 | 7 | 15 | | | | | | | |
| AT2G05170 | | vacuolar protein sorting 11 | 106 kDa | 0 | 0 | 8 | 7 | 15 | | | | | | | |
| AT2G39670 | AT2G39670.1;AT2G39670.2 | Radical SAM superfamily protein | 47 kDa | 0 | 0 | 8 | 7 | 15 | | | | 9.9E+05 | 1.1E+06 | 6.3E+05 | 2.8E+06 |
| AT2G40980 | | Protein kinase superfamily protein | 69 kDa | 0 | 0 | 8 | 7 | 15 | | | | | | | |
| AT3G11500 | AT3G11500.2;AT3G11500.1 | Small nuclear ribonucleoprotein family protein | 9 kDa | 1 | 0 | 8 | 7 | 16 | | | | 1.1E+07 | 1.2E+07 | 6.6E+05 | 2.4E+07 |
| AT3G25780 | AT3G25780.1 | allene oxide cyclase 3 | 28 kDa | 0 | 0 | 8 | 7 | 15 | | 0.37 | 0.37 | | | | |
| AT4G39960 | AT4G39960.2;AT4G39960.1 | Molecular chaperone Hsp40/DnaJ family protein | 48 kDa | 0 | 0 | 8 | 7 | 15 | | | | 4.3E+05 | 0.0E+00 | 1.6E+05 | 5.9E+05 |
| AT1G10850 | | Leucine-rich repeat protein kinase family protein | 73 kDa | 0 | 0 | 8 | 6 | 14 | | | | | | | |
| AT2G34585 | AT2G34585.1 | unknown protein; FUNCTIONS IN: molecular_function | 9 kDa | 6 | 0 | 8 | 6 | 20 | | 1.58 | 1.58 | 7.3E+06 | 0.0E+00 | 1.1E+06 | 8.3E+06 |
| AT2G45240 | AT2G45240.1;AT2G45240.2 | methionine aminopeptidase 1A | 44 kDa | 0 | 1 | 8 | 6 | 15 | | | | 2.2E+06 | 1.9E+06 | 1.8E+06 | 5.9E+06 |
| AT4G21390 | | S-locus lectin protein kinase family protein | 95 kDa | 0 | 0 | 8 | 6 | 14 | | | | | | | |
| AT5G05000 | AT5G05000.3;AT5G05000.2;AT5G05000.1 | translocon at the outer envelope membrane of chlorop | 35 kDa | 6 | 2 | 8 | 6 | 22 | 0.10 | 0.30 | 0.20 | 1.6E+06 | 4.1E+06 | 1.9E+05 | 5.9E+06 |
| AT5G05460 | | Glycosyl hydrolase family 85 | 76 kDa | 0 | 0 | 8 | 6 | 14 | | | | | | | |
| AT5G14590 | | Isocitrate/isopropylmalate dehydrogenase family prote | 54 kDa | 0 | 1 | 8 | 6 | 15 | | | | | | | |
| AT5G23210 | AT5G23210.4;AT5G23210.2;AT5G23210.5;AT5G23210.3;AT5G23210.1 | serine carboxypeptidase-like 34 | 56 kDa | 0 | 0 | 8 | 6 | 14 | | | | 1.1E+06 | 1.2E+06 | 0.0E+00 | 2.3E+06 |
| AT5G41210 | AT5G41210.1 | glutathione S-transferase THETA 1 | 28 kDa | 0 | 5 | 8 | 6 | 19 | | | | 1.0E+07 | 1.0E+07 | 2.4E+06 | 2.3E+07 |
| AT5G53480 | | ARM repeat superfamily protein | 96 kDa | 0 | 0 | 8 | 6 | 14 | | | | | | | |
| AT5G54910 | AT5G54910.1 | DEA(D/H)-box RNA helicase family protein | 84 kDa | 0 | 0 | 8 | 6 | 14 | | | | 1.2E+05 | 1.4E+05 | 3.9E+05 | 6.5E+05 |
| AT1G06760 | AT1G06760.1 | winged-helix DNA-binding transcription factor family p | 29 kDa | 0 | 0 | 8 | 5 | 13 | | | | 9.6E+05 | 1.5E+06 | 3.1E+05 | 2.7E+06 |
| AT1G09850 | | xylem bark cysteine peptidase 3 | 48 kDa | 2 | 0 | 8 | 5 | 15 | | 0.10 | 0.10 | | | | |
| AT1G13690 | AT1G13690.1 | ATPase E1 | 20 kDa | 0 | 0 | 8 | 5 | 13 | | | | 2.2E+07 | 3.8E+07 | 4.5E+06 | 6.5E+07 |
| AT1G14710 | AT1G14710.2;AT1G14710.1 | hydroxyproline-rich glycoprotein family protein | 66 kDa | 0 | 0 | 8 | 5 | 13 | | | | 1.0E+06 | 3.5E+05 | 4.1E+05 | 1.8E+06 |
| AT1G20930 | | cyclin-dependent kinase B2;2 | 36 kDa | 0 | 3 | 8 | 5 | 16 | | | | | | | |
| AT1G27930 | | Protein of unknown function (DUF579) | 32 kDa | 2 | 7 | 8 | 5 | 22 | | | | | | | |
| AT1G28380 | | MAC/Perforin domain-containing protein | 68 kDa | 0 | 0 | 8 | 5 | 13 | | | | | | | |
| AT1G54090 | | exocyst subunit exo70 family protein D2 | 71 kDa | 0 | 0 | 8 | 5 | 13 | | | | | | | |
| AT1G79930 | | heat shock protein 91 | 92 kDa | 0 | 0 | 8 | 5 | 13 | | | | | | | |
| AT2G17290 | | Calcium-dependent protein kinase family protein | 61 kDa | 0 | 0 | 8 | 5 | 13 | | | | | | | |
| AT2G21290 | AT2G21290.1 | unknown protein; FUNCTIONS IN: molecular_function | 11 kDa | 0 | 0 | 8 | 5 | 13 | | | | 2.8E+07 | 5.0E+07 | 2.1E+06 | 8.0E+07 |
| AT2G35840 | AT2G35840.4;AT2G35840.3;AT2G35840.2;AT2G35840.1 | Sucrose-6F-phosphate phosphohydrolase family protei | 48 kDa | 0 | 0 | 8 | 5 | 13 | | | | 4.2E+06 | 2.4E+06 | 8.6E+04 | 6.7E+06 |
| AT3G18050 | | unknown protein; BEST Arabidopsis thaliana protein m | ? | 0 | 0 | 8 | 5 | 13 | | | | | | | |
| AT3G46530 | | NB-ARC domain-containing disease resistance protein | 97 kDa | 0 | 0 | 8 | 5 | 13 | | | | | | | |
| AT3G56640 | | exocyst complex component sec15A | 90 kDa | 0 | 0 | 8 | 5 | 13 | | | | | | | |
| AT3G60240 | AT3G60240.2;AT3G60240.3;AT3G60240.4 | eukaryotic translation initiation factor 4G | 188 kDa | 0 | 0 | 8 | 5 | 13 | | 0.25 | 0.25 | 1.0E+05 | 1.5E+05 | 8.7E+04 | 3.4E+05 |
| AT4G26750 | | hydroxyproline-rich glycoprotein family protein | 46 kDa | 0 | 0 | 8 | 5 | 13 | | | | | | | |
| AT4G39260 | | cold, circadian rhythm, and RNA binding 1 | 17 kDa | 4 | 0 | 8 | 5 | 17 | | 0.23 | 0.23 | | | | |
| AT5G06860 | | polygalacturonase inhibiting protein 1 | 37 kDa | 0 | 0 | 8 | 5 | 13 | | | | | | | |
| AT5G08420 | AT5G08420.1 | RNA-binding KH domain-containing protein | 45 kDa | 0 | 0 | 8 | 5 | 13 | | | | 0.0E+00 | 7.0E+05 | 8.6E+05 | 1.6E+06 |
| AT5G13260 | | unknown protein; BEST Arabidopsis thaliana protein m | 60 kDa | 0 | 0 | 8 | 5 | 13 | | | | | | | |
| AT5G19050 | | alpha/beta-Hydrolases superfamily protein | 38 kDa | 2 | 1 | 8 | 5 | 16 | | | | | | | |
| AT1G72470 | | exocyst subunit exo70 family protein D1 | 71 kDa | 0 | 0 | 8 | 4 | 12 | | | | | | | |

| | | | | | | | | | | | | | | | |
|---|---|---|---|---|---|---|---|---|---|---|---|---|---|---|---|
| AT1G80640 | | Protein kinase superfamily protein | 47 kDa | 0 | 0 | 8 | 4 | 12 | | | | | | | |
| AT2G30880 | AT2G30880.2;AT2G30880.1 | Pleckstrin homology (PH) domain-containing protein | 56 kDa | 0 | 0 | 8 | 4 | 12 | | | | 1.2E+06 | 1.4E+06 | 0.0E+00 | 2.7E+06 |
| AT2G40280 | | S-adenosyl-L-methionine-dependent methyltransferase | 67 kDa | 0 | 0 | 8 | 4 | 12 | | | | | | | |
| AT2G47420 | | Ribosomal RNA adenine dimethylase family protein | 40 kDa | 0 | 0 | 8 | 4 | 12 | | | | | | | |
| AT3G43610 | | Spc97 / Spc98 family of spindle pole body (SBP) compo | 137 kDa | 0 | 0 | 8 | 4 | 12 | | | | | | | |
| AT5G41950 | | Tetratricopeptide repeat (TPR)-like superfamily protein | 62 kDa | 1 | 0 | 8 | 4 | 13 | | | | | | | |
| AT5G42790 | AT5G42790.1;AT1G47250.1 | proteasome alpha subunit F1 | 30 kDa | 0 | 0 | 8 | 4 | 12 | | | | 5.5E+06 | 5.9E+06 | 0.0E+00 | 1.1E+07 |
| AT5G48240 | AT5G48240.2;AT5G48240.1;AT5G48240.3 | unknown protein; FUNCTIONS IN: molecular_function | 34 kDa | 0 | 0 | 8 | 4 | 12 | | | | 0.0E+00 | 1.4E+06 | 1.5E+06 | 2.9E+06 |
| ATCG00750 | | ribosomal protein S11 | 15 kDa | 0 | 0 | 8 | 4 | 12 | | | | | | | |
| AT1G12310 | AT1G12310.1 | Calcium-binding EF-hand family protein | 17 kDa | 4 | 0 | 8 | 3 | 15 | 1.18 | 0.45 | 0.82 | 1.8E+06 | 1.7E+06 | 0.0E+00 | 3.4E+06 |
| AT1G15130 | | Endosomal targeting BRO1-like domain-containing pro | 95 kDa | 0 | 0 | 8 | 3 | 11 | | | | | | | |
| AT1G78630 | | Ribosomal protein L13 family protein | 27 kDa | 0 | 2 | 8 | 3 | 13 | | | | | | | |
| AT2G33330 | | plasmodesmata-located protein 3 | 33 kDa | 0 | 2 | 8 | 3 | 13 | | | | | | | |
| AT2G38230 | | pyridoxine biosynthesis 1.1 | 33 kDa | 0 | 0 | 8 | 3 | 11 | | | | | | | |
| AT2G39700 | | expansin A4 | 28 kDa | 0 | 0 | 8 | 3 | 11 | | | | | | | |
| AT2G46800 | AT2G46800.5;AT2G46800.4;AT2G46800.3;AT2G46800.2;AT2G46800.1 | zinc transporter of Arabidopsis thaliana | 44 kDa | 0 | 0 | 8 | 3 | 11 | | | | | | | |
| AT3G12800 | AT3G12800.1 | short-chain dehydrogenase-reductase B | ? | 0 | 0 | 8 | 3 | 11 | 0.13 | 0.31 | 0.22 | 6.6E+07 | 6.3E+07 | 6.9E+06 | 1.4E+08 |
| AT3G49601 | AT3G49601.1 | FUNCTIONS IN: molecular_function unknown; INVOLVI | 68 kDa | 0 | 0 | 8 | 3 | 11 | | | | 1.2E+06 | 1.3E+06 | 5.2E+05 | 3.0E+06 |
| AT5G26030 | AT5G26030.2;AT5G26030.1 | ferrochelatase 1 | 52 kDa | 0 | 0 | 8 | 3 | 11 | | | | 7.5E+05 | 1.6E+06 | 0.0E+00 | 2.3E+06 |
| AT5G47310 | | PPPDE putative thiol peptidase family protein | 27 kDa | 3 | 7 | 8 | 3 | 21 | | 6.95 | 6.95 | | | | |
| AT5G48620 | | Disease resistance protein (CC-NBS-LRR class) family | 104 kDa | 0 | 0 | 8 | 3 | 11 | | | | | | | |
| AT1G19880 | AT1G19880.1 | Regulator of chromosome condensation (RCC1) family | 58 kDa | 0 | 0 | 8 | 2 | 10 | | | | 1.6E+05 | 1.7E+06 | 1.5E+05 | 2.0E+06 |
| AT1G75950 | AT1G75950.1 | S phase kinase-associated protein 1 | 18 kDa | 0 | 2 | 8 | 2 | 12 | | | | 1.8E+07 | 1.1E+07 | 2.7E+05 | 3.0E+07 |
| AT2G31570 | | glutathione peroxidase 2 | 19 kDa | 0 | 0 | 8 | 2 | 10 | | | | | | | |
| AT3G02600 | | lipid phosphate phosphatase 3 | 35 kDa | 0 | 2 | 8 | 2 | 12 | | | | | | | |
| AT3G10090 | | Nucleic acid-binding, OB-fold-like protein | 7 kDa | 0 | 0 | 8 | 2 | 10 | 0.16 | 0.07 | 0.12 | | | | |
| AT3G22950 | | ADP-ribosylation factor C1 | 21 kDa | 0 | 0 | 8 | 2 | 10 | | | | | | | |
| AT4G04340 | | ERD (early-responsive to dehydration stress) family prot | 88 kDa | 0 | 0 | 8 | 2 | 10 | | | | | | | |
| AT4G29350 | AT4G29350.1 | profilin 2 | 14 kDa | 3 | 0 | 8 | 2 | 13 | 5.88 | | 5.88 | 1.7E+07 | 7.4E+06 | 0.0E+00 | 2.5E+07 |
| AT5G10810 | AT5G10810.1 | enhancer of rudimentary protein, putative | 13 kDa | 0 | 0 | 8 | 2 | 10 | | | | 3.0E+06 | 4.5E+06 | 3.6E+06 | 1.1E+07 |
| AT5G15750 | AT5G15750.1 | Alpha-L RNA-binding motif/Ribosomal protein S4 famil | 22 kDa | 0 | 0 | 8 | 2 | 10 | 0.09 | | 0.09 | 1.9E+06 | 2.5E+06 | 2.8E+05 | 4.6E+06 |
| AT5G22650 | AT5G22650.2;AT5G22650.1 | histone deacetylase 2B | 32 kDa | 0 | 0 | 8 | 2 | 10 | | 0.16 | 0.16 | 6.4E+07 | 9.9E+07 | 2.0E+07 | 1.8E+08 |
| AT1G31360 | AT1G31360.8;AT1G31360.4;AT1G31360.7;AT1G31360.5;AT1G31360.6;AT1G31360.1;AT1G31360.2;AT1G31360.3 | RECQ helicase L2 | 79 kDa | 0 | 0 | 8 | 1 | 9 | | | | 1.2E+06 | 6.8E+05 | 0.0E+00 | 1.9E+06 |
| AT2G41620 | | Nucleoporin interacting component (Nup93/Nic96-lik | 97 kDa | 0 | 0 | 8 | 1 | 9 | | | | | | | |
| AT3G06320 | | Ribosomal protein L33 family protein | 7 kDa | 0 | 0 | 8 | 1 | 9 | | | | | | | |
| AT3G10650 | | BEST Arabidopsis thaliana protein match is: nucleopori | 136 kDa | 0 | 0 | 8 | 1 | 9 | | | | | | | |
| AT3G59500 | | Integral membrane HRF1 family protein | 30 kDa | 0 | 0 | 8 | 1 | 9 | | | | | | | |
| AT5G14320 | | Ribosomal protein S13/S18 family | 19 kDa | 0 | 0 | 8 | 1 | 9 | | | | | | | |
| AT5G62600 | | ARM repeat superfamily protein | 107 kDa | 0 | 0 | 8 | 1 | 9 | | | | | | | |
| AT1G16740 | AT1G16740.1 | Ribosomal protein L20 | 15 kDa | 2 | 0 | 8 | 0 | 10 | | | | 2.2E+07 | 3.0E+07 | 1.4E+06 | 5.4E+07 |
| AT1G63770 | | Peptidase M1 family protein | 111 kDa | 0 | 0 | 7 | 20 | 27 | | | | | | | |
| AT2G39360 | | Protein kinase superfamily protein | 91 kDa | 0 | 0 | 7 | 19 | 26 | | | | | | | |
| AT5G52640 | | heat shock protein 90.1 | 81 kDa | 0 | 0 | 7 | 18 | 25 | | | | | | | |
| AT5G17000 | | Zinc-binding dehydrogenase family protein | 38 kDa | 0 | 0 | 7 | 17 | 24 | | | | | | | |
| AT1G47710 | | Serine protease inhibitor (SERPIN) family protein | 43 kDa | 0 | 0 | 7 | 14 | 21 | | | | | | | |
| AT2G42590 | AT2G42590.2;AT2G42590.1;AT2G42590.3 | general regulatory factor 9 | 30 kDa | 0 | 2 | 7 | 14 | 23 | | 3.25 | 3.25 | 1.1E+06 | 5.2E+05 | 0.0E+00 | 1.6E+06 |
| AT5G46290 | AT5G46290.3;AT5G46290.1;AT5G46290.2 | 3-ketoacyl-acyl carrier protein synthase I | 50 kDa | 2 | 1 | 7 | 14 | 24 | | | | 1.1E+07 | 9.8E+06 | 2.4E+05 | 2.1E+07 |
| AT1G54370 | | sodium hydrogen exchanger 5 | 57 kDa | 0 | 0 | 7 | 13 | 20 | | | | | | | |
| AT1G74020 | AT1G74020.1 | strictosidine synthase 2 | 35 kDa | 2 | 0 | 7 | 13 | 22 | | | | 2.7E+06 | 2.0E+06 | 3.6E+05 | 5.0E+06 |
| AT1G35470 | AT1G35470.1;AT1G35470.2 | SPla/RYanodine receptor (SPRY) domain-containing pro | 52 kDa | 0 | 0 | 7 | 12 | 19 | | | | 2.5E+06 | 1.5E+06 | 3.4E+04 | 4.1E+06 |
| AT2G38290 | AT2G38290.1 | ammonium transporter 2 | 51 kDa | 0 | 0 | 7 | 12 | 19 | | | | 1.8E+06 | 8.2E+05 | 0.0E+00 | 2.6E+06 |
| AT5G46330 | | Leucine-rich receptor-like protein kinase family protein | 129 kDa | 2 | 0 | 7 | 12 | 21 | | | | | | | |
| AT5G65620 | | Zincin-like metalloproteases family protein | 89 kDa | 1 | 0 | 7 | 12 | 20 | | | | | | | |
| AT1G02080 | | transcription regulators | 270 kDa | 0 | 0 | 7 | 11 | 18 | | | | | | | |
| AT1G44575 | AT1G44575.1;AT1G44575.3;AT1G44575.2 | Chlorophyll A-B binding family protein | 28 kDa | 0 | 0 | 7 | 11 | 18 | | 0.36 | 0.36 | 5.6E+06 | 1.4E+06 | 0.0E+00 | 7.1E+06 |
| AT3G22660 | AT3G22660.1 | rRNA processing protein-related | 33 kDa | 0 | 1 | 7 | 11 | 19 | 0.15 | 0.47 | 0.31 | 4.4E+06 | 5.8E+06 | 1.7E+06 | 1.2E+07 |
| AT5G39320 | | UDP-glucose 6-dehydrogenase family protein | 53 kDa | 0 | 2 | 7 | 11 | 20 | | | | | | | |
| AT5G50020 | AT5G50020.1;AT5G50020.3;AT5G50020.2;AT5G41060.4;AT5G41060.3;AT5G41060.2;AT4G24630.1;AT5G41060.1;AT3G48760.1 | DHHC-type zinc finger family protein | 47 kDa | 0 | 0 | 7 | 11 | 18 | | | | 4.9E+05 | 0.0E+00 | 0.0E+00 | 4.9E+05 |
| AT1G28400 | | unknown protein; FUNCTIONS IN: molecular_function | ? | 0 | 0 | 7 | 10 | 17 | | | | | | | |
| AT1G30630 | AT1G30630.1 | Coatomer epsilon subunit | 33 kDa | 0 | 8 | 7 | 10 | 25 | | | | 9.2E+06 | 6.9E+06 | 3.1E+05 | 1.6E+07 |
| AT2G44060 | | Late embryogenesis abundant protein, group 2 | 36 kDa | 1 | 3 | 7 | 10 | 21 | | | | | | | |
| AT2G44160 | | methylenetetrahydrofolate reductase 2 | 67 kDa | 0 | 0 | 7 | 10 | 17 | | | | | | | |
| AT3G52920 | | Family of unknown function (DUF662) | 21 kDa | 3 | 1 | 7 | 10 | 21 | | | | | | | |
| AT3G53030 | AT3G53030.1 | ser/arg-rich protein kinase 4 | 59 kDa | 0 | 0 | 7 | 10 | 17 | | | | 1.8E+06 | 1.3E+06 | 0.0E+00 | 3.1E+06 |
| AT3G55320 | | P-glycoprotein 20 | 155 kDa | 0 | 0 | 7 | 10 | 17 | | | | | | | |
| AT5G27470 | | seryl-tRNA synthetase / serine--tRNA ligase | 52 kDa | 0 | 0 | 7 | 10 | 17 | | | | | | | |
| AT2G37170 | | plasma membrane intrinsic protein 2 | 30 kDa | 0 | 1 | 7 | 9 | 17 | | | | | | | |
| AT2G39480 | | P-glycoprotein 6 | 156 kDa | 0 | 0 | 7 | 9 | 16 | | | | | | | |
| AT2G46560 | | transducin family protein / WD-40 repeat family protei | 276 kDa | 0 | 0 | 7 | 9 | 16 | | | | | | | |
| AT3G07540 | | Actin-binding FH2 (formin homology 2) family protein | 93 kDa | 0 | 0 | 7 | 9 | 16 | | | | | | | |
| AT4G18030 | AT4G18030.1 | S-adenosyl-L-methionine-dependent methyltransferase | 70 kDa | 0 | 3 | 7 | 9 | 19 | | | | | | | |
| AT1G11380 | | PLAC8 family protein | 28 kDa | 0 | 4 | 7 | 8 | 19 | | | | | | | |
| AT1G28290 | AT1G28290.2;AT1G28290.1 | arabinogalactan protein 31 | 38 kDa | 0 | 0 | 7 | 8 | 15 | | 0.10 | 0.10 | 1.4E+07 | 2.3E+07 | 9.2E+05 | 3.8E+07 |
| AT2G30870 | AT2G30870.1 | glutathione S-transferase PHI 10 | 24 kDa | 0 | 2 | 7 | 8 | 17 | | | | 3.2E+06 | 2.2E+06 | 0.0E+00 | 5.4E+06 |
| AT2G33210 | AT2G33210.2;AT2G33210.1 | heat shock protein 60-2 | 62 kDa | 0 | 0 | 7 | 8 | 15 | | | | 2.1E+06 | 2.2E+06 | 0.0E+00 | 4.4E+06 |
| AT3G06010 | | Homeotic gene regulator | 127 kDa | 0 | 2 | 7 | 8 | 17 | | | | | | | |
| AT3G09100 | | mRNA capping enzyme family protein | 79 kDa | 0 | 0 | 7 | 8 | 15 | | | | | | | |
| AT3G18000 | | S-adenosyl-L-methionine-dependent methyltransferase | 56 kDa | 0 | 0 | 7 | 8 | 15 | | | | | | | |
| AT3G51260 | AT3G51260.1;AT3G51260.2;AT5G66140.1 | 20S proteasome alpha subunit PAD1 | 27 kDa | 0 | 3 | 7 | 8 | 18 | 0.12 | | 0.12 | 9.6E+06 | 1.1E+07 | 5.1E+05 | 2.1E+07 |
| AT4G17830 | | Peptidase M20/M25/M40 family protein | 48 kDa | 0 | 1 | 7 | 8 | 16 | | | | | | | |
| AT5G07440 | | glutamate dehydrogenase 2 | 45 kDa | 0 | 0 | 7 | 8 | 15 | | | | | | | |
| AT5G22010 | | replication factor C1 | 104 kDa | 0 | 0 | 7 | 8 | 15 | | | | | | | |
| AT5G27670 | AT5G27670.1 | histone H2A 7 | 16 kDa | 5 | 0 | 7 | 8 | 20 | 0.17 | | 0.17 | 4.4E+07 | 6.3E+07 | 2.5E+05 | 1.1E+08 |
| AT5G63640 | | ENTH/VHS/GAT family protein | 49 kDa | 1 | 1 | 7 | 8 | 17 | | | | | | | |
| AT1G09970 | | Leucine-rich receptor-like protein kinase family protein | 107 kDa | 0 | 1 | 7 | 7 | 15 | | | | | | | |
| AT1G30440 | | Phototropic-responsive NPH3 family protein | 73 kDa | 0 | 0 | 7 | 7 | 14 | | | | | | | |

| | | | | | | | | | | | | | | | |
|---|---|---|---|---|---|---|---|---|---|---|---|---|---|---|---|
| AT1G55910 | | zinc transporter 11 precursor | 35 kDa | 0 | 0 | 7 | 7 | 14 | | | | | | | |
| AT3G13235 | | ubiquitin family protein | 45 kDa | 0 | 0 | 7 | 7 | 14 | | | | | | | |
| AT3G16840 | AT3G16840.1;AT3G16840.2 | P-loop containing nucleoside triphosphate hydrolases | 94 kDa | 0 | 0 | 7 | 7 | 14 | | | | 2.0E+05 | 1.5E+05 | 0.0E+00 | 3.6E+05 |
| AT3G53810 | | Concanavalin A-like lectin protein kinase family protein | 76 kDa | 0 | 0 | 7 | 7 | 14 | | | | | | | |
| AT3G55460 | AT3G55460.1 | SC35-like splicing factor 30 | 30 kDa | 0 | 1 | 7 | 7 | 15 | 0.08 | | 0.08 | 3.1E+06 | 2.5E+06 | 1.2E+06 | 6.8E+06 |
| AT4G38740 | | rotamase CYP 1 | 18 kDa | 1 | 0 | 7 | 7 | 15 | | | | | | | |
| AT5G01760 | | ENTH/VHS/GAT family protein | 60 kDa | 0 | 0 | 7 | 7 | 14 | | | | | | | |
| AT5G19600 | | sulfate transporter 3;5 | 70 kDa | 0 | 0 | 7 | 7 | 14 | | | | | | | |
| AT5G24520 | | Transducin/WD40 repeat-like superfamily protein | 38 kDa | 0 | 1 | 7 | 7 | 15 | | | | | | | |
| AT5G35590 | AT5G35590.1 | proteasome alpha subunit A1 | 27 kDa | 0 | 1 | 7 | 7 | 15 | 0.12 | 0.60 | 0.36 | 1.7E+06 | 9.9E+05 | 0.0E+00 | 2.6E+06 |
| AT5G66710 | | Protein kinase superfamily protein | 46 kDa | 0 | 2 | 7 | 7 | 16 | | | | | | | |
| ATCG00740 | ATCG00740.1 | RNA polymerase subunit alpha | 38 kDa | 0 | 2 | 7 | 7 | 16 | | | | 8.7E+05 | 0.0E+00 | 1.5E+05 | 1.0E+06 |
| AT1G08190 | AT1G08190.1 | vacuolar protein sorting 41 | 110 kDa | 0 | 0 | 7 | 6 | 13 | | | | | | | |
| AT1G12370 | AT1G12370.2;AT1G12370.1;AT1G12370.3 | photolyase 1 | 56 kDa | 0 | 0 | 7 | 6 | 13 | | | | 0.0E+00 | 1.8E+05 | 5.0E+04 | 2.3E+05 |
| AT1G25320 | | Leucine-rich repeat protein kinase family protein | 76 kDa | 0 | 0 | 7 | 6 | 13 | | | | | | | |
| AT1G30460 | AT1G30460.1;AT1G30460.2;AT1G30460.3 | cleavage and polyadenylation specificity factor 30 | 70 kDa | 0 | 0 | 7 | 6 | 13 | | | | 4.7E+05 | 1.3E+06 | 1.6E+05 | 1.9E+06 |
| AT1G49730 | | Protein kinase superfamily protein | 76 kDa | 0 | 0 | 7 | 6 | 13 | | | | | | | |
| AT1G51380 | | DEA(D/H)-box RNA helicase family protein | 44 kDa | 0 | 0 | 7 | 6 | 13 | | | | | | | |
| AT1G51660 | | mitogen-activated protein kinase kinase 4 | 40 kDa | 0 | 0 | 7 | 6 | 13 | | | | | | | |
| AT1G70330 | | equilibrative nucleotide transporter 1 | 49 kDa | 0 | 0 | 7 | 6 | 13 | | | | | | | |
| AT2G30490 | | cinnamate-4-hydroxylase | 58 kDa | 0 | 0 | 7 | 6 | 13 | | | | | | | |
| AT2G36950 | | Heavy metal transport/detoxification superfamily prot | 40 kDa | 0 | 0 | 7 | 6 | 13 | | | | | | | |
| AT2G44710 | AT2G44710.2;AT2G44710.1;AT2G44710.4;AT2G44710.3 | RNA-binding (RRM/RBD/RNP motifs) family protein | 90 kDa | 0 | 0 | 7 | 6 | 13 | | | | 1.1E+06 | 1.0E+06 | 1.2E+05 | 2.3E+06 |
| AT3G05050 | | Protein kinase superfamily protein | 67 kDa | 0 | 0 | 7 | 6 | 13 | | | | | | | |
| AT3G27850 | AT3G27850.1;AT3G27830.2;AT3G27830.1 | ribosomal protein L12-C | 20 kDa | 2 | 0 | 7 | 6 | 15 | | | | 8.2E+06 | 8.0E+06 | 7.1E+06 | 2.3E+07 |
| AT3G29160 | | SNF1 kinase homolog 11 | 59 kDa | 0 | 0 | 7 | 6 | 13 | | | | | | | |
| AT3G53390 | AT3G53390.1 | Transducin/WD40 repeat-like superfamily protein | 60 kDa | 0 | 0 | 7 | 6 | 13 | | | | 1.5E+05 | 1.9E+05 | 0.0E+00 | 3.3E+05 |
| AT3G63410 | AT3G63410.1 | S-adenosyl-L-methionine-dependent methyltransferase | 38 kDa | 4 | 4 | 7 | 6 | 21 | 0.07 | 0.45 | 0.26 | 2.9E+06 | 3.4E+06 | 0.0E+00 | 6.3E+06 |
| AT4G28480 | | DNAJ heat shock family protein | 38 kDa | 0 | 1 | 7 | 6 | 14 | | | | | | | |
| AT5G01500 | | thylakoid ATP/ADP carrier | 45 kDa | 0 | 0 | 7 | 6 | 13 | 0.18 | | 0.18 | | | | |
| AT5G13300 | | ARF GTPase-activating protein | 93 kDa | 1 | 0 | 7 | 6 | 14 | | | | | | | |
| AT5G17900 | AT5G17900.1;AT4G08580.2;AT4G08580.1 | microfibrillar-associated protein-related | 51 kDa | 0 | 0 | 7 | 6 | 13 | | | | 2.1E+05 | 5.0E+05 | 0.0E+00 | 7.0E+05 |
| AT5G58150 | | Leucine-rich repeat protein kinase family protein | 87 kDa | 0 | 0 | 7 | 6 | 13 | | | | | | | |
| AT1G13160 | | ARM repeat superfamily protein | 89 kDa | 0 | 0 | 7 | 5 | 12 | | | | | | | |
| AT1G29820 | | Magnesium transporter CorA-like family protein | 61 kDa | 0 | 0 | 7 | 5 | 12 | | | | | | | |
| AT1G48410 | AT1G48410.1;AT1G48410.3;AT1G48410.2 | Stabilizer of iron transporter SufD / Polynucleotidyl tra | 116 kDa | 0 | 0 | 7 | 5 | 12 | | 0.64 | 0.64 | 3.3E+06 | 3.5E+06 | 0.0E+00 | 6.8E+06 |
| AT2G20300 | | Protein kinase superfamily protein | 82 kDa | 0 | 0 | 7 | 5 | 12 | | | | | | | |
| AT2G29400 | | type one protein phosphatase 1 | 36 kDa | 0 | 3 | 7 | 5 | 15 | | | | | | | |
| AT3G04640 | | glycine-rich protein | 16 kDa | 1 | 0 | 7 | 5 | 13 | | | | | | | |
| AT3G13320 | | cation exchanger 2 | 48 kDa | 0 | 0 | 7 | 5 | 12 | | | | | | | |
| AT3G42725 | AT3G42725.1 | Putative membrane lipoprotein | 23 kDa | 0 | 0 | 7 | 5 | 12 | | | | 7.5E+06 | 4.1E+06 | 0.0E+00 | 1.2E+07 |
| AT4G09200 | | SPla/RYanodine receptor (SPRY) domain-containing pro | 44 kDa | 0 | 0 | 7 | 5 | 12 | | | | | | | |
| AT4G21100 | AT4G21100.1 | damaged DNA binding protein 1B | 121 kDa | 0 | 0 | 7 | 5 | 12 | | | | 0.0E+00 | 0.0E+00 | 2.7E+05 | 2.7E+05 |
| AT4G21110 | AT4G21110.2;AT4G21110.1 | G10 family protein | 17 kDa | 3 | 0 | 7 | 5 | 15 | | 0.26 | 0.26 | 1.0E+07 | 1.4E+07 | 2.5E+06 | 2.7E+07 |
| AT5G21090 | | Leucine-rich repeat (LRR) family protein | 24 kDa | 1 | 3 | 7 | 5 | 16 | | | | | | | |
| AT5G41190 | AT5G41190.1 | CONTAINS InterPro DOMAIN/s: Nin one binding (NOB1) | 67 kDa | 0 | 0 | 7 | 5 | 12 | | | | 1.8E+06 | 1.6E+06 | 1.9E+05 | 3.5E+06 |
| AT1G65980 | AT1G65980.1 | thioredoxin-dependent peroxidase 1 | 17 kDa | 2 | 0 | 7 | 4 | 13 | | | | 1.3E+07 | 4.5E+06 | 0.0E+00 | 1.8E+07 |
| AT1G76540 | AT1G76540.1 | cyclin-dependent kinase B2;1 | 36 kDa | 0 | 1 | 7 | 4 | 12 | | | | 1.2E+06 | 9.6E+05 | 4.2E+05 | 2.5E+06 |
| AT2G25280 | | CONTAINS InterPro DOMAIN/s: UPF0103/Mediator of E | 33 kDa | 0 | 5 | 7 | 4 | 16 | | | | | | | |
| AT2G28290 | AT2G28290.2;AT2G28290.4;AT2G28290.3;AT2G28290.6;AT2G28290.1;AT2G28290.5 | P-loop containing nucleoside triphosphate hydrolases | 390 kDa | 0 | 0 | 7 | 4 | 11 | | | | 4.0E+05 | 1.2E+06 | 3.3E+04 | 1.6E+06 |
| AT2G28390 | | SAND family protein | 67 kDa | 0 | 0 | 7 | 4 | 11 | | | | | | | |
| AT3G06730 | | Thioredoxin z | 21 kDa | 0 | 0 | 7 | 4 | 11 | | 0.26 | 0.26 | | | | |
| AT3G57280 | AT3G57280.1 | Transmembrane proteins 14C | 24 kDa | 0 | 0 | 7 | 4 | 11 | | 0.59 | 0.59 | 8.1E+05 | 1.2E+06 | 0.0E+00 | 2.0E+06 |
| AT4G02520 | AT4G02520.1;AT2G02930.1 | glutathione S-transferase PHI 2 | 24 kDa | 0 | 2 | 7 | 4 | 13 | | | | 1.3E+07 | 5.1E+06 | 0.0E+00 | 1.8E+07 |
| AT4G10730 | | Protein kinase superfamily protein | 78 kDa | 0 | 0 | 7 | 4 | 11 | | | | | | | |
| AT4G25550 | AT4G25550.1 | Cleavage/polyadenylation specificity factor, 25kDa sub | 23 kDa | 0 | 5 | 7 | 4 | 16 | | | | 1.5E+06 | 3.8E+06 | 0.0E+00 | 5.2E+06 |
| AT5G06910 | AT5G06910.1;AT3G12170.1;AT3G12170.2 | J-domain protein 6 | 33 kDa | 0 | 0 | 7 | 4 | 11 | | | | | | | |
| AT5G18580 | | tonneau 2 (TON2) | 55 kDa | 0 | 0 | 7 | 4 | 11 | | | | | | | |
| AT5G51130 | | S-adenosyl-L-methionine-dependent methyltransferase | 37 kDa | 0 | 0 | 7 | 4 | 11 | | | | | | | |
| AT5G59730 | | exocyst subunit exo70 family protein H7 | 71 kDa | 0 | 0 | 7 | 4 | 11 | | | | | | | |
| AT5G59950 | AT5G59950.1;AT5G59950.4;AT5G59950.3;AT5G59950.5;AT5G59950.2 | RNA-binding (RRM/RBD/RNP motifs) family protein | 26 kDa | 0 | 4 | 7 | 4 | 15 | | | | 7.0E+06 | 1.1E+07 | 2.0E+06 | 2.0E+07 |
| AT5G63750 | | RING/U-box superfamily protein | 62 kDa | 0 | 0 | 7 | 4 | 11 | | | | | | | |
| AT5G63880 | AT5G63880.1;AT5G63880.2 | SNF7 family protein | 25 kDa | 2 | 0 | 7 | 4 | 13 | | | | 1.1E+06 | 5.0E+05 | 3.9E+05 | 2.0E+06 |
| ATCG00770 | ATCG00770.1 | ribosomal protein S8 | 15 kDa | 0 | 0 | 7 | 4 | 11 | | 0.45 | 0.45 | 4.9E+06 | 2.2E+06 | 4.5E+05 | 7.5E+06 |
| AT1G01320 | AT1G01320.2;AT1G01320.3;AT1G01320.1 | Tetratricopeptide repeat (TPR)-like superfamily protein | 199 kDa | 0 | 0 | 7 | 3 | 10 | | | | 4.0E+05 | 2.1E+05 | 0.0E+00 | 6.2E+05 |
| AT1G18250 | AT1G18250.1;AT1G18250.2 | Pathogenesis-related thaumatin superfamily protein | 26 kDa | 0 | 0 | 7 | 3 | 10 | | | | 5.9E+06 | 6.4E+06 | 4.6E+06 | 1.7E+07 |
| AT1G55540 | | Nuclear pore complex protein | 193 kDa | 0 | 1 | 7 | 3 | 11 | | | | | | | |
| AT1G65580 | | Endonuclease/exonuclease/phosphatase family protein | 122 kDa | 0 | 1 | 7 | 3 | 11 | | | | | | | |
| AT1G80530 | | Major facilitator superfamily protein | 61 kDa | 0 | 0 | 7 | 3 | 10 | | | | | | | |
| AT2G18630 | | Protein of unknown function (DUF677) | 45 kDa | 0 | 0 | 7 | 3 | 10 | | | | | | | |
| AT2G27020 | | 20S proteasome alpha subunit G1 | 27 kDa | 0 | 0 | 7 | 3 | 10 | | | | | | | |
| AT3G19120 | | PIF / Ping-Pong family of plant transposases | 49 kDa | 0 | 0 | 7 | 3 | 10 | | | | | | | |
| AT3G22300 | AT3G22300.1 | ribosomal protein S10 | 27 kDa | 0 | 0 | 7 | 3 | 10 | | | | 2.2E+07 | 2.6E+07 | 1.5E+06 | 5.0E+07 |
| AT3G22440 | AT3G22440.1 | FRIGIDA-like protein | 59 kDa | 0 | 0 | 7 | 3 | 10 | | | | 7.8E+05 | 4.0E+05 | 6.9E+05 | 1.9E+06 |
| AT3G49560 | AT3G49560.1 | Mitochondrial import inner membrane translocase sub | 28 kDa | 0 | 1 | 7 | 3 | 11 | | | | 4.0E+06 | 4.4E+06 | 0.0E+00 | 8.4E+06 |
| AT4G17510 | AT4G17510.1 | ubiquitin C-terminal hydrolase 3 | 26 kDa | 0 | 2 | 7 | 3 | 12 | | | | 4.6E+06 | 3.1E+06 | 0.0E+00 | 7.7E+06 |
| AT5G06160 | | splicing factor-related | 59 kDa | 0 | 0 | 7 | 3 | 10 | | 0.32 | 0.32 | | | | |
| AT5G14680 | AT5G14680.1 | Adenine nucleotide alpha hydrolases-like superfamily p | 20 kDa | 0 | 0 | 7 | 3 | 10 | | | | 6.9E+04 | 1.1E+05 | 0.0E+00 | 1.8E+05 |
| AT5G19820 | AT5G19820.1 | ARM repeat superfamily protein | 124 kDa | 0 | 0 | 7 | 3 | 10 | | | | 7.5E+05 | 4.0E+05 | 0.0E+00 | 1.1E+06 |
| AT5G58950 | | Protein kinase superfamily protein | 59 kDa | 0 | 0 | 7 | 3 | 10 | | | | | | | |
| AT5G63960 | AT5G63960.1;AT5G63960.2 | DNA binding;nucleotide binding;nucleic acid binding;I | 123 kDa | 0 | 0 | 7 | 3 | 10 | | | | 7.2E+06 | 5.6E+06 | 1.0E+05 | 1.3E+07 |
| ATCG00065 | | ribosomal protein S12A | 14 kDa | 0 | 0 | 7 | 3 | 10 | | | | | | | |
| AT1G41830 | | SKU5-similar 6 | 61 kDa | 0 | 0 | 7 | 2 | 9 | | | | | | | |
| AT1G55190 | | PRA1 (Prenylated rab acceptor) family protein | 21 kDa | 0 | 0 | 7 | 2 | 9 | | | | | | | |
| AT1G62820 | | Calcium-binding EF-hand family protein | 17 kDa | 2 | 0 | 7 | 2 | 11 | 2.69 | | 2.69 | | | | |
| AT2G27210 | | BRI1 suppressor 1 (BSU1)-like 3 | 108 kDa | 0 | 0 | 7 | 2 | 9 | | | | | | | |
| AT3G02950 | AT3G02950.2;AT3G02950.1 | Tho complex subunit 7/Mft1p | 27 kDa | 0 | 0 | 7 | 2 | 9 | | | | 1.5E+06 | 1.5E+06 | 3.0E+05 | 3.3E+06 |
| AT3G13230 | AT3G13230.2;AT3G13230.1 | RNA-binding KH domain-containing protein | 24 kDa | 0 | 0 | 7 | 2 | 9 | | | | 3.0E+06 | 2.8E+06 | 1.5E+06 | 7.3E+06 |

| | | | | | | | | | | | | | | | |
|---|---|---|---|---|---|---|---|---|---|---|---|---|---|---|---|
| AT3G13530 | AT3G13530.1 | mitogen-activated protein kinase kinase kinase 7 | 151 kDa | 0 | 0 | 7 | 2 | 9 | | | | | | | |
| AT3G15480 | | Protein of unknown function (DUF1218) | 19 kDa | 5 | 0 | 7 | 2 | 14 | | | | | | | |
| AT3G55610 | | delta 1-pyrroline-5-carboxylate synthase 2 | 79 kDa | 0 | 0 | 7 | 2 | 9 | | | | | | | |
| AT4G12735 | | unknown protein; FUNCTIONS IN: molecular_function | ? | 0 | 0 | 7 | 2 | 9 | | | | | | | |
| AT4G34320 | | Protein of unknown function (DUF677) | 42 kDa | 0 | 0 | 7 | 2 | 9 | | | | | | | |
| AT4G37460 | | Tetratricopeptide repeat (TPR)-like superfamily protein | 118 kDa | 0 | 0 | 7 | 2 | 9 | | | | | | | |
| AT5G37720 | AT5G37720.2;AT5G37720.1 | ALWAYS EARLY 4 | 30 kDa | 0 | 0 | 7 | 2 | 9 | | | | 9.0E+05 | 9.5E+05 | 1.0E+06 | 2.9E+06 |
| AT5G40040 | AT5G40040.1 | 60S acidic ribosomal protein family | 12 kDa | 0 | 0 | 7 | 2 | 9 | | | | 3.9E+07 | 4.7E+07 | 1.4E+08 | 2.2E+08 |
| AT1G67680 | | SRP72 RNA-binding domain | 73 kDa | 0 | 0 | 7 | 1 | 8 | | | | | | | |
| AT2G01980 | | sodium proton exchanger, putative (NHX7) (SOS1) | 127 kDa | 0 | 0 | 7 | 1 | 8 | | | | | | | |
| AT3G12170 | | Chaperone DnaJ-domain superfamily protein | 30 kDa | 0 | 0 | 7 | 1 | 8 | | | | | | | |
| AT5G06600 | | ubiquitin-specific protease 12 | 131 kDa | 0 | 0 | 7 | 1 | 8 | | | | | | | |
| AT5G08120 | | movement protein binding protein 2C | 37 kDa | 0 | 0 | 7 | 1 | 8 | | | | | | | |
| AT5G19230 | | Glycoprotein membrane precursor GPI-anchored | 21 kDa | 4 | 8 | 7 | 1 | 20 | | | | | | | |
| AT5G48650 | AT5G48650.1 | Nuclear transport factor 2 (NTF2) family protein with R | 50 kDa | 0 | 0 | 7 | 1 | 8 | | | | 2.2E+06 | 1.3E+06 | 1.9E+05 | 3.7E+06 |
| AT1G02930 | AT1G02930.1;AT1G02930.2;AT1G02920.1 | glutathione S-transferase 6 | 23 kDa | 0 | 1 | 7 | 0 | 8 | | | | 2.8E+06 | 1.3E+07 | 7.9E+04 | 1.6E+07 |
| AT2G34520 | AT2G34520.1 | mitochondrial ribosomal protein S14 | 19 kDa | 1 | 0 | 7 | 0 | 8 | | 6.36 | 6.36 | 1.0E+07 | 1.9E+07 | 1.2E+06 | 3.0E+07 |
| AT5G63905 | AT5G63905.1;AT5G63905.2 | unknown protein; FUNCTIONS IN: molecular_function | 8 kDa | 2 | 0 | 7 | 0 | 9 | | | | | | | |
| AT3G13772 | | transmembrane nine 7 | 74 kDa | 0 | 0 | 6 | 19 | 25 | | | | | | | |
| AT1G26730 | | EXS (ERD1/XPR1/SYG1) family protein | 87 kDa | 0 | 0 | 6 | 17 | 23 | | | | | | | |
| AT2G47710 | AT2G47710.1 | Adenine nucleotide alpha hydrolases-like superfamily p | ? | 5 | 1 | 6 | 15 | 27 | | | | 5.1E+06 | 3.5E+06 | 4.9E+05 | 9.2E+06 |
| AT1G66970 | AT1G66970.3;AT1G66970.1;AT1G66970.2 | SHV3-like 2 | 84 kDa | 0 | 2 | 6 | 14 | 22 | | | | 9.6E+05 | 0.0E+00 | 0.0E+00 | 9.6E+05 |
| AT2G01720 | | Ribophorin I | 52 kDa | 0 | 0 | 6 | 14 | 20 | | 0.51 | 0.51 | | | | |
| AT1G43800 | AT1G43800.1 | Plant stearoyl-acyl-carrier-protein desaturase family pr | 44 kDa | 0 | 3 | 6 | 13 | 22 | | | | | | | |
| AT1G77110 | | Auxin efflux carrier family protein | 62 kDa | 0 | 0 | 6 | 13 | 19 | | | | | | | |
| AT3G02050 | AT3G02050.1 | K+ uptake transporter 3 | 87 kDa | 0 | 0 | 6 | 13 | 19 | | | | | | | |
| AT1G06020 | | pfkB-like carbohydrate kinase family protein | 38 kDa | 0 | 0 | 6 | 11 | 17 | | | | | | | |
| AT1G17070 | AT1G17070.1 | GC-rich sequence DNA-binding factor-like protein with | 97 kDa | 0 | 0 | 6 | 11 | 17 | | | | 0.0E+00 | 8.7E+04 | 4.4E+04 | 1.3E+05 |
| AT1G19110 | | inter-alpha-trypsin inhibitor heavy chain-related | ? | 0 | 0 | 6 | 11 | 17 | | | | | | | |
| AT5G60490 | AT5G60490.1 | FASCICLIN-like arabinogalactan-protein 12 | 26 kDa | 0 | 0 | 6 | 11 | 17 | | | | 7.3E+06 | 0.0E+00 | 0.0E+00 | 7.3E+06 |
| AT1G12000 | AT1G12000.1 | Phosphofructokinase family protein | 61 kDa | 0 | 0 | 6 | 10 | 16 | | | | 1.9E+06 | 1.9E+06 | 0.0E+00 | 3.8E+06 |
| AT2G39750 | | S-adenosyl-L-methionine-dependent methyltransferase | 78 kDa | 0 | 1 | 6 | 10 | 17 | | | | | | | |
| AT3G48710 | | DEK domain-containing chromatin associated protein | 52 kDa | 0 | 0 | 6 | 10 | 16 | | | | | | | |
| AT1G48900 | AT1G48900.2;AT1G48900.1;AT5G49500.1 | Signal recognition particle, SRP54 subunit protein | 55 kDa | 0 | 0 | 6 | 9 | 15 | | | | 9.5E+05 | 2.7E+06 | 5.3E+05 | 4.2E+06 |
| AT2G32810 | AT2G32810.2;AT2G32810.1 | beta galactosidase 9 | 99 kDa | 0 | 0 | 6 | 9 | 15 | | | | 4.5E+05 | 3.8E+05 | 0.0E+00 | 8.3E+05 |
| AT3G02360 | | 6-phosphogluconate dehydrogenase family protein | 54 kDa | 0 | 0 | 6 | 9 | 15 | | | | | | | |
| AT3G10960 | | AZA-guanine resistant1 | 62 kDa | 0 | 0 | 6 | 9 | 15 | | | | | | | |
| AT4G17620 | AT4G17620.1;AT4G17620.2;AT4G17620.3 | glycine-rich protein | 60 kDa | 0 | 1 | 6 | 9 | 16 | | | | 3.2E+05 | 4.9E+05 | 7.1E+05 | 1.5E+06 |
| AT4G24220 | AT4G24220.2;AT4G24220.1 | NAD(P)-binding Rossmann-fold superfamily protein | 44 kDa | 0 | 0 | 6 | 9 | 15 | | | | 4.0E+05 | 2.9E+05 | 0.0E+00 | 6.9E+05 |
| AT4G25500 | | arginine/serine-rich splicing factor 35 | 40 kDa | 0 | 0 | 6 | 9 | 15 | | | | | | | |
| AT5G62200 | AT5G62200.1 | Embryo-specific protein 3, (ATS3) | 21 kDa | 4 | 1 | 6 | 9 | 20 | | 0.41 | 0.41 | | | | |
| ATCG00170 | | DNA-directed RNA polymerase family protein | 156 kDa | 0 | 0 | 6 | 9 | 15 | | | | | | | |
| AT1G03060 | | Beige/BEACH domain ;WD domain, G-beta repeat prote | 401 kDa | 0 | 0 | 6 | 8 | 14 | | | | | | | |
| AT1G12110 | | nitrate transporter 1.1 | 65 kDa | 0 | 0 | 6 | 8 | 14 | | | | | | | |
| AT2G41530 | AT2G41530.1 | S-formylglutathione hydrolase | 32 kDa | 1 | 3 | 6 | 8 | 18 | | | | | | | |
| AT3G12490 | AT3G12490.1;AT3G12490.2 | cystatin B | 22 kDa | 0 | 1 | 6 | 8 | 15 | 0.52 | 3.77 | 2.15 | 3.6E+06 | 2.3E+06 | 0.0E+00 | 5.9E+06 |
| AT4G18905 | AT4G18905.1;AT4G18905.2 | Transducin/WD40 repeat-like superfamily protein | 54 kDa | 0 | 0 | 6 | 8 | 14 | | | | 6.3E+05 | 3.9E+05 | 0.0E+00 | 1.0E+06 |
| AT4G39910 | AT4G39910.1 | ubiquitin-specific protease 3 | 42 kDa | 0 | 0 | 6 | 8 | 14 | | | | 1.4E+06 | 5.5E+05 | 0.0E+00 | 2.0E+06 |
| AT5G07350 | | TUDOR-SN protein 1 | 108 kDa | 0 | 0 | 6 | 8 | 14 | | | | | | | |
| AT5G48300 | | ADP glucose pyrophosphorylase 1 | 57 kDa | 0 | 0 | 6 | 8 | 14 | | | | | | | |
| AT1G11090 | AT1G11090.1 | alpha/beta-Hydrolases superfamily protein | 37 kDa | 0 | 3 | 6 | 7 | 16 | | | | 4.5E+05 | 4.7E+05 | 0.0E+00 | 9.2E+05 |
| AT2G21660 | AT2G21660.2;AT2G21660.1 | cold, circadian rhythm, and rna binding 2 | 17 kDa | 2 | 0 | 6 | 7 | 15 | | 0.69 | 0.69 | 7.5E+05 | 6.5E+05 | 5.1E+05 | 1.9E+06 |
| AT2G37560 | | origin recognition complex second largest subunit 2 | 41 kDa | 0 | 0 | 6 | 7 | 13 | | | | | | | |
| AT3G06270 | | Protein phosphatase 2C family protein | 38 kDa | 0 | 1 | 6 | 7 | 14 | | 2.82 | 2.82 | | | | |
| AT3G15000 | AT3G15000.1 | cobalt ion binding | ? | 0 | 7 | 6 | 7 | 20 | | 0.36 | 0.36 | 1.0E+07 | 1.7E+07 | 2.8E+06 | 2.9E+07 |
| AT3G18440 | | aluminum-activated malate transporter 9 | 67 kDa | 0 | 0 | 6 | 7 | 13 | | | | | | | |
| AT3G21220 | | MAP kinase kinase 5 | 38 kDa | 0 | 0 | 6 | 7 | 13 | | | | | | | |
| AT5G52280 | | Myosin heavy chain-related protein | 99 kDa | 0 | 0 | 6 | 7 | 13 | | 0.46 | 0.46 | | | | |
| AT5G60270 | | Concanavalin A-like lectin protein kinase family protein | 74 kDa | 0 | 2 | 6 | 7 | 15 | | | | | | | |
| AT1G03190 | | RAD3-like DNA-binding helicase protein | 86 kDa | 0 | 0 | 6 | 6 | 12 | | | | | | | |
| AT1G36310 | | S-adenosyl-L-methionine-dependent methyltransferase | 45 kDa | 0 | 0 | 6 | 6 | 12 | | | | | | | |
| AT2G01080 | AT2G01080.1 | Late embryogenesis abundant (LEA) hydroxyproline-ric | 25 kDa | 0 | 0 | 6 | 6 | 12 | | | | 0.0E+00 | 9.4E+05 | 0.0E+00 | 9.4E+05 |
| AT3G22750 | | Protein kinase superfamily protein | 43 kDa | 0 | 1 | 6 | 6 | 13 | | | | | | | |
| AT3G23300 | | S-adenosyl-L-methionine-dependent methyltransferase | 69 kDa | 0 | 0 | 6 | 6 | 12 | | | | | | | |
| AT3G58500 | | protein phosphatase 2A-4 | 36 kDa | 0 | 0 | 6 | 6 | 12 | | | | | | | |
| AT3G58600 | | Adaptin ear-binding coat-associated protein 1 NECAP-1 | 34 kDa | 0 | 0 | 6 | 6 | 12 | | | | | | | |
| AT3G63490 | | Ribosomal protein L1p/L10e family | 38 kDa | 0 | 1 | 6 | 6 | 13 | | | | | | | |
| AT4G23740 | | Leucine-rich repeat protein kinase family protein | 71 kDa | 0 | 0 | 6 | 6 | 12 | | | | | | | |
| AT4G26630 | AT4G26630.2;AT4G26630.1 | DEK domain-containing chromatin associated protein | 85 kDa | 0 | 1 | 6 | 6 | 13 | | | | 2.7E+05 | 1.7E+05 | 2.2E+05 | 6.6E+05 |
| AT5G14210 | | Leucine-rich repeat protein kinase family protein | 86 kDa | 0 | 0 | 6 | 6 | 12 | | | | | | | |
| AT5G16750 | AT5G16750.1 | Transducin family protein / WD-40 repeat family protei | 97 kDa | 0 | 0 | 6 | 6 | 12 | | | | 7.4E+05 | 3.2E+05 | 1.1E+05 | 1.2E+06 |
| AT5G50520 | | Major facilitator superfamily protein | 59 kDa | 0 | 0 | 6 | 6 | 12 | | | | | | | |
| AT5G57830 | | Protein of unknown function, DUF593 | 44 kDa | 0 | 0 | 6 | 6 | 12 | | | | | | | |
| AT5G58410 | AT5G58410.2;AT5G58410.1 | HEAT/U-box domain-containing protein | 209 kDa | 0 | 0 | 6 | 6 | 12 | | | | 2.0E+05 | 2.2E+05 | 0.0E+00 | 4.1E+05 |
| AT5G61240 | | Leucine-rich repeat (LRR) family protein | 37 kDa | 0 | 3 | 6 | 6 | 15 | | | | | | | |
| AT5G65220 | AT5G65220.1 | Ribosomal L29 family protein | 19 kDa | 2 | 0 | 6 | 6 | 14 | | 1.11 | 1.11 | | | | |
| AT1G08770 | | prenylated RAB acceptor 1.E | 22 kDa | 0 | 0 | 6 | 5 | 11 | 0.83 | 1.18 | 1.00 | | | | |
| AT1G30230 | AT1G30230.1;AT1G30230.2 | Glutathione S-transferase, C-terminal-like;Translation e | 25 kDa | 0 | 0 | 6 | 5 | 11 | | | | 1.6E+07 | 2.4E+07 | 0.0E+00 | 4.0E+07 |
| AT1G30960 | AT1G30960.1 | GTP-binding family protein | 50 kDa | 0 | 0 | 6 | 5 | 11 | | | | 1.2E+06 | 1.8E+06 | 0.0E+00 | 3.0E+06 |
| AT1G48940 | AT1G48940.1 | early nodulin-like protein 6 | 20 kDa | 0 | 0 | 6 | 5 | 11 | | | | 1.3E+07 | 1.2E+07 | 4.7E+05 | 2.5E+07 |
| AT1G51760 | | peptidase M20/M25/M40 family protein | 48 kDa | 0 | 0 | 6 | 5 | 11 | | | | | | | |
| AT1G56590 | | Clathrin adaptor complexes medium subunit family pr | 46 kDa | 0 | 0 | 6 | 5 | 11 | | | | | | | |
| AT1G80620 | AT1G80620.1 | S15/NS1, RNA-binding protein | 47 kDa | 0 | 2 | 6 | 5 | 13 | | | | 1.0E+07 | 1.7E+07 | 5.4E+05 | 2.8E+07 |
| AT2G23890 | AT2G23890.3;AT2G23890.1;AT2G23890.2 | HAD-superfamily hydrolase, subfamily IG, 5'-nucleotida | 64 kDa | 0 | 2 | 6 | 5 | 13 | | | | | | | |
| AT2G28600 | AT2G28600.1 | P-loop containing nucleoside triphosphate hydrolases | 55 kDa | 0 | 0 | 6 | 5 | 11 | | | | 9.5E+04 | 0.0E+00 | 4.6E+04 | 1.4E+05 |
| AT3G56490 | | HIS triad family protein 3 | 16 kDa | 2 | 0 | 6 | 5 | 13 | | | | | | | |
| AT3G56940 | | dicarboxylate diiron protein, putative (Crd1) | 48 kDa | 0 | 1 | 6 | 5 | 12 | | | | | | | |
| AT4G28770 | AT4G28770.1;AT4G28770.2 | Tetraspanin family protein | 36 kDa | 0 | 0 | 6 | 5 | 11 | | | | | | | |
| AT4G33400 | | Vacuolar import/degradation, Vid27-related protein | 72 kDa | 0 | 0 | 6 | 5 | 11 | | | | | | | |
| AT4G38540 | | FAD/NAD(P)-binding oxidoreductase family protein | 46 kDa | 0 | 0 | 6 | 5 | 11 | | | | | | | |
| AT4G39955 | | alpha/beta-Hydrolases superfamily protein | 38 kDa | 0 | 0 | 6 | 5 | 11 | | | | | | | |
| AT5G27150 | AT5G27150.1;AT3G05030.4;AT3G05030.2;AT3G05030.1 | Na+/H+ exchanger 1 | 60 kDa | 0 | 0 | 6 | 5 | 11 | | | | 1.9E+06 | 4.7E+05 | 7.4E+04 | 2.5E+06 |
| AT5G42860 | | unknown protein; FUNCTIONS IN: molecular_function | ? | 0 | 0 | 6 | 5 | 11 | | | | | | | |
| AT5G51400 | AT5G51400.1 | PLAC8 family protein | 27 kDa | 2 | 6 | 6 | 5 | 19 | | | | 2.3E+06 | 2.1E+06 | 3.3E+06 | 7.6E+06 |

| | | | | | | | | | | | | | | | |
|---|---|---|---|---|---|---|---|---|---|---|---|---|---|---|---|
| AT5G60040 | AT5G60040.1;AT5G60040.2 | nuclear RNA polymerase C1 | 155 kDa | 0 | 0 | 6 | 5 | 11 | | | | 6.6E+04 | 2.8E+04 | 0.0E+00 | 9.4E+04 |
| AT1G01960 | AT1G01960.1 | SEC7-like guanine nucleotide exchange family protein | 195 kDa | 0 | 0 | 6 | 4 | 10 | | | | 4.3E+04 | 4.4E+04 | 0.0E+00 | 8.7E+04 |
| AT1G17500 | | ATPase E1-E2 type family protein / haloacid dehalogena | 138 kDa | 0 | 0 | 6 | 4 | 10 | | | | | | | |
| AT1G35580 | | cytosolic invertase 1 | 63 kDa | 0 | 0 | 6 | 4 | 10 | | | | | | | |
| AT1G47290 | | 3beta-hydroxysteroid-dehydrogenase/decarboxylase is | 48 kDa | 0 | 0 | 6 | 4 | 10 | | | | | | | |
| AT1G78040 | | Pollen Ole e 1 allergen and extensin family protein | 19 kDa | 4 | 0 | 6 | 4 | 14 | | | | | | | |
| AT2G27970 | AT2G27970.1;AT2G27960.1 | CDK-subunit 2 | 10 kDa | 0 | 0 | 6 | 4 | 10 | | | | | | | |
| AT2G35920 | AT2G35920.2;AT2G35920.1;AT2G35920.3 | RNA helicase family protein | 111 kDa | 0 | 0 | 6 | 4 | 10 | | | | 4.6E+06 | 3.0E+06 | 7.6E+05 | 8.4E+06 |
| AT2G46150 | | Late embryogenesis abundant (LEA) hydroxyproline-ric | 24 kDa | 0 | 2 | 6 | 4 | 12 | | | | | | | |
| AT3G05970 | | long-chain acyl-CoA synthetase 6 | 77 kDa | 0 | 0 | 6 | 4 | 10 | | | | | | | |
| AT3G11050 | AT3G11050.1 | ferritin 2 | 28 kDa | 0 | 0 | 6 | 4 | 10 | | | | 1.2E+07 | 4.4E+06 | 2.1E+05 | 1.7E+07 |
| AT3G20800 | | Cell differentiation, Rcd1-like protein | 35 kDa | 0 | 0 | 6 | 4 | 10 | | | | | | | |
| AT3G50930 | | cytochrome BC1 synthesis | 66 kDa | 0 | 0 | 6 | 4 | 10 | | | | | | | |
| AT5G19090 | | Heavy metal transport/detoxification superfamily prot | 60 kDa | 0 | 0 | 6 | 4 | 10 | | | | | | | |
| AT5G63910 | | farnesylcysteine lyase | 55 kDa | 0 | 0 | 6 | 4 | 10 | | | | | | | |
| ATCG00800 | ATCG00800.1 | structural constituent of ribosome | 25 kDa | 0 | 0 | 6 | 4 | 10 | | | | 2.4E+06 | 1.6E+06 | 1.9E+05 | 4.2E+06 |
| AT1G07210 | AT1G07210.1 | Ribosomal protein S18 | 30 kDa | 0 | 1 | 6 | 3 | 10 | | 3.74 | 3.74 | 2.2E+07 | 3.3E+07 | 1.8E+06 | 5.7E+07 |
| AT1G37130 | AT1G37130.1 | nitrate reductase 2 | 103 kDa | 0 | 0 | 6 | 3 | 9 | | | | 1.5E+06 | 1.2E+06 | 0.0E+00 | 2.7E+06 |
| AT1G65610 | | Six-hairpin glycosidases superfamily protein | 70 kDa | 1 | 0 | 6 | 3 | 10 | | | | | | | |
| AT1G69270 | | receptor-like protein kinase 1 | 60 kDa | 0 | 0 | 6 | 3 | 9 | | | | | | | |
| AT2G26600 | | Glycosyl hydrolase superfamily protein | 43 kDa | 0 | 0 | 6 | 3 | 9 | | | | | | | |
| AT2G46140 | AT2G46140.1 | Late embryogenesis abundant protein | 18 kDa | 2 | 0 | 6 | 3 | 11 | | | | 4.0E+06 | 1.8E+06 | 0.0E+00 | 5.8E+06 |
| AT3G26080 | | plastid-lipid associated protein PAP / fibrillin family pr | 26 kDa | 0 | 0 | 6 | 3 | 9 | | | | | | | |
| AT3G26420 | AT3G26420.1 | RNA-binding (RRM/RBD/RNP motifs) family protein wit | 27 kDa | 0 | 0 | 6 | 3 | 9 | | | | 5.9E+06 | 1.9E+07 | 2.2E+06 | 2.7E+07 |
| AT3G46960 | AT3G46960.1 | RNA helicase, ATP-dependent, SK12/DOB1 protein | 151 kDa | 0 | 0 | 6 | 3 | 9 | | | | | | | |
| AT4G22240 | AT4G22240.1 | Plastid-lipid associated protein PAP / fibrillin family pr | 34 kDa | 0 | 1 | 6 | 3 | 10 | 0.07 | 0.17 | 0.12 | | | | |
| AT4G27650 | | Eukaryotic release factor 1 (eRF1) family protein | 43 kDa | 0 | 0 | 6 | 3 | 9 | | | | | | | |
| AT5G16690 | | origin recognition complex subunit 3 | 83 kDa | 0 | 0 | 6 | 3 | 9 | | | | | | | |
| AT5G26667 | | P-loop containing nucleoside triphosphate hydrolases | 22 kDa | 0 | 2 | 6 | 3 | 11 | | | | | | | |
| AT5G37710 | | alpha/beta-Hydrolases superfamily protein | 49 kDa | 0 | 0 | 6 | 3 | 9 | | | | | | | |
| AT5G66750 | | chromatin remodeling 1 | 87 kDa | 0 | 0 | 6 | 3 | 9 | | | | | | | |
| ATCG00160 | ATCG00160.1 | ribosomal protein S2 | 27 kDa | 0 | 3 | 6 | 3 | 12 | | 0.23 | 0.23 | | | | |
| ATCG00780 | | ribosomal protein L14 | 14 kDa | 0 | 0 | 6 | 3 | 9 | | 0.30 | 0.30 | | | | |
| AT1G07140 | | Pleckstrin homology (PH) domain superfamily protein | 26 kDa | 0 | 1 | 6 | 2 | 9 | | | | | | | |
| AT1G10140 | | Uncharacterised conserved protein UCP031279 | 18 kDa | 0 | 0 | 6 | 2 | 8 | | | | | | | |
| AT1G15200 | AT1G15200.1;AT1G15200.2;AT1G15200.4;AT1G15200.3 | protein-protein interaction regulator family protein | 49 kDa | 0 | 0 | 6 | 2 | 8 | | | | 1.4E+05 | 7.4E+04 | 4.0E+05 | 6.1E+05 |
| AT1G16520 | | unknown protein; FUNCTIONS IN: molecular_function | ? | 0 | 0 | 6 | 2 | 8 | | | | | | | |
| AT1G20220 | | Alba DNA/RNA-binding protein | 34 kDa | 0 | 0 | 6 | 2 | 8 | | 0.48 | 0.48 | | | | |
| AT1G23860 | AT1G23860.3;AT1G23860.4;AT1G23860.2;AT1G23860.1 | RS-containing zinc finger protein 21 | 22 kDa | 2 | 6 | 6 | 2 | 16 | | | | 1.5E+06 | 0.0E+00 | 1.6E+05 | 1.7E+06 |
| AT2G33610 | | switch subunit 3 | 52 kDa | 0 | 0 | 6 | 2 | 8 | | | | | | | |
| AT3G11910 | | ubiquitin-specific protease 13 | 131 kDa | 1 | 0 | 6 | 2 | 9 | | | | | | | |
| AT4G36680 | AT4G36680.1 | Tetratricopeptide repeat (TPR)-like superfamily protein | 47 kDa | 0 | 0 | 6 | 2 | 8 | | | | | | | |
| AT4G39680 | | SAP domain-containing protein | 69 kDa | 0 | 1 | 6 | 2 | 9 | | | | | | | |
| AT5G45140 | AT5G45140.1;AT5G45140.2;AT5G45140.3 | nuclear RNA polymerase C2 | 130 kDa | 0 | 0 | 6 | 2 | 8 | | | | 1.9E+06 | 1.3E+06 | 1.8E+05 | 3.5E+06 |
| ATCG00830 | | ribosomal protein L2 | 30 kDa | 0 | 0 | 6 | 2 | 8 | | | | | | | |
| AT1G07170 | | PHF5-like protein | 12 kDa | 0 | 0 | 6 | 1 | 7 | | | | | | | |
| AT1G51650 | AT1G51650.1 | ATP synthase epsilon chain, mitochondrial | 8 kDa | 0 | 0 | 6 | 1 | 7 | | 0.26 | 0.26 | 7.0E+05 | 3.8E+06 | 0.0E+00 | 4.5E+06 |
| AT1G61620 | AT1G61620.1 | phosphoinositide binding | 35 kDa | 0 | 0 | 6 | 1 | 7 | | | | 1.7E+06 | 1.9E+06 | 8.2E+04 | 3.6E+06 |
| AT1G64600 | AT1G64600.1 | methyltransferases;copper ion binding | 62 kDa | 0 | 0 | 6 | 1 | 7 | | | | 1.6E+06 | 3.1E+06 | 7.0E+04 | 4.7E+06 |
| AT1G79210 | AT1G79210.3;AT1G79210.2;AT1G79210.1;AT1G16470.2;AT1G16470.1 | N-terminal nucleophile aminohydrolases (Ntn hydrolas | 26 kDa | 0 | 0 | 6 | 1 | 7 | | | | 3.1E+06 | 3.6E+06 | 0.0E+00 | 6.7E+06 |
| AT2G28900 | | outer plastid envelope protein 16-1 | 15 kDa | 2 | 0 | 6 | 1 | 9 | | | | | | | |
| AT2G29210 | AT2G29210.2;AT2G29210.1 | splicing factor PWI domain-containing protein | 100 kDa | 1 | 0 | 6 | 1 | 8 | | | | 5.2E+05 | 5.2E+05 | 0.0E+00 | 1.0E+06 |
| AT2G40930 | | ubiquitin-specific protease 5 | 104 kDa | 0 | 0 | 6 | 1 | 7 | | | | | | | |
| AT3G26980 | | membrane-anchored ubiquitin-fold protein 4 precurso | 13 kDa | 0 | 0 | 6 | 1 | 7 | | | | | | | |
| AT3G62130 | | Pyridoxal phosphate (PLP)-dependent transferases sup | 51 kDa | 0 | 0 | 6 | 1 | 7 | | | | | | | |
| AT4G14210 | | phytoene desaturase 3 | 63 kDa | 0 | 1 | 6 | 1 | 8 | | | | | | | |
| AT4G37090 | | unknown protein; FUNCTIONS IN: molecular_function | 22 kDa | 0 | 0 | 6 | 1 | 7 | | | | | | | |
| AT5G25270 | | Ubiquitin-like superfamily protein | 69 kDa | 0 | 0 | 6 | 1 | 7 | | | | | | | |
| AT5G58600 | | Plant protein of unknown function (DUF828) | 45 kDa | 0 | 0 | 6 | 1 | 7 | | | | | | | |
| AT1G08580 | | unknown protein; Has 39 Blast hits to 39 proteins in 1! | 14 kDa | 0 | 1 | 6 | 0 | 7 | | | | | | | |
| AT1G03270 | | CBS domain-containing protein with a domain of unkn | 55 kDa | 0 | 0 | 5 | 16 | 21 | | | | | | | |
| AT1G43130 | | like COV 2 | 29 kDa | 4 | 4 | 5 | 14 | 27 | | | | | | | |
| AT1G79710 | | Major facilitator superfamily protein | 55 kDa | 0 | 0 | 5 | 14 | 19 | | | | | | | |
| AT2G20130 | | like COV 1 | 28 kDa | 2 | 4 | 5 | 13 | 24 | | | | | | | |
| AT2G40120 | | Protein kinase superfamily protein | 65 kDa | 0 | 0 | 5 | 13 | 18 | | | | | | | |
| AT4G22670 | AT4G22670.1 | HSP70-interacting protein 1 | 47 kDa | 0 | 0 | 5 | 13 | 18 | | | | 2.1E+06 | 1.0E+06 | 0.0E+00 | 3.2E+06 |
| AT4G24120 | | YELLOW STRIPE like 1 | 74 kDa | 0 | 0 | 5 | 13 | 18 | | | | | | | |
| AT3G57560 | AT3G57560.1 | N-acetyl-l-glutamate kinase | 37 kDa | 0 | 9 | 5 | 11 | 25 | 0.12 | 0.83 | 0.47 | 1.7E+06 | 9.3E+05 | 8.6E+05 | 3.5E+06 |
| AT4G19880 | | Glutathione S-transferase family protein | 41 kDa | 0 | 0 | 5 | 11 | 16 | | | | | | | |
| AT5G16280 | | Tetratricopeptide repeat (TPR)-like superfamily protein | 144 kDa | 0 | 0 | 5 | 11 | 16 | | 0.18 | 0.18 | | | | |
| AT5G16760 | | Inositol 1,3,4-trisphosphate 5/6-kinase family protein | 36 kDa | 0 | 0 | 5 | 11 | 16 | | | | | | | |
| AT1G14850 | | nucleoporin 155 | 160 kDa | 0 | 2 | 5 | 10 | 17 | | | | | | | |
| AT1G63180 | AT1G63180.2;AT1G63180.1 | UDP-D-glucose/UDP-D-galactose 4-epimerase 3 | 39 kDa | 0 | 1 | 5 | 10 | 16 | | | | 2.9E+06 | 3.2E+06 | 0.0E+00 | 6.1E+06 |
| AT1G68400 | | leucine-rich repeat transmembrane protein kinase fami | 74 kDa | 0 | 0 | 5 | 10 | 15 | | | | | | | |
| AT3G25660 | | Amidase family protein | 57 kDa | 0 | 2 | 5 | 10 | 17 | | | | | | | |
| AT5G22070 | | Core-2/I-branching beta-1,6-N-acetylglucosaminyltran | 42 kDa | 0 | 0 | 5 | 10 | 15 | | | | | | | |
| AT1G69480 | | EXS (ERD1/XPR1/SYG1) family protein | 91 kDa | 0 | 0 | 5 | 9 | 14 | | | | | | | |
| AT3G22200 | AT3G22200.1;AT3G22200.2 | Pyridoxal phosphate (PLP)-dependent transferases sup | 56 kDa | 1 | 1 | 5 | 9 | 16 | | 0.43 | 0.43 | 4.9E+06 | 1.2E+07 | 0.0E+00 | 1.7E+07 |
| AT5G01390 | AT5G01390.4;AT5G01390.3;AT5G01390.2;AT5G01390.1 | DNAJ heat shock family protein | 37 kDa | 0 | 0 | 5 | 9 | 14 | | | | | | | |
| AT1G31817 | AT1G31817.1 | Ribosomal L18p/L5e family protein | 34 kDa | 0 | 0 | 5 | 8 | 13 | | | | 3.0E+06 | 1.9E+06 | 4.8E+06 | 9.7E+06 |
| AT2G34470 | AT2G34470.1;AT2G34470.2 | urease accessory protein G | 30 kDa | 0 | 7 | 5 | 8 | 20 | | 0.41 | 0.41 | 7.2E+06 | 5.0E+06 | 2.3E+06 | 1.4E+07 |
| AT2G36250 | | Tubulin/FtsZ family protein | 51 kDa | 0 | 0 | 5 | 8 | 13 | | | | | | | |
| AT3G03920 | AT3G03920.1 | H/ACA ribonucleoprotein complex, subunit Gar1/Naf1 | 21 kDa | 1 | 5 | 5 | 8 | 19 | 0.05 | 0.55 | 0.30 | 3.4E+07 | 4.9E+07 | 9.4E+06 | 9.3E+07 |
| AT3G16630 | | P-loop containing nucleoside triphosphate hydrolases | 89 kDa | 0 | 0 | 5 | 8 | 13 | | | | | | | |
| AT3G44330 | | INVOLVED IN: protein processing; LOCATED IN: mitoch | 62 kDa | 0 | 2 | 5 | 8 | 15 | | | | | | | |
| AT4G10790 | | UBX domain-containing protein | 53 kDa | 0 | 0 | 5 | 8 | 13 | | | | | | | |
| AT4G27870 | | Vacuolar iron transporter (VIT) family protein | 83 kDa | 0 | 0 | 5 | 8 | 13 | | | | | | | |
| AT5G12470 | AT5G12470.1 | Protein of unknown function (DUF3411) | 41 kDa | 0 | 0 | 5 | 8 | 13 | | 0.95 | 0.95 | 2.3E+06 | 2.7E+06 | 0.0E+00 | 5.0E+06 |
| AT5G13520 | AT5G13520.1 | peptidase M1 family protein | 69 kDa | 0 | 0 | 5 | 8 | 13 | | | | 0.0E+00 | 2.7E+05 | 0.0E+00 | 2.7E+05 |
| AT5G14570 | | high affinity nitrate transporter 2.7 | 53 kDa | 0 | 0 | 5 | 8 | 13 | | | | | | | |
| AT5G38990 | | Malectin/receptor-like protein kinase family protein | 98 kDa | 0 | 0 | 5 | 8 | 13 | | | | | | | |
| AT5G58300 | | Leucine-rich repeat protein kinase family protein | 72 kDa | 0 | 0 | 5 | 8 | 13 | | | | | | | |
| AT1G53240 | | Lactate/malate dehydrogenase family protein | 36 kDa | 0 | 0 | 5 | 7 | 12 | | | | | | | |

| | | | | | | | | | | | | | | | |
|---|---|---|---|---|---|---|---|---|---|---|---|---|---|---|---|
| AT1G56450 | AT1G56450.1 | 20S proteasome beta subunit G1 | 28 kDa | 0 | 0 | 5 | 7 | 12 | | | | 2.2E+06 | 1.6E+06 | 0.0E+00 | 3.8E+06 |
| AT1G70070 | | DEAD/DEAH box helicase, putative | 132 kDa | 0 | 0 | 5 | 7 | 12 | | | | | | | |
| AT1G76180 | AT1G76180.1;AT1G76180.2 | Dehydrin family protein | 21 kDa | 7 | 0 | 5 | 7 | 19 | | | | 1.7E+07 | 1.0E+07 | 0.0E+00 | 2.7E+07 |
| AT1G80500 | | SNARE-like superfamily protein | 15 kDa | 2 | 0 | 5 | 7 | 14 | | 0.39 | 0.39 | | | | |
| AT3G13200 | AT3G13200.1 | Cwf15 / Cwc15 cell cycle control family protein | 27 kDa | 0 | 0 | 5 | 7 | 12 | | | | 9.8E+06 | 1.6E+07 | 1.3E+05 | 2.5E+07 |
| AT3G46510 | | plant U-box 13 | 72 kDa | 0 | 0 | 5 | 7 | 12 | | | | | | | |
| AT4G35920 | | PLAC8 family protein | 48 kDa | 0 | 0 | 5 | 7 | 12 | | | | | | | |
| AT4G36180 | | Leucine-rich receptor-like protein kinase family protein | 124 kDa | 0 | 0 | 5 | 7 | 12 | | | | | | | |
| AT5G16880 | | Target of Myb protein 1 | 45 kDa | 0 | 3 | 5 | 7 | 15 | | | | | | | |
| AT1G29980 | | Protein of unknown function, DUF642 | 44 kDa | 0 | 0 | 5 | 6 | 11 | | | | | | | |
| AT1G54280 | | ATPase E1-E2 type family protein / haloacid dehalogena | 140 kDa | 0 | 0 | 5 | 6 | 11 | | | | | | | |
| AT2G44690 | | Arabidopsis RAC-like 9 | 23 kDa | 0 | 1 | 5 | 6 | 12 | | | | | | | |
| AT3G05200 | | RING/U-box superfamily protein | 43 kDa | 0 | 0 | 5 | 6 | 11 | | | | | | | |
| AT3G43230 | | RING/FYVE/PHD-type zinc finger family protein | 52 kDa | 0 | 0 | 5 | 6 | 11 | | | | | | | |
| AT4G27500 | AT4G27500.2;AT4G27500.1 | proton pump interactor 1 | ? | 0 | 0 | 5 | 6 | 11 | | | | | | | |
| AT5G02790 | AT5G02790.1 | Glutathione S-transferase family protein | 27 kDa | 0 | 0 | 5 | 6 | 11 | | | | | | | |
| AT5G08560 | AT5G08560.3;AT5G08560.2;AT5G08560.1 | transducin family protein / WD-40 repeat family protei | 66 kDa | 0 | 0 | 5 | 6 | 11 | | | | 1.7E+06 | 2.3E+06 | 3.8E+05 | 4.3E+06 |
| AT5G41060 | | DHHC-type zinc finger family protein | 46 kDa | 0 | 0 | 5 | 6 | 11 | | | | | | | |
| AT5G47780 | | galacturonosyltransferase 4 | 71 kDa | 0 | 0 | 5 | 6 | 11 | | | | | | | |
| AT5G59160 | AT5G59160.3;AT5G59160.2;AT5G59160.1;AT3G46820.1;AT2G29400.1 | type one serine/threonine protein phosphatase 2 | 36 kDa | 0 | 3 | 5 | 6 | 14 | | | | 2.9E+05 | 0.0E+00 | 5.0E+05 | 7.8E+05 |
| AT5G64120 | | Peroxidase superfamily protein | 35 kDa | 0 | 3 | 5 | 6 | 14 | | | | | | | |
| AT1G15710 | | prephenate dehydrogenase family protein | 41 kDa | 0 | 3 | 5 | 5 | 13 | | | | | | | |
| AT1G31930 | | extra-large GTP-binding protein 3 | 96 kDa | 0 | 0 | 5 | 5 | 10 | | | | | | | |
| AT1G49320 | | unknown seed protein like 1 | 32 kDa | 1 | 5 | 5 | 5 | 16 | | | | | | | |
| AT1G58230 | | binding | 293 kDa | 0 | 1 | 5 | 5 | 11 | | | | | | | |
| AT1G59580 | | mitogen-activated protein kinase homolog 2 | 43 kDa | 0 | 0 | 5 | 5 | 10 | | | | | | | |
| AT1G60650 | AT1G60650.4;AT1G60650.3;AT1G60650.2;AT1G60650.1 | RNA-binding (RRM/RBD/RNP motifs) family protein wit | 34 kDa | 0 | 2 | 5 | 5 | 12 | | | | 0.0E+00 | 0.0E+00 | 1.2E+05 | 1.2E+05 |
| AT1G70770 | | Protein of unknown function DUF2359, transmembrar | 67 kDa | 0 | 0 | 5 | 5 | 10 | | 0.14 | 0.14 | | | | |
| AT1G73250 | | GDP-4-keto-6-deoxymannose-3,5-epimerase-4-reducta | 36 kDa | 0 | 6 | 5 | 5 | 16 | | | | | | | |
| AT2G31510 | | IBR domain-containing protein | 64 kDa | 0 | 0 | 5 | 5 | 10 | | | | | | | |
| AT2G39090 | AT2G39090.2;AT2G39090.1 | tetratricopeptide repeat (TPR)-containing protein | 62 kDa | 0 | 0 | 5 | 5 | 10 | | | | 5.9E+05 | 6.7E+05 | 2.2E+05 | 1.5E+06 |
| AT2G41410 | | Calcium-binding EF-hand family protein | 23 kDa | 1 | 3 | 5 | 5 | 14 | | 7.22 | 7.22 | | | | |
| AT3G02630 | | Plant stearoyl-acyl-carrier-protein desaturase family pr | 45 kDa | 0 | 0 | 5 | 5 | 10 | | | | | | | |
| AT3G16640 | AT3G16640.1 | translationally controlled tumor protein | 19 kDa | 0 | 6 | 5 | 5 | 16 | | | | 2.8E+06 | 1.4E+06 | 3.8E+05 | 4.6E+06 |
| AT3G20650 | AT3G20650.3;AT3G20650.2;AT3G20650.1 | mRNA capping enzyme family protein | 42 kDa | 0 | 1 | 5 | 5 | 11 | | | | 6.3E+05 | 4.7E+05 | 6.0E+04 | 1.2E+06 |
| AT3G46440 | AT3G46440.2;AT3G46440.1 | UDP-XYL synthase 5 | 38 kDa | 1 | 6 | 5 | 5 | 17 | | | | | | | |
| AT4G02720 | AT4G02720.1 | unknown protein; FUNCTIONS IN: molecular_function | 48 kDa | 0 | 0 | 5 | 5 | 10 | | | | 9.8E+05 | 2.0E+06 | 3.0E+05 | 3.3E+06 |
| AT4G21960 | | Peroxidase superfamily protein | 37 kDa | 0 | 1 | 5 | 5 | 11 | | | | | | | |
| AT4G39150 | | DNAJ heat shock N-terminal domain-containing protei | 38 kDa | 0 | 0 | 5 | 5 | 10 | | | | | | | |
| AT5G06740 | AT5G06740.1;AT5G06740.2;AT4G01370.1;AT5G01540.1;AT3G08870.2;AT5G03140.1;AT5G01550.1;AT3G08870.1;AT5G01560.1;AT5G59270.2;AT5G59270.3;AT5G55830.1;AT5G10530.1;AT3G53810.1;AT5G65600.1;AT2G37710.1;AT5G59260.1;AT4G02410.1;AT5G59270.1;AT3G53380.1 | Concanavalin A-like lectin protein kinase family protein | 73 kDa | 0 | 0 | 5 | 5 | 10 | | | | 1.2E+06 | 9.0E+05 | 0.0E+00 | 2.1E+06 |
| AT5G09850 | | Transcription elongation factor (TFIIS) family protein | 40 kDa | 0 | 0 | 5 | 5 | 10 | | | | | | | |
| AT5G22750 | AT5G22750.1 | DNA/RNA helicase protein | 114 kDa | 0 | 0 | 5 | 5 | 10 | | | | 6.5E+04 | 9.5E+04 | 1.6E+04 | 1.8E+05 |
| AT5G57330 | | Galactose mutarotase-like superfamily protein | 35 kDa | 0 | 0 | 5 | 5 | 10 | | | | | | | |
| AT5G64290 | | dicarboxylate transport 2.1 | 60 kDa | 0 | 0 | 5 | 5 | 10 | | | | | | | |
| AT1G04040 | | HAD superfamily, subfamily IIIB acid phosphatase | 31 kDa | 0 | 0 | 5 | 4 | 9 | | | | | | | |
| AT1G34120 | | inositol polyphosphate 5-phosphatase I | 67 kDa | 0 | 0 | 5 | 4 | 9 | | | | | | | |
| AT1G34320 | | Protein of unknown function (DUF668) | 73 kDa | 0 | 0 | 5 | 4 | 9 | | | | | | | |
| AT1G35680 | | Ribosomal protein L21 | 24 kDa | 0 | 0 | 5 | 4 | 9 | | 0.34 | 0.34 | | | | |
| AT1G64450 | AT1G64450.1 | Glycine-rich protein family | 36 kDa | 0 | 0 | 5 | 4 | 9 | | | | 5.5E+06 | 4.4E+06 | 2.4E+05 | 1.0E+07 |
| AT2G06850 | AT2G06850.1;AT2G06850.2 | xyloglucan endotransglucosylase/hydrolase 4 | 34 kDa | 0 | 0 | 5 | 4 | 9 | | | | 2.6E+07 | 2.7E+07 | 2.1E+06 | 5.5E+07 |
| AT2G43610 | | Chitinase family protein | 30 kDa | 0 | 0 | 5 | 4 | 9 | | | | | | | |
| AT3G48195 | | Phox (PX) domain-containing protein | 104 kDa | 0 | 0 | 5 | 4 | 9 | | | | | | | |
| AT3G57300 | | INO80 ortholog | 172 kDa | 0 | 0 | 5 | 4 | 9 | | | | | | | |
| AT4G00340 | | receptor-like protein kinase 4 | 91 kDa | 0 | 0 | 5 | 4 | 9 | | | | | | | |
| AT4G01790 | AT4G01790.1 | Ribosomal protein L7Ae/L30e/S12e/Gadd45 family prc | 19 kDa | 0 | 0 | 5 | 4 | 9 | | | | 1.8E+06 | 7.3E+05 | 3.5E+05 | 2.8E+06 |
| AT4G14360 | | S-adenosyl-L-methionine-dependent methyltransferase | 69 kDa | 0 | 0 | 5 | 4 | 9 | | | | | | | |
| AT4G15415 | | Protein phosphatase 2A regulatory B subunit family prc | 59 kDa | 0 | 0 | 5 | 4 | 9 | | | | | | | |
| AT4G16566 | | histidine triad nucleotide-binding 4 | 17 kDa | 0 | 0 | 5 | 4 | 9 | | 0.26 | 0.26 | | | | |
| AT4G22900 | | Protein of unknown function (DUF1191) | 38 kDa | 0 | 0 | 5 | 4 | 9 | | | | | | | |
| AT5G22370 | | P-loop containing nucleoside triphosphate hydrolases | 34 kDa | 1 | 1 | 5 | 4 | 11 | | | | | | | |
| AT5G60540 | | pyridoxine biosynthesis 2 | 27 kDa | 0 | 2 | 5 | 4 | 11 | | | | | | | |
| ATCG00660 | | ribosomal protein L20 | 14 kDa | 0 | 0 | 5 | 4 | 9 | | | | | | | |
| AT1G26110 | AT1G26110.2;AT1G26110.1 | decapping 5 | 64 kDa | 0 | 0 | 5 | 3 | 8 | | | | 5.7E+06 | 5.5E+06 | 0.0E+00 | 1.1E+07 |
| AT1G27970 | | nuclear transport factor 2B | 14 kDa | 1 | 0 | 5 | 3 | 9 | | | | | | | |
| AT1G66680 | | S-adenosyl-L-methionine-dependent methyltransferase | 39 kDa | 0 | 0 | 5 | 3 | 8 | | | | | | | |
| AT1G74970 | AT1G74970.1 | ribosomal protein S9 | 23 kDa | 0 | 0 | 5 | 3 | 8 | | | | 3.0E+06 | 2.8E+06 | 0.0E+00 | 5.8E+06 |
| AT1G76700 | | DNAJ heat shock N-terminal domain-containing protei | 45 kDa | 0 | 0 | 5 | 3 | 8 | | | | | | | |
| AT2G39140 | | pseudouridine synthase family protein | 45 kDa | 0 | 0 | 5 | 3 | 8 | | | | | | | |
| AT2G41050 | | PQ-loop repeat family protein / transmembrane family | 42 kDa | 0 | 0 | 5 | 3 | 8 | | | | | | | |
| AT2G44730 | AT2G44730.1 | Alcohol dehydrogenase transcription factor Myb/SANT | 41 kDa | 0 | 0 | 5 | 3 | 8 | | | | 1.7E+05 | 3.1E+05 | 0.0E+00 | 4.8E+05 |
| AT3G04470 | | Ankyrin repeat family protein | 72 kDa | 0 | 0 | 5 | 3 | 8 | | | | | | | |
| AT3G12290 | AT3G12290.1 | Amino acid dehydrogenase family protein | 32 kDa | 0 | 0 | 5 | 3 | 8 | | | | 3.4E+06 | 1.2E+06 | 0.0E+00 | 4.6E+06 |
| AT3G14090 | AT3G14090.1 | exocyst subunit exo70 family protein D3 | 71 kDa | 0 | 1 | 5 | 3 | 9 | | | | | | | |
| AT3G56510 | AT3G56510.5;AT3G56510.4;AT3G56510.3;AT3G56510.2;AT3G56510.1 | RNA-binding (RRM/RBD/RNP motifs) family protein | 30 kDa | 0 | 0 | 5 | 3 | 8 | | | | 5.8E+05 | 8.5E+05 | 0.0E+00 | 1.4E+06 |
| AT4G27750 | | binding | 33 kDa | 3 | 1 | 5 | 3 | 12 | | | | | | | |
| AT4G32720 | AT4G32720.2;AT4G32720.1 | La protein 1 | 48 kDa | 0 | 0 | 5 | 3 | 8 | | | | | | | |
| AT5G11750 | | Ribosomal protein L19 family protein | 26 kDa | 0 | 0 | 5 | 3 | 8 | | | | | | | |
| AT5G25240 | | unknown protein; FUNCTIONS IN: molecular_function | 15 kDa | 0 | 0 | 5 | 3 | 8 | | | | | | | |
| AT5G42960 | AT5G42960.1 | unknown protein; FUNCTIONS IN: molecular_function | ? | 0 | 0 | 5 | 3 | 8 | | | | 8.5E+06 | 1.1E+07 | 3.3E+05 | 1.9E+07 |
| AT5G47020 | | unknown protein; FUNCTIONS IN: molecular_function | 153 kDa | 0 | 0 | 5 | 3 | 8 | | | | | | | |
| AT5G51290 | | Diacylglycerol kinase family protein | 68 kDa | 0 | 0 | 5 | 3 | 8 | | | | | | | |
| AT5G59880 | | actin depolymerizing factor 3 | 16 kDa | 4 | 0 | 5 | 3 | 12 | | | | | | | |
| AT5G60980 | AT5G60980.3;AT5G60980.2;AT5G60980.4;AT5G60980.1 | Nuclear transport factor 2 (NTF2) family protein with R | 49 kDa | 2 | 1 | 5 | 3 | 11 | 0.05 | 0.51 | 0.28 | 5.5E+06 | 7.2E+06 | 1.3E+06 | 1.4E+07 |

| | | | | | | | | | | | | | | | |
|---|---|---|---|---|---|---|---|---|---|---|---|---|---|---|---|
| AT1G02280 | AT1G02280.2;AT1G02280.1 | translocon at the outer envelope membrane of chlorop | 33 kDa | 0 | 2 | 5 | 2 | 9 | | 1.28 | 1.28 | 1.4E+06 | 1.3E+06 | 0.0E+00 | 2.7E+06 |
| AT1G03910 | | EXPRESSED IN: 25 plant structures; EXPRESSED DURING | ? | 0 | 0 | 5 | 2 | 7 | | | | | | | |
| AT1G08050 | | Zinc finger (C3HC4-type RING finger) family protein | 70 kDa | 0 | 0 | 5 | 2 | 7 | | | | | | | |
| AT1G10520 | | DNA polymerase lambda (POLL) | 60 kDa | 0 | 0 | 5 | 2 | 7 | | | | | | | |
| AT1G15270 | | Translation machinery associated TMA7 | 7 kDa | 0 | 0 | 5 | 2 | 7 | | | | | | | |
| AT1G16020 | | Protein of unknown function (DUF1712) | 56 kDa | 0 | 0 | 5 | 2 | 7 | | | | | | | |
| AT1G31460 | | unknown protein; FUNCTIONS IN: molecular_function | ? | 0 | 0 | 5 | 2 | 7 | | | | | | | |
| AT2G05830 | | NagB/RpiA/CoA transferase-like superfamily protein | 40 kDa | 0 | 0 | 5 | 2 | 7 | | | | | | | |
| AT3G15590 | AT3G15590.2;AT3G15590.1 | Tetratricopeptide repeat (TPR)-like superfamily protein | 69 kDa | 0 | 0 | 5 | 2 | 7 | | | | 2.6E+05 | 2.9E+05 | 3.7E+04 | 5.9E+05 |
| AT3G30300 | | O-fucosyltransferase family protein | 77 kDa | 0 | 0 | 5 | 2 | 7 | | | | | | | |
| AT4G08520 | | SNARE-like superfamily protein | 20 kDa | 0 | 0 | 5 | 2 | 7 | | | | | | | |
| AT4G12950 | | Fasciclin-like arabinogalactan family protein | 20 kDa | 0 | 0 | 5 | 2 | 7 | | | | | | | |
| AT4G25900 | | Galactose mutarotase-like superfamily protein | 36 kDa | 0 | 0 | 5 | 2 | 7 | | | | | | | |
| AT4G30430 | AT4G30430.1 | tetraspanin9 | 31 kDa | 0 | 8 | 5 | 2 | 15 | | | | 1.8E+06 | 9.6E+05 | 0.0E+00 | 2.8E+06 |
| AT4G32830 | | ataurora1 | 34 kDa | 0 | 0 | 5 | 2 | 7 | | | | | | | |
| AT4G35830 | | aconitase 1 | 98 kDa | 0 | 0 | 5 | 2 | 7 | | | | | | | |
| AT4G37980 | | elicitor-activated gene 3-1 | 38 kDa | 0 | 2 | 5 | 2 | 9 | | | | | | | |
| AT5G04870 | | calcium dependent protein kinase 1 | 68 kDa | 0 | 0 | 5 | 2 | 7 | | | | | | | |
| AT5G08200 | | peptidoglycan-binding LysM domain-containing protei | 45 kDa | 0 | 0 | 5 | 2 | 7 | | | | | | | |
| AT5G14230 | | CONTAINS InterPro DOMAIN/s: Ankyrin repeat-containi | 81 kDa | 0 | 0 | 5 | 2 | 7 | | | | | | | |
| AT5G19510 | AT5G19510.1 | Translation elongation factor EF1B/ribosomal protein | 24 kDa | 2 | 3 | 5 | 2 | 12 | | | | 4.0E+06 | 6.7E+06 | 0.0E+00 | 1.1E+07 |
| AT5G26110 | AT5G26110.3;AT5G26110.1;AT5G26110.2 | Protein kinase superfamily protein | ? | 0 | 1 | 5 | 2 | 8 | | | | 5.8E+06 | 2.0E+06 | 4.5E+05 | 8.2E+06 |
| AT5G40640 | | unknown protein; LOCATED IN: plasma membrane; EXP | ? | 0 | 0 | 5 | 2 | 7 | | | | | | | |
| AT5G45590 | | Ribosomal protein L35 | 20 kDa | 0 | 0 | 5 | 2 | 7 | | | | | | | |
| AT5G48810 | | cytochrome B5 isoform D | 15 kDa | 1 | 0 | 5 | 2 | 8 | 0.13 | | 0.13 | | | | |
| AT5G55130 | | co-factor for nitrate, reductase and xanthine dehydrog | 51 kDa | 0 | 0 | 5 | 2 | 7 | | | | | | | |
| AT5G56760 | AT5G56760.1 | serine acetyltransferase 1;1 | 33 kDa | 0 | 2 | 5 | 2 | 9 | | | | 9.5E+06 | 8.1E+06 | 3.0E+04 | 1.8E+07 |
| AT5G58450 | AT5G58450.1;AT5G58450.2 | Tetratricopeptide repeat (TPR)-like superfamily protein | 121 kDa | 0 | 0 | 5 | 2 | 7 | | | | 5.4E+06 | 3.2E+06 | 3.7E+06 | 1.2E+07 |
| AT1G15370 | AT1G15370.1 | SNARE-like superfamily protein | 17 kDa | 0 | 0 | 5 | 1 | 6 | | | | | | | |
| AT1G26550 | | FKBP-like peptidyl-prolyl cis-trans isomerase family pro | 15 kDa | 1 | 0 | 5 | 1 | 7 | | | | | | | |
| AT1G29900 | | carbamoyl phosphate synthetase B | 130 kDa | 0 | 0 | 5 | 1 | 6 | | | | | | | |
| AT1G32190 | | alpha/beta-Hydrolases superfamily protein | 46 kDa | 0 | 0 | 5 | 1 | 6 | | | | | | | |
| AT1G73940 | AT1G73940.1 | unknown protein; BEST Arabidopsis thaliana protein m | ? | 0 | 0 | 5 | 1 | 6 | | | | 7.1E+06 | 8.3E+06 | 2.3E+06 | 1.8E+07 |
| AT2G01710 | AT2G01710.1;AT2G01710.2 | Chaperone DnaJ-domain superfamily protein | 35 kDa | 0 | 0 | 5 | 1 | 6 | | | | 6.8E+05 | 6.8E+05 | 0.0E+00 | 1.4E+06 |
| AT2G19490 | AT2G19490.1 | recA DNA recombination family protein | 46 kDa | 0 | 1 | 5 | 1 | 7 | | | | 1.5E+06 | 1.5E+06 | 9.4E+04 | 3.1E+06 |
| AT2G33800 | | Ribosomal protein S5 family protein | 33 kDa | 0 | 0 | 5 | 1 | 6 | | | | | | | |
| AT2G47630 | | alpha/beta-Hydrolases superfamily protein | 40 kDa | 0 | 0 | 5 | 1 | 6 | | | | | | | |
| AT3G02190 | AT3G02190.1 | Ribosomal protein L39 family protein | 6 kDa | 0 | 0 | 5 | 1 | 6 | | | | 1.5E+08 | 1.8E+08 | 0.0E+00 | 3.3E+08 |
| AT3G02900 | | unknown protein; FUNCTIONS IN: molecular_function | 17 kDa | 1 | 0 | 5 | 1 | 7 | | 0.10 | 0.10 | | | | |
| AT4G28250 | AT4G28250.3;AT4G28250.4;AT4G28250.2;AT4G28250.1 | expansin B3 | 28 kDa | 0 | 2 | 5 | 1 | 8 | | | | 2.4E+06 | 3.0E+06 | 0.0E+00 | 5.4E+06 |
| AT4G35060 | | Heavy metal transport/detoxification superfamily prot | 17 kDa | 1 | 0 | 5 | 1 | 7 | | | | | | | |
| AT5G11270 | | overexpressor of cationic peroxidase 3 | 39 kDa | 0 | 0 | 5 | 1 | 6 | | | | | | | |
| AT5G41360 | | homolog of Xeroderma pigmentosum complementatic | 87 kDa | 0 | 0 | 5 | 1 | 6 | | | | | | | |
| AT5G41610 | | cation/H+ exchanger 18 | 80 kDa | 0 | 0 | 5 | 1 | 6 | | | | | | | |
| AT5G49900 | | Beta-glucosidase, GBA2 type family protein | 107 kDa | 0 | 0 | 5 | 1 | 6 | | | | | | | |
| AT5G52450 | | MATE efflux family protein | 52 kDa | 0 | 0 | 5 | 1 | 6 | | | | | | | |
| AT5G57000 | | unknown protein; FUNCTIONS IN: molecular_function | 21 kDa | 0 | 2 | 5 | 1 | 8 | | | | | | | |
| AT5G62460 | | RING/FYVE/PHD zinc finger superfamily protein | 34 kDa | 0 | 1 | 5 | 1 | 7 | | | | | | | |
| ATCG00330 | ATCG00330.1 | chloroplast ribosomal protein S14 | 12 kDa | 0 | 0 | 5 | 1 | 6 | | | | 8.1E+06 | 7.8E+06 | 1.4E+06 | 1.7E+07 |
| ATMG00510 | | NADH dehydrogenase subunit 7 | 45 kDa | 0 | 0 | 5 | 1 | 6 | | | | | | | |
| AT3G54230 | | suppressor of abi3-5 | 113 kDa | 0 | 1 | 5 | 0 | 6 | | | | | | | |
| AT5G40450 | AT5G40450.2;AT5G40450.1 | unknown protein; FUNCTIONS IN: molecular_function | 323 kDa | 0 | 0 | 4 | 27 | 31 | | 0.88 | 0.88 | 2.3E+06 | 1.1E+06 | 1.4E+06 | 4.8E+06 |
| AT5G13710 | | sterol methyltransferase 1 | 38 kDa | 0 | 0 | 4 | 17 | 21 | | 0.98 | 0.98 | | | | |
| AT4G27010 | AT4G27010.1;AT4G27010.2 | INVOLVED IN: biological_process unknown; LOCATED II | 270 kDa | 0 | 0 | 4 | 12 | 16 | | | | 5.2E+04 | 2.5E+04 | 4.6E+04 | 1.2E+05 |
| AT2G05755 | | Nodulin MtN21 /EamA-like transporter family protein | 44 kDa | 0 | 1 | 4 | 11 | 16 | | | | | | | |
| AT3G05100 | | S-adenosyl-L-methionine-dependent methyltransferase | 38 kDa | 0 | 8 | 4 | 11 | 23 | | | | | | | |
| AT2G37540 | | NAD(P)-binding Rossmann-fold superfamily protein | 35 kDa | 0 | 3 | 4 | 10 | 17 | | | | | | | |
| AT3G55480 | | protein affected trafﬁcking 2 | 123 kDa | 0 | 0 | 4 | 10 | 14 | | | | | | | |
| AT5G14060 | | Aspartate kinase family protein | 60 kDa | 0 | 0 | 4 | 10 | 14 | | | | | | | |
| AT5G65020 | | annexin 2 | 36 kDa | 0 | 0 | 4 | 10 | 14 | | | | | | | |
| AT1G71850 | AT1G71850.1 | Ubiquitin carboxyl-terminal hydrolase family protein | 54 kDa | 0 | 0 | 4 | 9 | 13 | | | | 3.4E+05 | 4.5E+05 | 1.9E+05 | 9.8E+05 |
| AT2G35630 | AT2G35630.1;AT2G35630.2 | ARM repeat superfamily protein | ? | 0 | 0 | 4 | 9 | 13 | | | | 1.1E+05 | 8.1E+04 | 3.6E+04 | 2.3E+05 |
| AT3G16860 | | COBRA-like protein 8 precursor | 72 kDa | 0 | 0 | 4 | 9 | 13 | | | | | | | |
| AT3G27020 | | YELLOW STRIPE like 6 | 74 kDa | 0 | 0 | 4 | 9 | 13 | | | | | | | |
| AT5G01990 | | Auxin efflux carrier family protein | 46 kDa | 0 | 0 | 4 | 9 | 13 | | | | | | | |
| AT5G15730 | AT5G15730.1;AT5G15730.3;AT5G15730.2 | Protein kinase superfamily protein | 49 kDa | 0 | 0 | 4 | 9 | 13 | | | | 2.1E+05 | 2.2E+05 | 0.0E+00 | 4.3E+05 |
| AT1G32220 | AT1G32220.1 | NAD(P)-binding Rossmann-fold superfamily protein | 32 kDa | 0 | 3 | 4 | 8 | 15 | | | | 1.1E+06 | 8.2E+05 | 0.0E+00 | 1.9E+06 |
| AT5G02240 | AT5G02240.1 | NAD(P)-binding Rossmann-fold superfamily protein | 27 kDa | 0 | 0 | 4 | 8 | 12 | | | | 3.0E+06 | 1.0E+06 | 0.0E+00 | 4.0E+06 |
| AT5G67330 | | natural resistance associated macrophage protein 4 | 56 kDa | 0 | 0 | 4 | 8 | 12 | | | | | | | |
| AT1G04430 | | S-adenosyl-L-methionine-dependent methyltransferase | 71 kDa | 0 | 0 | 4 | 7 | 11 | | | | | | | |
| AT1G09270 | | importin alpha isoform 4 | 59 kDa | 0 | 3 | 4 | 7 | 14 | | 0.33 | 0.33 | | | | |
| AT1G63830 | | PLAC8 family protein | 26 kDa | 1 | 0 | 4 | 7 | 12 | | | | | | | |
| AT1G69220 | | Protein kinase superfamily protein | 93 kDa | 0 | 0 | 4 | 7 | 11 | | | | | | | |
| AT2G25790 | | Leucine-rich receptor-like protein kinase family proteir | 106 kDa | 0 | 0 | 4 | 7 | 11 | | | | | | | |
| AT2G28120 | AT2G28120.1 | Major facilitator superfamily protein | 64 kDa | 0 | 0 | 4 | 7 | 11 | | | | | | | |
| AT2G36360 | | Galactose oxidase/kelch repeat superfamily protein | 55 kDa | 0 | 0 | 4 | 7 | 11 | | | | | | | |
| AT5G14450 | | GDSL-like Lipase/Acylhydrolase superfamily protein | 44 kDa | 0 | 0 | 4 | 7 | 11 | | | | | | | |
| AT1G47330 | | CBS domain-containing protein with a domain of unkn | 58 kDa | 0 | 0 | 4 | 6 | 10 | | | | | | | |
| AT1G68070 | | Zinc finger, C3HC4 type (RING finger) family protein | 38 kDa | 0 | 0 | 4 | 6 | 10 | | | | | | | |
| AT3G09880 | | Protein phosphatase 2A regulatory B subunit family pro | 57 kDa | 0 | 0 | 4 | 6 | 10 | | | | | | | |
| AT3G13870 | | Root hair defective 3 GTP-binding protein (RHD3) | 89 kDa | 1 | 0 | 4 | 6 | 11 | 0.23 | 0.29 | 0.26 | | | | |
| AT3G19950 | AT3G19950.1;AT3G19950.3;AT3G19950.2 | RING/U-box superfamily protein | ? | 1 | 4 | 4 | 6 | 15 | | | | 2.9E+06 | 4.4E+05 | 4.3E+04 | 3.3E+06 |
| AT3G49000 | | RNA polymerase III subunit RPC82 family protein | 59 kDa | 0 | 0 | 4 | 6 | 10 | | | | | | | |
| AT3G53380 | | Concanavalin A-like lectin protein kinase family proteir | 78 kDa | 0 | 0 | 4 | 6 | 10 | | | | | | | |
| AT3G58170 | | BET1P/SFT1P-like protein 14A | 14 kDa | 8 | 0 | 4 | 6 | 18 | | | | | | | |
| AT3G60860 | | SEC7-like guanine nucleotide exchange family protein | 200 kDa | 0 | 0 | 4 | 6 | 10 | | | | | | | |
| AT4G14880 | AT4G14880.5;AT4G14880.4;AT4G14880.3;AT4G14880.2;AT4G14880.1 | O-acetylserine (thiol) lyase (OAS-TL) isoform A1 | 34 kDa | 0 | 1 | 4 | 6 | 11 | | | | 1.0E+07 | 2.6E+06 | 0.0E+00 | 1.3E+07 |
| AT4G24990 | | Ubiquitin family protein | 13 kDa | 0 | 0 | 4 | 6 | 10 | | 2.52 | 2.52 | | | | |
| AT4G38800 | | methylthioadenosine nucleosidase 1 | 28 kDa | 0 | 2 | 4 | 6 | 12 | 0.48 | | 0.48 | | | | |
| AT5G01750 | | Protein of unknown function (DUF567) | 24 kDa | 0 | 2 | 4 | 6 | 12 | | 4.03 | 4.03 | | | | |
| AT5G18170 | | glutamate dehydrogenase 1 | 45 kDa | 0 | 0 | 4 | 6 | 10 | | | | | | | |
| AT1G29530 | | unknown protein; BEST Arabidopsis thaliana protein m | ? | 0 | 0 | 4 | 5 | 9 | | | | | | | |
| AT1G59990 | | DEA(D/H)-box RNA helicase family protein | 65 kDa | 0 | 0 | 4 | 5 | 9 | | | | | | | |
| AT1G65020 | | FUNCTIONS IN: molecular_function unknown; INVOLVI | 36 kDa | 0 | 0 | 4 | 5 | 9 | | | | | | | |
| AT1G69070 | AT1G69070.1;AT1G69070.2 | FUNCTIONS IN: molecular_function unknown; INVOLVI | 104 kDa | 0 | 0 | 4 | 5 | 9 | | | | 2.7E+05 | 7.5E+05 | 0.0E+00 | 1.0E+06 |

| | | | | | | | | | | | | | | | |
|---|---|---|---|---|---|---|---|---|---|---|---|---|---|---|---|
| AT1G74010 | AT1G74010.1 | Calcium-dependent phosphotriesterase superfamily pr | 34 kDa | 0 | 1 | 4 | 5 | 10 | | | | 6.4E+06 | 5.1E+06 | 0.0E+00 | 1.2E+07 |
| AT1G75280 | | NmrA-like negative transcriptional regulator family pro | 34 kDa | 0 | 4 | 4 | 5 | 13 | | 4.58 | 4.58 | | | | |
| AT2G21250 | | NAD(P)-linked oxidoreductase superfamily protein | 35 kDa | 0 | 1 | 4 | 5 | 10 | | | | | | | |
| AT2G22360 | | DNAJ heat shock family protein | 48 kDa | 0 | 0 | 4 | 5 | 9 | | | | | | | |
| AT2G24765 | | ADP-ribosylation factor 3 | 20 kDa | 1 | 1 | 4 | 5 | 11 | | | | | | | |
| AT2G27500 | | Glycosyl hydrolase superfamily protein | 44 kDa | 0 | 0 | 4 | 5 | 9 | | | | | | | |
| AT2G34310 | | unknown protein; FUNCTIONS IN: molecular_function | 26 kDa | 0 | 0 | 4 | 5 | 9 | | | | | | | |
| AT2G37790 | | NAD(P)-linked oxidoreductase superfamily protein | 35 kDa | 0 | 0 | 4 | 5 | 9 | | | | | | | |
| AT3G08943 | | ARM repeat superfamily protein | 96 kDa | 0 | 0 | 4 | 5 | 9 | | | | | | | |
| AT3G17590 | AT3G17590.2;AT3G17590.1;AT3G17590.3 | transcription regulatory protein SNF5, putative (BSH) | 27 kDa | 0 | 0 | 4 | 5 | 9 | | | | 0.0E+00 | 0.0E+00 | 3.2E+05 | 3.2E+05 |
| AT3G24040 | | Core-2/I-branching beta-1,6-N-acetylglucosaminyltran | 47 kDa | 0 | 0 | 4 | 5 | 9 | | | | | | | |
| AT3G52230 | AT3G52230.1 | unknown protein; FUNCTIONS IN: molecular_function | 16 kDa | 2 | 0 | 4 | 5 | 11 | | | | 2.2E+06 | 4.2E+06 | 0.0E+00 | 6.4E+06 |
| AT3G61670 | | Protein of unknown function (DUF3133) | 87 kDa | 0 | 0 | 4 | 5 | 9 | | | | | | | |
| AT3G62470 | | Pentatricopeptide repeat (PPR) superfamily protein | 69 kDa | 0 | 0 | 4 | 5 | 9 | | | | | | | |
| AT4G12070 | | unknown protein; INVOLVED IN: biological_process un | 53 kDa | 0 | 0 | 4 | 5 | 9 | | | | | | | |
| AT4G24780 | | Pectin lyase-like superfamily protein | 45 kDa | 0 | 0 | 4 | 5 | 9 | | | | | | | |
| AT5G14040 | AT5G14040.1 | phosphate transporter 3;1 | 40 kDa | 0 | 0 | 4 | 5 | 9 | 0.60 | 0.13 | 0.36 | 9.8E+05 | 9.7E+05 | 3.3E+04 | 2.0E+06 |
| AT5G22480 | AT5G22480.1 | ZPR1 zinc-finger domain protein | 54 kDa | 0 | 0 | 4 | 5 | 9 | | | | 1.5E+06 | 2.7E+06 | 0.0E+00 | 4.2E+06 |
| AT5G23300 | | pyrimidine d | 49 kDa | 0 | 0 | 4 | 5 | 9 | | | | | | | |
| AT5G56130 | | Transducin/WD40 repeat-like superfamily protein | 35 kDa | 0 | 0 | 4 | 5 | 9 | | | | | | | |
| AT1G16280 | | RNA helicase 36 | 55 kDa | 0 | 0 | 4 | 4 | 8 | | | | | | | |
| AT1G43860 | AT1G43860.2;AT1G43860.1 | sequence-specific DNA binding transcription factors | 42 kDa | 0 | 0 | 4 | 4 | 8 | | | | 6.4E+05 | 3.7E+05 | 2.6E+05 | 1.3E+06 |
| AT1G61010 | | cleavage and polyadenylation specificity factor 73-I | 77 kDa | 0 | 0 | 4 | 4 | 8 | | | | | | | |
| AT1G64385 | | unknown protein; FUNCTIONS IN: molecular_function | 38 kDa | 0 | 0 | 4 | 4 | 8 | | | | | | | |
| AT1G79280 | | nuclear pore anchor | 239 kDa | 0 | 0 | 4 | 4 | 8 | | | | | | | |
| AT2G11000 | AT2G11000.3;AT2G11000.2;AT2G11000.1;AT2G11000.4 | MAK10 homologue | 81 kDa | 0 | 0 | 4 | 4 | 8 | | | | 8.0E+05 | 6.3E+05 | 5.8E+04 | 1.5E+06 |
| AT2G19430 | | DWD (DDB1-binding WD40 protein) hypersensitive to / | 40 kDa | 0 | 0 | 4 | 4 | 8 | | | | | | | |
| AT2G19870 | AT2G19870.1 | tRNA/rRNA methyltransferase (SpoU) family protein | 66 kDa | 0 | 0 | 4 | 4 | 8 | | | | 4.7E+05 | 9.6E+05 | 4.1E+05 | 1.8E+06 |
| AT2G20020 | | RNA-binding CRS1 / YhbY (CRM) domain-containing pro | 79 kDa | 0 | 0 | 4 | 4 | 8 | | | | | | | |
| AT2G43040 | | tetratricopeptide repeat (TPR)-containing protein | 78 kDa | 0 | 0 | 4 | 4 | 8 | | | | | | | |
| AT3G02130 | | receptor-like protein kinase 2 | 125 kDa | 0 | 0 | 4 | 4 | 8 | | | | | | | |
| AT3G11900 | | aromatic and neutral transporter 1 | 46 kDa | 0 | 0 | 4 | 4 | 8 | | | | | | | |
| AT3G19870 | | unknown protein; Has 84 Blast hits to 64 proteins in 35 | 122 kDa | 0 | 0 | 4 | 4 | 8 | | | | | | | |
| AT3G45240 | | geminivirus rep interacting kinase 1 | 45 kDa | 0 | 1 | 4 | 4 | 9 | | | | | | | |
| AT3G54690 | | Sugar isomerase (SIS) family protein | 38 kDa | 0 | 0 | 4 | 4 | 8 | | | | | | | |
| AT3G56050 | | Protein kinase family protein | 55 kDa | 0 | 0 | 4 | 4 | 8 | | | | | | | |
| AT4G02290 | | glycosyl hydrolase 9B13 | 57 kDa | 0 | 1 | 4 | 4 | 9 | | | | | | | |
| AT4G16360 | | 5'-AMP-activated protein kinase beta-2 subunit protein | 29 kDa | 0 | 0 | 4 | 4 | 8 | | | | | | | |
| AT4G18530 | | Protein of unknown function (DUF707) | 45 kDa | 0 | 0 | 4 | 4 | 8 | | | | | | | |
| AT4G26300 | | Arginyl-tRNA synthetase, class Ic | 72 kDa | 0 | 0 | 4 | 4 | 8 | | | | | | | |
| AT4G31540 | | exocyst subunit exo70 family protein G1 | 78 kDa | 0 | 0 | 4 | 4 | 8 | | | | | | | |
| AT5G04060 | | S-adenosyl-L-methionine-dependent methyltransferase | 68 kDa | 0 | 0 | 4 | 4 | 8 | | | | | | | |
| AT5G11900 | AT5G11900.1 | Translation initiation factor SUI1 family protein | 22 kDa | 0 | 0 | 4 | 4 | 8 | | | | 6.3E+06 | 2.0E+06 | 3.3E+06 | 1.2E+07 |
| AT5G12260 | | BEST Arabidopsis thaliana protein match is: glycosyltra | 71 kDa | 0 | 0 | 4 | 4 | 8 | | | | | | | |
| AT5G63550 | AT5G63550.1;AT5G63550.2 | DEK domain-containing chromatin associated protein | 59 kDa | 0 | 0 | 4 | 4 | 8 | | | | 1.5E+06 | 1.1E+06 | 0.0E+00 | 2.7E+06 |
| AT1G03140 | | splicing factor Prp18 family protein | ? | 0 | 0 | 4 | 3 | 7 | | 0.56 | 0.56 | | | | |
| AT1G14510 | | alfin-like 7 | 28 kDa | 0 | 0 | 4 | 3 | 7 | | | | | | | |
| AT1G26150 | | proline-rich extensin-like receptor kinase 10 | 81 kDa | 0 | 0 | 4 | 3 | 7 | | | | | | | |
| AT1G29790 | | S-adenosyl-L-methionine-dependent methyltransferase | 42 kDa | 0 | 0 | 4 | 3 | 7 | | | | | | | |
| AT1G33990 | | methyl esterase 14 | 39 kDa | 0 | 0 | 4 | 3 | 7 | | | | | | | |
| AT1G57990 | AT1G57990.1 | purine permease 18 | 44 kDa | 0 | 0 | 4 | 3 | 7 | | | | | | | |
| AT2G01970 | | Endomembrane protein 70 protein family | 68 kDa | 0 | 0 | 4 | 3 | 7 | | | | | | | |
| AT2G18230 | AT2G18230.1 | pyrophosphorylase 2 | 25 kDa | 0 | 2 | 4 | 3 | 9 | | | | 5.9E+06 | 2.6E+06 | 0.0E+00 | 8.5E+06 |
| AT2G18470 | AT2G18470.3;AT2G18470.2;AT2G18470.1;AT2G18470.4 | roline-rich extensin-like receptor kinase 4 | 67 kDa | 0 | 0 | 4 | 3 | 7 | | | | 1.8E+06 | 1.7E+06 | 0.0E+00 | 3.5E+06 |
| AT2G35460 | | Late embryogenesis abundant (LEA) hydroxyproline-ric | 27 kDa | 0 | 0 | 4 | 3 | 7 | | | | | | | |
| AT2G41500 | AT2G41500.2;AT2G41500.1 | WD-40 repeat family protein / small nuclear ribonucle | 62 kDa | 0 | 0 | 4 | 3 | 7 | | | | 7.1E+05 | 2.1E+06 | 3.3E+05 | 3.1E+06 |
| AT3G02640 | | unknown protein; BEST Arabidopsis thaliana protein m | ? | 0 | 0 | 4 | 3 | 7 | | | | | | | |
| AT3G17210 | | heat stable protein 1 | 12 kDa | 1 | 0 | 4 | 3 | 8 | | | | | | | |
| AT3G17850 | | Protein kinase superfamily protein | 144 kDa | 0 | 0 | 4 | 3 | 7 | | | | | | | |
| AT3G22850 | | Aluminium induced protein with YGL and LRDR motifs | 27 kDa | 0 | 0 | 4 | 3 | 7 | | | | | | | |
| AT3G25070 | AT3G25070.1;AT3G25070.2 | RPM1 interacting protein 4 | 23 kDa | 1 | 3 | 4 | 3 | 11 | | | | | | | |
| AT3G51160 | | NAD(P)-binding Rossmann-fold superfamily protein | 42 kDa | 0 | 0 | 4 | 3 | 7 | | | | | | | |
| AT4G00300 | | fringe-related protein | 89 kDa | 0 | 0 | 4 | 3 | 7 | | | | | | | |
| AT4G02050 | | sugar transporter protein 7 | 56 kDa | 0 | 0 | 4 | 3 | 7 | | | | | | | |
| AT4G14290 | | alpha/beta-Hydrolases superfamily protein | 62 kDa | 0 | 0 | 4 | 3 | 7 | | | | | | | |
| AT4G23430 | | NAD(P)-binding Rossmann-fold superfamily protein | 34 kDa | 0 | 1 | 4 | 3 | 8 | | | | | | | |
| AT4G28350 | | Concanavalin A-like lectin protein kinase family proteir | 72 kDa | 0 | 0 | 4 | 3 | 7 | | | | | | | |
| AT5G22950 | | SNF7 family protein | 26 kDa | 0 | 1 | 4 | 3 | 8 | | | | | | | |
| AT5G39550 | | Zinc finger (C3HC4-type RING finger) family protein | 68 kDa | 0 | 0 | 4 | 3 | 7 | | | | | | | |
| AT5G44290 | | Protein kinase superfamily protein | 73 kDa | 0 | 0 | 4 | 3 | 7 | | | | | | | |
| AT5G48020 | | 2-oxoglutarate (2OG) and Fe(II)-dependent oxygenase s | 40 kDa | 0 | 0 | 4 | 3 | 7 | | | | | | | |
| AT5G48230 | | acetoacetyl-CoA thiolase 2 | 41 kDa | 0 | 0 | 4 | 3 | 7 | | | | | | | |
| AT5G49530 | | SIN-like family protein | 78 kDa | 0 | 0 | 4 | 3 | 7 | | | | | | | |
| AT5G51590 | | AT hook motif DNA-binding family protein | 45 kDa | 0 | 0 | 4 | 3 | 7 | | | | | | | |
| AT5G57940 | | cyclic nucleotide gated channel 5 | 82 kDa | 0 | 0 | 4 | 3 | 7 | | | | | | | |
| AT1G03280 | | Transcription factor TFIIE, alpha subunit | 54 kDa | 0 | 0 | 4 | 2 | 6 | | | | | | | |
| AT1G05520 | | Sec23/Sec24 protein transport family protein | 86 kDa | 0 | 0 | 4 | 2 | 6 | | | | | | | |
| AT1G07380 | | Neutral/alkaline non-lysosomal ceramidase | 87 kDa | 0 | 0 | 4 | 2 | 6 | | | | | | | |
| AT1G09620 | | ATP binding;leucine-tRNA ligases;aminoacyl-tRNA ligas | 123 kDa | 0 | 0 | 4 | 2 | 6 | | | | | | | |
| AT1G15740 | | Leucine-rich repeat family protein | 64 kDa | 0 | 0 | 4 | 2 | 6 | | | | | | | |
| AT1G45170 | | unknown protein; BEST Arabidopsis thaliana protein m | ? | 0 | 0 | 4 | 2 | 6 | | | | | | | |
| AT1G48570 | AT1G48570.1 | zinc finger (Ran-binding) family protein | 52 kDa | 0 | 0 | 4 | 2 | 6 | | 0.93 | 0.93 | 2.7E+05 | 1.9E+05 | 8.2E+04 | 5.4E+05 |
| AT1G68690 | | Protein kinase superfamily protein | 75 kDa | 0 | 0 | 4 | 2 | 6 | | | | | | | |
| AT1G69200 | | fructokinase-like 2 | 69 kDa | 0 | 0 | 4 | 2 | 6 | | | | | | | |
| AT1G70180 | | Sterile alpha motif (SAM) domain-containing protein | 51 kDa | 0 | 0 | 4 | 2 | 6 | | | | | | | |
| AT1G76010 | | Alba DNA/RNA-binding protein | 37 kDa | 0 | 0 | 4 | 2 | 6 | | | | | | | |
| AT2G04050 | | MATE efflux family protein | 51 kDa | 0 | 0 | 4 | 2 | 6 | | | | | | | |
| AT2G04520 | | Nucleic acid-binding, OB-fold-like protein | 17 kDa | 1 | 0 | 4 | 2 | 7 | | | | | | | |
| AT2G20560 | | DNAJ heat shock family protein | 37 kDa | 0 | 0 | 4 | 2 | 6 | | | | | | | |
| AT2G24580 | | FAD-dependent oxidoreductase family protein | 46 kDa | 0 | 0 | 4 | 2 | 6 | | | | | | | |
| AT2G29050 | | RHOMBOID-like 1 | 43 kDa | 0 | 0 | 4 | 2 | 6 | | | | | | | |
| AT2G30930 | | unknown protein; FUNCTIONS IN: molecular_function | ? | 0 | 0 | 4 | 2 | 6 | | | | | | | |
| AT2G38280 | AT2G38280.2;AT2G38280.1 | AMP deaminase, putative / myoadenylate deaminase, p | 95 kDa | 0 | 0 | 4 | 2 | 6 | | | | | | | |
| AT2G43970 | | RNA-binding protein | 61 kDa | 0 | 0 | 4 | 2 | 6 | | | | | | | |
| AT3G05000 | | Transport protein particle (TRAPP) component | 20 kDa | 0 | 1 | 4 | 2 | 7 | | | | | | | |
| AT3G06540 | | Rab escort protein | 62 kDa | 0 | 0 | 4 | 2 | 6 | | | | | | | |
| AT3G08030 | AT3G08030.2;AT3G08030.1 | Protein of unknown function, DUF642 | 39 kDa | 0 | 0 | 4 | 2 | 6 | | | | 1.6E+07 | 1.8E+07 | 1.2E+06 | 3.5E+07 |
| AT3G13350 | | HMG (high mobility group) box protein with ARID/BRIC | 36 kDa | 0 | 0 | 4 | 2 | 6 | | | | | | | |

| | | | | | | | | | | | | | | | |
|---|---|---|---|---|---|---|---|---|---|---|---|---|---|---|---|
| AT3G17740 | AT3G17740.1;AT3G17712.3;AT3G17712.1;AT3G17712.2 | unknown protein; FUNCTIONS IN: molecular_function | ? | 0 | 0 | 4 | 2 | 6 | | | | 7.1E+05 | 2.9E+05 | 0.0E+00 | 1.0E+06 |
| AT3G46220 | | unknown protein; CONTAINS InterPro DOMAIN/s: Prot | 89 kDa | 0 | 0 | 4 | 2 | 6 | | | | | | | |
| AT3G50520 | | Phosphoglycerate mutase family protein | 26 kDa | 0 | 0 | 4 | 2 | 6 | | | | | | | |
| AT3G53190 | | Pectin lyase-like superfamily protein | 54 kDa | 0 | 0 | 4 | 2 | 6 | | | | | | | |
| AT3G59700 | | lectin-receptor kinase | 74 kDa | 0 | 0 | 4 | 2 | 6 | | | | | | | |
| AT3G62240 | | RING/U-box superfamily protein | 90 kDa | 0 | 0 | 4 | 2 | 6 | | | | | | | |
| AT4G01610 | | Cysteine proteinases superfamily protein | 39 kDa | 0 | 0 | 4 | 2 | 6 | | | | | | | |
| AT4G01660 | | ABC transporter 1 | 69 kDa | 0 | 1 | 4 | 2 | 7 | | | | | | | |
| AT4G23530 | | Protein of unknown function (DUF793) | 44 kDa | 0 | 0 | 4 | 2 | 6 | | | | | | | |
| AT4G25880 | | pumilio 6 | 96 kDa | 0 | 0 | 4 | 2 | 6 | | | | | | | |
| AT4G32940 | | gamma vacuolar processing enzyme | 54 kDa | 0 | 0 | 4 | 2 | 6 | | | | | | | |
| AT4G33350 | | Tic22-like family protein | 27 kDa | 0 | 0 | 4 | 2 | 6 | | | | | | | |
| AT4G35785 | | RNA-binding (RRM/RBD/RNP motifs) family protein | 24 kDa | 0 | 1 | 4 | 2 | 7 | | 0.80 | 0.80 | | | | |
| AT5G10070 | AT5G10070.1;AT5G10070.2 | RNase L inhibitor protein-related | 29 kDa | 0 | 0 | 4 | 2 | 6 | | | | 1.5E+06 | 2.4E+06 | 0.0E+00 | 3.9E+06 |
| AT5G26210 | | alfin-like 4 | 29 kDa | 0 | 3 | 4 | 2 | 9 | | | | | | | |
| AT5G30510 | | ribosomal protein S1 | 45 kDa | 0 | 1 | 4 | 2 | 7 | | | | | | | |
| AT5G36880 | AT5G36880.4;AT5G36880.6;AT5G36880.5;AT5G36880.3;AT5G36880.1;AT5G36880.2 | acetyl-CoA synthetase | 82 kDa | 0 | 0 | 4 | 2 | 6 | | | | 1.2E+05 | 5.2E+04 | 0.0E+00 | 1.7E+05 |
| AT5G43430 | | electron transfer flavoprotein beta | 27 kDa | 0 | 0 | 4 | 2 | 6 | | | | | | | |
| AT5G61970 | | signal recognition particle-related / SRP-related | 69 kDa | 0 | 0 | 4 | 2 | 6 | | | | | | | |
| AT5G64350 | AT5G64350.1 | FK506-binding protein 12 | 12 kDa | 1 | 0 | 4 | 2 | 7 | | | | 2.8E+07 | 1.7E+07 | 0.0E+00 | 4.5E+07 |
| AT1G03400 | | 2-oxoglutarate (2OG) and Fe(II)-dependent oxygenase s | 39 kDa | 0 | 2 | 4 | 1 | 7 | | | | | | | |
| AT1G06190 | | Rho termination factor | 45 kDa | 0 | 0 | 4 | 1 | 5 | | | | | | | |
| AT1G06650 | | 2-oxoglutarate (2OG) and Fe(II)-dependent oxygenase s | 42 kDa | 0 | 0 | 4 | 1 | 5 | | | | | | | |
| AT1G09060 | | Zinc finger, RING-type;Transcription factor jumonji/asp | 107 kDa | 0 | 0 | 4 | 1 | 5 | | | | | | | |
| AT1G15940 | | Tudor/PWWP/MBT superfamily protein | ? | 0 | 0 | 4 | 1 | 5 | | | | | | | |
| AT1G24150 | | formin homologue 4 | 80 kDa | 0 | 0 | 4 | 1 | 5 | | | | | | | |
| AT1G33970 | | P-loop containing nucleoside triphosphate hydrolases | 39 kDa | 0 | 0 | 4 | 1 | 5 | | | | | | | |
| AT1G54250 | | RNA polymerase Rpb8 | 17 kDa | 0 | 0 | 4 | 1 | 5 | | | | | | | |
| AT1G61570 | AT1G61570.1 | translocase of the inner mitochondrial membrane 13 | 9 kDa | 0 | 0 | 4 | 1 | 5 | | 0.18 | 0.18 | 2.3E+06 | 4.2E+06 | 3.9E+06 | 1.0E+07 |
| AT1G66240 | | homolog of anti-oxidant 1 | 11 kDa | 0 | 0 | 4 | 1 | 5 | | | | | | | |
| AT1G68300 | | Adenine nucleotide alpha hydrolases-like superfamily p | 17 kDa | 0 | 0 | 4 | 1 | 5 | | | | | | | |
| AT1G71040 | | Cupredoxin superfamily protein | 66 kDa | 0 | 1 | 4 | 1 | 6 | | | | | | | |
| AT1G75670 | AT1G75670.1;AT1G75670.2;AT1G75670.3 | DNA-directed RNA polymerases | 22 kDa | 0 | 0 | 4 | 1 | 5 | | | | 9.2E+05 | 2.4E+06 | 1.3E+06 | 4.7E+06 |
| AT1G77710 | | CONTAINS InterPro DOMAIN/s: Ubiquitin-like, Ufm1 (In | 10 kDa | 0 | 0 | 4 | 1 | 5 | | | | | | | |
| AT1G80950 | | Phospholipid/glycerol acyltransferase family protein | 45 kDa | 0 | 0 | 4 | 1 | 5 | | | | | | | |
| AT2G16640 | AT2G16640.3;AT2G16640.2;AT2G16640.1 | multimeric translocon complex in the outer envelope r | 132 kDa | 0 | 0 | 4 | 1 | 5 | | 0.10 | 0.10 | 1.6E+06 | 2.0E+06 | 6.3E+04 | 3.6E+06 |
| AT2G34250 | | SecY protein transport family protein | 52 kDa | 0 | 0 | 4 | 1 | 5 | | | | | | | |
| AT2G40430 | AT2G40430.3;AT2G40430.1;AT2G40430.2 | CONTAINS InterPro DOMAIN/s: P60-like (InterPro:IPR0 | 50 kDa | 0 | 0 | 4 | 1 | 5 | | | | 1.9E+06 | 2.3E+06 | 8.9E+05 | 5.1E+06 |
| AT2G46520 | | cellular apoptosis susceptibility protein, putative / imp | 109 kDa | 0 | 0 | 4 | 1 | 5 | | | | | | | |
| AT3G06860 | AT3G06860.1 | multifunctional protein 2 | 79 kDa | 0 | 0 | 4 | 1 | 5 | | | | 2.0E+06 | 4.7E+06 | 0.0E+00 | 6.7E+06 |
| AT3G08640 | | Protein of unknown function (DUF3411) | 35 kDa | 0 | 0 | 4 | 1 | 5 | | | | | | | |
| AT3G10550 | | Myotubularin-like phosphatases II superfamily | 93 kDa | 0 | 0 | 4 | 1 | 5 | | | | | | | |
| AT3G11420 | | Protein of unknown function (DUF604) | 58 kDa | 0 | 0 | 4 | 1 | 5 | | | | | | | |
| AT3G12400 | | Ubiquitin-conjugating enzyme/RWD-like protein | 45 kDa | 0 | 0 | 4 | 1 | 5 | | | | | | | |
| AT3G22370 | | alternative oxidase 1A | 40 kDa | 10 | 0 | 4 | 1 | 15 | 0.11 | 0.12 | 0.11 | | | | |
| AT3G24180 | | Beta-glucosidase, GBA2 type family protein | 106 kDa | 0 | 0 | 4 | 1 | 5 | | | | | | | |
| AT3G42790 | AT3G42790.1 | alfin-like 3 | 28 kDa | 0 | 0 | 4 | 1 | 5 | | | | 5.4E+06 | 2.7E+06 | 4.7E+05 | 8.6E+06 |
| AT3G54850 | | plant U-box 14 | 69 kDa | 0 | 0 | 4 | 1 | 5 | | | | | | | |
| AT4G09730 | AT4G09730.1 | RH39 | 69 kDa | 0 | 0 | 4 | 1 | 5 | | | | 8.4E+05 | 8.7E+05 | 2.1E+05 | 1.9E+06 |
| AT4G15850 | | RNA helicase 1 | 58 kDa | 0 | 0 | 4 | 1 | 5 | | | | | | | |
| AT4G16180 | | unknown protein; FUNCTIONS IN: molecular_function | 92 kDa | 0 | 0 | 4 | 1 | 5 | | | | | | | |
| AT4G22320 | | unknown protein; FUNCTIONS IN: molecular_function | 26 kDa | 0 | 0 | 4 | 1 | 5 | | | | | | | |
| AT4G22740 | | glycine-rich protein | 39 kDa | 0 | 0 | 4 | 1 | 5 | | | | | | | |
| AT4G24760 | | alpha/beta-Hydrolases superfamily protein | 42 kDa | 0 | 0 | 4 | 1 | 5 | | | | | | | |
| AT4G31120 | | SHK1 binding protein 1 | 72 kDa | 0 | 0 | 4 | 1 | 5 | | | | | | | |
| AT4G32660 | | Protein kinase superfamily protein | 46 kDa | 0 | 0 | 4 | 1 | 5 | | | | | | | |
| AT4G34412 | AT4G34412.1 | CONTAINS InterPro DOMAIN/s: Kinase binding protein ( | 19 kDa | 0 | 0 | 4 | 1 | 5 | | | | 6.1E+06 | 3.4E+06 | 0.0E+00 | 9.5E+06 |
| AT5G03880 | AT5G03880.1 | Thioredoxin family protein | 37 kDa | 0 | 1 | 4 | 1 | 6 | | | | 2.9E+05 | 3.1E+05 | 0.0E+00 | 5.9E+05 |
| AT5G23140 | AT5G23140.1 | nuclear-encoded CLP protease P7 | 26 kDa | 1 | 6 | 4 | 1 | 12 | | | | 1.6E+07 | 1.7E+07 | 2.5E+06 | 3.5E+07 |
| AT5G38560 | | Protein kinase superfamily protein | 72 kDa | 0 | 0 | 4 | 1 | 5 | | | | | | | |
| AT5G45160 | | Root hair defective 3 GTP-binding protein (RHD3) | 94 kDa | 0 | 0 | 4 | 1 | 5 | | | | | | | |
| AT5G52340 | | exocyst subunit exo70 family protein A2 | 72 kDa | 0 | 1 | 4 | 1 | 6 | | | | | | | |
| AT5G62620 | | Galactosyltransferase family protein | 78 kDa | 0 | 0 | 4 | 1 | 5 | | | | | | | |
| AT1G14340 | | RNA-binding (RRM/RBD/RNP motifs) family protein | 26 kDa | 0 | 2 | 4 | 0 | 6 | | | | | | | |
| AT3G45900 | | Ribonuclease P protein subunit P38-related | 45 kDa | 0 | 1 | 4 | 0 | 5 | | | | | | | |
| AT3G62530 | AT3G62530.1 | ARM repeat superfamily protein | ? | 1 | 0 | 4 | 0 | 5 | | | | 3.4E+06 | 4.8E+06 | 5.9E+05 | 8.8E+06 |
| AT1G77260 | | S-adenosyl-L-methionine-dependent methyltransferase | 74 kDa | 0 | 0 | 3 | 15 | 18 | | | | | | | |
| AT4G01690 | | Flavin containing amine oxidoreductase family | 58 kDa | 0 | 0 | 3 | 11 | 14 | | | | | | | |
| AT1G19920 | | Pseudouridine synthase/archaeosine transglycosylase-l | 54 kDa | 0 | 1 | 3 | 10 | 14 | | | | | | | |
| AT5G08550 | AT5G08550.1 | GC-rich sequence DNA-binding factor-like protein | 101 kDa | 0 | 0 | 3 | 10 | 13 | | | | 1.7E+05 | 1.4E+05 | 0.0E+00 | 3.1E+05 |
| AT5G25265 | | unknown protein; FUNCTIONS IN: molecular_function | ? | 0 | 2 | 3 | 10 | 15 | | | | | | | |
| AT1G11840 | AT1G11840.5;AT1G11840.4;AT1G11840.3;AT1G11840.2;AT1G11840.1;AT1G11840.6 | glyoxalase I homolog | 36 kDa | 1 | 0 | 3 | 9 | 13 | | | | | | | |
| AT3G06580 | | Mevalonate/galactokinase family protein | 54 kDa | 0 | 1 | 3 | 9 | 13 | | | | | | | |
| AT4G31180 | | Class II aminoacyl-tRNA and biotin synthetases superfa | 63 kDa | 0 | 0 | 3 | 9 | 12 | | | | | | | |
| AT5G47110 | AT5G47110.1 | Chlorophyll A-B binding family protein | 29 kDa | 0 | 0 | 3 | 9 | 12 | | | | 1.2E+06 | 1.3E+06 | 0.0E+00 | 2.5E+06 |
| AT1G02840 | AT1G02840.5;AT1G02840.4;AT1G02840.3;AT1G02840.2;AT1G02840.1 | RNA-binding (RRM/RBD/RNP motifs) family protein | 34 kDa | 0 | 0 | 3 | 8 | 11 | | | | 3.6E+06 | 4.7E+06 | 0.0E+00 | 8.3E+06 |
| AT1G63160 | | replication factor C 2 | 37 kDa | 0 | 0 | 3 | 8 | 11 | | | | | | | |
| AT1G78610 | AT1G78610.1 | mechanosensitive channel of small conductance-like 6 | 97 kDa | 0 | 0 | 3 | 8 | 11 | | | | 1.1E+06 | 6.7E+05 | 0.0E+00 | 1.8E+06 |
| AT3G21250 | AT3G21250.6;AT3G21250.3;AT3G21250.1;AT3G21250.5;AT3G21250.2;AT3G21250.4 | multidrug resistance-associated protein 6 | 161 kDa | 0 | 0 | 3 | 8 | 11 | | | | 1.4E+05 | 5.2E+04 | 0.0E+00 | 1.9E+05 |
| AT5G05200 | | Protein kinase superfamily protein | 60 kDa | 0 | 0 | 3 | 8 | 11 | | | | | | | |
| AT1G07320 | AT1G07320.4;AT1G07320.3;AT1G07320.2;AT1G07320.1 | ribosomal protein L4 | 31 kDa | 0 | 1 | 3 | 7 | 11 | | 1.60 | 1.60 | 9.9E+05 | 0.0E+00 | 1.2E+06 | 2.2E+06 |
| AT2G02400 | | NAD(P)-binding Rossmann-fold superfamily protein | 35 kDa | 0 | 0 | 3 | 7 | 10 | | | | | | | |
| AT2G42650 | | Ribosomal protein L1p/L10e family | 42 kDa | 0 | 0 | 3 | 7 | 10 | | | | | | | |
| AT2G44020 | AT2G44020.1 | Mitochondrial transcription termination factor family | 58 kDa | 0 | 0 | 3 | 7 | 10 | | | | | | | |
| AT3G09770 | | RING/U-box superfamily protein | 43 kDa | 0 | 0 | 3 | 7 | 10 | | | | | | | |
| AT3G48090 | | alpha/beta-Hydrolases superfamily protein | 72 kDa | 0 | 0 | 3 | 7 | 10 | | | | | | | |
| AT4G31750 | | HOPW1-1-interacting 2 | 33 kDa | 0 | 0 | 3 | 7 | 10 | | | | | | | |

| | | | | | | | | | | | | | | | |
|---|---|---|---|---|---|---|---|---|---|---|---|---|---|---|---|
| AT5G16620 | | hydroxyproline-rich glycoprotein family protein | 49 kDa | 0 | 1 | 3 | 7 | 11 | | | | | | | |
| AT5G60550 | | geminivirus rep interacting kinase 2 | 46 kDa | 0 | 1 | 3 | 7 | 11 | | | | | | | |
| AT1G16180 | AT1G16180.2;AT1G16180.1 | Serinc-domain containing serine and sphingolipid bios | 46 kDa | 0 | 0 | 3 | 6 | 9 | | | | 1.1E+06 | 7.5E+05 | 0.0E+00 | 1.8E+06 |
| AT1G34220 | | Regulator of Vps4 activity in the MVB pathway protein | 69 kDa | 0 | 0 | 3 | 6 | 9 | | | | | | | |
| AT1G59830 | AT1G59830.2;AT1G59830.1;AT1G10430.2;AT1G10430.1 | protein phosphatase 2A-2 | 35 kDa | 0 | 0 | 3 | 6 | 9 | | | | 6.3E+05 | 2.9E+05 | 0.0E+00 | 9.2E+05 |
| AT1G61150 | AT1G61150.5;AT1G61150.9;AT1G61150.8;AT1G61150.6;AT1G61150.4;AT1G61150.2;AT1G61150.11;AT1G61150.10;AT1G61150.1;AT1G61150.7;AT1G61150.3 | LisH and RanBPM domains containing protein | ? | 0 | 1 | 3 | 6 | 10 | | 2.02 | 2.02 | 1.8E+07 | 2.3E+07 | 1.7E+06 | 4.3E+07 |
| AT2G46170 | | Reticulon family protein | 29 kDa | 3 | 0 | 3 | 6 | 12 | | | | | | | |
| AT3G43300 | AT3G43300.1;AT3G43300.3;AT3G43300.2 | HOPM interactor 7 | 191 kDa | 4 | 5 | 3 | 6 | 18 | | | | 1.6E+06 | 7.8E+05 | 0.0E+00 | 2.4E+06 |
| AT3G51460 | | Phosphoinositide phosphatase family protein | 68 kDa | 0 | 0 | 3 | 6 | 9 | | | | | | | |
| AT4G13430 | AT4G13430.1 | isopropyl malate isomerase large subunit 1 | 55 kDa | 0 | 0 | 3 | 6 | 9 | | | | 4.4E+06 | 6.4E+06 | 4.7E+04 | 1.1E+07 |
| AT4G24630 | | DHHC-type zinc finger family protein | 47 kDa | 0 | 0 | 3 | 6 | 9 | | | | | | | |
| AT4G31990 | | aspartate aminotransferase 5 | 50 kDa | 0 | 0 | 3 | 6 | 9 | | | | | | | |
| AT4G32400 | AT4G32400.1 | Mitochondrial substrate carrier family protein | 43 kDa | 0 | 0 | 3 | 6 | 9 | | | | 4.6E+05 | 6.9E+05 | 3.6E+05 | 1.5E+06 |
| AT5G21326 | | Ca2+regulated serine-threonine protein kinase family p | 50 kDa | 0 | 1 | 3 | 6 | 10 | | | | | | | |
| AT1G09070 | | soybean gene regulated by cold-2 | 34 kDa | 0 | 0 | 3 | 5 | 8 | | | | | | | |
| AT1G18070 | AT1G18070.1;AT1G18070.2;AT1G18070.3 | Translation elongation factor EF1A/initiation factor IF2 | 59 kDa | 0 | 0 | 3 | 5 | 8 | | | | 2.0E+05 | 3.4E+05 | 0.0E+00 | 5.4E+05 |
| AT1G19130 | | CONTAINS InterPro DOMAIN/s: Protein of unknown fun | 22 kDa | 0 | 0 | 3 | 5 | 8 | | | | | | | |
| AT1G80830 | | natural resistance-associated macrophage protein 1 | 58 kDa | 0 | 0 | 3 | 5 | 8 | | | | | | | |
| AT2G16780 | AT2G16780.1 | Transducin family protein / WD-40 repeat family protei | 47 kDa | 0 | 0 | 3 | 5 | 8 | | | | 3.1E+06 | 5.4E+06 | 7.2E+04 | 8.6E+06 |
| AT2G19080 | | metaxin-related | 36 kDa | 0 | 0 | 3 | 5 | 8 | | | | | | | |
| AT2G29570 | | proliferating cell nuclear antigen 2 | 29 kDa | 0 | 0 | 3 | 5 | 8 | | | | | | | |
| AT2G39130 | | Transmembrane amino acid transporter family protein | 60 kDa | 0 | 0 | 3 | 5 | 8 | | | | | | | |
| AT2G41520 | | Heat shock protein DnaJ with tetratricopeptide repeat | 123 kDa | 0 | 0 | 3 | 5 | 8 | | | | | | | |
| AT3G12050 | | Aha1 domain-containing protein | 40 kDa | 0 | 0 | 3 | 5 | 8 | | | | | | | |
| AT3G18860 | | transducin family protein / WD-40 repeat family protei | 84 kDa | 0 | 0 | 3 | 5 | 8 | | | | | | | |
| AT3G26670 | | Protein of unknown function (DUF803) | 48 kDa | 0 | 0 | 3 | 5 | 8 | | | | | | | |
| AT4G01310 | | Ribosomal L5P family protein | 28 kDa | 0 | 0 | 3 | 5 | 8 | | | | | | | |
| AT4G28490 | | Leucine-rich receptor-like protein kinase family proteir | 109 kDa | 0 | 1 | 3 | 5 | 9 | | | | | | | |
| AT5G02740 | AT5G02740.1 | Ribosomal protein S24e family protein | 25 kDa | 0 | 7 | 3 | 5 | 15 | | | | 1.3E+06 | 2.5E+06 | 0.0E+00 | 3.7E+06 |
| AT5G54870 | | unknown protein; FUNCTIONS IN: molecular_function | ? | 0 | 0 | 3 | 5 | 8 | | | | | | | |
| AT5G56680 | | Class II aminoacyl-tRNA and biotin synthetases superfar | 64 kDa | 0 | 0 | 3 | 5 | 8 | | | | | | | |
| AT1G05850 | | Chitinase family protein | 36 kDa | 0 | 0 | 3 | 4 | 7 | | | | | | | |
| AT1G10510 | | RNI-like superfamily protein | 65 kDa | 0 | 0 | 3 | 4 | 7 | | | | | | | |
| AT1G16220 | | Protein phosphatase 2C family protein | 54 kDa | 0 | 0 | 3 | 4 | 7 | | | | | | | |
| AT1G16610 | AT1G16610.2;AT1G16610.1;AT1G16610.3;AT1G16610.4;AT1G16610.5;AT1G16610.7;AT1G16610.6 | arginine/serine-rich 45 | 45 kDa | 0 | 0 | 3 | 4 | 7 | | | | 3.0E+06 | 3.0E+06 | 2.3E+05 | 6.2E+06 |
| AT1G31770 | | ATP-binding cassette 14 | 73 kDa | 0 | 0 | 3 | 4 | 7 | | | | | | | |
| AT1G31800 | | cytochrome P450, family 97, subfamily A, polypeptide | 67 kDa | 0 | 0 | 3 | 4 | 7 | | | | | | | |
| AT1G43670 | | Inositol monophosphatase family protein | 37 kDa | 0 | 0 | 3 | 4 | 7 | | | | | | | |
| AT1G66260 | AT1G66260.2;AT1G66260.1 | RNA-binding (RRM/RBD/RNP motifs) family protein | 31 kDa | 0 | 0 | 3 | 4 | 7 | | | | 8.3E+06 | 1.3E+07 | 2.2E+06 | 2.4E+07 |
| AT2G20550 | | HSP40/DnaJ peptide-binding protein | 31 kDa | 0 | 0 | 3 | 4 | 7 | | | | | | | |
| AT2G20790 | | clathrin adaptor complexes medium subunit family prc | 67 kDa | 0 | 0 | 3 | 4 | 7 | | | | | | | |
| AT2G22250 | | aspartate aminotransferase | 51 kDa | 0 | 0 | 3 | 4 | 7 | | | | | | | |
| AT2G23740 | | nucleic acid binding;sequence-specific DNA binding tra | 156 kDa | 0 | 0 | 3 | 4 | 7 | | | | | | | |
| AT2G26770 | | plectin-related | 55 kDa | 0 | 0 | 3 | 4 | 7 | | | | | | | |
| AT2G27860 | | UDP-D-apiose/UDP-D-xylose synthase 1 | 44 kDa | 0 | 1 | 3 | 4 | 8 | | | | | | | |
| AT2G36200 | AT2G36200.1;AT2G36200.2 | P-loop containing nucleoside triphosphate hydrolases | 117 kDa | 0 | 0 | 3 | 4 | 7 | | | | 2.2E+05 | 0.0E+00 | 3.6E+05 | 5.8E+05 |
| AT2G40815 | | Calcium-dependent lipid-binding (CaLB domain) family | 40 kDa | 0 | 0 | 3 | 4 | 7 | | | | | | | |
| AT2G46020 | AT2G46020.6;AT2G46020.5;AT2G46020.1;AT2G46020.4;AT2G46020.3;AT2G46020.2 | transcription regulatory protein SNF2, putative | 245 kDa | 0 | 0 | 3 | 4 | 7 | | | | | | | |
| AT2G47580 | AT2G47580.1 | spliceosomal protein U1A | 28 kDa | 0 | 2 | 3 | 4 | 9 | | | | | | | |
| AT3G49120 | AT3G49120.1 | peroxidase CB | 39 kDa | 0 | 0 | 3 | 4 | 7 | | | | 7.0E+05 | 7.5E+05 | 0.0E+00 | 1.5E+06 |
| AT3G54080 | | Concanavalin A-like lectin family protein | 40 kDa | 0 | 0 | 3 | 4 | 7 | | | | | | | |
| AT3G56460 | | GroES-like zinc-binding alcohol dehydrogenase family p | 37 kDa | 0 | 0 | 3 | 4 | 7 | | | | | | | |
| AT4G00620 | AT4G00620.1 | Amino acid dehydrogenase family protein | 39 kDa | 0 | 0 | 3 | 4 | 7 | | | | 2.3E+06 | 2.3E+06 | 0.0E+00 | 4.6E+06 |
| AT4G18700 | | CBL-interacting protein kinase 12 | 55 kDa | 0 | 0 | 3 | 4 | 7 | | | | | | | |
| AT5G13740 | | zinc induced facilitator 1 | 53 kDa | 0 | 0 | 3 | 4 | 7 | | | | | | | |
| AT5G15460 | | membrane-anchored ubiquitin-fold protein 2 | 14 kDa | 1 | 0 | 3 | 4 | 8 | | 11.25 | 11.25 | | | | |
| AT5G37630 | | ARM repeat superfamily protein | 117 kDa | 0 | 0 | 3 | 4 | 7 | | | | | | | |
| AT5G38640 | | NagB/RpiA/CoA transferase-like superfamily protein | 69 kDa | 0 | 0 | 3 | 4 | 7 | | | | | | | |
| AT5G40270 | AT5G40270.1;AT5G40290.1 | HD domain-containing metal-dependent phosphohydr | 55 kDa | 0 | 0 | 3 | 4 | 7 | | | | 3.5E+05 | 7.5E+04 | 2.2E+05 | 6.5E+05 |
| AT5G47080 | | casein kinase II beta chain 1 | 32 kDa | 0 | 0 | 3 | 4 | 7 | | | | | | | |
| AT5G53370 | | pectin methylesterase PCR fragment F | 64 kDa | 0 | 0 | 3 | 4 | 7 | | | | | | | |
| AT5G55540 | | tornado 1 | 157 kDa | 0 | 0 | 3 | 4 | 7 | | | | | | | |
| AT5G63840 | | Glycosyl hydrolases family 31 protein | 104 kDa | 0 | 0 | 3 | 4 | 7 | | | | | | | |
| AT1G04190 | AT1G04190.1 | Tetratricopeptide repeat (TPR)-like superfamily protein | 37 kDa | 0 | 0 | 3 | 3 | 6 | | | | 1.0E+06 | 4.4E+05 | 5.2E+05 | 2.0E+06 |
| AT1G04200 | | FUNCTIONS IN: molecular_function unknown; INVOLVE | 83 kDa | 0 | 0 | 3 | 3 | 6 | | | | | | | |
| AT1G08200 | | UDP-D-apiose/UDP-D-xylose synthase 2 | 44 kDa | 0 | 0 | 3 | 3 | 6 | | | | | | | |
| AT1G08210 | | Eukaryotic aspartyl protease family protein | 53 kDa | 0 | 0 | 3 | 3 | 6 | | | | | | | |
| AT1G09870 | AT1G09870.1 | histidine acid phosphatase family protein | 55 kDa | 0 | 0 | 3 | 3 | 6 | | | | 3.4E+05 | 4.7E+05 | 0.0E+00 | 8.1E+05 |
| AT1G11680 | | CYTOCHROME P450 51G1 | 55 kDa | 0 | 0 | 3 | 3 | 6 | | | | | | | |
| AT1G13930 | AT1G13930.3;AT1G13930.2;AT1G13930.1 | Involved in response to salt stress. Knockout mutants a | 16 kDa | 0 | 4 | 3 | 3 | 10 | | | | 7.0E+06 | 2.0E+06 | 0.0E+00 | 9.0E+06 |
| AT1G14230 | | GDA1/CD39 nucleoside phosphatase family protein | 55 kDa | 0 | 1 | 3 | 3 | 7 | | | | | | | |
| AT1G16340 | | Aldolase superfamily protein | 32 kDa | 0 | 0 | 3 | 3 | 6 | | | | | | | |
| AT1G18670 | | Protein kinase superfamily protein | 80 kDa | 0 | 0 | 3 | 3 | 6 | | | | | | | |
| AT1G27650 | AT1G27650.1;AT1G27650.2 | U2 snRNP auxiliary factor small subunit, putative | 29 kDa | 0 | 2 | 3 | 3 | 8 | | | | 1.4E+06 | 8.7E+05 | 7.5E+05 | 3.0E+06 |
| AT1G49740 | | PLC-like phosphodiesterases superfamily protein | 40 kDa | 0 | 0 | 3 | 3 | 6 | | | | | | | |
| AT1G54080 | | oligouridylate-binding protein 1A | 47 kDa | 0 | 0 | 3 | 3 | 6 | | | | | | | |
| AT1G60940 | | SNF1-related protein kinase 2.10 | 41 kDa | 0 | 0 | 3 | 3 | 6 | | | | | | | |
| AT1G64640 | | early nodulin-like protein 8 | 20 kDa | 0 | 0 | 3 | 3 | 6 | | | | | | | |
| AT1G73100 | | SU(VAR)3-9 homolog 3 | 73 kDa | 0 | 0 | 3 | 3 | 6 | | | | | | | |
| AT2G02790 | AT2G02790.3;AT2G02790.2;AT2G02790.1 | IQ-domain 29 | 70 kDa | 0 | 0 | 3 | 3 | 6 | | | | 4.3E+05 | 7.0E+05 | 2.3E+04 | 1.2E+06 |
| AT2G13370 | | chromatin remodeling 5 | 197 kDa | 0 | 0 | 3 | 3 | 6 | | | | | | | |
| AT2G18170 | | MAP kinase 7 | 42 kDa | 0 | 0 | 3 | 3 | 6 | | | | | | | |
| AT2G27880 | | Argonaute family protein | 111 kDa | 0 | 0 | 3 | 3 | 6 | | | | | | | |
| AT2G29440 | AT2G29440.1 | glutathione S-transferase tau 6 | 26 kDa | 0 | 0 | 3 | 3 | 6 | | | | 4.2E+06 | 1.5E+06 | 0.0E+00 | 5.7E+06 |
| AT2G32520 | | alpha/beta-Hydrolases superfamily protein | 26 kDa | 0 | 0 | 3 | 3 | 6 | | | | | | | |

| | | | | | | | | | | | | | | | |
|---|---|---|---|---|---|---|---|---|---|---|---|---|---|---|---|
| AT2G39400 | | alpha/beta-Hydrolases superfamily protein | 35 kDa | 0 | 0 | 3 | 3 | 6 | | | | | | | |
| AT2G39840 | | type one serine/threonine protein phosphatase 4 | 36 kDa | 0 | 0 | 3 | 3 | 6 | | | | | | | |
| AT2G47900 | | tubby like protein 3 | 45 kDa | 0 | 0 | 3 | 3 | 6 | | | | | | | |
| AT3G11040 | | Glycosyl hydrolase family 85 | 79 kDa | 0 | 0 | 3 | 3 | 6 | | | | | | | |
| AT3G11330 | | plant intracellular ras group-related LRR 9 | 56 kDa | 2 | 0 | 3 | 3 | 8 | | | | | | | |
| AT3G14110 | AT3G14110.2;AT3G14110.1;AT3G14110.3 | Tetratricopeptide repeat (TPR)-like superfamily protein | 35 kDa | 0 | 0 | 3 | 3 | 6 | | | | 2.6E+06 | 1.2E+06 | 0.0E+00 | 3.8E+06 |
| AT3G18420 | | Protein prenylyltransferase superfamily protein | 36 kDa | 0 | 0 | 3 | 3 | 6 | | 0.43 | 0.43 | | | | |
| AT3G22630 | | 20S proteasome beta subunit D1 | 23 kDa | 0 | 0 | 3 | 3 | 6 | | | | | | | |
| AT3G43520 | AT3G43520.1 | Transmembrane proteins 14C | 25 kDa | 3 | 2 | 3 | 3 | 11 | 0.07 | 0.29 | 0.18 | | | | |
| AT3G51370 | | Protein phosphatase 2C family protein | 42 kDa | 0 | 0 | 3 | 3 | 6 | | | | | | | |
| AT3G60750 | AT3G60750.2;AT3G60750.1 | Transketolase | 80 kDa | 0 | 0 | 3 | 3 | 6 | | | | 2.2E+06 | 6.6E+06 | 0.0E+00 | 8.8E+06 |
| AT4G00630 | AT4G00630.1;AT4G00630.2 | K+ efflux antiporter 2 | 126 kDa | 0 | 0 | 3 | 3 | 6 | | 0.23 | 0.23 | 3.8E+05 | 1.5E+05 | 0.0E+00 | 5.3E+05 |
| AT4G16190 | | Papain family cysteine protease | 41 kDa | 0 | 0 | 3 | 3 | 6 | | | | | | | |
| AT4G20130 | | plastid transcriptionally active 14 | 56 kDa | 0 | 0 | 3 | 3 | 6 | | | | | | | |
| AT4G25180 | | RNA polymerase III RPC4 | 34 kDa | 0 | 0 | 3 | 3 | 6 | | | | | | | |
| AT5G03455 | | Rhodanese/Cell cycle control phosphatase superfamily | 16 kDa | 0 | 0 | 3 | 3 | 6 | | | | | | | |
| AT5G08300 | AT5G08300.1 | Succinyl-CoA ligase, alpha subunit | 36 kDa | 0 | 0 | 3 | 3 | 6 | | | | 6.3E+06 | 9.6E+06 | 0.0E+00 | 1.6E+07 |
| AT5G10950 | AT5G10950.1 | Tudor/PWWP/MBT superfamily protein | 44 kDa | 0 | 0 | 3 | 3 | 6 | | | | 9.4E+05 | 1.9E+06 | 0.0E+00 | 2.9E+06 |
| AT5G12080 | | mechanosensitive channel of small conductance-like 1( | 83 kDa | 0 | 0 | 3 | 3 | 6 | | | | | | | |
| AT5G17380 | | Thiamine pyrophosphate dependent pyruvate decarbo | 61 kDa | 0 | 0 | 3 | 3 | 6 | | | | | | | |
| AT5G21160 | AT5G21160.1;AT5G21160.2;AT5G21160.3 | LA RNA-binding protein | 91 kDa | 0 | 0 | 3 | 3 | 6 | | | | 2.6E+05 | 1.9E+05 | 3.5E+04 | 4.8E+05 |
| AT5G24490 | AT5G24490.1 | 30S ribosomal protein, putative | 35 kDa | 0 | 1 | 3 | 3 | 7 | | | | 5.6E+06 | 4.2E+06 | 0.0E+00 | 9.8E+06 |
| AT5G39940 | | FAD/NAD(P)-binding oxidoreductase family protein | 52 kDa | 0 | 0 | 3 | 3 | 6 | | | | | | | |
| AT5G54750 | | Transport protein particle (TRAPP) component | 21 kDa | 0 | 0 | 3 | 3 | 6 | 0.08 | | 0.08 | | | | |
| AT5G57030 | | Lycopene beta/epsilon cyclase protein | 58 kDa | 0 | 0 | 3 | 3 | 6 | | | | | | | |
| AT5G61130 | AT5G61130.1 | plasmodesmata callose-binding protein 1 | 20 kDa | 0 | 1 | 3 | 3 | 7 | | | | 2.9E+06 | 3.1E+06 | 1.2E+06 | 7.3E+06 |
| AT5G64610 | | histone acetyltransferase of the MYST family 1 | 51 kDa | 0 | 0 | 3 | 3 | 6 | | | | | | | |
| AT1G06410 | | trehalose-phosphatase/synthase 7 | 97 kDa | 0 | 0 | 3 | 2 | 5 | | | | | | | |
| AT1G09140 | | SERINE-ARGININE PROTEIN 30 | 30 kDa | 0 | 0 | 3 | 2 | 5 | | | | | | | |
| AT1G10270 | AT1G10270.1 | glutamine-rich protein 23 | 102 kDa | 0 | 0 | 3 | 2 | 5 | | | | 4.5E+04 | 8.7E+04 | 9.8E+04 | 2.3E+05 |
| AT1G17690 | | FUNCTIONS IN: molecular_function unknown; INVOLVE | 87 kDa | 0 | 0 | 3 | 2 | 5 | | | | | | | |
| AT1G18480 | | Calcineurin-like metallo-phosphoesterase superfamily | | 43 kDa | 0 | 0 | 3 | 2 | 5 | | | | | | |
| AT1G18500 | | methylthioalkylmalate synthase-like 4 | 69 kDa | 0 | 0 | 3 | 2 | 5 | | | | | | | |
| AT1G21720 | | proteasome beta subunit C1 | 23 kDa | 0 | 0 | 3 | 2 | 5 | 0.11 | | 0.11 | | | | |
| AT1G26340 | AT1G26340.1 | cytochrome B5 isoform A | 15 kDa | 2 | 0 | 3 | 2 | 7 | 0.17 | | 0.17 | | | | |
| AT1G29690 | | MAC/Perforin domain-containing protein | 62 kDa | 0 | 0 | 3 | 2 | 5 | | | | | | | |
| AT1G32170 | | xyloglucan endotransglucosylase/hydrolase 30 | 40 kDa | 0 | 0 | 3 | 2 | 5 | | | | | | | |
| AT1G32730 | | unknown protein; LOCATED IN: chloroplast; EXPRESSEI | 36 kDa | 0 | 0 | 3 | 2 | 5 | | | | | | | |
| AT1G32750 | | HAC13 protein (HAC13) | 217 kDa | 0 | 0 | 3 | 2 | 5 | | | | | | | |
| AT1G61350 | | ARM repeat superfamily protein | 63 kDa | 0 | 0 | 3 | 2 | 5 | | | | | | | |
| AT1G61980 | | Mitochondrial transcription termination factor family | 47 kDa | 0 | 0 | 3 | 2 | 5 | | | | | | | |
| AT1G71840 | | transducin family protein / WD-40 repeat family protei | 44 kDa | 0 | 0 | 3 | 2 | 5 | | | | | | | |
| AT1G73990 | | signal peptide peptidase | 75 kDa | 0 | 0 | 3 | 2 | 5 | | | | | | | |
| AT1G75170 | | Sec14p-like phosphatidylinositol transfer family protei | 34 kDa | 0 | 1 | 3 | 2 | 6 | | | | | | | |
| AT1G76630 | AT1G76630.3;AT1G76630.2;AT1G76630.1 | Tetratricopeptide repeat (TPR)-like superfamily protein | 130 kDa | 0 | 0 | 3 | 2 | 5 | | | | 5.9E+05 | 3.2E+05 | 3.9E+05 | 1.3E+06 |
| AT1G78060 | | Glycosyl hydrolase family protein | 84 kDa | 0 | 0 | 3 | 2 | 5 | | | | | | | |
| AT1G79570 | | Protein kinase superfamily protein with octicosapeptic | 137 kDa | 0 | 0 | 3 | 2 | 5 | | | | | | | |
| AT1G80870 | | Protein kinase superfamily protein | 79 kDa | 0 | 0 | 3 | 2 | 5 | | | | | | | |
| AT2G18910 | | hydroxyproline-rich glycoprotein family protein | 14 kDa | 7 | 0 | 3 | 2 | 12 | | | | | | | |
| AT2G22780 | | peroxisomal NAD-malate dehydrogenase 1 | 37 kDa | 0 | 0 | 3 | 2 | 5 | | 0.31 | 0.31 | | | | |
| AT2G30260 | | U2 small nuclear ribonucleoprotein B | 26 kDa | 0 | 3 | 3 | 2 | 8 | | | | | | | |
| AT2G36290 | | alpha/beta-Hydrolases superfamily protein | 40 kDa | 0 | 0 | 3 | 2 | 5 | | | | | | | |
| AT2G38750 | | annexin 4 | 36 kDa | 0 | 1 | 3 | 2 | 6 | | | | | | | |
| AT2G40410 | | Staphylococcal nuclease homologue | 25 kDa | 0 | 2 | 3 | 2 | 7 | | | | | | | |
| AT2G41040 | | S-adenosyl-L-methionine-dependent methyltransferase | 39 kDa | 0 | 0 | 3 | 2 | 5 | | | | | | | |
| AT2G45000 | | structural constituent of nuclear pore | 73 kDa | 0 | 0 | 3 | 2 | 5 | | | | | | | |
| AT2G45030 | | Translation elongation factor EFG/EF2 protein | 83 kDa | 0 | 0 | 3 | 2 | 5 | | | | | | | |
| AT2G47840 | | Uncharacterised conserved protein ycf60 | ? | 0 | 0 | 3 | 2 | 5 | | | | | | | |
| AT3G02220 | AT3G02220.1 | unknown protein; CONTAINS InterPro DOMAIN/s: Prote | 26 kDa | 0 | 0 | 3 | 2 | 5 | | | | 1.2E+06 | 8.5E+05 | 3.4E+05 | 2.3E+06 |
| AT3G05070 | AT3G05070.1 | CONTAINS InterPro DOMAIN/s: mRNA splicing factor, C | 16 kDa | 0 | 0 | 3 | 2 | 5 | | | | 3.3E+06 | 5.0E+06 | 2.2E+06 | 1.0E+07 |
| AT3G05630 | | phospholipase D P2 | 119 kDa | 0 | 0 | 3 | 2 | 5 | | | | | | | |
| AT3G05810 | AT3G05810.2;AT3G05810.1 | FUNCTIONS IN: molecular_function unknown; INVOLVE | ? | 0 | 0 | 3 | 2 | 5 | | 0.81 | 0.81 | 6.6E+06 | 1.0E+07 | 0.0E+00 | 1.7E+07 |
| AT3G07800 | | Thymidine kinase | 26 kDa | 0 | 0 | 3 | 2 | 5 | | | | | | | |
| AT3G11780 | | MD-2-related lipid recognition domain-containing pro | 17 kDa | 2 | 0 | 3 | 2 | 7 | | | | | | | |
| AT3G13720 | | PRA1 (Prenylated rab acceptor) family protein | 21 kDa | 0 | 0 | 3 | 2 | 5 | | | | | | | |
| AT3G16940 | | calmodulin binding;transcription regulators | 96 kDa | 0 | 0 | 3 | 2 | 5 | | | | | | | |
| AT3G19630 | | Radical SAM superfamily protein | 41 kDa | 0 | 0 | 3 | 2 | 5 | | | | | | | |
| AT3G20390 | | endoribonuclease L-PSP family protein | 20 kDa | 0 | 0 | 3 | 2 | 5 | | | | | | | |
| AT3G23670 | | phragmoplast-associated kinesin-related protein, puta | 147 kDa | 0 | 0 | 3 | 2 | 5 | | | | | | | |
| AT3G24090 | | glutamine-fructose-6-phosphate transaminase (isomeri | 75 kDa | 0 | 0 | 3 | 2 | 5 | | | | | | | |
| AT3G26380 | AT3G26380.2;AT3G26380.1 | Melibiase family protein | 72 kDa | 0 | 0 | 3 | 2 | 5 | | | | 7.1E+05 | 7.0E+05 | 3.3E+05 | 1.7E+06 |
| AT3G48150 | | anaphase-promoting complex subunit 8 | 67 kDa | 0 | 0 | 3 | 2 | 5 | | | | | | | |
| AT3G51390 | | DHHC-type zinc finger family protein | 39 kDa | 0 | 0 | 3 | 2 | 5 | | | | | | | |
| AT3G56820 | | unknown protein; Has 34 Blast hits to 34 proteins in 13 | 24 kDa | 0 | 1 | 3 | 2 | 6 | | | | | | | |
| AT3G61600 | | POZ/BTB containin G-protein 1 | 63 kDa | 0 | 1 | 3 | 2 | 6 | | | | | | | |
| AT4G00970 | | cysteine-rich RLK (RECEPTOR-like protein kinase) 41 | 75 kDa | 0 | 0 | 3 | 2 | 5 | | | | | | | |
| AT4G14070 | | acyl-activating enzyme 15 | 81 kDa | 0 | 0 | 3 | 2 | 5 | | | | | | | |
| AT4G15930 | | Dynein light chain type 1 family protein | 14 kDa | 0 | 0 | 3 | 2 | 5 | | | | | | | |
| AT4G16490 | | ARM repeat superfamily protein | 50 kDa | 0 | 0 | 3 | 2 | 5 | | | | | | | |
| AT4G19180 | | GDA1/CD39 nucleoside phosphatase family protein | 82 kDa | 0 | 0 | 3 | 2 | 5 | | | | | | | |
| AT4G22720 | AT4G22720.2;AT4G22720.1 | Actin-like ATPase superfamily protein | 39 kDa | 0 | 0 | 3 | 2 | 5 | | | | | | | |
| AT4G29910 | | origin recognition complex protein 5 | 61 kDa | 0 | 0 | 3 | 2 | 5 | | | | | | | |
| AT4G34200 | AT4G34200.1 | D-3-phosphoglycerate dehydrogenase | 63 kDa | 0 | 0 | 3 | 2 | 5 | | | | 3.2E+05 | 1.1E+06 | 0.0E+00 | 1.4E+06 |
| AT5G02530 | AT5G02530.2;AT5G02530.1 | RNA-binding (RRM/RBD/RNP motifs) family protein | 31 kDa | 0 | 0 | 3 | 2 | 5 | | | | 3.9E+06 | 4.9E+05 | 0.0E+00 | 4.4E+06 |
| AT5G03900 | AT5G03900.2 | Iron-sulphur cluster biosynthesis family protein | 60 kDa | 0 | 0 | 3 | 2 | 5 | | | | 3.7E+05 | 4.4E+05 | 0.0E+00 | 8.0E+05 |
| AT5G07830 | AT5G07830.1 | glucuronidase 2 | 60 kDa | 0 | 0 | 3 | 2 | 5 | | | | 3.1E+06 | 3.6E+05 | 0.0E+00 | 3.5E+06 |
| AT5G13200 | | GRAM domain family protein | 30 kDa | 0 | 1 | 3 | 2 | 6 | | | | | | | |
| AT5G19485 | AT5G19485.1 | transferases;nucleotidyltransferases | 49 kDa | 0 | 0 | 3 | 2 | 5 | | | | 9.1E+05 | 6.1E+05 | 1.5E+05 | 1.7E+06 |
| AT5G23150 | | Tudor/PWWP/MBT domain-containing protein | 151 kDa | 0 | 0 | 3 | 2 | 5 | | | | | | | |
| AT5G24020 | | septum site-determining protein (MIND) | 36 kDa | 0 | 0 | 3 | 2 | 5 | | | | | | | |
| AT5G28040 | | DNA-binding storekeeper protein-related transcription | 47 kDa | 0 | 0 | 3 | 2 | 5 | | | | | | | |
| AT5G41970 | | Metal-dependent protein hydrolase | 42 kDa | 0 | 0 | 3 | 2 | 5 | | | | | | | |
| AT5G43970 | | translocase of outer membrane 22-V | 10 kDa | 0 | 0 | 3 | 2 | 5 | | 0.13 | 0.13 | | | | |
| AT5G46920 | | Intron maturase, type II family protein | 85 kDa | 0 | 0 | 3 | 2 | 5 | | | | | | | |
| AT5G53060 | | RNA-binding KH domain-containing protein | 71 kDa | 0 | 0 | 3 | 2 | 5 | | | | | | | |
| AT5G60160 | AT5G60160.1 | Zn-dependent exopeptidases superfamily protein | 52 kDa | 0 | 0 | 3 | 2 | 5 | | | | 1.0E+06 | 9.7E+05 | 0.0E+00 | 2.0E+06 |
| AT1G01910 | | P-loop containing nucleoside triphosphate hydrolases | 40 kDa | 0 | 0 | 3 | 1 | 4 | | | | | | | |
| AT1G03330 | | Small nuclear ribonucleoprotein family protein | 11 kDa | 0 | 0 | 3 | 1 | 4 | | | | | | | |

| | | | | | | | | | | | | | | | |
|---|---|---|---|---|---|---|---|---|---|---|---|---|---|---|---|
| AT1G04030 | | unknown protein; BEST Arabidopsis thaliana protein m | ? | 0 | 0 | 3 | 1 | 4 | | | | | | | |
| AT1G04780 | | Ankyrin repeat family protein | 74 kDa | 0 | 0 | 3 | 1 | 4 | | | | | | | |
| AT1G05205 | | unknown protein; Has 44 Blast hits to 44 proteins in 1€ | 11 kDa | 0 | 0 | 3 | 1 | 4 | | | | | | | |
| AT1G09900 | | Pentatricopeptide repeat (PPR-like) superfamily protei | 66 kDa | 0 | 0 | 3 | 1 | 4 | | | | | | | |
| AT1G48920 | AT1G48920.1 | nucleolin like 1 | ? | 0 | 0 | 3 | 1 | 4 | | | | 1.4E+06 | 2.7E+06 | 2.9E+05 | 4.4E+06 |
| AT1G51680 | | 4-coumarate:CoA ligase 1 | 59 kDa | 0 | 0 | 3 | 1 | 4 | | | | | | | |
| AT1G54290 | | Translation initiation factor SUI1 family protein | 13 kDa | 0 | 0 | 3 | 1 | 4 | | 0.05 | 0.05 | | | | |
| AT1G79150 | AT1G79150.1 | binding | 95 kDa | 0 | 0 | 3 | 1 | 4 | | | | 0.0E+00 | 1.1E+05 | 8.0E+04 | 1.9E+05 |
| AT2G03150 | | ATP/GTP-binding protein family | 151 kDa | 0 | 0 | 3 | 1 | 4 | | | | | | | |
| AT2G16060 | AT2G16060.1 | hemoglobin 1 | 18 kDa | 0 | 0 | 3 | 1 | 4 | | | | 1.6E+06 | 9.7E+05 | 0.0E+00 | 2.6E+06 |
| AT2G25737 | | Sulfite exporter TauE/SafE family protein | 52 kDa | 0 | 0 | 3 | 1 | 4 | | | | | | | |
| AT2G30950 | | FtsH extracellular protease family | 74 kDa | 0 | 1 | 3 | 1 | 5 | | | | | | | |
| AT2G35490 | | Plastid-lipid associated protein PAP / fibrillin family pr | 41 kDa | 0 | 0 | 3 | 1 | 4 | | | | | | | |
| AT2G35980 | | Late embryogenesis abundant (LEA) hydroxyproline-ric | 26 kDa | 0 | 0 | 3 | 1 | 4 | | | | | | | |
| AT2G42400 | | vascular plant one zinc finger protein 2 | 51 kDa | 0 | 0 | 3 | 1 | 4 | | | | | | | |
| AT2G46230 | AT2G46230.1 | PIN domain-like family protein | 22 kDa | 0 | 0 | 3 | 1 | 4 | | | | 8.8E+05 | 1.0E+06 | 3.1E+05 | 2.2E+06 |
| AT3G05910 | | Pectinacetylesterase family protein | 46 kDa | 0 | 0 | 3 | 1 | 4 | | | | | | | |
| AT3G06960 | | pigment defective 320 | ? | 0 | 0 | 3 | 1 | 4 | | | | | | | |
| AT3G07230 | AT3G07230.1 | wound-responsive protein-related | 5 kDa | 0 | 0 | 3 | 1 | 4 | | 0.16 | 0.16 | 3.8E+08 | 5.0E+08 | 9.2E+07 | 9.8E+08 |
| AT3G10920 | AT3G10920.2;AT3G10920.1 | manganese superoxide dismutase 1 | 25 kDa | 0 | 0 | 3 | 1 | 4 | | | | 8.3E+06 | 1.2E+07 | 0.0E+00 | 2.0E+07 |
| AT3G11070 | | Outer membrane OMP85 family protein | 57 kDa | 0 | 0 | 3 | 1 | 4 | | | | | | | |
| AT3G44340 | | clone eighty-four | 115 kDa | 0 | 0 | 3 | 1 | 4 | | | | | | | |
| AT3G44530 | | homolog of histone chaperone HIRA | 116 kDa | 0 | 0 | 3 | 1 | 4 | | | | | | | |
| AT3G46940 | | DUTP-PYROPHOSPHATASE-LIKE 1 | 18 kDa | 0 | 0 | 3 | 1 | 4 | | | | | | | |
| AT3G55340 | AT3G55340.1 | phragmoplastin interacting protein 1 | 66 kDa | 0 | 0 | 3 | 1 | 4 | | | | | | | |
| AT3G55510 | AT3G55510.2;AT3G55510.1 | Noc2p family | 68 kDa | 0 | 0 | 3 | 1 | 4 | | | | | | | |
| AT3G57180 | | P-loop containing nucleoside triphosphate hydrolases | 73 kDa | 0 | 0 | 3 | 1 | 4 | | | | | | | |
| AT4G17260 | AT4G17260.1 | Lactate/malate dehydrogenase family protein | 38 kDa | 0 | 0 | 3 | 1 | 4 | | | | | | | |
| AT4G22305 | | alpha/beta-Hydrolases superfamily protein | 25 kDa | 0 | 1 | 3 | 1 | 5 | | | | | | | |
| AT4G31130 | | Protein of unknown function (DUF1218) | 21 kDa | 0 | 0 | 3 | 1 | 4 | | | | | | | |
| AT4G35110 | | Arabidopsis phospholipase-like protein (PEARLI 4) fami | 45 kDa | 0 | 0 | 3 | 1 | 4 | | | | | | | |
| AT4G35240 | | Protein of unknown function (DUF630 and DUF632) | 96 kDa | 0 | 0 | 3 | 1 | 4 | | | | | | | |
| AT4G39460 | | S-adenosylmethionine carrier 1 | 35 kDa | 0 | 0 | 3 | 1 | 4 | | 0.28 | 0.28 | | | | |
| AT5G08660 | | Protein of unknown function (DUF668) | 73 kDa | 0 | 0 | 3 | 1 | 4 | | | | | | | |
| AT5G11030 | | aberrant lateral root formation 4 | 64 kDa | 0 | 0 | 3 | 1 | 4 | | | | | | | |
| AT5G23250 | AT5G23250.1;AT5G23250.2;AT5G23250.3 | Succinyl-CoA ligase, alpha subunit | 35 kDa | 0 | 0 | 3 | 1 | 4 | | | | 2.6E+06 | 2.3E+06 | 0.0E+00 | 4.9E+06 |
| AT5G27550 | AT5G27550.2;AT5G27550.1 | P-loop containing nucleoside triphosphate hydrolases | 85 kDa | 0 | 0 | 3 | 1 | 4 | | | | 1.3E+05 | 1.3E+05 | 0.0E+00 | 2.6E+05 |
| AT5G34930 | | arogenate dehydrogenase | 72 kDa | 0 | 0 | 3 | 1 | 4 | | | | | | | |
| AT5G37370 | AT5G37370.7;AT5G37370.6;AT5G37370.5;AT5G37370.4;AT5G37370.2;AT5G37370.1 | PRP38 family protein | 46 kDa | 0 | 0 | 3 | 1 | 4 | | | | 1.8E+05 | 2.2E+05 | 0.0E+00 | 3.9E+05 |
| AT5G43090 | | pumilio 13 | 60 kDa | 0 | 0 | 3 | 1 | 4 | | | | | | | |
| AT5G47870 | AT5G47870.1 | FUNCTIONS IN: molecular_function unknown; INVOLVE | 21 kDa | 0 | 0 | 3 | 1 | 4 | | | | | | | |
| AT5G53230 | AT5G53230.1 | Protein of unknown function (DUF295) | 40 kDa | 0 | 0 | 3 | 1 | 4 | | | | | | | |
| AT5G55210 | | unknown protein; FUNCTIONS IN: molecular_function | ? | 0 | 0 | 3 | 1 | 4 | | | | | | | |
| AT5G58320 | | Kinase interacting (KIP1-like) family protein | 60 kDa | 0 | 0 | 3 | 1 | 4 | | | | | | | |
| AT5G61020 | | evolutionarily conserved C-terminal region 3 | 55 kDa | 0 | 0 | 3 | 1 | 4 | | | | | | | |
| AT5G61070 | | histone deacetylase of the RPD3/HDA1 superfamily 18 | 77 kDa | 0 | 0 | 3 | 1 | 4 | | | | | | | |
| AT5G64300 | AT5G64300.1 | GTP cyclohydrolase II | 59 kDa | 0 | 0 | 3 | 1 | 4 | | | | 1.6E+05 | 1.5E+05 | 0.0E+00 | 3.1E+05 |
| AT5G65200 | | plant U-box 38 | 62 kDa | 0 | 0 | 3 | 1 | 4 | | | | | | | |
| AT1G14870 | | PLANT CADMIUM RESISTANCE 2 | 17 kDa | 7 | 0 | 3 | 0 | 10 | | | | | | | |
| AT1G48090 | AT1G48090.5;AT1G48090.6;AT1G48090.4;AT1G48090.1;AT1G48090.3;AT1G48090.2 | calcium-dependent lipid-binding family protein | 462 kDa | 0 | 0 | 2 | 13 | 15 | | | | 3.2E+04 | 1.8E+04 | 0.0E+00 | 4.9E+04 |
| AT1G64650 | | Major facilitator superfamily protein | 51 kDa | 0 | 0 | 2 | 12 | 14 | | | | | | | |
| AT3G15990 | | sulfate transporter 3;4 | 71 kDa | 0 | 0 | 2 | 12 | 14 | | | | | | | |
| AT4G02450 | AT4G02450.2;AT4G02450.1 | HSP20-like chaperones superfamily protein | 25 kDa | 2 | 5 | 2 | 10 | 19 | | 2.14 | 2.14 | 9.2E+06 | 5.7E+06 | 0.0E+00 | 1.5E+07 |
| AT1G23820 | | spermidine synthase 1 | 42 kDa | 0 | 0 | 2 | 9 | 11 | | | | | | | |
| AT2G39770 | AT2G39770.3;AT2G39770.2;AT2G39770.1 | Glucose-1-phosphate adenylyltransferase family protei | 40 kDa | 0 | 0 | 2 | 9 | 11 | | | | 1.0E+06 | 8.1E+05 | 0.0E+00 | 1.8E+06 |
| AT5G41790 | AT5G41790.2;AT5G41790.1 | COP1-interactive protein 1 | 182 kDa | 0 | 0 | 2 | 9 | 11 | | | | 3.1E+05 | 2.0E+05 | 0.0E+00 | 5.1E+05 |
| AT5G61210 | | soluble N-ethylmaleimide-sensitive factor adaptor prot | 34 kDa | 5 | 0 | 2 | 9 | 16 | | | | | | | |
| AT2G01070 | | Lung seven transmembrane receptor family protein | 56 kDa | 0 | 0 | 2 | 8 | 10 | | | | | | | |
| AT4G28320 | | Glycosyl hydrolase superfamily protein | 50 kDa | 0 | 2 | 2 | 8 | 12 | | | | | | | |
| AT2G17440 | | plant intracellular ras group-related LRR 5 | 59 kDa | 0 | 0 | 2 | 7 | 9 | | | | | | | |
| AT3G19840 | | pre-mRNA-processing protein 40C | 93 kDa | 0 | 0 | 2 | 7 | 9 | | | | | | | |
| AT4G25770 | | alpha/beta-Hydrolases superfamily protein | 47 kDa | 0 | 0 | 2 | 7 | 9 | | | | | | | |
| AT1G51690 | AT1G51690.2;AT1G51690.4;AT1G51690.1;AT1G51690.3;AT1G51690.5 | protein phosphatase 2A 55 kDa regulatory subunit B al | 57 kDa | 0 | 0 | 2 | 6 | 8 | | | | 2.3E+06 | 1.6E+06 | 0.0E+00 | 3.9E+06 |
| AT1G68410 | | Protein phosphatase 2C family protein | 47 kDa | 0 | 0 | 2 | 6 | 8 | | | | | | | |
| AT2G01180 | | phosphatidic acid phosphatase 1 | 37 kDa | 2 | 0 | 2 | 6 | 10 | | | | | | | |
| AT2G16860 | AT2G16860.2;AT2G16860.1 | GCIP-interacting family protein | 35 kDa | 0 | 0 | 2 | 6 | 8 | | | | 7.6E+05 | 1.0E+06 | 0.0E+00 | 1.8E+06 |
| AT2G19560 | | proteasome family protein | 48 kDa | 0 | 0 | 2 | 6 | 8 | | | | | | | |
| AT2G37180 | | Aquaporin-like superfamily protein | 30 kDa | 0 | 0 | 2 | 6 | 8 | | | | | | | |
| AT3G03220 | | expansin A13 | 29 kDa | 0 | 4 | 2 | 6 | 12 | | | | | | | |
| AT3G04080 | | apyrase 1 | 51 kDa | 0 | 1 | 2 | 6 | 9 | | | | | | | |
| AT3G48430 | AT3G48430.1 | relative of early flowering 6 | 153 kDa | 0 | 0 | 2 | 6 | 8 | | | | 2.2E+06 | 1.2E+06 | 1.7E+05 | 3.6E+06 |
| AT3G54090 | | fructokinase-like 1 | 54 kDa | 0 | 0 | 2 | 6 | 8 | | | | | | | |
| AT3G60260 | | ELMO/CED-12 family protein | 30 kDa | 0 | 0 | 2 | 6 | 8 | | | | | | | |
| AT3G60450 | | Phosphoglycerate mutase family protein | 31 kDa | 0 | 0 | 2 | 6 | 8 | | | | | | | |
| AT4G15570 | | P-loop containing nucleoside triphosphate hydrolases | 92 kDa | 0 | 0 | 2 | 6 | 8 | | | | | | | |
| AT4G17430 | | O-fucosyltransferase family protein | 57 kDa | 0 | 0 | 2 | 6 | 8 | | | | | | | |
| AT4G21534 | | Diacylglycerol kinase family protein | 54 kDa | 0 | 0 | 2 | 6 | 8 | | | | | | | |
| AT5G13850 | | nascent polypeptide-associated complex subunit alpha | 22 kDa | 0 | 1 | 2 | 6 | 9 | | | | | | | |
| AT5G16440 | | isopentenyl diphosphate isomerase 1 | 33 kDa | 0 | 1 | 2 | 6 | 9 | | | | | | | |
| AT5G51830 | | pfkB-like carbohydrate kinase family protein | 37 kDa | 0 | 0 | 2 | 6 | 8 | | | | | | | |
| AT5G56350 | | Pyruvate kinase family protein | 54 kDa | 0 | 0 | 2 | 6 | 8 | | | | | | | |
| AT1G05840 | | Eukaryotic aspartyl protease family protein | 53 kDa | 0 | 0 | 2 | 5 | 7 | | | | | | | |
| AT1G11180 | | Secretory carrier membrane protein (SCAMP) family pr | 33 kDa | 0 | 0 | 2 | 5 | 7 | | | | | | | |
| AT1G11310 | | Seven transmembrane MLO family protein | 66 kDa | 0 | 0 | 2 | 5 | 7 | | | | | | | |
| AT1G15750 | | Transducin family protein / WD-40 repeat family protei | 124 kDa | 0 | 0 | 2 | 5 | 7 | | 0.32 | 0.32 | | | | |
| AT1G16010 | | magnesium transporter 2 | 50 kDa | 0 | 0 | 2 | 5 | 7 | | | | | | | |
| AT1G20630 | AT1G20630.1 | catalase 1 | 57 kDa | 0 | 1 | 2 | 5 | 8 | | | | 2.6E+06 | 2.2E+06 | 0.0E+00 | 4.8E+06 |
| AT1G50250 | | FTSH protease 1 | 77 kDa | 0 | 0 | 2 | 5 | 7 | | | | | | | |
| AT1G52310 | | protein kinase family protein / C-type lectin domain-co | 62 kDa | 0 | 0 | 2 | 5 | 7 | | | | | | | |
| AT1G67940 | | non-intrinsic ABC protein 3 | 29 kDa | 0 | 2 | 2 | 5 | 9 | 0.12 | 22.50 | 11.31 | | | | |
| AT2G37160 | | Transducin/WD40 repeat-like superfamily protein | 58 kDa | 0 | 0 | 2 | 5 | 7 | | | | | | | |
| AT2G45670 | | calcineurin B subunit-related | 61 kDa | 0 | 2 | 2 | 5 | 9 | | | | | | | |
| AT3G11700 | | FASCICLIN-like arabinogalactan protein 18 precursor | 51 kDa | 0 | 0 | 2 | 5 | 7 | | | | | | | |

| | | | | | | | | | | | | | | | |
|---|---|---|---|---|---|---|---|---|---|---|---|---|---|---|---|
| AT3G13410 | | unknown protein; FUNCTIONS IN: molecular_function | ? | 0 | 6 | 2 | 5 | 13 | | | | | | | |
| AT3G19370 | | Plant protein of unknown function (DUF869) | 82 kDa | 0 | 0 | 2 | 5 | 7 | | | | | | | |
| AT4G11850 | | phospholipase D gamma 1 | 96 kDa | 0 | 0 | 2 | 5 | 7 | | | | | | | |
| AT4G18070 | | unknown protein; BEST Arabidopsis thaliana protein m | 28 kDa | 0 | 1 | 2 | 5 | 8 | | | | | | | |
| AT4G27350 | | Protein of unknown function (DUF1223) | 30 kDa | 0 | 4 | 2 | 5 | 11 | | 3.44 | 3.44 | | | | |
| AT4G29810 | | MAP kinase kinase 2 | 40 kDa | 0 | 0 | 2 | 5 | 7 | | | | | | | |
| AT4G32350 | | Regulator of Vps4 activity in the MVB pathway protein | 84 kDa | 0 | 0 | 2 | 5 | 7 | | | | | | | |
| AT5G01220 | | sulfoquinovosyldiacylglycerol 2 | 57 kDa | 0 | 0 | 2 | 5 | 7 | | | | | | | |
| AT5G05180 | | unknown protein; BEST Arabidopsis thaliana protein m | 47 kDa | 0 | 0 | 2 | 5 | 7 | | | | | | | |
| AT5G06730 | | Peroxidase superfamily protein | 37 kDa | 0 | 0 | 2 | 5 | 7 | | | | | | | |
| AT5G17930 | | MIF4G domain-containing protein / MA3 domain-cont | 90 kDa | 0 | 0 | 2 | 5 | 7 | | | | | | | |
| AT5G37600 | | glutamine synthase clone R1 | 39 kDa | 0 | 0 | 2 | 5 | 7 | | | | | | | |
| AT5G55670 | AT5G55670.2;AT5G55670.1 | RNA-binding (RRM/RBD/RNP motifs) family protein | 76 kDa | 0 | 0 | 2 | 5 | 7 | | | | | | | |
| AT1G04970 | | lipid-binding serum glycoprotein family protein | 53 kDa | 0 | 0 | 2 | 4 | 6 | | | | | | | |
| AT1G17100 | | SOUL heme-binding family protein | 25 kDa | 0 | 1 | 2 | 4 | 7 | | | | | | | |
| AT1G18210 | AT1G18210.2;AT1G18210.1 | Calcium-binding EF-hand family protein | 18 kDa | 2 | 0 | 2 | 4 | 8 | | | | 1.7E+07 | 6.1E+06 | 0.0E+00 | 2.3E+07 |
| AT1G27480 | | alpha/beta-Hydrolases superfamily protein | 48 kDa | 0 | 0 | 2 | 4 | 6 | | | | | | | |
| AT1G27760 | | interferon-related developmental regulator family prot | 49 kDa | 0 | 0 | 2 | 4 | 6 | | | | | | | |
| AT1G55460 | | DNA/RNA-binding protein Kin17, conserved region | 47 kDa | 0 | 0 | 2 | 4 | 6 | | | | | | | |
| AT1G65650 | | Peptidase C12, ubiquitin carboxyl-terminal hydrolase : | 37 kDa | 0 | 0 | 2 | 4 | 6 | | | | | | | |
| AT1G76970 | | Target of Myb protein 1 | 49 kDa | 0 | 1 | 2 | 4 | 7 | | | | | | | |
| AT1G77220 | | Protein of unknown function (DUF300) | 55 kDa | 0 | 0 | 2 | 4 | 6 | | | | | | | |
| AT2G01350 | | quinolinate phoshoribosyltransferase | 38 kDa | 0 | 0 | 2 | 4 | 6 | | | | | | | |
| AT2G26430 | | arginine-rich cyclin 1 | 48 kDa | 0 | 0 | 2 | 4 | 6 | | | | | | | |
| AT2G27060 | AT2G27060.3;AT2G27060.1 | Leucine-rich repeat protein kinase family protein | 110 kDa | 1 | 0 | 2 | 4 | 7 | | | | 3.5E+05 | 1.1E+05 | 0.0E+00 | 4.7E+05 |
| AT2G37110 | | PLAC8 family protein | 27 kDa | 0 | 3 | 2 | 4 | 9 | | | | | | | |
| AT3G03270 | | Adenine nucleotide alpha hydrolases-like superfamily p | 23 kDa | 0 | 0 | 2 | 4 | 6 | | | | | | | |
| AT3G04600 | | Nucleotidylyl transferase superfamily protein | 46 kDa | 0 | 0 | 2 | 4 | 6 | | | | | | | |
| AT3G07360 | | plant U-box 9 | 51 kDa | 0 | 0 | 2 | 4 | 6 | | | | | | | |
| AT3G07570 | | Cytochrome b561/ferric reductase transmembrane wit | 40 kDa | 0 | 0 | 2 | 4 | 6 | | | | | | | |
| AT3G12010 | | FUNCTIONS IN: molecular_function unknown; INVOLVE | 76 kDa | 0 | 0 | 2 | 4 | 6 | | | | | | | |
| AT3G12480 | | nuclear factor Y, subunit C11 | 32 kDa | 0 | 0 | 2 | 4 | 6 | | | | | | | |
| AT3G13940 | AT3G13940.1 | DNA binding;DNA-directed RNA polymerases | 51 kDa | 0 | 0 | 2 | 4 | 6 | | | | 7.1E+05 | 4.7E+05 | 0.0E+00 | 1.2E+06 |
| AT3G20320 | | trigalactosyldiacylglycerol2 | 42 kDa | 0 | 0 | 2 | 4 | 6 | | | | | | | |
| AT3G23600 | | alpha/beta-Hydrolases superfamily protein | 26 kDa | 0 | 1 | 2 | 4 | 7 | | | | | | | |
| AT3G48890 | | membrane-associated progesterone binding protein 3 | 25 kDa | 1 | 0 | 2 | 4 | 7 | 0.13 | 0.24 | 0.18 | | | | |
| AT3G49360 | | 6-phosphogluconolactonase 2 | 29 kDa | 0 | 2 | 2 | 4 | 8 | | | | | | | |
| AT3G53970 | | proteasome inhibitor-related | 32 kDa | 0 | 0 | 2 | 4 | 6 | | | | | | | |
| AT3G62600 | | DNAJ heat shock family protein | 39 kDa | 0 | 0 | 2 | 4 | 6 | | | | | | | |
| AT4G05470 | | RNI-like superfamily protein | ? | 0 | 0 | 2 | 4 | 6 | | | | | | | |
| AT4G24610 | | unknown protein; FUNCTIONS IN: molecular_function | 130 kDa | 0 | 1 | 2 | 4 | 7 | | | | | | | |
| AT4G27585 | | SPFH/Band 7/PHB domain-containing membrane-assoc | 45 kDa | 0 | 0 | 2 | 4 | 6 | 0.08 | 0.22 | 0.15 | | | | |
| AT4G29160 | | SNF7 family protein | 24 kDa | 0 | 0 | 2 | 4 | 6 | | | | | | | |
| AT4G31200 | | SWAP (Suppressor-of-White-APricot)/surp RNA-binding | 72 kDa | 0 | 0 | 2 | 4 | 6 | | | | | | | |
| AT5G03720 | | heat shock transcription factor A3 | 46 kDa | 0 | 0 | 2 | 4 | 6 | | | | | | | |
| AT5G08160 | | serine/threonine protein kinase 3 | 39 kDa | 0 | 0 | 2 | 4 | 6 | | | | | | | |
| AT5G14470 | | GHMP kinase family protein | 41 kDa | 0 | 0 | 2 | 4 | 6 | | | | | | | |
| AT5G14720 | | Protein kinase superfamily protein | 75 kDa | 0 | 0 | 2 | 4 | 6 | | | | | | | |
| AT5G23080 | AT5G23080.2;AT5G23080.1 | SWAP (Suppressor-of-White-APricot)/surp domain-con | 105 kDa | 0 | 0 | 2 | 4 | 6 | | | | | | | |
| AT5G24690 | | Protein of unknown function (DUF3411) | 57 kDa | 0 | 0 | 2 | 4 | 6 | | | | | | | |
| AT5G48480 | | Lactoylglutathione lyase / glyoxalase I family protein | 18 kDa | 0 | 0 | 2 | 4 | 6 | | | | | | | |
| AT5G57280 | | S-adenosyl-L-methionine-dependent methyltransferase | 32 kDa | 0 | 0 | 2 | 4 | 6 | | | | | | | |
| AT1G04910 | | O-fucosyltransferase family protein | 59 kDa | 0 | 0 | 2 | 3 | 5 | | | | | | | |
| AT1G07530 | | SCARECROW-like 14 | 86 kDa | 0 | 0 | 2 | 3 | 5 | | | | | | | |
| AT1G19570 | AT1G19570.1;AT1G19550.1 | dehydroascorbate reductase | 24 kDa | 0 | 0 | 2 | 3 | 5 | | | | | | | |
| AT1G20620 | | catalase 3 | 57 kDa | 1 | 2 | 2 | 3 | 8 | | | | | | | |
| AT1G26370 | | RNA helicase family protein | 81 kDa | 0 | 0 | 2 | 3 | 5 | | | | | | | |
| AT1G27460 | | no pollen germination related 1 | 77 kDa | 0 | 0 | 2 | 3 | 5 | | | | | | | |
| AT1G48420 | | D-cysteine desulfhydrase | 44 kDa | 0 | 0 | 2 | 3 | 5 | | | | | | | |
| AT1G49820 | | S-methyl-5-thioribose kinase | 48 kDa | 0 | 0 | 2 | 3 | 5 | | | | | | | |
| AT1G52760 | AT1G52760.1 | lysophospholipase 2 | 37 kDa | 0 | 0 | 2 | 3 | 5 | | | | | | | |
| AT1G55210 | AT1G55210.2;AT1G55210.1 | Disease resistance-responsive (dirigent-like protein) fan | 21 kDa | 0 | 0 | 2 | 3 | 5 | 0.12 | | 0.12 | 2.5E+06 | 1.7E+06 | 0.0E+00 | 4.2E+06 |
| AT1G58360 | AT1G58360.1 | amino acid permease 1 | 53 kDa | 0 | 0 | 2 | 3 | 5 | | | | | | | |
| AT1G58848 | | Disease resistance protein (CC-NBS-LRR class) family | 121 kDa | 0 | 0 | 2 | 3 | 5 | | | | | | | |
| AT1G60440 | | pantothenate kinase 1 | 42 kDa | 0 | 0 | 2 | 3 | 5 | | | | | | | |
| AT1G63250 | AT1G63250.1;AT2G07750.1 | DEA(D/H)-box RNA helicase family protein | 90 kDa | 0 | 0 | 2 | 3 | 5 | | | | 3.6E+05 | 1.1E+06 | 1.1E+05 | 1.6E+06 |
| AT1G64970 | | gamma-tocopherol methyltransferase | 38 kDa | 0 | 0 | 2 | 3 | 5 | | | | | | | |
| AT1G77420 | | alpha/beta-Hydrolases superfamily protein | 43 kDa | 0 | 0 | 2 | 3 | 5 | | | | | | | |
| AT1G79500 | | Aldolase-type TIM barrel family protein | 32 kDa | 0 | 1 | 2 | 3 | 6 | | | | | | | |
| AT1G79880 | AT1G79880.2;AT1G79880.3;AT1G79880.1 | RNA recognition motif (RRM)-containing protein | 45 kDa | 0 | 0 | 2 | 3 | 5 | | | | 1.2E+06 | 9.9E+05 | 0.0E+00 | 2.2E+06 |
| AT2G14120 | | dynamin related protein | 87 kDa | 0 | 0 | 2 | 3 | 5 | | | | | | | |
| AT2G15490 | | UDP-glycosyltransferase 73B4 | 54 kDa | 0 | 0 | 2 | 3 | 5 | | | | | | | |
| AT2G19090 | | Protein of unknown function (DUF630 and DUF632) | 92 kDa | 0 | 0 | 2 | 3 | 5 | | | | | | | |
| AT2G20050 | | protein serine/threonine phosphatases;protein kinases | 121 kDa | 0 | 0 | 2 | 3 | 5 | | | | | | | |
| AT2G25840 | | Nucleotidylyl transferase superfamily protein | 46 kDa | 0 | 0 | 2 | 3 | 5 | | | | | | | |
| AT2G27090 | | Protein of unknown function (DUF630 and DUF632) | 84 kDa | 0 | 0 | 2 | 3 | 5 | | | | | | | |
| AT2G29420 | | glutathione S-transferase tau 7 | 26 kDa | 0 | 0 | 2 | 3 | 5 | | | | | | | |
| AT2G30530 | | unknown protein; LOCATED IN: cellular_component ur | 41 kDa | 0 | 0 | 2 | 3 | 5 | | | | | | | |
| AT2G34750 | | RNA polymerase I specific transcription initiation facto | 69 kDa | 0 | 0 | 2 | 3 | 5 | | | | | | | |
| AT2G36480 | | ENTH/VHS family protein | 97 kDa | 0 | 0 | 2 | 3 | 5 | | | | | | | |
| AT2G38000 | | chaperone protein dnaJ-related | 46 kDa | 0 | 0 | 2 | 3 | 5 | | | | | | | |
| AT3G03720 | | cationic amino acid transporter 4 | 86 kDa | 0 | 0 | 2 | 3 | 5 | | | | | | | |
| AT3G07790 | AT3G07790.1 | DGCR14-related | 57 kDa | 0 | 0 | 2 | 3 | 5 | | | | 3.2E+05 | 0.0E+00 | 0.0E+00 | 3.2E+05 |
| AT3G19640 | | magnesium transporter 4 | 53 kDa | 0 | 0 | 2 | 3 | 5 | | | | | | | |
| AT3G19960 | | myosin 1 | 131 kDa | 0 | 0 | 2 | 3 | 5 | | | | | | | |
| AT3G28730 | | high mobility group | 72 kDa | 0 | 0 | 2 | 3 | 5 | | | | | | | |
| AT3G45970 | | expansin-like A1 | 29 kDa | 0 | 0 | 2 | 3 | 5 | | 0.27 | 0.27 | | | | |
| AT3G47200 | | Plant protein of unknown function (DUF247) | 54 kDa | 0 | 2 | 2 | 3 | 7 | | | | | | | |
| AT3G51730 | | saposin B domain-containing protein | 24 kDa | 0 | 4 | 2 | 3 | 9 | | | | | | | |
| AT3G58180 | | ARM repeat superfamily protein | 34 kDa | 0 | 0 | 2 | 3 | 5 | | | | | | | |
| AT3G60440 | | Phosphoglycerate mutase family protein | 32 kDa | 0 | 0 | 2 | 3 | 5 | | | | | | | |
| AT3G60820 | AT3G60820.1;AT3G60820.3;AT3G60820.2 | N-terminal nucleophile aminohydrolases (Ntn hydrolas | 25 kDa | 0 | 2 | 2 | 3 | 7 | | | | 2.5E+06 | 3.6E+06 | 0.0E+00 | 6.1E+06 |
| AT3G61180 | | RING/U-box superfamily protein | 42 kDa | 0 | 0 | 2 | 3 | 5 | | | | | | | |
| AT3G62940 | | Cysteine proteinases superfamily protein | 36 kDa | 0 | 0 | 2 | 3 | 5 | | | | | | | |
| AT4G02030 | AT4G02030.1;AT4G02030.2 | Vps51/Vps67 family (components of vesicular transpor | 89 kDa | 0 | 0 | 2 | 3 | 5 | | | | 2.9E+05 | 1.3E+05 | 0.0E+00 | 4.2E+05 |
| AT4G08455 | AT4G08455.1 | BTB/POZ domain-containing protein | 27 kDa | 0 | 4 | 2 | 3 | 9 | | | | 1.1E+06 | 1.3E+06 | 0.0E+00 | 2.5E+06 |
| AT4G18710 | | Protein kinase superfamily protein | 43 kDa | 0 | 0 | 2 | 3 | 5 | | | | | | | |
| AT4G19710 | | aspartate kinase-homoserine dehydrogenase ii | 94 kDa | 0 | 0 | 2 | 3 | 5 | | | | | | | |
| AT4G37100 | | Pyridoxal phosphate (PLP)-dependent transferases supe | 101 kDa | 0 | 0 | 2 | 3 | 5 | | | | | | | |

| | | | | | | | | | | | | | | | |
|---|---|---|---|---|---|---|---|---|---|---|---|---|---|---|---|
| AT5G03380 | | Heavy metal transport/detoxification superfamily prot | 42 kDa | 0 | 0 | 2 | 3 | 5 | | | | | | | |
| AT5G14030 | | translocon-associated protein beta (TRAPB) family prot | 21 kDa | 0 | 0 | 2 | 3 | 5 | 0.08 | 0.37 | 0.23 | | | | |
| AT5G20140 | | SOUL heme-binding family protein | 43 kDa | 0 | 0 | 2 | 3 | 5 | | | | | | | |
| AT5G20280 | AT5G20280.1 | sucrose phosphate synthase 1F | 117 kDa | 0 | 0 | 2 | 3 | 5 | | | | 1.4E+06 | 9.2E+05 | 0.0E+00 | 2.4E+06 |
| AT5G22740 | | cellulose synthase-like A02 | 62 kDa | 0 | 0 | 2 | 3 | 5 | | | | | | | |
| AT5G37510 | AT5G37510.1;AT5G37510.2 | NADH-ubiquinone dehydrogenase, mitochondrial, put | 81 kDa | 0 | 0 | 2 | 3 | 5 | | | | 8.3E+05 | 1.4E+06 | 0.0E+00 | 2.3E+06 |
| AT5G40770 | AT5G40770.1 | prohibitin 3 | 30 kDa | 0 | 0 | 2 | 3 | 5 | 0.12 | 0.23 | 0.17 | 9.7E+05 | 9.1E+05 | 0.0E+00 | 1.9E+06 |
| AT5G42560 | | Abscisic acid-responsive (TB2/DP1, HVA22) family prot | 33 kDa | 0 | 0 | 2 | 3 | 5 | | | | | | | |
| AT5G42650 | | allene oxide synthase | 58 kDa | 0 | 0 | 2 | 3 | 5 | | | | | | | |
| AT5G43280 | AT5G43280.1 | delta(3,5),delta(2,4)-dienoyl-CoA isomerase 1 | 30 kDa | 0 | 0 | 2 | 3 | 5 | | | | 9.9E+05 | 1.3E+06 | 0.0E+00 | 2.3E+06 |
| AT5G54430 | AT5G54430.4;AT5G54430.1;AT5G54430.5;AT5G54430.2;AT5G54430.6;AT5G54430.3 | Adenine nucleotide alpha hydrolases-like superfamily p | 26 kDa | 0 | 0 | 2 | 3 | 5 | | | | 1.6E+07 | 2.0E+07 | 7.3E+05 | 3.7E+07 |
| AT5G55630 | | Outward rectifying potassium channel protein | 41 kDa | 0 | 0 | 2 | 3 | 5 | | | | | | | |
| AT5G57410 | | Afadin/alpha-actinin-binding protein | 45 kDa | 0 | 0 | 2 | 3 | 5 | | | | | | | |
| AT5G57490 | | voltage dependent anion channel 4 | 30 kDa | 0 | 0 | 2 | 3 | 5 | | 0.23 | 0.23 | | | | |
| AT5G64560 | | magnesium transporter 9 | 44 kDa | 0 | 0 | 2 | 3 | 5 | | | | | | | |
| AT5G65760 | | Serine carboxypeptidase S28 family protein | 59 kDa | 0 | 0 | 2 | 3 | 5 | | | | | | | |
| AT5G66470 | | RNA binding;GTP binding | 49 kDa | 0 | 0 | 2 | 3 | 5 | | | | | | | |
| AT5G66680 | | dolichyl-diphosphooligosaccharide-protein glycosyltra | 49 kDa | 0 | 0 | 2 | 3 | 5 | | 0.19 | 0.19 | | | | |
| AT1G01710 | | Acyl-CoA thioesterase family protein | 48 kDa | 0 | 0 | 2 | 2 | 4 | | | | | | | |
| AT1G04900 | | Protein of unknown function (DUF185) | 51 kDa | 0 | 0 | 2 | 2 | 4 | | | | | | | |
| AT1G07740 | | Tetratricopeptide repeat (TPR)-like superfamily protein | 52 kDa | 0 | 0 | 2 | 2 | 4 | | | | | | | |
| AT1G14610 | | valyl-tRNA synthetase / valine--tRNA ligase (VALRS) | 126 kDa | 0 | 0 | 2 | 2 | 4 | | | | | | | |
| AT1G18270 | AT1G18270.4;AT1G18270.1;AT1G18270.2;AT1G18270.3 | ketose-bisphosphate aldolase class-II family protein | 147 kDa | 0 | 0 | 2 | 2 | 4 | | | | 6.1E+05 | 7.3E+05 | 0.0E+00 | 1.3E+06 |
| AT1G22050 | | membrane-anchored ubiquitin-fold protein 6 precurso | 14 kDa | 0 | 0 | 2 | 2 | 4 | | | | | | | |
| AT1G31230 | | aspartate kinase-homoserine dehydrogenase i | 99 kDa | 0 | 0 | 2 | 2 | 4 | | 0.24 | 0.24 | | | | |
| AT1G32790 | | CTC-interacting domain 11 | 39 kDa | 0 | 0 | 2 | 2 | 4 | | | | | | | |
| AT1G36280 | | L-Aspartase-like family protein | 59 kDa | 0 | 0 | 2 | 2 | 4 | | | | | | | |
| AT1G42540 | | glutamate receptor 3.3 | 105 kDa | 0 | 0 | 2 | 2 | 4 | | | | | | | |
| AT1G48380 | | root hair initiation protein root hairless 1 (RHL1) | 39 kDa | 0 | 0 | 2 | 2 | 4 | | | | | | | |
| AT1G50030 | | target of rapamycin | 279 kDa | 0 | 0 | 2 | 2 | 4 | | | | | | | |
| AT1G50510 | AT1G50510.1 | indigoidine synthase A family protein | 35 kDa | 0 | 0 | 2 | 2 | 4 | | | | 4.4E+05 | 6.0E+05 | 0.0E+00 | 1.0E+06 |
| AT1G55930 | | CBS domain-containing protein / transporter associate | 73 kDa | 0 | 0 | 2 | 2 | 4 | | | | | | | |
| AT1G60170 | AT1G60170.1 | pre-mRNA processing ribonucleoprotein binding regio | ? | 0 | 0 | 2 | 2 | 4 | | | | 3.9E+05 | 2.1E+05 | 0.0E+00 | 6.0E+05 |
| AT1G60860 | | ARF-GAP domain 2 | 88 kDa | 0 | 0 | 2 | 2 | 4 | | | | | | | |
| AT1G63680 | | acid-amino acid ligases;ligases;ATP binding;ATP bindin | 86 kDa | 0 | 0 | 2 | 2 | 4 | | | | | | | |
| AT1G65150 | | TRAF-like family protein | 34 kDa | 0 | 0 | 2 | 2 | 4 | | | | | | | |
| AT1G66530 | | Arginyl-tRNA synthetase, class Ic | 66 kDa | 0 | 0 | 2 | 2 | 4 | | | | | | | |
| AT1G69420 | | DHHC-type zinc finger family protein | 66 kDa | 0 | 0 | 2 | 2 | 4 | | | | | | | |
| AT1G69700 | | HVA22 homologue C | 22 kDa | 3 | 0 | 2 | 2 | 7 | | | | | | | |
| AT1G71270 | | Vps52 / Sac2 family | 81 kDa | 0 | 0 | 2 | 2 | 4 | | | | | | | |
| AT1G75640 | | Leucine-rich receptor-like protein kinase family protei | 124 kDa | 0 | 0 | 2 | 2 | 4 | | | | | | | |
| AT1G79050 | | recA DNA recombination family protein | 48 kDa | 0 | 0 | 2 | 2 | 4 | | | | | | | |
| AT2G04030 | | Chaperone protein htpG family protein | 89 kDa | 0 | 0 | 2 | 2 | 4 | | | | | | | |
| AT2G20770 | | GCR2-like 2 | 44 kDa | 0 | 0 | 2 | 2 | 4 | | | | | | | |
| AT2G24180 | | cytochrome p450 71b6 | 57 kDa | 0 | 0 | 2 | 2 | 4 | | | | | | | |
| AT2G24820 | | translocon at the inner envelope membrane of chlorop | 61 kDa | 0 | 0 | 2 | 2 | 4 | | 0.38 | 0.38 | | | | |
| AT2G30270 | | Protein of unknown function (DUF567) | 20 kDa | 0 | 0 | 2 | 2 | 4 | | | | | | | |
| AT2G31660 | AT2G31660.1 | ARM repeat superfamily protein | 119 kDa | 0 | 0 | 2 | 2 | 4 | | | | | | | |
| AT2G32600 | AT2G32600.1 | hydroxyproline-rich glycoprotein family protein | 31 kDa | 0 | 0 | 2 | 2 | 4 | | | | 3.9E+06 | 1.9E+06 | 1.1E+06 | 6.9E+06 |
| AT2G34570 | AT2G34570.1 | PIN domain-like family protein | 32 kDa | 0 | 0 | 2 | 2 | 4 | | | | | | | |
| AT2G37690 | | phosphoribosylaminoimidazole carboxylase, putative , | 70 kDa | 0 | 0 | 2 | 2 | 4 | | | | | | | |
| AT2G38730 | AT2G38730.1 | Cyclophilin-like peptidyl-prolyl cis-trans isomerase fam | 21 kDa | 0 | 0 | 2 | 2 | 4 | | | | 1.1E+06 | 4.2E+06 | 1.9E+06 | 7.2E+06 |
| AT2G38960 | | endoplasmic reticulum oxidoreductins 2 | 53 kDa | 0 | 0 | 2 | 2 | 4 | | | | | | | |
| AT2G41190 | | Transmembrane amino acid transporter family protein | 59 kDa | 0 | 0 | 2 | 2 | 4 | | | | | | | |
| AT2G43430 | | glyoxalase 2-1 | 35 kDa | 0 | 0 | 2 | 2 | 4 | | | | | | | |
| AT2G46060 | | transmembrane protein-related | 90 kDa | 0 | 0 | 2 | 2 | 4 | | | | | | | |
| AT3G01650 | | RING domain ligase1 | 53 kDa | 0 | 0 | 2 | 2 | 4 | | | | | | | |
| AT3G03110 | | exportin 1B | 123 kDa | 0 | 0 | 2 | 2 | 4 | | | | | | | |
| AT3G05220 | AT3G05220.2;AT3G05220.1 | Heavy metal transport/detoxification superfamily prot | 59 kDa | 0 | 0 | 2 | 2 | 4 | | | | 6.0E+05 | 4.8E+05 | 0.0E+00 | 1.1E+06 |
| AT3G06670 | | binding | 100 kDa | 0 | 0 | 2 | 2 | 4 | | | | | | | |
| AT3G12760 | AT3G12760.1 | CONTAINS InterPro DOMAIN/s: Defective-in-cullin nedc | 29 kDa | 0 | 1 | 2 | 2 | 5 | | | | 1.0E+06 | 0.0E+00 | 1.6E+05 | 1.2E+06 |
| AT3G14067 | | Subtilase family protein | 82 kDa | 0 | 0 | 2 | 2 | 4 | | | | | | | |
| AT3G14940 | | phosphoenolpyruvate carboxylase 3 | 110 kDa | 0 | 0 | 2 | 2 | 4 | | | | | | | |
| AT3G24490 | | Alcohol dehydrogenase transcription factor Myb/SANT | 39 kDa | 0 | 1 | 2 | 2 | 5 | | | | | | | |
| AT3G26730 | | RING/U-box superfamily protein | 85 kDa | 0 | 0 | 2 | 2 | 4 | | | | | | | |
| AT3G27570 | | Sucrase/ferredoxin-like family protein | 42 kDa | 0 | 0 | 2 | 2 | 4 | | | | | | | |
| AT3G27770 | | unknown protein; FUNCTIONS IN: molecular_function | 37 kDa | 0 | 0 | 2 | 2 | 4 | | | | | | | |
| AT3G28480 | | Oxoglutarate/iron-dependent oxygenase | 36 kDa | 0 | 0 | 2 | 2 | 4 | | | | | | | |
| AT3G29400 | | exocyst subunit exo70 family protein E1 | 75 kDa | 0 | 0 | 2 | 2 | 4 | | | | | | | |
| AT3G44300 | | nitrilase 2 | 37 kDa | 0 | 0 | 2 | 2 | 4 | | | | | | | |
| AT3G45620 | | Transducin/WD40 repeat-like superfamily protein | 58 kDa | 0 | 0 | 2 | 2 | 4 | | | | | | | |
| AT3G49590 | | Autophagy-related protein 13 | 67 kDa | 0 | 0 | 2 | 2 | 4 | | | | | | | |
| AT3G49720 | | unknown protein; FUNCTIONS IN: molecular_function | 29 kDa | 0 | 1 | 2 | 2 | 5 | | | | | | | |
| AT3G54190 | | Transducin/WD40 repeat-like superfamily protein | 53 kDa | 0 | 0 | 2 | 2 | 4 | | | | | | | |
| AT3G58140 | AT3G58140.1 | phenylalanyl-tRNA synthetase class IIc family protein | 49 kDa | 0 | 0 | 2 | 2 | 4 | | | | 1.5E+06 | 1.2E+06 | 0.0E+00 | 2.6E+06 |
| AT3G60320 | | Protein of unknown function (DUF630 and DUF632) | 90 kDa | 0 | 0 | 2 | 2 | 4 | | | | | | | |
| AT3G61310 | | AT hook motif DNA-binding family protein | 37 kDa | 0 | 0 | 2 | 2 | 4 | | | | | | | |
| AT4G08170 | | Inositol 1,3,4-trisphosphate 5/6-kinase family protein | 40 kDa | 0 | 0 | 2 | 2 | 4 | | | | | | | |
| AT4G11260 | AT4G11260.1 | phosphatase-related | 40 kDa | 0 | 0 | 2 | 2 | 4 | | | | | | | |
| AT4G11860 | | Protein of unknown function (DUF544) | 74 kDa | 0 | 0 | 2 | 2 | 4 | | | | | | | |
| AT4G16760 | AT4G16760.2;AT4G16760.1 | acyl-CoA oxidase 1 | 74 kDa | 0 | 0 | 2 | 2 | 4 | | | | 4.7E+05 | 0.0E+00 | 0.0E+00 | 4.7E+05 |
| AT4G21790 | | tobamovirus multiplication 1 | ? | 0 | 0 | 2 | 2 | 4 | | | | | | | |
| AT4G24690 | | ubiquitin-associated (UBA)/TS-N domain-containing pr | 76 kDa | 0 | 0 | 2 | 2 | 4 | | | | | | | |
| AT4G30950 | | fatty acid desaturase 6 | 51 kDa | 0 | 0 | 2 | 2 | 4 | | | | | | | |
| AT4G31010 | | RNA-binding CRS1 / YhbY (CRM) domain-containing prc | 46 kDa | 0 | 0 | 2 | 2 | 4 | | | | | | | |
| AT4G31300 | AT4G31300.3;AT4G31300.1;AT4G31300.2 | N-terminal nucleophile aminohydrolases (Ntn hydrolas | 25 kDa | 0 | 0 | 2 | 2 | 4 | | 0.12 | 0.12 | 4.0E+06 | 5.5E+06 | 0.0E+00 | 9.5E+06 |
| AT4G32050 | | neurochondrin family protein | 68 kDa | 0 | 0 | 2 | 2 | 4 | | | | | | | |
| AT4G32470 | | Cytochrome bd ubiquinol oxidase, 14kDa subunit | 15 kDa | 0 | 0 | 2 | 2 | 4 | 0.05 | 0.05 | 0.05 | | | | |
| AT4G33250 | | eukaryotic translation initiation factor 3K | 26 kDa | 0 | 0 | 2 | 2 | 4 | | | | | | | |
| AT4G34240 | | aldehyde dehydrogenase 3I1 | 60 kDa | 0 | 0 | 2 | 2 | 4 | | | | | | | |
| AT4G39730 | | Lipase/lipooxygenase, PLAT/LH2 family protein | 20 kDa | 0 | 0 | 2 | 2 | 4 | | | | | | | |
| AT5G03740 | AT5G03740.1 | histone deacetylase 2C | 32 kDa | 0 | 0 | 2 | 2 | 4 | | | | 4.5E+06 | 7.1E+06 | 0.0E+00 | 1.2E+07 |
| AT5G09960 | AT5G09960.3;AT5G09960.1 | FUNCTIONS IN: molecular_function unknown; INVOLVE | ? | 0 | 0 | 2 | 2 | 4 | | | | 6.4E+06 | 1.7E+06 | 0.0E+00 | 8.1E+06 |
| AT5G13420 | | Aldolase-type TIM barrel family protein | 48 kDa | 0 | 0 | 2 | 2 | 4 | | | | | | | |
| AT5G16310 | | Peptidase C12, ubiquitin carboxyl-terminal hydrolase : | 39 kDa | 0 | 0 | 2 | 2 | 4 | | | | | | | |
| AT5G16420 | | Pentatricopeptide repeat (PPR-like) superfamily protei | 61 kDa | 0 | 0 | 2 | 2 | 4 | | | | | | | |
| AT5G19940 | | Plastid-lipid associated protein PAP / fibrillin family pr | 26 kDa | 0 | 2 | 2 | 2 | 6 | | | | | | | |

| | | | | | | | | | | | | | | | |
|---|---|---|---|---|---|---|---|---|---|---|---|---|---|---|---|
| AT5G22350 | | Protein of unknown function (DUF1022) | 48 kDa | 0 | 0 | 2 | 2 | 4 | | | | | | | |
| AT5G23450 | | long-chain base (LCB) kinase 1 | 85 kDa | 0 | 0 | 2 | 2 | 4 | | | | | | | |
| AT5G25050 | | Major facilitator superfamily protein | 55 kDa | 0 | 0 | 2 | 2 | 4 | | | | | | | |
| AT5G27730 | | Protein of unknown function (DUF1624) | 54 kDa | 0 | 0 | 2 | 2 | 4 | | | | | | | |
| AT5G27840 | | Calcineurin-like metallo-phosphoesterase superfamily | 37 kDa | 0 | 1 | 2 | 2 | 5 | | | | | | | |
| AT5G38720 | | unknown protein; Has 13419 Blast hits to 9093 protei | 35 kDa | 0 | 0 | 2 | 2 | 4 | | | | | | | |
| AT5G45550 | | Mob1/phocein family protein | 25 kDa | 0 | 1 | 2 | 2 | 5 | | | | | | | |
| AT5G47435 | | formyltetrahydrofolate deformylase, putative | 36 kDa | 0 | 0 | 2 | 2 | 4 | | | | | | | |
| AT5G48460 | | Actin binding Calponin homology (CH) domain-contair | 74 kDa | 0 | 0 | 2 | 2 | 4 | | | | | | | |
| AT5G61140 | | U5 small nuclear ribonucleoprotein helicase | 243 kDa | 0 | 0 | 2 | 2 | 4 | | 0.26 | 0.26 | | | | |
| AT5G63400 | | adenylate kinase 1 | 27 kDa | 2 | 0 | 2 | 2 | 6 | 0.17 | 0.21 | 0.19 | | | | |
| AT5G63620 | AT5G63620.1;AT5G63620.2;AT5G63620.3 | GroES-like zinc-binding alcohol dehydrogenase family | 45 kDa | 0 | 0 | 2 | 2 | 4 | | | | 1.4E+06 | 2.1E+06 | 0.0E+00 | 3.6E+06 |
| AT1G01730 | | unknown protein; FUNCTIONS IN: molecular_function | 24 kDa | 0 | 0 | 2 | 1 | 3 | | | | | | | |
| AT1G07230 | | non-specific phospholipase C1 | 60 kDa | 0 | 0 | 2 | 1 | 3 | | | | | | | |
| AT1G12470 | AT1G12470.1;AT1G18370.1 | zinc ion binding | 112 kDa | 0 | 0 | 2 | 1 | 3 | | | | 2.8E+05 | 7.3E+06 | 0.0E+00 | 7.5E+06 |
| AT1G13060 | AT1G13060.1;AT1G13060.3;AT1G13060.2;AT3G26340.1 | 20S proteasome beta subunit E1 | 30 kDa | 1 | 0 | 2 | 1 | 4 | | | | 3.9E+06 | 4.7E+06 | 0.0E+00 | 8.5E+06 |
| AT1G15860 | | Domain of unknown function (DUF298) | 27 kDa | 0 | 0 | 2 | 1 | 3 | | | | | | | |
| AT1G17860 | | Kunitz family trypsin and protease inhibitor protein | 22 kDa | 0 | 0 | 2 | 1 | 3 | | | | | | | |
| AT1G23010 | | Cupredoxin superfamily protein | 66 kDa | 0 | 0 | 2 | 1 | 3 | | | | | | | |
| AT1G24120 | | ARG1-like 1 | 49 kDa | 0 | 0 | 2 | 1 | 3 | | | | | | | |
| AT1G30825 | | Arp2/3 complex, 34 kD subunit p34-Arc | 36 kDa | 0 | 0 | 2 | 1 | 3 | | | | | | | |
| AT1G33390 | AT1G33390.2;AT1G33390.1 | RNA helicase family protein | 139 kDa | 0 | 0 | 2 | 1 | 3 | | | | 1.4E+05 | 1.1E+05 | 0.0E+00 | 2.5E+05 |
| AT1G48970 | | NagB/RpiA/CoA transferase-like superfamily protein | 73 kDa | 0 | 0 | 2 | 1 | 3 | | | | | | | |
| AT1G50710 | | unknown protein; Has 109 Blast hits to 109 proteins in | 48 kDa | 0 | 0 | 2 | 1 | 3 | | | | | | | |
| AT1G55520 | | TATA binding protein 2 | 22 kDa | 0 | 0 | 2 | 1 | 3 | | | | | | | |
| AT1G59910 | | Actin-binding FH2 (formin homology 2) family protein | 99 kDa | 0 | 0 | 2 | 1 | 3 | | | | | | | |
| AT1G71400 | | receptor like protein 12 | 95 kDa | 0 | 0 | 2 | 1 | 3 | | | | | | | |
| AT1G74640 | AT1G74640.1 | alpha/beta-Hydrolases superfamily protein | 41 kDa | 0 | 1 | 2 | 1 | 4 | | | | 5.1E+05 | 5.7E+05 | 0.0E+00 | 1.1E+06 |
| AT1G80270 | | PENTATRICOPEPTIDE REPEAT 596 | 67 kDa | 0 | 0 | 2 | 1 | 3 | | | | | | | |
| AT2G03350 | | Protein of unknown function, DUF538 | 20 kDa | 0 | 0 | 2 | 1 | 3 | | | | | | | |
| AT2G21440 | | RNA-binding (RRM/RBD/RNP motifs) family protein | 112 kDa | 0 | 0 | 2 | 1 | 3 | | | | | | | |
| AT2G22230 | | Thioesterase superfamily protein | 24 kDa | 0 | 0 | 2 | 1 | 3 | | | | | | | |
| AT2G28070 | | ABC-2 type transporter family protein | 81 kDa | 0 | 0 | 2 | 1 | 3 | | | | | | | |
| AT2G32550 | | Cell differentiation, Rcd1-like protein | 37 kDa | 0 | 2 | 2 | 1 | 5 | | | | | | | |
| AT2G35690 | | acyl-CoA oxidase 5 | 74 kDa | 0 | 0 | 2 | 1 | 3 | | | | | | | |
| AT2G46450 | | cyclic nucleotide-gated channel 12 | 72 kDa | 0 | 0 | 2 | 1 | 3 | | | | | | | |
| AT3G05990 | | Leucine-rich repeat (LRR) family protein | 56 kDa | 0 | 0 | 2 | 1 | 3 | | | | | | | |
| AT3G07220 | | SMAD/FHA domain-containing protein | 36 kDa | 0 | 0 | 2 | 1 | 3 | | | | | | | |
| AT3G07750 | | 3'-5'-exoribonuclease family protein | 30 kDa | 0 | 0 | 2 | 1 | 3 | | | | | | | |
| AT3G12080 | | GTP-binding family protein | 73 kDa | 0 | 0 | 2 | 1 | 3 | | | | | | | |
| AT3G20230 | | Ribosomal L18p/L5e family protein | 21 kDa | 0 | 0 | 2 | 1 | 3 | | | | | | | |
| AT3G20500 | | purple acid phosphatase 18 | 50 kDa | 0 | 0 | 2 | 1 | 3 | | | | | | | |
| AT3G20790 | | NAD(P)-binding Rossmann-fold superfamily protein | 39 kDa | 0 | 0 | 2 | 1 | 3 | | | | | | | |
| AT3G24170 | AT3G24170.3;AT3G24170.2;AT3G24170.1 | glutathione-disulfide reductase | 54 kDa | 1 | 0 | 2 | 1 | 4 | | | | 5.9E+05 | 3.2E+06 | 0.0E+00 | 3.7E+06 |
| AT3G26410 | | methyltransferases;nucleic acid binding | 55 kDa | 0 | 0 | 2 | 1 | 3 | | | | | | | |
| AT3G26600 | | armadillo repeat only 4 | 68 kDa | 0 | 0 | 2 | 1 | 3 | | | | | | | |
| AT3G28200 | | Peroxidase superfamily protein | 35 kDa | 0 | 0 | 2 | 1 | 3 | | | | | | | |
| AT3G44050 | | P-loop containing nucleoside triphosphate hydrolases | 143 kDa | 0 | 0 | 2 | 1 | 3 | | | | | | | |
| AT3G44890 | | ribosomal protein L9 | 22 kDa | 0 | 0 | 2 | 1 | 3 | | 0.13 | 0.13 | | | | |
| AT3G46600 | | GRAS family transcription factor | 67 kDa | 0 | 0 | 2 | 1 | 3 | | | | | | | |
| AT3G47390 | | cytidine/deoxycytidylate deaminase family protein | 66 kDa | 0 | 0 | 2 | 1 | 3 | | | | | | | |
| AT3G49660 | | Transducin/WD40 repeat-like superfamily protein | 35 kDa | 0 | 0 | 2 | 1 | 3 | | | | | | | |
| AT3G55020 | AT3G55020.2;AT3G55020.3;AT3G55020.1 | Ypt/Rab-GAP domain of gyp1p superfamily protein | 87 kDa | 3 | 0 | 2 | 1 | 6 | | | | | | | |
| AT3G56370 | | Leucine-rich repeat protein kinase family protein | 104 kDa | 0 | 0 | 2 | 1 | 3 | | | | | | | |
| AT3G58520 | | Ubiquitin carboxyl-terminal hydrolase family protein | 49 kDa | 0 | 0 | 2 | 1 | 3 | | | | | | | |
| AT3G61450 | | syntaxin of plants 73 | 30 kDa | 0 | 0 | 2 | 1 | 3 | | | | | | | |
| AT3G61710 | | AUTOPHAGY 6 | 59 kDa | 0 | 0 | 2 | 1 | 3 | | | | | | | |
| AT3G62060 | | Pectinacetylesterase family protein | 46 kDa | 0 | 0 | 2 | 1 | 3 | | | | | | | |
| AT4G01150 | | unknown protein; FUNCTIONS IN: molecular_function | ? | 0 | 0 | 2 | 1 | 3 | | | | | | | |
| AT4G02500 | | UDP-xylosyltransferase 2 | 53 kDa | 0 | 0 | 2 | 1 | 3 | | | | | | | |
| AT4G04610 | | APS reductase 1 | 52 kDa | 0 | 0 | 2 | 1 | 3 | | | | | | | |
| AT4G09080 | | Outer membrane OMP85 family protein | 44 kDa | 0 | 0 | 2 | 1 | 3 | | | | | | | |
| AT4G13360 | | ATP-dependent caseinolytic (Clp) protease/crotonase fa | 46 kDa | 0 | 0 | 2 | 1 | 3 | | | | | | | |
| AT4G16210 | | enoyl-CoA hydratase/isomerase A | 29 kDa | 0 | 0 | 2 | 1 | 3 | | | | | | | |
| AT4G16500 | | Cystatin/monellin superfamily protein | 13 kDa | 0 | 0 | 2 | 1 | 3 | | | | | | | |
| AT4G18260 | | Cytochrome b561/ferric reductase transmembrane pro | 62 kDa | 0 | 0 | 2 | 1 | 3 | | | | | | | |
| AT4G22850 | | SNARE associated Golgi protein family | 33 kDa | 0 | 0 | 2 | 1 | 3 | | | | | | | |
| AT4G22980 | | FUNCTIONS IN: molecular_function unknown; INVOLVE | 64 kDa | 0 | 0 | 2 | 1 | 3 | | | | | | | |
| AT4G23620 | | Ribosomal protein L25/Gln-tRNA synthetase, anti-codo | 29 kDa | 0 | 0 | 2 | 1 | 3 | | | | | | | |
| AT4G35730 | | Regulator of Vps4 activity in the MVB pathway protein | 52 kDa | 0 | 0 | 2 | 1 | 3 | | | | | | | |
| AT5G06950 | | bZIP transcription factor family protein | 37 kDa | 0 | 0 | 2 | 1 | 3 | | | | | | | |
| AT5G10050 | | NAD(P)-binding Rossmann-fold superfamily protein | 31 kDa | 0 | 0 | 2 | 1 | 3 | | | | | | | |
| AT5G12350 | | Regulator of chromosome condensation (RCC1) family | 118 kDa | 0 | 0 | 2 | 1 | 3 | | | | | | | |
| AT5G13450 | | delta subunit of Mt ATP synthase | 26 kDa | 0 | 0 | 2 | 1 | 3 | 0.04 | 0.14 | 0.09 | | | | |
| AT5G16715 | | ATP binding;valine-tRNA ligases;aminoacyl-tRNA ligase | 111 kDa | 0 | 3 | 2 | 1 | 6 | | | | | | | |
| AT5G18200 | | UTP:galactose-1-phosphate uridylyltransferases;ribose- | 39 kDa | 0 | 0 | 2 | 1 | 3 | | | | | | | |
| AT5G23310 | | Fe superoxide dismutase 3 | 30 kDa | 0 | 0 | 2 | 1 | 3 | | | | | | | |
| AT5G24310 | | ABL interactor-like protein 3 | 37 kDa | 0 | 0 | 2 | 1 | 3 | | | | | | | |
| AT5G24680 | | Peptidase C78, ubiquitin fold modifier-specific peptida | 44 kDa | 0 | 0 | 2 | 1 | 3 | | | | | | | |
| AT5G26670 | | Pectinacetylesterase family protein | 47 kDa | 0 | 1 | 2 | 1 | 4 | | | | | | | |
| AT5G37010 | | unknown protein; FUNCTIONS IN: molecular_function | ? | 0 | 0 | 2 | 1 | 3 | | | | | | | |
| AT5G41270 | AT5G41270.1 | CONTAINS InterPro DOMAIN/s: RNAse P, Rpr2/Rpp21 su | 29 kDa | 0 | 0 | 2 | 1 | 3 | | | | 0.0E+00 | 0.0E+00 | 2.7E+05 | 2.7E+05 |
| AT5G50340 | | ATP-dependent peptidases;nucleotide binding;serine-t | 63 kDa | 0 | 0 | 2 | 1 | 3 | | | | | | | |
| AT5G50870 | | ubiquitin-conjugating enzyme 27 | 23 kDa | 0 | 0 | 2 | 1 | 3 | | | | | | | |
| AT5G51350 | | Leucine-rich repeat transmembrane protein kinase fam | 99 kDa | 0 | 0 | 2 | 1 | 3 | | | | | | | |
| AT5G52580 | | RabGAP/TBC domain-containing protein | 78 kDa | 0 | 0 | 2 | 1 | 3 | | | | | | | |
| AT5G56650 | | IAA-leucine resistant (ILR)-like 1 | 48 kDa | 0 | 0 | 2 | 1 | 3 | | | | | | | |
| AT5G58270 | | ABC transporter of the mitochondrion 3 | 80 kDa | 0 | 0 | 2 | 1 | 3 | | | | | | | |
| AT5G58490 | | NAD(P)-binding Rossmann-fold superfamily protein | 36 kDa | 0 | 1 | 2 | 1 | 4 | | | | | | | |
| AT5G58760 | | damaged DNA binding 2 | 63 kDa | 0 | 0 | 2 | 1 | 3 | | | | | | | |
| AT5G59180 | | DNA-directed RNA polymerase II | 19 kDa | 0 | 0 | 2 | 1 | 3 | | | | | | | |
| ATCG00130 | ATCG00130.1 | ATPase, F0 complex, subunit B/B', bacterial/chloroplas | 21 kDa | 0 | 0 | 2 | 1 | 3 | | | | 1.6E+06 | 2.0E+06 | 0.0E+00 | 3.6E+06 |
| AT1G02560 | | nuclear encoded CLP protease 5 | 32 kDa | 0 | 2 | 2 | 0 | 4 | | | | | | | |
| AT1G15080 | | lipid phosphate phosphatase 2 | 33 kDa | 0 | 1 | 2 | 0 | 3 | | | | | | | |
| AT3G23400 | | Plastid-lipid associated protein PAP / fibrillin family pro | 30 kDa | 0 | 3 | 2 | 0 | 5 | | 0.21 | 0.21 | | | | |
| AT1G19870 | | IQ-domain 32 | 87 kDa | 0 | 0 | 1 | 20 | 21 | | | | | | | |
| AT1G70710 | AT1G70710.1 | glycosyl hydrolase 9B1 | 55 kDa | 0 | 0 | 1 | 13 | 14 | | | | | | | |
| AT4G27720 | | Major facilitator superfamily protein | 51 kDa | 0 | 0 | 1 | 11 | 12 | | | | | | | |

| | | | | | | | | | | | | | | | |
|---|---|---|---|---|---|---|---|---|---|---|---|---|---|---|---|
| AT3G09470 | | Major facilitator superfamily protein | 50 kDa | 0 | 0 | 1 | 10 | 11 | | | | | | | |
| AT5G11560 | | catalytics | 109 kDa | 0 | 0 | 1 | 10 | 11 | | | | | | | |
| AT3G19420 | | PTEN 2 | 66 kDa | 0 | 0 | 1 | 8 | 9 | | | | | | | |
| AT1G15950 | | cinnamoyl coa reductase 1 | 37 kDa | 0 | 0 | 1 | 7 | 8 | | | | | | | |
| AT2G17420 | | NADPH-dependent thioredoxin reductase A | 40 kDa | 0 | 0 | 1 | 7 | 8 | | | | | | | |
| AT5G14430 | | S-adenosyl-L-methionine-dependent methyltransferase | 70 kDa | 0 | 0 | 1 | 7 | 8 | | | | | | | |
| AT5G16720 | | Protein of unknown function, DUF593 | 77 kDa | 0 | 0 | 1 | 7 | 8 | | | | | | | |
| AT5G20840 | | Phosphoinositide phosphatase family protein | 94 kDa | 0 | 0 | 1 | 7 | 8 | | | | | | | |
| AT1G13980 | | sec7 domain-containing protein | 163 kDa | 0 | 0 | 1 | 6 | 7 | | | | | | | |
| AT1G22750 | | unknown protein; FUNCTIONS IN: molecular_function | 27 kDa | 0 | 0 | 1 | 6 | 7 | | | | | | | |
| AT2G21050 | | like AUXIN RESISTANT 2 | 55 kDa | 0 | 0 | 1 | 6 | 7 | | | | | | | |
| AT3G09300 | | OSBP(oxysterol binding protein)-related protein 3B | 52 kDa | 0 | 0 | 1 | 6 | 7 | | | | | | | |
| AT5G51550 | | EXORDIUM like 3 | 37 kDa | 0 | 0 | 1 | 6 | 7 | | | | | | | |
| AT1G15170 | | MATE efflux family protein | 52 kDa | 0 | 0 | 1 | 5 | 6 | | | | | | | |
| AT1G16460 | | rhodanese homologue 2 | 35 kDa | 0 | 0 | 1 | 5 | 6 | | | | | | | |
| AT1G32330 | | heat shock transcription factor A1D | 55 kDa | 0 | 0 | 1 | 5 | 6 | | | | | | | |
| AT1G48270 | | G-protein-coupled receptor 1 | 37 kDa | 0 | 0 | 1 | 5 | 6 | | | | | | | |
| AT1G60740 | | Thioredoxin superfamily protein | 17 kDa | 1 | 0 | 1 | 5 | 7 | | | | | | | |
| AT1G73030 | | SNF7 family protein | 23 kDa | 0 | 0 | 1 | 5 | 6 | | | | | | | |
| AT2G24190 | | NAD(P)-binding Rossmann-fold superfamily protein | 33 kDa | 2 | 2 | 1 | 5 | 10 | | | | | | | |
| AT2G35040 | AT2G35040.2;AT2G35040.1 | AICARFT/IMPCHase bienzyme family protein | 59 kDa | 0 | 0 | 1 | 5 | 6 | | | | 8.9E+06 | 1.3E+07 | 0.0E+00 | 2.2E+07 |
| AT2G37400 | | Tetratricopeptide repeat (TPR)-like superfamily protein | 38 kDa | 0 | 0 | 1 | 5 | 6 | | | | | | | |
| AT2G47470 | | thioredoxin family protein | 39 kDa | 0 | 0 | 1 | 5 | 6 | | | | | | | |
| AT3G30841 | | Cofactor-independent phosphoglycerate mutase | 53 kDa | 0 | 0 | 1 | 5 | 6 | | | | | | | |
| AT3G48690 | AT3G48690.1 | alpha/beta-Hydrolases superfamily protein | 36 kDa | 0 | 0 | 1 | 5 | 6 | | | | 2.9E+06 | 1.9E+06 | 0.0E+00 | 4.8E+06 |
| AT5G08450 | AT5G08450.3;AT5G08450.2;AT5G08450.1 | CONTAINS InterPro DOMAIN/s: Histone deacetylation p | 104 kDa | 0 | 1 | 1 | 5 | 7 | | | | 1.1E+06 | 9.9E+05 | 0.0E+00 | 2.0E+06 |
| AT5G15120 | AT5G15120.1 | Protein of unknown function (DUF1637) | 33 kDa | 0 | 0 | 1 | 5 | 6 | | | | | | | |
| AT5G25440 | | Protein kinase superfamily protein | 35 kDa | 0 | 5 | 1 | 5 | 11 | | | | | | | |
| AT5G42740 | | Sugar isomerase (SIS) family protein | 62 kDa | 0 | 0 | 1 | 5 | 6 | | | | | | | |
| AT5G49160 | AT5G49160.2;AT5G49160.1 | methyltransferase 1 | 172 kDa | 0 | 0 | 1 | 5 | 6 | | 0.30 | 0.30 | 0.0E+00 | 1.2E+05 | 4.7E+05 | 6.0E+05 |
| AT5G52240 | AT5G52240.2;AT5G52240.1;AT3G48890.1 | membrane steroid binding protein 1 | 24 kDa | 0 | 0 | 1 | 5 | 6 | | | | 1.6E+06 | 3.1E+06 | 0.0E+00 | 4.8E+06 |
| AT5G67270 | | end binding protein 1C | 36 kDa | 0 | 0 | 1 | 5 | 6 | | | | | | | |
| AT1G13870 | | calmodulin binding;purine nucleotide binding | 34 kDa | 0 | 2 | 1 | 4 | 7 | | | | | | | |
| AT1G35620 | | PDI-like 5-2 | 50 kDa | 0 | 0 | 1 | 4 | 5 | | | | | | | |
| AT1G48370 | | YELLOW STRIPE like 8 | 80 kDa | 0 | 0 | 1 | 4 | 5 | | | | | | | |
| AT1G74920 | | aldehyde dehydrogenase 10A8 | 54 kDa | 0 | 0 | 1 | 4 | 5 | | | | | | | |
| AT2G17390 | | ankyrin repeat-containing 2B | 37 kDa | 0 | 0 | 1 | 4 | 5 | | | | | | | |
| AT2G17820 | | histidine kinase 1 | 135 kDa | 0 | 0 | 1 | 4 | 5 | | | | | | | |
| AT2G38170 | | cation exchanger 1 | 50 kDa | 0 | 0 | 1 | 4 | 5 | | | | | | | |
| AT2G45260 | | Plant protein of unknown function (DUF641) | 48 kDa | 0 | 0 | 1 | 4 | 5 | | | | | | | |
| AT3G08910 | | DNAJ heat shock family protein | 36 kDa | 0 | 0 | 1 | 4 | 5 | | | | | | | |
| AT3G13860 | AT3G13860.1 | heat shock protein 60-3A | 60 kDa | 0 | 0 | 1 | 4 | 5 | | | | 6.6E+05 | 6.5E+05 | 0.0E+00 | 1.3E+06 |
| AT3G20630 | | ubiquitin-specific protease 14 | 88 kDa | 0 | 0 | 1 | 4 | 5 | | | | | | | |
| AT3G24530 | | AAA-type ATPase family protein / ankyrin repeat family | 54 kDa | 0 | 0 | 1 | 4 | 5 | | | | | | | |
| AT3G56740 | | Ubiquitin-associated (UBA) protein | 32 kDa | 0 | 0 | 1 | 4 | 5 | | | | | | | |
| AT3G61140 | | 26S proteasome, regulatory subunit Rpn7;Proteasome | 51 kDa | 0 | 0 | 1 | 4 | 5 | | | | | | | |
| AT4G15550 | | indole-3-acetate beta-D-glucosyltransferase | 54 kDa | 0 | 0 | 1 | 4 | 5 | | | | | | | |
| AT4G32290 | | Core-2/I-branching beta-1,6-N-acetylglucosaminyltran | 44 kDa | 0 | 0 | 1 | 4 | 5 | | | | | | | |
| AT4G38370 | | Phosphoglycerate mutase family protein | 25 kDa | 0 | 1 | 1 | 4 | 6 | | | | | | | |
| AT5G03160 | | homolog of mamallian P58IPK | 54 kDa | 0 | 0 | 1 | 4 | 5 | | | | | | | |
| AT5G10710 | | INVOLVED IN: chromosome segregation, cell division; L | 36 kDa | 0 | 0 | 1 | 4 | 5 | | | | | | | |
| AT5G13030 | | unknown protein; FUNCTIONS IN: molecular_function | 71 kDa | 0 | 0 | 1 | 4 | 5 | | | | | | | |
| AT5G13630 | | magnesium-chelatase subunit chlH, chloroplast, putati | 154 kDa | 0 | 0 | 1 | 4 | 5 | | | | | | | |
| AT5G23530 | | carboxyesterase 18 | 38 kDa | 0 | 0 | 1 | 4 | 5 | | | | | | | |
| AT5G47390 | | myb-like transcription factor family protein | 40 kDa | 0 | 2 | 1 | 4 | 7 | | | | | | | |
| AT1G01800 | | NAD(P)-binding Rossmann-fold superfamily protein | 32 kDa | 0 | 0 | 1 | 3 | 4 | | | | | | | |
| AT1G05350 | | NAD(P)-binding Rossmann-fold superfamily protein | 47 kDa | 0 | 0 | 1 | 3 | 4 | | | | | | | |
| AT1G10020 | | Protein of unknown function (DUF1005) | 49 kDa | 0 | 0 | 1 | 3 | 4 | | | | | | | |
| AT1G12760 | | Zinc finger, C3HC4 type (RING finger) family protein | 45 kDa | 0 | 0 | 1 | 3 | 4 | | | | | | | |
| AT1G19835 | | Plant protein of unknown function (DUF869) | 110 kDa | 0 | 0 | 1 | 3 | 4 | | | | | | | |
| AT1G20110 | | RING/FYVE/PHD zinc finger superfamily protein | 65 kDa | 0 | 0 | 1 | 3 | 4 | | | | | | | |
| AT1G20840 | | tonoplast monosaccharide transporter1 | 79 kDa | 0 | 0 | 1 | 3 | 4 | | | | | | | |
| AT1G50380 | | Prolyl oligopeptidase family protein | 81 kDa | 0 | 0 | 1 | 3 | 4 | | | | | | | |
| AT1G50500 | | Membrane trafficking VPS53 family protein | 95 kDa | 0 | 0 | 1 | 3 | 4 | | | | | | | |
| AT1G53880 | | Eukaryotic translation initiation factor 2B (eIF-2B) fami | 65 kDa | 0 | 0 | 1 | 3 | 4 | | | | | | | |
| AT1G54060 | | 6B-interacting protein 1-like 1 | 42 kDa | 0 | 0 | 1 | 3 | 4 | | | | | | | |
| AT1G54360 | | TBP-ASSOCIATED FACTOR 6B | 56 kDa | 0 | 0 | 1 | 3 | 4 | | | | | | | |
| AT1G56290 | | CwfJ-like family protein | 79 kDa | 0 | 0 | 1 | 3 | 4 | | | | | | | |
| AT1G58060 | | RNA helicase family protein | 164 kDa | 0 | 0 | 1 | 3 | 4 | | | | | | | |
| AT1G58400 | | Disease resistance protein (CC-NBS-LRR class) family | 105 kDa | 0 | 0 | 1 | 3 | 4 | | | | | | | |
| AT1G74330 | | Protein kinase superfamily protein | 78 kDa | 0 | 0 | 1 | 3 | 4 | | | | | | | |
| AT1G80910 | | Protein of unknown function (DUF1712) | 55 kDa | 0 | 0 | 1 | 3 | 4 | | | | | | | |
| AT2G21160 | | Translocon-associated protein (TRAP), alpha subunit | 28 kDa | 0 | 1 | 1 | 3 | 5 | | 0.28 | 0.28 | | | | |
| AT2G23520 | | Pyridoxal phosphate (PLP)-dependent transferases supe | 100 kDa | 0 | 0 | 1 | 3 | 4 | | | | | | | |
| AT3G18524 | | MUTS homolog 2 | 106 kDa | 0 | 0 | 1 | 3 | 4 | 0.04 | | 0.04 | | | | |
| AT3G19460 | | Reticulon family protein | 22 kDa | 0 | 0 | 1 | 3 | 4 | | | | | | | |
| AT3G19670 | | pre-mRNA-processing protein 40B | 114 kDa | 0 | 0 | 1 | 3 | 4 | | | | | | | |
| AT3G22180 | | DHHC-type zinc finger family protein | 77 kDa | 0 | 0 | 1 | 3 | 4 | | | | | | | |
| AT3G26370 | | O-fucosyltransferase family protein | 63 kDa | 0 | 0 | 1 | 3 | 4 | | | | | | | |
| AT3G30390 | | Transmembrane amino acid transporter family protein | 50 kDa | 0 | 0 | 1 | 3 | 4 | | | | | | | |
| AT3G43600 | | aldehyde oxidase 2 | 145 kDa | 0 | 0 | 1 | 3 | 4 | | | | | | | |
| AT3G52050 | | 5'-3' exonuclease family protein | 47 kDa | 0 | 0 | 1 | 3 | 4 | | | | | | | |
| AT3G52060 | | Core-2/I-branching beta-1,6-N-acetylglucosaminyltran | 40 kDa | 0 | 0 | 1 | 3 | 4 | | | | | | | |
| AT3G54110 | | plant uncoupling mitochondrial protein 1 | 33 kDa | 0 | 0 | 1 | 3 | 4 | 0.12 | 0.14 | 0.13 | | | | |
| AT3G66658 | | aldehyde dehydrogenase 22A1 | 66 kDa | 0 | 0 | 1 | 3 | 4 | | | | | | | |
| AT4G10480 | | Nascent polypeptide-associated complex (NAC), alpha s | 23 kDa | 0 | 0 | 1 | 3 | 4 | | | | | | | |
| AT4G17420 | | Tryptophan RNA-binding attenuator protein-like | 31 kDa | 0 | 0 | 1 | 3 | 4 | | | | | | | |
| AT4G24490 | | RAB geranylgeranyl transferase alpha subunit 1 | 77 kDa | 0 | 0 | 1 | 3 | 4 | | | | | | | |
| AT4G31340 | | myosin heavy chain-related | 50 kDa | 0 | 0 | 1 | 3 | 4 | | | | | | | |
| AT4G31770 | | debranching enzyme 1 | 48 kDa | 0 | 0 | 1 | 3 | 4 | | | | | | | |
| AT4G35000 | | ascorbate peroxidase 3 | 32 kDa | 5 | 2 | 1 | 3 | 11 | 0.10 | | 0.10 | | | | |
| AT4G35630 | | phosphoserine aminotransferase | 47 kDa | 0 | 0 | 1 | 3 | 4 | | | | | | | |
| AT4G39230 | | NmrA-like negative transcriptional regulator family pro | 34 kDa | 0 | 0 | 1 | 3 | 4 | | | | | | | |
| AT5G10470 | | kinesin like protein for actin based chloroplast movem | 141 kDa | 0 | 1 | 1 | 3 | 5 | | | | | | | |
| AT5G11850 | | Protein kinase superfamily protein | 98 kDa | 0 | 0 | 1 | 3 | 4 | | | | | | | |
| AT5G12890 | | UDP-Glycosyltransferase superfamily protein | 55 kDa | 0 | 0 | 1 | 3 | 4 | | | | | | | |
| AT5G13020 | | Emsy N Terminus (ENT)/ plant Tudor-like domains-cont | 44 kDa | 0 | 0 | 1 | 3 | 4 | | | | | | | |
| AT5G14800 | AT5G14800.2;AT5G14800.1 | pyrroline-5- carboxylate (P5C) reductase | 29 kDa | 0 | 12 | 1 | 3 | 16 | | | | 9.8E+05 | 1.2E+06 | 0.0E+00 | 2.2E+06 |
| AT5G17770 | | NADH:cytochrome B5 reductase 1 | 31 kDa | 0 | 0 | 1 | 3 | 4 | 0.08 | | 0.08 | | | | |
| AT5G19070 | | SNARE associated Golgi protein family | 30 kDa | 0 | 1 | 1 | 3 | 5 | | | | | | | |

| | | | | | | | | | | | | | | | |
|---|---|---|---|---|---|---|---|---|---|---|---|---|---|---|---|
| AT5G22830 | | magnesium (Mg) transporter 10 | 51 kDa | 0 | 0 | 1 | 3 | 4 | | | | | | | |
| AT5G51180 | | alpha/beta-Hydrolases superfamily protein | 40 kDa | 0 | 0 | 1 | 3 | 4 | | | | | | | |
| AT5G51980 | | Transducin/WD40 repeat-like superfamily protein | 47 kDa | 0 | 0 | 1 | 3 | 4 | | | | | | | |
| AT5G58380 | | SOS3-interacting protein 1 | 55 kDa | 0 | 0 | 1 | 3 | 4 | | | | | | | |
| AT5G64680 | AT5G64680.1;AT5G64680.2;AT5G64680.3 | unknown protein; FUNCTIONS IN: molecular_function | 23 kDa | 0 | 0 | 1 | 3 | 4 | | | | 1.6E+06 | 3.4E+06 | 2.2E+05 | 5.2E+06 |
| AT5G66950 | | Pyridoxal phosphate (PLP)-dependent transferases supe | 97 kDa | 0 | 0 | 1 | 3 | 4 | | | | | | | |
| AT1G01630 | | Sec14p-like phosphatidylinositol transfer family protei | 29 kDa | 0 | 0 | 1 | 2 | 3 | | | | | | | |
| AT1G02170 | | metacaspase 1 | 40 kDa | 0 | 0 | 1 | 2 | 3 | | | | | | | |
| AT1G08110 | | lactoylglutathione lyase family protein / glyoxalase I fai | 26 kDa | 0 | 0 | 1 | 2 | 3 | | | | | | | |
| AT1G11545 | | xyloglucan endotransglucosylase/hydrolase 8 | 36 kDa | 0 | 0 | 1 | 2 | 3 | | | | | | | |
| AT1G22920 | | COP9 signalosome 5A | 40 kDa | 0 | 0 | 1 | 2 | 3 | | | | | | | |
| AT1G24100 | | UDP-glucosyl transferase 74B1 | 51 kDa | 0 | 0 | 1 | 2 | 3 | | | | | | | |
| AT1G28200 | | FH interacting protein 1 | 28 kDa | 0 | 0 | 1 | 2 | 3 | | | | | | | |
| AT1G50110 | | D-aminoacid aminotransferase-like PLP-dependent enz | 39 kDa | 0 | 0 | 1 | 2 | 3 | | | | | | | |
| AT1G54520 | | unknown protein; FUNCTIONS IN: molecular_function | 43 kDa | 0 | 0 | 1 | 2 | 3 | | | | | | | |
| AT1G62640 | | 3-ketoacyl-acyl carrier protein synthase III | 43 kDa | 0 | 0 | 1 | 2 | 3 | | | | | | | |
| AT1G63850 | | BTB/POZ domain-containing protein | ? | 0 | 0 | 1 | 2 | 3 | | | | | | | |
| AT1G66940 | | protein kinase-related | 37 kDa | 0 | 0 | 1 | 2 | 3 | | | | | | | |
| AT1G67690 | | Zincin-like metalloproteases family protein | 80 kDa | 0 | 0 | 1 | 2 | 3 | | | | | | | |
| AT1G76510 | | ARID/BRIGHT DNA-binding domain-containing protein | 48 kDa | 0 | 0 | 1 | 2 | 3 | | | | | | | |
| AT2G01120 | | origin recognition complex subunit 4 | 47 kDa | 0 | 0 | 1 | 2 | 3 | | | | | | | |
| AT2G01270 | | quiescin-sulfhydryl oxidase 2 | 56 kDa | 0 | 0 | 1 | 2 | 3 | | | | | | | |
| AT2G01320 | AT2G01320.7;AT2G01320.6;AT2G01320.8;AT2G01320.5;AT2G01320.4;AT2G01320.1;AT2G01320.2;AT2G01320.3 | ABC-2 type transporter family protein | 79 kDa | 0 | 0 | 1 | 2 | 3 | | | | | | | |
| AT2G01470 | | SEC12P-like 2 protein | 43 kDa | 0 | 0 | 1 | 2 | 3 | | | | | | | |
| AT2G18245 | | alpha/beta-Hydrolases superfamily protein | 45 kDa | 0 | 0 | 1 | 2 | 3 | | | | | | | |
| AT2G20635 | | ATP binding;protein kinases;protein serine/threonine k | 61 kDa | 0 | 0 | 1 | 2 | 3 | | | | | | | |
| AT2G31670 | AT2G31670.1 | Stress responsive alpha-beta barrel domain protein | 29 kDa | 0 | 1 | 1 | 2 | 4 | | | | 3.4E+06 | 1.5E+06 | 0.0E+00 | 4.9E+06 |
| AT2G32260 | | phosphorylcholine cytidylyltransferase | 38 kDa | 0 | 0 | 1 | 2 | 3 | | | | | | | |
| AT2G35110 | | transcription activators | 159 kDa | 0 | 0 | 1 | 2 | 3 | | | | | | | |
| AT2G38760 | | annexin 3 | 36 kDa | 0 | 0 | 1 | 2 | 3 | | | | | | | |
| AT2G39780 | | ribonuclease 2 | 29 kDa | 0 | 0 | 1 | 2 | 3 | | | | | | | |
| AT2G45790 | | phosphomannomutase | 28 kDa | 0 | 4 | 1 | 2 | 7 | | | | | | | |
| AT3G01310 | | Phosphoglycerate mutase-like family protein | 119 kDa | 0 | 0 | 1 | 2 | 3 | | | | | | | |
| AT3G01400 | | ARM repeat superfamily protein | 38 kDa | 0 | 0 | 1 | 2 | 3 | | | | | | | |
| AT3G02710 | AT3G02710.1 | ARM repeat superfamily protein | 60 kDa | 0 | 0 | 1 | 2 | 3 | | | | | | | |
| AT3G03570 | | Protein of unknown function (DUF3550/UPF0682) | 69 kDa | 0 | 0 | 1 | 2 | 3 | | | | | | | |
| AT3G04880 | | DNA-damage-repair/toleration protein (DRT102) | 33 kDa | 0 | 0 | 1 | 2 | 3 | | | | | | | |
| AT3G10220 | | tubulin folding cofactor B | 27 kDa | 0 | 1 | 1 | 2 | 4 | | | | | | | |
| AT3G10880 | | unknown protein; BEST Arabidopsis thaliana protein m | ? | 0 | 1 | 1 | 2 | 4 | | | | | | | |
| AT3G15260 | | Protein phosphatase 2C family protein | 32 kDa | 0 | 1 | 1 | 2 | 4 | | | | | | | |
| AT3G15410 | AT3G15410.1;AT3G15410.2 | Leucine-rich repeat (LRR) family protein | 64 kDa | 0 | 0 | 1 | 2 | 3 | | | | 3.4E+05 | 2.1E+05 | 0.0E+00 | 5.5E+05 |
| AT3G18210 | | 2-oxoglutarate (2OG) and Fe(II)-dependent oxygenase s | 45 kDa | 0 | 0 | 1 | 2 | 3 | | | | | | | |
| AT3G18490 | | Eukaryotic aspartyl protease family protein | 53 kDa | 0 | 0 | 1 | 2 | 3 | | | | | | | |
| AT3G20170 | | ARM repeat superfamily protein | 52 kDa | 0 | 0 | 1 | 2 | 3 | | | | | | | |
| AT3G20890 | AT3G20890.1;AT3G20890.2 | RNA-binding (RRM/RBD/RNP motifs) family protein | 32 kDa | 0 | 1 | 1 | 2 | 4 | | | | 6.4E+06 | 6.8E+05 | 1.2E+06 | 8.3E+06 |
| AT3G22220 | | hAT transposon superfamily | 86 kDa | 0 | 0 | 1 | 2 | 3 | | | | | | | |
| AT3G24480 | AT3G24480.1 | Leucine-rich repeat (LRR) family protein | 55 kDa | 0 | 0 | 1 | 2 | 3 | | | | 8.3E+06 | 6.2E+06 | 1.7E+06 | 1.6E+07 |
| AT3G27320 | | alpha/beta-Hydrolases superfamily protein | 51 kDa | 0 | 0 | 1 | 2 | 3 | | | | | | | |
| AT3G29770 | | methyl esterase 11 | 43 kDa | 0 | 0 | 1 | 2 | 3 | | | | | | | |
| AT3G51010 | AT3G51010.1 | unknown protein; FUNCTIONS IN: molecular_function | 22 kDa | 0 | 0 | 1 | 2 | 3 | | | | 4.7E+06 | 8.3E+06 | 2.4E+06 | 1.5E+07 |
| AT3G52630 | | Nucleic acid-binding, OB-fold-like protein | 12 kDa | 0 | 0 | 1 | 2 | 3 | | | | | | | |
| AT3G60360 | AT3G60360.1 | embryo sac development arrest 14 | 27 kDa | 0 | 0 | 1 | 2 | 3 | | | | | | | |
| AT4G04920 | | sensitive to freezing 6 | 138 kDa | 0 | 0 | 1 | 2 | 3 | | | | | | | |
| AT4G11090 | | TRICHOME BIREFRINGENCE-LIKE 23 | 49 kDa | 0 | 0 | 1 | 2 | 3 | | | | | | | |
| AT4G22745 | | methyl-CPG-binding domain 1 | 23 kDa | 0 | 0 | 1 | 2 | 3 | | | | | | | |
| AT4G28510 | | prohibitin 1 | 32 kDa | 0 | 0 | 1 | 2 | 3 | | | | | | | |
| AT4G31860 | | Protein phosphatase 2C family protein | 39 kDa | 0 | 0 | 1 | 2 | 3 | | | | | | | |
| AT4G31890 | | ARM repeat superfamily protein | 56 kDa | 0 | 0 | 1 | 2 | 3 | | | | | | | |
| AT4G35870 | | early-responsive to dehydration stress protein (ERD4) | 93 kDa | 0 | 0 | 1 | 2 | 3 | | | | | | | |
| AT4G38470 | | ACT-like protein tyrosine kinase family protein | 65 kDa | 0 | 0 | 1 | 2 | 3 | | | | | | | |
| AT5G01210 | | HXXXD-type acyl-transferase family protein | 52 kDa | 0 | 0 | 1 | 2 | 3 | | | | | | | |
| AT5G03070 | | importin alpha isoform 9 | 57 kDa | 0 | 0 | 1 | 2 | 3 | | | | | | | |
| AT5G04920 | | EAP30/Vps36 family protein | 49 kDa | 0 | 0 | 1 | 2 | 3 | | | | | | | |
| AT5G07360 | | Amidase family protein | 73 kDa | 0 | 0 | 1 | 2 | 3 | | | | | | | |
| AT5G10630 | AT5G10630.1;AT5G10630.5;AT5G10630.2;AT5G10630.3;AT5G10630.4 | Translation elongation factor EF1A/initiation factor IF2 | 72 kDa | 0 | 0 | 1 | 2 | 3 | | | | 6.0E+05 | 1.6E+05 | 0.0E+00 | 7.6E+05 |
| AT5G11010 | | Pre-mRNA cleavage complex II protein family | 36 kDa | 0 | 0 | 1 | 2 | 3 | | | | | | | |
| AT5G12150 | | Rho GTPase activation protein (RhoGAP) with PH doma | 85 kDa | 0 | 0 | 1 | 2 | 3 | | | | | | | |
| AT5G13360 | | Auxin-responsive GH3 family protein | 76 kDa | 0 | 0 | 1 | 2 | 3 | | | | | | | |
| AT5G17510 | | unknown protein; BEST Arabidopsis thaliana protein m | ? | 0 | 0 | 1 | 2 | 3 | | | | | | | |
| AT5G19210 | | P-loop containing nucleoside triphosphate hydrolases | 53 kDa | 0 | 0 | 1 | 2 | 3 | | | | | | | |
| AT5G20060 | | alpha/beta-Hydrolases superfamily protein | 27 kDa | 0 | 0 | 1 | 2 | 3 | | | | | | | |
| AT5G23510 | | unknown protein; BEST Arabidopsis thaliana protein m | 30 kDa | 0 | 0 | 1 | 2 | 3 | | | | | | | |
| AT5G30495 | | Fcf2 pre-rRNA processing protein | 23 kDa | 0 | 0 | 1 | 2 | 3 | | | | | | | |
| AT5G42370 | | Calcineurin-like metallo-phosphoesterase superfamily | | 50 kDa | 0 | 0 | 1 | 2 | 3 | | | | | | |
| AT5G42770 | | Maf-like protein | 23 kDa | 0 | 0 | 1 | 2 | 3 | | | | | | | |
| AT5G43400 | | Uncharacterised conserved protein UCP015417, vWA | 75 kDa | 0 | 0 | 1 | 2 | 3 | | | | | | | |
| AT5G49810 | | methionine S-methyltransferase | 119 kDa | 0 | 0 | 1 | 2 | 3 | | | | | | | |
| AT5G51410 | | LUC7 N_terminus domain-containing protein | 39 kDa | 0 | 0 | 1 | 2 | 3 | | | | | | | |
| AT5G54160 | | O-methyltransferase 1 | 40 kDa | 0 | 0 | 1 | 2 | 3 | | | | | | | |
| AT5G55000 | | potassium channel tetramerisation domain-containing | 33 kDa | 0 | 0 | 1 | 2 | 3 | | | | | | | |
| AT5G57160 | | DNA ligase IV | 138 kDa | 0 | 0 | 1 | 2 | 3 | | | | | | | |
| AT5G58370 | | P-loop containing nucleoside triphosphate hydrolases | 52 kDa | 0 | 0 | 1 | 2 | 3 | | | | | | | |
| AT5G63530 | | farnesylated protein 3 | 39 kDa | 0 | 0 | 1 | 2 | 3 | | | | | | | |
| AT5G65470 | | O-fucosyltransferase family protein | 57 kDa | 0 | 0 | 1 | 2 | 3 | | | | | | | |
| AT5G66570 | | PS II oxygen-evolving complex 1 | 35 kDa | 0 | 0 | 1 | 2 | 3 | | | | | | | |
| ATCG00490 | | ribulose-bisphosphate carboxylases | 53 kDa | 0 | 0 | 1 | 2 | 3 | | | | | | | |
| AT2G03440 | | nodulin-related protein 1 | 20 kDa | 0 | 4 | 1 | 0 | 5 | | | | | | | |
| AT2G21640 | AT2G21640.1 | Encodes a protein of unknown function that is a marke | 12 kDa | 3 | 0 | 1 | 0 | 4 | 0.14 | | 0.14 | 8.5E+05 | 1.3E+06 | 0.0E+00 | 2.1E+06 |
| AT5G64730 | AT5G64730.1 | Transducin/WD40 repeat-like superfamily protein | 33 kDa | 0 | 2 | 1 | 0 | 3 | | | | 2.4E+06 | 2.0E+06 | 1.3E+06 | 5.7E+06 |
| AT5G01330 | AT5G01330.1 | pyruvate decarboxylase-3 | 64 kDa | 0 | 38 | 0 | 173 | 211 | | | | 2.1E+06 | 6.8E+06 | 0.0E+00 | 8.9E+06 |
| AT1G60730 | AT1G60730.1;AT1G60710.1;AT1G60730.3;AT1G60680.2;AT1G60750.1;AT1G60690.1;AT1G60680.1 | NAD(P)-linked oxidoreductase superfamily protein | 38 kDa | 0 | 2 | 0 | 11 | 13 | | 3.70 | 3.70 | | | | |
| AT1G73690 | | cyclin-dependent kinase D1;1 | 45 kDa | 0 | 2 | 0 | 7 | 9 | | | | | | | |

| | | | | | | | | | | | | | | | |
|---|---|---|---|---|---|---|---|---|---|---|---|---|---|---|---|
| AT4G32840 | AT4G32840.1;AT5G56630.1;AT4G26270.1;AT4G29220.1;AT5G61580.2;AT5G61580.1 | phosphofructokinase 6 | 51 kDa | 0 | 1 | 0 | 7 | 8 | | | | 1.6E+06 | 1.6E+06 | 0.0E+00 | 3.2E+06 |
| AT5G56630 | | phosphofructokinase 7 | 53 kDa | 0 | 1 | 0 | 7 | 8 | | | | | | | |
| AT1G15880 | | golgi snare 11 | 25 kDa | 0 | 3 | 0 | 5 | 8 | | | | | | | |
| AT4G24280 | | chloroplast heat shock protein 70-1 | 77 kDa | 0 | 1 | 0 | 5 | 6 | | | | | | | |
| AT1G53850 | AT1G53850.2;AT1G53850.1 | 20S proteasome alpha subunit E1 | 26 kDa | 0 | 3 | 0 | 4 | 7 | 0.12 | | 0.12 | 0.0E+00 | 0.0E+00 | 0.0E+00 | 0.0E+00 |
| AT3G12500 | | basic chitinase | 36 kDa | 0 | 3 | 0 | 4 | 7 | | | | | | | |
| AT3G48990 | | AMP-dependent synthetase and ligase family protein | 56 kDa | 0 | 1 | 0 | 4 | 5 | | | | | | | |
| AT1G62730 | | Terpenoid synthases superfamily protein | 34 kDa | 0 | 2 | 0 | 3 | 5 | | | | | | | |
| AT3G25530 | | glyoxylate reductase 1 | 31 kDa | 0 | 1 | 0 | 3 | 4 | | | | | | | |
| AT3G60340 | | alpha/beta-Hydrolases superfamily protein | 37 kDa | 0 | 1 | 0 | 3 | 4 | | | | | | | |
| AT5G16840 | | binding partner of acd11 1 | 27 kDa | 0 | 1 | 0 | 3 | 4 | | 0.22 | 0.22 | | | | |
| AT5G66410 | | phosducin-like protein 3 homolog | 26 kDa | 0 | 1 | 0 | 3 | 4 | | | | | | | |
| AT1G47200 | | WPP domain protein 2 | 19 kDa | 0 | 2 | 0 | 2 | 4 | | | | | | | |
| AT2G22300 | | signal responsive 1 | 116 kDa | 5 | 1 | 0 | 2 | 8 | | | | | | | |
| AT3G26090 | | G-protein coupled receptors;GTPase activators | 53 kDa | 0 | 1 | 0 | 2 | 3 | | | | | | | |
| AT4G27760 | | NAD(P)-binding Rossmann-fold superfamily protein | 41 kDa | 0 | 2 | 0 | 2 | 4 | | | | | | | |
| AT5G51220 | | ubiquinol-cytochrome C chaperone family protein | 32 kDa | 0 | 1 | 0 | 2 | 3 | | | | | | | |
| AT4G38150 | | Pentatricopeptide repeat (PPR) superfamily protein | 33 kDa | 0 | 2 | 0 | 1 | 3 | | | | | | | |

**Supplemental Dataset 2. LC/MS-MS data corresponding to CCVs digested in solution.** Complete list of protein groups identified by MaxQuant in LC/MS-MS dataset corresponding to CCVs digested in solution and not within gel. Only protein groups with peptides identified in at least two of three biological replicate were retained. Groups are sorted by the iBAQ (intensity based absolute quantitation) values of the first protein listed in the majority protein ID column from high abundance to low. Column header descriptions are listed below.

[illegible]

| | | | | | |
|---|---|---|---|---|---|
| AT1G27090.1 | AT1G27090.1 | 17 | 17 | 17 | glycine-rich protein \| 201606 |
| AT5G07090.3;AT5G07090.2;AT5G07090.1 | AT5G07090.3;AT5G07090 | 29;29;29;28;20 | 29;29;29;28;20 | 2;2;2;2;2 | Ribosomal protein S4 (RPS4A) family protein \| 201606; Ribosomal protein S4 (RPS4A) [illegible] |
| AT2G38040.2;AT2G38040.1 | AT2G38040.2;AT2G38040 | 35;35 | 35;35 | 35;35 | CAC3 \| acetyl Co-enzyme a carboxylase carboxyltransferase alpha subunit \| 201606; [illegible] |
| AT5G44750.2;AT5G44750.1 | AT5G44750.2;AT5G44750 | 7;7 | 7;7 | 7;7 | ATHD2A, HD2A, HDA3, HDT1 \| histone deacetylase 3 \| 201606; ATHD2A, HD2A, HDA3 [illegible] |
| AT2G26780.2;AT2G26780.1 | AT2G26780.2;AT2G26780 | 66;65 | 66;65 | 66;65 | ARM repeat superfamily protein \| 201606; ARM repeat superfamily protein \| 201606 |
| AT1G62390.1 | AT1G62390.1 | 28 | 28 | 28 | CLMP1, Phox2 \| Octicosapeptide/Phox/Bem1p (PB1) domain-containing protein / tet[illegible] |
| AT1G65930.1;AT5G14590.1;AT1G54340.1 | AT1G65930.1 | 21;4;3;2 | 21;4;3;2 | 21;4;3;2 | cICDH \| cytosolic NADP+-dependent isocitrate dehydrogenase \| 201606 |
| AT5G27700.1 | AT5G27700.1 | 3 | 1 | 1 | EVR1L1, RPS21C \| Ribosomal protein S21e \| 201606 |
| AT4G20360.2;AT4G20360.1 | AT4G20360.2;AT4G20360 | 20;20 | 20;20 | 20;20 | ATRAB8D, ATRABE1B, RAB8b \| RAB GTPase homolog E1B \| 201606; ATRAB8D, ATRA[illegible] |
| AT4G23460.1;AT4G23460.2 | AT4G23460.1;AT4G23460 | 59;47 | 19;19 | 19;19 | Adaptin family protein \| 201606; Adaptin family protein \| 201606 |
| AT4G30190.1;AT4G30190.2;AT5G47950.1 | AT4G30190.1;AT4G30190 | [illegible] | [illegible] | [illegible] | AHA2, AHA2, HA2, PMA2 \| H(+)-ATPase 2 \| 201606; AHA2, AHA2, HA2, PMA2 \| H(+[illegible] |
| AT5G39740.2;AT5G39740.1 | AT5G39740.2;AT5G39740 | 32;32 | 32;32 | 6;6 | OLI7, RPL5B \| ribosomal protein L5 B \| 201606; OLI7, RPL5B \| ribosomal protein L5 B [illegible] |
| AT2G33340.1;AT2G33340.2;AT2G33340.3 | AT2G33340.1;AT2G33340 | 24;24;21 | 24;24;21 | 22;22;09 | MAC3B \| MOS4-associated complex 3B \| 201606; MAC3B \| MOS4-associated compl[illegible] |
| AT5G22440.2;AT5G22440.1 | AT5G22440.2;AT5G22440 | 11;11 | 5;5 | 5;5 | Ribosomal protein L1p/L10e family \| 201606; Ribosomal protein L1p/L10e family \| [illegible] |
| AT3G06610.1 | AT3G06610.1 | 4 | 4 | 4 | DNA-binding enhancer protein-like protein \| 201606 |
| AT5G53430.1 | AT5G53430.1 | 12 | 12 | 2 | Ribosomal protein L11 family protein \| 201606 |
| AT5G64350.1 | AT5G64350.1 | 2 | 2 | 2 | ATFKBP12, FKBP12, FKP12 \| FK506-binding protein 12 \| 201606 |
| AT5G49430.2;AT5G49430.7;AT5G49430.4 | AT5G49430.2;AT5G49430 | 8;8;8;8;8;8;8 | 8;8;8;8;8;8;8 | 7;7;7;7;7;7;7 | At-SR34a, SR34a, SRp34a \| SER/ARG-rich protein 34A \| 201606; At-SR34a, SR34a, SRp[illegible] |
| AT5G18180.1 | AT5G18180.1 | 31 | 31 | 8 | RACK1C, RACK1C_AT \| receptor for activated C kinase 1C \| 201606 |
| AT2G01600.1 | AT2G01600.1 | 14 | 14 | 11 | ENTH/ANTH/VHS superfamily protein \| 201606 |
| AT5G02880.1 | AT5G02880.1 | 20 | 20 | 16 | KIN7 \| Leucine-rich repeat protein kinase family protein \| 201606 |
| AT4G19210.1 | AT4G19210.1 | 15 | 15 | 15 | ABCE2, ATRLI2, RLI2 \| RNAse l inhibitor protein 2 \| 201606 |
| AT3G13580.9;AT3G13580.8;AT3G13580.3 | AT3G13580.9;AT3G1358 | [illegible] | [illegible] | 5;5;5;5;5;5;5;5;5 | Ribosomal protein L30/L7 family protein \| 201606; Ribosomal protein L30/L7 family [illegible] |
| AT5G46630.1;AT5G46630.2 | AT5G46630.1;AT5G46630 | 19;18 | 19;18 | 19;18 | AP2M \| Clathrin adaptor complexes medium subunit family protein \| 201606; AP2M [illegible] |
| AT3G55220.1;AT3G55220.3;AT3G55220.2 | AT3G55220.1;AT3G55220 | 41;41;41;41 | 41;41;41;41 | 41;41;41;41 | ATSAP130b, SAP130b \| Cleavage and polyadenylation specificity factor (CPSF) A subu[illegible] |
| AT5G51480.1 | AT5G51480.1 | 13 | 11 | 11 | SKS2 \| SKU5 similar 2 \| 201606 |
| AT1G20960.2;AT1G20960.1;AT2G42270.1 | AT1G20960.2;AT1G20960 | 88;88;16;1;1 | 88;88;16;1;1 | 88;88;16;1;1 | BRR2, emb1507 \| U5 small nuclear ribonucleoprotein helicase \| 201606; BRR2, emb1[illegible] |
| AT3G13920.1;AT3G13920.4;AT3G13920.3 | AT3G13920.1;AT3G1392 | [illegible] | [illegible] | [illegible] | EIF4A1, RH4, TIF4A1 \| eukaryotic translation initiation factor 4A1 \| 201606; EIF4A1, R[illegible] |
| AT1G15810.1 | AT1G15810.1 | 18 | 18 | 16 | S15/NS1, RNA-binding protein \| 201606 |
| AT2G40590.1 | AT2G40590.1 | 10 | 10 | 3 | Ribosomal protein S26e family protein \| 201606 |
| AT5G13780.1 | AT5G13780.1 | 8 | 8 | 8 | NAA10 \| Acyl-CoA N-acyltransferases (NAT) superfamily protein \| 201606 |
| AT1G10580.1 | AT1G10580.1 | 25 | 25 | 25 | Transducin/WD40 repeat-like superfamily protein \| 201606 |
| AT2G29550.1 | AT2G29550.1 | 35 | 14 | 6 | TBB7, TUB7 \| tubulin beta-7 chain \| 201606 |
| AT1G61150.5;AT1G61150.9;AT1G61150.3 | AT1G61150.5;AT1G6115 | [illegible] | [illegible] | [illegible] | LisH and RanBPM domains containing protein \| 201606; LisH and RanBPM domains [illegible] |
| AT1G64750.1;AT1G64750.2;AT1G64750.3 | AT1G64750.1;AT1G6475 | 2;2;1 | 2;2;1 | 2;2;1 | ATDSS1(I), DSS1(I) \| deletion of SUV3 suppressor 1(I) \| 201606; ATDSS1(I), DSS1(I) [illegible] |
| AT5G05470.1 | AT5G05470.1 | 14 | 11 | 11 | ATEIF2-A2, EIF2 ALPHA, EIF2-A2 \| eukaryotic translation initiation factor 2 alpha subu[illegible] |
| AT2G17120.1 | AT2G17120.1 | 8 | 8 | 8 | CL-1, LYM2, LYP1 \| lysm domain GPI-anchored protein 2 precursor \| 201606 |
| AT3G64550.1 | AT3G64550.1 | 25 | 25 | 25 | ABCF3, ATABCF3, ATGCN20, ATGCN3, GCN20, GCN3, SCORD5 \| general control non-r[illegible] |
| AT5G09770.1;AT5G64650.1 | AT5G09770.1;AT5G64650 | 5;5 | 5;5 | 5;5 | Ribosomal protein L17 family protein \| 201606; Ribosomal protein L17 family protein [illegible] |
| AT3G54200.1 | AT3G54200.1 | 5 | 5 | 5 | Late embryogenesis abundant (LEA) hydroxyproline-rich glycoprotein family \| 201606 |
| ATCG00500.1 | ATCG00500.1 | 22 | 22 | 22 | ACCD \| acetyl-CoA carboxylase carboxyl transferase subunit beta \| 201606 |
| AT2G45820.1 | AT2G45820.1 | 7 | 7 | 7 | Rem1.3 \| Remorin family protein \| 201606 |
| AT2G01660.2;AT2G01660.3;AT2G01660.1 | AT2G01660.2;AT2G01660 | 3;3;3 | 3;3;3 | 3;3;3 | PDLP6 \| plasmodesmata-located protein 6 \| 201606; PDLP6 \| plasmodesmata-locate[illegible] |
| AT1G42440.1 | AT1G42440.1 | 31 | 31 | 31 | pre-rRNA-processing TSR1-like protein \| 201606 |
| AT2G36130.1 | AT2G36130.1 | 5 | 5 | 5 | Cyclophilin-like peptidyl-prolyl cis-trans isomerase family protein \| 201606 |
| AT4G20830.2;AT4G20830.1;AT1G30760.1 | AT4G20830.2;AT4G2083 | 25;25;1;1;1 | 25;25;1;1;1 | 21;21;0;0;0 | ATBBE19, ATBBE20, OGOX1 \| FAD-binding Berberine family protein \| 201606; ATBBE[illegible] |
| AT3G07050.1 | AT3G07050.1 | 21 | 21 | 21 | NSN1 \| GTP-binding family protein \| 201606 |
| AT3G53890.2;AT3G53890.1 | AT3G53890.2;AT3G53890 | 4;4 | 4;4 | 2;2 | EVR1, RPS21B \| Ribosomal protein S21e \| 201606; EVR1, RPS21B \| Ribosomal protei[illegible] |
| AT3G55440.1;AT2G21170.3;AT2G21170.1 | AT3G55440.1 | 9;1;1 | 9;1;1 | 9;1;1 | ATCTIMC, CYTOTPI, TPI \| triosephosphate isomerase \| 201606 |

[illegible]

| | | | | | |
|---|---|---|---|---|---|
| AT5G12140.1 | AT5G12140.1 | 3 | 2 | 2 | ATCYS1, CYS1 \| cystatin-1 \| 201606 |
| AT5G58710.1 | AT5G58710.1 | 5 | 5 | 4 | ROC7 \| rotamase CYP 7 \| 201606 |
| AT3G22590.1 | AT3G22590.1 | 7 | 7 | 7 | CDC73, PHP \| PLANT HOMOLOGOUS TO PARAFIBROMIN \| 201606 |
| AT4G31480.9;AT4G31480.8;AT4G31480.[illegible] | AT4G31480.9;AT4G3148[illegible] | [illegible] | [illegible] | [illegible] | Coatomer, beta subunit \| 201606; Coatomer, beta [illegible] |
| AT3G22320.1 | AT3G22320.1 | 4 | 4 | 4 | ATRPABC24.3, NRPB5, NRPD5, RPB5A \| Eukaryotic rpb5 RNA polymerase subunit fam[illegible] |
| AT2G44640.1 | AT2G44640.1 | 7 | 7 | 7 | TRIGALACTOSYLDIACYLGLYCEROL-like protein \| 201606 |
| AT5G54160.2;AT5G54160.1 | AT5G54160.2;AT5G5416[illegible] | 8;8 | 8;8 | 8;8 | ATNPF8.1, ATPTR1, NPF8.1, PTR1 \| peptide transporter 1 \| 201606;ATNPF8.1, ATPTR[illegible] |
| AT3G51740.1 | AT3G51740.1 | 9 | 9 | 9 | IMK2 \| inflorescence meristem receptor-like kinase 2 \| 201606 |
| AT3G11900.1 | AT3G11900.1 | 8 | 8 | 8 | Translation initiation factor SUI1 family protein \| 201606 |
| AT5G03740.1 | AT5G03740.1 | 6 | 6 | 6 | HD2C, HDT3 \| histone deacetylase 2C \| 201606 |
| AT2G33040.1 | AT2G33040.1 | 11 | 11 | 11 | ATP3 \| gamma subunit of Mt ATP synthase \| 201606 |
| AT1G74010.1 | AT1G74010.1 | 3 | 2 | 2 | Calcium-dependent phosphotriesterase superfamily protein \| 201606 |
| AT1G22280.3;AT1G22280.1;AT1G22280.[illegible] | AT1G22280.3;AT1G2228[illegible] | 8;8;8 | 8;8;8 | 8;8;8 | PAPP2C \| phytochrome-associated protein phosphatase type 2C \| 201606;PAPP2C \| [illegible] |
| AT3G42725.1 | AT3G42725.1 | 2 | 2 | 2 | Putative membrane lipoprotein \| 201606 |
| AT5G64400.1;AT5G64400.2 | AT5G64400.1;AT5G6440[illegible] | 4;4 | 4;4 | 4;4 | CHCH domain protein \| 201606;CHCH domain protein \| 201606 |
| [illegible] | [illegible] | [illegible] | [illegible] | [illegible] | [illegible] |

| | | | | | | |
|---|---|---|---|---|---|---|
| AT5G47690.1;AT5G62500.1;AT5G67270.1 | AT5G47690.1;AT5G6250 | 2;1;1 | 2;1;1 | 2;1;1 | ATEB1a, ATEB1H2, EB1a \| microtubule end binding protein EB1A \| 201606; ATEB1, AT | [illegible] |
| AT1G12000.1 | AT1G12000.1 | 2 | 2 | 2 | Phosphofructokinase family protein \| 201606 | [illegible] |
| AT5G18040.1;AT1G29590.1;AT1G29550.1 | AT5G18040.1;AT1G2959 | 2;1;1 | 2;1;1 | 2;1;1 | AT.EIF4E1, CUM1, EIF4E, eIF4E1 \| eukaryotic translation initiation factor 4E \| 201606; | [illegible] |
| AT5G16250.1 | AT5G16250.1 | 1 | 1 | 1 | transmembrane protein \| 201606 | [illegible] |
| AT1G36240.1 | AT1G36240.1 | 10 | 2 | 2 | Ribosomal protein L7Ae/L30e/S12e/Gadd45 family protein \| 201606 | [illegible] |
| AT2G18040.1 | AT2G18040.1 | 2 | 2 | 2 | PIN1AT \| peptidylprolyl cis/trans isomerase, NIMA-interacting 1 \| 201606 | [illegible] |
| AT2G33730.1;AT1G28180.2;AT1G28180.1 | AT2G33730.1 | 7;1;1 | 7;1;1 | 7;1;1 | P-loop containing nucleoside triphosphate hydrolases superfamily protein \| 201606 | [illegible] |
| AT1G56450.1 | AT1G56450.1 | 2 | 2 | 2 | PBG1 \| 20S proteasome beta subunit G1 \| 201606 | [illegible] |
| AT5G02500.1 | AT5G02500.1 | 35 | 7 | 6 | ATHSP70, HSC70-1, HSP70 \| heat shock protein 70 \| 201606 | [illegible] |
| AT3G14110.2;AT3G14110.1;AT3G14110.3 | AT3G14110.2;AT3G1411 | 3;3;3 | 3;3;3 | 3;3;3 | FLU \| Tetratricopeptide repeat (TPR)-like superfamily protein \| 201606; FLU \| Tetratric | [illegible] |
| AT2G24360.1 | AT2G24360.1 | 6 | 5 | 4 | STYK \| Protein kinase superfamily protein \| 201606 | [illegible] |
| AT3G24170.3;AT3G24170.2;AT3G24170.1 | AT3G24170.3;AT3G2417 | 2;2;2 | 2;2;2 | 2;2;2 | ATGR1, GR1 \| glutathione-disulfide reductase \| 201606; ATGR1, GR1 \| glutathione-di | [illegible] |
| AT5G02740.1 | AT5G02740.1 | 1 | 1 | 1 | Ribosomal protein S24e family protein \| 201606 | [illegible] |
| AT1G65730.1 | AT1G65730.1 | 4 | 4 | 4 | YSL7 \| YELLOW STRIPE like 7 \| 201606 | [illegible] |
| AT5G49460.2;AT5G49460.1 | AT5G49460.2;AT5G4946 | 20;20 | 20;20 | 6;6 | ACLB-2 \| ATP citrate lyase subunit B 2 \| 201606; ACLB-2 \| ATP citrate lyase subunit B | [illegible] |
| AT1G13650.1;AT5G13650.2 | AT5G13650.1;AT5G1365 | 4;4 | 4;4 | 4;4 | SVR3 \| elongation factor family protein \| 201606; SVR3 \| elongation factor family pro | [illegible] |
| AT5G14050.1 | AT5G14050.1 | 4 | 4 | 4 | Transducin/WD40 repeat-like superfamily protein \| 201606 | [illegible] |
| AT3G21465.1;AT3G21465.2 | AT3G21465.1;AT3G2146 | 8;7 | 6;5 | 6;5 | adenylyl cyclase \| 201606; adenylyl cyclase \| 201606 | [illegible] |
| AT3G57990.1 | AT3G57990.1 | 6 | 6 | 6 | OXP40 \| hypothetical protein \| 201606 | [illegible] |
| AT1G54610.3;AT1G54610.2;AT1G54610.1 | AT1G54610.3;AT1G5461 | [illegible] | [illegible] | [illegible] | Protein kinase superfamily protein \| 201606; Protein kinase superfamily protein \| 201 | [illegible] |
| AT4G36750.1 | AT4G36750.1 | 1 | 1 | 1 | Quinone reductase family protein \| 201606 | [illegible] |
| AT3G23990.1 | AT3G23990.1 | 10 | 4 | 4 | HSP60, HSP60-3B \| heat shock protein 60 \| 201606 | [illegible] |
| AT3G12280.2;AT3G12280.1 | AT3G12280.2;AT3G1228 | 8;8 | 8;8 | 8;8 | ATRBR1, RB, RB1, RBR, RBR1 \| retinoblastoma-related 1 \| 201606; ATRBR1, RB, RB1, | [illegible] |
| AT2G16570.1 | AT2G16570.1 | 4 | 4 | 4 | ASE1, ATASE, ATASE1 \| GLN phosphoribosyl pyrophosphate amidotransferase 1 \| 201 | [illegible] |
| AT4G29900.2;AT4G29900.1 | AT4G29900.2;AT4G2990 | 9;9 | 8;8 | 8;8 | ACA10, ATACA10, CIF1 \| autoinhibited Ca(2+)-ATPase 10 \| 201606; ACA10, ATACA10, | [illegible] |
| AT5G48650.1 | AT5G48650.1 | 4 | 4 | 4 | Nuclear transport factor 2 (NTF2) family protein with RNA binding (RRM-RBD-RNP m | [illegible] |
| AT4G36945.1 | AT4G36945.1 | 3 | 3 | 3 | PLC-like phosphodiesterases superfamily protein \| 201606 | [illegible] |
| AT5G04885.3;AT5G04885.1;AT5G04885.2 | AT5G04885.3;AT5G0488 | 3;3;3 | 3;3;3 | 3;3;3 | BGLC3 \| Glycosyl hydrolase family protein \| 201606; BGLC3 \| Glycosyl hydrolase fam | [illegible] |
| AT1G61620.1 | AT1G61620.1 | 3 | 3 | 3 | CSU1 \| phosphoinositide binding protein \| 201606 | [illegible] |
| AT2G16640.3;AT2G16640.2;AT2G16640.1 | AT2G16640.3;AT2G1664 | 8;8;8;2 | 8;8;8;2 | 8;8;8;2 | ATTOC132, TOC132 \| multimeric translocon complex in the outer envelope membrane | [illegible] |
| ATCG00130.1 | ATCG00130.1 | 2 | 2 | 2 | ATPF \| ATPase, F0 complex, subunit B/B', bacterial/chloroplast \| 201606 | [illegible] |
| AT2G20280.1 | AT2G20280.1 | 7 | 7 | 7 | Zinc finger C-x8-C-x5-C-x3-H type family protein \| 201606 | [illegible] |
| AT5G62310.1;AT5G62310.3;AT5G62310.2 | AT5G62310.1;AT5G6231 | 17;15;15 | 17;15;15 | 7;6;6 | RNA helicase family protein \| 201606; RNA helicase family protein \| 201606; RNA heli | [illegible] |
| AT3G48430.1;AT3G48430.2 | AT3G48430.1 | 3;1 | 3;1 | 3;1 | JMJ12, REF6 \| relative of early flowering 6 \| 201606 | [illegible] |
| AT5G46800.1;AT5G46800.2 | AT5G46800.1;AT5G4680 | 6;5 | 6;5 | 6;5 | RNA-binding (RRM/RBD/RNP motifs) family protein \| 201606; RNA-binding (RRM/RB | [illegible] |
| AT5G63620.1;AT5G63620.2;AT5G63620.3 | AT5G63620.1;AT5G6362 | 4;3;2 | 4;3;2 | 4;3;2 | HXR2 \| GroES-like zinc-binding alcohol dehydrogenase family protein \| 201606; HXR2 | [illegible] |
| AT5G41190.1 | AT5G41190.1 | 5 | 5 | 5 | ATNOB1, NOB1 \| RNA-binding NOB1-like protein \| 201606 | [illegible] |
| AT1G24280.1 | AT1G24280.1 | 15 | 4 | 4 | G6PD3 \| glucose-6-phosphate dehydrogenase 3 \| 201606 | [illegible] |
| AT2G30740.9;AT2G30740.8;AT2G30740.1 | AT2G30740.9;AT2G3074 | [illegible] | [illegible] | [illegible] | Protein kinase superfamily protein \| 201606; Protein kinase superfamily protein \| 201 | [illegible] |
| AT2G24590.1 | AT2G24590.1 | 4 | 1 | 1 | AT-RSZ22a, RSZ22a \| RNA recognition motif and CCHC-type zinc finger domains conta | [illegible] |
| AT2G18470.3;AT2G18470.2;AT2G18470.1 | AT2G18470.3;AT2G1847 | 5;5;5;4 | 5;5;5;4 | 3;3;3;2 | ATPERK4, PERK4 \| roline-rich extensin-like receptor kinase 4 \| 201606; ATPERK4, PERK | [illegible] |
| AT5G66760.1;AT2G18450.1 | AT5G66760.1 | 3;1 | 3;1 | 3;1 | SDH1-1 \| succinate dehydrogenase 1-1 \| 201606 | [illegible] |
| AT5G64816.4;AT5G64816.3;AT5G64816.2 | AT5G64816.4;AT5G6481 | 1;1;1;1 | 1;1;1;1 | 1;1;1;1 | thionin-like protein \| 201606; thionin-like protein \| 201606; thionin-like protein \| 20 | [illegible] |
| AT5G07830.1 | AT5G07830.1 | 4 | 4 | 4 | AtGUS2, GUS2 \| glucuronidase 2 \| 201606 | [illegible] |
| AT5G06810.1 | AT5G06810.1 | 7 | 7 | 7 | IBR3 \| acyl-CoA dehydrogenase-like protein \| 201606 | [illegible] |
| AT2G03430.1 | AT2G03430.1 | 2 | 2 | 2 | Ankyrin repeat family protein \| 201606 | [illegible] |
| AT1G76680.1;AT1G76680.2;AT1G76690.1 | AT1G76680.1;AT1G7668 | 4;4;2 | 4;4;2 | 4;4;2 | ATOPR1, OPR1 \| 12-oxophytodienoate reductase 1 \| 201606; ATOPR1, OPR1 \| 12-ox | [illegible] |
| AT3G45140.1;AT3G45140.2;AT3G45140.3 | AT3G45140.1;AT3G4514 | 6;6;6 | 6;6;6 | 6;6;6 | NRPC2 \| nuclear RNA polymerase C2 \| 201606; NRPC2 \| nuclear RNA polymerase C2 | [illegible] |
| AT3G57560.1 | AT3G57560.1 | 2 | 2 | 2 | NAGK \| N-acetyl-l-glutamate kinase \| 201606 | [illegible] |
| [illegible] | [illegible] | [illegible] | [illegible] | [illegible] | [illegible] | [illegible] |

| | | | | | | | | | | | | |
|---|---|---|---|---|---|---|---|---|---|---|---|---|
| AT2G17790.1 | AT2G17790.1 | 3 | 3 | 3 | VPS35A, ZIP3 \| vacuolar protein sorting-associated 35A-like protein \| 201606 | 1 | 3 | 3 | 3 | 2 | 1 | [illegible] |
| AT3G10050.1 | AT3G10050.1 | 3 | 3 | 3 | OMR1 \| L-O-methylthreonine resistant 1 \| 201606 | 1 | 3 | 3 | 3 | 1 | 2 | [illegible] |
| AT3G01800.1 | AT3G01800.1 | 1 | 1 | 1 | Ribosome recycling factor \| 201606 | 1 | 1 | 1 | 1 | 1 | 1 | [illegible] |
| AT5G26980.3;AT5G26980.2;AT5G26980.[illegible] | AT5G26980.3;AT5G2698[illegible] | 2;2;2 | 2;2;2 | 2;2;2 | ATSYP41, ATTLG2A, SYP41 \| syntaxin of plants 41 \| 201606;ATSYP41, ATTLG2A, SYP4[illegible] | 3 | 2 | 2 | 2 | 0 | 1 | [illegible] |
| AT5G13990.2;AT5G13990.1 | AT5G13990.2;AT5G1399[illegible] | 3;3 | 3;3 | 3;3 | dentin sialophosphoprotein, putative (DUF1296) \| 201606;dentin sialophosphopro[illegible] | 2 | 3 | 3 | 3 | 2 | 2 | [illegible] |
| AT1G66970.3;AT1G66970.1;AT1G66970.[illegible] | AT1G66970.3;AT1G6697[illegible] | 3;3;3 | 2;2;2 | 2;2;2 | GDPDL1, SVL2 \| SHV3-like 2 \| 201606;GDPDL1, SVL2 \| SHV3-like 2 \| 201606;GDPD[illegible] | 3 | 3 | 3 | 2 | 2 | 3 | [illegible] |
| AT4G08950.1 | AT4G08950.1 | 1 | 1 | 1 | EXO \| Phosphate-responsive 1 family protein \| 201606 | 1 | 1 | 1 | 1 | 1 | 1 | [illegible] |
| AT1G16970.1 | AT1G16970.1 | 1 | 1 | 1 | ATKU70, KU70 \| ATP-dependent DNA helicase 2 subunit Ku70-like protein \| 201606 | 1 | 1 | 1 | 1 | 1 | 1 | [illegible] |
| AT2G01080.1 | AT2G01080.1 | 1 | 1 | 1 | Late embryogenesis abundant (LEA) hydroxyproline-rich glycoprotein family \| 201606 | 1 | 1 | 1 | 1 | 1 | 1 | [illegible] |
| AT5G50380.1 | AT5G50380.1 | 3 | 3 | 3 | ATEXO70F1, EXO70F1 \| exocyst subunit exo70 family protein F1 \| 201606 | 1 | 3 | 3 | 3 | 1 | 2 | [illegible] |
| AT4G25350.1 | AT4G25350.1 | 2 | 2 | 2 | SHB1 \| EXS (ERD1/XPR1/SYG1) family protein \| 201606 | 1 | 2 | 2 | 2 | 1 | 0 | [illegible] |
| AT1G11090.1 | AT1G11090.1 | 2 | 2 | 2 | MAGL1 \| alpha/beta-Hydrolases superfamily protein \| 201606 | 1 | 2 | 2 | 2 | 1 | 2 | [illegible] |
| AT1G59830.2;AT1G59830.1;AT1G10430.[illegible] | AT1G59830.2;AT1G5983[illegible] | 1;1;1;1 | 1;1;1;1 | 1;1;1;1 | PP2A-1 \| protein phosphatase 2A-2 \| 201606;PP2A-1 \| protein phosphatase 2A-2 \| [illegible] | 4 | 1 | 1 | 1 | 1 | 1 | [illegible] |
| AT4G17180.1 | AT4G17180.1 | 1 | 1 | 1 | O-Glycosyl hydrolases family 17 protein \| 201606 | 1 | 1 | 1 | 1 | 1 | 1 | [illegible] |
| AT5G13330.1 | AT5G13330.1 | 1 | 1 | 1 | Rap2.6L \| related to AP2 6l \| 201606 | 1 | 1 | 1 | 1 | 1 | 1 | [illegible] |
| AT4G16630.1 | AT4G16630.1 | 4 | 4 | 4 | DEA(D/H)-box RNA helicase family protein \| 201606 | 1 | 4 | 4 | 4 | 2 | 2 | [illegible] |
| AT5G15570.1 | AT5G15570.1 | 1 | 1 | 1 | Bromodomain transcription factor \| 201606 | 1 | 1 | 1 | 1 | 1 | 1 | [illegible] |
| AT1G22620.1 | AT1G22620.1 | 1 | 1 | 1 | ATSAC1, SAC1 \| Phosphoinositide phosphatase family protein \| 201606 | 1 | 1 | 1 | 1 | 1 | 1 | [illegible] |
| AT5G22790.1 | AT5G22790.1 | 3 | 1 | 1 | RER1 \| reticulata-related 1 \| 201606 | 1 | 3 | 1 | 1 | 3 | 3 | [illegible] |
| AT5G41260.1 | AT5G41260.1 | 4 | 1 | 1 | BSK8 \| kinase with tetratricopeptide repeat domain-containing protein \| 201606 | 1 | 4 | 1 | 1 | 3 | 3 | [illegible] |
| AT4G06634.2;AT4G06634.1 | AT4G06634.2;AT4G0663[illegible] | 1;1 | 1;1 | 1;1 | AtYY1, YY1 \| zinc finger (C2H2 type) family protein \| 201606;AtYY1, YY1 \| zinc finger [illegible] | 2 | 1 | 1 | 1 | 1 | 1 | [illegible] |
| AT5G02310.2;AT5G02310.3;AT5G02310.[illegible] | AT5G02310.2;AT5G0231[illegible] | 4;4;4 | 4;4;4 | 4;4;4 | GED1, PRT6 \| proteolysis 6 \| 201606;GED1, PRT6 \| proteolysis 6 \| 201606;GED1, P[illegible] | 3 | 4 | 4 | 4 | 3 | 2 | [illegible] |
| AT1G72730.1 | AT1G72730.1 | 17 | 1 | 1 | DEA(D/H)-box RNA helicase family protein \| 201606 | 1 | 17 | 1 | 1 | 16 | 15 | [illegible] |
| AT5G12370.3;AT5G12370.2;AT5G12370.[illegible] | AT5G12370.3;AT5G1237[illegible] | 5;5;5 | 5;5;5 | 5;5;5 | SEC10 \| exocyst complex component sec10 \| 201606;SEC10 \| exocyst complex comp[illegible] | 3 | 5 | 5 | 5 | 3 | 5 | [illegible] |
| AT5G42920.2;AT5G42920.1 | AT5G42920.2;AT5G4292[illegible] | 6;6 | 6;6 | 5;5 | AtTHO5, THO5 \| THO complex, subunit 5 \| 201606;AtTHO5, THO5 \| THO complex, s[illegible] | 2 | 6 | 6 | 5 | 2 | 2 | [illegible] |
| AT5G13510.1 | AT5G13510.1 | 1 | 1 | 1 | EMB3136 \| Ribosomal protein L10 family protein \| 201606 | 1 | 1 | 1 | 1 | 1 | 1 | [illegible] |
| AT5G35200.2;AT5G35200.1 | AT5G35200.2;AT5G3520[illegible] | 2;2 | 2;2 | 2;2 | ENTH/ANTH/VHS superfamily protein \| 201606;ENTH/ANTH/VHS superfamily protein [illegible] | 2 | 2 | 2 | 2 | 1 | 2 | [illegible] |
| AT5G52920.1 | AT5G52920.1 | 2 | 1 | 1 | PKP1, PKP2, PKP-BETA1 \| plastidic pyruvate kinase beta subunit 1 \| 201606 | 1 | 2 | 1 | 1 | 2 | 2 | [illegible] |
| AT1G74040.1;AT1G18500.1;AT1G74040.[illegible] | AT1G74040.1;AT1G1850[illegible] | 2;2;1;1 | 2;2;1;1 | 2;2;1;1 | IMS1, IPMS2, MAML-3 \| 2-isopropylmalate synthase 1 \| 201606;IPMS1, MAML-4 \| [illegible] | 4 | 2 | 2 | 2 | 1 | 2 | [illegible] |
| AT3G60415.1;AT3G60420.1;AT3G60450.[illegible] | AT3G60415.1;AT3G6042[illegible] | 1;1;1;1 | 1;1;1;1 | 1;1;1;1 | phosphoglycerate mutase family protein \| 201606;phosphoglycerate mutase family p[illegible] | 4 | 1 | 1 | 1 | 1 | 1 | [illegible] |
| AT2G25170.4;AT2G25170.1;AT2G25170.[illegible] | AT2G25170.4;AT2G2517[illegible] | 3;3;3;3 | 3;3;3;3 | 3;3;3;3 | CHD3, CHR6, CKH2, EPP1, GYM, PKL, SSL2 \| chromatin remodeling factor CHD3 (PICK[illegible] | 4 | 3 | 3 | 3 | 1 | 2 | [illegible] |
| AT5G59980.1;AT5G59980.3;AT5G59980.[illegible] | AT5G59980.1;AT5G5998[illegible] | 3;3;3;3 | 3;3;3;3 | 3;3;3;3 | AtRPP30, GAF1, RPP30 \| Polymerase/histidinol phosphatase-like protein \| 201606;A[illegible] | 4 | 3 | 3 | 3 | 1 | 2 | [illegible] |
| AT2G32810.2;AT2G32810.1 | AT2G32810.2;AT2G3281[illegible] | 2;2 | 2;2 | 2;2 | BGAL9, BGAL9 \| beta galactosidase 9 \| 201606;BGAL9, BGAL9 \| beta galactosidase 9 [illegible] | 2 | 2 | 2 | 2 | 2 | 2 | [illegible] |
| AT1G71820.1;AT1G71820.2 | AT1G71820.1;AT1G7182[illegible] | 2;2 | 2;2 | 2;2 | SEC6 \| SEC6 \| 201606;SEC6 \| SEC6 \| 201606 | 2 | 2 | 2 | 2 | 2 | 2 | [illegible] |
| AT1G30570.1 | AT1G30570.1 | 5 | 4 | 3 | HERK2 \| hercules receptor kinase 2 \| 201606 | 1 | 5 | 4 | 3 | 5 | 3 | [illegible] |
| AT5G28540.1 | AT5G28540.1 | 33 | 2 | 1 | BIP1 \| heat shock protein 70 (Hsp 70) family protein \| 201606 | 1 | 33 | 2 | 1 | 25 | 28 | [illegible] |
| AT1G17370.2;AT1G17370.1;AT1G54080.[illegible] | AT1G17370.2;AT1G1737[illegible] | 1;1;1;1;1 | 1;1;1;1;1 | 1;1;1;1;1 | UBP1B \| oligouridylate binding protein 1B \| 201606;UBP1B \| oligouridylate binding [illegible] | 5 | 1 | 1 | 1 | 1 | 1 | [illegible] |
| AT1G09870.1 | AT1G09870.1 | 1 | 1 | 1 | histidine acid phosphatase family protein \| 201606 | 1 | 1 | 1 | 1 | 1 | 1 | [illegible] |
| AT5G03900.2 | AT5G03900.2 | 1 | 1 | 1 | Iron-sulfur cluster biosynthesis family protein \| 201606 | 1 | 1 | 1 | 1 | 1 | 1 | [illegible] |
| [illegible] | [illegible] | [illegible] | [illegible] | [illegible] | [illegible] | [illegible] | [illegible] | [illegible] | [illegible] | [illegible] | [illegible] | [illegible] |

D9,1
2949;2950;2951;2952;2953;

**Supplemental Dataset 3. CCV dimethyl labeling LC/MS-MS data.** Complete list of accession numbers and protein labels corresponding to those proteins (n = 1,109) which were identified in at least one of two reciprocal dimethyl labeling replicates. The ratios indicating enrichment of the protein between the DFGL and CCV stages of the purification process are indicated for each replicate. Depletion between DFGL and CCVs is defined as 1 / fold enrichment.

| Accession Number | Label | Ratio Rep1 | Ratio Rep 2 | Average Fold Enrichment (CCV:DFGL) | Average Fold Depletion (CCV:DFGL) |
|---|---|---|---|---|---|
| AT5G25060 | AT5G25060.1 \| Symbols: \| RNA recognition motif (RRM)-containing protein \| | 58.136 | | 58.136 | 0.017201046 |
| AT4G33150 | AT4G33150.1 \| Symbols: \| lysine-ketoglutarate reductase/saccharopine | 40.652 | | 40.652 | 0.024599036 |
| AT3G59920 | AT3G59920.1 \| Symbols: ATGDI2, GDI2 \| RAB GDP dissociation inhibitor 2 \| chr3:22135157-22138221 FORWARD LENGTH=444 | | 35.79610538 | 35.79610538 | 0.027936 |
| AT3G49750 | AT3G49750.1 \| Symbols: AtRLP44, RLP44 \| receptor like protein 44 \| chr3:18450604-18451428 REVERSE LENGTH=274;AT5G65830.1 \| Symbols: ATRLP57, RLP57 \| receptor like protein 57 \| chr5:26342396-26343235 REVERSE LENGTH=279 | | 25.94101014 | 25.94101014 | 0.038549 |
| AT2G40060 | AT2G40060.1 \| Symbols: \| Clathrin light chain protein \| chr2:16726564-16728001 FORWARD LENGTH=258 | 4.4661 | 31.8745418 | 18.1703209 | 0.055034801 |
| AT5G11890 | AT5G11890.1 \| Symbols: \| FUNCTIONS IN: molecular_function unknown; LOCATED IN: plasma membrane; EXPRESSED IN: 12 plant structures; EXPRESSED DURING: 6 growth stages; BEST Arabidopsis thaliana protein match is: Late embryogenesis abundant (LEA) hydroxyproline-rich glycoprotein family (TAIR:AT1G17620.1); Has 1807 Blast hits to 1807 proteins in 277 species: Archae - 0; Bacteria - 0; Metazoa - 736; Fungi - 347; Plants - 385; Viruses - 0; Other Eukaryotes - 339 (source: NCBI BLink). \| chr5:3831770-3832633 FORWARD LENGTH=287 | | 15.77809685 | 15.77809685 | 0.063379 |
| AT5G18770 | AT5G18770.1 \| Symbols: \| F-box/FBD-like domains containing protein \| chr5:6261426-6263172 FORWARD LENGTH=481 | | 14.98531439 | 14.98531439 | 0.066732 |
| AT1G71410 | AT1G71410.1 \| Symbols: \| ARM repeat superfamily protein \| chr1:26913070-26917515 REVERSE LENGTH=909;AT1G22870.1 \| Symbols: \| Protein kinase family protein with ARM repeat domain \| chr1:8089501-8094173 FORWARD LENGTH=913 | | 14.93718912 | 14.93718912 | 0.066947 |
| AT1G22280 | AT1G22280.3 \| Symbols: PAPP2C \| phytochrome-associated protein phosphatase type 2C \| chr1:7874236-7875496 FORWARD LENGTH=287;AT1G22280.1 \| Symbols: PAPP2C \| phytochrome-associated protein phosphatase type 2C \| chr1:7874236-7875496 FORWARD LENGTH=281;AT1G22280.2 \| Symbols: \| phytochrome-associated protein phosphatase type 2C \| chr1:7874236-7875054 FORWARD LENGTH=199 | | 14.24947989 | 14.24947989 | 0.070178 |
| AT4G26120 | AT4G26120.1 \| Symbols: \| Ankyrin repeat family protein / BTB/POZ domain-containing protein \| chr4:13236448-13238487 FORWARD LENGTH=600 | | 12.92190004 | 12.92190004 | 0.077388 |
| AT4G27590 | AT4G27590.2 \| Symbols: \| Heavy metal transport/detoxification superfamily protein \| chr4:13771224-13772594 FORWARD LENGTH=332 | | 12.71278016 | 12.71278016 | 0.078661 |
| AT5G26980 | AT5G26980.1 \| Symbols: SYP41, ATSYP41, ATTLG2A \| syntaxin of plants 41 \| chr5:9488724-9490604 REVERSE LENGTH=322;AT5G26980.2 \| Symbols: SYP41, ATSYP41, ATTLG2A \| syntaxin of plants 41 \| chr5:9488724-9490604 REVERSE LENGTH=322 | | 12.54594954 | 12.54594954 | 0.079707 |
| AT1G69295 | AT1G69295.1 \| Symbols: PDCB4 \| plasmodesmata callose-binding protein 4 \| chr1:26050492-26051843 REVERSE LENGTH=222;AT1G26450.1 \| Symbols: \| Carbohydrate-binding X8 domain superfamily protein \| chr1:9149003-9150063 REVERSE LENGTH=197 | | 12.42637374 | 12.42637374 | 0.080474 |
| AT4G31140 | AT4G31140.1 \| Symbols: \| O-Glycosyl hydrolases family 17 protein \| chr4:15141581-15143188 FORWARD LENGTH=484;AT5G20870.1 \| Symbols: \| O-Glycosyl hydrolases family 17 protein \| chr5:7080037-7081739 REVERSE LENGTH=501 | | 11.82270669 | 11.82270669 | 0.084583 |
| AT1G67940 | AT1G67940.1 \| Symbols: ATNAP3, AtSTAR1, NAP3 \| non-intrinsic ABC protein 3 \| chr1:25477805-25478667 FORWARD LENGTH=263 | 0.12351 | 22.50427581 | 11.31389291 | 0.088386907 |

| | | | | | |
|---|---|---|---|---|---|
| AT5G15460 | AT5G15460.1 \| Symbols: MUB2 \| membrane-anchored ubiquitin-fold protein 2 \| chr5:5018947-5020105 REVERSE LENGTH=124;AT5G15460.2 \| Symbols: MUB2 \| membrane-anchored ubiquitin-fold protein 2 \| chr5:5018947-5020105 REVERSE LENGTH=124 | | 11.25163149 | 11.25163149 | 0.088876 |
| AT3G17420 | AT3G17420.1 \| Symbols: GPK1 \| glyoxysomal protein kinase 1 \| chr3:5959462-5961313 REVERSE LENGTH=467 | | 10.86732088 | 10.86732088 | 0.092019 |
| AT4G17615 | AT4G17615.1 \| Symbols: CBL1, ATCBL1, SCABP5 \| calcineurin B-like protein 1 \| chr4:9817142-9818995 FORWARD LENGTH=213;AT5G47100.1 \| Symbols: CBL9, ATCBL9 \| calcineurin B-like protein 9 \| chr5:19129896-19131727 REVERSE LENGTH=213;AT4G17615.2 \| Symbols: CBL1, ATCBL1, SCABP5 \| calcineurin B-like protein 1 \| chr4:9817981-9818995 FORWARD LENGTH=171 | | 10.68832835 | 10.68832835 | 0.09356 |
| AT1G78200 | AT1G78200.1 \| Symbols: \| Protein phosphatase 2C family protein \| chr1:29420483-29421650 FORWARD LENGTH=283;AT1G78200.2 \| Symbols: \| Protein phosphatase 2C family protein \| chr1:29420483-29421650 FORWARD LENGTH=283 | | 10.56568688 | 10.56568688 | 0.094646 |
| AT1G70520 | AT1G70520.1 \| Symbols: CRK2 \| cysteine-rich RLK (RECEPTOR-like protein kinase) 2 \| chr1:26584888-26587334 REVERSE LENGTH=649 | | 10.34821752 | 10.34821752 | 0.096635 |
| AT4G35410 | AT4G35410.2 \| Symbols: \| Clathrin adaptor complex small chain family protein \| chr4:16832572-16833796 FORWARD LENGTH=162;AT2G17380.1 \| Symbols: AP19 \| associated protein 19 \| chr2:7553122-7554887 FORWARD LENGTH=161;AT4G35410.1 \| Symbols: \| Clathrin adaptor complex small chain family protein \| chr4:16832572-16833476 FORWARD LENGTH=124 | 10.334 | | 10.334 | 0.09676795 |
| AT2G21720 | AT2G21720.1 \| Symbols: \| Plant protein of unknown function (DUF639) \| chr2:9273696-9276802 FORWARD LENGTH=734 | | 9.88533017 | 9.88533017 | 0.10116 |
| AT1G48210 | AT1G48210.1 \| Symbols: \| Protein kinase superfamily protein \| chr1:17799551-17801798 FORWARD LENGTH=363;AT1G48210.2 \| Symbols: \| Protein kinase superfamily protein \| chr1:17799551-17801798 FORWARD LENGTH=363 | | 9.670244657 | 9.670244657 | 0.10341 |
| AT3G02740 | AT3G02740.1 \| Symbols: \| Eukaryotic aspartyl protease family protein \| chr3:590561-593089 FORWARD LENGTH=488 | | 9.627418889 | 9.627418889 | 0.10387 |
| AT2G23810 | AT2G23810.1 \| Symbols: TET8 \| tetraspanin8 \| chr2:10135859-10137352 REVERSE LENGTH=273 | | 9.410878976 | 9.410878976 | 0.10626 |
| AT2G26550 | AT2G26550.1 \| Symbols: HO2 \| heme oxygenase 2 \| chr2:11291584-11293426 REVERSE LENGTH=354;AT2G26550.3 \| Symbols: HO2 \| heme oxygenase 2 \| chr2:11291584-11293426 REVERSE LENGTH=314;AT2G26550.2 \| Symbols: HO2 \| heme oxygenase 2 \| chr2:11291584-11293426 REVERSE LENGTH=299 | 9.122 | | 9.122 | 0.109625082 |
| AT4G28050 | AT4G28050.1 \| Symbols: TET7 \| tetraspanin7 \| chr4:13942566-13943621 REVERSE LENGTH=263 | | 9.098353198 | 9.098353198 | 0.10991 |
| AT3G62220 | AT3G62220.1 \| Symbols: \| Protein kinase superfamily protein \| chr3:23029276-23030864 REVERSE LENGTH=361;AT2G47060.4 \| Symbols: \| Protein kinase superfamily protein \| chr2:19333116-19334759 REVERSE LENGTH=397;AT2G47060.1 \| Symbols: \| Protein kinase superfamily protein \| chr2:19333116-19334759 REVERSE LENGTH=365;AT2G47060.2 \| Symbols: \| Protein kinase superfamily protein \| chr2:19333116-19334759 REVERSE LENGTH=365;AT2G47060.3 \| Symbols: \| Protein kinase superfamily protein \| chr2:19333116-19334393 REVERSE LENGTH=266 | | 8.954956568 | 8.954956568 | 0.11167 |
| AT1G64760 | AT1G64760.1 \| Symbols: \| O-Glycosyl hydrolases family 17 protein \| chr1:24054220-24056194 REVERSE LENGTH=481;AT1G64760.2 \| Symbols: \| O-Glycosyl hydrolases family 17 protein \| chr1:24054220-24056194 REVERSE LENGTH=481 | | 8.865248227 | 8.865248227 | 0.1128 |
| AT3G51890 | AT3G51890.1 \| Symbols: \| Clathrin light chain protein \| chr3:19249686-19250890 REVERSE LENGTH=258 | 11.073 | 6.485925542 | 8.779462771 | 0.113902186 |
| AT3G05710 | AT3G05710.1 \| Symbols: SYP43, ATSYP43 \| syntaxin of plants 43 \| chr3:1685262-1687229 FORWARD LENGTH=330;AT3G05710.2 \| Symbols: SYP43, ATSYP43 \| syntaxin of plants 43 \| chr3:1685262-1687229 FORWARD LENGTH=331 | | 8.724480893 | 8.724480893 | 0.11462 |

| | | | | | |
|---|---|---|---|---|---|
| AT5G43980 | AT5G43980.1 \| Symbols: PDLP1, PDLP1A \| plasmodesmata-located protein 1 \| chr5:17694817-17696458 REVERSE LENGTH=303 | | 8.528057309 | 8.528057309 | 0.11726 |
| AT3G55440 | AT3G55440.1 \| Symbols: ATCTIMC, TPI, CYTOTPI \| triosephosphate isomerase \| chr3:20553794-20556078 FORWARD LENGTH=254 | 3.237 | 13.47654408 | 8.356772038 | 0.119663429 |
| AT5G20230 | AT5G20230.1 \| Symbols: ATBCB, BCB, SAG14 \| blue-copper-binding protein \| chr5:6826626-6827408 FORWARD LENGTH=196 | 4.6751 | 12.0045137 | 8.339806849 | 0.119906854 |
| AT2G25270 | AT2G25270.1 \| Symbols: \| unknown protein; LOCATED IN: plasma membrane; EXPRESSED IN: 18 plant structures; EXPRESSED DURING: 13 growth stages; BEST Arabidopsis thaliana protein match is: unknown protein (TAIR:AT2G12400.1); Has 35333 Blast hits to 34131 proteins in 2444 species: Archae - 798; Bacteria - 22429; Metazoa - 974; Fungi - 991; Plants - 531; Viruses - 0; Other Eukaryotes - 9610 (source: NCBI BLink). \| chr2:10759779-10762358 FORWARD LENGTH=545 | | 8.321544479 | 8.321544479 | 0.12017 |
| AT4G34660 | AT4G34660.1 \| Symbols: \| SH3 domain-containing protein \| chr4:16545595-16548294 REVERSE LENGTH=368;AT4G34660.2 \| Symbols: \| SH3 domain-containing protein \| chr4:16545595-16548294 REVERSE LENGTH=345;AT4G34660.3 \| Symbols: \| SH3 domain-containing protein \| chr4:16545595-16548294 REVERSE LENGTH=317 | 7.1336 | 9.399379641 | 8.26648982 | 0.12097033 |
| AT3G58100 | AT3G58100.1 \| Symbols: PDCB5 \| plasmodesmata callose-binding protein 5 \| chr3:21515397-21516329 REVERSE LENGTH=180 | 13.013 | 3.416467373 | 8.214733686 | 0.121732492 |
| AT2G20760 | AT2G20760.1 \| Symbols: \| Clathrin light chain protein \| chr2:8943279-8945108 REVERSE LENGTH=338 | 3.5658 | 12.77481828 | 8.170309139 | 0.12239439 |
| AT3G45600 | AT3G45600.1 \| Symbols: TET3 \| tetraspanin3 \| chr3:16733973-16735854 REVERSE LENGTH=285 | 3.9072 | 12.41665322 | 8.161926608 | 0.122520092 |
| AT3G52470 | AT3G52470.1 \| Symbols: \| Late embryogenesis abundant (LEA) hydroxyproline-rich glycoprotein family \| chr3:19450750-19451376 FORWARD LENGTH=208 | | 8.095854922 | 8.095854922 | 0.12352 |
| AT3G54200 | AT3G54200.1 \| Symbols: \| Late embryogenesis abundant (LEA) hydroxyproline-rich glycoprotein family \| chr3:20065731-20066438 FORWARD LENGTH=235 | 3.0449 | 13.08232708 | 8.063613542 | 0.12401388 |
| AT3G17410 | AT3G17410.1 \| Symbols: \| Protein kinase superfamily protein \| chr3:5956601-5958882 FORWARD LENGTH=364;AT2G47060.5 \| Symbols: \| Protein kinase superfamily protein \| chr2:19333436-19334759 REVERSE LENGTH=322 | | 8.033419023 | 8.033419023 | 0.12448 |
| AT1G10730 | AT1G10730.1 \| Symbols: \| Clathrin adaptor complexes medium subunit family protein \| chr1:3565463-3567779 FORWARD LENGTH=428 | | 8.019246191 | 8.019246191 | 0.1247 |
| AT2G01630 | AT2G01630.1 \| Symbols: \| O-Glycosyl hydrolases family 17 protein \| chr2:279541-281955 REVERSE LENGTH=501;AT2G01630.2 \| Symbols: \| O-Glycosyl hydrolases family 17 protein \| chr2:279564-281955 REVERSE LENGTH=391 | | 7.954816641 | 7.954816641 | 0.12571 |
| AT4G29920 | AT4G29920.1 \| Symbols: \| Double Clp-N motif-containing P-loop nucleoside triphosphate hydrolases superfamily protein \| chr4:14632653-14635885 REVERSE LENGTH=1017 | | 7.742335088 | 7.742335088 | 0.12916 |
| AT1G72230 | AT1G72230.1 \| Symbols: \| Cupredoxin superfamily protein \| chr1:27188287-27189093 FORWARD LENGTH=181 | | 7.704160247 | 7.704160247 | 0.1298 |
| AT3G07040 | AT3G07040.1 \| Symbols: RPM1, RPS3 \| NB-ARC domain-containing disease resistance protein \| chr3:2226244-2229024 REVERSE LENGTH=926 | 7.6826 | | 7.6826 | 0.130164267 |
| AT5G67130 | AT5G67130.1 \| Symbols: \| PLC-like phosphodiesterases superfamily protein \| chr5:26791599-26793649 FORWARD LENGTH=426 | | 7.676364474 | 7.676364474 | 0.13027 |
| AT5G08080 | AT5G08080.3 \| Symbols: SYP132 \| syntaxin of plants 132 \| chr5:2588532-2591106 FORWARD LENGTH=315;AT5G08080.1 \| Symbols: SYP132, ATSYP132 \| syntaxin of plants 132 \| chr5:2588532-2591106 FORWARD LENGTH=304;AT5G08080.2 \| Symbols: SYP132, ATSYP132 \| syntaxin of plants 132 \| chr5:2588532-2590488 FORWARD LENGTH=223 | | 7.630093087 | 7.630093087 | 0.13106 |
| AT3G03610 | AT3G03610.1 \| Symbols: \| ELMO/CED-12 family protein \| chr3:869479-871687 FORWARD LENGTH=323 | | 7.256367462 | 7.256367462 | 0.13781 |
| AT2G41410 | AT2G41410.1 \| Symbols: \| Calcium-binding EF-hand family protein \| chr2:17262085-17262735 REVERSE LENGTH=216 | | 7.224389539 | 7.224389539 | 0.13842 |
| AT3G47730 | AT3G47730.1 \| Symbols: ATATH1, ATH1, ABCA2 \| ATP-binding cassette A2 \| chr3:17594342-17598828 REVERSE LENGTH=983 | | 7.151029748 | 7.151029748 | 0.13984 |

| | | | | | |
|---|---|---|---|---|---|
| AT2G39450 | AT2G39450.1 \| Symbols: MTP11, ATMTP11 \| Cation efflux family protein \| chr2:16471744-16473735 REVERSE LENGTH=394 | | 7.114905727 | 7.114905727 | 0.14055 |
| AT5G38480 | AT5G38480.1 \| Symbols: GRF3, RCI1 \| general regulatory factor 3 \| chr5:15410277-15411285 FORWARD LENGTH=255;AT5G38480.2 \| Symbols: GRF3, RCI1 \| general regulatory factor 3 \| chr5:15410277-15411285 FORWARD LENGTH=254 | | 7.015574576 | 7.015574576 | 0.14254 |
| AT2G17380 | AT2G17380.1 \| Symbols: AP19 \| associated protein 19 \| chr2:7553122-7554887 FORWARD LENGTH=161;AT4G35410.2 \| Symbols: \| Clathrin adaptor complex small chain family protein \| chr4:16832572-16833796 FORWARD LENGTH=162;AT4G35410.1 \| Symbols: \| Clathrin adaptor complex small chain family protein \| chr4:16832572-16833476 FORWARD LENGTH=124 | | 7.010656197 | 7.010656197 | 0.14264 |
| AT5G47310 | AT5G47310.1 \| Symbols: \| PPPDE putative thiol peptidase family protein \| chr5:19201325-19202674 FORWARD LENGTH=245;AT4G17486.1 \| Symbols: \| PPPDE putative thiol peptidase family protein \| chr4:9749992-9751201 REVERSE LENGTH=224;AT4G17486.2 \| Symbols: \| PPPDE putative thiol peptidase family protein \| chr4:9750063-9751201 REVERSE LENGTH=199 | | 6.949270327 | 6.949270327 | 0.1439 |
| AT1G53840 | AT1G53840.1 \| Symbols: ATPME1, PME1 \| pectin methylesterase 1 \| chr1:20101533-20103458 FORWARD LENGTH=586 | | 6.937699459 | 6.937699459 | 0.14414 |
| AT4G10740 | AT4G10740.1 \| Symbols: \| F-box and associated interaction domains-containing protein \| chr4:6617143-6618426 REVERSE LENGTH=427 | | 6.908462867 | 6.908462867 | 0.14475 |
| AT3G11820 | AT3G11820.1 \| Symbols: SYP121, AT-SYR1, ATSYP121, SYR1, ATSYR1, PEN1 \| syntaxin of plants 121 \| chr3:3729540-3730868 REVERSE LENGTH=346;AT3G11820.2 \| Symbols: SYP121, AT-SYR1, ATSYP121, SYR1, ATSYR1, PEN1 \| syntaxin of plants 121 \| chr3:3729540-3730775 REVERSE LENGTH=315 | | 6.862005078 | 6.862005078 | 0.14573 |
| AT2G42910 | AT2G42910.1 \| Symbols: \| Phosphoribosyltransferase family protein \| chr2:17856396-17858394 FORWARD LENGTH=337 | | 6.852600562 | 6.852600562 | 0.14593 |
| AT1G76680 | AT1G76680.2 \| Symbols: OPR1 \| 12-oxophytodienoate reductase 1 \| chr1:28776982-28778271 FORWARD LENGTH=397;AT1G76680.1 \| Symbols: OPR1, ATOPR1 \| 12-oxophytodienoate reductase 1 \| chr1:28776982-28778271 FORWARD LENGTH=372;AT1G76690.1 \| Symbols: OPR2, ATOPR2 \| 12-oxophytodienoate reductase 2 \| chr1:28778976-28780355 FORWARD LENGTH=374 | | 6.825006825 | 6.825006825 | 0.14652 |
| AT1G06700 | AT1G06700.1 \| Symbols: \| Protein kinase superfamily protein \| chr1:2052750-2054552 REVERSE LENGTH=361;AT1G06700.2 \| Symbols: \| Protein kinase superfamily protein \| chr1:2052750-2054552 REVERSE LENGTH=361;AT2G43230.2 \| Symbols: \| Protein kinase superfamily protein \| chr2:17966475-17968446 FORWARD LENGTH=440;AT2G43230.1 \| Symbols: \| Protein kinase superfamily protein \| chr2:17966475-17968446 FORWARD LENGTH=406;AT2G30730.1 \| Symbols: \| Protein kinase superfamily protein \| chr2:13093145-13094677 FORWARD LENGTH=338;AT4G13190.1 \| Symbols: \| Protein kinase superfamily protein \| chr4:7659435-7661106 REVERSE LENGTH=389;AT2G41970.1 \| Symbols: \| Protein kinase superfamily protein \| chr2:17520517-17522304 REVERSE LENGTH=365 | | 6.763611769 | 6.763611769 | 0.14785 |
| AT5G07910 | AT5G07910.1 \| Symbols: \| Leucine-rich repeat (LRR) family protein \| chr5:2521937-2523769 REVERSE LENGTH=262 | | 6.758126647 | 6.758126647 | 0.14797 |
| AT3G16270 | AT3G16270.1 \| Symbols: \| ENTH/VHS family protein \| chr3:5513701-5516540 FORWARD LENGTH=690 | | 6.741724533 | 6.741724533 | 0.14833 |
| AT4G13350 | AT4G13350.1 \| Symbols: NIG \| NSP (nuclear shuttle protein)-interacting GTPase \| chr4:7770170-7773321 REVERSE LENGTH=602;AT4G13350.2 \| Symbols: NIG \| NSP (nuclear shuttle protein)-interacting GTPase \| chr4:7770170-7773321 REVERSE LENGTH=602 | | 6.615506748 | 6.615506748 | 0.15116 |
| AT1G26670 | AT1G26670.1 \| Symbols: VTI12, ATVTI12, VTI1B \| Vesicle transport v-SNARE family protein \| chr1:9216103-9217793 FORWARD LENGTH=222 | 6.5906 | | 6.5906 | 0.151731254 |

| | | | | | |
|---|---|---|---|---|---|
| AT2G01660 | AT2G01660.1 \| Symbols: PDLP6 \| plasmodesmata-located protein 6 \| chr2:290973-292496 REVERSE LENGTH=288;AT2G01660.2 \| Symbols: PDLP6 \| plasmodesmata-located protein 6 \| chr2:291717-292496 REVERSE LENGTH=259 | | 6.50026001 | 6.50026001 | 0.15384 |
| AT3G61260 | AT3G61260.1 \| Symbols: \| Remorin family protein \| chr3:22675403-22676701 REVERSE LENGTH=212 | | 6.4499484 | 6.4499484 | 0.15504 |
| AT3G11650 | AT3G11650.1 \| Symbols: NHL2 \| NDR1/HIN1-like 2 \| chr3:3676264-3676986 REVERSE LENGTH=240 | | 6.393453104 | 6.393453104 | 0.15641 |
| AT4G32760 | AT4G32760.2 \| Symbols: \| ENTH/VHS/GAT family protein \| chr4:15799376-15803832 FORWARD LENGTH=676;AT4G32760.1 \| Symbols: \| ENTH/VHS/GAT family protein \| chr4:15799376-15803832 FORWARD LENGTH=675 | 8.8761 | 3.859513701 | 6.367806851 | 0.157039939 |
| AT2G34520 | AT2G34520.1 \| Symbols: RPS14 \| mitochondrial ribosomal protein S14 \| chr2:14548218-14548712 REVERSE LENGTH=164 | | 6.364562118 | 6.364562118 | 0.15712 |
| AT1G22300 | AT1G22300.1 \| Symbols: GRF10, 14-3-3EPSILON, GF14 EPSILON \| general regulatory factor 10 \| chr1:7879146-7881103 REVERSE LENGTH=254;AT1G22300.2 \| Symbols: GRF10, GF14 EPSILON \| general regulatory factor 10 \| chr1:7879244-7881103 REVERSE LENGTH=254;AT1G22300.3 \| Symbols: GRF10, GF14 EPSILON \| general regulatory factor 10 \| chr1:7879601-7881103 REVERSE LENGTH=251 | | 6.35768326 | 6.35768326 | 0.15729 |
| AT4G12420 | AT4G12420.1 \| Symbols: SKU5 \| Cupredoxin superfamily protein \| chr4:7349941-7352868 REVERSE LENGTH=587;AT4G12420.2 \| Symbols: SKU5 \| Cupredoxin superfamily protein \| chr4:7349941-7352868 REVERSE LENGTH=587 | 4.8614 | 7.8015291 | 6.33146455 | 0.157941341 |
| AT5G55730 | AT5G55730.1 \| Symbols: FLA1 \| FASCICLIN-like arabinogalactan 1 \| chr5:22558375-22560392 REVERSE LENGTH=424;AT5G55730.2 \| Symbols: FLA1 \| FASCICLIN-like arabinogalactan 1 \| chr5:22558375-22560392 REVERSE LENGTH=424 | | 6.313928526 | 6.313928526 | 0.15838 |
| AT1G08560 | AT1G08560.1 \| Symbols: SYP111, KN, ATSYP111 \| syntaxin of plants 111 \| chr1:2709778-2710710 REVERSE LENGTH=310 | 1.4149 | 11.15262365 | 6.283761827 | 0.159140341 |
| AT5G11490 | AT5G11490.2 \| Symbols: \| adaptin family protein \| chr5:3671964-3676263 FORWARD LENGTH=850;AT5G11490.1 \| Symbols: \| adaptin family protein \| chr5:3671964-3676144 FORWARD LENGTH=841 | | 6.271558482 | 6.271558482 | 0.15945 |
| AT3G15890 | AT3G15890.1 \| Symbols: \| Protein kinase superfamily protein \| chr3:5374389-5376114 FORWARD LENGTH=361;AT1G52540.1 \| Symbols: \| Protein kinase superfamily protein \| chr1:19570298-19571884 REVERSE LENGTH=350 | | 6.255082254 | 6.255082254 | 0.15987 |
| AT5G51480 | AT5G51480.1 \| Symbols: SKS2 \| SKU5 similar 2 \| chr5:20910433-20913153 FORWARD LENGTH=592 | 5.212 | 7.214486689 | 6.213243345 | 0.160946537 |
| AT2G25430 | AT2G25430.1 \| Symbols: \| epsin N-terminal homology (ENTH) domain-containing protein / clathrin assembly protein-related \| chr2:10822716-10824677 FORWARD LENGTH=653 | 3.7784 | 8.608815427 | 6.193607713 | 0.161456787 |
| AT2G45010 | AT2G45010.1 \| Symbols: \| PLAC8 family protein \| chr2:18568045-18569615 FORWARD LENGTH=244;AT2G45010.2 \| Symbols: \| PLAC8 family protein \| chr2:18568782-18569615 FORWARD LENGTH=198 | | 6.164848036 | 6.164848036 | 0.16221 |
| AT4G16370 | AT4G16370.1 \| Symbols: ATOPT3, OPT3 \| oligopeptide transporter \| chr4:9247514-9250071 REVERSE LENGTH=737 | 6.6996 | 5.611987205 | 6.155793602 | 0.162448591 |
| AT3G11130 | AT3G11130.1 \| Symbols: \| Clathrin, heavy chain \| chr3:3482575-3491667 REVERSE LENGTH=1705 | 7.34 | 4.878524734 | 6.109262367 | 0.163685882 |
| AT1G03870 | AT1G03870.1 \| Symbols: FLA9 \| FASCICLIN-like arabinoogalactan 9 \| chr1:982625-983368 REVERSE LENGTH=247 | | 6.069434329 | 6.069434329 | 0.16476 |
| AT3G17440 | AT3G17440.1 \| Symbols: NPSN13, ATNPSN13 \| novel plant snare 13 \| chr3:5969909-5972290 REVERSE LENGTH=269;AT1G48240.1 \| Symbols: ATNPSN12, NPSN12 \| novel plant snare 12 \| chr1:17809223-17811656 REVERSE LENGTH=265;AT3G17440.2 \| Symbols: NPSN13, ATNPSN13 \| novel plant snare 13 \| chr3:5970153-5972290 REVERSE LENGTH=216 | | 6.054368227 | 6.054368227 | 0.16517 |
| AT3G58460 | AT3G58460.2 \| Symbols: RBL15 \| RHOMBOID-like protein 15 \| chr3:21623374-21626642 REVERSE LENGTH=426;AT3G58460.1 \| Symbols: ATRBL15, RBL15 \| RHOMBOID-like protein 15 \| chr3:21623374-21626348 REVERSE LENGTH=403 | 10.151 | 1.946926776 | 6.048963388 | 0.165317582 |

| | | | | | |
|---|---|---|---|---|---|
| AT5G24010 | AT5G24010.1 \| Symbols: \| Protein kinase superfamily protein \| chr5:8113910-8116384 FORWARD LENGTH=824 | 7.6016 | 4.492564805 | 6.047082403 | 0.165369005 |
| AT1G62380 | AT1G62380.1 \| Symbols: ACO2, ATACO2 \| ACC oxidase 2 \| chr1:23082340-23084068 FORWARD LENGTH=320 | 2.2532 | 9.639483324 | 5.946341662 | 0.168170626 |
| AT4G32285 | AT4G32285.1 \| Symbols: \| ENTH/ANTH/VHS superfamily protein \| chr4:15586003-15587910 FORWARD LENGTH=635;AT4G32285.2 \| Symbols: \| ENTH/ANTH/VHS superfamily protein \| chr4:15586003-15587910 FORWARD LENGTH=635 | 6.0415 | 5.77700751 | 5.909253755 | 0.169226106 |
| AT4G25240 | AT4G25240.1 \| Symbols: SKS1 \| SKU5 similar 1 \| chr4:12930539-12933563 FORWARD LENGTH=589 | 3.9573 | 7.805791898 | 5.881545949 | 0.170023325 |
| AT4G29350 | AT4G29350.1 \| Symbols: PFN2, PRO2, PRF2 \| profilin 2 \| chr4:14450135-14451119 FORWARD LENGTH=131 | 5.8811 | | 5.8811 | 0.170036218 |
| AT4G36750 | AT4G36750.1 \| Symbols: \| Quinone reductase family protein \| chr4:17324642-17326215 FORWARD LENGTH=273 | | 5.849663644 | 5.849663644 | 0.17095 |
| AT3G01290 | AT3G01290.1 \| Symbols: \| SPFH/Band 7/PHB domain-containing membrane-associated protein family \| chr3:88252-89356 REVERSE LENGTH=285 | | 5.828185103 | 5.828185103 | 0.17158 |
| AT1G75680 | AT1G75680.1 \| Symbols: AtGH9B7, GH9B7 \| glycosyl hydrolase 9B7 \| chr1:28417215-28419231 REVERSE LENGTH=525 | | 5.782686636 | 5.782686636 | 0.17293 |
| AT5G11680 | AT5G11680.1 \| Symbols: \| FUNCTIONS IN: molecular_function unknown; INVOLVED IN: biological_process unknown; LOCATED IN: cytosol, plasma membrane; EXPRESSED IN: 26 plant structures; EXPRESSED DURING: 15 growth stages; CONTAINS InterPro DOMAIN/s: WW-domain-binding protein (InterPro:IPR018826); Has 1807 Blast hits to 1807 proteins in 277 species: Archae - 0; Bacteria - 0; Metazoa - 736; Fungi - 347; Plants - 385; Viruses - 0; Other Eukaryotes - 339 (source: NCBI BLink). \| chr5:3759463-3760634 FORWARD LENGTH=207 | | 5.767677933 | 5.767677933 | 0.17338 |
| AT4G29360 | AT4G29360.1 \| Symbols: \| O-Glycosyl hydrolases family 17 protein \| chr4:14451562-14453687 REVERSE LENGTH=534;AT4G29360.2 \| Symbols: \| O-Glycosyl hydrolases family 17 protein \| chr4:14451881-14453687 REVERSE LENGTH=488 | 3.4091 | 8.074935401 | 5.7420177 | 0.17415481 |
| AT1G47830 | AT1G47830.1 \| Symbols: \| SNARE-like superfamily protein \| chr1:17613346-17614784 REVERSE LENGTH=142 | | 5.714612264 | 5.714612264 | 0.17499 |
| AT1G07750 | AT1G07750.1 \| Symbols: \| RmlC-like cupins superfamily protein \| chr1:2404300-2405863 REVERSE LENGTH=356;AT2G28680.1 \| Symbols: \| RmlC-like cupins superfamily protein \| chr2:12303118-12304747 REVERSE LENGTH=356 | | 5.70711106 | 5.70711106 | 0.17522 |
| AT1G60070 | AT1G60070.2 \| Symbols: \| Adaptor protein complex AP-1, gamma subunit \| chr1:22142944-22149296 REVERSE LENGTH=898;AT1G60070.1 \| Symbols: \| Adaptor protein complex AP-1, gamma subunit \| chr1:22142944-22149296 REVERSE LENGTH=862 | 5.057 | 6.19885941 | 5.627929705 | 0.177685233 |
| AT1G28490 | AT1G28490.1 \| Symbols: SYP61, ATSYP61, OSM1 \| syntaxin of plants 61 \| chr1:10016433-10017842 FORWARD LENGTH=245;AT1G28490.2 \| Symbols: SYP61, ATSYP61, OSM1 \| syntaxin of plants 61 \| chr1:10016634-10017842 FORWARD LENGTH=206 | 3.6537 | 7.53806724 | 5.59588362 | 0.178702787 |
| AT4G11380 | AT4G11380.2 \| Symbols: \| Adaptin family protein \| chr4:6920608-6925444 FORWARD LENGTH=916;AT4G11380.1 \| Symbols: \| Adaptin family protein \| chr4:6920608-6925444 FORWARD LENGTH=894 | 3.5253 | 7.645259939 | 5.585279969 | 0.179042054 |
| AT1G05150 | AT1G05150.1 \| Symbols: \| Calcium-binding tetratricopeptide family protein \| chr1:1484280-1486706 REVERSE LENGTH=808 | | 5.58347292 | 5.58347292 | 0.1791 |
| AT1G48480 | AT1G48480.1 \| Symbols: RKL1 \| receptor-like kinase 1 \| chr1:17918475-17920743 FORWARD LENGTH=655 | | 5.566069242 | 5.566069242 | 0.17966 |
| AT3G09740 | AT3G09740.1 \| Symbols: SYP71, ATSYP71 \| syntaxin of plants 71 \| chr3:2989615-2991354 FORWARD LENGTH=266 | 2.6579 | 8.4324142 | 5.5451571 | 0.180337542 |

| | | | | | |
|---|---|---|---|---|---|
| AT3G59770 | AT3G59770.3 \| Symbols: SAC9 \| sacI homology domain-containing protein / WW domain-containing protein \| chr3:22079281-22085674 REVERSE LENGTH=1646;AT3G59770.1 \| Symbols: SAC9 \| sacI homology domain-containing protein / WW domain-containing protein \| chr3:22079281-22085674 REVERSE LENGTH=1630;AT3G59770.2 \| Symbols: SAC9 \| sacI homology domain-containing protein / WW domain-containing protein \| chr3:22079281-22084785 REVERSE LENGTH=1405 | 5.8017 | 5.250721974 | 5.526210987 | 0.180955813 |
| AT3G24550 | AT3G24550.1 \| Symbols: ATPERK1, PERK1 \| proline extensin-like receptor kinase 1 \| chr3:8960411-8963303 FORWARD LENGTH=652 | | 5.503577325 | 5.503577325 | 0.1817 |
| AT4G07400 | AT4G07400.1 \| Symbols: VFB3 \| VIER F-box proteine 3 \| chr4:4197847-4199511 REVERSE LENGTH=554 | 5.4969 | | 5.4969 | 0.181920719 |
| AT1G35720 | AT1G35720.1 \| Symbols: ANNAT1, OXY5, ATOXY5 \| annexin 1 \| chr1:13225304-13226939 FORWARD LENGTH=317 | | 5.494203615 | 5.494203615 | 0.18201 |
| AT3G53990 | AT3G53990.1 \| Symbols: \| Adenine nucleotide alpha hydrolases-like superfamily protein \| chr3:19989658-19991019 REVERSE LENGTH=160;AT3G53990.2 \| Symbols: \| Adenine nucleotide alpha hydrolases-like superfamily protein \| chr3:19990558-19991019 REVERSE LENGTH=126 | 8.8876 | 2.067824648 | 5.477712324 | 0.182557962 |
| AT1G07890 | AT1G07890.1 \| Symbols: APX1, MEE6, CS1, ATAPX1, ATAPX01 \| ascorbate peroxidase 1 \| chr1:2438005-2439435 FORWARD LENGTH=250;AT1G07890.2 \| Symbols: APX1, MEE6, CS1, ATAPX1, ATAPX01 \| ascorbate peroxidase 1 \| chr1:2438005-2439435 FORWARD LENGTH=250;AT1G07890.3 \| Symbols: APX1, MEE6, CS1, ATAPX1, ATAPX01 \| ascorbate peroxidase 1 \| chr1:2438005-2439435 FORWARD LENGTH=250;AT1G07890.4 \| Symbols: APX1, MEE6, CS1, ATAPX1, ATAPX01 \| ascorbate peroxidase 1 \| chr1:2438005-2439435 FORWARD LENGTH=250;AT1G07890.5 \| Symbols: APX1, MEE6, CS1, ATAPX1, ATAPX01 \| ascorbate peroxidase 1 \| chr1:2438005-2439435 FORWARD LENGTH=250;AT1G07890.7 \| Symbols: APX1, MEE6, CS1, ATAPX1, ATAPX01 \| ascorbate peroxidase 1 \| chr1:2438005-2439435 FORWARD LENGTH=250;AT1G07890.8 \| Symbols: APX1, MEE6, CS1, ATAPX1, ATAPX01 \| ascorbate peroxidase 1 \| chr1:2438005-2439435 FORWARD LENGTH=250;AT1G07890.6 \| Symbols: APX1, MEE6, CS1, ATAPX1, ATAPX01 \| ascorbate peroxidase 1 \| chr1:2438005-2439349 FORWARD LENGTH=249 | 0.86587 | 9.978048294 | 5.421959147 | 0.184435178 |
| AT1G71860 | AT1G71860.1 \| Symbols: PTP1, ATPTP1 \| protein tyrosine phosphatase 1 \| chr1:27026866-27028675 FORWARD LENGTH=340;AT1G71860.3 \| Symbols: PTP1, ATPTP1 \| protein tyrosine phosphatase 1 \| chr1:27026866-27028675 FORWARD LENGTH=340;AT1G71860.2 \| Symbols: PTP1, ATPTP1 \| protein tyrosine phosphatase 1 \| chr1:27026866-27028395 FORWARD LENGTH=277 | | 5.419466724 | 5.419466724 | 0.18452 |
| AT4G04720 | AT4G04720.1 \| Symbols: CPK21 \| calcium-dependent protein kinase 21 \| chr4:2394817-2397631 REVERSE LENGTH=531;AT4G21940.2 \| Symbols: CPK15 \| calcium-dependent protein kinase 15 \| chr4:11640847-11643487 FORWARD LENGTH=561;AT4G21940.1 \| Symbols: CPK15 \| calcium-dependent protein kinase 15 \| chr4:11640847-11643387 FORWARD LENGTH=554;AT3G20410.1 \| Symbols: CPK9 \| calmodulin-domain protein kinase 9 \| chr3:7116388-7118824 FORWARD LENGTH=541 | 3.9014 | 6.802721088 | 5.352060544 | 0.186843925 |
| AT2G19440 | AT2G19440.1 \| Symbols: \| O-Glycosyl hydrolases family 17 protein \| chr2:8418164-8419806 REVERSE LENGTH=478 | | 5.345878328 | 5.345878328 | 0.18706 |
| AT1G11130 | AT1G11130.1 \| Symbols: SUB, SCM, SRF9 \| Leucine-rich repeat protein kinase family protein \| chr1:3723135-3727178 FORWARD LENGTH=768 | | 5.321979776 | 5.321979776 | 0.1879 |
| AT3G59760 | AT3G59760.1 \| Symbols: OASC, ATCS-C \| O-acetylserine (thiol) lyase isoform C \| chr3:22072119-22075345 REVERSE LENGTH=433;AT3G59760.2 \| Symbols: OASC, ATCS-C \| O-acetylserine (thiol) lyase isoform C \| chr3:22072661-22075345 REVERSE LENGTH=432;AT3G59760.3 \| Symbols: OASC, ATCS-C \| O-acetylserine (thiol) lyase isoform C \| chr3:22072668-22075345 REVERSE LENGTH=430 | 5.3123 | | 5.3123 | 0.188242381 |

| | | | | | |
|---|---|---|---|---|---|
| AT3G54300 | AT3G54300.1 \| Symbols: ATVAMP727, VAMP727 \| vesicle-associated membrane protein 727 \| chr3:20108355-20110127 REVERSE LENGTH=240;AT3G54300.2 \| Symbols: ATVAMP727, VAMP727 \| vesicle-associated membrane protein 727 \| chr3:20108355-20110127 REVERSE LENGTH=240 | 2.5416 | 8.06321561 | 5.302407805 | 0.188593567 |
| AT4G38580 | AT4G38580.1 \| Symbols: ATFP6, HIPP26, FP6 \| farnesylated protein 6 \| chr4:18034596-18035693 FORWARD LENGTH=153 | | 5.290725358 | 5.290725358 | 0.18901 |
| AT5G43330 | AT5G43330.1 \| Symbols: \| Lactate/malate dehydrogenase family protein \| chr5:17390552-17392449 FORWARD LENGTH=332 | | 5.262603936 | 5.262603936 | 0.19002 |
| AT4G22753 | AT4G22753.2 \| Symbols: SMO1-3 \| sterol 4-alpha methyl oxidase 1-3 \| chr4:11949366-11953609 REVERSE LENGTH=581;AT4G22750.2 \| Symbols: \| DHHC-type zinc finger family protein \| chr4:11949366-11951335 REVERSE LENGTH=324;AT4G22750.1 \| Symbols: \| DHHC-type zinc finger family protein \| chr4:11949366-11951335 REVERSE LENGTH=302 | 1.8869 | 8.611780916 | 5.249340458 | 0.190500122 |
| AT1G77140 | AT1G77140.1 \| Symbols: VPS45, ATVPS45 \| vacuolar protein sorting 45 \| chr1:28984163-28987681 FORWARD LENGTH=569 | | 5.247966413 | 5.247966413 | 0.19055 |
| AT3G08530 | AT3G08530.1 \| Symbols: \| Clathrin, heavy chain \| chr3:2587171-2595411 REVERSE LENGTH=1703;REV__AT1G69270.1 \| Symbols: RPK1 \| receptor-like protein kinase 1 \| chr1:26040877-26042499 REVERSE LENGTH=540 | 6.8171 | 3.673094582 | 5.245097291 | 0.190654233 |
| AT5G42720 | AT5G42720.1 \| Symbols: \| Glycosyl hydrolase family 17 protein \| chr5:17130535-17132763 FORWARD LENGTH=438 | | 5.235602094 | 5.235602094 | 0.191 |
| AT5G47480 | AT5G47480.1 \| Symbols: \| RGPR-related \| chr5:19257420-19263132 FORWARD LENGTH=1350 | | 5.175715543 | 5.175715543 | 0.19321 |
| AT3G52400 | AT3G52400.1 \| Symbols: SYP122, ATSYP122 \| syntaxin of plants 122 \| chr3:19425835-19427032 REVERSE LENGTH=341 | | 5.152779925 | 5.152779925 | 0.19407 |
| AT1G54920 | AT1G54920.3 \| Symbols: \| unknown protein; FUNCTIONS IN: molecular_function unknown; INVOLVED IN: biological_process unknown; EXPRESSED IN: 23 plant structures; EXPRESSED DURING: 15 growth stages. \| chr1:20470916-20474898 FORWARD LENGTH=901;AT1G54920.2 \| Symbols: \| unknown protein; FUNCTIONS IN: molecular_function unknown; INVOLVED IN: biological_process unknown; LOCATED IN: membrane; EXPRESSED IN: 23 plant structures; EXPRESSED DURING: 15 growth stages; Has 35333 Blast hits to 34131 proteins in 2444 species: Archae - 798; Bacteria - 22429; Metazoa - 974; Fungi - 991; Plants - 531; Viruses - 0; Other Eukaryotes - 9610 (source: NCBI BLink). \| chr1:20470916-20474898 FORWARD LENGTH=890;AT1G54920.1 \| Symbols: \| unknown protein; FUNCTIONS IN: molecular_function unknown; INVOLVED IN: biological_process unknown; LOCATED IN: membrane; EXPRESSED IN: 23 plant structures; EXPRESSED DURING: 15 growth stages; Has 30201 Blast hits to 17322 proteins in 780 species: Archae - 12; Bacteria - 1396; Metazoa - 17338; Fungi - 3422; Plants - 5037; Viruses - 0; Other Eukaryotes - 2996 (source: NCBI BLink). \| chr1:20470916-20474421 FORWARD LENGTH=766 | 4.8317 | 5.4007345 | 5.11621725 | 0.195456907 |
| AT3G49490 | AT3G49490.1 \| Symbols: \| unknown protein; Has 722 Blast hits to 186 proteins in 64 species: Archae - 0; Bacteria - 30; Metazoa - 72; Fungi - 48; Plants - 38; Viruses - 0; Other Eukaryotes - 534 (source: NCBI BLink). \| chr3:18344926-18348648 REVERSE LENGTH=953 | | 5.081559022 | 5.081559022 | 0.19679 |
| AT4G09000 | AT4G09000.2 \| Symbols: GRF1, GF14 CHI \| general regulatory factor 1 \| chr4:5775387-5777157 FORWARD LENGTH=318;AT4G09000.1 \| Symbols: GRF1, GF14 CHI \| general regulatory factor 1 \| chr4:5775387-5777157 FORWARD LENGTH=267 | | 5.076399817 | 5.076399817 | 0.19699 |
| AT5G11710 | AT5G11710.1 \| Symbols: \| ENTH/VHS family protein \| chr5:3772981-3776316 FORWARD LENGTH=560 | 3.3084 | 6.655574043 | 4.981987022 | 0.200723124 |
| AT1G79550 | AT1G79550.1 \| Symbols: PGK \| phosphoglycerate kinase \| chr1:29924347-29926295 REVERSE LENGTH=401;AT1G79550.2 \| Symbols: PGK \| phosphoglycerate kinase \| chr1:29924347-29926295 REVERSE LENGTH=401 | 2.058 | 7.854225573 | 4.956112787 | 0.201771034 |
| AT1G74790 | AT1G74790.1 \| Symbols: \| catalytics \| chr1:28098912-28101673 FORWARD LENGTH=695 | | 4.93485985 | 4.93485985 | 0.20264 |

| | | | | | |
|---|---|---|---|---|---|
| AT5G06320 | AT5G06320.1 \| Symbols: NHL3 \| NDR1/HIN1-like 3 \| chr5:1931016-1931711 REVERSE LENGTH=231 | 2.8496 | 7.014590348 | 4.932095174 | 0.202753589 |
| AT1G23900 | AT1G23900.1 \| Symbols: GAMMA-ADAPTIN 1, Gamma-ADR \| gamma-adaptin 1 \| chr1:8441379-8447152 FORWARD LENGTH=876;AT1G23900.2 \| Symbols: GAMMA-ADAPTIN 1 \| gamma-adaptin 1 \| chr1:8441379-8447152 FORWARD LENGTH=876;AT1G23940.1 \| Symbols: \| ARM repeat superfamily protein \| chr1:8466094-8469346 FORWARD LENGTH=495 | 4.2495 | 5.605695387 | 4.927597693 | 0.202938645 |
| AT2G16520 | AT2G16520.1 \| Symbols: \| FUNCTIONS IN: molecular_function unknown; INVOLVED IN: biological_process unknown; LOCATED IN: endomembrane system; BEST Arabidopsis thaliana protein match is: zinc finger (C3HC4-type RING finger) family protein (TAIR:AT3G45560.1); Has 72 Blast hits to 69 proteins in 3 species: Archae - 0; Bacteria - 0; Metazoa - 0; Fungi - 0; Plants - 72; Viruses - 0; Other Eukaryotes - 0 (source: NCBI BLink). \| chr2:7161922-7162384 REVERSE LENGTH=99 | 4.8855 | | 4.8855 | 0.20468734 |
| AT1G14140 | AT1G14140.1 \| Symbols: \| Mitochondrial substrate carrier family protein \| chr1:4838131-4839602 REVERSE LENGTH=305 | 4.8606 | | 4.8606 | 0.205735917 |
| AT5G58090 | AT5G58090.1 \| Symbols: \| O-Glycosyl hydrolases family 17 protein \| chr5:23505556-23507193 REVERSE LENGTH=477 | | 4.842615012 | 4.842615012 | 0.2065 |
| AT1G17620 | AT1G17620.1 \| Symbols: \| Late embryogenesis abundant (LEA) hydroxyproline-rich glycoprotein family \| chr1:6062313-6063107 FORWARD LENGTH=264 | | 4.789501413 | 4.789501413 | 0.20879 |
| AT3G47670 | AT3G47670.1 \| Symbols: \| Plant invertase/pectin methylesterase inhibitor superfamily protein \| chr3:17575019-17575849 REVERSE LENGTH=276 | 1.2124 | 8.304268394 | 4.758334197 | 0.21015758 |
| AT2G01600 | AT2G01600.1 \| Symbols: \| ENTH/ANTH/VHS superfamily protein \| chr2:268975-272356 FORWARD LENGTH=571 | 2.6564 | 6.791632709 | 4.724016354 | 0.21168428 |
| AT2G19580 | AT2G19580.1 \| Symbols: TET2 \| tetraspanin2 \| chr2:8472393-8475021 REVERSE LENGTH=270 | | 4.716981132 | 4.716981132 | 0.212 |
| AT1G66250 | AT1G66250.1 \| Symbols: \| O-Glycosyl hydrolases family 17 protein \| chr1:24693063-24695416 FORWARD LENGTH=505 | | 4.700131604 | 4.700131604 | 0.21276 |
| AT1G65240 | AT1G65240.1 \| Symbols: \| Eukaryotic aspartyl protease family protein \| chr1:24230963-24233349 REVERSE LENGTH=475 | | 4.667662435 | 4.667662435 | 0.21424 |
| AT1G78300 | AT1G78300.1 \| Symbols: GRF2, 14-3-3OMEGA, GF14 OMEGA \| general regulatory factor 2 \| chr1:29461883-29463052 FORWARD LENGTH=259;AT1G35160.2 \| Symbols: GF14 PHI \| GF14 protein phi chain \| chr1:12867264-12868514 FORWARD LENGTH=295;AT1G35160.1 \| Symbols: GRF4, 14-3-3PHI, GF14 PHI \| GF14 protein phi chain \| chr1:12867264-12868514 FORWARD LENGTH=267;AT1G26480.1 \| Symbols: GRF12, GF14 IOTA \| general regulatory factor 12 \| chr1:9156573-9157845 REVERSE LENGTH=268;AT5G65430.3 \| Symbols: GRF8 \| general regulatory factor 8 \| chr5:26148630-26150255 REVERSE LENGTH=260;AT1G34760.1 \| Symbols: GRF11, GF14 OMICRON, RHS5 \| general regulatory factor 11 \| chr1:12744062-12745603 REVERSE LENGTH=255;AT1G34760.2 \| Symbols: GRF11, GF14 OMICRON, RHS5 \| general regulatory factor 11 \| chr1:12743981-12745603 REVERSE LENGTH=252;AT5G65430.1 \| Symbols: GRF8, 14-3-3KAPPA, GF14 KAPPA \| general regulatory factor 8 \| chr5:26148546-26150255 REVERSE LENGTH=248;AT5G65430.2 \| Symbols: GRF8, GF14 KAPPA \| general regulatory factor 8 \| chr5:26148546-26150255 REVERSE LENGTH=246 | | 4.636068614 | 4.636068614 | 0.2157 |

| | | | | | |
|---|---|---|---|---|---|
| AT2G33120 | AT2G33120.2 \| Symbols: SAR1, VAMP722, ATVAMP722 \| synaptobrevin-related protein 1 \| chr2:14043785-14045337 REVERSE LENGTH=229;AT2G33120.1 \| Symbols: SAR1, VAMP722, ATVAMP722 \| synaptobrevin-related protein 1 \| chr2:14043785-14045337 REVERSE LENGTH=221;AT1G04750.1 \| Symbols: VAMP7B, VAMP721, ATVAMP721, AT VAMP7B \| vesicle-associated membrane protein 721 \| chr1:1331857-1333426 REVERSE LENGTH=219;AT1G04750.2 \| Symbols: VAMP7B, VAMP721, ATVAMP721, AT VAMP7B \| vesicle-associated membrane protein 721 \| chr1:1331857-1333426 REVERSE LENGTH=181;AT2G32670.1 \| Symbols: ATVAMP725, VAMP725 \| vesicle-associated membrane protein 725 \| chr2:13857941-13859346 FORWARD LENGTH=285;AT1G04760.1 \| Symbols: ATVAMP726, VAMP726 \| vesicle-associated membrane protein 726 \| chr1:1334760-1336070 FORWARD LENGTH=220;AT2G33120.3 \| Symbols: SAR1 \| synaptobrevin-related protein 1 \| chr2:14043785-14045186 REVERSE LENGTH=139 | 2.0579 | 7.203054095 | 4.630477047 | 0.21596047 |
| AT1G31730 | AT1G31730.1 \| Symbols: \| Adaptin family protein \| chr1:11359907-11363916 FORWARD LENGTH=938 | 3.3004 | 5.862695667 | 4.581547834 | 0.218266847 |
| AT1G75280 | AT1G75280.1 \| Symbols: \| NmrA-like negative transcriptional regulator family protein \| chr1:28252030-28253355 FORWARD LENGTH=310 | | 4.580012824 | 4.580012824 | 0.21834 |
| AT5G49720 | AT5G49720.1 \| Symbols: ATGH9A1, TSD1, DEC, KOR, RSW2, IRX2, KOR1, GH9A1 \| glycosyl hydrolase 9A1 \| chr5:20197765-20200168 REVERSE LENGTH=621 | | 4.52857531 | 4.52857531 | 0.22082 |
| AT2G43160 | AT2G43160.1 \| Symbols: \| ENTH/VHS family protein \| chr2:17948884-17953267 FORWARD LENGTH=895;AT2G43160.2 \| Symbols: \| ENTH/VHS family protein \| chr2:17948884-17953267 FORWARD LENGTH=895;AT2G43160.3 \| Symbols: \| ENTH/VHS family protein \| chr2:17948884-17953267 FORWARD LENGTH=895;AT2G43160.4 \| Symbols: \| ENTH/VHS family protein \| chr2:17948884-17952920 FORWARD LENGTH=646;AT3G59290.1 \| Symbols: \| ENTH/VHS family protein \| chr3:21912796-21917474 FORWARD LENGTH=1024 | 4.3221 | 4.675737598 | 4.498918799 | 0.222275628 |
| AT2G13820 | AT2G13820.1 \| Symbols: \| Bifunctional inhibitor/lipid-transfer protein/seed storage 2S albumin superfamily protein \| chr2:5774295-5776279 REVERSE LENGTH=169;AT2G13820.2 \| Symbols: \| Bifunctional inhibitor/lipid-transfer protein/seed storage 2S albumin superfamily protein \| chr2:5775805-5776279 REVERSE LENGTH=129 | | 4.483500717 | 4.483500717 | 0.22304 |
| AT4G20720 | AT4G20720.1 \| Symbols: \| dentin sialophosphoprotein-related \| chr4:11105409-11107841 FORWARD LENGTH=729;AT1G05090.1 \| Symbols: \| dentin sialophosphoprotein-related \| chr1:1463202-1465622 FORWARD LENGTH=706 | 3.6046 | 5.350454789 | 4.477527394 | 0.22333755 |
| AT5G07120 | AT5G07120.1 \| Symbols: SNX2b \| sorting nexin 2B \| chr5:2207065-2209355 REVERSE LENGTH=572 | | 4.476075377 | 4.476075377 | 0.22341 |
| AT4G23460 | AT4G23460.1 \| Symbols: \| Adaptin family protein \| chr4:12243899-12248898 REVERSE LENGTH=893 | 4.1252 | 4.80977346 | 4.46748673 | 0.223839501 |
| AT3G51550 | AT3G51550.1 \| Symbols: FER \| Malectin/receptor-like protein kinase family protein \| chr3:19117877-19120564 REVERSE LENGTH=895 | 3.2852 | 5.625562556 | 4.455381278 | 0.224447682 |
| AT5G61820 | AT5G61820.1 \| Symbols: \| FUNCTIONS IN: molecular_function unknown; INVOLVED IN: biological_process unknown; LOCATED IN: vacuole; EXPRESSED IN: 24 plant structures; EXPRESSED DURING: 15 growth stages; CONTAINS InterPro DOMAIN/s: Stress up-regulated Nod 19 (InterPro:IPR011692); Has 30201 Blast hits to 17322 proteins in 780 species: Archae - 12; Bacteria - 1396; Metazoa - 17338; Fungi - 3422; Plants - 5037; Viruses - 0; Other Eukaryotes - 2996 (source: NCBI BLink). \| chr5:24834505-24836223 REVERSE LENGTH=475 | | 4.413452202 | 4.413452202 | 0.22658 |
| AT3G05280 | AT3G05280.1 \| Symbols: \| Integral membrane Yip1 family protein \| chr3:1503992-1505554 REVERSE LENGTH=281 | | 4.398117606 | 4.398117606 | 0.22737 |
| AT3G02520 | AT3G02520.1 \| Symbols: GRF7, GF14 NU \| general regulatory factor 7 \| chr3:526800-527915 REVERSE LENGTH=265 | | 4.360148245 | 4.360148245 | 0.22935 |
| AT5G55480 | AT5G55480.1 \| Symbols: SVL1 \| SHV3-like 1 \| chr5:22474277-22477819 FORWARD LENGTH=766 | 2.8215 | 5.802483463 | 4.311991731 | 0.231911391 |

| | | | | | |
|---|---|---|---|---|---|
| AT4G20830 | AT4G20830.1 \| Symbols: \| FAD-binding Berberine family protein \| chr4:11155486-11157577 FORWARD LENGTH=570;AT4G20830.2 \| Symbols: \| FAD-binding Berberine family protein \| chr4:11155486-11157108 FORWARD LENGTH=540;AT1G01980.1 \| Symbols: \| FAD-binding Berberine family protein \| chr1:340374-341999 REVERSE LENGTH=541 | 2.4323 | 6.002040694 | 4.217170347 | 0.237125826 |
| AT3G02880 | AT3G02880.1 \| Symbols: \| Leucine-rich repeat protein kinase family protein \| chr3:634819-636982 FORWARD LENGTH=627 | 2.4881 | 5.942829976 | 4.215464988 | 0.237221754 |
| AT5G42100 | AT5G42100.1 \| Symbols: ATBG_PPAP, ATBG_PAP, BG_PPAP \| beta-1,3-glucanase_putative \| chr5:16829460-16831168 REVERSE LENGTH=425;AT5G42100.2 \| Symbols: ATBG_PPAP, ATBG_PAP \| beta-1,3-glucanase_putative \| chr5:16829897-16831168 REVERSE LENGTH=423 | 3.4845 | 4.926593753 | 4.205546877 | 0.237781204 |
| AT2G43030 | AT2G43030.1 \| Symbols: \| Ribosomal protein L3 family protein \| chr2:17894898-17895713 FORWARD LENGTH=271 | | 4.16389074 | 4.16389074 | 0.24016 |
| AT5G01750 | AT5G01750.2 \| Symbols: \| Protein of unknown function (DUF567) \| chr5:290034-291109 FORWARD LENGTH=217;AT5G01750.1 \| Symbols: \| Protein of unknown function (DUF567) \| chr5:290034-290685 FORWARD LENGTH=174 | | 4.027386226 | 4.027386226 | 0.2483 |
| AT1G61250 | AT1G61250.1 \| Symbols: SC3 \| secretory carrier 3 \| chr1:22586035-22588664 FORWARD LENGTH=289;AT1G61250.2 \| Symbols: SC3 \| secretory carrier 3 \| chr1:22586035-22588519 FORWARD LENGTH=274;AT1G11180.2 \| Symbols: \| Secretory carrier membrane protein (SCAMP) family protein \| chr1:3745829-3748307 FORWARD LENGTH=304;AT1G11180.1 \| Symbols: \| Secretory carrier membrane protein (SCAMP) family protein \| chr1:3745829-3748307 FORWARD LENGTH=291 | | 3.999840006 | 3.999840006 | 0.25001 |
| AT1G61870 | AT1G61870.1 \| Symbols: PPR336 \| pentatricopeptide repeat 336 \| chr1:22865326-22866552 REVERSE LENGTH=408;AT1G11630.1 \| Symbols: \| Tetratricopeptide repeat (TPR)-like superfamily protein \| chr1:3913168-3914385 REVERSE LENGTH=405 | | 3.99904023 | 3.99904023 | 0.25006 |
| AT4G34870 | AT4G34870.1 \| Symbols: ROC5, ATCYP1 \| rotamase cyclophilin 5 \| chr4:16614451-16614969 FORWARD LENGTH=172 | 3.9887 | | 3.9887 | 0.250708251 |
| AT5G44130 | AT5G44130.1 \| Symbols: FLA13 \| FASCICLIN-like arabinogalactan protein 13 precursor \| chr5:17761128-17761871 FORWARD LENGTH=247 | | 3.984857541 | 3.984857541 | 0.25095 |
| AT1G02180 | AT1G02180.1 \| Symbols: \| ferredoxin-related \| chr1:413619-414505 REVERSE LENGTH=226 | 2.9198 | 5.020584396 | 3.970192198 | 0.251876975 |
| AT3G17840 | AT3G17840.1 \| Symbols: RLK902 \| receptor-like kinase 902 \| chr3:6106092-6108430 FORWARD LENGTH=647 | | 3.952725404 | 3.952725404 | 0.25299 |
| AT5G16970 | AT5G16970.1 \| Symbols: AT-AER, AER \| alkenal reductase \| chr5:5576291-5578001 REVERSE LENGTH=345;AT5G17000.1 \| Symbols: \| Zinc-binding dehydrogenase family protein \| chr5:5584983-5586991 REVERSE LENGTH=345;AT5G16990.1 \| Symbols: \| Zinc-binding dehydrogenase family protein \| chr5:5581831-5583849 REVERSE LENGTH=343;AT5G16980.2 \| Symbols: \| Zinc-binding dehydrogenase family protein \| chr5:5579202-5580889 REVERSE LENGTH=305;AT5G16980.1 \| Symbols: \| Zinc-binding dehydrogenase family protein \| chr5:5579202-5580383 REVERSE LENGTH=239 | | 3.946641408 | 3.946641408 | 0.25338 |
| AT4G09320 | AT4G09320.1 \| Symbols: NDPK1 \| Nucleoside diphosphate kinase family protein \| chr4:5923424-5924366 FORWARD LENGTH=169 | | 3.93452943 | 3.93452943 | 0.25416 |
| AT3G16760 | AT3G16760.1 \| Symbols: \| Tetratricopeptide repeat (TPR)-like superfamily protein \| chr3:5703213-5705080 FORWARD LENGTH=475 | 2.3137 | 5.53464689 | 3.924173445 | 0.254830734 |
| AT2G03450 | AT2G03450.1 \| Symbols: PAP9, ATPAP9 \| purple acid phosphatase 9 \| chr2:1041528-1043569 FORWARD LENGTH=651 | | 3.875818767 | 3.875818767 | 0.25801 |

| | | | | | |
|---|---|---|---|---|---|
| AT4G12780 | AT4G12780.1 \| Symbols: \| Chaperone DnaJ-domain superfamily protein \| chr4:7512200-7516655 FORWARD LENGTH=904;AT4G12780.2 \| Symbols: \| Chaperone DnaJ-domain superfamily protein \| chr4:7512200-7516655 FORWARD LENGTH=894;AT4G12770.1 \| Symbols: \| Chaperone DnaJ-domain superfamily protein \| chr4:7506736-7511408 REVERSE LENGTH=891;AT4G12770.2 \| Symbols: \| Chaperone DnaJ-domain superfamily protein \| chr4:7506736-7511408 REVERSE LENGTH=889 | 3.98 | 3.747704531 | 3.863852265 | 0.258809067 |
| AT2G40380 | AT2G40380.1 \| Symbols: PRA1.B2 \| prenylated RAB acceptor 1.B2 \| chr2:16864734-16865375 REVERSE LENGTH=213 | | 3.86368905 | 3.86368905 | 0.25882 |
| AT3G19930 | AT3G19930.1 \| Symbols: STP4, ATSTP4 \| sugar transporter 4 \| chr3:6935048-6936841 FORWARD LENGTH=514 | 4.7548 | 2.892430509 | 3.823615255 | 0.26153259 |
| AT5G06140 | AT5G06140.1 \| Symbols: SNX1, ATSNX1 \| sorting nexin 1 \| chr5:1856212-1858752 REVERSE LENGTH=402 | | 3.786157807 | 3.786157807 | 0.26412 |
| AT1G60780 | AT1G60780.1 \| Symbols: HAP13 \| Clathrin adaptor complexes medium subunit family protein \| chr1:22369289-22371885 REVERSE LENGTH=428;AT1G10730.1 \| Symbols: \| Clathrin adaptor complexes medium subunit family protein \| chr1:3565463-3567779 FORWARD LENGTH=428 | 3.0692 | 4.498020871 | 3.783610435 | 0.264297823 |
| AT1G75270 | AT1G75270.1 \| Symbols: DHAR2 \| dehydroascorbate reductase 2 \| chr1:28250255-28251237 REVERSE LENGTH=213 | 3.7821 | | 3.7821 | 0.264403374 |
| AT1G71820 | AT1G71820.2 \| Symbols: SEC6 \| SEC6 \| chr1:27010022-27016745 FORWARD LENGTH=774;AT1G71820.1 \| Symbols: SEC6 \| SEC6 \| chr1:27010022-27016745 FORWARD LENGTH=752 | | 3.78028957 | 3.78028957 | 0.26453 |
| AT2G20010 | AT2G20010.2 \| Symbols: \| Protein of unknown function (DUF810) \| chr2:8637977-8641184 REVERSE LENGTH=952;AT2G20010.1 \| Symbols: \| Protein of unknown function (DUF810) \| chr2:8637977-8640830 REVERSE LENGTH=834 | | 3.776292436 | 3.776292436 | 0.26481 |
| AT2G38410 | AT2G38410.1 \| Symbols: \| ENTH/VHS/GAT family protein \| chr2:16086912-16090047 REVERSE LENGTH=671 | 3.5177 | 4.034210102 | 3.775955051 | 0.264833661 |
| AT1G21660 | AT1G21660.1 \| Symbols: \| Chaperone DnaJ-domain superfamily protein \| chr1:7605924-7608835 FORWARD LENGTH=523 | 2.5623 | 4.982561036 | 3.772430518 | 0.265081092 |
| AT3G24660 | AT3G24660.1 \| Symbols: TMKL1 \| transmembrane kinase-like 1 \| chr3:9003641-9005751 FORWARD LENGTH=674 | | 3.748125937 | 3.748125937 | 0.2668 |
| AT4G08850 | AT4G08850.1 \| Symbols: \| Leucine-rich repeat receptor-like protein kinase family protein \| chr4:5636693-5640496 REVERSE LENGTH=1045;AT4G08850.2 \| Symbols: \| Leucine-rich repeat receptor-like protein kinase family protein \| chr4:5637467-5640496 REVERSE LENGTH=1009 | 2.6665 | 4.82043866 | 3.74346933 | 0.26713188 |
| AT1G07210 | AT1G07210.1 \| Symbols: \| Ribosomal protein S18 \| chr1:2215320-2216668 FORWARD LENGTH=261 | | 3.740135393 | 3.740135393 | 0.26737 |
| AT1G09330 | AT1G09330.1 \| Symbols: \| unknown protein; FUNCTIONS IN: molecular_function unknown; INVOLVED IN: biological_process unknown; LOCATED IN: integral to membrane; EXPRESSED IN: 24 plant structures; EXPRESSED DURING: 15 growth stages; CONTAINS InterPro DOMAIN/s: Protein of unknown function DUF846, eukaryotic (InterPro:IPR008564); Has 518 Blast hits to 518 proteins in 206 species: Archae - 0; Bacteria - 0; Metazoa - 193; Fungi - 145; Plants - 73; Viruses - 0; Other Eukaryotes - 107 (source: NCBI BLink). \| chr1:3013003-3014903 REVERSE LENGTH=186 | 3.0066 | 4.472671974 | 3.739635987 | 0.267405706 |
| AT4G22570 | AT4G22570.1 \| Symbols: APT3 \| adenine phosphoribosyl transferase 3 \| chr4:11882310-11885250 REVERSE LENGTH=183 | | 3.718992897 | 3.718992897 | 0.26889 |

| | | | | | |
|---|---|---|---|---|---|
| AT1G60730 | AT1G60730.3 \| Symbols: \| NAD(P)-linked oxidoreductase superfamily protein \| chr1:22358327-22360082 REVERSE LENGTH=365;AT1G60680.1 \| Symbols: \| NAD(P)-linked oxidoreductase superfamily protein \| chr1:22347824-22349236 REVERSE LENGTH=346;AT1G60710.1 \| Symbols: ATB2 \| NAD(P)-linked oxidoreductase superfamily protein \| chr1:22355073-22356627 REVERSE LENGTH=345;AT1G60730.1 \| Symbols: \| NAD(P)-linked oxidoreductase superfamily protein \| chr1:22358327-22360082 REVERSE LENGTH=345;AT1G60750.1 \| Symbols: \| NAD(P)-linked oxidoreductase superfamily protein \| chr1:22362293-22363854 REVERSE LENGTH=330;AT1G60730.2 \| Symbols: \| NAD(P)-linked oxidoreductase superfamily protein \| chr1:22358613-22360082 REVERSE LENGTH=251 | | 3.701647233 | 3.701647233 | 0.27015 |
| AT1G34750 | AT1G34750.1 \| Symbols: \| Protein phosphatase 2C family protein \| chr1:12736386-12737727 REVERSE LENGTH=282 | | 3.635306093 | 3.635306093 | 0.27508 |
| AT2G30050 | AT2G30050.1 \| Symbols: \| transducin family protein / WD-40 repeat family protein \| chr2:12825540-12826448 FORWARD LENGTH=302 | | 3.575642722 | 3.575642722 | 0.27967 |
| AT3G28715 | AT3G28715.1 \| Symbols: \| ATPase, V0/A0 complex, subunit C/D \| chr3:10778025-10780350 FORWARD LENGTH=351;AT3G28715.2 \| Symbols: \| ATPase, V0/A0 complex, subunit C/D \| chr3:10778025-10780350 FORWARD LENGTH=343 | | 3.565062389 | 3.565062389 | 0.2805 |
| AT5G62630 | AT5G62630.1 \| Symbols: HIPL2 \| hipl2 protein precursor \| chr5:25143719-25146390 REVERSE LENGTH=696 | | 3.559225513 | 3.559225513 | 0.28096 |
| AT4G18060 | AT4G18060.1 \| Symbols: \| SH3 domain-containing protein \| chr4:10027668-10029662 REVERSE LENGTH=351 | | 3.549497746 | 3.549497746 | 0.28173 |
| AT3G53780 | AT3G53780.2 \| Symbols: ATRBL4, RBL4 \| RHOMBOID-like protein 4 \| chr3:19924729-19926648 REVERSE LENGTH=394;AT3G53780.1 \| Symbols: ATRBL4, RBL4 \| RHOMBOID-like protein 4 \| chr3:19924729-19926025 REVERSE LENGTH=270 | | 3.490279571 | 3.490279571 | 0.28651 |
| AT1G64880 | AT1G64880.1 \| Symbols: \| Ribosomal protein S5 family protein \| chr1:24105713-24108780 FORWARD LENGTH=515 | | 3.462244227 | 3.462244227 | 0.28883 |
| AT4G27350 | AT4G27350.1 \| Symbols: \| Protein of unknown function (DUF1223) \| chr4:13690979-13692621 REVERSE LENGTH=278 | | 3.439499209 | 3.439499209 | 0.29074 |
| AT1G01980 | AT1G01980.1 \| Symbols: \| FAD-binding Berberine family protein \| chr1:340374-341999 REVERSE LENGTH=541 | | 3.437134804 | 3.437134804 | 0.29094 |
| AT5G56590 | AT5G56590.1 \| Symbols: \| O-Glycosyl hydrolases family 17 protein \| chr5:22907521-22909436 FORWARD LENGTH=506 | | 3.359424866 | 3.359424866 | 0.29767 |
| AT5G17310 | AT5G17310.2 \| Symbols: UGP2 \| UDP-glucose pyrophosphorylase 2 \| chr5:5696955-5700845 REVERSE LENGTH=470;AT3G03250.1 \| Symbols: UGP, UGP1, AtUGP1 \| UDP-GLUCOSE PYROPHOSPHORYLASE 1 \| chr3:749761-754014 REVERSE LENGTH=469;AT5G17310.1 \| Symbols: UGP2, AtUGP2 \| UDP-glucose pyrophosphorylase 2 \| chr5:5696955-5699671 REVERSE LENGTH=390 | | 3.34694424 | 3.34694424 | 0.29878 |
| AT3G52850 | AT3G52850.1 \| Symbols: VSR1, BP-80, ATELP, BP80, BP80B, ATELP1, ATVSR1, GFS1, VSR1;1, BP80-1;1 \| vacuolar sorting receptor homolog 1 \| chr3:19587999-19591690 FORWARD LENGTH=623 | 3.6801 | 2.989894158 | 3.334997079 | 0.299850338 |
| AT3G07390 | AT3G07390.1 \| Symbols: AIR12 \| auxin-responsive family protein \| chr3:2365452-2366273 FORWARD LENGTH=273 | 2.9243 | 3.724672229 | 3.324486114 | 0.300798369 |
| AT2G25800 | AT2G25800.1 \| Symbols: \| Protein of unknown function (DUF810) \| chr2:11006138-11009728 REVERSE LENGTH=987 | 3.2191 | 3.424188467 | 3.321644234 | 0.301055721 |
| AT5G43100 | AT5G43100.1 \| Symbols: \| Eukaryotic aspartyl protease family protein \| chr5:17299264-17302718 FORWARD LENGTH=631 | | 3.312025966 | 3.312025966 | 0.30193 |

| | | | | | |
|---|---|---|---|---|---|
| AT4G35100 | AT4G35100.1 \| Symbols: PIP3, PIP3A, PIP2;7, SIMIP \| plasma membrane intrinsic protein 3 \| chr4:16708672-16709958 FORWARD LENGTH=280;AT4G35100.2 \| Symbols: PIP3 \| plasma membrane intrinsic protein 3 \| chr4:16708672-16709958 FORWARD LENGTH=280;AT5G60660.1 \| Symbols: PIP2F, PIP2;4 \| plasma membrane intrinsic protein 2;4 \| chr5:24375673-24376939 REVERSE LENGTH=291;AT2G16850.1 \| Symbols: PIP3B, PIP2;8 \| plasma membrane intrinsic protein 2;8 \| chr2:7301647-7303180 FORWARD LENGTH=278 | | 3.295435821 | 3.295435821 | 0.30345 |
| AT3G17020 | AT3G17020.1 \| Symbols: \| Adenine nucleotide alpha hydrolases-like superfamily protein \| chr3:5802728-5804063 REVERSE LENGTH=163 | 4.1649 | 2.389429165 | 3.277164583 | 0.305141831 |
| AT5G59520 | AT5G59520.1 \| Symbols: ZIP2 \| ZRT/IRT-like protein 2 \| chr5:23991447-23992762 REVERSE LENGTH=353 | | 3.272572569 | 3.272572569 | 0.30557 |
| AT5G03700 | AT5G03700.1 \| Symbols: \| D-mannose binding lectin protein with Apple-like carbohydrate-binding domain \| chr5:965874-967322 REVERSE LENGTH=482 | | 3.269469692 | 3.269469692 | 0.30586 |
| AT2G42590 | AT2G42590.3 \| Symbols: GRF9, GF14 MU \| general regulatory factor 9 \| chr2:17732164-17733775 REVERSE LENGTH=276;AT2G42590.1 \| Symbols: GRF9, GF14 MU \| general regulatory factor 9 \| chr2:17732118-17733775 REVERSE LENGTH=263;AT2G42590.2 \| Symbols: GRF9, GF14 MU \| general regulatory factor 9 \| chr2:17732118-17733775 REVERSE LENGTH=262 | | 3.25414904 | 3.25414904 | 0.3073 |
| AT1G04410 | AT1G04410.1 \| Symbols: \| Lactate/malate dehydrogenase family protein \| chr1:1189418-1191267 REVERSE LENGTH=332;AT5G43330.1 \| Symbols: \| Lactate/malate dehydrogenase family protein \| chr5:17390552-17392449 FORWARD LENGTH=332;AT5G56720.1 \| Symbols: \| Lactate/malate dehydrogenase family protein \| chr5:22945537-22946718 FORWARD LENGTH=339 | 0.82848 | 5.64557105 | 3.237025525 | 0.308925584 |
| AT4G26690 | AT4G26690.1 \| Symbols: SHV3, MRH5, GPDL2 \| PLC-like phosphodiesterase family protein \| chr4:13456793-13459890 REVERSE LENGTH=759 | | 3.225598348 | 3.225598348 | 0.31002 |
| AT2G01820 | AT2G01820.1 \| Symbols: \| Leucine-rich repeat protein kinase family protein \| chr2:357664-360681 REVERSE LENGTH=943;AT1G24650.1 \| Symbols: \| Leucine-rich repeat protein kinase family protein \| chr1:8734570-8737315 FORWARD LENGTH=886 | 2.9257 | 3.514938489 | 3.220319244 | 0.310528219 |
| AT4G20410 | AT4G20410.1 \| Symbols: GSNAP, GAMMA-SNAP \| gamma-soluble NSF attachment protein \| chr4:11014099-11016454 REVERSE LENGTH=291 | | 3.209448617 | 3.209448617 | 0.31158 |
| AT3G62370 | AT3G62370.1 \| Symbols: \| heme binding \| chr3:23080789-23083234 FORWARD LENGTH=361 | | 3.204203916 | 3.204203916 | 0.31209 |
| AT4G20260 | AT4G20260.4 \| Symbols: ATPCAP1, PCAP1 \| plasma-membrane associated cation-binding protein 1 \| chr4:10941593-10943227 FORWARD LENGTH=226;AT4G20260.1 \| Symbols: ATPCAP1, PCAP1 \| plasma-membrane associated cation-binding protein 1 \| chr4:10941593-10943227 FORWARD LENGTH=225;AT4G20260.2 \| Symbols: ATPCAP1, PCAP1 \| plasma-membrane associated cation-binding protein 1 \| chr4:10941593-10943227 FORWARD LENGTH=225;AT4G20260.3 \| Symbols: ATPCAP1, PCAP1 \| plasma-membrane associated cation-binding protein 1 \| chr4:10941593-10943227 FORWARD LENGTH=225;AT4G20260.5 \| Symbols: PCAP1 \| plasma-membrane associated cation-binding protein 1 \| chr4:10941593-10943120 FORWARD LENGTH=167;AT4G20260.6 \| Symbols: PCAP1 \| plasma-membrane associated cation-binding protein 1 \| chr4:10941593-10943227 FORWARD LENGTH=166 | 0.76545 | 5.634437683 | 3.199943842 | 0.312505484 |
| AT4G04700 | AT4G04700.1 \| Symbols: CPK27 \| calcium-dependent protein kinase 27 \| chr4:2385276-2387986 REVERSE LENGTH=485 | | 3.198567042 | 3.198567042 | 0.31264 |
| AT4G24550 | AT4G24550.2 \| Symbols: \| Clathrin adaptor complexes medium subunit family protein \| chr4:12675873-12678903 FORWARD LENGTH=451;AT4G24550.3 \| Symbols: \| Clathrin adaptor complexes medium subunit family protein \| chr4:12675873-12678689 FORWARD LENGTH=385;AT4G24550.1 \| Symbols: \| Clathrin adaptor complexes medium subunit family protein \| chr4:12675873-12678526 FORWARD LENGTH=380 | 3.4695 | 2.875629044 | 3.172564522 | 0.315202415 |

| | | | | | |
|---|---|---|---|---|---|
| AT5G19440 | AT5G19440.1 \| Symbols: \| NAD(P)-binding Rossmann-fold superfamily protein \| chr5:6556493-6558123 FORWARD LENGTH=326 | | 3.167463812 | 3.167463812 | 0.31571 |
| AT3G44150 | AT3G44150.1 \| Symbols: \| unknown protein; FUNCTIONS IN: molecular_function unknown; INVOLVED IN: biological_process unknown; LOCATED IN: plasma membrane; EXPRESSED IN: cultured cell; BEST Arabidopsis thaliana protein match is: unknown protein (TAIR:AT3G11800.1); Has 76 Blast hits to 75 proteins in 19 species: Archae - 0; Bacteria - 0; Metazoa - 0; Fungi - 0; Plants - 74; Viruses - 0; Other Eukaryotes - 2 (source: NCBI BLink). \| chr3:15887617-15888728 FORWARD LENGTH=246;AT3G11800.1 \| Symbols: \| unknown protein; FUNCTIONS IN: molecular_function unknown; INVOLVED IN: biological_process unknown; LOCATED IN: endomembrane system; EXPRESSED IN: 24 plant structures; EXPRESSED DURING: 15 growth stages; BEST Arabidopsis thaliana protein match is: unknown protein (TAIR:AT3G44150.1); Has 74 Blast hits to 73 proteins in 18 species: Archae - 0; Bacteria - 0; Metazoa - 0; Fungi - 0; Plants - 72; Viruses - 0; Other Eukaryotes - 2 (source: NCBI BLink). \| chr3:3726154-3727562 FORWARD LENGTH=246 | | 3.162555345 | 3.162555345 | 0.3162 |
| AT3G46290 | AT3G46290.1 \| Symbols: HERK1 \| hercules receptor kinase 1 \| chr3:17013009-17015501 FORWARD LENGTH=830 | | 3.155470007 | 3.155470007 | 0.31691 |
| AT5G62740 | AT5G62740.1 \| Symbols: HIR1, ATHIR1 \| SPFH/Band 7/PHB domain-containing membrane-associated protein family \| chr5:25201320-25202535 FORWARD LENGTH=286;AT3G01290.1 \| Symbols: \| SPFH/Band 7/PHB domain-containing membrane-associated protein family \| chr3:88252-89356 REVERSE LENGTH=285 | 1.0246 | 5.268148773 | 3.146374386 | 0.317826132 |
| AT5G24430 | AT5G24430.1 \| Symbols: \| Calcium-dependent protein kinase (CDPK) family protein \| chr5:8339390-8342913 REVERSE LENGTH=594 | | 3.129890454 | 3.129890454 | 0.3195 |
| AT3G56190 | AT3G56190.1 \| Symbols: ALPHA-SNAP2, ASNAP \| alpha-soluble NSF attachment protein 2 \| chr3:20846119-20848356 REVERSE LENGTH=289;AT3G56190.2 \| Symbols: ALPHA-SNAP2, ASNAP \| alpha-soluble NSF attachment protein 2 \| chr3:20846119-20848213 REVERSE LENGTH=240;AT3G56450.1 \| Symbols: ALPHA-SNAP1 \| alpha-soluble NSF attachment protein 1 \| chr3:20930557-20932869 FORWARD LENGTH=381 | | 3.108679433 | 3.108679433 | 0.32168 |
| AT2G19760 | AT2G19760.1 \| Symbols: PFN1, PRF1 \| profilin 1 \| chr2:8517074-8518067 REVERSE LENGTH=131 | | 3.088898499 | 3.088898499 | 0.32374 |
| AT1G04260 | AT1G04260.1 \| Symbols: MPI7, MPIP7, PRA1.D \| CAMV movement protein interacting protein 7 \| chr1:1140749-1141297 REVERSE LENGTH=182 | | 3.070121577 | 3.070121577 | 0.32572 |
| AT4G28360 | AT4G28360.1 \| Symbols: \| Ribosomal protein L22p/L17e family protein \| chr4:14029294-14030926 REVERSE LENGTH=271 | | 3.049431281 | 3.049431281 | 0.32793 |
| AT2G36460 | AT2G36460.1 \| Symbols: \| Aldolase superfamily protein \| chr2:15296929-15298387 REVERSE LENGTH=358;AT3G52930.1 \| Symbols: \| Aldolase superfamily protein \| chr3:19627383-19628874 REVERSE LENGTH=358;AT2G36460.2 \| Symbols: \| Aldolase superfamily protein \| chr2:15296929-15297732 REVERSE LENGTH=267 | | 3.015317814 | 3.015317814 | 0.33164 |
| AT1G30360 | AT1G30360.1 \| Symbols: ERD4 \| Early-responsive to dehydration stress protein (ERD4) \| chr1:10715892-10718799 FORWARD LENGTH=724 | 4.7088 | 1.279885322 | 2.994342661 | 0.333963114 |
| AT5G16590 | AT5G16590.1 \| Symbols: LRR1 \| Leucine-rich repeat protein kinase family protein \| chr5:5431862-5433921 FORWARD LENGTH=625 | | 2.98445101 | 2.98445101 | 0.33507 |
| AT5G40370 | AT5G40370.2 \| Symbols: \| Glutaredoxin family protein \| chr5:16147826-16148900 REVERSE LENGTH=136;AT5G40370.1 \| Symbols: \| Glutaredoxin family protein \| chr5:16147826-16149052 REVERSE LENGTH=111 | | 2.981959147 | 2.981959147 | 0.33535 |

| | | | | | |
|---|---|---|---|---|---|
| AT5G62880 | AT5G62880.1 \| Symbols: ARAC10, ATRAC10, ATROP11, RAC10 \| RAC-like 10 \| chr5:25237236-25238939 FORWARD LENGTH=215;AT2G44690.1 \| Symbols: ARAC9, ATROP8, ROP8 \| Arabidopsis RAC-like 9 \| chr2:18429276-18430636 FORWARD LENGTH=209;AT4G28950.1 \| Symbols: ARAC7, ATROP9, ATRAC7, RAC7, ROP9 \| RHO-related protein from plants 9 \| chr4:14278289-14279705 FORWARD LENGTH=209;AT3G48040.1 \| Symbols: ARAC8, ATROP10, ROP10, ATRAC8 \| RHO-related protein from plants 10 \| chr3:17731561-17733241 FORWARD LENGTH=208;AT5G45970.1 \| Symbols: ARAC2, RAC2, ROP7, ATRAC2, ATROP7 \| RAC-like 2 \| chr5:18643761-18645758 FORWARD LENGTH=201;AT4G35020.1 \| Symbols: ARAC3, ROP6, RHO1PS, ATROP6, RAC3 \| RAC-like 3 \| chr4:16673176-16674540 FORWARD LENGTH=198;AT4G35020.2 \| Symbols: RAC3 \| RAC-like 3 \| chr4:16673176-16674540 FORWARD LENGTH=198;AT4G35020.3 \| Symbols: RAC3 \| RAC-like 3 \| chr4:16673176-16674540 FORWARD LENGTH=198;AT2G17800.1 \| Symbols: ARAC1, ATGP2, ATRAC1, ROP3, ATROP3 \| Arabidopsis RAC-like 1 \| chr2:7740313-7741942 FORWARD LENGTH=197;AT2G17800.2 \| Symbols: ARAC1, ATGP2, ATRAC1, ROP3, ATROP3 \| Arabidopsis RAC-like 1 \| chr2:7740313-7741942 FORWARD LENGTH=197;AT3G51300.1 \| Symbols: ARAC11, ROP1, ROP1AT, ATRAC11, ATROP1 \| RHO-related protein from plants 1 \| chr3:19043197-19044215 FORWARD LENGTH=197;AT4G35950.1 \| Symbols: ARAC6, RAC2, ATROP5, ATRAC6, ROP5, RAC6 \| RAC-like 6 \| chr4:17024051-17025514 REVERSE LENGTH=197;AT1G75840.1 \| Symbols: ARAC5, ATGP3, ROP4, ATROP4 \| RAC-like GTP binding protein 5 \| chr1:28475964-28477377 FORWARD LENGTH=196;AT1G20090.1 \| Symbols: ARAC4, ROP2, ATROP2, ATRAC4 \| RHO-related protein from plants 2 \| chr1:6967223-6968603 FORWARD LENGTH=195 | | 2.941176471 | 2.941176471 | 0.34 |
| AT2G28520 | AT2G28520.1 \| Symbols: VHA-A1 \| vacuolar proton ATPase A1 \| chr2:12210026-12215532 FORWARD LENGTH=817 | 3.9194 | 1.906904903 | 2.913152451 | 0.343270741 |
| AT5G46700 | AT5G46700.1 \| Symbols: TET1, TRN2 \| Tetraspanin family protein \| chr5:18951035-18952439 FORWARD LENGTH=269 | | 2.909937436 | 2.909937436 | 0.34365 |
| AT3G11530 | AT3G11530.1 \| Symbols: \| Vacuolar protein sorting 55 (VPS55) family protein \| chr3:3628801-3629885 REVERSE LENGTH=113;AT3G11530.2 \| Symbols: \| Vacuolar protein sorting 55 (VPS55) family protein \| chr3:3628801-3629885 REVERSE LENGTH=113 | 3.3682 | 2.422304581 | 2.89525229 | 0.345393043 |
| AT5G46630 | AT5G46630.1 \| Symbols: \| Clathrin adaptor complexes medium subunit family protein \| chr5:18920580-18923252 FORWARD LENGTH=438;AT5G46630.2 \| Symbols: \| Clathrin adaptor complexes medium subunit family protein \| chr5:18920580-18923060 FORWARD LENGTH=441 | | 2.887002714 | 2.887002714 | 0.34638 |
| AT3G15060 | AT3G15060.1 \| Symbols: AtRABA1g, RABA1g \| RAB GTPase homolog A1G \| chr3:5069239-5070025 FORWARD LENGTH=217;AT1G16920.1 \| Symbols: RAB11, ATRABA1B, RABA1b \| RAB GTPase homolog A1B \| chr1:5787489-5789147 REVERSE LENGTH=216;AT2G33870.1 \| Symbols: ArRABA1h \| RAB GTPase homolog A1H \| chr2:14337366-14338251 REVERSE LENGTH=218;AT1G09630.1 \| Symbols: ATRAB11C, ATRABA2A, ATRAB-A2A, RAB-A2A, RAB11c \| RAB GTPase 11C \| chr1:3118350-3119571 REVERSE LENGTH=217 | | 2.86377044 | 2.86377044 | 0.34919 |
| AT4G18800 | AT4G18800.1 \| Symbols: ATHSGBP, ATRAB11B, ATRABA1D, RABA1d \| RAB GTPase homolog A1D \| chr4:10320156-10321339 REVERSE LENGTH=214 | | 2.86377044 | 2.86377044 | 0.34919 |
| AT4G12730 | AT4G12730.1 \| Symbols: FLA2 \| FASCICLIN-like arabinogalactan 2 \| chr4:7491598-7492809 REVERSE LENGTH=403 | | 2.836074872 | 2.836074872 | 0.3526 |
| AT3G06270 | AT3G06270.1 \| Symbols: \| Protein phosphatase 2C family protein \| chr3:1896763-1897887 FORWARD LENGTH=348 | | 2.817060116 | 2.817060116 | 0.35498 |

| | | | | | |
|---|---|---|---|---|---|
| AT1G69840 | AT1G69840.1 \| Symbols: \| SPFH/Band 7/PHB domain-containing membrane-associated protein family \| chr1:26293932-26295150 REVERSE LENGTH=286;AT1G69840.2 \| Symbols: \| SPFH/Band 7/PHB domain-containing membrane-associated protein family \| chr1:26293932-26295150 REVERSE LENGTH=286;AT1G69840.3 \| Symbols: \| SPFH/Band 7/PHB domain-containing membrane-associated protein family \| chr1:26293932-26295150 REVERSE LENGTH=286;AT1G69840.4 \| Symbols: \| SPFH/Band 7/PHB domain-containing membrane-associated protein family \| chr1:26293932-26295150 REVERSE LENGTH=286;AT1G69840.5 \| Symbols: \| SPFH/Band 7/PHB domain-containing membrane-associated protein family \| chr1:26293932-26295150 REVERSE LENGTH=286;AT1G69840.6 \| Symbols: \| SPFH/Band 7/PHB domain-containing membrane-associated protein family \| chr1:26293932-26295150 REVERSE LENGTH=286;AT1G69840.7 \| Symbols: \| SPFH/Band 7/PHB domain-containing membrane-associated protein family \| chr1:26293932-26295150 REVERSE LENGTH=286 | 1.3597 | 4.203800235 | 2.781750118 | 0.35948592 |
| AT1G45145 | AT1G45145.1 \| Symbols: ATTRX5, ATH5, LIV1, TRX5 \| thioredoxin H-type 5 \| chr1:17075264-17076256 REVERSE LENGTH=118 | | 2.765639692 | 2.765639692 | 0.36158 |
| AT2G14720 | AT2G14720.1 \| Symbols: VSR4, VSR2;1, BP80-2;1, MTV4 \| vacuolar sorting receptor 4 \| chr2:6300878-6304156 REVERSE LENGTH=628;AT2G14720.2 \| Symbols: VSR4, VSR2;1, BP80-2;1, MTV4 \| vacuolar sorting receptor 4 \| chr2:6300878-6304156 REVERSE LENGTH=628;AT2G14740.1 \| Symbols: ATVSR3, VSR3, VSR2;2, BP80-2;2 \| vaculolar sorting receptor 3 \| chr2:6308895-6312303 FORWARD LENGTH=628;AT2G14740.2 \| Symbols: ATVSR3, VSR3, VSR2;2, BP80-2;2 \| vaculolar sorting receptor 3 \| chr2:6308895-6312303 FORWARD LENGTH=628 | 1.4719 | 4.029658285 | 2.750779142 | 0.363533366 |
| AT4G28400 | AT4G28400.1 \| Symbols: \| Protein phosphatase 2C family protein \| chr4:14048499-14050118 FORWARD LENGTH=283;AT5G10740.1 \| Symbols: \| Protein phosphatase 2C family protein \| chr5:3393797-3395848 REVERSE LENGTH=354;AT2G20630.2 \| Symbols: PIA1 \| PP2C induced by AVRRPM1 \| chr2:8897335-8899648 REVERSE LENGTH=290;AT3G15260.1 \| Symbols: \| Protein phosphatase 2C family protein \| chr3:5138842-5140242 FORWARD LENGTH=289;AT3G15260.2 \| Symbols: \| Protein phosphatase 2C family protein \| chr3:5138842-5140242 FORWARD LENGTH=289;AT2G20630.1 \| Symbols: PIA1 \| PP2C induced by AVRRPM1 \| chr2:8897826-8899648 REVERSE LENGTH=279 | | 2.731270314 | 2.731270314 | 0.36613 |
| AT3G63250 | AT3G63250.1 \| Symbols: HMT-2, ATHMT-2, HMT2 \| homocysteine methyltransferase 2 \| chr3:23370575-23372587 REVERSE LENGTH=333;AT3G63250.2 \| Symbols: HMT-2, ATHMT-2, HMT2 \| homocysteine methyltransferase 2 \| chr3:23370575-23371974 REVERSE LENGTH=293 | | 2.687521836 | 2.687521836 | 0.37209 |
| AT1G62820 | AT1G62820.1 \| Symbols: \| Calcium-binding EF-hand family protein \| chr1:23263822-23264268 REVERSE LENGTH=148 | 2.6862 | | 2.6862 | 0.3722731 |
| AT4G23650 | AT4G23650.1 \| Symbols: CDPK6, CPK3 \| calcium-dependent protein kinase 6 \| chr4:12324967-12327415 REVERSE LENGTH=529 | | 2.641449628 | 2.641449628 | 0.37858 |
| AT5G22780 | AT5G22780.1 \| Symbols: \| Adaptor protein complex AP-2, alpha subunit \| chr5:7590100-7597828 REVERSE LENGTH=1013;AT5G22770.1 \| Symbols: alpha-ADR \| alpha-adaptin \| chr5:7579844-7588026 REVERSE LENGTH=1012;AT5G22770.2 \| Symbols: alpha-ADR \| alpha-adaptin \| chr5:7579844-7588026 REVERSE LENGTH=1012;AT5G22770.3 \| Symbols: alpha-ADR \| alpha-adaptin \| chr5:7579844-7588026 REVERSE LENGTH=1012 | 1.5563 | 3.681071928 | 2.618685964 | 0.381870913 |
| AT4G11220 | AT4G11220.1 \| Symbols: BTI2, RTNLB2 \| VIRB2-interacting protein 2 \| chr4:6838176-6839578 REVERSE LENGTH=271 | | 2.583645524 | 2.583645524 | 0.38705 |
| AT3G02650 | AT3G02650.1 \| Symbols: \| Tetratricopeptide repeat (TPR)-like superfamily protein \| chr3:568135-569865 FORWARD LENGTH=576 | | 2.556302564 | 2.556302564 | 0.39119 |

| | | | | | |
|---|---|---|---|---|---|
| AT4G13730 | AT4G13730.1 \| Symbols: \| Ypt/Rab-GAP domain of gyp1p superfamily protein \| chr4:7970299-7974055 FORWARD LENGTH=449;AT4G13730.3 \| Symbols: \| Ypt/Rab-GAP domain of gyp1p superfamily protein \| chr4:7970299-7974055 FORWARD LENGTH=438;AT4G13730.2 \| Symbols: \| Ypt/Rab-GAP domain of gyp1p superfamily protein \| chr4:7970299-7973901 FORWARD LENGTH=408 | | 2.556302564 | 2.556302564 | 0.39119 |
| AT1G31340 | [illegible] chr1:11218076-11219417 REVERSE LENGTH=156;AT2G35635.1 \| Symbols: UBQ7, RUB2 \| ubiquitin 7 \| chr2:14981044-14981943 FORWARD LENGTH=154;AT3G09790.1 \| Symbols: UBQ8 \| ubiquitin 8 \| chr3:3004111-3006006 REVERSE LENGTH=631;AT4G05320.2 \| Symbols: UBQ10 \| polyubiquitin 10 \| chr4:2718559-2719932 FORWARD LENGTH=457;AT4G05320.4 \| Symbols: UBQ10 \| polyubiquitin 10 \| chr4:2718559-2719932 FORWARD LENGTH=457;AT5G20620.1 \| Symbols: UBQ4 \| ubiquitin 4 \| chr5:6973315-6974463 REVERSE LENGTH=382;AT4G05320.1 \| Symbols: UBQ10 \| polyubiquitin 10 \| chr4:2718559-2719932 FORWARD LENGTH=381;AT4G05320.3 \| Symbols: UBQ10 \| polyubiquitin 10 \| chr4:2718787-2719932 FORWARD LENGTH=381;AT4G05320.6 \| Symbols: UBQ10 \| polyubiquitin 10 \| chr4:2718559-2719932 FORWARD LENGTH=381;AT1G65350.1 \| Symbols: UBQ13 \| ubiquitin 13 \| chr1:24272518-24277275 REVERSE LENGTH=319;AT5G03240.1 \| Symbols: UBQ3 \| polyubiquitin 3 \| chr5:771976-772896 REVERSE LENGTH=306;AT5G03240.2 \| Symbols: UBQ3 \| polyubiquitin 3 \| chr5:771976-772896 REVERSE LENGTH=306;AT5G03240.3 \| Symbols: UBQ3 \| polyubiquitin 3 \| chr5:771976-772896 REVERSE LENGTH=306;AT4G02890.3 \| Symbols: UBQ14 \| Ubiquitin family protein \| chr4:1278747-1279664 REVERSE LENGTH=305;AT4G02890.4 \| Symbols: UBQ14 \| Ubiquitin family protein \| chr4:1278747-1279664 REVERSE LENGTH=305;AT4G05320.5 \| Symbols: UBQ10 \| polyubiquitin 10 \| chr4:2718559-2719932 FORWARD LENGTH=305;AT1G55060.1 \| Symbols: UBQ12 \| ubiquitin 12 \| chr1:20549533-20550225 FORWARD LENGTH=230;AT4G02890.1 \| Symbols: UBQ14 \| Ubiquitin family protein \| chr4:1278747-1279664 REVERSE LENGTH=229;AT4G02890.2 \| Symbols: UBQ14 \| Ubiquitin family protein \| chr4:1278747-1279664 REVERSE LENGTH=229;AT4G05050.1 \| Symbols: UBQ11 \| ubiquitin 11 \| chr4:2588271-2588960 REVERSE LENGTH=229;AT4G05050.2 \| Symbols: UBQ11 \| ubiquitin 11 \| | | 2.528700754 | 2.528700754 | 0.39546 |
| AT4G24990 | AT4G24990.1 \| Symbols: ATGP4 \| Ubiquitin family protein \| chr4:12849973-12851249 REVERSE LENGTH=118 | | 2.5231499 | 2.5231499 | 0.39633 |
| AT5G54500 | AT5G54500.1 \| Symbols: FQR1 \| flavodoxin-like quinone reductase 1 \| chr5:22124674-22126256 FORWARD LENGTH=204;AT5G54500.2 \| Symbols: FQR1 \| flavodoxin-like quinone reductase 1 \| chr5:22124674-22126435 FORWARD LENGTH=244;AT4G27270.1 \| Symbols: \| Quinone reductase family protein \| chr4:13661458-13663243 REVERSE LENGTH=205 | 1.9285 | 3.092050339 | 2.510275169 | 0.398362702 |
| AT5G51570 | AT5G51570.1 \| Symbols: \| SPFH/Band 7/PHB domain-containing membrane-associated protein family \| chr5:20949511-20951234 FORWARD LENGTH=292 | 0.47481 | 4.418522446 | 2.446666223 | 0.40871942 |
| AT5G01600 | AT5G01600.1 \| Symbols: ATFER1, FER1 \| ferretin 1 \| chr5:228149-229594 REVERSE LENGTH=255 | 2.427 | | 2.427 | 0.412031314 |
| AT5G16830 | AT5G16830.1 \| Symbols: SYP21, ATSYP21, PEP12, ATPEP12, PEP12P \| syntaxin of plants 21 \| chr5:5533076-5535152 REVERSE LENGTH=279 | | 2.357100766 | 2.357100766 | 0.42425 |
| AT3G18760 | AT3G18760.1 \| Symbols: \| Translation elongation factor EF1B/ribosomal protein S6 family protein \| chr3:6457930-6459558 REVERSE LENGTH=139;AT3G18760.2 \| Symbols: \| Translation elongation factor EF1B/ribosomal protein S6 family protein \| chr3:6457930-6459558 REVERSE LENGTH=134 | | 2.335193704 | 2.335193704 | 0.42823 |
| AT5G23060 | AT5G23060.1 \| Symbols: CaS \| calcium sensing receptor \| chr5:7736760-7738412 REVERSE LENGTH=387 | | 2.312352588 | 2.312352588 | 0.43246 |
| AT1G07870 | AT1G07870.2 \| Symbols: \| Protein kinase superfamily protein \| chr1:2428942-2431843 REVERSE LENGTH=538;AT1G07870.1 \| Symbols: \| Protein kinase superfamily protein \| chr1:2429933-2431843 REVERSE LENGTH=423 | | 2.307337333 | 2.307337333 | 0.4334 |
| AT4G18430 | AT4G18430.1 \| Symbols: AtRABA1e, RABA1e \| RAB GTPase homolog A1E \| chr4:10183903-10185223 REVERSE LENGTH=217 | | 2.275727095 | 2.275727095 | 0.43942 |

| | | | | | |
|---|---|---|---|---|---|
| AT2G40300 | AT2G40300.1 \| Symbols: ATFER4, FER4 \| ferritin 4 \| chr2:16831501-16833214 REVERSE LENGTH=259 | 1.5016 | 3.039790862 | 2.270695431 | 0.440393717 |
| AT4G05320 | [illegible] FORWARD LENGTH=457;AT4G05320.4 \| Symbols: UBQ10 \| polyubiquitin 10 \| chr4:2718559-2719932 FORWARD LENGTH=457;AT5G20620.1 \| Symbols: UBQ4 \| ubiquitin 4 \| chr5:6973315-6974463 REVERSE LENGTH=382;AT4G05320.1 \| Symbols: UBQ10 \| polyubiquitin 10 \| chr4:2718559-2719932 FORWARD LENGTH=381;AT4G05320.3 \| Symbols: UBQ10 \| polyubiquitin 10 \| chr4:2718787-2719932 FORWARD LENGTH=381;AT4G05320.6 \| Symbols: UBQ10 \| polyubiquitin 10 \| chr4:2718559-2719932 FORWARD LENGTH=381;AT5G37640.1 \| Symbols: UBQ9 \| ubiquitin 9 \| chr5:14952782-14953750 REVERSE LENGTH=322;AT1G65350.1 \| Symbols: UBQ13 \| ubiquitin 13 \| chr1:24272518-24277275 REVERSE LENGTH=319;AT5G03240.1 \| Symbols: UBQ3 \| polyubiquitin 3 \| chr5:771976-772896 REVERSE LENGTH=306;AT5G03240.2 \| Symbols: UBQ3 \| polyubiquitin 3 \| chr5:771976-772896 REVERSE LENGTH=306;AT5G03240.3 \| Symbols: UBQ3 \| polyubiquitin 3 \| chr5:771976-772896 REVERSE LENGTH=306;AT4G02890.3 \| Symbols: UBQ14 \| Ubiquitin family protein \| chr4:1278747-1279664 REVERSE LENGTH=305;AT4G02890.4 \| Symbols: UBQ14 \| Ubiquitin family protein \| chr4:1278747-1279664 REVERSE LENGTH=305;AT4G05320.5 \| Symbols: UBQ10 \| polyubiquitin 10 \| chr4:2718559-2719932 FORWARD LENGTH=305;AT1G55060.1 \| Symbols: UBQ12 \| ubiquitin 12 \| chr1:20549533-20550225 FORWARD LENGTH=230;AT4G02890.1 \| Symbols: UBQ14 \| Ubiquitin family protein \| chr4:1278747-1279664 REVERSE LENGTH=229;AT4G02890.2 \| Symbols: UBQ14 \| Ubiquitin family protein \| chr4:1278747-1279664 REVERSE LENGTH=229;AT4G05050.1 \| Symbols: UBQ11 \| ubiquitin 11 \| chr4:2588271-2588960 REVERSE LENGTH=229;AT4G05050.2 \| Symbols: UBQ11 \| ubiquitin 11 \| chr4:2588271-2588960 REVERSE LENGTH=229;AT4G05050.3 \| Symbols: UBQ11 \| ubiquitin 11 \| chr4:2588271-2588960 REVERSE LENGTH=229;AT2G47110.1 \| Symbols: UBQ6 \| ubiquitin 6 \| chr2:19344701-19345174 FORWARD LENGTH=157;AT2G47110.2 \| Symbols: | 2.2553 | | 2.2553 | 0.443399991 |
| AT3G07020 | AT3G07020.1 \| Symbols: \| UDP-Glycosyltransferase superfamily protein \| chr3:2217841-2221590 REVERSE LENGTH=637;AT3G07020.2 \| Symbols: \| UDP-Glycosyltransferase superfamily protein \| chr3:2218120-2221590 REVERSE LENGTH=637 | | 2.230947707 | 2.230947707 | 0.44824 |
| AT3G28450 | AT3G28450.1 \| Symbols: \| Leucine-rich repeat protein kinase family protein \| chr3:10667359-10669176 FORWARD LENGTH=605 | | 2.226130318 | 2.226130318 | 0.44921 |
| AT3G54030 | AT3G54030.1 \| Symbols: \| Protein kinase protein with tetratricopeptide repeat domain \| chr3:20011162-20013490 FORWARD LENGTH=490 | 1.533 | 2.902167919 | 2.21758396 | 0.450941213 |
| AT1G13440 | AT1G13440.1 \| Symbols: GAPC-2, GAPC2 \| glyceraldehyde-3-phosphate dehydrogenase C2 \| chr1:4608465-4610494 REVERSE LENGTH=338;AT3G04120.1 \| Symbols: GAPC, GAPC-1, GAPC1 \| glyceraldehyde-3-phosphate dehydrogenase C subunit 1 \| chr3:1081077-1083131 FORWARD LENGTH=338;AT1G13440.2 \| Symbols: GAPC-2, GAPC2 \| glyceraldehyde-3-phosphate dehydrogenase C2 \| chr1:4608465-4610494 REVERSE LENGTH=310 | 0.60361 | 3.819126184 | 2.211368092 | 0.45220875 |
| AT3G28710 | AT3G28710.1 \| Symbols: \| ATPase, V0/A0 complex, subunit C/D \| chr3:10773144-10775594 REVERSE LENGTH=351;AT3G28715.1 \| Symbols: \| ATPase, V0/A0 complex, subunit C/D \| chr3:10778025-10780350 FORWARD LENGTH=351;AT3G28715.2 \| Symbols: \| ATPase, V0/A0 complex, subunit C/D \| chr3:10778025-10780350 FORWARD LENGTH=343 | 0.5239 | 3.815774411 | 2.169837206 | 0.460864067 |
| AT1G31130 | AT1G31130.1 \| Symbols: \| unknown protein; FUNCTIONS IN: molecular_function unknown; INVOLVED IN: biological_process unknown; LOCATED IN: cellular_component unknown; EXPRESSED IN: 23 plant structures; EXPRESSED DURING: 13 growth stages; BEST Arabidopsis thaliana protein match is: unknown protein (TAIR:AT4G19950.1); Has 246 Blast hits to 244 proteins in 29 species: Archae - 2; Bacteria - 16; Metazoa - 0; Fungi - 0; Plants - 222; Viruses - 0; Other Eukaryotes - 6 (source: NCBI BLink). \| chr1:11114963-11115928 REVERSE LENGTH=321 | 2.8842 | 1.433897333 | 2.159048666 | 0.463166957 |

| | | | | | |
|---|---|---|---|---|---|
| AT5G16050 | AT5G16050.1 \| Symbols: GRF5, GF14 UPSILON \| general regulatory factor 5 \| chr5:5244008-5245402 REVERSE LENGTH=268;AT3G02520.1 \| Symbols: GRF7, GF14 NU \| general regulatory factor 7 \| chr3:526800-527915 REVERSE LENGTH=265 | 0.59901 | 3.707960992 | 2.153485496 | 0.464363471 |
| AT3G12490 | AT3G12490.2 \| Symbols: ATCYSB, ATCYS6, CYSB \| cystatin B \| chr3:3960523-3961876 REVERSE LENGTH=234;AT3G12490.1 \| Symbols: ATCYSB, ATCYS6, CYSB \| cystatin B \| chr3:3960523-3961777 REVERSE LENGTH=201 | 0.52124 | 3.770454717 | 2.145847358 | 0.466016372 |
| AT4G02450 | AT4G02450.1 \| Symbols: \| HSP20-like chaperones superfamily protein \| chr4:1073987-1075765 REVERSE LENGTH=241;AT4G02450.2 \| Symbols: \| HSP20-like chaperones superfamily protein \| chr4:1073987-1075765 REVERSE LENGTH=240 | | 2.143668675 | 2.143668675 | 0.46649 |
| AT1G28340 | AT1G28340.1 \| Symbols: AtRLP4, RLP4 \| receptor like protein 4 \| chr1:9940175-9943252 FORWARD LENGTH=626 | | 2.143255176 | 2.143255176 | 0.46658 |
| AT4G31810 | AT4G31810.1 \| Symbols: \| ATP-dependent caseinolytic (Clp) protease/crotonase family protein \| chr4:15387365-15390290 REVERSE LENGTH=409 | | 2.138579983 | 2.138579983 | 0.4676 |
| AT2G26300 | AT2G26300.1 \| Symbols: GPA1, GP ALPHA 1, ATGPA1 \| G protein alpha subunit 1 \| chr2:11198087-11200822 FORWARD LENGTH=383 | | 2.117836418 | 2.117836418 | 0.47218 |
| AT5G67380 | AT5G67380.1 \| Symbols: CKA1, ATCKA1 \| casein kinase alpha 1 \| chr5:26881156-26883383 REVERSE LENGTH=409;AT3G50000.1 \| Symbols: CKA2, ATCKA2 \| casein kinase II, alpha chain 2 \| chr3:18534487-18536743 FORWARD LENGTH=403;AT5G67380.2 \| Symbols: CKA1, ATCKA1 \| casein kinase alpha 1 \| chr5:26881156-26883383 REVERSE LENGTH=376;AT2G23080.1 \| Symbols: \| Protein kinase superfamily protein \| chr2:9827228-9829343 FORWARD LENGTH=333;AT2G23080.2 \| Symbols: \| Protein kinase superfamily protein \| chr2:9827228-9829057 FORWARD LENGTH=307 | | 2.108903792 | 2.108903792 | 0.47418 |
| AT5G42980 | AT5G42980.1 \| Symbols: ATTRX3, ATH3, ATTRXH3, TRXH3, TRX3 \| thioredoxin 3 \| chr5:17242772-17243718 FORWARD LENGTH=118 | | 2.094855036 | 2.094855036 | 0.47736 |
| AT2G37340 | AT2G37340.1 \| Symbols: RSZ33, ATRSZ33, RS2Z33, AT-RS2Z33 \| arginine/serine-rich zinc knuckle-containing protein 33 \| chr2:15670372-15672331 REVERSE LENGTH=290;AT2G37340.3 \| Symbols: RSZ33, RS2Z33, AT-RS2Z33 \| arginine/serine-rich zinc knuckle-containing protein 33 \| chr2:15670372-15671584 REVERSE LENGTH=249 | 0.36532 | 3.754599384 | 2.059959692 | 0.485446392 |
| AT5G51100 | AT5G51100.1 \| Symbols: FSD2 \| Fe superoxide dismutase 2 \| chr5:20773357-20775635 REVERSE LENGTH=305 | | 2.056090139 | 2.056090139 | 0.48636 |
| AT1G61150 | AT1G61150.1 \| Symbols: \| LisH and RanBPM domains containing protein \| chr1:22542928-22544355 FORWARD LENGTH=243;AT1G61150.2 \| Symbols: \| LisH and RanBPM domains containing protein \| chr1:22542979-22544355 FORWARD LENGTH=226;AT1G61150.4 \| Symbols: \| LisH and RanBPM domains containing protein \| chr1:22542979-22544355 FORWARD LENGTH=226;AT1G61150.6 \| Symbols: \| LisH and RanBPM domains containing protein \| chr1:22542979-22544355 FORWARD LENGTH=226;AT1G61150.5 \| Symbols: \| LisH and RanBPM domains containing protein \| chr1:22543122-22544355 FORWARD LENGTH=220;AT1G61150.3 \| Symbols: \| LisH and RanBPM domains containing protein \| chr1:22543304-22544355 FORWARD LENGTH=193;AT1G61150.7 \| Symbols: \| LisH and RanBPM domains containing protein \| chr1:22543304-22544355 FORWARD LENGTH=193 | | 2.018041289 | 2.018041289 | 0.49553 |
| AT3G53500 | AT3G53500.2 \| Symbols: RSZ32, RS2Z32, At-RS2Z \| RNA-binding (RRM/RBD/RNP motifs) family protein with retrovirus zinc finger-like domain \| chr3:19834557-19836507 REVERSE LENGTH=284;AT3G53500.1 \| Symbols: RSZ32, RS2Z32, At-RS2Z \| RNA-binding (RRM/RBD/RNP motifs) family protein with retrovirus zinc finger-like domain \| chr3:19834557-19835896 REVERSE LENGTH=243 | | 2.01134398 | 2.01134398 | 0.49718 |
| AT4G38630 | AT4G38630.1 \| Symbols: RPN10, MCB1, ATMCB1, MBP1 \| regulatory particle non-ATPase 10 \| chr4:18057357-18059459 REVERSE LENGTH=386 | | 1.984599508 | 1.984599508 | 0.50388 |

| | | | | | |
|---|---|---|---|---|---|
| AT2G25060 | AT2G25060.1 \| Symbols: ENODL14, AtENODL14 \| early nodulin-like protein 14 \| chr2:10662308-10662930 FORWARD LENGTH=182 | | 1.971064769 | 1.971064769 | 0.50734 |
| AT4G19006 | AT4G19006.1 \| Symbols: \| Proteasome component (PCI) domain protein \| chr4:10409349-10411347 REVERSE LENGTH=386 | | 1.963170914 | 1.963170914 | 0.50938 |
| AT5G48450 | AT5G48450.1 \| Symbols: sks3 \| SKU5 similar 3 \| chr5:19632791-19635612 REVERSE LENGTH=621 | 1.961 | 1.855115481 | 1.90805774 | 0.524093154 |
| AT3G18240 | AT3G18240.1 \| Symbols: \| Ribosomal protein S24/S35, mitochondrial \| chr3:6255810-6257695 FORWARD LENGTH=419;AT3G18240.2 \| Symbols: \| Ribosomal protein S24/S35, mitochondrial \| chr3:6255810-6257695 FORWARD LENGTH=419;AT4G21460.1 \| Symbols: \| Ribosomal protein S24/S35, mitochondrial \| chr4:11428530-11430390 FORWARD LENGTH=415 | | 1.886116298 | 1.886116298 | 0.53019 |
| AT5G12140 | AT5G12140.1 \| Symbols: ATCYS1, CYS1 \| cystatin-1 \| chr5:3923295-3923936 REVERSE LENGTH=101 | | 1.87892225 | 1.87892225 | 0.53222 |
| AT5G25040 | AT5G25040.2 \| Symbols: \| Major facilitator superfamily protein \| chr5:8622185-8623965 FORWARD LENGTH=517;AT5G25040.1 \| Symbols: \| Major facilitator superfamily protein \| chr5:8622185-8623965 FORWARD LENGTH=457 | | 1.847814036 | 1.847814036 | 0.54118 |
| AT5G10450 | AT5G10450.4 \| Symbols: GRF6 \| G-box regulating factor 6 \| chr5:3284476-3286261 REVERSE LENGTH=273;AT5G10450.3 \| Symbols: GRF6 \| G-box regulating factor 6 \| chr5:3284615-3286261 REVERSE LENGTH=258;AT5G10450.1 \| Symbols: GRF6, AFT1, 14-3-3lambda \| G-box regulating factor 6 \| chr5:3284452-3286261 REVERSE LENGTH=248;AT5G10450.2 \| Symbols: GRF6, AFT1, 14-3-3lambda \| G-box regulating factor 6 \| chr5:3284452-3286261 REVERSE LENGTH=246 | | 1.842604706 | 1.842604706 | 0.54271 |
| AT1G41880 | AT1G41880.1 \| Symbols: \| Ribosomal protein L35Ae family protein \| chr1:15651585-15652427 REVERSE LENGTH=111;AT1G07070.1 \| Symbols: \| Ribosomal protein L35Ae family protein \| chr1:2168652-2169703 FORWARD LENGTH=112 | | 1.816233495 | 1.816233495 | 0.55059 |
| AT5G64390 | AT5G64390.3 \| Symbols: HEN4 \| RNA-binding KH domain-containing protein \| chr5:25743442-25747380 FORWARD LENGTH=866;AT5G64390.1 \| Symbols: HEN4 \| RNA-binding KH domain-containing protein \| chr5:25743442-25747380 FORWARD LENGTH=857;AT5G64390.2 \| Symbols: HEN4 \| RNA-binding KH domain-containing protein \| chr5:25743442-25746742 FORWARD LENGTH=824 | 1.8146 | | 1.8146 | 0.551085639 |
| AT3G53870 | AT3G53870.1 \| Symbols: \| Ribosomal protein S3 family protein \| chr3:19951547-19952782 FORWARD LENGTH=249 | | 1.814486863 | 1.814486863 | 0.55112 |
| AT1G06840 | AT1G06840.1 \| Symbols: \| Leucine-rich repeat protein kinase family protein \| chr1:2097854-2103208 REVERSE LENGTH=953 | | 1.810413498 | 1.810413498 | 0.55236 |
| AT3G57990 | AT3G57990.1 \| Symbols: \| unknown protein; Has 1497 Blast hits to 1323 proteins in 52 species: Archae - 0; Bacteria - 4; Metazoa - 23; Fungi - 34; Plants - 61; Viruses - 0; Other Eukaryotes - 1375 (source: NCBI BLink). \| chr3:21470244-21471347 REVERSE LENGTH=367 | | 1.762860064 | 1.762860064 | 0.56726 |
| AT1G16870 | AT1G16870.1 \| Symbols: \| mitochondrial 28S ribosomal protein S29-related \| chr1:5770964-5773157 REVERSE LENGTH=480 | | 1.757500132 | 1.757500132 | 0.56899 |
| AT1G55890 | AT1G55890.1 \| Symbols: \| Tetratricopeptide repeat (TPR)-like superfamily protein \| chr1:20901364-20902560 FORWARD LENGTH=398 | | 1.751374829 | 1.751374829 | 0.57098 |
| AT3G02230 | AT3G02230.1 \| Symbols: RGP1, ATRGP1 \| reversibly glycosylated polypeptide 1 \| chr3:415463-417304 FORWARD LENGTH=357 | | 1.734304544 | 1.734304544 | 0.5766 |
| AT5G45620 | AT5G45620.1 \| Symbols: \| Proteasome component (PCI) domain protein \| chr5:18501590-18503868 FORWARD LENGTH=386;AT5G45620.2 \| Symbols: \| Proteasome component (PCI) domain protein \| chr5:18501590-18503422 FORWARD LENGTH=350 | | 1.729894303 | 1.729894303 | 0.57807 |

| | | | | | |
|---|---|---|---|---|---|
| AT3G47070 | AT3G47070.1 \| Symbols: \| LOCATED IN: thylakoid, chloroplast thylakoid membrane, chloroplast, chloroplast envelope; EXPRESSED IN: 22 plant structures; EXPRESSED DURING: 13 growth stages; CONTAINS InterPro DOMAIN/s: Thylakoid soluble phosphoprotein TSP9 (InterPro:IPR021584); Has 37 Blast hits to 37 proteins in 10 species: Archae - 0; Bacteria - 0; Metazoa - 0; Fungi - 0; Plants - 37; Viruses - 0; Other Eukaryotes - 0 (source: NCBI BLink). \| chr3:17337205-17337507 REVERSE LENGTH=100 | | 1.7188628 | 1.7188628 | 0.58178 |
| AT1G63780 | AT1G63780.1 \| Symbols: IMP4 \| Ribosomal RNA processing Brix domain protein \| chr1:23665045-23667243 REVERSE LENGTH=294 | | 1.662869781 | 1.662869781 | 0.60137 |
| AT2G40660 | AT2G40660.1 \| Symbols: \| Nucleic acid-binding, OB-fold-like protein \| chr2:16966011-16968866 FORWARD LENGTH=389 | | 1.630044989 | 1.630044989 | 0.61348 |
| AT3G61860 | AT3G61860.1 \| Symbols: ATRSP31, RSP31, At-RS31, RS31 \| RNA-binding (RRM/RBD/RNP motifs) family protein \| chr3:22900311-22902159 REVERSE LENGTH=264;AT2G46610.1 \| Symbols: RS31a, At-RS31a \| RNA-binding (RRM/RBD/RNP motifs) family protein \| chr2:19136769-19138492 REVERSE LENGTH=250;AT2G46610.2 \| Symbols: RS31a, At-RS31a \| RNA-binding (RRM/RBD/RNP motifs) family protein \| chr2:19136769-19137858 REVERSE LENGTH=224 | 0.13477 | 3.099814011 | 1.617292006 | 0.618317531 |
| AT2G37970 | AT2G37970.1 \| Symbols: SOUL-1 \| SOUL heme-binding family protein \| chr2:15891027-15891704 FORWARD LENGTH=225 | 0.43543 | 2.781099647 | 1.608264823 | 0.621788144 |
| AT1G07320 | AT1G07320.1 \| Symbols: RPL4 \| ribosomal protein L4 \| chr1:2249190-2250189 FORWARD LENGTH=282;AT1G07320.2 \| Symbols: RPL4 \| ribosomal protein L4 \| chr1:2249190-2250189 FORWARD LENGTH=280;AT1G07320.3 \| Symbols: RPL4 \| ribosomal protein L4 \| chr1:2249190-2250026 FORWARD LENGTH=278;AT1G07320.4 \| Symbols: RPL4 \| ribosomal protein L4 \| chr1:2249190-2250173 FORWARD LENGTH=278 | | 1.598491024 | 1.598491024 | 0.62559 |
| AT2G34585 | AT2G34585.1 \| Symbols: \| unknown protein; FUNCTIONS IN: molecular_function unknown; INVOLVED IN: biological_process unknown; LOCATED IN: plasma membrane, vacuole; EXPRESSED IN: 24 plant structures; EXPRESSED DURING: 15 growth stages; Has 43 Blast hits to 43 proteins in 9 species: Archae - 0; Bacteria - 0; Metazoa - 0; Fungi - 0; Plants - 43; Viruses - 0; Other Eukaryotes - 0 (source: NCBI BLink). \| chr2:14568227-14568472 REVERSE LENGTH=81 | | 1.580128306 | 1.580128306 | 0.63286 |
| AT3G60830 | AT3G60830.1 \| Symbols: ATARP7, ARP7 \| actin-related protein 7 \| chr3:22474298-22476000 FORWARD LENGTH=363 | | 1.576416805 | 1.576416805 | 0.63435 |
| AT5G62890 | AT5G62890.1 \| Symbols: \| Xanthine/uracil permease family protein \| chr5:25243723-25247075 FORWARD LENGTH=532;AT5G62890.2 \| Symbols: \| Xanthine/uracil permease family protein \| chr5:25243723-25247075 FORWARD LENGTH=532;AT5G62890.3 \| Symbols: \| Xanthine/uracil permease family protein \| chr5:25243723-25247075 FORWARD LENGTH=532;AT1G60030.1 \| Symbols: ATNAT7, NAT7 \| nucleobase-ascorbate transporter 7 \| chr1:22113993-22116648 REVERSE LENGTH=538;AT5G62890.4 \| Symbols: \| Xanthine/uracil permease family protein \| chr5:25243723-25246748 FORWARD LENGTH=476 | | 1.563403843 | 1.563403843 | 0.63963 |
| AT5G04950 | AT5G04950.1 \| Symbols: NAS1, ATNAS1 \| nicotianamine synthase 1 \| chr5:1457876-1458838 REVERSE LENGTH=320 | | 1.521583664 | 1.521583664 | 0.65721 |
| AT2G36530 | AT2G36530.1 \| Symbols: LOS2, ENO2 \| Enolase \| chr2:15321081-15323786 REVERSE LENGTH=444 | 0.49285 | 2.537555826 | 1.515202913 | 0.659977612 |
| AT3G18790 | AT3G18790.1 \| Symbols: \| FUNCTIONS IN: molecular_function unknown; INVOLVED IN: biological_process unknown; LOCATED IN: chloroplast; EXPRESSED IN: 22 plant structures; EXPRESSED DURING: 13 growth stages; CONTAINS InterPro DOMAIN/s: Isy1-like splicing (InterPro:IPR009360); Has 1147 Blast hits to 965 proteins in 236 species: Archae - 12; Bacteria - 13; Metazoa - 351; Fungi - 230; Plants - 49; Viruses - 9; Other Eukaryotes - 483 (source: NCBI BLink). \| chr3:6477853-6478755 FORWARD LENGTH=300 | | 1.491802545 | 1.491802545 | 0.67033 |

| | | | | | |
|---|---|---|---|---|---|
| AT1G64200 | AT1G64200.1 \| Symbols: VHA-E3 \| vacuolar H+-ATPase subunit E isoform 3 \| chr1:23828537-23830002 REVERSE LENGTH=237 | | 1.484053841 | 1.484053841 | 0.67383 |
| AT2G15430 | AT2G15430.1 \| Symbols: RBP36A, RPB35.5A, NRPB3, NRPD3, NRPE3A \| DNA-directed RNA polymerase family protein \| chr2:6733661-6735482 FORWARD LENGTH=319 | | 1.481108462 | 1.481108462 | 0.67517 |
| AT3G01390 | AT3G01390.1 \| Symbols: VMA10, AVMA10 \| vacuolar membrane ATPase 10 \| chr3:150265-150922 REVERSE LENGTH=110;AT3G01390.2 \| Symbols: VMA10, AVMA10 \| vacuolar membrane ATPase 10 \| chr3:150265-150922 REVERSE LENGTH=110 | 1.6914 | 1.189626457 | 1.440513229 | 0.694197027 |
| AT5G64080 | AT5G64080.1 \| Symbols: \| Bifunctional inhibitor/lipid-transfer protein/seed storage 2S albumin superfamily protein \| chr5:25645475-25646638 REVERSE LENGTH=182;AT5G64080.2 \| Symbols: \| Bifunctional inhibitor/lipid-transfer protein/seed storage 2S albumin superfamily protein \| chr5:25645475-25646638 REVERSE LENGTH=178 | | 1.413467519 | 1.413467519 | 0.70748 |
| ATMG00090 | ATMG00090.1 \| Symbols: \| structural constituent of ribosome;protein binding \| chrM:25482-28733 REVERSE LENGTH=556 | | 1.395537072 | 1.395537072 | 0.71657 |
| AT3G15760 | AT3G15760.1 \| Symbols: \| unknown protein; FUNCTIONS IN: molecular_function unknown; INVOLVED IN: biological_process unknown; LOCATED IN: endomembrane system; EXPRESSED IN: 22 plant structures; EXPRESSED DURING: 10 growth stages; BEST Arabidopsis thaliana protein match is: unknown protein (TAIR:AT1G52565.1); Has 42 Blast hits to 42 proteins in 10 species: Archae - 0; Bacteria - 0; Metazoa - 0; Fungi - 0; Plants - 42; Viruses - 0; Other Eukaryotes - 0 (source: NCBI BLink). \| chr3:5337734-5338219 FORWARD LENGTH=135 | | 1.39215659 | 1.39215659 | 0.71831 |
| AT5G57460 | AT5G57460.1 \| Symbols: \| unknown protein; FUNCTIONS IN: molecular_function unknown; INVOLVED IN: biological_process unknown; LOCATED IN: plasma membrane, chloroplast; EXPRESSED IN: 23 plant structures; EXPRESSED DURING: 13 growth stages; Has 166 Blast hits to 166 proteins in 41 species: Archae - 0; Bacteria - 0; Metazoa - 112; Fungi - 4; Plants - 36; Viruses - 0; Other Eukaryotes - 14 (source: NCBI BLink). \| chr5:23275321-23277261 FORWARD LENGTH=646 | | 1.379082083 | 1.379082083 | 0.72512 |
| AT3G49400 | AT3G49400.1 \| Symbols: \| Transducin/WD40 repeat-like superfamily protein \| chr3:18319249-18324373 REVERSE LENGTH=892;AT3G49400.2 \| Symbols: \| Transducin/WD40 repeat-like superfamily protein \| chr3:18319249-18324373 REVERSE LENGTH=874 | | 1.347454658 | 1.347454658 | 0.74214 |
| AT3G63520 | AT3G63520.1 \| Symbols: CCD1, ATCCD1, ATNCED1, NCED1 \| carotenoid cleavage dioxygenase 1 \| chr3:23452940-23455896 FORWARD LENGTH=538 | | 1.322873811 | 1.322873811 | 0.75593 |
| AT5G19990 | AT5G19990.1 \| Symbols: RPT6A, ATSUG1 \| regulatory particle triple-A ATPase 6A \| chr5:6752144-6754918 FORWARD LENGTH=419;AT5G20000.1 \| Symbols: \| AAA-type ATPase family protein \| chr5:6756915-6759550 FORWARD LENGTH=419 | | 1.302948573 | 1.302948573 | 0.76749 |
| AT2G07360 | AT2G07360.2 \| Symbols: \| SH3 domain-containing protein \| chr2:3047623-3057099 REVERSE LENGTH=1198;AT2G07360.1 \| Symbols: \| SH3 domain-containing protein \| chr2:3047623-3057099 REVERSE LENGTH=1196 | | 1.293510458 | 1.293510458 | 0.77309 |
| AT5G62270 | AT5G62270.2 \| Symbols: \| FUNCTIONS IN: molecular_function unknown; INVOLVED IN: biological_process unknown; LOCATED IN: cellular_component unknown; BEST Arabidopsis thaliana protein match is: mucin-related (TAIR:AT2G02880.1). \| chr5:25012227-25014486 FORWARD LENGTH=420;AT5G62270.1 \| Symbols: \| BEST Arabidopsis thaliana protein match is: mucin-related (TAIR:AT2G02880.1); Has 35333 Blast hits to 34131 proteins in 2444 species: Archae - 798; Bacteria - 22429; Metazoa - 974; Fungi - 991; Plants - 531; Viruses - 0; Other Eukaryotes - 9610 (source: NCBI BLink). \| chr5:25012227-25014007 FORWARD LENGTH=383 | | 1.289673584 | 1.289673584 | 0.77539 |

| | | | | | |
|---|---|---|---|---|---|
| AT3G58730 | AT3G58730.1 \| Symbols: \| vacuolar ATP synthase subunit D (VATD) / V-ATPase D subunit / vacuolar proton pump D subunit (VATPD) \| chr3:21718495-21719280 REVERSE LENGTH=261 | 0.30973 | 2.26326272 | 1.28649636 | 0.777304959 |
| AT1G02280 | AT1G02280.1 \| Symbols: TOC33, ATTOC33, PPI1 \| translocon at the outer envelope membrane of chloroplasts 33 \| chr1:448665-450246 REVERSE LENGTH=297;AT1G02280.2 \| Symbols: TOC33, ATTOC33, PPI1 \| translocon at the outer envelope membrane of chloroplasts 33 \| chr1:448665-450246 REVERSE LENGTH=297 | | 1.279868941 | 1.279868941 | 0.78133 |
| AT5G56030 | AT5G56030.2 \| Symbols: HSP81-2 \| heat shock protein 81-2 \| chr5:22686923-22689433 FORWARD LENGTH=728;AT5G56000.1 \| Symbols: Hsp81.4, AtHsp90.4 \| HEAT SHOCK PROTEIN 81.4 \| chr5:22677602-22680067 REVERSE LENGTH=699;AT5G56010.1 \| Symbols: HSP81-3, Hsp81.3, AtHsp90-3, AtHsp90.3 \| heat shock protein 81-3 \| chr5:22681410-22683911 FORWARD LENGTH=699;AT5G56030.1 \| Symbols: HSP81-2, ERD8, HSP90.2, AtHsp90.2 \| heat shock protein 81-2 \| chr5:22686923-22689433 FORWARD LENGTH=699;AT5G52640.1 \| Symbols: HSP81-1, ATHS83, HSP81.1, HSP83, ATHSP90.1, AtHsp90-1, HSP90.1 \| heat shock protein 90.1 \| chr5:21352542-21355147 FORWARD LENGTH=705 | | 1.267523513 | 1.267523513 | 0.78894 |
| AT1G77750 | AT1G77750.1 \| Symbols: \| Ribosomal protein S13/S18 family \| chr1:29230346-29231490 REVERSE LENGTH=154 | | 1.263743207 | 1.263743207 | 0.7913 |
| AT5G52040 | AT5G52040.2 \| Symbols: ATRSP41, RS41, At-RS41 \| RNA-binding (RRM/RBD/RNP motifs) family protein \| chr5:21131081-21133318 FORWARD LENGTH=357;AT5G52040.1 \| Symbols: ATRSP41, RS41, At-RS41 \| RNA-binding (RRM/RBD/RNP motifs) family protein \| chr5:21131081-21133318 FORWARD LENGTH=356;AT5G52040.3 \| Symbols: RS41, At-RS41 \| RNA-binding (RRM/RBD/RNP motifs) family protein \| chr5:21131791-21133318 FORWARD LENGTH=329;AT5G52040.4 \| Symbols: RS41, At-RS41 \| RNA-binding (RRM/RBD/RNP motifs) family protein \| chr5:21131881-21133318 FORWARD LENGTH=324 | | 1.258589876 | 1.258589876 | 0.79454 |
| AT3G17465 | AT3G17465.1 \| Symbols: RPL3P \| ribosomal protein L3 plastid \| chr3:5978059-5979572 REVERSE LENGTH=324 | | 1.255776572 | 1.255776572 | 0.79632 |
| AT5G09310 | AT5G09310.1 \| Symbols: \| CONTAINS InterPro DOMAIN/s: Gamma-secretase aspartyl protease complex, presenilin enhancer-2 subunit (InterPro:IPR019379); Has 168 Blast hits to 168 proteins in 71 species: Archae - 0; Bacteria - 0; Metazoa - 126; Fungi - 0; Plants - 36; Viruses - 0; Other Eukaryotes - 6 (source: NCBI BLink). \| chr5:2887403-2888501 REVERSE LENGTH=146 | | 1.252850234 | 1.252850234 | 0.79818 |
| AT2G04390 | AT2G04390.1 \| Symbols: \| Ribosomal S17 family protein \| chr2:1527911-1528336 FORWARD LENGTH=141;AT5G04800.1 \| Symbols: \| Ribosomal S17 family protein \| chr5:1389217-1389642 FORWARD LENGTH=141;AT5G04800.2 \| Symbols: \| Ribosomal S17 family protein \| chr5:1389217-1389642 FORWARD LENGTH=141;AT5G04800.3 \| Symbols: \| Ribosomal S17 family protein \| chr5:1389217-1389642 FORWARD LENGTH=141;AT5G04800.4 \| Symbols: \| Ribosomal S17 family protein \| chr5:1389217-1389642 FORWARD LENGTH=141;AT2G05220.1 \| Symbols: \| Ribosomal S17 family protein \| chr2:1894757-1895179 REVERSE LENGTH=140;AT2G05220.2 \| Symbols: \| Ribosomal S17 family protein \| chr2:1894757-1895179 REVERSE LENGTH=140;AT3G10610.1 \| Symbols: \| Ribosomal S17 family protein \| chr3:3319459-3319881 FORWARD LENGTH=140 | 1.8346 | 0.650956907 | 1.242778453 | 0.804648646 |
| AT3G24160 | AT3G24160.1 \| Symbols: PMP \| putative type 1 membrane protein \| chr3:8726241-8729023 FORWARD LENGTH=364 | 1.6559 | 0.793839803 | 1.224869902 | 0.816413236 |
| AT4G26570 | AT4G26570.2 \| Symbols: ATCBL3, CBL3 \| calcineurin B-like 3 \| chr4:13408608-13409998 REVERSE LENGTH=230;AT4G26570.1 \| Symbols: ATCBL3, CBL3 \| calcineurin B-like 3 \| chr4:13408608-13409998 REVERSE LENGTH=226;AT5G55990.1 \| Symbols: CBL2, ATCBL2 \| calcineurin B-like protein 2 \| chr5:22672189-22673579 FORWARD LENGTH=226 | 0.30435 | 2.076800066 | 1.190575033 | 0.839930262 |

| | | | | | |
|---|---|---|---|---|---|
| AT1G64520 | AT1G64520.1 \| Symbols: RPN12a \| regulatory particle non-ATPase 12A \| chr1:23956459-23958120 FORWARD LENGTH=267 | 0.21553 | 2.149659279 | 1.18259464 | 0.845598286 |
| AT2G37990 | AT2G37990.1 \| Symbols: \| ribosome biogenesis regulatory protein (RRS1) family protein \| chr2:15900713-15903028 FORWARD LENGTH=318 | 0.16077 | 2.203468259 | 1.18211913 | 0.84593843 |
| AT2G01650 | AT2G01650.1 \| Symbols: PUX2 \| plant UBX domain-containing protein 2 \| chr2:284851-286394 REVERSE LENGTH=458 | | 1.167351513 | 1.167351513 | 0.85664 |
| AT5G15490 | AT5G15490.1 \| Symbols: \| UDP-glucose 6-dehydrogenase family protein \| chr5:5027872-5029314 REVERSE LENGTH=480;AT1G26570.1 \| Symbols: UGD1, ATUGD1 \| UDP-glucose dehydrogenase 1 \| chr1:9182801-9184246 FORWARD LENGTH=481;AT3G29360.1 \| Symbols: \| UDP-glucose 6-dehydrogenase family protein \| chr3:11267375-11268817 REVERSE LENGTH=480;AT3G29360.2 \| Symbols: \| UDP-glucose 6-dehydrogenase family protein \| chr3:11267375-11268817 REVERSE LENGTH=480;AT5G39320.1 \| Symbols: \| UDP-glucose 6-dehydrogenase family protein \| chr5:15743254-15744696 FORWARD LENGTH=480 | | 1.161912508 | 1.161912508 | 0.86065 |
| AT5G47320 | AT5G47320.1 \| Symbols: RPS19 \| ribosomal protein S19 \| chr5:19203801-19204951 FORWARD LENGTH=212 | | 1.147038919 | 1.147038919 | 0.87181 |
| AT2G38620 | AT2G38620.2 \| Symbols: CDKB1;2 \| cyclin-dependent kinase B1;2 \| chr2:16152551-16153866 FORWARD LENGTH=311;AT3G54180.1 \| Symbols: CDC2B, CDKB1;1 \| cyclin-dependent kinase B1;1 \| chr3:20059882-20061250 FORWARD LENGTH=309;AT2G38620.1 \| Symbols: CDKB1;2 \| cyclin-dependent kinase B1;2 \| chr2:16152551-16153406 FORWARD LENGTH=257 | | 1.146670642 | 1.146670642 | 0.87209 |
| AT1G56630 | AT1G56630.1 \| Symbols: \| alpha/beta-Hydrolases superfamily protein \| chr1:21223593-21226943 REVERSE LENGTH=485 | 1.1442 | | 1.1442 | 0.873973082 |
| AT4G31840 | AT4G31840.1 \| Symbols: ENODL15, AtENODL15 \| early nodulin-like protein 15 \| chr4:15401798-15402426 FORWARD LENGTH=177 | 1.1376 | | 1.1376 | 0.879043601 |
| AT5G05780 | AT5G05780.1 \| Symbols: RPN8A, AE3, ATHMOV34 \| RP non-ATPase subunit 8A \| chr5:1735862-1738176 FORWARD LENGTH=308;AT5G05780.2 \| Symbols: RPN8A \| RP non-ATPase subunit 8A \| chr5:1735862-1738176 FORWARD LENGTH=305 | 0.1344 | 2.135520106 | 1.134960053 | 0.881088279 |
| AT4G13010 | AT4G13010.1 \| Symbols: \| Oxidoreductase, zinc-binding dehydrogenase family protein \| chr4:7600682-7602567 FORWARD LENGTH=329 | | 1.124935316 | 1.124935316 | 0.88894 |
| AT5G64200 | AT5G64200.1 \| Symbols: ATSC35, SC35, At-SC35 \| ortholog of human splicing factor SC35 \| chr5:25681849-25683553 REVERSE LENGTH=303;AT5G64200.2 \| Symbols: ATSC35, SC35, At-SC35 \| ortholog of human splicing factor SC35 \| chr5:25681849-25683553 REVERSE LENGTH=303 | | 1.113139499 | 1.113139499 | 0.89836 |
| AT5G65220 | AT5G65220.1 \| Symbols: \| Ribosomal L29 family protein \| chr5:26061301-26062506 FORWARD LENGTH=173 | | 1.111531023 | 1.111531023 | 0.89966 |
| AT5G19550 | AT5G19550.1 \| Symbols: ASP2, AAT2 \| aspartate aminotransferase 2 \| chr5:6598201-6601597 FORWARD LENGTH=405 | 0.36273 | 1.860292066 | 1.111511033 | 0.89967618 |
| AT1G12840 | AT1G12840.1 \| Symbols: DET3, ATVHA-C \| vacuolar ATP synthase subunit C (VATC) / V-ATPase C subunit / vacuolar proton pump C subunit (DET3) \| chr1:4375584-4378220 FORWARD LENGTH=375 | 0.22394 | 1.99832141 | 1.111130705 | 0.899984129 |
| AT5G25754 | AT5G25754.1 \| Symbols: \| RNA polymerase I-associated factor PAF67 \| chr5:8953564-8955511 FORWARD LENGTH=514;AT5G25757.1 \| Symbols: \| RNA polymerase I-associated factor PAF67 \| chr5:8965515-8967460 REVERSE LENGTH=514 | | 1.099614035 | 1.099614035 | 0.90941 |
| AT4G33360 | AT4G33360.1 \| Symbols: FLDH \| NAD(P)-binding Rossmann-fold superfamily protein \| chr4:16067989-16069374 REVERSE LENGTH=344;AT4G33360.3 \| Symbols: FLDH \| NAD(P)-binding Rossmann-fold superfamily protein \| chr4:16067917-16069338 REVERSE LENGTH=327;AT4G33360.2 \| Symbols: FLDH \| NAD(P)-binding Rossmann-fold superfamily protein \| chr4:16068123-16069374 REVERSE LENGTH=305 | | 1.089633229 | 1.089633229 | 0.91774 |

| | | | | | |
|---|---|---|---|---|---|
| AT3G11270 | AT3G11270.1 \| Symbols: MEE34 \| Mov34/MPN/PAD-1 family protein \| chr3:3528989-3531317 FORWARD LENGTH=310;AT3G11270.2 \| Symbols: MEE34 \| Mov34/MPN/PAD-1 family protein \| chr3:3528989-3531317 FORWARD LENGTH=307 | 0.20288 | 1.957445143 | 1.080162571 | 0.925786568 |
| AT5G15610 | AT5G15610.1 \| Symbols: \| Proteasome component (PCI) domain protein \| chr5:5079579-5081929 FORWARD LENGTH=442;AT5G15610.2 \| Symbols: \| Proteasome component (PCI) domain protein \| chr5:5079579-5081929 FORWARD LENGTH=413 | | 1.080076901 | 1.080076901 | 0.92586 |
| AT3G23620 | AT3G23620.1 \| Symbols: \| Ribosomal RNA processing Brix domain protein \| chr3:8480147-8481873 FORWARD LENGTH=314 | | 1.077179943 | 1.077179943 | 0.92835 |
| AT4G33070 | AT4G33070.1 \| Symbols: \| Thiamine pyrophosphate dependent pyruvate decarboxylase family protein \| chr4:15952519-15954676 REVERSE LENGTH=607;AT5G01320.1 \| Symbols: \| Thiamine pyrophosphate dependent pyruvate decarboxylase family protein \| chr5:129484-131625 REVERSE LENGTH=603;AT5G01330.1 \| Symbols: PDC3 \| pyruvate decarboxylase-3 \| chr5:132538-134807 REVERSE LENGTH=592 | 0.48876 | 1.654095541 | 1.07142777 | 0.933334031 |
| AT2G18960 | AT2G18960.1 \| Symbols: AHA1, PMA, OST2, HA1 \| H(+)-ATPase 1 \| chr2:8221858-8227268 FORWARD LENGTH=949;AT4G30190.2 \| Symbols: HA2 \| H(+)-ATPase 2 \| chr4:14770820-14775920 REVERSE LENGTH=981;AT4G30190.1 \| Symbols: AHA2, PMA2, HA2 \| H(+)-ATPase 2 \| chr4:14770820-14775920 REVERSE LENGTH=948;AT3G42640.1 \| Symbols: AHA8, HA8 \| H(+)-ATPase 8 \| chr3:14724309-14728062 FORWARD LENGTH=948;AT5G62670.1 \| Symbols: AHA11, HA11 \| H(+)-ATPase 11 \| chr5:25159495-25164957 FORWARD LENGTH=956;AT3G47950.1 \| Symbols: AHA4, HA4 \| H(+)-ATPase 4 \| chr3:17693015-17697801 FORWARD LENGTH=960;AT5G57350.1 \| Symbols: AHA3, ATAHA3, HA3 \| H(+)-ATPase 3 \| chr5:23231208-23236381 REVERSE LENGTH=949;AT5G57350.2 \| Symbols: HA3 \| H(+)-ATPase 3 \| chr5:23231208-23236381 REVERSE LENGTH=949;AT1G17260.1 \| Symbols: AHA10 \| autoinhibited H(+)-ATPase isoform 10 \| chr1:5904058-5908898 FORWARD LENGTH=947;AT2G24520.1 \| Symbols: AHA5, HA5 \| H(+)-ATPase 5 \| chr2:10415522-10419730 FORWARD LENGTH=931;AT1G80660.1 \| Symbols: AHA9, HA9 \| H(+)-ATPase 9 \| chr1:30316227-30319948 REVERSE LENGTH=954;AT2G07560.1 \| Symbols: AHA6, HA6 \| H(+)-ATPase 6 \| chr2:3170394-3173952 REVERSE LENGTH=949;AT1G80660.2 \| Symbols: HA9 \| H(+)-ATPase 9 \| chr1:30316227-30319948 REVERSE LENGTH=945;AT3G60330.1 \| Symbols: AHA7, HA7 \| H(+)-ATPase 7 \| chr3:22298763-22303509 FORWARD LENGTH=961;AT3G60330.2 \| Symbols: HA7 \| H(+)-ATPase 7 \| chr3:22298763-22303509 FORWARD LENGTH=961 | 1.6078 | 0.443517985 | 1.025658992 | 0.974982921 |
| AT1G08770 | AT1G08770.1 \| Symbols: PRA1.E \| prenylated RAB acceptor 1.E \| chr1:2808933-2809562 FORWARD LENGTH=209 | 0.82799 | 1.180776951 | 1.004383476 | 0.995635655 |
| AT2G37230 | AT2G37230.1 \| Symbols: \| Tetratricopeptide repeat (TPR)-like superfamily protein \| chr2:15637177-15639450 REVERSE LENGTH=757 | | 1.000130017 | 1.000130017 | 0.99987 |
| AT4G11150 | AT4G11150.1 \| Symbols: TUF, emb2448, TUFF, VHA-E1 \| vacuolar ATP synthase subunit E1 \| chr4:6800091-6801696 FORWARD LENGTH=230;AT3G08560.1 \| Symbols: VHA-E2 \| vacuolar H+-ATPase subunit E isoform 2 \| chr3:2600130-2601907 FORWARD LENGTH=235;AT1G64200.1 \| Symbols: VHA-E3 \| vacuolar H+-ATPase subunit E isoform 3 \| chr1:23828537-23830002 REVERSE LENGTH=237 | 0.32746 | 1.670536743 | 0.998998372 | 1.001002633 |
| AT5G39510 | AT5G39510.1 \| Symbols: VTI11, ATVTI1A, ATVTI11, ZIG, SGR4, VTI1A, ZIG1 \| Vesicle transport v-SNARE family protein \| chr5:15822035-15823591 FORWARD LENGTH=221 | 0.3907 | 1.603155009 | 0.996927505 | 1.003081965 |

| | | | | | |
|---|---|---|---|---|---|
| AT4G09720 | AT4G09720.3 \| Symbols: ATRABG3A, RABG3A \| RAB GTPase homolog G3A \| chr4:6133101-6134959 FORWARD LENGTH=217;AT4G09720.4 \| Symbols: RABG3A \| RAB GTPase homolog G3A \| chr4:6133101-6134959 FORWARD LENGTH=211;AT4G09720.1 \| Symbols: ATRABG3A, RABG3A \| RAB GTPase homolog G3A \| chr4:6133101-6134959 FORWARD LENGTH=206;AT4G09720.2 \| Symbols: ATRABG3A, RABG3A \| RAB GTPase homolog G3A \| chr4:6133405-6134959 FORWARD LENGTH=172;AT2G21880.1 \| Symbols: ATRAB7A, ATRABG2, RAB7A \| RAB GTPase homolog 7A \| chr2:9324899-9326170 REVERSE LENGTH=212;AT2G21880.2 \| Symbols: ATRAB7A, ATRABG2, RAB7A \| RAB GTPase homolog 7A \| chr2:9324899-9326170 REVERSE LENGTH=204 | | 0.989315394 | 0.989315394 | 1.0108 |
| AT5G15980 | AT5G15980.1 \| Symbols: \| Pentatricopeptide repeat (PPR) superfamily protein \| chr5:5213290-5215296 FORWARD LENGTH=668 | | 0.988533017 | 0.988533017 | 1.0116 |
| AT5G13710 | AT5G13710.1 \| Symbols: SMT1, CPH \| sterol methyltransferase 1 \| chr5:4424048-4426866 REVERSE LENGTH=336;AT5G13710.2 \| Symbols: SMT1, CPH \| sterol methyltransferase 1 \| chr5:4424048-4426866 REVERSE LENGTH=336 | | 0.980969198 | 0.980969198 | 1.0194 |
| AT2G26200 | AT2G26200.1 \| Symbols: \| S-adenosyl-L-methionine-dependent methyltransferases superfamily protein \| chr2:11152875-11156330 FORWARD LENGTH=565 | 0.98055 | | 0.98055 | 1.019835806 |
| AT5G12470 | AT5G12470.1 \| Symbols: \| Protein of unknown function (DUF3411) \| chr5:4044950-4047290 REVERSE LENGTH=386 | | 0.947777462 | 0.947777462 | 1.0551 |
| AT4G35650 | AT4G35650.1 \| Symbols: IDH-III \| isocitrate dehydrogenase III \| chr4:16908584-16909973 FORWARD LENGTH=368;AT2G17130.1 \| Symbols: IDH2, IDH-II \| isocitrate dehydrogenase subunit 2 \| chr2:7461062-7462466 REVERSE LENGTH=367;AT4G35260.1 \| Symbols: IDH1, IDH-I \| isocitrate dehydrogenase 1 \| chr4:16774494-16776233 REVERSE LENGTH=367;AT2G17130.2 \| Symbols: IDH2, IDH-II \| isocitrate dehydrogenase subunit 2 \| chr2:7461062-7462466 REVERSE LENGTH=363 | | 0.932748811 | 0.932748811 | 1.0721 |
| AT1G48570 | AT1G48570.1 \| Symbols: \| zinc finger (Ran-binding) family protein \| chr1:17955455-17957661 REVERSE LENGTH=455 | | 0.930232558 | 0.930232558 | 1.075 |
| AT5G55660 | AT5G55660.1 \| Symbols: \| DEK domain-containing chromatin associated protein \| chr5:22539375-22543142 FORWARD LENGTH=778 | | 0.913158616 | 0.913158616 | 1.0951 |
| AT4G26900 | AT4G26900.1 \| Symbols: AT-HF, HISN4 \| HIS HF \| chr4:13515514-13519608 FORWARD LENGTH=592 | | 0.912741877 | 0.912741877 | 1.0956 |
| AT3G05380 | AT3G05380.2 \| Symbols: ALY2, ATALY2 \| DIRP ;Myb-like DNA-binding domain \| chr3:1540562-1546139 FORWARD LENGTH=1052;AT3G05380.4 \| Symbols: ALY2 \| DIRP ;Myb-like DNA-binding domain \| chr3:1540562-1546139 FORWARD LENGTH=1052;AT3G05380.5 \| Symbols: ALY2 \| DIRP ;Myb-like DNA-binding domain \| chr3:1540562-1546139 FORWARD LENGTH=1052;AT3G05380.1 \| Symbols: ALY2, ATALY2 \| DIRP ;Myb-like DNA-binding domain \| chr3:1540562-1546139 FORWARD LENGTH=1051;AT3G05380.3 \| Symbols: ALY2, ATALY2 \| DIRP ;Myb-like DNA-binding domain \| chr3:1540961-1546139 FORWARD LENGTH=974 | | 0.905961225 | 0.905961225 | 1.1038 |
| AT3G56090 | AT3G56090.1 \| Symbols: ATFER3, FER3 \| ferritin 3 \| chr3:20814350-20815984 REVERSE LENGTH=259 | 0.51181 | 1.286521118 | 0.899165559 | 1.112142241 |
| AT5G10730 | AT5G10730.1 \| Symbols: \| NAD(P)-binding Rossmann-fold superfamily protein \| chr5:3390822-3392947 REVERSE LENGTH=287 | | 0.892936869 | 0.892936869 | 1.1199 |
| AT1G25260 | AT1G25260.1 \| Symbols: \| Ribosomal protein L10 family protein \| chr1:8854163-8855766 REVERSE LENGTH=235 | | 0.885269122 | 0.885269122 | 1.1296 |

| | | | | | |
|---|---|---|---|---|---|
| AT5G40450 | AT5G40450.1 \| Symbols: \| unknown protein; FUNCTIONS IN: molecular_function unknown; INVOLVED IN: biological_process unknown; LOCATED IN: chloroplast, plasma membrane; EXPRESSED IN: 24 plant structures; EXPRESSED DURING: 13 growth stages; Has 30201 Blast hits to 17322 proteins in 780 species: Archae - 12; Bacteria - 1396; Metazoa - 17338; Fungi - 3422; Plants - 5037; Viruses - 0; Other Eukaryotes - 2996 (source: NCBI BLink). \| chr5:16184799-16195513 REVERSE LENGTH=2889 | | 0.881989769 | 0.881989769 | 1.1338 |
| AT1G75990 | AT1G75990.1 \| Symbols: \| PAM domain (PCI/PINT associated module) protein \| chr1:28524623-28526718 REVERSE LENGTH=487 | | 0.879198171 | 0.879198171 | 1.1374 |
| AT2G17790 | AT2G17790.1 \| Symbols: VPS35A, ZIP3 \| VPS35 homolog A \| chr2:7733685-7739344 FORWARD LENGTH=787 | | 0.877654906 | 0.877654906 | 1.1394 |
| AT4G20440 | AT4G20440.1 \| Symbols: smB \| small nuclear ribonucleoprotein associated protein B \| chr4:11022843-11023616 REVERSE LENGTH=257;AT4G20440.2 \| Symbols: smB \| small nuclear ribonucleoprotein associated protein B \| chr4:11022843-11023616 REVERSE LENGTH=257;AT4G20440.3 \| Symbols: smB \| small nuclear ribonucleoprotein associated protein B \| chr4:11022843-11023616 REVERSE LENGTH=257;AT4G20440.4 \| Symbols: \| small nuclear ribonucleoprotein associated protein B \| chr4:11022843-11023616 REVERSE LENGTH=257;AT5G44500.1 \| Symbols: \| Small nuclear ribonucleoprotein family protein \| chr5:17927505-17928269 FORWARD LENGTH=254;AT5G44500.2 \| Symbols: \| Small nuclear ribonucleoprotein family protein \| chr5:17927505-17928269 FORWARD LENGTH=254 | | 0.873820343 | 0.873820343 | 1.1444 |
| AT1G14810 | AT1G14810.1 \| Symbols: \| semialdehyde dehydrogenase family protein \| chr1:5102684-5104633 REVERSE LENGTH=375 | | 0.861252261 | 0.861252261 | 1.1611 |
| AT4G26910 | AT4G26910.1 \| Symbols: \| Dihydrolipoamide succinyltransferase \| chr4:13520127-13522889 REVERSE LENGTH=464;AT5G55070.1 \| Symbols: \| Dihydrolipoamide succinyltransferase \| chr5:22347637-22350409 FORWARD LENGTH=464;AT4G26910.2 \| Symbols: \| Dihydrolipoamide succinyltransferase \| chr4:13520127-13522889 REVERSE LENGTH=463;AT4G26910.3 \| Symbols: \| Dihydrolipoamide succinyltransferase \| chr4:13520127-13522055 REVERSE LENGTH=365 | | 0.860363073 | 0.860363073 | 1.1623 |
| AT1G10840 | AT1G10840.1 \| Symbols: TIF3H1 \| translation initiation factor 3 subunit H1 \| chr1:3607885-3610299 REVERSE LENGTH=337 | | 0.852224305 | 0.852224305 | 1.1734 |
| AT1G60850 | AT1G60850.1 \| Symbols: ATRPAC42, AAC42 \| DNA-directed RNA polymerase family protein \| chr1:22398078-22400155 REVERSE LENGTH=375;AT1G60850.2 \| Symbols: ATRPAC42, AAC42 \| DNA-directed RNA polymerase family protein \| chr1:22398078-22400155 REVERSE LENGTH=375;AT1G60850.3 \| Symbols: ATRPAC42, AAC42 \| DNA-directed RNA polymerase family protein \| chr1:22398588-22400155 REVERSE LENGTH=302 | | 0.833402784 | 0.833402784 | 1.1999 |
| ATCG00650 | ATCG00650.1 \| Symbols: RPS18 \| ribosomal protein S18 \| chrC:67917-68222 FORWARD LENGTH=101 | 0.57194 | 1.091333719 | 0.831636859 | 1.202447906 |
| AT5G60670 | AT5G60670.1 \| Symbols: \| Ribosomal protein L11 family protein \| chr5:24381066-24381566 REVERSE LENGTH=166 | | 0.827198279 | 0.827198279 | 1.2089 |
| AT1G12310 | AT1G12310.1 \| Symbols: \| Calcium-binding EF-hand family protein \| chr1:4187500-4187946 REVERSE LENGTH=148 | 1.1847 | 0.454277018 | 0.819488509 | 1.220273364 |
| AT2G38360 | AT2G38360.1 \| Symbols: PRA1.B4 \| prenylated RAB acceptor 1.B4 \| chr2:16069840-16070502 REVERSE LENGTH=220 | 0.069219 | 1.564480045 | 0.816849523 | 1.224215688 |
| AT2G18040 | AT2G18040.1 \| Symbols: PIN1AT \| peptidylprolyl cis/trans isomerase, NIMA-interacting 1 \| chr2:7842346-7843537 FORWARD LENGTH=119 | | 0.815926893 | 0.815926893 | 1.2256 |
| AT3G13570 | AT3G13570.1 \| Symbols: SCL30A, At-SCL30A \| SC35-like splicing factor 30A \| chr3:4429564-4431602 REVERSE LENGTH=262;AT1G55310.1 \| Symbols: SR33, ATSCL33, SCL33, At-SCL33 \| SC35-like splicing factor 33 \| chr1:20630676-20632695 FORWARD LENGTH=287;AT1G55310.2 \| Symbols: SR33, ATSCL33, SCL33, At-SCL33 \| SC35-like splicing factor 33 \| chr1:20630676-20632567 FORWARD LENGTH=220 | | 0.813868316 | 0.813868316 | 1.2287 |

| | | | | | |
|---|---|---|---|---|---|
| AT3G05810 | AT3G05810.1 \| Symbols: \| FUNCTIONS IN: molecular_function unknown; INVOLVED IN: biological_process unknown; LOCATED IN: mitochondrion; EXPRESSED IN: 22 plant structures; EXPRESSED DURING: 13 growth stages; CONTAINS InterPro DOMAIN/s: IGR protein motif (InterPro:IPR019083); BEST Arabidopsis thaliana protein match is: unknown protein (TAIR:AT5G26800.1); Has 62 Blast hits to 62 proteins in 14 species: Archae - 0; Bacteria - 0; Metazoa - 0; Fungi - 0; Plants - 62; Viruses - 0; Other Eukaryotes - 0 (source: NCBI BLink). \| chr3:1731259-1732321 REVERSE LENGTH=112 | | 0.812545706 | 0.812545706 | 1.2307 |
| AT5G60790 | AT5G60790.1 \| Symbols: ATGCN1, GCN1 \| ABC transporter family protein \| chr5:24453760-24455767 REVERSE LENGTH=595 | | 0.805023346 | 0.805023346 | 1.2422 |
| AT5G60960 | AT5G60960.1 \| Symbols: \| Pentatricopeptide repeat (PPR) superfamily protein \| chr5:24528423-24529988 REVERSE LENGTH=521 | | 0.80070462 | 0.80070462 | 1.2489 |
| AT4G35785 | AT4G35785.5 \| Symbols: \| RNA-binding (RRM/RBD/RNP motifs) family protein \| chr4:16953404-16955127 REVERSE LENGTH=244;AT4G35785.2 \| Symbols: \| RNA-binding (RRM/RBD/RNP motifs) family protein \| chr4:16953211-16955127 REVERSE LENGTH=239;AT4G35785.1 \| Symbols: \| RNA-binding (RRM/RBD/RNP motifs) family protein \| chr4:16953211-16955127 REVERSE LENGTH=238;AT4G35785.3 \| Symbols: \| RNA-binding (RRM/RBD/RNP motifs) family protein \| chr4:16953708-16955127 REVERSE LENGTH=207;AT4G35785.4 \| Symbols: \| RNA-binding (RRM/RBD/RNP motifs) family protein \| chr4:16953404-16955127 REVERSE LENGTH=201 | | 0.79649542 | 0.79649542 | 1.2555 |
| AT2G37620 | AT2G37620.1 \| Symbols: ACT1, AAc1 \| actin 1 \| chr2:15779761-15781241 FORWARD LENGTH=377;AT2G37620.2 \| Symbols: ACT1, AAc1 \| actin 1 \| chr2:15779761-15781241 FORWARD LENGTH=377;AT3G53750.1 \| Symbols: ACT3 \| actin 3 \| chr3:19915924-19917371 FORWARD LENGTH=377 | 0.3207 | 1.269792897 | 0.795246448 | 1.257471822 |
| AT3G57290 | AT3G57290.1 \| Symbols: EIF3E, TIF3E1, ATEIF3E-1, INT-6, ATINT6, INT6 \| eukaryotic translation initiation factor 3E \| chr3:21196786-21199073 REVERSE LENGTH=441 | | 0.789639924 | 0.789639924 | 1.2664 |
| AT5G14670 | AT5G14670.1 \| Symbols: ATARFA1B, ARFA1B \| ADP-ribosylation factor A1B \| chr5:4729319-4730495 FORWARD LENGTH=188;AT1G10630.1 \| Symbols: ATARFA1F, ARFA1F \| ADP-ribosylation factor A1F \| chr1:3513189-3514230 REVERSE LENGTH=181;AT1G23490.1 \| Symbols: ATARFA1A, ATARF1, ATARF, ARF1 \| ADP-ribosylation factor 1 \| chr1:8337232-8338373 FORWARD LENGTH=181;AT1G70490.1 \| Symbols: ATARFA1D, ARFA1D \| Ras-related small GTP-binding family protein \| chr1:26564162-26565152 REVERSE LENGTH=181;AT1G70490.2 \| Symbols: ATARFA1D, ARFA1D \| Ras-related small GTP-binding family protein \| chr1:26564162-26565152 REVERSE LENGTH=181;AT1G70490.3 \| Symbols: ATARFA1D, ARFA1D \| Ras-related small GTP-binding family protein \| chr1:26564162-26565152 REVERSE LENGTH=181;AT2G47170.1 \| Symbols: ARF1A1C \| Ras-related small GTP-binding family protein \| chr2:19367264-19368518 FORWARD LENGTH=181;AT3G62290.1 \| Symbols: ATARFA1E, ARFA1E \| ADP-ribosylation factor A1E \| chr3:23052287-23053545 FORWARD LENGTH=181;AT3G62290.2 \| Symbols: ARFA1E \| ADP-ribosylation factor A1E \| chr3:23052287-23053545 FORWARD LENGTH=181;AT3G62290.3 \| Symbols: ARFA1E \| ADP-ribosylation factor A1E \| chr3:23052287-23053545 FORWARD LENGTH=181 | 1.085 | 0.440955993 | 0.762977996 | 1.310653787 |
| AT2G06990 | AT2G06990.1 \| Symbols: HEN2 \| RNA helicase, ATP-dependent, SK12/DOB1 protein \| chr2:2895135-2900909 FORWARD LENGTH=995 | 0.75942 | | 0.75942 | 1.316794396 |
| AT3G03960 | AT3G03960.1 \| Symbols: \| TCP-1/cpn60 chaperonin family protein \| chr3:1024432-1027604 FORWARD LENGTH=549 | | 0.758955677 | 0.758955677 | 1.3176 |
| AT5G54960 | AT5G54960.1 \| Symbols: PDC2 \| pyruvate decarboxylase-2 \| chr5:22310858-22312681 REVERSE LENGTH=607 | 0.10052 | 1.381730756 | 0.741125378 | 1.349299362 |
| AT1G27190 | AT1G27190.1 \| Symbols: \| Leucine-rich repeat protein kinase family protein \| chr1:9446923-9448728 REVERSE LENGTH=601 | | 0.740740741 | 0.740740741 | 1.35 |

| | | | | | |
|---|---|---|---|---|---|
| AT1G32400 | AT1G32400.1 \| Symbols: TOM2A \| tobamovirus multiplication 2A \| chr1:11689393-11690873 REVERSE LENGTH=280;AT1G32400.2 \| Symbols: TOM2A \| tobamovirus multiplication 2A \| chr1:11689393-11690873 REVERSE LENGTH=280;AT1G32400.3 \| Symbols: TOM2A \| tobamovirus multiplication 2A \| chr1:11689393-11690873 REVERSE LENGTH=280 | 0.4753 | 0.971345313 | 0.723322657 | 1.382508886 |
| AT5G40870 | AT5G40870.1 \| Symbols: ATUK/UPRT1, UKL1, UK/UPRT1 \| uridine kinase/uracil phosphoribosyltransferase 1 \| chr5:16375021-16378384 FORWARD LENGTH=486;AT3G27190.1 \| Symbols: UKL2 \| uridine kinase-like 2 \| chr3:10039504-10042917 REVERSE LENGTH=483;AT3G27440.1 \| Symbols: UKL5 \| uridine kinase-like 5 \| chr3:10155555-10157931 FORWARD LENGTH=465 | | 0.711490573 | 0.711490573 | 1.4055 |
| AT2G16440 | AT2G16440.1 \| Symbols: MCM4 \| Minichromosome maintenance (MCM2/3/5) family protein \| chr2:7126536-7130665 REVERSE LENGTH=847 | | 0.709572128 | 0.709572128 | 1.4093 |
| AT2G35390 | AT2G35390.2 \| Symbols: \| Phosphoribosyltransferase family protein \| chr2:14895528-14897581 REVERSE LENGTH=403;AT2G35390.1 \| Symbols: \| Phosphoribosyltransferase family protein \| chr2:14895528-14897098 REVERSE LENGTH=352;AT2G35390.3 \| Symbols: \| Phosphoribosyltransferase family protein \| chr2:14895808-14897581 REVERSE LENGTH=344 | | 0.709521782 | 0.709521782 | 1.4094 |
| AT4G29410 | AT4G29410.1 \| Symbols: \| Ribosomal L28e protein family \| chr4:14468439-14469964 REVERSE LENGTH=143;AT4G29410.2 \| Symbols: \| Ribosomal L28e protein family \| chr4:14468439-14469964 REVERSE LENGTH=143 | | 0.70856657 | 0.70856657 | 1.4113 |
| AT3G25520 | AT3G25520.1 \| Symbols: ATL5, PGY3, OLI5, RPL5A \| ribosomal protein L5 \| chr3:9269573-9271327 REVERSE LENGTH=301;AT5G39740.1 \| Symbols: OLI7, RPL5B \| ribosomal protein L5 B \| chr5:15903365-15905185 FORWARD LENGTH=301;AT5G39740.2 \| Symbols: RPL5B \| ribosomal protein L5 B \| chr5:15903484-15905185 FORWARD LENGTH=301;AT3G25520.2 \| Symbols: ATL5 \| ribosomal protein L5 \| chr3:9269573-9270434 REVERSE LENGTH=190 | 0.22531 | 1.188806201 | 0.7070581 | 1.414310931 |
| AT4G39520 | AT4G39520.1 \| Symbols: \| GTP-binding protein-related \| chr4:18371329-18374000 REVERSE LENGTH=369 | | 0.704076604 | 0.704076604 | 1.4203 |
| AT5G59840 | AT5G59840.1 \| Symbols: \| Ras-related small GTP-binding family protein \| chr5:24107450-24109049 REVERSE LENGTH=216;AT3G53610.1 \| Symbols: ATRAB8, AtRab8B, AtRABE1a, RAB8 \| RAB GTPase homolog 8 \| chr3:19876531-19878264 REVERSE LENGTH=216;AT3G53610.2 \| Symbols: ATRAB8, RAB8 \| RAB GTPase homolog 8 \| chr3:19876531-19878264 REVERSE LENGTH=216;AT3G53610.3 \| Symbols: ATRAB8, RAB8 \| RAB GTPase homolog 8 \| chr3:19876531-19878264 REVERSE LENGTH=216 | | 0.694444444 | 0.694444444 | 1.44 |
| AT1G73650 | AT1G73650.3 \| Symbols: \| Protein of unknown function (DUF1295) \| chr1:27688409-27690165 REVERSE LENGTH=302;AT1G73650.2 \| Symbols: \| Protein of unknown function (DUF1295) \| chr1:27688549-27690165 REVERSE LENGTH=291;AT1G73650.4 \| Symbols: \| Protein of unknown function (DUF1295) \| chr1:27688420-27689606 REVERSE LENGTH=219;AT1G73650.1 \| Symbols: \| Protein of unknown function (DUF1295) \| chr1:27688449-27689606 REVERSE LENGTH=208 | | 0.691993634 | 0.691993634 | 1.4451 |
| AT2G21660 | AT2G21660.1 \| Symbols: ATGRP7, CCR2, GR-RBP7, GRP7 \| cold, circadian rhythm, and rna binding 2 \| chr2:9265477-9266316 REVERSE LENGTH=176;AT2G21660.2 \| Symbols: ATGRP7, CCR2 \| cold, circadian rhythm, and rna binding 2 \| chr2:9265477-9266316 REVERSE LENGTH=159 | | 0.687757909 | 0.687757909 | 1.454 |
| AT1G29940 | AT1G29940.1 \| Symbols: NRPA2 \| nuclear RNA polymerase A2 \| chr1:10479322-10486670 REVERSE LENGTH=1178 | | 0.678012069 | 0.678012069 | 1.4749 |

| | | | | | |
|---|---|---|---|---|---|
| AT5G17920 | AT5G17920.1 \| Symbols: ATCIMS, ATMETS, ATMS1 \| Cobalamin-independent synthase family protein \| chr5:5935771-5939195 FORWARD LENGTH=765;AT5G17920.2 \| Symbols: ATCIMS \| Cobalamin-independent synthase family protein \| chr5:5935771-5939195 FORWARD LENGTH=765;AT3G03780.1 \| Symbols: ATMS2, MS2 \| methionine synthase 2 \| chr3:957602-960740 FORWARD LENGTH=765;AT3G03780.2 \| Symbols: ATMS2, MS2 \| methionine synthase 2 \| chr3:957602-960740 FORWARD LENGTH=765;AT3G03780.3 \| Symbols: ATMS2, MS2 \| methionine synthase 2 \| chr3:957602-960740 FORWARD LENGTH=765;AT5G20980.1 \| Symbols: ATMS3, MS3 \| methionine synthase 3 \| chr5:7124397-7128353 REVERSE LENGTH=812;AT5G20980.2 \| Symbols: ATMS3, MS3 \| methionine synthase 3 \| chr5:7124397-7128353 REVERSE LENGTH=812 | 0.44666 | 0.901306895 | 0.673983447 | 1.483715963 |
| AT2G18510 | AT2G18510.1 \| Symbols: emb2444 \| RNA-binding (RRM/RBD/RNP motifs) family protein \| chr2:8031554-8033517 REVERSE LENGTH=363 | | 0.66436354 | 0.66436354 | 1.5052 |
| AT1G44910 | AT1G44910.1 \| Symbols: ATPRP40A, PRP40A \| pre-mRNA-processing protein 40A \| chr1:16975930-16982818 FORWARD LENGTH=958;AT1G44910.2 \| Symbols: ATPRP40A, PRP40A \| pre-mRNA-processing protein 40A \| chr1:16975930-16982647 FORWARD LENGTH=926 | | 0.661769572 | 0.661769572 | 1.5111 |
| AT4G14960 | AT4G14960.2 \| Symbols: TUA6 \| Tubulin/FtsZ family protein \| chr4:8548769-8550319 REVERSE LENGTH=450;AT4G14960.1 \| Symbols: TUA6 \| Tubulin/FtsZ family protein \| chr4:8548753-8550319 REVERSE LENGTH=427 | | 0.661769572 | 0.661769572 | 1.5111 |
| AT4G15900 | AT4G15900.1 \| Symbols: PRL1 \| pleiotropic regulatory locus 1 \| chr4:9023775-9027443 FORWARD LENGTH=486 | | 0.653637493 | 0.653637493 | 1.5299 |
| AT1G29150 | AT1G29150.1 \| Symbols: ATS9, RPN6 \| non-ATPase subunit 9 \| chr1:10181240-10182499 FORWARD LENGTH=419 | 0.18581 | 1.120059139 | 0.65293457 | 1.531547029 |
| AT2G47610 | AT2G47610.1 \| Symbols: \| Ribosomal protein L7Ae/L30e/S12e/Gadd45 family protein \| chr2:19529854-19531401 FORWARD LENGTH=257 | | 0.648802959 | 0.648802959 | 1.5413 |
| AT3G21280 | AT3G21280.1 \| Symbols: UBP7 \| ubiquitin-specific protease 7 \| chr3:7478166-7482037 REVERSE LENGTH=532;AT1G51710.1 \| Symbols: UBP6, ATUBP6 \| ubiquitin-specific protease 6 \| chr1:19175805-19179894 REVERSE LENGTH=482;AT1G51710.2 \| Symbols: UBP6 \| ubiquitin-specific protease 6 \| chr1:19175805-19179399 REVERSE LENGTH=443 | | 0.646495992 | 0.646495992 | 1.5468 |
| AT5G37640 | AT5G37640.1 \| Symbols: UBQ9 \| ubiquitin 9 \| chr5:14952782-14953750 REVERSE LENGTH=322 | | 0.646370629 | 0.646370629 | 1.5471 |
| AT1G48410 | AT1G48410.2 \| Symbols: AGO1 \| Stabilizer of iron transporter SufD / Polynucleotidyl transferase \| chr1:17886285-17891892 REVERSE LENGTH=1050;AT1G48410.3 \| Symbols: AGO1 \| Stabilizer of iron transporter SufD / Polynucleotidyl transferase \| chr1:17886285-17891892 REVERSE LENGTH=1050;AT1G48410.1 \| Symbols: AGO1 \| Stabilizer of iron transporter SufD / Polynucleotidyl transferase \| chr1:17886285-17891892 REVERSE LENGTH=1048;AT2G27880.1 \| Symbols: AGO5 \| Argonaute family protein \| chr2:11871488-11876712 FORWARD LENGTH=997;AT5G43810.1 \| Symbols: ZLL, PNH, AGO10 \| Stabilizer of iron transporter SufD / Polynucleotidyl transferase \| chr5:17611939-17616562 FORWARD LENGTH=988;AT5G43810.2 \| Symbols: ZLL \| Stabilizer of iron transporter SufD / Polynucleotidyl transferase \| chr5:17611939-17616562 FORWARD LENGTH=988 | | 0.635081926 | 0.635081926 | 1.5746 |
| AT1G05910 | AT1G05910.1 \| Symbols: \| cell division cycle protein 48-related / CDC48-related \| chr1:1790796-1796503 FORWARD LENGTH=1210 | | 0.630119723 | 0.630119723 | 1.587 |
| AT3G63460 | AT3G63460.1 \| Symbols: \| transducin family protein / WD-40 repeat family protein \| chr3:23431009-23437241 REVERSE LENGTH=1104;AT3G63460.2 \| Symbols: \| transducin family protein / WD-40 repeat family protein \| chr3:23431009-23437241 REVERSE LENGTH=1102;AT3G63460.3 \| Symbols: \| transducin family protein / WD-40 repeat family protein \| chr3:23431009-23437241 REVERSE LENGTH=1094 | | 0.628298567 | 0.628298567 | 1.5916 |

| | | | | | |
|---|---|---|---|---|---|
| AT3G53110 | AT3G53110.1 \| Symbols: LOS4 \| P-loop containing nucleoside triphosphate hydrolases superfamily protein \| chr3:19687968-19690423 FORWARD LENGTH=496 | | 0.624336642 | 0.624336642 | 1.6017 |
| AT1G65260 | AT1G65260.1 \| Symbols: PTAC4, VIPP1 \| plastid transcriptionally active 4 \| chr1:24236329-24240428 FORWARD LENGTH=330 | 0.27973 | 0.957487553 | 0.618608776 | 1.616530574 |
| AT4G11600 | AT4G11600.1 \| Symbols: ATGPX6, PHGPX, LSC803, GPX6 \| glutathione peroxidase 6 \| chr4:7010021-7011330 REVERSE LENGTH=232 | | 0.61716966 | 0.61716966 | 1.6203 |
| AT2G20190 | AT2G20190.1 \| Symbols: ATCLASP, CLASP \| CLIP-associated protein \| chr2:8711862-8718813 REVERSE LENGTH=1439 | 0.1174 | 1.115088259 | 0.61624413 | 1.622733511 |
| AT1G45000 | AT1G45000.1 \| Symbols: \| AAA-type ATPase family protein \| chr1:17009220-17011607 FORWARD LENGTH=399;AT5G43010.1 \| Symbols: RPT4A \| regulatory particle triple-A ATPase 4A \| chr5:17248563-17251014 REVERSE LENGTH=399;AT1G45000.2 \| Symbols: \| AAA-type ATPase family protein \| chr1:17009220-17011607 FORWARD LENGTH=335 | 0.21197 | 1.016859531 | 0.614414766 | 1.627565052 |
| AT2G37600 | AT2G37600.1 \| Symbols: \| Ribosomal protein L36e family protein \| chr2:15774410-15775203 REVERSE LENGTH=113;AT2G37600.2 \| Symbols: \| Ribosomal protein L36e family protein \| chr2:15774410-15775203 REVERSE LENGTH=113;AT3G53740.2 \| Symbols: \| Ribosomal protein L36e family protein \| chr3:19913921-19914813 REVERSE LENGTH=112;AT3G53740.3 \| Symbols: \| Ribosomal protein L36e family protein \| chr3:19913921-19914813 REVERSE LENGTH=112;AT3G53740.4 \| Symbols: \| Ribosomal protein L36e family protein \| chr3:19913921-19914813 REVERSE LENGTH=112;AT3G53740.1 \| Symbols: \| Ribosomal protein L36e family protein \| chr3:19913921-19914813 REVERSE LENGTH=103 | | 0.607754953 | 0.607754953 | 1.6454 |
| AT1G19520 | AT1G19520.1 \| Symbols: NFD5 \| pentatricopeptide (PPR) repeat-containing protein \| chr1:6760032-6762581 FORWARD LENGTH=725 | | 0.606612072 | 0.606612072 | 1.6485 |
| AT1G76810 | AT1G76810.1 \| Symbols: \| eukaryotic translation initiation factor 2 (eIF-2) family protein \| chr1:28831366-28836310 REVERSE LENGTH=1294;AT1G76820.1 \| Symbols: \| eukaryotic translation initiation factor 2 (eIF-2) family protein \| chr1:28837796-28842343 FORWARD LENGTH=1166 | | 0.606060606 | 0.606060606 | 1.65 |
| AT5G02610 | AT5G02610.2 \| Symbols: \| Ribosomal L29 family protein \| chr5:587611-588547 FORWARD LENGTH=146;AT2G39390.1 \| Symbols: \| Ribosomal L29 family protein \| chr2:16450803-16451762 REVERSE LENGTH=123;AT5G02610.1 \| Symbols: \| Ribosomal L29 family protein \| chr5:587611-588547 FORWARD LENGTH=123 | 0.60332 | | 0.60332 | 1.657495193 |
| AT3G57280 | AT3G57280.1 \| Symbols: \| Transmembrane proteins 14C \| chr3:21193960-21195547 FORWARD LENGTH=226 | | 0.594494977 | 0.594494977 | 1.6821 |
| AT2G04795 | AT2G04795.1 \| Symbols: \| unknown protein; BEST Arabidopsis thaliana protein match is: unknown protein (TAIR:AT5G35732.1); Has 18 Blast hits to 18 proteins in 6 species: Archae - 0; Bacteria - 0; Metazoa - 0; Fungi - 0; Plants - 18; Viruses - 0; Other Eukaryotes - 0 (source: NCBI BLink). \| chr2:1683027-1683314 FORWARD LENGTH=95 | | 0.591715976 | 0.591715976 | 1.69 |
| AT3G08590 | AT3G08590.1 \| Symbols: \| Phosphoglycerate mutase, 2,3-bisphosphoglycerate-independent \| chr3:2608683-2611237 REVERSE LENGTH=560;AT3G08590.2 \| Symbols: \| Phosphoglycerate mutase, 2,3-bisphosphoglycerate-independent \| chr3:2608683-2611237 REVERSE LENGTH=560 | | 0.583498658 | 0.583498658 | 1.7138 |
| AT3G46060 | AT3G46060.1 \| Symbols: ARA3, ARA-3, ATRABE1C, ATRAB8A, RAB8A \| RAB GTPase homolog 8A \| chr3:16917908-16919740 FORWARD LENGTH=216;AT3G46060.2 \| Symbols: ARA3 \| RAB GTPase homolog 8A \| chr3:16917908-16919740 FORWARD LENGTH=216;AT3G46060.3 \| Symbols: ARA3 \| RAB GTPase homolog 8A \| chr3:16917908-16919740 FORWARD LENGTH=216 | 0.57768 | | 0.57768 | 1.73106218 |

| | | | | | |
|---|---|---|---|---|---|
| AT4G15640 | AT4G15640.1 \| Symbols: \| unknown protein; BEST Arabidopsis thaliana protein match is: unknown protein (TAIR:AT3G21465.1); Has 38 Blast hits to 38 proteins in 14 species: Archae - 0; Bacteria - 0; Metazoa - 0; Fungi - 0; Plants - 38; Viruses - 0; Other Eukaryotes - 0 (source: NCBI BLink). \| chr4:8919245-8921852 REVERSE LENGTH=390 | | 0.572409845 | 0.572409845 | 1.747 |
| AT3G55430 | AT3G55430.1 \| Symbols: \| O-Glycosyl hydrolases family 17 protein \| chr3:20549806-20552004 REVERSE LENGTH=449 | | 0.570092925 | 0.570092925 | 1.7541 |
| AT4G25740 | AT4G25740.1 \| Symbols: \| RNA binding Plectin/S10 domain-containing protein \| chr4:13107488-13108751 REVERSE LENGTH=177;AT4G25740.2 \| Symbols: \| RNA binding Plectin/S10 domain-containing protein \| chr4:13107488-13108751 REVERSE LENGTH=149 | | 0.567504682 | 0.567504682 | 1.7621 |
| ATCG00900 | ATCG00900.1 \| Symbols: RPS7.1, RPS7 \| Ribosomal protein S7p/S5e family protein \| chrC:97478-97945 REVERSE LENGTH=155;ATCG01240.1 \| Symbols: RPS7.2 \| ribosomal protein S7 \| chrC:140704-141171 FORWARD LENGTH=155 | | 0.565866908 | 0.565866908 | 1.7672 |
| ATCG00500 | ATCG00500.1 \| Symbols: ACCD \| acetyl-CoA carboxylase carboxyl transferase subunit beta \| chrC:57075-58541 FORWARD LENGTH=488 | | 0.564588979 | 0.564588979 | 1.7712 |
| AT1G03140 | AT1G03140.1 \| Symbols: \| splicing factor Prp18 family protein \| chr1:754471-756223 REVERSE LENGTH=420 | | 0.563475517 | 0.563475517 | 1.7747 |
| AT4G02620 | AT4G02620.1 \| Symbols: \| vacuolar ATPase subunit F family protein \| chr4:1149419-1151132 REVERSE LENGTH=128 | 0.50753 | 0.61447708 | 0.56100354 | 1.782519946 |
| AT4G17140 | AT4G17140.3 \| Symbols: \| pleckstrin homology (PH) domain-containing protein \| chr4:9613617-9636618 REVERSE LENGTH=4219;AT4G17140.2 \| Symbols: \| pleckstrin homology (PH) domain-containing protein \| chr4:9613617-9636618 REVERSE LENGTH=4218;AT4G17140.1 \| Symbols: \| pleckstrin homology (PH) domain-containing protein \| chr4:9613617-9636618 REVERSE LENGTH=4216 | | 0.557413601 | 0.557413601 | 1.794 |
| AT5G26800 | AT5G26800.1 \| Symbols: \| FUNCTIONS IN: molecular_function unknown; INVOLVED IN: biological_process unknown; LOCATED IN: mitochondrion; CONTAINS InterPro DOMAIN/s: IGR protein motif (InterPro:IPR019083); BEST Arabidopsis thaliana protein match is: unknown protein (TAIR:AT3G05810.1); Has 1807 Blast hits to 1807 proteins in 277 species: Archae - 0; Bacteria - 0; Metazoa - 736; Fungi - 347; Plants - 385; Viruses - 0; Other Eukaryotes - 339 (source: NCBI BLink). \| chr5:9425224-9426485 FORWARD LENGTH=112 | 0.48307 | 0.630477271 | 0.556773636 | 1.796062055 |
| AT5G28740 | AT5G28740.1 \| Symbols: \| Tetratricopeptide repeat (TPR)-like superfamily protein \| chr5:10780774-10783772 FORWARD LENGTH=917 | | 0.555926173 | 0.555926173 | 1.7988 |
| AT2G36850 | AT2G36850.1 \| Symbols: ATGSL08, GSL8, GSL08, ATGSL8, CHOR \| glucan synthase-like 8 \| chr2:15454935-15469666 REVERSE LENGTH=1904 | 0.48155 | 0.629207827 | 0.555378914 | 1.800572502 |
| AT5G09900 | AT5G09900.3 \| Symbols: EMB2107, RPN5A, MSA \| 26S proteasome regulatory subunit, putative (RPN5) \| chr5:3089406-3092434 REVERSE LENGTH=462;AT5G09900.1 \| Symbols: EMB2107, RPN5A, MSA \| 26S proteasome regulatory subunit, putative (RPN5) \| chr5:3089462-3092434 REVERSE LENGTH=442;AT5G09900.2 \| Symbols: EMB2107, RPN5A, MSA \| 26S proteasome regulatory subunit, putative (RPN5) \| chr5:3089462-3092434 REVERSE LENGTH=442;AT5G64760.1 \| Symbols: RPN5B \| regulatory particle non-ATPase subunit 5B \| chr5:25894027-25896445 REVERSE LENGTH=442;AT5G64760.2 \| Symbols: RPN5B \| regulatory particle non-ATPase subunit 5B \| chr5:25894027-25896445 REVERSE LENGTH=442 | 0.12729 | 0.975134081 | 0.55121204 | 1.814183883 |
| AT1G54220 | AT1G54220.1 \| Symbols: \| Dihydrolipoamide acetyltransferase, long form protein \| chr1:20246460-20250208 REVERSE LENGTH=539;AT1G54220.2 \| Symbols: \| Dihydrolipoamide acetyltransferase, long form protein \| chr1:20246460-20250208 REVERSE LENGTH=539 | 0.056215 | 1.003562647 | 0.529888824 | 1.887188322 |
| AT3G01780 | AT3G01780.1 \| Symbols: TPLATE \| ARM repeat superfamily protein \| chr3:279171-283399 FORWARD LENGTH=1176 | | 0.52759312 | 0.52759312 | 1.8954 |
| AT4G23710 | AT4G23710.1 \| Symbols: VHA-G2, VAG2, VATG2 \| vacuolar ATP synthase subunit G2 \| chr4:12350577-12351354 FORWARD LENGTH=106 | 0.36504 | 0.685588921 | 0.52531446 | 1.903621688 |

| | | | | | |
|---|---|---|---|---|---|
| AT1G31290 | AT1G31290.1 \| Symbols: AGO3 \| ARGONAUTE 3 \| chr1:11188293-11192317 FORWARD LENGTH=1194 | | 0.524191435 | 0.524191435 | 1.9077 |
| AT3G49870 | AT3G49870.1 \| Symbols: ATARLA1C, ARLA1C \| ADP-ribosylation factor-like A1C \| chr3:18492674-18494021 REVERSE LENGTH=184 | 0.52045 | | 0.52045 | 1.921414161 |
| AT1G16820 | AT1G16820.1 \| Symbols: \| vacuolar ATP synthase catalytic subunit-related / V-ATPase-related / vacuolar proton pump-related \| chr1:5757199-5758464 REVERSE LENGTH=93;AT1G16820.2 \| Symbols: \| vacuolar ATP synthase catalytic subunit-related / V-ATPase-related / vacuolar proton pump-related \| chr1:5757199-5758202 REVERSE LENGTH=66 | 0.13952 | 0.894134478 | 0.516827239 | 1.934882539 |
| AT4G25530 | AT4G25530.1 \| Symbols: FWA, HDG6 \| FLOWERING WAGENINGEN \| chr4:13039312-13042242 FORWARD LENGTH=686 | | 0.512373828 | 0.512373828 | 1.9517 |
| AT2G01720 | AT2G01720.1 \| Symbols: \| Ribophorin I \| chr2:317193-320016 REVERSE LENGTH=464 | | 0.51064699 | 0.51064699 | 1.9583 |
| AT2G23930 | AT2G23930.1 \| Symbols: SNRNP-G \| probable small nuclear ribonucleoprotein G \| chr2:10182353-10183026 FORWARD LENGTH=80;AT3G11500.1 \| Symbols: \| Small nuclear ribonucleoprotein family protein \| chr3:3621626-3622474 REVERSE LENGTH=79;AT2G23930.2 \| Symbols: SNRNP-G \| probable small nuclear ribonucleoprotein G \| chr2:10182496-10183026 FORWARD LENGTH=67 | 0.80491 | 0.208420175 | 0.506665088 | 1.973690362 |
| AT4G04640 | AT4G04640.1 \| Symbols: ATPC1 \| ATPase, F1 complex, gamma subunit protein \| chr4:2350761-2351882 REVERSE LENGTH=373 | | 0.503423278 | 0.503423278 | 1.9864 |
| AT3G51800 | AT3G51800.2 \| Symbols: ATG2, EBP1, ATEBP1 \| metallopeptidase M24 family protein \| chr3:19211261-19213568 REVERSE LENGTH=401;AT3G51800.1 \| Symbols: ATG2, EBP1, ATEBP1 \| metallopeptidase M24 family protein \| chr3:19211261-19213568 REVERSE LENGTH=392;AT3G51800.3 \| Symbols: ATG2 \| metallopeptidase M24 family protein \| chr3:19211261-19213568 REVERSE LENGTH=385 | | 0.502411576 | 0.502411576 | 1.9904 |
| AT2G21580 | AT2G21580.1 \| Symbols: \| Ribosomal protein S25 family protein \| chr2:9236629-9237510 FORWARD LENGTH=108;AT2G21580.2 \| Symbols: \| Ribosomal protein S25 family protein \| chr2:9236629-9237510 FORWARD LENGTH=107 | | 0.501328521 | 0.501328521 | 1.9947 |
| AT1G50480 | AT1G50480.1 \| Symbols: THFS \| 10-formyltetrahydrofolate synthetase \| chr1:18702064-18704687 FORWARD LENGTH=634 | | 0.499151443 | 0.499151443 | 2.0034 |
| AT2G44610 | AT2G44610.1 \| Symbols: RAB6, ATRABH1B, ATRAB6A, RAB6A \| Ras-related small GTP-binding family protein \| chr2:18411778-18413883 REVERSE LENGTH=208;AT4G39890.1 \| Symbols: AtRABH1c, RABH1c \| RAB GTPase homolog H1C \| chr4:18506112-18507459 FORWARD LENGTH=214;AT2G22290.1 \| Symbols: ATRABH1D, ATRAB-H1D, RAB-H1D, ATRAB6, RABH1d \| RAB GTPase homolog H1D \| chr2:9466568-9467688 FORWARD LENGTH=207;AT5G10260.1 \| Symbols: AtRABH1e, RABH1e \| RAB GTPase homolog H1E \| chr5:3219991-3221301 FORWARD LENGTH=207;AT5G64990.1 \| Symbols: AtRABH1a, RABH1a \| RAB GTPase homolog H1A \| chr5:25963562-25964792 REVERSE LENGTH=206 | | 0.499051802 | 0.499051802 | 2.0038 |
| AT2G06210 | AT2G06210.1 \| Symbols: ELF8, VIP6 \| binding \| chr2:2429108-2436588 REVERSE LENGTH=1091 | | 0.496943796 | 0.496943796 | 2.0123 |
| AT3G59650 | AT3G59650.2 \| Symbols: \| mitochondrial ribosomal protein L51/S25/CI-B8 family protein \| chr3:22033216-22033928 FORWARD LENGTH=146;AT3G59650.1 \| Symbols: \| mitochondrial ribosomal protein L51/S25/CI-B8 family protein \| chr3:22033216-22033928 FORWARD LENGTH=119 | | 0.49689441 | 0.49689441 | 2.0125 |
| AT4G35360 | AT4G35360.1 \| Symbols: \| Uncharacterised conserved protein (UCP030210) \| chr4:16812303-16814656 REVERSE LENGTH=367 | | 0.496130185 | 0.496130185 | 2.0156 |
| AT2G05620 | AT2G05620.1 \| Symbols: PGR5 \| proton gradient regulation 5 \| chr2:2081204-2081687 REVERSE LENGTH=133 | | 0.495613818 | 0.495613818 | 2.0177 |
| AT3G19760 | AT3G19760.1 \| Symbols: EIF4A-III \| eukaryotic initiation factor 4A-III \| chr3:6863790-6866242 FORWARD LENGTH=408 | | 0.492465281 | 0.492465281 | 2.0306 |

| | | | | | |
|---|---|---|---|---|---|
| AT2G37190 | AT2G37190.1 \| Symbols: \| Ribosomal protein L11 family protein \| chr2:15619559-15620059 REVERSE LENGTH=166;AT3G53430.1 \| Symbols: \| Ribosomal protein L11 family protein \| chr3:19809895-19810395 REVERSE LENGTH=166;AT5G60670.1 \| Symbols: \| Ribosomal protein L11 family protein \| chr5:24381066-24381566 REVERSE LENGTH=166 | 0.1545 | 0.826787929 | 0.490643964 | 2.038137779 |
| AT1G26880 | AT1G26880.1 \| Symbols: \| Ribosomal protein L34e superfamily protein \| chr1:9315640-9316681 REVERSE LENGTH=120;AT1G69620.1 \| Symbols: RPL34 \| ribosomal protein L34 \| chr1:26189900-26191081 FORWARD LENGTH=119;AT3G28900.1 \| Symbols: \| Ribosomal protein L34e superfamily protein \| chr3:10902864-10904415 REVERSE LENGTH=120;AT1G26880.2 \| Symbols: \| Ribosomal protein L34e superfamily protein \| chr1:9315798-9316681 REVERSE LENGTH=96 | | 0.490364341 | 0.490364341 | 2.0393 |
| AT1G53750 | AT1G53750.1 \| Symbols: RPT1A \| regulatory particle triple-A 1A \| chr1:20065921-20068324 REVERSE LENGTH=426 | | 0.489212857 | 0.489212857 | 2.0441 |
| AT1G07830 | AT1G07830.1 \| Symbols: \| ribosomal protein L29 family protein \| chr1:2422549-2423392 FORWARD LENGTH=144 | | 0.489069301 | 0.489069301 | 2.0447 |
| AT2G38040 | AT2G38040.1 \| Symbols: CAC3 \| acetyl Co-enzyme a carboxylase carboxyltransferase alpha subunit \| chr2:15917612-15920749 FORWARD LENGTH=769;AT2G38040.2 \| Symbols: CAC3 \| acetyl Co-enzyme a carboxylase carboxyltransferase alpha subunit \| chr2:15917612-15920749 FORWARD LENGTH=769 | | 0.487662148 | 0.487662148 | 2.0506 |
| AT5G23540 | AT5G23540.1 \| Symbols: \| Mov34/MPN/PAD-1 family protein \| chr5:7937772-7939339 FORWARD LENGTH=308;AT5G23540.2 \| Symbols: \| Mov34/MPN/PAD-1 family protein \| chr5:7938109-7939339 FORWARD LENGTH=259 | 0.14805 | 0.82088327 | 0.484466635 | 2.064125633 |
| AT1G20220 | AT1G20220.1 \| Symbols: \| Alba DNA/RNA-binding protein \| chr1:7005090-7007285 REVERSE LENGTH=315 | | 0.483629153 | 0.483629153 | 2.0677 |
| AT4G26600 | AT4G26600.1 \| Symbols: \| S-adenosyl-L-methionine-dependent methyltransferases superfamily protein \| chr4:13419629-13423418 FORWARD LENGTH=671 | | 0.482299605 | 0.482299605 | 2.0734 |
| AT3G16950 | AT3G16950.2 \| Symbols: LPD1, ptlpd1 \| lipoamide dehydrogenase 1 \| chr3:5786508-5790383 REVERSE LENGTH=623;AT3G16950.1 \| Symbols: LPD1, ptlpd1 \| lipoamide dehydrogenase 1 \| chr3:5786761-5790383 REVERSE LENGTH=570 | | 0.480399693 | 0.480399693 | 2.0816 |
| AT4G38800 | AT4G38800.1 \| Symbols: ATMTN1, ATMTAN1, MTN1, MTAN1 \| methylthioadenosine nucleosidase 1 \| chr4:18113355-18114996 REVERSE LENGTH=267 | 0.47764 | | 0.47764 | 2.093626999 |
| AT3G02200 | AT3G02200.2 \| Symbols: \| Proteasome component (PCI) domain protein \| chr3:406692-408919 FORWARD LENGTH=417;AT3G02200.1 \| Symbols: \| Proteasome component (PCI) domain protein \| chr3:406692-408675 FORWARD LENGTH=364 | 0.10881 | 0.844309355 | 0.476559677 | 2.098373084 |
| AT5G15810 | AT5G15810.1 \| Symbols: \| N2,N2-dimethylguanosine tRNA methyltransferase \| chr5:5157798-5161121 FORWARD LENGTH=691;AT3G02320.1 \| Symbols: \| N2,N2-dimethylguanosine tRNA methyltransferase \| chr3:470097-473492 FORWARD LENGTH=589 | | 0.476553565 | 0.476553565 | 2.0984 |
| AT3G57560 | AT3G57560.1 \| Symbols: NAGK \| N-acetyl-l-glutamate kinase \| chr3:21311164-21312207 REVERSE LENGTH=347 | 0.12008 | 0.828363154 | 0.474221577 | 2.108718894 |
| AT5G58290 | AT5G58290.1 \| Symbols: RPT3 \| regulatory particle triple-A ATPase 3 \| chr5:23569155-23571116 FORWARD LENGTH=408 | 0.14113 | 0.807037366 | 0.474083683 | 2.109332247 |
| AT2G07696 | AT2G07696.1 \| Symbols: \| Ribosomal protein S7p/S5e family protein \| chr2:3351340-3351786 REVERSE LENGTH=148;ATMG01270.1 \| Symbols: RPS7 \| mitochondrial ribosomal protein S7 \| chrM:314627-315073 FORWARD LENGTH=148 | 0.48535 | 0.459263342 | 0.472306671 | 2.117268423 |
| AT5G63030 | AT5G63030.1 \| Symbols: \| Thioredoxin superfamily protein \| chr5:25286352-25287517 FORWARD LENGTH=125 | | 0.471653618 | 0.471653618 | 2.1202 |

| | | | | |
|---|---|---|---|---|
| AT3G58570 | AT3G58570.1 \| Symbols: \| P-loop containing nucleoside triphosphate hydrolases superfamily protein \| chr3:21657099-21660352 FORWARD LENGTH=646 | 0.471564652 | 0.471564652 | 2.1206 |
| AT2G36130 | AT2G36130.1 \| Symbols: \| Cyclophilin-like peptidyl-prolyl cis-trans isomerase family protein \| chr2:15166863-15168259 FORWARD LENGTH=164 | 0.470987189 | 0.470987189 | 2.1232 |
| AT1G27090 | AT1G27090.1 \| Symbols: \| glycine-rich protein \| chr1:9404041-9406098 REVERSE LENGTH=420 | 0.470588235 | 0.470588235 | 2.125 |
| AT3G18190 | AT3G18190.1 \| Symbols: \| TCP-1/cpn60 chaperonin family protein \| chr3:6232226-6233836 FORWARD LENGTH=536 | 0.46999107 | 0.46999107 | 2.1277 |
| AT5G57110 | AT5G57110.1 \| Symbols: ACA8, AT-ACA8 \| autoinhibited Ca2+-ATPase, isoform 8 \| chr5:23109729-23116857 REVERSE LENGTH=1074;AT5G57110.2 \| Symbols: ACA8, AT-ACA8 \| autoinhibited Ca2+-ATPase, isoform 8 \| chr5:23109729-23116857 REVERSE LENGTH=1074;AT3G21180.1 \| Symbols: ACA9, ATACA9 \| autoinhibited Ca(2+)-ATPase 9 \| chr3:7425770-7431941 FORWARD LENGTH=1086;AT4G29900.1 \| Symbols: ACA10, CIF1, ATACA10 \| autoinhibited Ca(2+)-ATPase 10 \| chr4:14611225-14618775 REVERSE LENGTH=1069 | 0.468142877 | 0.468142877 | 2.1361 |
| AT2G33040 | AT2G33040.1 \| Symbols: ATP3 \| gamma subunit of Mt ATP synthase \| chr2:14018978-14021047 REVERSE LENGTH=325 | 0.4629201 | 0.4629201 | 2.1602 |
| AT5G52280 | AT5G52280.1 \| Symbols: \| Myosin heavy chain-related protein \| chr5:21226959-21230109 FORWARD LENGTH=853 | 0.461275889 | 0.461275889 | 2.1679 |
| AT3G12580 | AT3G12580.1 \| Symbols: HSP70, ATHSP70 \| heat shock protein 70 \| chr3:3991487-3993689 REVERSE LENGTH=650 | 0.460765793 | 0.460765793 | 2.1703 |
| AT3G01130 | AT3G01130.1 \| Symbols: \| unknown protein; FUNCTIONS IN: molecular_function unknown; INVOLVED IN: biological_process unknown; LOCATED IN: endomembrane system; EXPRESSED IN: male gametophyte, pollen tube; EXPRESSED DURING: M germinated pollen stage; BEST Arabidopsis thaliana protein match is: unknown protein (TAIR:AT5G15320.1); Has 64 Blast hits to 64 proteins in 11 species: Archae - 0; Bacteria - 0; Metazoa - 0; Fungi - 0; Plants - 64; Viruses - 0; Other Eukaryotes - 0 (source: NCBI BLink). \| chr3:44714-45522 REVERSE LENGTH=53;AT3G01130.2 \| Symbols: \| unknown protein; FUNCTIONS IN: molecular_function unknown; INVOLVED IN: biological_process unknown; EXPRESSED IN: male gametophyte, pollen tube; EXPRESSED DURING: M germinated pollen stage; BEST Arabidopsis thaliana protein match is: unknown protein (TAIR:AT5G15320.1); Has 64 Blast hits to 64 proteins in 11 species: Archae - 0; Bacteria - 0; Metazoa - 0; Fungi - 0; Plants - 64; Viruses - 0; Other Eukaryotes - 0 (source: NCBI BLink). \| chr3:44911-45522 REVERSE LENGTH=50 | 0.458694555 | 0.458694555 | 2.1801 |
| AT1G30120 | AT1G30120.1 \| Symbols: PDH-E1 BETA \| pyruvate dehydrogenase E1 beta \| chr1:10584350-10586477 REVERSE LENGTH=406;AT2G34590.1 \| Symbols: \| Transketolase family protein \| chr2:14568956-14570844 REVERSE LENGTH=406 | 0.458547322 | 0.458547322 | 2.1808 |

| | | | | | |
|---|---|---|---|---|---|
| AT5G65270 | AT5G65270.1 \| Symbols: AtRABA4a, RABA4a \| RAB GTPase homolog A4A \| chr5:26083437-26084550 FORWARD LENGTH=226;AT4G39990.1 \| Symbols: ATRABA4B, ATRAB11G, ATGB3, RABA4B \| RAB GTPase homolog A4B \| chr4:18542722-18543779 FORWARD LENGTH=224;AT5G47960.1 \| Symbols: SMG1, ATRABA4C, RABA4C \| RAB GTPase homolog A4C \| chr5:19421533-19422473 REVERSE LENGTH=223;AT3G12160.1 \| Symbols: ATRABA4D, RABA4D \| RAB GTPase homolog A4D \| chr3:3879495-3880437 REVERSE LENGTH=222;AT1G05810.1 \| Symbols: ARA, ARA-1, ATRAB11D, ATRABA5E, RABA5E \| RAB GTPase homolog A5E \| chr1:1748314-1749350 FORWARD LENGTH=261;AT1G01200.1 \| Symbols: ATRABA3, ATRAB-A3, RABA3 \| RAB GTPase homolog A3 \| chr1:86715-88145 REVERSE LENGTH=237;AT5G47520.1 \| Symbols: AtRABA5a, RABA5a \| RAB GTPase homolog A5A \| chr5:19277596-19278366 REVERSE LENGTH=221;AT2G31680.1 \| Symbols: AtRABA5d, RABA5d \| RAB GTPase homolog A5D \| chr2:13473781-13474957 REVERSE LENGTH=219;AT3G07410.1 \| Symbols: AtRABA5b, RABA5b \| RAB GTPase homolog A5B \| chr3:2372485-2373482 REVERSE LENGTH=217;AT2G43130.1 \| Symbols: ARA4, ATRAB11F, ATRABA5C, ARA-4, RABA5C \| P-loop containing nucleoside triphosphate hydrolases superfamily protein \| chr2:17929899-17930904 REVERSE LENGTH=214 | | 0.458547322 | 0.458547322 | 2.1808 |
| AT2G43070 | AT2G43070.1 \| Symbols: SPPL3, ATSPPL3 \| SIGNAL PEPTIDE PEPTIDASE-LIKE 3 \| chr2:17911233-17914776 REVERSE LENGTH=540 | | 0.45712196 | 0.45712196 | 2.1876 |
| AT5G28060 | AT5G28060.1 \| Symbols: \| Ribosomal protein S24e family protein \| chr5:10069791-10070792 REVERSE LENGTH=133 | 0.33714 | 0.572409845 | 0.454774923 | 2.198889934 |
| AT3G17390 | AT3G17390.1 \| Symbols: MTO3, SAMS3, MAT4 \| S-adenosylmethionine synthetase family protein \| chr3:5952484-5953665 REVERSE LENGTH=393;AT1G02500.1 \| Symbols: SAM1, SAM-1, MAT1, AtSAM1 \| S-adenosylmethionine synthetase 1 \| chr1:519037-520218 FORWARD LENGTH=393;AT1G02500.2 \| Symbols: SAM1, SAM-1, MAT1, AtSAM1 \| S-adenosylmethionine synthetase 1 \| chr1:519037-520218 FORWARD LENGTH=393;AT4G01850.1 \| Symbols: SAM-2, MAT2, SAM2, AtSAM2 \| S-adenosylmethionine synthetase 2 \| chr4:796298-797479 REVERSE LENGTH=393;AT4G01850.2 \| Symbols: SAM-2, MAT2, SAM2, AtSAM2 \| S-adenosylmethionine synthetase 2 \| chr4:796298-797479 REVERSE LENGTH=393 | | 0.454669455 | 0.454669455 | 2.1994 |
| ATCG00770 | ATCG00770.1 \| Symbols: RPS8 \| ribosomal protein S8 \| chrC:80068-80472 REVERSE LENGTH=134 | | 0.454628114 | 0.454628114 | 2.1996 |
| AT5G09810 | AT5G09810.1 \| Symbols: ACT7 \| actin 7 \| chr5:3052809-3054220 FORWARD LENGTH=377;AT3G12110.1 \| Symbols: ACT11 \| actin-11 \| chr3:3858116-3859609 FORWARD LENGTH=377;AT3G46520.1 \| Symbols: ACT12 \| actin-12 \| chr3:17128567-17129981 FORWARD LENGTH=377;AT5G59370.1 \| Symbols: ACT4 \| actin 4 \| chr5:23950109-23951586 FORWARD LENGTH=377;AT5G59370.2 \| Symbols: ACT4 \| actin 4 \| chr5:23950109-23951586 FORWARD LENGTH=377;CON__P60712;AT2G42100.1 \| Symbols: \| Actin-like ATPase superfamily protein \| chr2:17560211-17561945 FORWARD LENGTH=378;AT2G42170.1 \| Symbols: \| Actin family protein \| chr2:17578683-17580222 FORWARD LENGTH=329 | 0.14306 | 0.765990042 | 0.454525021 | 2.200098902 |
| AT4G38510 | AT4G38510.5 \| Symbols: \| ATPase, V1 complex, subunit B protein \| chr4:18011155-18014897 REVERSE LENGTH=494;AT4G38510.1 \| Symbols: \| ATPase, V1 complex, subunit B protein \| chr4:18011155-18014789 REVERSE LENGTH=487;AT4G38510.2 \| Symbols: \| ATPase, V1 complex, subunit B protein \| chr4:18011155-18014789 REVERSE LENGTH=487;AT4G38510.3 \| Symbols: \| ATPase, V1 complex, subunit B protein \| chr4:18011155-18014789 REVERSE LENGTH=487;AT4G38510.4 \| Symbols: \| ATPase, V1 complex, subunit B protein \| chr4:18011155-18014789 REVERSE LENGTH=487 | 0.18792 | 0.719683339 | 0.45380167 | 2.203605819 |
| ATCG00820 | ATCG00820.1 \| Symbols: RPS19 \| ribosomal protein S19 \| chrC:84005-84283 REVERSE LENGTH=92 | | 0.453535308 | 0.453535308 | 2.2049 |

| | | | | | |
|---|---|---|---|---|---|
| AT3G61820 | AT3G61820.1 \| Symbols: \| Eukaryotic aspartyl protease family protein \| chr3:22880074-22881525 REVERSE LENGTH=483;AT1G01300.1 \| Symbols: \| Eukaryotic aspartyl protease family protein \| chr1:117065-118522 FORWARD LENGTH=485 | | 0.449478605 | 0.449478605 | 2.2248 |
| AT3G56450 | AT3G56450.1 \| Symbols: ALPHA-SNAP1 \| alpha-soluble NSF attachment protein 1 \| chr3:20930557-20932869 FORWARD LENGTH=381;AT3G56190.1 \| Symbols: ALPHA-SNAP2, ASNAP \| alpha-soluble NSF attachment protein 2 \| chr3:20846119-20848356 REVERSE LENGTH=289;AT3G56190.2 \| Symbols: ALPHA-SNAP2, ASNAP \| alpha-soluble NSF attachment protein 2 \| chr3:20846119-20848213 REVERSE LENGTH=240 | 0.44754 | | 0.44754 | 2.234437145 |
| AT5G67630 | AT5G67630.1 \| Symbols: \| P-loop containing nucleoside triphosphate hydrolases superfamily protein \| chr5:26967535-26969306 REVERSE LENGTH=469 | | 0.447087227 | 0.447087227 | 2.2367 |
| AT5G39850 | AT5G39850.1 \| Symbols: \| Ribosomal protein S4 \| chr5:15950053-15951171 FORWARD LENGTH=197 | 0.44441 | | 0.44441 | 2.250174389 |
| AT2G21410 | AT2G21410.1 \| Symbols: VHA-A2 \| vacuolar proton ATPase A2 \| chr2:9162703-9168141 FORWARD LENGTH=821 | 0.67338 | 0.20666708 | 0.44002354 | 2.272605688 |
| AT1G71880 | AT1G71880.1 \| Symbols: SUC1, ATSUC1 \| sucrose-proton symporter 1 \| chr1:27054334-27056100 FORWARD LENGTH=513;AT1G71890.1 \| Symbols: SUC5, ATSUC5 \| Major facilitator superfamily protein \| chr1:27058492-27060573 FORWARD LENGTH=512 | | 0.437445319 | 0.437445319 | 2.286 |
| AT5G05470 | AT5G05470.1 \| Symbols: EIF2 ALPHA, ATEIF2-A2, EIF2-A2 \| eukaryotic translation initiation factor 2 alpha subunit \| chr5:1620579-1622214 FORWARD LENGTH=344 | 0.22428 | 0.647542576 | 0.435911288 | 2.294044746 |
| AT3G16240 | AT3G16240.1 \| Symbols: DELTA-TIP, TIP2;1, DELTA-TIP1, AQP1, ATTIP2;1 \| delta tonoplast integral protein \| chr3:5505534-5506788 FORWARD LENGTH=250 | 0.22335 | 0.647542576 | 0.435446288 | 2.296494488 |
| AT2G07690 | AT2G07690.1 \| Symbols: MCM5 \| Minichromosome maintenance (MCM2/3/5) family protein \| chr2:3523379-3527388 REVERSE LENGTH=727;AT2G07690.2 \| Symbols: MCM5 \| Minichromosome maintenance (MCM2/3/5) family protein \| chr2:3523379-3527388 REVERSE LENGTH=725 | | 0.434669217 | 0.434669217 | 2.3006 |
| AT3G11560 | AT3G11560.2 \| Symbols: \| LETM1-like protein \| chr3:3639553-3645506 FORWARD LENGTH=872;AT3G11560.3 \| Symbols: \| LETM1-like protein \| chr3:3639553-3645506 FORWARD LENGTH=872;AT3G11560.4 \| Symbols: \| LETM1-like protein \| chr3:3639553-3645506 FORWARD LENGTH=872;AT3G11560.1 \| Symbols: \| LETM1-like protein \| chr3:3639553-3643450 FORWARD LENGTH=619 | | 0.434555884 | 0.434555884 | 2.3012 |

| | | | | | |
|---|---|---|---|---|---|
| AT4G17530 | AT4G17530.1 \| Symbols: RAB1C, ATRAB1C, ATRABD2C \| RAB GTPase homolog 1C \| chr4:9773721-9775424 REVERSE LENGTH=202;AT5G47200.1 \| Symbols: ATRABD2B, ATRAB1A, RAB1A \| RAB GTPase homolog 1A \| chr5:19167029-19168718 FORWARD LENGTH=202;AT3G09900.1 \| Symbols: ATRABE1E, ATRAB8E, RABE1e \| RAB GTPase homolog E1E \| chr3:3034687-3036379 FORWARD LENGTH=218;AT3G53610.1 \| Symbols: ATRAB8, AtRab8B, AtRABE1a, RAB8 \| RAB GTPase homolog 8 \| chr3:19876531-19878264 REVERSE LENGTH=216;AT3G53610.2 \| Symbols: ATRAB8, RAB8 \| RAB GTPase homolog 8 \| chr3:19876531-19878264 REVERSE LENGTH=216;AT3G53610.3 \| Symbols: ATRAB8, RAB8 \| RAB GTPase homolog 8 \| chr3:19876531-19878264 REVERSE LENGTH=216;AT5G03520.1 \| Symbols: ATRAB8C, ATRABE1D, ATRAB-E1D, RAB-E1D, RAB8C \| RAB GTPase homolog 8C \| chr5:883679-885158 FORWARD LENGTH=216;AT5G59840.1 \| Symbols: \| Ras-related small GTP-binding family protein \| chr5:24107450-24109049 REVERSE LENGTH=216;AT5G03530.1 \| Symbols: ATRAB ALPHA, ATRAB, ATRAB18B, ATRABC2A, RABC2A \| RAB GTPase homolog C2A \| chr5:885741-887061 REVERSE LENGTH=210;AT3G09910.1 \| Symbols: ATRAB18C, ATRABC2B, RABC2b \| RAB GTPase homolog C2B \| chr3:3036864-3038121 REVERSE LENGTH=205;AT3G09910.2 \| Symbols: RABC2b \| RAB GTPase homolog C2B \| chr3:3036864-3038121 REVERSE LENGTH=205;AT3G09910.3 \| Symbols: RABC2b \| RAB GTPase homolog C2B \| chr3:3036864-3038121 REVERSE LENGTH=205;AT5G39620.1 \| Symbols: AtRABG1, RABG1 \| RAB GTPase homolog G1 \| chr5:15864166-15865782 REVERSE LENGTH=204;AT1G02130.1 \| Symbols: ATRAB1B, ARA5, ARA-5, ATRABD2A, RABD2A, RA-5 \| RAS 5 \| chr1:400350-401788 REVERSE LENGTH=203 | 0.31723 | 0.548395942 | 0.432812971 | 2.3104668 |
| AT3G22200 | AT3G22200.2 \| Symbols: POP2 \| Pyridoxal phosphate (PLP)-dependent transferases superfamily protein \| chr3:7835457-7838863 FORWARD LENGTH=513;AT3G22200.1 \| Symbols: POP2, GABA-T, HER1 \| Pyridoxal phosphate (PLP)-dependent transferases superfamily protein \| chr3:7835286-7838863 FORWARD LENGTH=504 | | 0.432806752 | 0.432806752 | 2.3105 |
| AT1G24280 | AT1G24280.1 \| Symbols: G6PD3 \| glucose-6-phosphate dehydrogenase 3 \| chr1:8609495-8612383 FORWARD LENGTH=599;AT5G13110.1 \| Symbols: G6PD2 \| glucose-6-phosphate dehydrogenase 2 \| chr5:4158952-4161640 FORWARD LENGTH=596;AT5G35790.1 \| Symbols: G6PD1 \| glucose-6-phosphate dehydrogenase 1 \| chr5:13956879-13959686 REVERSE LENGTH=576 | | 0.431127398 | 0.431127398 | 2.3195 |
| AT3G06720 | AT3G06720.1 \| Symbols: AT-IMP, ATKAP ALPHA, AIMP ALPHA, IMPA-1, IMPA1 \| importin alpha isoform 1 \| chr3:2120559-2123555 FORWARD LENGTH=532;AT3G06720.2 \| Symbols: AT-IMP, ATKAP ALPHA, AIMP ALPHA \| importin alpha isoform 1 \| chr3:2120559-2123555 FORWARD LENGTH=532 | | 0.430811649 | 0.430811649 | 2.3212 |
| AT3G18420 | AT3G18420.1 \| Symbols: \| Protein prenylyltransferase superfamily protein \| chr3:6324771-6325721 REVERSE LENGTH=316 | | 0.428192173 | 0.428192173 | 2.3354 |
| AT3G22320 | AT3G22320.1 \| Symbols: ATRPABC24.3, RPB5A, NRPB5, NRPD5 \| Eukaryotic rpb5 RNA polymerase subunit family protein \| chr3:7891045-7892094 REVERSE LENGTH=205 | | 0.428192173 | 0.428192173 | 2.3354 |
| AT5G22330 | AT5G22330.1 \| Symbols: ATTIP49A, RIN1 \| P-loop containing nucleoside triphosphate hydrolases superfamily protein \| chr5:7391026-7394071 REVERSE LENGTH=458 | 0.25733 | 0.592241635 | 0.424785817 | 2.354127561 |
| AT2G42790 | AT2G42790.1 \| Symbols: CSY3 \| citrate synthase 3 \| chr2:17803132-17805991 REVERSE LENGTH=509;AT3G58740.1 \| Symbols: CSY1 \| citrate synthase 1 \| chr3:21721066-21723726 REVERSE LENGTH=480 | | 0.422404325 | 0.422404325 | 2.3674 |
| AT2G31610 | AT2G31610.1 \| Symbols: \| Ribosomal protein S3 family protein \| chr2:13450384-13451669 FORWARD LENGTH=250;AT3G53870.1 \| Symbols: \| Ribosomal protein S3 family protein \| chr3:19951547-19952782 FORWARD LENGTH=249;AT5G35530.1 \| Symbols: \| Ribosomal protein S3 family protein \| chr5:13710355-13712192 REVERSE LENGTH=248 | 0.10789 | 0.735186002 | 0.421538001 | 2.372265365 |

| | | | | | |
|---|---|---|---|---|---|
| AT5G12190 | AT5G12190.1 \| Symbols: \| RNA-binding (RRM/RBD/RNP motifs) family protein \| chr5:3940731-3941105 REVERSE LENGTH=124 | 0.63647 | 0.205326161 | 0.42089808 | 2.375872086 |
| AT1G49670 | AT1G49670.2 \| Symbols: NQR \| ARP protein (REF) \| chr1:18381591-18386021 REVERSE LENGTH=652;AT1G49670.1 \| Symbols: NQR \| ARP protein (REF) \| chr1:18381591-18386021 REVERSE LENGTH=629 | | 0.420238696 | 0.420238696 | 2.3796 |
| AT2G29550 | AT2G29550.1 \| Symbols: TUB7 \| tubulin beta-7 chain \| chr2:12644258-12645932 REVERSE LENGTH=449;AT5G23860.1 \| Symbols: TUB8 \| tubulin beta 8 \| chr5:8042962-8044528 FORWARD LENGTH=449;AT5G23860.2 \| Symbols: TUB8 \| tubulin beta 8 \| chr5:8042962-8044528 FORWARD LENGTH=449;AT5G12250.1 \| Symbols: TUB6 \| beta-6 tubulin \| chr5:3961317-3962971 REVERSE LENGTH=449;AT1G75780.1 \| Symbols: TUB1 \| tubulin beta-1 chain \| chr1:28451378-28453602 REVERSE LENGTH=447;AT1G20010.1 \| Symbols: TUB5 \| tubulin beta-5 chain \| chr1:6938033-6940481 REVERSE LENGTH=449 | | 0.419621501 | 0.419621501 | 2.3831 |
| AT1G42440 | AT1G42440.1 \| Symbols: \| FUNCTIONS IN: molecular_function unknown; INVOLVED IN: ribosome biogenesis; LOCATED IN: nucleus; EXPRESSED IN: 22 plant structures; EXPRESSED DURING: 13 growth stages; CONTAINS InterPro DOMAIN/s: AARP2CN (InterPro:IPR012948), Protein of unknown function DUF663 (InterPro:IPR007034); BEST Arabidopsis thaliana protein match is: P-loop containing nucleoside triphosphate hydrolases superfamily protein (TAIR:AT1G06720.1); Has 2741 Blast hits to 2088 proteins in 291 species: Archae - 2; Bacteria - 131; Metazoa - 833; Fungi - 650; Plants - 171; Viruses - 49; Other Eukaryotes - 905 (source: NCBI BLink). \| chr1:15895528-15899939 REVERSE LENGTH=793 | | 0.418182579 | 0.418182579 | 2.3913 |
| AT4G20840 | AT4G20840.1 \| Symbols: \| FAD-binding Berberine family protein \| chr4:11157916-11159535 FORWARD LENGTH=539 | 0.41712 | | 0.41712 | 2.397391638 |
| AT2G01690 | AT2G01690.2 \| Symbols: \| ARM repeat superfamily protein \| chr2:309144-313499 REVERSE LENGTH=744;AT2G01690.1 \| Symbols: \| ARM repeat superfamily protein \| chr2:309144-313499 REVERSE LENGTH=743 | 0.28453 | 0.549088513 | 0.416809257 | 2.399178963 |
| AT1G24180 | AT1G24180.1 \| Symbols: IAR4 \| Thiamin diphosphate-binding fold (THDP-binding) superfamily protein \| chr1:8560777-8563382 REVERSE LENGTH=393 | 0.12304 | 0.70666384 | 0.41485192 | 2.410498667 |
| AT4G10750 | AT4G10750.1 \| Symbols: \| Phosphoenolpyruvate carboxylase family protein \| chr4:6618724-6619800 FORWARD LENGTH=358 | | 0.414010102 | 0.414010102 | 2.4154 |
| AT4G02930 | AT4G02930.1 \| Symbols: \| GTP binding Elongation factor Tu family protein \| chr4:1295751-1298354 REVERSE LENGTH=454 | 0.14294 | 0.682081713 | 0.412510857 | 2.424178622 |
| AT5G22100 | AT5G22100.1 \| Symbols: \| RNA cyclase family protein \| chr5:7329015-7330718 FORWARD LENGTH=375 | | 0.41064389 | 0.41064389 | 2.4352 |
| AT1G43710 | AT1G43710.1 \| Symbols: emb1075 \| Pyridoxal phosphate (PLP)-dependent transferases superfamily protein \| chr1:16486534-16488298 REVERSE LENGTH=482 | | 0.410407945 | 0.410407945 | 2.4366 |
| AT5G62200 | AT5G62200.1 \| Symbols: \| Embryo-specific protein 3, (ATS3) \| chr5:24984463-24985674 REVERSE LENGTH=190 | | 0.410104987 | 0.410104987 | 2.4384 |
| AT2G34470 | AT2G34470.2 \| Symbols: UREG, PSKF109 \| urease accessory protein G \| chr2:14530900-14532411 REVERSE LENGTH=276;AT2G34470.1 \| Symbols: UREG, PSKF109 \| urease accessory protein G \| chr2:14530900-14532411 REVERSE LENGTH=275 | | 0.40950041 | 0.40950041 | 2.442 |
| AT5G36230 | AT5G36230.1 \| Symbols: \| ARM repeat superfamily protein \| chr5:14273519-14276773 FORWARD LENGTH=411 | | 0.409081612 | 0.409081612 | 2.4445 |
| AT1G43890 | AT1G43890.1 \| Symbols: ATRAB18, ATRABC1, RAB18-1, RABC1, ATRAB-C1, RAB18 \| RAB GTPASE HOMOLOG B18 \| chr1:16646934-16648395 FORWARD LENGTH=212;AT1G43890.2 \| Symbols: ATRAB18, ATRABC1, RAB18-1, RABC1, ATRAB-C1, RAB18 \| RAB GTPASE HOMOLOG B18 \| chr1:16646934-16648395 FORWARD LENGTH=212;AT1G43890.3 \| Symbols: ATRAB18, ATRABC1, RAB18-1, RABC1, ATRAB-C1, RAB18 \| RAB GTPASE HOMOLOG B18 \| chr1:16646934-16648395 FORWARD LENGTH=212 | 0.15376 | 0.663173951 | 0.408466975 | 2.448178336 |

| | | | | | |
|---|---|---|---|---|---|
| AT1G30580 | AT1G30580.1 \| Symbols: \| GTP binding \| chr1:10831953-10835454 REVERSE LENGTH=394 | 0.3007 | 0.515995872 | 0.408347936 | 2.448892015 |
| AT3G06610 | AT3G06610.1 \| Symbols: \| DNA-binding enhancer protein-related \| chr3:2060837-2061845 FORWARD LENGTH=115 | | 0.407946804 | 0.407946804 | 2.4513 |
| AT3G05530 | AT3G05530.1 \| Symbols: RPT5A, ATS6A.2 \| regulatory particle triple-A ATPase 5A \| chr3:1603540-1605993 FORWARD LENGTH=424;AT1G09100.1 \| Symbols: RPT5B \| 26S proteasome AAA-ATPase subunit RPT5B \| chr1:2936675-2939258 REVERSE LENGTH=423;AT5G19990.1 \| Symbols: RPT6A, ATSUG1 \| regulatory particle triple-A ATPase 6A \| chr5:6752144-6754918 FORWARD LENGTH=419;AT5G20000.1 \| Symbols: \| AAA-type ATPase family protein \| chr5:6756915-6759550 FORWARD LENGTH=419;AT5G53540.1 \| Symbols: \| P-loop containing nucleoside triphosphate hydrolases superfamily protein \| chr5:21749561-21751099 REVERSE LENGTH=403;AT4G27680.1 \| Symbols: \| P-loop containing nucleoside triphosphate hydrolases superfamily protein \| chr4:13821263-13823083 FORWARD LENGTH=398 | 0.15832 | 0.653167864 | 0.405743932 | 2.464608638 |
| AT5G46430 | AT5G46430.1 \| Symbols: \| Ribosomal protein L32e \| chr5:18833429-18834409 FORWARD LENGTH=133;AT5G46430.2 \| Symbols: \| Ribosomal protein L32e \| chr5:18833429-18834409 FORWARD LENGTH=133 | | 0.405153553 | 0.405153553 | 2.4682 |
| AT2G27100 | AT2G27100.1 \| Symbols: SE \| C2H2 zinc-finger protein SERRATE (SE) \| chr2:11572587-11576357 FORWARD LENGTH=720 | 0.29414 | 0.505740151 | 0.399940075 | 2.500374585 |
| AT3G59140 | AT3G59140.1 \| Symbols: ATMRP14, MRP14, ABCC10 \| multidrug resistance-associated protein 14 \| chr3:21863519-21868701 REVERSE LENGTH=1453 | | 0.398708185 | 0.398708185 | 2.5081 |
| AT3G62150 | AT3G62150.1 \| Symbols: PGP21 \| P-glycoprotein 21 \| chr3:23008755-23013579 REVERSE LENGTH=1296;AT2G47000.1 \| Symbols: MDR4, PGP4, ABCB4, ATPGP4 \| ATP binding cassette subfamily B4 \| chr2:19310008-19314750 REVERSE LENGTH=1286;AT2G36910.1 \| Symbols: ATPGP1, PGP1, ABCB1 \| ATP binding cassette subfamily B1 \| chr2:15502162-15507050 FORWARD LENGTH=1286;AT1G02520.1 \| Symbols: PGP11 \| P-glycoprotein 11 \| chr1:524134-528745 FORWARD LENGTH=1278;AT1G02530.1 \| Symbols: PGP12 \| P-glycoprotein 12 \| chr1:529836-534542 FORWARD LENGTH=1273;AT3G28860.1 \| Symbols: ATMDR1, ATMDR11, PGP19, MDR11, MDR1, ATPGP19, ABCB19, ATABCB19 \| ATP binding cassette subfamily B19 \| chr3:10870287-10877286 REVERSE LENGTH=1252;AT5G46540.1 \| Symbols: PGP7 \| P-glycoprotein 7 \| chr5:18877192-18882347 REVERSE LENGTH=1248;AT4G18050.1 \| Symbols: PGP9 \| P-glycoprotein 9 \| chr4:10022205-10027280 FORWARD LENGTH=1236;AT4G01830.1 \| Symbols: PGP5 \| P-glycoprotein 5 \| chr4:785683-790447 REVERSE LENGTH=1230;AT4G01820.1 \| Symbols: PGP3, MDR3 \| P-glycoprotein 3 \| chr4:780734-785329 REVERSE LENGTH=1229 | | 0.394104201 | 0.394104201 | 2.5374 |
| AT5G55920 | AT5G55920.1 \| Symbols: OLI2 \| S-adenosyl-L-methionine-dependent methyltransferases superfamily protein \| chr5:22645742-22649383 REVERSE LENGTH=682 | | 0.392079984 | 0.392079984 | 2.5505 |
| AT5G23740 | AT5G23740.1 \| Symbols: RPS11-BETA \| ribosomal protein S11-beta \| chr5:8008251-8009330 REVERSE LENGTH=159 | | 0.391328168 | 0.391328168 | 2.5554 |
| AT1G80500 | AT1G80500.1 \| Symbols: \| SNARE-like superfamily protein \| chr1:30271195-30272431 FORWARD LENGTH=135 | | 0.389559797 | 0.389559797 | 2.567 |
| AT5G02870 | AT5G02870.1 \| Symbols: \| Ribosomal protein L4/L1 family \| chr5:657830-659526 FORWARD LENGTH=407;AT5G02870.2 \| Symbols: \| Ribosomal protein L4/L1 family \| chr5:657830-659526 FORWARD LENGTH=406 | 0.10685 | 0.670061646 | 0.388455823 | 2.574295303 |
| AT1G43680 | AT1G43680.1 \| Symbols: \| BEST Arabidopsis thaliana protein match is: nucleic acid binding;zinc ion binding (TAIR:AT5G36228.1); Has 59 Blast hits to 57 proteins in 2 species: Archae - 0; Bacteria - 0; Metazoa - 0; Fungi - 0; Plants - 59; Viruses - 0; Other Eukaryotes - 0 (source: NCBI BLink). \| chr1:16472205-16473266 FORWARD LENGTH=247 | 0.38816 | | 0.38816 | 2.576257214 |

| | | | | | |
|---|---|---|---|---|---|
| AT4G00810 | AT4G00810.1 \| Symbols: \| 60S acidic ribosomal protein family \| chr4:346179-346957 REVERSE LENGTH=113;AT4G00810.2 \| Symbols: \| 60S acidic ribosomal protein family \| chr4:346179-346957 REVERSE LENGTH=113 | | 0.387596899 | 0.387596899 | 2.58 |
| AT1G75500 | AT1G75500.1 \| Symbols: WAT1 \| Walls Are Thin 1 \| chr1:28338282-28340091 REVERSE LENGTH=389;AT1G75500.2 \| Symbols: WAT1 \| Walls Are Thin 1 \| chr1:28338282-28340091 REVERSE LENGTH=389 | 0.39898 | 0.372897789 | 0.385938894 | 2.591083756 |
| AT1G31280 | AT1G31280.1 \| Symbols: AGO2 \| Argonaute family protein \| chr1:11181777-11185112 FORWARD LENGTH=1014 | | 0.385297064 | 0.385297064 | 2.5954 |
| AT2G17510 | AT2G17510.2 \| Symbols: EMB2763 \| ribonuclease II family protein \| chr2:7609395-7616242 REVERSE LENGTH=1012;AT2G17510.1 \| Symbols: EMB2763 \| ribonuclease II family protein \| chr2:7609395-7615881 REVERSE LENGTH=933 | | 0.384186868 | 0.384186868 | 2.6029 |
| AT5G02450 | AT5G02450.1 \| Symbols: \| Ribosomal protein L36e family protein \| chr5:533501-534394 FORWARD LENGTH=108;AT3G53740.2 \| Symbols: \| Ribosomal protein L36e family protein \| chr3:19913921-19914813 REVERSE LENGTH=112;AT3G53740.3 \| Symbols: \| Ribosomal protein L36e family protein \| chr3:19913921-19914813 REVERSE LENGTH=112;AT3G53740.4 \| Symbols: \| Ribosomal protein L36e family protein \| chr3:19913921-19914813 REVERSE LENGTH=112;AT3G53740.1 \| Symbols: \| Ribosomal protein L36e family protein \| chr3:19913921-19914813 REVERSE LENGTH=103 | 0.19934 | 0.568343279 | 0.38384164 | 2.605241059 |
| AT1G08130 | AT1G08130.1 \| Symbols: ATLIG1, LIG1 \| DNA ligase 1 \| chr1:2542913-2547815 REVERSE LENGTH=790 | | 0.382511571 | 0.382511571 | 2.6143 |
| AT2G24820 | AT2G24820.1 \| Symbols: TIC55-II \| translocon at the inner envelope membrane of chloroplasts 55-II \| chr2:10575038-10576829 FORWARD LENGTH=539 | | 0.38197097 | 0.38197097 | 2.618 |
| AT3G24830 | AT3G24830.1 \| Symbols: \| Ribosomal protein L13 family protein \| chr3:9064613-9065871 FORWARD LENGTH=206 | | 0.377500944 | 0.377500944 | 2.649 |
| AT1G70870 | AT1G70870.1 \| Symbols: \| Polyketide cyclase/dehydrase and lipid transport superfamily protein \| chr1:26721020-26721563 REVERSE LENGTH=139 | 0.37382 | | 0.37382 | 2.675084265 |
| AT3G25780 | AT3G25780.1 \| Symbols: AOC3 \| allene oxide cyclase 3 \| chr3:9409362-9410409 FORWARD LENGTH=258 | | 0.373245745 | 0.373245745 | 2.6792 |
| AT3G62870 | AT3G62870.1 \| Symbols: \| Ribosomal protein L7Ae/L30e/S12e/Gadd45 family protein \| chr3:23242862-23244273 REVERSE LENGTH=256;AT2G47610.1 \| Symbols: \| Ribosomal protein L7Ae/L30e/S12e/Gadd45 family protein \| chr2:19529854-19531401 FORWARD LENGTH=257 | 0.087641 | 0.65681445 | 0.372227725 | 2.686527448 |
| AT5G02490 | AT5G02490.1 \| Symbols: \| Heat shock protein 70 (Hsp 70) family protein \| chr5:550296-552565 REVERSE LENGTH=653;AT1G56410.1 \| Symbols: ERD2, HSP70T-1 \| heat shock protein 70 (Hsp 70) family protein \| chr1:21117147-21119241 FORWARD LENGTH=617 | | 0.372093023 | 0.372093023 | 2.6875 |
| AT1G04270 | AT1G04270.1 \| Symbols: RPS15 \| cytosolic ribosomal protein S15 \| chr1:1141852-1142960 REVERSE LENGTH=152;AT1G04270.2 \| Symbols: RPS15 \| cytosolic ribosomal protein S15 \| chr1:1141852-1142960 REVERSE LENGTH=151;AT5G09510.1 \| Symbols: \| Ribosomal protein S19 family protein \| chr5:2955698-2956554 REVERSE LENGTH=152;AT5G09500.1 \| Symbols: \| Ribosomal protein S19 family protein \| chr5:2954044-2954850 REVERSE LENGTH=150 | 0.16637 | 0.57700075 | 0.371685375 | 2.690447532 |
| ATCG01120 | ATCG01120.1 \| Symbols: RPS15 \| chloroplast ribosomal protein S15 \| chrC:123296-123562 REVERSE LENGTH=88 | | 0.369412634 | 0.369412634 | 2.707 |
| AT1G29320 | AT1G29320.1 \| Symbols: \| Transducin/WD40 repeat-like superfamily protein \| chr1:10255355-10258238 FORWARD LENGTH=468 | | 0.366945545 | 0.366945545 | 2.7252 |
| ATCG00790 | ATCG00790.1 \| Symbols: RPL16 \| ribosomal protein L16 \| chrC:81189-82652 REVERSE LENGTH=135;AT2G28830.1 \| Symbols: PUB12, AtPUB12 \| PLANT U-BOX 12 \| chr2:12367001-12370608 REVERSE LENGTH=962 | | 0.366555478 | 0.366555478 | 2.7281 |
| AT5G23900 | AT5G23900.1 \| Symbols: \| Ribosomal protein L13e family protein \| chr5:8064176-8065081 REVERSE LENGTH=206 | 0.24543 | 0.487020893 | 0.366225447 | 2.730558483 |

| | | | | | |
|---|---|---|---|---|---|
| AT2G17930 | AT2G17930.1 \| Symbols: \| Phosphatidylinositol 3- and 4-kinase family protein with FAT domain \| chr2:7784455-7802230 REVERSE LENGTH=3858;AT4G36080.1 \| Symbols: \| phosphotransferases, alcohol group as acceptor;binding;inositol or phosphatidylinositol kinases \| chr4:17059996-17077628 REVERSE LENGTH=3834;AT4G36080.3 \| Symbols: \| phosphotransferases, alcohol group as acceptor;binding;inositol or phosphatidylinositol kinases \| chr4:17059996-17077628 REVERSE LENGTH=3809;AT4G36080.2 \| Symbols: \| phosphotransferases, alcohol group as acceptor;binding;inositol or phosphatidylinositol kinases \| chr4:17059996-17077628 REVERSE LENGTH=3804 | | 0.364537766 | 0.364537766 | 2.7432 |
| AT5G14040 | AT5G14040.1 \| Symbols: PHT3;1 \| phosphate transporter 3;1 \| chr5:4531059-4532965 REVERSE LENGTH=375 | 0.59937 | 0.127335006 | 0.363352503 | 2.752148374 |
| AT4G29830 | AT4G29830.1 \| Symbols: VIP3 \| Transducin/WD40 repeat-like superfamily protein \| chr4:14597728-14599157 FORWARD LENGTH=321 | | 0.362818373 | 0.362818373 | 2.7562 |
| AT1G44575 | AT1G44575.1 \| Symbols: NPQ4, PSBS \| Chlorophyll A-B binding family protein \| chr1:16871768-16873194 FORWARD LENGTH=265;AT1G44575.2 \| Symbols: NPQ4, PSBS \| Chlorophyll A-B binding family protein \| chr1:16871768-16872548 FORWARD LENGTH=205 | | 0.36272625 | 0.36272625 | 2.7569 |
| AT2G26780 | AT2G26780.1 \| Symbols: \| ARM repeat superfamily protein \| chr2:11410125-11423598 FORWARD LENGTH=1826 | | 0.362581581 | 0.362581581 | 2.758 |
| AT5G18620 | AT5G18620.2 \| Symbols: CHR17 \| chromatin remodeling factor17 \| chr5:6196190-6202058 REVERSE LENGTH=1072;AT5G18620.1 \| Symbols: CHR17 \| chromatin remodeling factor17 \| chr5:6196190-6202058 REVERSE LENGTH=1069;AT3G06400.3 \| Symbols: CHR11 \| chromatin-remodeling protein 11 \| chr3:1941066-1946700 FORWARD LENGTH=1057;AT3G06400.2 \| Symbols: CHR11 \| chromatin-remodeling protein 11 \| chr3:1941066-1946700 FORWARD LENGTH=1056;AT3G06400.1 \| Symbols: CHR11 \| chromatin-remodeling protein 11 \| chr3:1941066-1946700 FORWARD LENGTH=1055 | | 0.362542146 | 0.362542146 | 2.7583 |
| AT3G15000 | AT3G15000.1 \| Symbols: \| cobalt ion binding \| chr3:5050321-5052121 FORWARD LENGTH=395 | | 0.3620827 | 0.3620827 | 2.7618 |
| AT2G47110 | AT2G47110.1 \| Symbols: UBQ6 \| ubiquitin 6 \| chr2:19344701-19345174 FORWARD LENGTH=157;AT2G47110.2 \| Symbols: UBQ6 \| ubiquitin 6 \| chr2:19344701-19345174 FORWARD LENGTH=157;AT3G62250.1 \| Symbols: UBQ5 \| ubiquitin 5 \| chr3:23037138-23037611 FORWARD LENGTH=157;AT1G23410.1 \| Symbols: \| Ribosomal protein S27a / Ubiquitin family protein \| chr1:8314940-8315410 FORWARD LENGTH=156 | | 0.361154249 | 0.361154249 | 2.7689 |
| AT2G39990 | AT2G39990.1 \| Symbols: EIF2, AteIF3f, eIF3F \| eukaryotic translation initiation factor 2 \| chr2:16698332-16699929 REVERSE LENGTH=293 | 0.099618 | 0.622471211 | 0.361044605 | 2.769740872 |
| AT5G67500 | AT5G67500.2 \| Symbols: VDAC2 \| voltage dependent anion channel 2 \| chr5:26935223-26937123 FORWARD LENGTH=303;AT5G67500.1 \| Symbols: VDAC2, ATVDAC2 \| voltage dependent anion channel 2 \| chr5:26935223-26937123 FORWARD LENGTH=276 | 0.077798 | 0.642549637 | 0.360173818 | 2.776437233 |
| AT3G52090 | AT3G52090.2 \| Symbols: NRPB11 \| DNA-directed RNA polymerase, RBP11-like \| chr3:19318069-19318998 REVERSE LENGTH=146;AT3G52090.1 \| Symbols: ATRPB13.6, NRPB11, NRPD11, NRPE11 \| DNA-directed RNA polymerase, RBP11-like \| chr3:19318069-19318998 REVERSE LENGTH=116 | | 0.359751052 | 0.359751052 | 2.7797 |
| AT3G53430 | AT3G53430.1 \| Symbols: \| Ribosomal protein L11 family protein \| chr3:19809895-19810395 REVERSE LENGTH=166 | | 0.359699291 | 0.359699291 | 2.7801 |
| AT5G33320 | AT5G33320.1 \| Symbols: CUE1, PPT, ARAPPT \| Glucose-6-phosphate/phosphate translocator-related \| chr5:12588950-12591408 FORWARD LENGTH=408 | | 0.359014863 | 0.359014863 | 2.7854 |

| | | | | | |
|---|---|---|---|---|---|
| AT2G37270 | AT2G37270.1 \| Symbols: ATRPS5B, RPS5B \| ribosomal protein 5B \| chr2:15647883-15649042 REVERSE LENGTH=207;AT2G37270.2 \| Symbols: ATRPS5B, RPS5B \| ribosomal protein 5B \| chr2:15647883-15649042 REVERSE LENGTH=207;AT3G11940.1 \| Symbols: ATRPS5A, AML1, RPS5A \| ribosomal protein 5A \| chr3:3778175-3779354 REVERSE LENGTH=207;AT3G11940.2 \| Symbols: ATRPS5A, AML1, RPS5A \| ribosomal protein 5A \| chr3:3778175-3779354 REVERSE LENGTH=207 | 0.20079 | 0.516155673 | 0.358472836 | 2.789611649 |
| AT5G35590 | AT5G35590.1 \| Symbols: PAA1 \| proteasome alpha subunit A1 \| chr5:13765417-13767768 REVERSE LENGTH=246 | 0.12113 | 0.595699053 | 0.358414526 | 2.790065486 |
| AT1G50920 | AT1G50920.1 \| Symbols: \| Nucleolar GTP-binding protein \| chr1:18870555-18872570 FORWARD LENGTH=671 | | 0.357935428 | 0.357935428 | 2.7938 |
| AT2G28830 | AT2G28830.1 \| Symbols: PUB12, AtPUB12 \| PLANT U-BOX 12 \| chr2:12367001-12370608 REVERSE LENGTH=962;ATCG00790.1 \| Symbols: RPL16 \| ribosomal protein L16 \| chrC:81189-82652 REVERSE LENGTH=135 | 0.35662 | | 0.35662 | 2.80410521 |
| AT4G37870 | AT4G37870.1 \| Symbols: PCK1, PEPCK \| phosphoenolpyruvate carboxykinase 1 \| chr4:17802974-17806332 REVERSE LENGTH=671;AT5G65690.1 \| Symbols: PCK2, PEPCK \| phosphoenolpyruvate carboxykinase 2 \| chr5:26266477-26269486 FORWARD LENGTH=670 | | 0.356353788 | 0.356353788 | 2.8062 |
| AT1G74470 | AT1G74470.1 \| Symbols: \| Pyridine nucleotide-disulphide oxidoreductase family protein \| chr1:27991248-27992845 FORWARD LENGTH=467 | | 0.355947889 | 0.355947889 | 2.8094 |
| AT1G20200 | AT1G20200.1 \| Symbols: EMB2719, HAP15 \| PAM domain (PCI/PINT associated module) protein \| chr1:7001409-7004154 REVERSE LENGTH=488;AT1G75990.1 \| Symbols: \| PAM domain (PCI/PINT associated module) protein \| chr1:28524623-28526718 REVERSE LENGTH=487 | 0.17726 | 0.533760342 | 0.355510171 | 2.812859046 |
| AT1G16350 | AT1G16350.1 \| Symbols: \| Aldolase-type TIM barrel family protein \| chr1:5590951-5592872 FORWARD LENGTH=502 | | 0.35501278 | 0.35501278 | 2.8168 |
| AT1G15420 | AT1G15420.1 \| Symbols: \| CONTAINS InterPro DOMAIN/s: Small-subunit processome, Utp12 (InterPro:IPR007148); Has 764 Blast hits to 656 proteins in 193 species: Archae - 0; Bacteria - 42; Metazoa - 237; Fungi - 154; Plants - 85; Viruses - 23; Other Eukaryotes - 223 (source: NCBI BLink). \| chr1:5301794-5303296 REVERSE LENGTH=278 | 0.060629 | 0.64913989 | 0.354884445 | 2.81781863 |
| AT4G38130 | AT4G38130.1 \| Symbols: HD1, ATHD1, HDA1, RPD3A, HDA19, ATHDA19 \| histone deacetylase 1 \| chr4:17896493-17899057 REVERSE LENGTH=501;AT4G38130.2 \| Symbols: HD1, ATHD1, HDA1, RPD3A, HDA19 \| histone deacetylase 1 \| chr4:17896663-17899057 REVERSE LENGTH=469 | | 0.352621743 | 0.352621743 | 2.8359 |
| AT2G29580 | AT2G29580.1 \| Symbols: \| CCCH-type zinc fingerfamily protein with RNA-binding domain \| chr2:12652011-12654158 FORWARD LENGTH=483;AT1G07360.1 \| Symbols: \| CCCH-type zinc fingerfamily protein with RNA-binding domain \| chr1:2260562-2262795 REVERSE LENGTH=481;AT5G07060.1 \| Symbols: \| CCCH-type zinc fingerfamily protein with RNA-binding domain \| chr5:2194394-2195872 REVERSE LENGTH=363;AT5G07060.2 \| Symbols: \| CCCH-type zinc fingerfamily protein with RNA-binding domain \| chr5:2194159-2195872 REVERSE LENGTH=355 | | 0.351951571 | 0.351951571 | 2.8413 |
| AT2G36880 | AT2G36880.1 \| Symbols: MAT3 \| methionine adenosyltransferase 3 \| chr2:15479721-15480893 REVERSE LENGTH=390;AT2G36880.2 \| Symbols: MAT3 \| methionine adenosyltransferase 3 \| chr2:15479721-15480893 REVERSE LENGTH=390 | 0.11703 | 0.586544665 | 0.351787333 | 2.842626516 |
| AT2G43460 | AT2G43460.1 \| Symbols: \| Ribosomal L38e protein family \| chr2:18046285-18047292 REVERSE LENGTH=69;AT3G59540.1 \| Symbols: \| Ribosomal L38e protein family \| chr3:21995897-21996742 REVERSE LENGTH=69 | 0.54455 | 0.157388608 | 0.350969304 | 2.849252024 |
| AT5G44340 | AT5G44340.1 \| Symbols: TUB4 \| tubulin beta chain 4 \| chr5:17859442-17860994 REVERSE LENGTH=444 | 0.053542 | 0.647081662 | 0.350311831 | 2.854599565 |
| AT3G55620 | AT3G55620.1 \| Symbols: emb1624 \| Translation initiation factor IF6 \| chr3:20634581-20636312 FORWARD LENGTH=245 | 0.23211 | 0.468427956 | 0.350268978 | 2.854948805 |
| AT2G20280 | AT2G20280.1 \| Symbols: \| Zinc finger C-x8-C-x5-C-x3-H type family protein \| chr2:8740054-8742349 REVERSE LENGTH=371 | | 0.347656793 | 0.347656793 | 2.8764 |

| | | | | | |
|---|---|---|---|---|---|
| AT4G16630 | AT4G16630.1 \| Symbols: \| DEA(D/H)-box RNA helicase family protein \| chr4:9362176-9366449 REVERSE LENGTH=789 | | 0.346488341 | 0.346488341 | 2.8861 |
| AT5G18380 | AT5G18380.1 \| Symbols: \| Ribosomal protein S5 domain 2-like superfamily protein \| chr5:6090253-6090693 REVERSE LENGTH=146;AT5G18380.2 \| Symbols: \| Ribosomal protein S5 domain 2-like superfamily protein \| chr5:6090128-6090693 REVERSE LENGTH=144;AT5G18380.3 \| Symbols: \| Ribosomal protein S5 domain 2-like superfamily protein \| chr5:6090128-6090693 REVERSE LENGTH=139;AT3G04230.1 \| Symbols: \| Ribosomal protein S5 domain 2-like superfamily protein \| chr3:1113169-1113609 REVERSE LENGTH=146 | 0.37647 | 0.315527088 | 0.345998544 | 2.890185573 |
| AT4G10450 | AT4G10450.1 \| Symbols: \| Ribosomal protein L6 family \| chr4:6463201-6464458 REVERSE LENGTH=194 | | 0.345673891 | 0.345673891 | 2.8929 |
| AT3G52580 | AT3G52580.1 \| Symbols: \| Ribosomal protein S11 family protein \| chr3:19503324-19504701 FORWARD LENGTH=150 | 0.22319 | 0.467398925 | 0.345294462 | 2.896078879 |
| AT1G59870 | AT1G59870.1 \| Symbols: PEN3, PDR8, ATPDR8, ABCG36, ATABCG36 \| ABC-2 and Plant PDR ABC-type transporter family protein \| chr1:22034661-22039844 FORWARD LENGTH=1469;AT1G15210.1 \| Symbols: PDR7, ATPDR7 \| pleiotropic drug resistance 7 \| chr1:5231552-5236573 REVERSE LENGTH=1442;AT1G66950.1 \| Symbols: PDR11, ATPDR11 \| pleiotropic drug resistance 11 \| chr1:24978239-24984461 FORWARD LENGTH=1454;AT2G36380.1 \| Symbols: PDR6, ATPDR6 \| pleiotropic drug resistance 6 \| chr2:15257583-15263627 FORWARD LENGTH=1453;AT2G29940.1 \| Symbols: PDR3, ATPDR3 \| pleiotropic drug resistance 3 \| chr2:12760139-12766455 FORWARD LENGTH=1426;AT1G15520.1 \| Symbols: PDR12, ATPDR12, ABCG40, ATABCG40 \| pleiotropic drug resistance 12 \| chr1:5331993-5338175 REVERSE LENGTH=1423;AT2G26910.1 \| Symbols: PDR4, ATPDR4 \| pleiotropic drug resistance 4 \| chr2:11481623-11487874 FORWARD LENGTH=1420;AT3G16340.1 \| Symbols: ATPDR1, PDR1 \| pleiotropic drug resistance 1 \| chr3:5539897-5546263 FORWARD LENGTH=1416;AT3G16340.2 \| Symbols: PDR1 \| pleiotropic drug resistance 1 \| chr3:5539897-5546263 FORWARD LENGTH=1411;AT3G30842.1 \| Symbols: PDR10, ATPDR10 \| pleiotropic drug resistance 10 \| chr3:12593959-12600432 REVERSE LENGTH=1406 | | 0.344186687 | 0.344186687 | 2.9054 |
| AT2G42520 | AT2G42520.1 \| Symbols: \| P-loop containing nucleoside triphosphate hydrolases superfamily protein \| chr2:17705382-17708744 FORWARD LENGTH=633 | | 0.343406593 | 0.343406593 | 2.912 |
| AT4G33510 | AT4G33510.1 \| Symbols: DHS2 \| 3-deoxy-d-arabino-heptulosonate 7-phosphate synthase \| chr4:16116496-16118549 FORWARD LENGTH=507;AT4G33510.2 \| Symbols: DHS2 \| 3-deoxy-d-arabino-heptulosonate 7-phosphate synthase \| chr4:16116496-16118233 FORWARD LENGTH=432 | | 0.342853225 | 0.342853225 | 2.9167 |
| AT5G20650 | AT5G20650.1 \| Symbols: COPT5 \| copper transporter 5 \| chr5:6985481-6985921 REVERSE LENGTH=146 | 0.52193 | 0.162022035 | 0.341976017 | 2.924181664 |
| AT5G63120 | AT5G63120.2 \| Symbols: \| P-loop containing nucleoside triphosphate hydrolases superfamily protein \| chr5:25318967-25322071 REVERSE LENGTH=591;AT5G63120.1 \| Symbols: \| P-loop containing nucleoside triphosphate hydrolases superfamily protein \| chr5:25319798-25322071 REVERSE LENGTH=484 | | 0.340878102 | 0.340878102 | 2.9336 |
| AT5G62390 | AT5G62390.1 \| Symbols: ATBAG7, BAG7 \| BCL-2-associated athanogene 7 \| chr5:25052377-25054170 REVERSE LENGTH=446 | | 0.339558574 | 0.339558574 | 2.945 |
| AT1G73180 | AT1G73180.1 \| Symbols: \| Eukaryotic translation initiation factor eIF2A family protein \| chr1:27518335-27521432 FORWARD LENGTH=513;AT1G73180.2 \| Symbols: \| Eukaryotic translation initiation factor eIF2A family protein \| chr1:27518335-27521014 FORWARD LENGTH=484 | | 0.33807769 | 0.33807769 | 2.9579 |

| | | | | | |
|---|---|---|---|---|---|
| AT3G42660 | AT3G42660.1 \| Symbols: \| transducin family protein / WD-40 repeat family protein \| chr3:14751280-14755701 FORWARD LENGTH=953 | | 0.337632521 | 0.337632521 | 2.9618 |
| AT5G27120 | AT5G27120.1 \| Symbols: \| NOP56-like pre RNA processing ribonucleoprotein \| chr5:9541287-9543684 FORWARD LENGTH=533 | | 0.337416068 | 0.337416068 | 2.9637 |
| AT1G35680 | AT1G35680.1 \| Symbols: \| Ribosomal protein L21 \| chr1:13208777-13210246 FORWARD LENGTH=220 | | 0.33724538 | 0.33724538 | 2.9652 |
| AT5G60860 | AT5G60860.1 \| Symbols: AtRABA1f, RABA1f \| RAB GTPase homolog A1F \| chr5:24484750-24485565 FORWARD LENGTH=217;AT5G45750.1 \| Symbols: AtRABA1c, RABA1c \| RAB GTPase homolog A1C \| chr5:18559318-18560639 FORWARD LENGTH=216;AT3G15060.1 \| Symbols: AtRABA1g, RABA1g \| RAB GTPase homolog A1G \| chr3:5069239-5070025 FORWARD LENGTH=217;AT1G16920.1 \| Symbols: RAB11, ATRABA1B, RABA1b \| RAB GTPase homolog A1B \| chr1:5787489-5789147 REVERSE LENGTH=216;AT4G18800.1 \| Symbols: ATHSGBP, ATRAB11B, ATRABA1D, RABA1d \| RAB GTPase homolog A1D \| chr4:10320156-10321339 REVERSE LENGTH=214;AT2G33870.1 \| Symbols: ArRABA1h \| RAB GTPase homolog A1H \| chr2:14337366-14338251 REVERSE LENGTH=218;AT1G09630.1 \| Symbols: ATRAB11C, ATRABA2A, ATRAB-A2A, RAB-A2A, RAB11c \| RAB GTPase 11C \| chr1:3118350-3119571 REVERSE LENGTH=217;AT1G28550.1 \| Symbols: AtRABA1i, RABA1i \| RAB GTPase homolog A1I \| chr1:10036966-10037698 REVERSE LENGTH=218 | 0.31533 | 0.359118006 | 0.337224003 | 2.965387964 |
| AT3G15010 | AT3G15010.1 \| Symbols: \| RNA-binding (RRM/RBD/RNP motifs) family protein \| chr3:5052844-5054058 FORWARD LENGTH=404;AT3G15010.2 \| Symbols: \| RNA-binding (RRM/RBD/RNP motifs) family protein \| chr3:5052844-5054058 FORWARD LENGTH=404 | | 0.336643663 | 0.336643663 | 2.9705 |
| AT4G22140 | AT4G22140.1 \| Symbols: EBS \| PHD finger family protein / bromo-adjacent homology (BAH) domain-containing protein \| chr4:11728556-11730230 REVERSE LENGTH=234;AT4G22140.2 \| Symbols: EBS \| PHD finger family protein / bromo-adjacent homology (BAH) domain-containing protein \| chr4:11728093-11730230 REVERSE LENGTH=224;AT4G04260.1 \| Symbols: \| Bromo-adjacent homology (BAH) domain-containing protein \| chr4:2054636-2055849 FORWARD LENGTH=193 | | 0.335649314 | 0.335649314 | 2.9793 |
| AT5G15200 | AT5G15200.1 \| Symbols: \| Ribosomal protein S4 \| chr5:4935124-4936334 REVERSE LENGTH=198;AT5G15200.2 \| Symbols: \| Ribosomal protein S4 \| chr5:4935602-4936334 REVERSE LENGTH=156 | 0.17227 | 0.496179418 | 0.334224709 | 2.991999013 |
| AT1G18540 | AT1G18540.1 \| Symbols: \| Ribosomal protein L6 family protein \| chr1:6377448-6378548 REVERSE LENGTH=233 | 0.12218 | 0.544217687 | 0.333198844 | 3.001210897 |
| AT3G61650 | AT3G61650.1 \| Symbols: TUBG1 \| gamma-tubulin \| chr3:22812601-22815011 REVERSE LENGTH=474;AT5G05620.1 \| Symbols: TUBG2, ATGCP2, GCP2 \| gamma-tubulin complex protein 2 \| chr5:1679340-1681719 FORWARD LENGTH=474 | 0.21799 | 0.441403664 | 0.329696832 | 3.033089504 |
| AT1G09270 | AT1G09270.1 \| Symbols: IMPA-4 \| importin alpha isoform 4 \| chr1:2994506-2997833 FORWARD LENGTH=538;AT1G09270.2 \| Symbols: IMPA-4 \| importin alpha isoform 4 \| chr1:2994506-2997833 FORWARD LENGTH=538;AT1G09270.3 \| Symbols: IMPA-4 \| importin alpha isoform 4 \| chr1:2995162-2997833 FORWARD LENGTH=456 | | 0.329565303 | 0.329565303 | 3.0343 |
| AT5G20290 | AT5G20290.1 \| Symbols: \| Ribosomal protein S8e family protein \| chr5:6851695-6853012 REVERSE LENGTH=222;AT5G59240.1 \| Symbols: \| Ribosomal protein S8e family protein \| chr5:23902626-23903670 REVERSE LENGTH=210 | 0.17315 | 0.484284953 | 0.328717477 | 3.042126054 |
| AT1G72370 | AT1G72370.1 \| Symbols: P40, AP40, RP40, RPSAA \| 40s ribosomal protein SA \| chr1:27243148-27244842 REVERSE LENGTH=298;AT1G72370.2 \| Symbols: P40, AP40, RP40, RPSAA \| 40s ribosomal protein SA \| chr1:27243148-27244842 REVERSE LENGTH=294 | 0.20758 | 0.449256481 | 0.32841824 | 3.044897869 |

| | | | | | |
|---|---|---|---|---|---|
| AT4G22690 | AT4G22690.1 \| Symbols: CYP706A1 \| cytochrome P450, family 706, subfamily A, polypeptide 1 \| chr4:11929847-11931520 FORWARD LENGTH=557;AT4G22710.1 \| Symbols: CYP706A2 \| cytochrome P450, family 706, subfamily A, polypeptide 2 \| chr4:11935038-11936618 FORWARD LENGTH=526;AT4G12320.1 \| Symbols: CYP706A6 \| cytochrome P450, family 706, subfamily A, polypeptide 6 \| chr4:7314939-7316647 REVERSE LENGTH=518 | 0.19139 | 0.465073016 | 0.328231508 | 3.046630122 |
| AT1G75220 | AT1G75220.1 \| Symbols: \| Major facilitator superfamily protein \| chr1:28229412-28232606 REVERSE LENGTH=487 | | 0.32776139 | 0.32776139 | 3.051 |
| AT1G72440 | AT1G72440.1 \| Symbols: EDA25, SWA2 \| CCAAT-binding factor \| chr1:27268325-27273596 REVERSE LENGTH=1043 | | 0.32709669 | 0.32709669 | 3.0572 |
| AT4G31580 | AT4G31580.1 \| Symbols: SRZ-22, SRZ22, RSZP22, RSZ22, At-RSZ22 \| serine/arginine-rich 22 \| chr4:15306983-15308064 FORWARD LENGTH=200;AT4G31580.2 \| Symbols: SRZ-22, SRZ22, RSZP22, RSZ22, At-RSZ22 \| serine/arginine-rich 22 \| chr4:15306983-15308064 FORWARD LENGTH=200;AT2G24590.1 \| Symbols: RSZ22a, At-RSZ22a \| RNA recognition motif and CCHC-type zinc finger domains containing protein \| chr2:10449837-10450860 FORWARD LENGTH=196 | 0.17845 | 0.471631373 | 0.325040686 | 3.076537928 |
| AT1G56330 | AT1G56330.1 \| Symbols: SAR1, ATSAR1, ATSARA1B, ATSAR1B, SAR1B \| secretion-associated RAS 1B \| chr1:21086845-21088478 REVERSE LENGTH=193;AT3G62560.1 \| Symbols: \| Ras-related small GTP-binding family protein \| chr3:23137539-23138880 FORWARD LENGTH=193;AT4G02080.1 \| Symbols: ASAR1, ATSARA1C, ATSAR2, SAR2 \| secretion-associated RAS super family 2 \| chr4:921554-922547 FORWARD LENGTH=193;AT1G09180.1 \| Symbols: ATSAR1, ATSARA1A, SARA1A \| secretion-associated RAS super family 1 \| chr1:2965147-2965941 FORWARD LENGTH=193;AT1G02620.1 \| Symbols: \| Ras-related small GTP-binding family protein \| chr1:557092-557668 FORWARD LENGTH=122 | 0.24328 | 0.406553645 | 0.324916822 | 3.077710759 |
| AT5G06160 | AT5G06160.1 \| Symbols: ATO \| splicing factor-related \| chr5:1862623-1866298 REVERSE LENGTH=504 | | 0.324696409 | 0.324696409 | 3.0798 |
| AT4G05410 | AT4G05410.1 \| Symbols: YAO \| Transducin/WD40 repeat-like superfamily protein \| chr4:2743229-2745521 REVERSE LENGTH=504 | | 0.323708403 | 0.323708403 | 3.0892 |
| AT1G05650 | AT1G05650.1 \| Symbols: \| Pectin lyase-like superfamily protein \| chr1:1690264-1692126 REVERSE LENGTH=394;AT1G05660.1 \| Symbols: \| Pectin lyase-like superfamily protein \| chr1:1694286-1696058 REVERSE LENGTH=394 | 0.32366 | | 0.32366 | 3.089661991 |
| AT5G12250 | AT5G62690.1 \| Symbols: TUB2 \| tubulin beta chain 2 \| chr5:25181560-25183501 FORWARD LENGTH=450;AT5G62700.1 \| Symbols: TUB3 \| tubulin beta chain 3 \| chr5:25184501-25186426 FORWARD LENGTH=450;AT5G23860.1 \| Symbols: TUB8 \| tubulin beta 8 \| chr5:8042962-8044528 FORWARD LENGTH=449;AT5G23860.2 \| Symbols: TUB8 \| tubulin beta 8 \| chr5:8042962-8044528 FORWARD LENGTH=449;AT5G12250.1 \| Symbols: TUB6 \| beta-6 tubulin \| chr5:3961317-3962971 REVERSE LENGTH=449;AT2G29550.1 \| Symbols: TUB7 \| tubulin beta-7 chain \| chr2:12644258-12645932 REVERSE LENGTH=449;AT4G20890.1 \| Symbols: TUB9 \| tubulin beta-9 chain \| chr4:11182218-11183840 FORWARD LENGTH=444;AT1G20010.1 \| Symbols: TUB5 \| tubulin beta-5 chain \| chr1:6938033-6940481 REVERSE LENGTH=449;AT1G75780.1 \| Symbols: TUB1 \| tubulin beta-1 chain \| chr1:28451378-28453602 REVERSE LENGTH=447 | 0.28023 | 0.366824401 | 0.323527201 | 3.090930216 |
| AT3G14990 | AT3G14990.1 \| Symbols: \| Class I glutamine amidotransferase-like superfamily protein \| chr3:5047510-5049621 FORWARD LENGTH=392;AT3G14990.2 \| Symbols: \| Class I glutamine amidotransferase-like superfamily protein \| chr3:5047683-5049621 FORWARD LENGTH=369;AT3G14990.3 \| Symbols: \| Class I glutamine amidotransferase-like superfamily protein \| chr3:5047683-5049621 FORWARD LENGTH=369 | 0.14722 | 0.499550405 | 0.323385202 | 3.092287442 |

| | | | | | |
|---|---|---|---|---|---|
| AT2G01250 | AT2G01250.1 \| Symbols: \| Ribosomal protein L30/L7 family protein \| chr2:132943-134264 REVERSE LENGTH=242;AT2G01250.2 \| Symbols: \| Ribosomal protein L30/L7 family protein \| chr2:133038-134264 REVERSE LENGTH=184;AT5G48130.1 \| Symbols: \| Phototropic-responsive NPH3 family protein \| chr5:19516291-19518450 FORWARD LENGTH=625 | 0.06408 | 0.581192607 | 0.322636304 | 3.099465215 |
| AT5G48370 | AT5G48370.1 \| Symbols: \| Thioesterase/thiol ester dehydrase-isomerase superfamily protein \| chr5:19598666-19600512 REVERSE LENGTH=438;AT5G48370.2 \| Symbols: \| Thioesterase/thiol ester dehydrase-isomerase superfamily protein \| chr5:19598666-19600512 REVERSE LENGTH=437;AT2G30720.1 \| Symbols: \| Thioesterase/thiol ester dehydrase-isomerase superfamily protein \| chr2:13089604-13091538 REVERSE LENGTH=455 | | 0.322362271 | 0.322362271 | 3.1021 |
| AT1G15750 | AT1G15750.1 \| Symbols: WSIP1, TPL \| Transducin family protein / WD-40 repeat family protein \| chr1:5415086-5420359 REVERSE LENGTH=1131;AT1G15750.2 \| Symbols: WSIP1, TPL \| Transducin family protein / WD-40 repeat family protein \| chr1:5415086-5420359 REVERSE LENGTH=1131;AT1G15750.3 \| Symbols: WSIP1, TPL \| Transducin family protein / WD-40 repeat family protein \| chr1:5415086-5420359 REVERSE LENGTH=1131;AT1G15750.4 \| Symbols: WSIP1, TPL \| Transducin family protein / WD-40 repeat family protein \| chr1:5415086-5420359 REVERSE LENGTH=1131;AT3G16830.1 \| Symbols: TPR2 \| TOPLESS-related 2 \| chr3:5731709-5737531 FORWARD LENGTH=1131;AT1G80490.2 \| Symbols: TPR1 \| TOPLESS-related 1 \| chr1:30261094-30266446 REVERSE LENGTH=1120;AT1G80490.1 \| Symbols: TPR1 \| TOPLESS-related 1 \| chr1:30261094-30266446 REVERSE LENGTH=1119;AT3G15880.2 \| Symbols: WSIP2, TPR4 \| WUS-interacting protein 2 \| chr3:5364454-5371869 REVERSE LENGTH=1137;AT3G15880.1 \| Symbols: WSIP2, TPR4 \| WUS-interacting protein 2 \| chr3:5364792-5371869 REVERSE LENGTH=1135;AT5G27030.2 \| Symbols: TPR3 \| TOPLESS-related 3 \| chr5:9508913-9515263 REVERSE LENGTH=1134;AT3G15880.3 \| Symbols: WSIP2 \| WUS-interacting protein 2 \| chr3:5364792-5371869 REVERSE LENGTH=1125;AT5G27030.1 \| Symbols: TPR3 \| TOPLESS-related 3 \| chr5:9508913-9515263 REVERSE LENGTH=1108 | | 0.319060685 | 0.319060685 | 3.1342 |
| AT5G41520 | AT5G41520.1 \| Symbols: \| RNA binding Plectin/S10 domain-containing protein \| chr5:16609377-16610583 REVERSE LENGTH=180;AT5G41520.2 \| Symbols: \| RNA binding Plectin/S10 domain-containing protein \| chr5:16609377-16610203 REVERSE LENGTH=134 | 0.34792 | 0.288184438 | 0.318052219 | 3.144137787 |
| AT3G26618 | AT3G26618.1 \| Symbols: ERF1-3 \| eukaryotic release factor 1-3 \| chr3:9788854-9790161 FORWARD LENGTH=435;AT1G12920.1 \| Symbols: ERF1-2 \| eukaryotic release factor 1-2 \| chr1:4396555-4397859 REVERSE LENGTH=434 | | 0.317158262 | 0.317158262 | 3.153 |
| AT1G53500 | AT1G53500.1 \| Symbols: MUM4, RHM2, ATRHM2, ATMUM4 \| NAD-dependent epimerase/dehydratase family protein \| chr1:19967157-19969239 REVERSE LENGTH=667;AT3G14790.1 \| Symbols: RHM3, ATRHM3 \| rhamnose biosynthesis 3 \| chr3:4964791-4966875 FORWARD LENGTH=664;AT1G78570.1 \| Symbols: RHM1, ROL1, ATRHM1 \| rhamnose biosynthesis 1 \| chr1:29550110-29552207 FORWARD LENGTH=669 | | 0.31702755 | 0.31702755 | 3.1543 |
| AT1G74050 | AT1G74050.1 \| Symbols: \| Ribosomal protein L6 family protein \| chr1:27847256-27848680 REVERSE LENGTH=233;AT1G74060.1 \| Symbols: \| Ribosomal protein L6 family protein \| chr1:27850033-27851299 REVERSE LENGTH=233 | 0.12021 | 0.510490581 | 0.315350291 | 3.171076829 |
| AT2G20450 | AT2G20450.1 \| Symbols: \| Ribosomal protein L14 \| chr2:8813923-8815071 FORWARD LENGTH=134 | 0.4786 | 0.151405039 | 0.315002519 | 3.174577784 |
| AT2G22780 | AT2G22780.1 \| Symbols: PMDH1 \| peroxisomal NAD-malate dehydrogenase 1 \| chr2:9689995-9691923 REVERSE LENGTH=354 | | 0.312871535 | 0.312871535 | 3.1962 |
| AT3G49240 | AT3G49240.1 \| Symbols: emb1796 \| Pentatricopeptide repeat (PPR) superfamily protein \| chr3:18256086-18257975 FORWARD LENGTH=629 | | 0.312627005 | 0.312627005 | 3.1987 |

| | | | | | |
|---|---|---|---|---|---|
| AT1G77180 | AT1G77180.1 \| Symbols: SKIP \| chromatin protein family \| chr1:28999791-29001632 REVERSE LENGTH=613;AT1G77180.2 \| Symbols: SKIP \| chromatin protein family \| chr1:28999791-29001632 REVERSE LENGTH=613;AT1G77180.3 \| Symbols: SKIP \| chromatin protein family \| chr1:28999791-29001632 REVERSE LENGTH=511 | | 0.312314563 | 0.312314563 | 3.2019 |
| AT3G42050 | AT3G42050.1 \| Symbols: \| vacuolar ATP synthase subunit H family protein \| chr3:14228846-14232228 REVERSE LENGTH=441 | 0.12019 | 0.49960032 | 0.30989516 | 3.226897769 |
| AT5G22360 | AT5G22360.1 \| Symbols: ATVAMP714, VAMP714 \| vesicle-associated membrane protein 714 \| chr5:7404379-7405654 REVERSE LENGTH=221 | | 0.309741366 | 0.309741366 | 3.2285 |
| AT3G22660 | AT3G22660.1 \| Symbols: \| rRNA processing protein-related \| chr3:8016237-8017118 REVERSE LENGTH=293 | 0.14968 | 0.469043152 | 0.309361576 | 3.232463491 |
| AT5G11240 | AT5G11240.1 \| Symbols: \| transducin family protein / WD-40 repeat family protein \| chr5:3582949-3586782 FORWARD LENGTH=615 | | 0.309300671 | 0.309300671 | 3.2331 |
| AT5G19390 | AT5G19390.1 \| Symbols: \| Rho GTPase activation protein (RhoGAP) with PH domain \| chr5:6531906-6538206 FORWARD LENGTH=870;AT5G19390.2 \| Symbols: \| Rho GTPase activation protein (RhoGAP) with PH domain \| chr5:6531906-6538206 FORWARD LENGTH=870;AT5G19390.3 \| Symbols: \| Rho GTPase activation protein (RhoGAP) with PH domain \| chr5:6531906-6538069 FORWARD LENGTH=822;AT5G19390.4 \| Symbols: \| Rho GTPase activation protein (RhoGAP) with PH domain \| chr5:6531906-6537143 FORWARD LENGTH=714 | | 0.307607124 | 0.307607124 | 3.2509 |
| AT1G02690 | AT1G02690.2 \| Symbols: IMPA-6 \| importin alpha isoform 6 \| chr1:584397-587036 FORWARD LENGTH=539;AT1G02690.1 \| Symbols: IMPA-6 \| importin alpha isoform 6 \| chr1:584397-587036 FORWARD LENGTH=538 | 0.0044736 | 0.609161793 | 0.306817697 | 3.259264413 |
| AT1G54270 | AT1G54270.1 \| Symbols: EIF4A-2 \| eif4a-2 \| chr1:20260495-20262018 FORWARD LENGTH=412;AT1G54270.2 \| Symbols: EIF4A-2 \| eif4a-2 \| chr1:20260495-20262018 FORWARD LENGTH=407 | | 0.306795521 | 0.306795521 | 3.2595 |
| AT5G39950 | AT5G39950.1 \| Symbols: ATTRX2, ATH2, ATTRXH2, TRXH2, TRX2 \| thioredoxin 2 \| chr5:15990885-15991881 REVERSE LENGTH=133 | | 0.305231671 | 0.305231671 | 3.2762 |
| AT4G23850 | AT4G23850.1 \| Symbols: LACS4 \| AMP-dependent synthetase and ligase family protein \| chr4:12403720-12408263 REVERSE LENGTH=666 | 0.11407 | 0.494413132 | 0.304241566 | 3.286861864 |
| AT2G30970 | AT2G30970.1 \| Symbols: ASP1 \| aspartate aminotransferase 1 \| chr2:13179012-13181686 FORWARD LENGTH=430;AT2G30970.2 \| Symbols: ASP1 \| aspartate aminotransferase 1 \| chr2:13179012-13181686 FORWARD LENGTH=430 | 0.090172 | 0.516822575 | 0.303497287 | 3.294922365 |
| AT2G20230 | AT2G20230.1 \| Symbols: \| Tetraspanin family protein \| chr2:8725762-8727388 FORWARD LENGTH=270 | | 0.30284676 | 0.30284676 | 3.302 |
| AT3G03920 | AT3G03920.1 \| Symbols: \| H/ACA ribonucleoprotein complex, subunit Gar1/Naf1 protein \| chr3:1009123-1010379 REVERSE LENGTH=202;AT5G18180.1 \| Symbols: \| H/ACA ribonucleoprotein complex, subunit Gar1/Naf1 protein \| chr5:6008561-6009605 REVERSE LENGTH=189 | 0.050747 | 0.554354454 | 0.302550727 | 3.30523086 |
| AT1G02780 | AT1G02780.1 \| Symbols: emb2386 \| Ribosomal protein L19e family protein \| chr1:608120-609391 REVERSE LENGTH=214;AT4G02230.1 \| Symbols: \| Ribosomal protein L19e family protein \| chr4:979391-980640 REVERSE LENGTH=208;AT3G16780.1 \| Symbols: \| Ribosomal protein L19e family protein \| chr3:5708982-5710249 FORWARD LENGTH=209 | 0.11069 | 0.493510339 | 0.30210017 | 3.31016034 |
| ATCG00780 | ATCG00780.1 \| Symbols: RPL14 \| ribosomal protein L14 \| chrC:80696-81064 REVERSE LENGTH=122 | | 0.301905021 | 0.301905021 | 3.3123 |
| AT4G02840 | AT4G02840.2 \| Symbols: \| Small nuclear ribonucleoprotein family protein \| chr4:1264726-1266253 FORWARD LENGTH=126;AT4G02840.1 \| Symbols: \| Small nuclear ribonucleoprotein family protein \| chr4:1264726-1266253 FORWARD LENGTH=116;AT3G07590.1 \| Symbols: \| Small nuclear ribonucleoprotein family protein \| chr3:2423146-2423958 FORWARD LENGTH=114;AT3G07590.2 \| Symbols: \| Small nuclear ribonucleoprotein family protein \| chr3:2423146-2423958 FORWARD LENGTH=114 | | 0.301486328 | 0.301486328 | 3.3169 |
| AT5G49160 | AT5G49160.1 \| Symbols: MET1, MET2, METI, DDM2, DMT01, DMT1 \| methyltransferase 1 \| chr5:19932501-19938186 FORWARD LENGTH=1534 | | 0.301213892 | 0.301213892 | 3.3199 |

| | | | | | |
|---|---|---|---|---|---|
| AT4G21150 | AT4G21150.1 \| Symbols: HAP6 \| ribophorin II (RPN2) family protein \| chr4:11278646-11283599 FORWARD LENGTH=691;AT4G21150.3 \| Symbols: HAP6 \| ribophorin II (RPN2) family protein \| chr4:11278646-11283599 FORWARD LENGTH=691;AT4G21150.2 \| Symbols: HAP6 \| ribophorin II (RPN2) family protein \| chr4:11278649-11283599 FORWARD LENGTH=678 | 0.11485 | 0.486263068 | 0.300556534 | 3.327161071 |
| AT3G22520 | AT3G22520.1 \| Symbols: \| unknown protein; INVOLVED IN: biological_process unknown; LOCATED IN: chloroplast stroma, chloroplast, chloroplast envelope; EXPRESSED IN: 24 plant structures; EXPRESSED DURING: 13 growth stages; BEST Arabidopsis thaliana protein match is: unknown protein (TAIR:AT4G14840.1); Has 717 Blast hits to 703 proteins in 179 species: Archae - 14; Bacteria - 134; Metazoa - 141; Fungi - 74; Plants - 209; Viruses - 0; Other Eukaryotes - 145 (source: NCBI BLink). \| chr3:7974984-7977406 FORWARD LENGTH=600 | | 0.299742222 | 0.299742222 | 3.3362 |
| AT4G12320 | AT4G12320.1 \| Symbols: CYP706A6 \| cytochrome P450, family 706, subfamily A, polypeptide 6 \| chr4:7314939-7316647 REVERSE LENGTH=518 | | 0.299571613 | 0.299571613 | 3.3381 |
| AT2G46280 | AT2G46280.1 \| Symbols: TRIP-1, TIF3I1 \| TGF-beta receptor interacting protein 1 \| chr2:19003656-19005393 REVERSE LENGTH=328;AT2G46280.2 \| Symbols: TRIP-1, TIF3I1 \| TGF-beta receptor interacting protein 1 \| chr2:19003656-19005393 REVERSE LENGTH=328;AT2G46280.3 \| Symbols: TRIP-1, TIF3I1 \| TGF-beta receptor interacting protein 1 \| chr2:19004179-19005393 REVERSE LENGTH=254 | 0.16281 | 0.435502134 | 0.299156067 | 3.34273682 |
| AT4G12650 | AT4G12650.1 \| Symbols: \| Endomembrane protein 70 protein family \| chr4:7468207-7470165 REVERSE LENGTH=652 | | 0.297619048 | 0.297619048 | 3.36 |
| AT3G49910 | AT3G49910.1 \| Symbols: \| Translation protein SH3-like family protein \| chr3:18504311-18504751 FORWARD LENGTH=146 | 0.11858 | 0.476009139 | 0.29729457 | 3.363667224 |
| AT4G39080 | AT4G39080.1 \| Symbols: VHA-A3 \| vacuolar proton ATPase A3 \| chr4:18209513-18214752 FORWARD LENGTH=821 | 0.38732 | 0.20718089 | 0.297250445 | 3.364166538 |
| AT3G60770 | AT3G60770.1 \| Symbols: \| Ribosomal protein S13/S15 \| chr3:22460525-22461656 REVERSE LENGTH=151;AT4G00100.1 \| Symbols: ATRPS13A, RPS13, PFL2, RPS13A \| ribosomal protein S13A \| chr4:37172-38123 FORWARD LENGTH=151 | 0.1275 | 0.466722673 | 0.297111337 | 3.365741648 |
| AT1G07660 | AT1G07660.1 \| Symbols: \| Histone superfamily protein \| chr1:2369212-2369523 FORWARD LENGTH=103;AT1G07820.1 \| Symbols: \| Histone superfamily protein \| chr1:2421435-2421746 REVERSE LENGTH=103;AT1G07820.2 \| Symbols: \| Histone superfamily protein \| chr1:2421435-2421746 REVERSE LENGTH=103;AT2G28740.1 \| Symbols: HIS4 \| histone H4 \| chr2:12329643-12329954 REVERSE LENGTH=103;AT3G45930.1 \| Symbols: \| Histone superfamily protein \| chr3:16883390-16883701 FORWARD LENGTH=103;AT3G46320.1 \| Symbols: \| Histone superfamily protein \| chr3:17020043-17020354 REVERSE LENGTH=103;AT3G53730.1 \| Symbols: \| Histone superfamily protein \| chr3:19912659-19912970 REVERSE LENGTH=103;AT5G59690.1 \| Symbols: \| Histone superfamily protein \| chr5:24051649-24051960 FORWARD LENGTH=103;AT5G59970.1 \| Symbols: \| Histone superfamily protein \| chr5:24146352-24146663 REVERSE LENGTH=103;AT1G07660.2 \| Symbols: \| Histone superfamily protein \| chr1:2369212-2369523 FORWARD LENGTH=86 | 0.23405 | 0.359634611 | 0.296842306 | 3.368792052 |
| AT3G22330 | AT3G22330.1 \| Symbols: PMH2, ATRH53 \| putative mitochondrial RNA helicase 2 \| chr3:7892641-7895145 FORWARD LENGTH=616;AT3G22310.1 \| Symbols: PMH1, ATRH9 \| putative mitochondrial RNA helicase 1 \| chr3:7887382-7889806 FORWARD LENGTH=610 | | 0.294134949 | 0.294134949 | 3.3998 |
| AT4G31700 | AT4G31700.1 \| Symbols: RPS6, RPS6A \| ribosomal protein S6 \| chr4:15346306-15347714 REVERSE LENGTH=250;AT5G10360.2 \| Symbols: EMB3010 \| Ribosomal protein S6e \| chr5:3258734-3260142 REVERSE LENGTH=197;AT4G31700.2 \| Symbols: RPS6, RPS6A \| ribosomal protein S6 \| chr4:15346306-15347075 REVERSE LENGTH=188 | 0.097561 | 0.490604916 | 0.294082958 | 3.400401054 |

| | | | | | |
|---|---|---|---|---|---|
| AT2G02040 | AT2G02040.1 \| Symbols: ATPTR2-B, NTR1, PTR2-B, PTR2, ATPTR2 \| peptide transporter 2 \| chr2:487542-489707 FORWARD LENGTH=585;AT5G01180.1 \| Symbols: PTR5, ATPTR5 \| peptide transporter 5 \| chr5:61257-63240 REVERSE LENGTH=570 | | 0.293979304 | 0.293979304 | 3.4016 |
| AT5G27770 | AT5G27770.1 \| Symbols: \| Ribosomal L22e protein family \| chr5:9836166-9837113 FORWARD LENGTH=124 | 0.42925 | 0.158596736 | 0.293923368 | 3.40224735 |
| AT3G10950 | AT3G10950.1 \| Symbols: \| Zinc-binding ribosomal protein family protein \| chr3:3423893-3424566 FORWARD LENGTH=92;AT3G60245.1 \| Symbols: \| Zinc-binding ribosomal protein family protein \| chr3:22268803-22269750 FORWARD LENGTH=92 | 0.29301 | | 0.29301 | 3.412852804 |
| AT1G32410 | AT1G32410.1 \| Symbols: \| Vacuolar protein sorting 55 (VPS55) family protein \| chr1:11694398-11695297 FORWARD LENGTH=140;AT1G32410.2 \| Symbols: \| Vacuolar protein sorting 55 (VPS55) family protein \| chr1:11694398-11695297 FORWARD LENGTH=140;AT1G32410.3 \| Symbols: \| Vacuolar protein sorting 55 (VPS55) family protein \| chr1:11694398-11695297 FORWARD LENGTH=140;AT1G32410.4 \| Symbols: \| Vacuolar protein sorting 55 (VPS55) family protein \| chr1:11694398-11695297 FORWARD LENGTH=140;AT1G32410.5 \| Symbols: \| Vacuolar protein sorting 55 (VPS55) family protein \| chr1:11694398-11695297 FORWARD LENGTH=140 | | 0.291919664 | 0.291919664 | 3.4256 |
| AT3G04920 | AT3G04920.1 \| Symbols: \| Ribosomal protein S24e family protein \| chr3:1360989-1362065 FORWARD LENGTH=133 | 0.29018 | 0.29156219 | 0.290871095 | 3.437949032 |
| AT1G78900 | AT1G78900.1 \| Symbols: VHA-A \| vacuolar ATP synthase subunit A \| chr1:29660463-29664575 FORWARD LENGTH=623;AT1G78900.2 \| Symbols: VHA-A \| vacuolar ATP synthase subunit A \| chr1:29660463-29664575 FORWARD LENGTH=623 | 0.12427 | 0.456475099 | 0.29037255 | 3.443851704 |
| AT1G60200 | AT1G60200.1 \| Symbols: \| splicing factor PWI domain-containing protein / RNA recognition motif (RRM)-containing protein \| chr1:22200882-22205241 REVERSE LENGTH=899 | | 0.29010734 | 0.29010734 | 3.447 |
| AT5G49930 | AT5G49930.1 \| Symbols: emb1441 \| zinc knuckle (CCHC-type) family protein \| chr5:20308260-20312736 REVERSE LENGTH=1080 | | 0.289335108 | 0.289335108 | 3.4562 |
| AT1G62020 | AT1G62020.1 \| Symbols: \| Coatomer, alpha subunit \| chr1:22919814-22923728 FORWARD LENGTH=1216 | | 0.289134332 | 0.289134332 | 3.4586 |
| AT2G31060 | AT2G31060.3 \| Symbols: \| elongation factor family protein \| chr2:13213496-13218544 REVERSE LENGTH=671;AT2G31060.2 \| Symbols: \| elongation factor family protein \| chr2:13213496-13218544 REVERSE LENGTH=667;AT2G31060.1 \| Symbols: \| elongation factor family protein \| chr2:13213496-13217686 REVERSE LENGTH=527 | | 0.287232515 | 0.287232515 | 3.4815 |
| AT5G46280 | AT5G46280.1 \| Symbols: MCM3 \| Minichromosome maintenance (MCM2/3/5) family protein \| chr5:18769902-18773606 REVERSE LENGTH=776 | | 0.286631507 | 0.286631507 | 3.4888 |
| AT5G58070 | AT5G58070.1 \| Symbols: ATTIL, TIL \| temperature-induced lipocalin \| chr5:23500512-23501156 REVERSE LENGTH=186 | 0.38788 | 0.185130332 | 0.286505166 | 3.490338462 |
| AT4G20850 | AT4G20850.1 \| Symbols: TPP2 \| tripeptidyl peptidase ii \| chr4:11160935-11169889 REVERSE LENGTH=1380 | 0.14891 | 0.422707867 | 0.285808933 | 3.498840951 |
| AT1G63660 | AT1G63660.1 \| Symbols: \| GMP synthase (glutamine-hydrolyzing), putative / glutamine amidotransferase, putative \| chr1:23604127-23607080 REVERSE LENGTH=534;AT1G63660.2 \| Symbols: \| GMP synthase (glutamine-hydrolyzing), putative / glutamine amidotransferase, putative \| chr1:23604874-23607080 REVERSE LENGTH=434 | | 0.284786695 | 0.284786695 | 3.5114 |
| AT5G16930 | AT5G16930.1 \| Symbols: \| AAA-type ATPase family protein \| chr5:5568578-5571565 FORWARD LENGTH=644 | 0.28415 | | 0.28415 | 3.519267992 |

| | | | | | |
|---|---|---|---|---|---|
| AT2G37860 | AT2G37860.3 \| Symbols: LCD1 \| Protein of unknown function (DUF3411) \| chr2:15856952-15858793 FORWARD LENGTH=432;AT2G37860.1 \| Symbols: LCD1 \| Protein of unknown function (DUF3411) \| chr2:15856952-15858351 FORWARD LENGTH=347;AT2G37860.2 \| Symbols: LCD1 \| Protein of unknown function (DUF3411) \| chr2:15856952-15858351 FORWARD LENGTH=347;AT5G22790.1 \| Symbols: RER1 \| reticulata-related 1 \| chr5:7599395-7601573 REVERSE LENGTH=433 | | 0.283840936 | 0.283840936 | 3.5231 |
| AT4G39460 | AT4G39460.1 \| Symbols: SAMC1, SAMT1 \| S-adenosylmethionine carrier 1 \| chr4:18356093-18358596 REVERSE LENGTH=325;AT4G39460.2 \| Symbols: SAMC1 \| S-adenosylmethionine carrier 1 \| chr4:18356093-18358596 REVERSE LENGTH=325 | | 0.28319787 | 0.28319787 | 3.5311 |
| AT5G60980 | AT5G60980.2 \| Symbols: \| Nuclear transport factor 2 (NTF2) family protein with RNA binding (RRM-RBD-RNP motifs) domain \| chr5:24543988-24546027 FORWARD LENGTH=460;AT5G60980.1 \| Symbols: \| Nuclear transport factor 2 (NTF2) family protein with RNA binding (RRM-RBD-RNP motifs) domain \| chr5:24543988-24546027 FORWARD LENGTH=459 | 0.051985 | 0.513927433 | 0.282956217 | 3.534115672 |
| AT1G80410 | AT1G80410.2 \| Symbols: EMB2753 \| tetratricopeptide repeat (TPR)-containing protein \| chr1:30227779-30234832 REVERSE LENGTH=911;AT1G80410.1 \| Symbols: EMB2753 \| tetratricopeptide repeat (TPR)-containing protein \| chr1:30227963-30234832 REVERSE LENGTH=897 | 0.11463 | 0.450734698 | 0.282682349 | 3.537539589 |
| AT1G56110 | AT1G56110.1 \| Symbols: NOP56 \| homolog of nucleolar protein NOP56 \| chr1:20984544-20986893 REVERSE LENGTH=522;AT3G12860.1 \| Symbols: \| NOP56-like pre RNA processing ribonucleoprotein \| chr3:4091678-4093921 FORWARD LENGTH=499 | | 0.281618745 | 0.281618745 | 3.5509 |
| ATCG00480 | ATCG00480.1 \| Symbols: ATPB, PB \| ATP synthase subunit beta \| chrC:52660-54156 REVERSE LENGTH=498 | 0.11578 | 0.447087227 | 0.281433613 | 3.553235834 |
| AT4G29130 | AT4G29130.1 \| Symbols: ATHXK1, GIN2, HXK1 \| hexokinase 1 \| chr4:14352338-14354865 REVERSE LENGTH=496 | | 0.280717514 | 0.280717514 | 3.5623 |
| AT2G34660 | AT2G34660.1 \| Symbols: ATMRP2, EST4, MRP2, ABCC2, AtABCC2 \| multidrug resistance-associated protein 2 \| chr2:14603267-14612387 FORWARD LENGTH=1623;AT2G34660.2 \| Symbols: MRP2, ABCC2, AtABCC2 \| multidrug resistance-associated protein 2 \| chr2:14603267-14612387 FORWARD LENGTH=1623 | | 0.27990819 | 0.27990819 | 3.5726 |
| AT2G19740 | AT2G19740.1 \| Symbols: \| Ribosomal protein L31e family protein \| chr2:8513577-8514346 FORWARD LENGTH=119;AT4G26230.1 \| Symbols: \| Ribosomal protein L31e family protein \| chr4:13286026-13286553 FORWARD LENGTH=119;AT5G56710.1 \| Symbols: \| Ribosomal protein L31e family protein \| chr5:22944003-22944767 REVERSE LENGTH=119;AT5G56710.2 \| Symbols: \| Ribosomal protein L31e family protein \| chr5:22944003-22944767 REVERSE LENGTH=85 | | 0.279759407 | 0.279759407 | 3.5745 |
| AT2G21160 | AT2G21160.1 \| Symbols: \| Translocon-associated protein (TRAP), alpha subunit \| chr2:9068428-9070207 FORWARD LENGTH=258;AT2G21160.2 \| Symbols: \| Translocon-associated protein (TRAP), alpha subunit \| chr2:9068428-9070310 FORWARD LENGTH=188 | | 0.279041214 | 0.279041214 | 3.5837 |
| AT1G80930 | AT1G80930.1 \| Symbols: \| MIF4G domain-containing protein / MA3 domain-containing protein \| chr1:30405774-30409499 REVERSE LENGTH=900 | | 0.278644672 | 0.278644672 | 3.5888 |
| AT3G52200 | AT3G52200.2 \| Symbols: LTA3 \| Dihydrolipoamide acetyltransferase, long form protein \| chr3:19360317-19366091 FORWARD LENGTH=713;AT3G52200.1 \| Symbols: LTA3 \| Dihydrolipoamide acetyltransferase, long form protein \| chr3:19360317-19366091 FORWARD LENGTH=637 | | 0.278148643 | 0.278148643 | 3.5952 |
| AT4G23630 | AT4G23630.1 \| Symbols: BTI1, RTNLB1 \| VIRB2-interacting protein 1 \| chr4:12318070-12319574 FORWARD LENGTH=275;AT4G11220.1 \| Symbols: BTI2, RTNLB2 \| VIRB2-interacting protein 2 \| chr4:6838176-6839578 REVERSE LENGTH=271 | 0.27704 | | 0.27704 | 3.609587063 |

| | | | | | |
|---|---|---|---|---|---|
| AT2G33340 | AT2G33340.1 \| Symbols: MAC3B \| MOS4-associated complex 3B \| chr2:14126584-14131000 REVERSE LENGTH=525;AT2G33340.2 \| Symbols: MAC3B \| MOS4-associated complex 3B \| chr2:14126584-14131000 REVERSE LENGTH=525;AT1G04510.1 \| Symbols: MAC3A \| MOS4-associated complex 3A \| chr1:1226749-1230592 FORWARD LENGTH=523;AT1G04510.2 \| Symbols: MAC3A \| MOS4-associated complex 3A \| chr1:1226749-1230592 FORWARD LENGTH=523;AT2G33340.3 \| Symbols: MAC3B \| MOS4-associated complex 3B \| chr2:14126703-14131000 REVERSE LENGTH=485 | | 0.276265989 | 0.276265989 | 3.6197 |
| AT4G35300 | AT4G35300.1 \| Symbols: TMT2 \| tonoplast monosaccharide transporter2 \| chr4:16796432-16799071 REVERSE LENGTH=739;AT4G35300.4 \| Symbols: TMT2 \| tonoplast monosaccharide transporter2 \| chr4:16796432-16799071 REVERSE LENGTH=739;AT4G35300.2 \| Symbols: TMT2 \| tonoplast monosaccharide transporter2 \| chr4:16796432-16799071 REVERSE LENGTH=729;AT4G35300.3 \| Symbols: \| tonoplast monosaccharide transporter2 \| chr4:16796432-16799071 REVERSE LENGTH=729;AT4G35300.5 \| Symbols: TMT2 \| tonoplast monosaccharide transporter2 \| chr4:16796432-16798332 REVERSE LENGTH=542 | | 0.276220203 | 0.276220203 | 3.6203 |
| AT4G00430 | AT4G00430.1 \| Symbols: TMP-C, PIP1;4, PIP1E \| plasma membrane intrinsic protein 1;4 \| chr4:186143-187531 REVERSE LENGTH=287;AT4G23400.1 \| Symbols: PIP1D, PIP1;5 \| plasma membrane intrinsic protein 1;5 \| chr4:12220792-12222155 FORWARD LENGTH=287;AT1G01620.1 \| Symbols: PIP1C, TMP-B, PIP1;3 \| plasma membrane intrinsic protein 1C \| chr1:225986-227176 REVERSE LENGTH=286;AT3G54820.1 \| Symbols: PIP2D, PIP2;5 \| plasma membrane intrinsic protein 2;5 \| chr3:20302117-20303738 FORWARD LENGTH=286;AT3G61430.1 \| Symbols: PIP1A, ATPIP1, PIP1, PIP1;1 \| plasma membrane intrinsic protein 1A \| chr3:22733657-22735113 FORWARD LENGTH=286;AT3G61430.2 \| Symbols: PIP1A, ATPIP1, PIP1, PIP1;1 \| plasma membrane intrinsic protein 1A \| chr3:22733657-22735113 FORWARD LENGTH=286;AT1G01620.2 \| Symbols: PIP1C, TMP-B, PIP1;3 \| plasma membrane intrinsic protein 1C \| chr1:225986-226960 REVERSE LENGTH=214 | | 0.275504863 | 0.275504863 | 3.6297 |
| AT1G58380 | AT1G58380.1 \| Symbols: XW6 \| Ribosomal protein S5 family protein \| chr1:21689115-21690085 FORWARD LENGTH=284;AT1G58684.1 \| Symbols: \| Ribosomal protein S5 family protein \| chr1:21770021-21771217 REVERSE LENGTH=284;AT1G58983.1 \| Symbols: \| Ribosomal protein S5 family protein \| chr1:21806279-21807475 REVERSE LENGTH=284;AT1G59359.1 \| Symbols: \| Ribosomal protein S5 family protein \| chr1:21842537-21843733 REVERSE LENGTH=284;AT3G57490.1 \| Symbols: \| Ribosomal protein S5 family protein \| chr3:21279824-21280887 REVERSE LENGTH=276 | 0.10398 | 0.444424692 | 0.274202346 | 3.646941808 |
| AT4G39880 | AT4G39880.1 \| Symbols: \| Ribosomal protein L23/L15e family protein \| chr4:18504601-18505137 FORWARD LENGTH=178 | | 0.273980109 | 0.273980109 | 3.6499 |
| AT3G45970 | AT3G45970.1 \| Symbols: ATEXLA1, EXPL1, ATEXPL1, ATHEXP BETA 2.1, EXLA1 \| expansin-like A1 \| chr3:16896238-16897189 FORWARD LENGTH=265 | | 0.273515495 | 0.273515495 | 3.6561 |
| AT4G20360 | AT4G20360.1 \| Symbols: ATRAB8D, ATRABE1B, RABE1b \| RAB GTPase homolog E1B \| chr4:10990036-10991466 FORWARD LENGTH=476 | 0.14784 | 0.398660501 | 0.27325025 | 3.65964898 |

| | | | | | |
|---|---|---|---|---|---|
| AT4G40030 | AT4G40030.2 \| Symbols: \| Histone superfamily protein \| chr4:18555840-18556827 REVERSE LENGTH=164;AT4G40030.1 \| Symbols: \| Histone superfamily protein \| chr4:18555840-18556417 REVERSE LENGTH=136;AT4G40030.3 \| Symbols: \| Histone superfamily protein \| chr4:18555840-18556417 REVERSE LENGTH=136;AT4G40040.1 \| Symbols: \| Histone superfamily protein \| chr4:18557539-18558236 REVERSE LENGTH=136;AT4G40040.2 \| Symbols: \| Histone superfamily protein \| chr4:18557539-18558236 REVERSE LENGTH=136;AT5G10980.1 \| Symbols: \| Histone superfamily protein \| chr5:3472591-3473349 REVERSE LENGTH=136;AT5G65350.1 \| Symbols: HTR11 \| histone 3 11 \| chr5:26119406-26119825 REVERSE LENGTH=139;AT1G09200.1 \| Symbols: \| Histone superfamily protein \| chr1:2971722-2972132 REVERSE LENGTH=136;AT1G13370.1 \| Symbols: \| Histone superfamily protein \| chr1:4587926-4588522 REVERSE LENGTH=136;AT3G27360.1 \| Symbols: \| Histone superfamily protein \| chr3:10129459-10129869 REVERSE LENGTH=136;AT5G10390.1 \| Symbols: \| Histone superfamily protein \| chr5:3269065-3269475 REVERSE LENGTH=136;AT5G10400.1 \| Symbols: \| Histone superfamily protein \| chr5:3270478-3270888 REVERSE LENGTH=136;AT5G65360.1 \| Symbols: \| Histone superfamily protein \| chr5:26120099-26120509 REVERSE LENGTH=136;AT1G75600.1 \| Symbols: \| Histone superfamily protein \| chr1:28390753-28391461 FORWARD LENGTH=136 | 0.20066 | 0.34524426 | 0.27295213 | 3.663646074 |
| AT5G02500 | AT5G02500.1 \| Symbols: HSC70-1, HSP70-1, AT-HSC70-1, HSC70 \| heat shock cognate protein 70-1 \| chr5:554055-556334 REVERSE LENGTH=651;AT3G09440.1 \| Symbols: \| Heat shock protein 70 (Hsp 70) family protein \| chr3:2903434-2905632 REVERSE LENGTH=649;AT3G09440.2 \| Symbols: \| Heat shock protein 70 (Hsp 70) family protein \| chr3:2903434-2905632 REVERSE LENGTH=649;AT5G02500.2 \| Symbols: HSC70-1, HSP70-1, AT-HSC70-1, HSC70 \| heat shock cognate protein 70-1 \| chr5:554055-556334 REVERSE LENGTH=521;AT5G02490.1 \| Symbols: \| Heat shock protein 70 (Hsp 70) family protein \| chr5:550296-552565 REVERSE LENGTH=653;AT3G12580.1 \| Symbols: HSP70, ATHSP70 \| heat shock protein 70 \| chr3:3991487-3993689 REVERSE LENGTH=650;AT1G56410.1 \| Symbols: ERD2, HSP70T-1 \| heat shock protein 70 (Hsp 70) family protein \| chr1:21117147-21119241 FORWARD LENGTH=617;AT1G16030.1 \| Symbols: Hsp70b \| heat shock protein 70B \| chr1:5502386-5504326 REVERSE LENGTH=646 | 0.2112 | 0.334135258 | 0.272667629 | 3.667468719 |
| AT1G49300 | AT1G49300.1 \| Symbols: ATRAB7, ATRABG3E, RABG3E \| RAB GTPase homolog G3E \| chr1:18234842-18236968 FORWARD LENGTH=206;AT1G49300.2 \| Symbols: ATRAB7, ATRABG3E, RABG3E \| RAB GTPase homolog G3E \| chr1:18234842-18236968 FORWARD LENGTH=206;AT1G52280.1 \| Symbols: AtRABG3d, RABG3d \| RAB GTPase homolog G3D \| chr1:19468150-19469449 REVERSE LENGTH=206;AT3G16100.1 \| Symbols: ATRABG3C, ATRAB7D, RABG3c \| RAB GTPase homolog G3C \| chr3:5459270-5460556 FORWARD LENGTH=206;AT3G18820.1 \| Symbols: ATRABG3F, ATRAB7B, RAB71, RABG3F, RAB7B \| RAB GTPase homolog G3F \| chr3:6484266-6486005 FORWARD LENGTH=206;AT1G22740.1 \| Symbols: RAB7, ATRABG3B, RAB75, RABG3B \| RAB GTPase homolog G3B \| chr1:8049247-8050494 FORWARD LENGTH=203 | 0.27246 | | 0.27246 | 3.670263525 |
| AT3G48930 | AT3G48930.1 \| Symbols: EMB1080 \| Nucleic acid-binding, OB-fold-like protein \| chr3:18141017-18142189 REVERSE LENGTH=160;AT4G30800.1 \| Symbols: \| Nucleic acid-binding, OB-fold-like protein \| chr4:15001216-15002392 FORWARD LENGTH=159;AT5G23740.1 \| Symbols: RPS11-BETA \| ribosomal protein S11-beta \| chr5:8008251-8009330 REVERSE LENGTH=159 | 0.21093 | 0.33214867 | 0.271539335 | 3.682707702 |

| | | | | | |
|---|---|---|---|---|---|
| AT2G47800 | AT2G47800.1 \| Symbols: ATMRP4, EST3, MRP4, ABCC4 \| multidrug resistance-associated protein 4 \| chr2:19574944-19580383 FORWARD LENGTH=1516;AT3G62700.1 \| Symbols: ATMRP10, MRP10, ABCC14 \| multidrug resistance-associated protein 10 \| chr3:23190428-23195727 REVERSE LENGTH=1539 | 0.33979 | 0.2028233 | 0.27130665 | 3.685866157 |
| AT4G31480 | AT4G31480.1 \| Symbols: \| Coatomer, beta subunit \| chr4:15264145-15267384 FORWARD LENGTH=948;AT4G31480.2 \| Symbols: \| Coatomer, beta subunit \| chr4:15264145-15267384 FORWARD LENGTH=948;AT4G31490.1 \| Symbols: \| Coatomer, beta subunit \| chr4:15269460-15272693 FORWARD LENGTH=948 | | 0.270321412 | 0.270321412 | 3.6993 |
| AT5G52820 | AT5G52820.1 \| Symbols: \| WD-40 repeat family protein / notchless protein, putative \| chr5:21401423-21404203 FORWARD LENGTH=473 | | 0.270182643 | 0.270182643 | 3.7012 |
| AT2G18720 | AT2G18720.1 \| Symbols: \| Translation elongation factor EF1A/initiation factor IF2gamma family protein \| chr2:8114920-8116759 REVERSE LENGTH=470;AT4G18330.2 \| Symbols: \| Translation elongation factor EF1A/initiation factor IF2gamma family protein \| chr4:10126747-10128587 FORWARD LENGTH=471 | | 0.269490932 | 0.269490932 | 3.7107 |
| ATCG00380 | ATCG00380.1 \| Symbols: RPS4 \| chloroplast ribosomal protein S4 \| chrC:45223-45828 REVERSE LENGTH=201 | | 0.269447363 | 0.269447363 | 3.7113 |
| AT1G59900 | AT1G59900.1 \| Symbols: AT-E1 ALPHA, E1 ALPHA \| pyruvate dehydrogenase complex E1 alpha subunit \| chr1:22051368-22053660 FORWARD LENGTH=389 | 0.18009 | 0.356353788 | 0.268221894 | 3.728256426 |
| AT3G28500 | AT3G28500.1 \| Symbols: \| 60S acidic ribosomal protein family \| chr3:10682204-10682551 FORWARD LENGTH=115 | 0.37623 | 0.157873133 | 0.267051567 | 3.744595146 |
| AT3G09440 | AT3G09440.1 \| Symbols: \| Heat shock protein 70 (Hsp 70) family protein \| chr3:2903434-2905632 REVERSE LENGTH=649;AT3G09440.2 \| Symbols: \| Heat shock protein 70 (Hsp 70) family protein \| chr3:2903434-2905632 REVERSE LENGTH=649;AT1G16030.1 \| Symbols: Hsp70b \| heat shock protein 70B \| chr1:5502386-5504326 REVERSE LENGTH=646 | | 0.26571011 | 0.26571011 | 3.7635 |
| AT1G22410 | AT1G22410.1 \| Symbols: \| Class-II DAHP synthetase family protein \| chr1:7912120-7914742 FORWARD LENGTH=527 | | 0.265118375 | 0.265118375 | 3.7719 |
| AT1G04480 | AT1G04480.1 \| Symbols: \| Ribosomal protein L14p/L23e family protein \| chr1:1216110-1217257 FORWARD LENGTH=140;AT2G33370.1 \| Symbols: \| Ribosomal protein L14p/L23e family protein \| chr2:14143718-14144644 REVERSE LENGTH=140;AT3G04400.1 \| Symbols: emb2171 \| Ribosomal protein L14p/L23e family protein \| chr3:1167339-1168308 FORWARD LENGTH=140;AT3G04400.2 \| Symbols: emb2171 \| Ribosomal protein L14p/L23e family protein \| chr3:1167611-1168308 FORWARD LENGTH=125 | | 0.264417356 | 0.264417356 | 3.7819 |
| AT5G18520 | AT5G18520.1 \| Symbols: \| Lung seven transmembrane receptor family protein \| chr5:6145027-6146349 FORWARD LENGTH=440 | | 0.264061262 | 0.264061262 | 3.787 |
| AT2G32060 | AT2G32060.1 \| Symbols: \| Ribosomal protein L7Ae/L30e/S12e/Gadd45 family protein \| chr2:13639228-13640104 REVERSE LENGTH=144;AT2G32060.2 \| Symbols: \| Ribosomal protein L7Ae/L30e/S12e/Gadd45 family protein \| chr2:13639228-13640104 REVERSE LENGTH=144;AT2G32060.3 \| Symbols: \| Ribosomal protein L7Ae/L30e/S12e/Gadd45 family protein \| chr2:13639228-13640104 REVERSE LENGTH=144 | | 0.262068243 | 0.262068243 | 3.8158 |
| AT5G26710 | AT5G26710.1 \| Symbols: \| Glutamyl/glutaminyl-tRNA synthetase, class Ic \| chr5:9305673-9308247 FORWARD LENGTH=719 | 0.16572 | 0.357449242 | 0.261584621 | 3.822854707 |
| AT3G02080 | AT3G02080.1 \| Symbols: \| Ribosomal protein S19e family protein \| chr3:364138-365161 REVERSE LENGTH=143;AT5G15520.1 \| Symbols: \| Ribosomal protein S19e family protein \| chr5:5037242-5038136 REVERSE LENGTH=143;AT5G61170.1 \| Symbols: \| Ribosomal protein S19e family protein \| chr5:24611158-24612202 FORWARD LENGTH=143 | 0.26631 | 0.256390534 | 0.261350267 | 3.826282679 |
| AT1G51650 | AT1G51650.1 \| Symbols: \| ATP synthase epsilon chain, mitochondrial \| chr1:19152680-19153641 FORWARD LENGTH=70 | | 0.261055709 | 0.261055709 | 3.8306 |

| | | | | | |
|---|---|---|---|---|---|
| AT3G13870 | AT3G13870.1 \| Symbols: RHD3 \| Root hair defective 3 GTP-binding protein (RHD3) \| chr3:4565762-4571109 REVERSE LENGTH=802;AT3G13870.2 \| Symbols: RHD3 \| Root hair defective 3 GTP-binding protein (RHD3) \| chr3:4565762-4570261 REVERSE LENGTH=738 | 0.23254 | 0.28952778 | 0.26103389 | 3.830920191 |
| AT4G05530 | AT4G05530.1 \| Symbols: IBR1, SDRA \| indole-3-butyric acid response 1 \| chr4:2816462-2818074 FORWARD LENGTH=254 | | 0.260538794 | 0.260538794 | 3.8382 |
| AT3G63410 | AT3G63410.1 \| Symbols: APG1, VTE3, IEP37, E37 \| S-adenosyl-L-methionine-dependent methyltransferases superfamily protein \| chr3:23415816-23417002 REVERSE LENGTH=338 | 0.0712 | 0.449599856 | 0.260399928 | 3.840246837 |
| AT4G21110 | AT4G21110.1 \| Symbols: \| G10 family protein \| chr4:11267673-11268774 FORWARD LENGTH=145 | | 0.260308205 | 0.260308205 | 3.8416 |
| AT1G27400 | AT1G27400.1 \| Symbols: \| Ribosomal protein L22p/L17e family protein \| chr1:9515230-9516725 FORWARD LENGTH=176;AT1G67430.1 \| Symbols: \| Ribosomal protein L22p/L17e family protein \| chr1:25262209-25263627 FORWARD LENGTH=175;AT1G67430.2 \| Symbols: \| Ribosomal protein L22p/L17e family protein \| chr1:25262209-25263627 FORWARD LENGTH=131 | 0.18209 | 0.337643921 | 0.25986696 | 3.848122896 |
| AT1G71380 | AT1G71380.1 \| Symbols: ATGH9B3, ATCEL3, CEL3 \| cellulase 3 \| chr1:26899989-26901749 REVERSE LENGTH=484;AT1G22880.1 \| Symbols: ATGH9B4, ATCEL5, CEL5 \| cellulase 5 \| chr1:8095770-8097539 FORWARD LENGTH=484 | | 0.258946605 | 0.258946605 | 3.8618 |
| AT1G80750 | AT1G80750.1 \| Symbols: \| Ribosomal protein L30/L7 family protein \| chr1:30349052-30350434 FORWARD LENGTH=247 | 0.067461 | 0.449519015 | 0.258490007 | 3.8686215 |
| AT1G53210 | AT1G53210.1 \| Symbols: \| sodium/calcium exchanger family protein / calcium-binding EF hand family protein \| chr1:19844790-19847533 FORWARD LENGTH=585 | 0.20078 | 0.314336906 | 0.257558453 | 3.882613783 |
| AT2G36620 | AT2G36620.1 \| Symbols: RPL24A \| ribosomal protein L24 \| chr2:15350548-15351819 REVERSE LENGTH=164;AT3G53020.1 \| Symbols: STV1, RPL24B, RPL24 \| Ribosomal protein L24e family protein \| chr3:19660749-19661912 REVERSE LENGTH=163 | 0.15057 | 0.364311997 | 0.257440998 | 3.884385184 |
| AT1G07810 | AT1G07810.1 \| Symbols: ECA1, ATECA1, ACA3 \| ER-type Ca2+-ATPase 1 \| chr1:2416681-2420572 FORWARD LENGTH=1061 | | 0.257267816 | 0.257267816 | 3.887 |
| AT3G45030 | AT3G45030.1 \| Symbols: \| Ribosomal protein S10p/S20e family protein \| chr3:16471606-16472312 REVERSE LENGTH=124;AT5G62300.1 \| Symbols: \| Ribosomal protein S10p/S20e family protein \| chr5:25021388-25022235 REVERSE LENGTH=124;AT5G62300.2 \| Symbols: \| Ribosomal protein S10p/S20e family protein \| chr5:25021388-25022235 REVERSE LENGTH=124 | 0.057861 | 0.455746969 | 0.256803985 | 3.894020575 |
| AT1G09590 | AT1G09590.1 \| Symbols: \| Translation protein SH3-like family protein \| chr1:3106549-3107606 FORWARD LENGTH=164;AT1G09690.1 \| Symbols: \| Translation protein SH3-like family protein \| chr1:3136407-3137430 REVERSE LENGTH=164;AT1G57660.1 \| Symbols: \| Translation protein SH3-like family protein \| chr1:21355567-21356364 FORWARD LENGTH=164;AT1G57860.1 \| Symbols: \| Translation protein SH3-like family protein \| chr1:21430034-21430827 REVERSE LENGTH=164 | 0.12235 | 0.39117509 | 0.256762545 | 3.894649043 |
| AT4G16566 | AT4G16566.1 \| Symbols: HINT4 \| histidine triad nucleotide-binding 4 \| chr4:9334873-9335884 REVERSE LENGTH=146 | | 0.256213169 | 0.256213169 | 3.903 |
| AT3G06730 | AT3G06730.1 \| Symbols: TRX P, TRX z \| Thioredoxin z \| chr3:2124276-2125845 FORWARD LENGTH=183 | | 0.255754476 | 0.255754476 | 3.91 |
| AT4G22010 | AT4G22010.1 \| Symbols: sks4 \| SKU5 similar 4 \| chr4:11663429-11666463 FORWARD LENGTH=541;AT1G41830.1 \| Symbols: SKS6 \| SKU5-similar 6 \| chr1:15603892-15607802 REVERSE LENGTH=542 | | 0.255649862 | 0.255649862 | 3.9116 |
| AT5G61140 | AT5G61140.2 \| Symbols: \| U5 small nuclear ribonucleoprotein helicase \| chr5:24589999-24603311 FORWARD LENGTH=2157;AT5G61140.1 \| Symbols: \| U5 small nuclear ribonucleoprotein helicase \| chr5:24589999-24603311 FORWARD LENGTH=2146 | | 0.255232261 | 0.255232261 | 3.918 |
| AT2G41475 | AT2G41475.1 \| Symbols: \| Embryo-specific protein 3, (ATS3) \| chr2:17295259-17296329 REVERSE LENGTH=179 | | 0.254660283 | 0.254660283 | 3.9268 |

| | | | | | |
|---|---|---|---|---|---|
| AT4G36130 | AT4G36130.1 \| Symbols: \| Ribosomal protein L2 family \| chr4:17097613-17098656 FORWARD LENGTH=258 | 0.013707 | 0.494853523 | 0.254280262 | 3.932668597 |
| AT1G34430 | AT1G34430.1 \| Symbols: EMB3003 \| 2-oxoacid dehydrogenases acyltransferase family protein \| chr1:12588027-12590084 REVERSE LENGTH=465 | 0.22909 | 0.279298402 | 0.254194201 | 3.934000049 |
| AT5G35680 | AT5G35680.3 \| Symbols: \| Nucleic acid-binding, OB-fold-like protein \| chr5:13858085-13858589 REVERSE LENGTH=150;AT2G04520.1 \| Symbols: \| Nucleic acid-binding, OB-fold-like protein \| chr2:1574802-1575239 REVERSE LENGTH=145;AT5G35680.1 \| Symbols: \| Nucleic acid-binding, OB-fold-like protein \| chr5:13858152-13858589 REVERSE LENGTH=145;AT5G35680.2 \| Symbols: \| Nucleic acid-binding, OB-fold-like protein \| chr5:13858152-13858589 REVERSE LENGTH=145 | | 0.254148982 | 0.254148982 | 3.9347 |
| AT1G52360 | AT1G52360.2 \| Symbols: \| Coatomer, beta' subunit \| chr1:19499420-19505397 FORWARD LENGTH=970;AT1G52360.1 \| Symbols: \| Coatomer, beta' subunit \| chr1:19499282-19505397 FORWARD LENGTH=926;AT1G79990.1 \| Symbols: \| structural molecules \| chr1:30084522-30091949 FORWARD LENGTH=1135;AT1G79990.3 \| Symbols: \| structural molecules \| chr1:30085910-30091949 FORWARD LENGTH=920;AT1G79990.5 \| Symbols: \| structural molecules \| chr1:30085910-30091949 FORWARD LENGTH=912 | | 0.253897324 | 0.253897324 | 3.9386 |
| AT4G39980 | AT4G39980.1 \| Symbols: DHS1 \| 3-deoxy-D-arabino-heptulosonate 7-phosphate synthase 1 \| chr4:18539654-18541832 FORWARD LENGTH=525 | | 0.253549696 | 0.253549696 | 3.944 |
| AT5G24710 | AT5G24710.1 \| Symbols: \| Transducin/WD40 repeat-like superfamily protein \| chr5:8459148-8467920 REVERSE LENGTH=1377 | | 0.253517556 | 0.253517556 | 3.9445 |
| AT2G27710 | AT2G27710.1 \| Symbols: \| 60S acidic ribosomal protein family \| chr2:11816929-11817670 FORWARD LENGTH=115;AT2G27710.2 \| Symbols: \| 60S acidic ribosomal protein family \| chr2:11816929-11817670 FORWARD LENGTH=115;AT2G27710.3 \| Symbols: \| 60S acidic ribosomal protein family \| chr2:11816929-11817670 FORWARD LENGTH=115;AT2G27710.4 \| Symbols: \| 60S acidic ribosomal protein family \| chr2:11816929-11817670 FORWARD LENGTH=98;AT2G27720.2 \| Symbols: \| 60S acidic ribosomal protein family \| chr2:11818696-11819370 FORWARD LENGTH=130;AT2G27720.3 \| Symbols: \| 60S acidic ribosomal protein family \| chr2:11818696-11819370 FORWARD LENGTH=127 | 0.28202 | 0.224982001 | 0.253501001 | 3.944757603 |
| AT4G03430 | AT4G03430.1 \| Symbols: STA1, EMB2770 \| pre-mRNA splicing factor-related \| chr4:1517411-1520500 REVERSE LENGTH=1029 | 0.25347 | | 0.25347 | 3.945240068 |
| AT2G22125 | AT2G22125.1 \| Symbols: CSI1 \| binding \| chr2:9406793-9414223 FORWARD LENGTH=2150 | | 0.251603975 | 0.251603975 | 3.9745 |
| AT1G04810 | AT1G04810.1 \| Symbols: \| 26S proteasome regulatory complex, non-ATPase subcomplex, Rpn2/Psmd1 subunit \| chr1:1350304-1355261 FORWARD LENGTH=1001 | | 0.250607724 | 0.250607724 | 3.9903 |
| AT4G32520 | AT4G32520.1 \| Symbols: SHM3, AtSHMT3 \| serine hydroxymethyltransferase 3 \| chr4:15689642-15692334 REVERSE LENGTH=529;AT4G32520.2 \| Symbols: SHM3 \| serine hydroxymethyltransferase 3 \| chr4:15689642-15692334 REVERSE LENGTH=529 | | 0.250438267 | 0.250438267 | 3.993 |
| AT2G01750 | AT2G01750.2 \| Symbols: MAP70-3 \| microtubule-associated proteins 70-3 \| chr2:328351-331848 FORWARD LENGTH=630;AT2G01750.1 \| Symbols: ATMAP70-3, MAP70-3 \| microtubule-associated proteins 70-3 \| chr2:328351-331848 FORWARD LENGTH=629;AT1G14840.1 \| Symbols: ATMAP70-4, MAP70-4 \| microtubule-associated proteins 70-4 \| chr1:5112415-5115299 REVERSE LENGTH=604;AT1G14840.2 \| Symbols: MAP70-4 \| microtubule-associated proteins 70-4 \| chr1:5112415-5115299 REVERSE LENGTH=589 | | 0.250168864 | 0.250168864 | 3.9973 |
| AT5G16130 | AT5G16130.1 \| Symbols: \| Ribosomal protein S7e family protein \| chr5:5268984-5269912 FORWARD LENGTH=190;AT3G02560.1 \| Symbols: \| Ribosomal protein S7e family protein \| chr3:542341-543168 FORWARD LENGTH=191;AT3G02560.2 \| Symbols: \| Ribosomal protein S7e family protein \| chr3:542341-543168 FORWARD LENGTH=191 | 0.13939 | 0.360646278 | 0.250018139 | 3.999709796 |

| | | | | | |
|---|---|---|---|---|---|
| AT3G09630 | AT3G09630.1 \| Symbols: \| Ribosomal protein L4/L1 family \| chr3:2953813-2955444 FORWARD LENGTH=406;AT3G09630.2 \| Symbols: \| Ribosomal protein L4/L1 family \| chr3:2953813-2955444 FORWARD LENGTH=405 | 0.104 | 0.395288165 | 0.249644083 | 4.005702798 |
| AT4G34450 | AT4G34450.1 \| Symbols: \| coatomer gamma-2 subunit, putative / gamma-2 coat protein, putative / gamma-2 COP, putative \| chr4:16471956-16476795 FORWARD LENGTH=886 | 0.16773 | 0.331411149 | 0.249570574 | 4.006882633 |
| AT3G56150 | AT3G56150.1 \| Symbols: EIF3C, ATEIF3C-1, EIF3C-1, ATTIF3C1, TIF3C1 \| eukaryotic translation initiation factor 3C \| chr3:20833790-20836820 REVERSE LENGTH=900;AT3G56150.2 \| Symbols: EIF3C \| eukaryotic translation initiation factor 3C \| chr3:20833790-20836820 REVERSE LENGTH=900 | 0.14384 | 0.354924579 | 0.249382289 | 4.009907853 |
| AT5G08540 | AT5G08540.1 \| Symbols: \| unknown protein; FUNCTIONS IN: molecular_function unknown; INVOLVED IN: biological_process unknown; LOCATED IN: chloroplast thylakoid membrane, chloroplast, chloroplast envelope; EXPRESSED IN: 24 plant structures; EXPRESSED DURING: 13 growth stages; Has 30201 Blast hits to 17322 proteins in 780 species: Archae - 12; Bacteria - 1396; Metazoa - 17338; Fungi - 3422; Plants - 5037; Viruses - 0; Other Eukaryotes - 2996 (source: NCBI BLink). \| chr5:2763914-2765431 FORWARD LENGTH=346 | | 0.249264669 | 0.249264669 | 4.0118 |
| AT5G19770 | AT5G19770.1 \| Symbols: TUA3 \| tubulin alpha-3 \| chr5:6682761-6684474 REVERSE LENGTH=450;AT5G19780.1 \| Symbols: TUA5 \| tubulin alpha-5 \| chr5:6687212-6688926 FORWARD LENGTH=450 | 0.09002 | 0.408329931 | 0.249174965 | 4.013244263 |
| AT3G18600 | AT3G18600.1 \| Symbols: \| P-loop containing nucleoside triphosphate hydrolases superfamily protein \| chr3:6399724-6403007 REVERSE LENGTH=568 | | 0.249121845 | 0.249121845 | 4.0141 |
| AT1G63810 | AT1G63810.1 \| Symbols: \| CONTAINS InterPro DOMAIN/s: Nrap protein (InterPro:IPR005554); Has 396 Blast hits to 382 proteins in 182 species: Archae - 3; Bacteria - 2; Metazoa - 142; Fungi - 146; Plants - 43; Viruses - 0; Other Eukaryotes - 60 (source: NCBI BLink). \| chr1:23669690-23675542 REVERSE LENGTH=1053 | | 0.248941997 | 0.248941997 | 4.017 |
| AT2G45810 | AT2G45810.1 \| Symbols: \| DEA(D/H)-box RNA helicase family protein \| chr2:18859836-18862318 FORWARD LENGTH=528;AT4G00660.1 \| Symbols: RH8, ATRH8 \| RNAhelicase-like 8 \| chr4:274638-277438 FORWARD LENGTH=505;AT4G00660.2 \| Symbols: RH8, ATRH8 \| RNAhelicase-like 8 \| chr4:274638-277438 FORWARD LENGTH=505;AT3G61240.1 \| Symbols: \| DEA(D/H)-box RNA helicase family protein \| chr3:22666590-22669154 FORWARD LENGTH=498;AT3G61240.2 \| Symbols: \| DEA(D/H)-box RNA helicase family protein \| chr3:22666590-22669154 FORWARD LENGTH=498 | | 0.248471898 | 0.248471898 | 4.0246 |
| AT3G04840 | AT3G04840.1 \| Symbols: \| Ribosomal protein S3Ae \| chr3:1329751-1331418 FORWARD LENGTH=262 | 0.090169 | 0.4058771 | 0.24802305 | 4.031883324 |
| AT3G60240 | AT3G60240.4 \| Symbols: EIF4G, CUM2 \| eukaryotic translation initiation factor 4G \| chr3:22261842-22268295 FORWARD LENGTH=1727;AT3G60240.3 \| Symbols: EIF4G, CUM2 \| eukaryotic translation initiation factor 4G \| chr3:22261842-22268295 FORWARD LENGTH=1725;AT3G60240.2 \| Symbols: EIF4G, CUM2 \| eukaryotic translation initiation factor 4G \| chr3:22261842-22268295 FORWARD LENGTH=1723 | | 0.246682125 | 0.246682125 | 4.0538 |
| AT2G30860 | AT2G30860.1 \| Symbols: ATGSTF9, GLUTTR, ATGSTF7, GSTF9 \| glutathione S-transferase PHI 9 \| chr2:13139132-13140057 FORWARD LENGTH=215;AT2G30860.2 \| Symbols: ATGSTF9, GLUTTR, ATGSTF7, GSTF9 \| glutathione S-transferase PHI 9 \| chr2:13139132-13140079 FORWARD LENGTH=166 | 0.24654 | | 0.24654 | 4.056136935 |
| AT3G57330 | AT3G57330.1 \| Symbols: ACA11 \| autoinhibited Ca2+-ATPase 11 \| chr3:21211655-21216375 REVERSE LENGTH=1025;AT2G41560.1 \| Symbols: ACA4 \| autoinhibited Ca(2+)-ATPase, isoform 4 \| chr2:17332256-17337179 REVERSE LENGTH=1030 | | 0.245603694 | 0.245603694 | 4.0716 |

| | | | | | |
|---|---|---|---|---|---|
| AT5G27850 | AT5G27850.1 \| Symbols: \| Ribosomal protein L18e/L15 superfamily protein \| chr5:9873169-9874297 FORWARD LENGTH=187;AT3G05590.1 \| Symbols: RPL18 \| ribosomal protein L18 \| chr3:1621511-1622775 FORWARD LENGTH=187;AT2G47570.1 \| Symbols: \| Ribosomal protein L18e/L15 superfamily protein \| chr2:19515898-19516717 REVERSE LENGTH=135;AT2G47570.2 \| Symbols: \| Ribosomal protein L18e/L15 superfamily protein \| chr2:19515898-19516717 REVERSE LENGTH=135 | 0.1356 | 0.355403917 | 0.245501958 | 4.073287264 |
| AT2G18900 | AT2G18900.1 \| Symbols: \| Transducin/WD40 repeat-like superfamily protein \| chr2:8188491-8192409 REVERSE LENGTH=804 | | 0.245074012 | 0.245074012 | 4.0804 |
| AT4G29430 | AT4G29430.1 \| Symbols: rps15ae \| ribosomal protein S15A E \| chr4:14472440-14473383 FORWARD LENGTH=129 | | 0.24473813 | 0.24473813 | 4.086 |
| AT2G20140 | AT2G20140.1 \| Symbols: \| AAA-type ATPase family protein \| chr2:8692736-8694837 FORWARD LENGTH=443;AT4G29040.1 \| Symbols: RPT2a \| regulatory particle AAA-ATPase 2A \| chr4:14312369-14314386 FORWARD LENGTH=443 | 0.046187 | 0.443203475 | 0.244695237 | 4.086716239 |
| AT2G16930 | AT2G16930.1 \| Symbols: \| Ribosomal protein L27 family protein \| chr2:7341492-7342118 FORWARD LENGTH=154;AT2G16930.2 \| Symbols: \| Ribosomal protein L27 family protein \| chr2:7341492-7342118 FORWARD LENGTH=154;AT2G16930.3 \| Symbols: \| Ribosomal protein L27 family protein \| chr2:7341492-7342118 FORWARD LENGTH=154;AT5G15220.1 \| Symbols: \| Ribosomal protein L27 family protein \| chr5:4941466-4942133 REVERSE LENGTH=154 | | 0.244039339 | 0.244039339 | 4.0977 |
| AT5G12860 | AT5G12860.1 \| Symbols: DiT1 \| dicarboxylate transporter 1 \| chr5:4059927-4061919 REVERSE LENGTH=557;AT5G12860.2 \| Symbols: DiT1 \| dicarboxylate transporter 1 \| chr5:4059850-4061919 REVERSE LENGTH=556 | | 0.24354603 | 0.24354603 | 4.106 |
| AT4G15000 | AT4G15000.1 \| Symbols: \| Ribosomal L27e protein family \| chr4:8571896-8572303 FORWARD LENGTH=135;AT4G15000.2 \| Symbols: \| Ribosomal L27e protein family \| chr4:8571896-8572387 FORWARD LENGTH=131;AT3G22230.1 \| Symbols: \| Ribosomal L27e protein family \| chr3:7844136-7844543 REVERSE LENGTH=135;AT2G32220.1 \| Symbols: \| Ribosomal L27e protein family \| chr2:13678945-13679352 FORWARD LENGTH=135 | 0.14275 | 0.344103782 | 0.243426891 | 4.108009581 |
| AT1G04820 | AT1G04820.1 \| Symbols: TUA4, TOR2 \| tubulin alpha-4 chain \| chr1:1356421-1358266 REVERSE LENGTH=450;AT1G50010.1 \| Symbols: TUA2 \| tubulin alpha-2 chain \| chr1:18517737-18519729 FORWARD LENGTH=450;AT4G14960.2 \| Symbols: TUA6 \| Tubulin/FtsZ family protein \| chr4:8548769-8550319 REVERSE LENGTH=450;AT4G14960.1 \| Symbols: TUA6 \| Tubulin/FtsZ family protein \| chr4:8548753-8550319 REVERSE LENGTH=427;CON__ENSEMBL:ENSBTAP00000016242 | 0.17842 | 0.308024026 | 0.243222013 | 4.111469961 |
| AT1G07770 | AT1G07770.1 \| Symbols: RPS15A \| ribosomal protein S15A \| chr1:2408413-2409065 REVERSE LENGTH=130;AT1G07770.2 \| Symbols: RPS15A \| ribosomal protein S15A \| chr1:2408413-2409065 REVERSE LENGTH=130;AT5G59850.1 \| Symbols: \| Ribosomal protein S8 family protein \| chr5:24112499-24113084 REVERSE LENGTH=130;AT2G39590.1 \| Symbols: \| Ribosomal protein S8 family protein \| chr2:16517588-16518265 REVERSE LENGTH=136;AT3G46040.1 \| Symbols: RPS15AD \| ribosomal protein S15A D \| chr3:16914891-16915529 FORWARD LENGTH=130 | 0.3134 | 0.17197496 | 0.24268748 | 4.120525703 |
| AT5G22060 | AT5G22060.1 \| Symbols: ATJ2, J2 \| DNAJ homologue 2 \| chr5:7303798-7305668 REVERSE LENGTH=419 | 0.099446 | 0.384644973 | 0.242045486 | 4.131454856 |
| AT5G62670 | AT5G62670.1 \| Symbols: AHA11, HA11 \| H(+)-ATPase 11 \| chr5:25159495-25164957 FORWARD LENGTH=956;AT1G80660.1 \| Symbols: AHA9, HA9 \| H(+)-ATPase 9 \| chr1:30316227-30319948 REVERSE LENGTH=954;AT1G80660.2 \| Symbols: HA9 \| H(+)-ATPase 9 \| chr1:30316227-30319948 REVERSE LENGTH=945 | | 0.241359336 | 0.241359336 | 4.1432 |
| AT2G27720 | AT2G27720.1 \| Symbols: \| 60S acidic ribosomal protein family \| chr2:11818696-11819370 FORWARD LENGTH=115 | 0.25119 | 0.231395779 | 0.24129289 | 4.144340935 |

| | | | | | |
|---|---|---|---|---|---|
| AT1G20260 | AT1G20260.1 \| Symbols: \| ATPase, V1 complex, subunit B protein \| chr1:7016971-7020290 FORWARD LENGTH=487;AT1G76030.1 \| Symbols: \| ATPase, V1 complex, subunit B protein \| chr1:28534134-28536916 FORWARD LENGTH=486 | 0.22659 | 0.255245291 | 0.240917645 | 4.150796005 |
| AT2G40360 | AT2G40360.1 \| Symbols: \| Transducin/WD40 repeat-like superfamily protein \| chr2:16853123-16856635 REVERSE LENGTH=753 | | 0.238920081 | 0.238920081 | 4.1855 |
| AT3G05560 | AT3G05560.1 \| Symbols: \| Ribosomal L22e protein family \| chr3:1614641-1615204 FORWARD LENGTH=124;AT3G05560.2 \| Symbols: \| Ribosomal L22e protein family \| chr3:1614641-1615204 FORWARD LENGTH=124;AT3G05560.3 \| Symbols: \| Ribosomal L22e protein family \| chr3:1614641-1615204 FORWARD LENGTH=124 | 0.34407 | 0.132996409 | 0.238533205 | 4.192288457 |
| AT1G48830 | AT1G48830.1 \| Symbols: \| Ribosomal protein S7e family protein \| chr1:18059854-18060935 REVERSE LENGTH=191;AT1G48830.2 \| Symbols: \| Ribosomal protein S7e family protein \| chr1:18059854-18060935 REVERSE LENGTH=191 | 0.20935 | 0.266773376 | 0.238061688 | 4.200591907 |
| AT3G60245 | AT3G60245.1 \| Symbols: \| Zinc-binding ribosomal protein family protein \| chr3:22268803-22269750 FORWARD LENGTH=92;AT3G10950.1 \| Symbols: \| Zinc-binding ribosomal protein family protein \| chr3:3423893-3424566 FORWARD LENGTH=92 | | 0.236837742 | 0.236837742 | 4.2223 |
| AT4G16450 | AT4G16450.1 \| Symbols: \| unknown protein; FUNCTIONS IN: molecular_function unknown; INVOLVED IN: photorespiration; LOCATED IN: mitochondrion, mitochondrial membrane, mitochondrial respiratory chain complex I, respiratory chain complex I, membrane; EXPRESSED IN: 25 plant structures; EXPRESSED DURING: 15 growth stages; Has 30201 Blast hits to 17322 proteins in 780 species: Archae - 12; Bacteria - 1396; Metazoa - 17338; Fungi - 3422; Plants - 5037; Viruses - 0; Other Eukaryotes - 2996 (source: NCBI BLink). \| chr4:9280132-9280541 FORWARD LENGTH=106;AT4G16450.2 \| Symbols: \| unknown protein; FUNCTIONS IN: molecular_function unknown; INVOLVED IN: photorespiration; LOCATED IN: mitochondrion, mitochondrial membrane, mitochondrial respiratory chain complex I, respiratory chain complex I; EXPRESSED IN: 25 plant structures; EXPRESSED DURING: 15 growth stages. \| chr4:9280132-9280541 FORWARD LENGTH=106 | | 0.236714404 | 0.236714404 | 4.2245 |
| AT4G15770 | AT4G15770.1 \| Symbols: \| RNA binding \| chr4:8978060-8978623 FORWARD LENGTH=187 | | 0.236524043 | 0.236524043 | 4.2279 |
| AT1G31230 | AT1G31230.1 \| Symbols: AK-HSDH I, AK-HSDH \| aspartate kinase-homoserine dehydrogenase i \| chr1:11158744-11163055 REVERSE LENGTH=911 | | 0.235943657 | 0.235943657 | 4.2383 |
| AT2G42740 | AT2G42740.1 \| Symbols: RPL16A \| ribosomal protein large subunit 16A \| chr2:17791794-17792946 FORWARD LENGTH=182;AT3G58700.1 \| Symbols: \| Ribosomal L5P family protein \| chr3:21711661-21712816 FORWARD LENGTH=182;AT4G18730.1 \| Symbols: RPL16B \| ribosomal protein L16B \| chr4:10302238-10303206 FORWARD LENGTH=182;AT5G45775.2 \| Symbols: \| Ribosomal L5P family protein \| chr5:18565281-18566496 REVERSE LENGTH=182;AT5G45775.1 \| Symbols: \| Ribosomal L5P family protein \| chr5:18565281-18566377 REVERSE LENGTH=172 | 0.22362 | 0.24691358 | 0.23526679 | 4.250493661 |
| AT2G43770 | AT2G43770.1 \| Symbols: \| Transducin/WD40 repeat-like superfamily protein \| chr2:18134272-18135303 REVERSE LENGTH=343 | | 0.234499578 | 0.234499578 | 4.2644 |
| AT5G57870 | AT5G57870.1 \| Symbols: eIFiso4G1 \| MIF4G domain-containing protein / MA3 domain-containing protein \| chr5:23439755-23443433 FORWARD LENGTH=780;AT5G57870.2 \| Symbols: eIFiso4G1 \| MIF4G domain-containing protein / MA3 domain-containing protein \| chr5:23439755-23443433 FORWARD LENGTH=776 | 0.26196 | 0.204666394 | 0.233313197 | 4.286084171 |
| AT2G20580 | AT2G20580.1 \| Symbols: RPN1A, ATRPN1A \| 26S proteasome regulatory subunit S2 1A \| chr2:8859211-8864699 FORWARD LENGTH=891;AT4G28470.1 \| Symbols: RPN1B, ATRPN1B \| 26S proteasome regulatory subunit S2 1B \| chr4:14067082-14072357 REVERSE LENGTH=891 | 0.19313 | 0.272895972 | 0.233012986 | 4.291606305 |

| | | | | | |
|---|---|---|---|---|---|
| AT3G49010 | AT3G49010.1 \| Symbols: ATBBC1, BBC1, RSU2 \| breast basic conserved 1 \| chr3:18166971-18168047 REVERSE LENGTH=206;AT3G49010.2 \| Symbols: ATBBC1, BBC1, RSU2 \| breast basic conserved 1 \| chr3:18166971-18168047 REVERSE LENGTH=206;AT3G49010.3 \| Symbols: ATBBC1, BBC1, RSU2 \| breast basic conserved 1 \| chr3:18166971-18168047 REVERSE LENGTH=206;AT3G49010.4 \| Symbols: ATBBC1, BBC1, RSU2 \| breast basic conserved 1 \| chr3:18166971-18168047 REVERSE LENGTH=204;AT3G48960.1 \| Symbols: \| Ribosomal protein L13e family protein \| chr3:18150444-18151412 REVERSE LENGTH=206;AT3G49010.5 \| Symbols: ATBBC1, BBC1, RSU2 \| breast basic conserved 1 \| chr3:18166971-18168047 REVERSE LENGTH=153;AT3G29796.1 \| Symbols: \| unknown protein; Has 28 Blast hits to 20 proteins in 8 species: Archae - 0; Bacteria - 0; Metazoa - 18; Fungi - 0; Plants - 1; Viruses - 0; Other Eukaryotes - 9 (source: NCBI BLink). \| chr3:11708561-11713028 FORWARD LENGTH=463 | 0.11447 | 0.351160586 | 0.232815293 | 4.295250487 |
| AT1G10490 | AT1G10490.1 \| Symbols: \| Domain of unknown function (DUF1726) ;Putative ATPase (DUF699) \| chr1:3453589-3459925 FORWARD LENGTH=1028 | | 0.232574366 | 0.232574366 | 4.2997 |
| AT1G43170 | AT1G43170.1 \| Symbols: ARP1, emb2207, RPL3A, RP1 \| ribosomal protein 1 \| chr1:16266992-16268631 FORWARD LENGTH=389;AT1G43170.2 \| Symbols: ARP1, RPL3A, RP1 \| ribosomal protein 1 \| chr1:16266992-16268631 FORWARD LENGTH=389;AT1G43170.3 \| Symbols: ARP1, RPL3A, RP1 \| ribosomal protein 1 \| chr1:16266992-16268631 FORWARD LENGTH=389;AT1G43170.5 \| Symbols: RP1 \| ribosomal protein 1 \| chr1:16266992-16268631 FORWARD LENGTH=389;AT1G43170.6 \| Symbols: RP1 \| ribosomal protein 1 \| chr1:16266992-16268631 FORWARD LENGTH=389;AT1G43170.7 \| Symbols: RP1 \| ribosomal protein 1 \| chr1:16266992-16268631 FORWARD LENGTH=389;AT1G43170.8 \| Symbols: RP1 \| ribosomal protein 1 \| chr1:16266992-16268631 FORWARD LENGTH=389;AT1G43170.9 \| Symbols: RP1 \| ribosomal protein 1 \| chr1:16266992-16268631 FORWARD LENGTH=389;AT1G43170.4 \| Symbols: ARP1, RPL3A, RP1 \| ribosomal protein 1 \| chr1:16267519-16268631 FORWARD LENGTH=306;AT1G61580.1 \| Symbols: RPL3B, ARP2 \| R-protein L3 B \| chr1:22720833-22722402 REVERSE LENGTH=390 | 0.13756 | 0.326882845 | 0.232221423 | 4.306234924 |
| AT3G07050 | AT3G07050.1 \| Symbols: \| GTP-binding family protein \| chr3:2229602-2232279 REVERSE LENGTH=582 | | 0.232120889 | 0.232120889 | 4.3081 |
| AT1G15500 | AT1G15500.1 \| Symbols: ATNTT2 \| TLC ATP/ADP transporter \| chr1:5326426-5328688 FORWARD LENGTH=618 | 0.3705 | 0.091945568 | 0.231222784 | 4.324833315 |
| AT5G14520 | AT5G14520.1 \| Symbols: \| pescadillo-related \| chr5:4680789-4684330 REVERSE LENGTH=590 | | 0.2310803 | 0.2310803 | 4.3275 |
| AT3G52990 | AT3G52990.1 \| Symbols: \| Pyruvate kinase family protein \| chr3:19649046-19652237 FORWARD LENGTH=527;AT3G52990.2 \| Symbols: \| Pyruvate kinase family protein \| chr3:19649336-19652237 FORWARD LENGTH=474;AT2G36580.1 \| Symbols: \| Pyruvate kinase family protein \| chr2:15339253-15342781 FORWARD LENGTH=527 | | 0.230802964 | 0.230802964 | 4.3327 |
| AT1G15440 | AT1G15440.1 \| Symbols: PWP2, ATPWP2 \| periodic tryptophan protein 2 \| chr1:5306159-5309460 REVERSE LENGTH=900;AT1G15440.2 \| Symbols: PWP2 \| periodic tryptophan protein 2 \| chr1:5306159-5309460 REVERSE LENGTH=860 | | 0.23019198 | 0.23019198 | 4.3442 |
| AT4G00630 | AT4G00630.2 \| Symbols: KEA2 \| K+ efflux antiporter 2 \| chr4:261655-267789 REVERSE LENGTH=1185;AT4G00630.1 \| Symbols: KEA2, ATKEA2 \| K+ efflux antiporter 2 \| chr4:261655-267789 REVERSE LENGTH=1174 | | 0.230022542 | 0.230022542 | 4.3474 |
| AT5G10360 | AT5G10360.1 \| Symbols: EMB3010, RPS6B \| Ribosomal protein S6e \| chr5:3258734-3260142 REVERSE LENGTH=249 | 0.074789 | 0.385089341 | 0.22993917 | 4.34897629 |
| AT5G06360 | AT5G06360.1 \| Symbols: \| Ribosomal protein S8e family protein \| chr5:1944835-1946512 FORWARD LENGTH=260 | | 0.229858637 | 0.229858637 | 4.3505 |

| | | | | | |
|---|---|---|---|---|---|
| AT3G13580 | AT3G13580.1 \| Symbols: \| Ribosomal protein L30/L7 family protein \| chr3:4433809-4435109 FORWARD LENGTH=244;AT3G13580.2 \| Symbols: \| Ribosomal protein L30/L7 family protein \| chr3:4433809-4435109 FORWARD LENGTH=244;AT3G13580.3 \| Symbols: \| Ribosomal protein L30/L7 family protein \| chr3:4433809-4435109 FORWARD LENGTH=244 | | 0.229436733 | 0.229436733 | 4.3585 |
| AT1G52740 | AT1G52740.1 \| Symbols: HTA9 \| histone H2A protein 9 \| chr1:19645409-19646221 FORWARD LENGTH=134;AT2G38810.1 \| Symbols: HTA8 \| histone H2A 8 \| chr2:16219444-16220679 REVERSE LENGTH=136;AT2G38810.2 \| Symbols: HTA8 \| histone H2A 8 \| chr2:16219444-16220679 REVERSE LENGTH=136;AT2G38810.3 \| Symbols: HTA8 \| histone H2A 8 \| chr2:16219444-16220679 REVERSE LENGTH=136;AT3G54560.1 \| Symbols: HTA11 \| histone H2A 11 \| chr3:20196532-20197466 FORWARD LENGTH=136;AT4G13570.2 \| Symbols: HTA4 \| histone H2A 4 \| chr4:7884582-7885664 FORWARD LENGTH=124;AT4G13570.1 \| Symbols: HTA4 \| histone H2A 4 \| chr4:7884516-7885664 FORWARD LENGTH=118 | | 0.228655051 | 0.228655051 | 4.3734 |
| AT3G27830 | AT3G27830.1 \| Symbols: RPL12-A, RPL12 \| ribosomal protein L12-A \| chr3:10318576-10319151 FORWARD LENGTH=191;AT3G27850.1 \| Symbols: RPL12-C \| ribosomal protein L12-C \| chr3:10324905-10325468 FORWARD LENGTH=187 | 0.069233 | 0.386772384 | 0.228002692 | 4.385913123 |
| AT5G57490 | AT5G57490.1 \| Symbols: VDAC4, ATVDAC4 \| voltage dependent anion channel 4 \| chr5:23283895-23285335 REVERSE LENGTH=274 | | 0.227671152 | 0.227671152 | 4.3923 |
| AT4G39260 | AT4G39260.1 \| Symbols: CCR1, ATGRP8, GR-RBP8, GRP8 \| cold, circadian rhythm, and RNA binding 1 \| chr4:18274166-18274958 REVERSE LENGTH=169;AT4G39260.2 \| Symbols: CCR1, ATGRP8, GR-RBP8, GRP8 \| cold, circadian rhythm, and RNA binding 1 \| chr4:18274166-18274958 REVERSE LENGTH=126;AT4G39260.3 \| Symbols: CCR1, ATGRP8, GR-RBP8, GRP8 \| cold, circadian rhythm, and RNA binding 1 \| chr4:18274166-18274958 REVERSE LENGTH=92;AT4G39260.4 \| Symbols: CCR1, ATGRP8, GR-RBP8, GRP8 \| cold, circadian rhythm, and RNA binding 1 \| chr4:18274166-18274958 REVERSE LENGTH=105 | | 0.227231412 | 0.227231412 | 4.4008 |
| AT3G55410 | AT3G55410.1 \| Symbols: \| 2-oxoglutarate dehydrogenase, E1 component \| chr3:20541897-20545728 FORWARD LENGTH=1017;AT5G65750.1 \| Symbols: \| 2-oxoglutarate dehydrogenase, E1 component \| chr5:26304212-26307947 FORWARD LENGTH=1025 | | 0.227050837 | 0.227050837 | 4.4043 |
| ATCG00160 | ATCG00160.1 \| Symbols: RPS2 \| ribosomal protein S2 \| chrC:15013-15723 REVERSE LENGTH=236 | | 0.226485177 | 0.226485177 | 4.4153 |
| AT2G27530 | AT2G27530.1 \| Symbols: PGY1 \| Ribosomal protein L1p/L10e family \| chr2:11763443-11764570 REVERSE LENGTH=216;AT2G27530.2 \| Symbols: PGY1 \| Ribosomal protein L1p/L10e family \| chr2:11763443-11764570 REVERSE LENGTH=216;AT5G22440.1 \| Symbols: \| Ribosomal protein L1p/L10e family \| chr5:7435328-7436486 REVERSE LENGTH=217;AT5G22440.2 \| Symbols: \| Ribosomal protein L1p/L10e family \| chr5:7435328-7436486 REVERSE LENGTH=217 | 0.086267 | 0.36668989 | 0.226478445 | 4.415431231 |
| AT5G20010 | AT5G20010.1 \| Symbols: RAN-1, RAN1, ATRAN1 \| RAS-related nuclear protein-1 \| chr5:6760364-6761747 FORWARD LENGTH=221;AT5G20020.1 \| Symbols: RAN2 \| RAS-related GTP-binding nuclear protein 2 \| chr5:6762817-6764381 FORWARD LENGTH=221;AT5G55190.1 \| Symbols: RAN3, ATRAN3 \| RAN GTPase 3 \| chr5:22392285-22393957 FORWARD LENGTH=221 | 0.14006 | 0.312119604 | 0.226089802 | 4.423021254 |
| AT5G52650 | AT5G52650.1 \| Symbols: \| RNA binding Plectin/S10 domain-containing protein \| chr5:21355781-21357003 REVERSE LENGTH=179;AT4G25740.1 \| Symbols: \| RNA binding Plectin/S10 domain-containing protein \| chr4:13107488-13108751 REVERSE LENGTH=177;AT4G25740.2 \| Symbols: \| RNA binding Plectin/S10 domain-containing protein \| chr4:13107488-13108751 REVERSE LENGTH=149 | 0.21527 | 0.236663984 | 0.225966992 | 4.425425103 |

| | | | | | |
|---|---|---|---|---|---|
| AT5G14030 | AT5G14030.1 \| Symbols: \| translocon-associated protein beta (TRAPB) family protein \| chr5:4526878-4528253 FORWARD LENGTH=195;AT5G14030.2 \| Symbols: \| translocon-associated protein beta (TRAPB) family protein \| chr5:4526878-4528253 FORWARD LENGTH=195;AT5G14030.3 \| Symbols: \| translocon-associated protein beta (TRAPB) family protein \| chr5:4526878-4528253 FORWARD LENGTH=195;AT5G14030.4 \| Symbols: \| translocon-associated protein beta (TRAPB) family protein \| chr5:4526878-4528253 FORWARD LENGTH=195 | 0.077021 | 0.373412995 | 0.225216997 | 4.440162206 |
| AT1G52280 | AT1G52280.1 \| Symbols: AtRABG3d, RABG3d \| RAB GTPase homolog G3D \| chr1:19468150-19469449 REVERSE LENGTH=206;AT3G16100.1 \| Symbols: ATRABG3C, ATRAB7D, RABG3c \| RAB GTPase homolog G3C \| chr3:5459270-5460556 FORWARD LENGTH=206;AT3G18820.1 \| Symbols: ATRABG3F, ATRAB7B, RAB71, RABG3F, RAB7B \| RAB GTPase homolog G3F \| chr3:6484266-6486005 FORWARD LENGTH=206;AT1G49300.1 \| Symbols: ATRAB7, ATRABG3E, RABG3E \| RAB GTPase homolog G3E \| chr1:18234842-18236968 FORWARD LENGTH=206;AT1G49300.2 \| Symbols: ATRAB7, ATRABG3E, RABG3E \| RAB GTPase homolog G3E \| chr1:18234842-18236968 FORWARD LENGTH=206;AT1G22740.1 \| Symbols: RAB7, ATRABG3B, RAB75, RABG3B \| RAB GTPase homolog G3B \| chr1:8049247-8050494 FORWARD LENGTH=203 | | 0.22462824 | 0.22462824 | 4.4518 |
| AT5G07090 | AT5G07090.1 \| Symbols: \| Ribosomal protein S4 (RPS4A) family protein \| chr5:2202410-2203805 FORWARD LENGTH=262;AT5G58420.1 \| Symbols: \| Ribosomal protein S4 (RPS4A) family protein \| chr5:23619599-23620896 FORWARD LENGTH=262;AT2G17360.1 \| Symbols: \| Ribosomal protein S4 (RPS4A) family protein \| chr2:7546598-7548138 FORWARD LENGTH=261;AT5G07090.2 \| Symbols: \| Ribosomal protein S4 (RPS4A) family protein \| chr5:2202783-2203805 FORWARD LENGTH=244;AT2G17360.2 \| Symbols: \| Ribosomal protein S4 (RPS4A) family protein \| chr2:7546598-7548041 FORWARD LENGTH=184 | 0.12086 | 0.327729165 | 0.224294582 | 4.458422445 |
| AT4G02510 | AT4G02510.1 \| Symbols: TOC159, TOC86, PPI2, TOC160, ATTOC159 \| translocon at the outer envelope membrane of chloroplasts 159 \| chr4:1104766-1109360 FORWARD LENGTH=1503 | | 0.223723657 | 0.223723657 | 4.4698 |
| AT5G48760 | AT5G48760.1 \| Symbols: \| Ribosomal protein L13 family protein \| chr5:19771315-19772686 REVERSE LENGTH=206;AT5G48760.2 \| Symbols: \| Ribosomal protein L13 family protein \| chr5:19771315-19772686 REVERSE LENGTH=206 | | 0.223658608 | 0.223658608 | 4.4711 |
| AT5G62190 | AT5G62190.1 \| Symbols: PRH75 \| DEAD box RNA helicase (PRH75) \| chr5:24980542-24983879 REVERSE LENGTH=671 | | 0.223468681 | 0.223468681 | 4.4749 |
| AT5G16840 | AT5G16840.2 \| Symbols: BPA1 \| binding partner of acd11 1 \| chr5:5536042-5538026 FORWARD LENGTH=260;AT5G16840.1 \| Symbols: BPA1 \| binding partner of acd11 1 \| chr5:5536042-5538026 FORWARD LENGTH=259;AT5G16840.3 \| Symbols: BPA1 \| binding partner of acd11 1 \| chr5:5536042-5538026 FORWARD LENGTH=257 | | 0.221258518 | 0.221258518 | 4.5196 |
| AT1G30400 | AT1G30400.1 \| Symbols: ATMRP1, EST1, ABCC1, ATABCC1, MRP1 \| multidrug resistance-associated protein 1 \| chr1:10728139-10737697 FORWARD LENGTH=1622;AT1G30400.2 \| Symbols: ATMRP1, EST1, ABCC1 \| multidrug resistance-associated protein 1 \| chr1:10728139-10737697 FORWARD LENGTH=1622 | 0.27268 | 0.16889039 | 0.220785195 | 4.529289202 |
| AT1G09770 | AT1G09770.1 \| Symbols: ATCDC5, CDC5, ATMYBCDC5 \| cell division cycle 5 \| chr1:3162002-3165122 FORWARD LENGTH=844 | 0.14032 | 0.300724747 | 0.220522373 | 4.534687274 |
| AT2G05920 | AT2G05920.1 \| Symbols: \| Subtilase family protein \| chr2:2269831-2272207 REVERSE LENGTH=754 | 0.18404 | 0.253923112 | 0.218981556 | 4.56659464 |
| AT3G49100 | AT3G49100.1 \| Symbols: \| Signal recognition particle, SRP9/SRP14 subunit \| chr3:18196944-18198298 FORWARD LENGTH=103 | | 0.21805495 | 0.21805495 | 4.586 |

| | | | | | |
|---|---|---|---|---|---|
| AT3G27300 | AT3G27300.1 \| Symbols: G6PD5 \| glucose-6-phosphate dehydrogenase 5 \| chr3:10083318-10086288 REVERSE LENGTH=516;AT3G27300.2 \| Symbols: G6PD5 \| glucose-6-phosphate dehydrogenase 5 \| chr3:10083318-10086288 REVERSE LENGTH=516;AT3G27300.3 \| Symbols: G6PD5 \| glucose-6-phosphate dehydrogenase 5 \| chr3:10083318-10086288 REVERSE LENGTH=516 | 0.091864 | 0.342876736 | 0.217370368 | 4.600443058 |
| AT3G12800 | AT3G12800.1 \| Symbols: SDRB, DECR \| short-chain dehydrogenase-reductase B \| chr3:4063463-4064757 REVERSE LENGTH=298 | 0.12897 | 0.305213039 | 0.217091519 | 4.60635221 |
| AT4G12600 | AT4G12600.2 \| Symbols: \| Ribosomal protein L7Ae/L30e/S12e/Gadd45 family protein \| chr4:7453296-7454208 FORWARD LENGTH=183;AT5G20160.2 \| Symbols: \| Ribosomal protein L7Ae/L30e/S12e/Gadd45 family protein \| chr5:6804075-6805102 REVERSE LENGTH=160;AT4G12600.1 \| Symbols: \| Ribosomal protein L7Ae/L30e/S12e/Gadd45 family protein \| chr4:7453296-7454208 FORWARD LENGTH=128;AT5G20160.1 \| Symbols: \| Ribosomal protein L7Ae/L30e/S12e/Gadd45 family protein \| chr5:6804075-6805102 REVERSE LENGTH=128;AT5G20160.3 \| Symbols: \| Ribosomal protein L7Ae/L30e/S12e/Gadd45 family protein \| chr5:6804653-6805102 REVERSE LENGTH=109 | | 0.216910329 | 0.216910329 | 4.6102 |
| AT3G05060 | AT3G05060.1 \| Symbols: \| NOP56-like pre RNA processing ribonucleoprotein \| chr3:1413174-1415564 REVERSE LENGTH=533 | | 0.215805602 | 0.215805602 | 4.6338 |
| AT3G58750 | AT3G58750.1 \| Symbols: CSY2 \| citrate synthase 2 \| chr3:21724564-21727458 REVERSE LENGTH=514 | | 0.215703192 | 0.215703192 | 4.636 |
| AT3G54670 | AT3G54670.3 \| Symbols: TTN8 \| Structural maintenance of chromosomes (SMC) family protein \| chr3:20235818-20243701 FORWARD LENGTH=1239;AT3G54670.1 \| Symbols: TTN8, SMC1, ATSMC1 \| Structural maintenance of chromosomes (SMC) family protein \| chr3:20235818-20243701 FORWARD LENGTH=1238 | | 0.215628774 | 0.215628774 | 4.6376 |
| AT1G31660 | AT1G31660.1 \| Symbols: \| CONTAINS InterPro DOMAIN/s: Bystin (InterPro:IPR007955); Has 475 Blast hits to 467 proteins in 210 species: Archae - 0; Bacteria - 9; Metazoa - 155; Fungi - 139; Plants - 55; Viruses - 0; Other Eukaryotes - 117 (source: NCBI BLink). \| chr1:11331176-11333440 REVERSE LENGTH=444 | | 0.215021395 | 0.215021395 | 4.6507 |
| AT4G16130 | AT4G16130.1 \| Symbols: ARA1, ISA1, ATISA1 \| arabinose kinase \| chr4:9120875-9127656 FORWARD LENGTH=1039 | | 0.213816844 | 0.213816844 | 4.6769 |
| AT2G40510 | AT2G40510.1 \| Symbols: \| Ribosomal protein S26e family protein \| chr2:16918506-16919623 FORWARD LENGTH=133;AT2G40590.1 \| Symbols: \| Ribosomal protein S26e family protein \| chr2:16945215-16946345 REVERSE LENGTH=131;AT3G56340.1 \| Symbols: \| Ribosomal protein S26e family protein \| chr3:20892309-20893343 REVERSE LENGTH=130 | 0.13884 | 0.287323296 | 0.213081648 | 4.693036726 |
| AT1G47530 | AT1G47530.1 \| Symbols: \| MATE efflux family protein \| chr1:17451724-17454110 FORWARD LENGTH=484 | | 0.212861066 | 0.212861066 | 4.6979 |
| AT3G02530 | AT3G02530.1 \| Symbols: \| TCP-1/cpn60 chaperonin family protein \| chr3:528806-532457 REVERSE LENGTH=535;AT5G16070.1 \| Symbols: \| TCP-1/cpn60 chaperonin family protein \| chr5:5247549-5251050 REVERSE LENGTH=535 | | 0.212165574 | 0.212165574 | 4.7133 |
| AT1G65030 | AT1G65030.1 \| Symbols: \| Transducin/WD40 repeat-like superfamily protein \| chr1:24156798-24158511 FORWARD LENGTH=345 | | 0.211184322 | 0.211184322 | 4.7352 |
| AT4G13930 | AT4G13930.1 \| Symbols: SHM4 \| serine hydroxymethyltransferase 4 \| chr4:8048013-8050021 REVERSE LENGTH=471 | 0.18319 | 0.239137193 | 0.211163597 | 4.735664748 |
| AT1G28110 | AT1G28110.1 \| Symbols: SCPL45 \| serine carboxypeptidase-like 45 \| chr1:9804153-9806832 REVERSE LENGTH=461;AT1G28110.2 \| Symbols: SCPL45 \| serine carboxypeptidase-like 45 \| chr1:9804153-9806832 REVERSE LENGTH=461 | 0.23141 | 0.190577832 | 0.210993916 | 4.739473152 |

| | | | | | |
|---|---|---|---|---|---|
| AT1G48630 | AT1G48630.1 \| Symbols: RACK1B_AT \| receptor for activated C kinase 1B \| chr1:17981977-17983268 REVERSE LENGTH=326;AT3G18130.1 \| Symbols: RACK1C_AT \| receptor for activated C kinase 1C \| chr3:6211109-6212371 REVERSE LENGTH=326 | | 0.210992721 | 0.210992721 | 4.7395 |
| AT3G23400 | AT3G23400.1 \| Symbols: FIB4 \| Plastid-lipid associated protein PAP / fibrillin family protein \| chr3:8376636-8378225 REVERSE LENGTH=284 | | 0.210677116 | 0.210677116 | 4.7466 |
| AT1G22780 | AT1G22780.1 \| Symbols: PFL, RPS18A, PFL1 \| Ribosomal protein S13/S18 family \| chr1:8067990-8069163 FORWARD LENGTH=152;AT1G34030.1 \| Symbols: \| Ribosomal protein S13/S18 family \| chr1:12370285-12371465 REVERSE LENGTH=152;AT4G09800.1 \| Symbols: RPS18C \| S18 ribosomal protein \| chr4:6173818-6174963 FORWARD LENGTH=152 | 0.20083 | 0.220196415 | 0.210513208 | 4.750295772 |
| AT3G54540 | AT3G54540.1 \| Symbols: ATGCN4, GCN4 \| general control non-repressible 4 \| chr3:20190393-20192564 FORWARD LENGTH=723 | | 0.210154674 | 0.210154674 | 4.7584 |
| AT2G29540 | AT2G29540.3 \| Symbols: ATRPC14, ATRPAC14, RPC14 \| RNApolymerase 14 kDa subunit \| chr2:12643288-12643828 FORWARD LENGTH=139;AT2G29540.1 \| Symbols: ATRPC14, ATRPAC14, RPC14 \| RNApolymerase 14 kDa subunit \| chr2:12642727-12643828 FORWARD LENGTH=122;AT2G29540.2 \| Symbols: ATRPC14, ATRPAC14, RPC14 \| RNApolymerase 14 kDa subunit \| chr2:12642727-12643828 FORWARD LENGTH=122 | | 0.209687566 | 0.209687566 | 4.769 |
| AT2G03270 | AT2G03270.1 \| Symbols: \| DNA-binding protein, putative \| chr2:994071-995990 FORWARD LENGTH=639 | 0.10551 | 0.312119604 | 0.208814802 | 4.788932537 |
| AT1G64550 | AT1G64550.1 \| Symbols: ATGCN3, GCN3 \| general control non-repressible 3 \| chr1:23968850-23973369 FORWARD LENGTH=715 | 0.15288 | 0.262556778 | 0.207718389 | 4.814210263 |
| AT1G33120 | AT1G33120.1 \| Symbols: \| Ribosomal protein L6 family \| chr1:12010986-12012223 FORWARD LENGTH=194;AT1G33140.1 \| Symbols: PGY2 \| Ribosomal protein L6 family \| chr1:12023360-12024502 FORWARD LENGTH=194;AT4G10450.1 \| Symbols: \| Ribosomal protein L6 family \| chr4:6463201-6464458 REVERSE LENGTH=194 | 0.17976 | 0.23557681 | 0.207668405 | 4.815369003 |
| AT4G11420 | AT4G11420.1 \| Symbols: EIF3A, ATEIF3A-1, EIF3A-1, ATTIF3A1, TIF3A1 \| eukaryotic translation initiation factor 3A \| chr4:6947834-6952053 REVERSE LENGTH=987 | | 0.207658443 | 0.207658443 | 4.8156 |
| AT5G11150 | AT5G11150.1 \| Symbols: ATVAMP713, VAMP713 \| vesicle-associated membrane protein 713 \| chr5:3546625-3547844 REVERSE LENGTH=221 | 0.23399 | 0.180378434 | 0.207184217 | 4.826622484 |
| AT1G69620 | AT1G69620.1 \| Symbols: RPL34 \| ribosomal protein L34 \| chr1:26189900-26191081 FORWARD LENGTH=119;AT1G26880.1 \| Symbols: \| Ribosomal protein L34e superfamily protein \| chr1:9315640-9316681 REVERSE LENGTH=120;AT3G28900.1 \| Symbols: \| Ribosomal protein L34e superfamily protein \| chr3:10902864-10904415 REVERSE LENGTH=120;AT1G26880.2 \| Symbols: \| Ribosomal protein L34e superfamily protein \| chr1:9315798-9316681 REVERSE LENGTH=96 | 0.20647 | | 0.20647 | 4.843318642 |
| AT3G12810 | AT3G12810.1 \| Symbols: PIE1, SRCAP, chr13 \| SNF2 domain-containing protein / helicase domain-containing protein \| chr3:4065636-4073992 FORWARD LENGTH=2055 | | 0.206428173 | 0.206428173 | 4.8443 |
| AT1G16470 | AT1G16470.1 \| Symbols: PAB1 \| proteasome subunit PAB1 \| chr1:5623122-5625439 FORWARD LENGTH=235;AT1G16470.2 \| Symbols: PAB1 \| proteasome subunit PAB1 \| chr1:5623122-5625439 FORWARD LENGTH=235;AT1G79210.1 \| Symbols: \| N-terminal nucleophile aminohydrolases (Ntn hydrolases) superfamily protein \| chr1:29796286-29798240 REVERSE LENGTH=235;AT1G79210.2 \| Symbols: \| N-terminal nucleophile aminohydrolases (Ntn hydrolases) superfamily protein \| chr1:29796286-29798240 REVERSE LENGTH=235;AT1G79210.3 \| Symbols: \| N-terminal nucleophile aminohydrolases (Ntn hydrolases) superfamily protein \| chr1:29796286-29798240 REVERSE LENGTH=235 | | 0.206334468 | 0.206334468 | 4.8465 |

| | | | | | |
|---|---|---|---|---|---|
| AT2G18020 | AT2G18020.1 \| Symbols: EMB2296 \| Ribosomal protein L2 family \| chr2:7837151-7838160 FORWARD LENGTH=258;AT3G51190.1 \| Symbols: \| Ribosomal protein L2 family \| chr3:19016606-19017547 REVERSE LENGTH=260 | 0.15443 | 0.256633989 | 0.205531994 | 4.865422551 |
| AT4G18040 | AT4G18040.1 \| Symbols: EIF4E, CUM1, AT.EIF4E1, eIF4E1 \| eukaryotic translation initiation factor 4E \| chr4:10016724-10018151 REVERSE LENGTH=235 | | 0.205300868 | 0.205300868 | 4.8709 |
| AT3G61110 | AT3G61110.1 \| Symbols: ARS27A, RS27A \| ribosomal protein S27 \| chr3:22611710-22612632 FORWARD LENGTH=86;AT2G45710.1 \| Symbols: \| Zinc-binding ribosomal protein family protein \| chr2:18831243-18831999 FORWARD LENGTH=84;AT3G61111.1 \| Symbols: \| Zinc-binding ribosomal protein family protein \| chr3:22613230-22613919 REVERSE LENGTH=86 | | 0.20366184 | 0.20366184 | 4.9101 |
| AT1G01100 | AT1G01100.1 \| Symbols: \| 60S acidic ribosomal protein family \| chr1:50284-50954 REVERSE LENGTH=112;AT1G01100.2 \| Symbols: \| 60S acidic ribosomal protein family \| chr1:50284-50954 REVERSE LENGTH=112;AT1G01100.4 \| Symbols: \| 60S acidic ribosomal protein family \| chr1:50284-50954 REVERSE LENGTH=112;AT1G01100.3 \| Symbols: \| 60S acidic ribosomal protein family \| chr1:50284-50954 REVERSE LENGTH=96;AT4G00810.1 \| Symbols: \| 60S acidic ribosomal protein family \| chr4:346179-346957 REVERSE LENGTH=113;AT4G00810.2 \| Symbols: \| 60S acidic ribosomal protein family \| chr4:346179-346957 REVERSE LENGTH=113 | 0.28207 | 0.124946898 | 0.203508449 | 4.913800906 |
| AT2G02470 | AT2G02470.1 \| Symbols: AL6 \| alfin-like 6 \| chr2:652837-654621 FORWARD LENGTH=256;AT1G14510.1 \| Symbols: AL7 \| alfin-like 7 \| chr1:4962171-4964154 REVERSE LENGTH=252;AT2G02470.2 \| Symbols: AL6 \| alfin-like 6 \| chr2:652837-654621 FORWARD LENGTH=247 | 0.20328 | | 0.20328 | 4.919323101 |
| AT1G15690 | AT1G15690.1 \| Symbols: AVP1, ATAVP3, AVP-3, AtVHP1;1 \| Inorganic H pyrophosphatase family protein \| chr1:5399115-5402185 FORWARD LENGTH=770;AT1G15690.2 \| Symbols: AVP1, ATAVP3, AVP-3, AtVHP1;1 \| Inorganic H pyrophosphatase family protein \| chr1:5399115-5402185 FORWARD LENGTH=642 | 0.22933 | 0.176956699 | 0.203143349 | 4.922632236 |
| AT3G62700 | AT3G62700.1 \| Symbols: ATMRP10, MRP10, ABCC14 \| multidrug resistance-associated protein 10 \| chr3:23190428-23195727 REVERSE LENGTH=1539 | | 0.202314478 | 0.202314478 | 4.9428 |
| AT5G14120 | AT5G14120.1 \| Symbols: \| Major facilitator superfamily protein \| chr5:4556308-4558447 FORWARD LENGTH=579 | | 0.202289922 | 0.202289922 | 4.9434 |
| AT1G51980 | AT1G51980.1 \| Symbols: \| Insulinase (Peptidase family M16) protein \| chr1:19323692-19326771 REVERSE LENGTH=503;AT3G16480.1 \| Symbols: MPPalpha \| mitochondrial processing peptidase alpha subunit \| chr3:5599906-5602716 FORWARD LENGTH=499;AT1G51980.2 \| Symbols: \| Insulinase (Peptidase family M16) protein \| chr1:19323929-19326771 REVERSE LENGTH=451 | | 0.202150885 | 0.202150885 | 4.9468 |
| AT3G52300 | AT3G52300.1 \| Symbols: ATPQ \| ATP synthase D chain, mitochondrial \| chr3:19396689-19398119 FORWARD LENGTH=168;AT3G52300.2 \| Symbols: ATPQ \| ATP synthase D chain, mitochondrial \| chr3:19396689-19397866 FORWARD LENGTH=122 | | 0.201515396 | 0.201515396 | 4.9624 |
| AT5G05000 | AT5G05000.1 \| Symbols: TOC34, ATTOC34, OEP34 \| translocon at the outer envelope membrane of chloroplasts 34 \| chr5:1474262-1475772 REVERSE LENGTH=313;AT5G05000.2 \| Symbols: TOC34, ATTOC34, OEP34 \| translocon at the outer envelope membrane of chloroplasts 34 \| chr5:1474262-1475772 REVERSE LENGTH=313;AT5G05000.3 \| Symbols: TOC34, ATTOC34, OEP34 \| translocon at the outer envelope membrane of chloroplasts 34 \| chr5:1474262-1475772 REVERSE LENGTH=313 | 0.1045 | 0.297902765 | 0.201201382 | 4.970144781 |
| AT3G50590 | AT3G50590.1 \| Symbols: \| Transducin/WD40 repeat-like superfamily protein \| chr3:18771292-18779220 FORWARD LENGTH=1614 | | 0.200625953 | 0.200625953 | 4.9844 |
| AT1G55150 | AT1G55150.1 \| Symbols: \| DEA(D/H)-box RNA helicase family protein \| chr1:20574634-20577141 FORWARD LENGTH=501 | | 0.200481155 | 0.200481155 | 4.988 |

| | | | | | |
|---|---|---|---|---|---|
| AT4G34670 | AT4G34670.1 \| Symbols: \| Ribosomal protein S3Ae \| chr4:16548724-16550222 FORWARD LENGTH=262 | 0.11514 | 0.285714286 | 0.200427143 | 4.989344186 |
| AT5G27640 | AT5G27640.2 \| Symbols: TIF3B1, EIF3B, ATEIF3B-1, EIF3B-1, ATTIF3B1 \| translation initiation factor 3B1 \| chr5:9781207-9784759 REVERSE LENGTH=738;AT5G25780.1 \| Symbols: EIF3B-2, EIF3B, ATEIF3B-2 \| eukaryotic translation initiation factor 3B-2 \| chr5:8973420-8976758 REVERSE LENGTH=714;AT5G27640.1 \| Symbols: TIF3B1, EIF3B, ATEIF3B-1, EIF3B-1, ATTIF3B1 \| translation initiation factor 3B1 \| chr5:9781207-9784759 REVERSE LENGTH=712 | 0.08102 | 0.319539863 | 0.200279931 | 4.993011499 |
| AT2G39390 | AT2G39390.1 \| Symbols: \| Ribosomal L29 family protein \| chr2:16450803-16451762 REVERSE LENGTH=123 | | 0.200048012 | 0.200048012 | 4.9988 |
| AT2G24050 | AT2G24050.1 \| Symbols: eIFiso4G2 \| MIF4G domain-containing protein / MA3 domain-containing protein \| chr2:10225500-10228456 REVERSE LENGTH=747 | | 0.199844122 | 0.199844122 | 5.0039 |
| AT1G76300 | AT1G76300.1 \| Symbols: SmD3 \| snRNP core protein SMD3 \| chr1:28625957-28627080 FORWARD LENGTH=128;AT1G20600.1 \| Symbols: \| AP2/B3-like transcriptional factor family protein \| chr1:7133111-7134237 FORWARD LENGTH=237 | | 0.199425654 | 0.199425654 | 5.0144 |
| AT5G52470 | AT5G52470.1 \| Symbols: FIB1, FBR1, ATFIB1, ATFBR1, SKIP7 \| fibrillarin 1 \| chr5:21294290-21296509 FORWARD LENGTH=308 | | 0.198444197 | 0.198444197 | 5.0392 |
| AT2G32180 | AT2G32180.1 \| Symbols: PTAC18 \| plastid transcriptionally active 18 \| chr2:13672362-13672781 REVERSE LENGTH=139;AT2G32650.1 \| Symbols: \| RmlC-like cupins superfamily protein \| chr2:13851547-13851966 FORWARD LENGTH=139;AT2G32650.2 \| Symbols: \| RmlC-like cupins superfamily protein \| chr2:13851547-13851966 FORWARD LENGTH=139 | | 0.198192485 | 0.198192485 | 5.0456 |
| AT2G41840 | AT2G41840.1 \| Symbols: \| Ribosomal protein S5 family protein \| chr2:17460016-17461398 REVERSE LENGTH=285 | 0.11784 | 0.276556321 | 0.19719816 | 5.071041222 |
| AT5G06110 | AT5G06110.1 \| Symbols: \| DnaJ domain ;Myb-like DNA-binding domain \| chr5:1841009-1843000 REVERSE LENGTH=663;AT5G06110.2 \| Symbols: \| DnaJ domain ;Myb-like DNA-binding domain \| chr5:1841009-1843000 REVERSE LENGTH=663 | | 0.197013279 | 0.197013279 | 5.0758 |
| AT3G25860 | AT3G25860.1 \| Symbols: LTA2, PLE2 \| 2-oxoacid dehydrogenases acyltransferase family protein \| chr3:9460632-9462585 FORWARD LENGTH=480 | 0.089154 | 0.304636569 | 0.196895284 | 5.0788418 |
| AT5G28540 | AT5G28540.1 \| Symbols: BIP1 \| heat shock protein 70 (Hsp 70) family protein \| chr5:10540665-10543274 REVERSE LENGTH=669;AT1G09080.1 \| Symbols: BIP3 \| Heat shock protein 70 (Hsp 70) family protein \| chr1:2929268-2931804 REVERSE LENGTH=675;AT1G09080.2 \| Symbols: BIP3 \| Heat shock protein 70 (Hsp 70) family protein \| chr1:2929268-2931804 REVERSE LENGTH=665 | 0.09834 | 0.293677131 | 0.196008566 | 5.101817854 |
| AT4G08950 | AT4G08950.1 \| Symbols: EXO \| Phosphate-responsive 1 family protein \| chr4:5740378-5741322 FORWARD LENGTH=314 | 0.19582 | | 0.19582 | 5.106730671 |
| AT1G49240 | AT1G49240.1 \| Symbols: ACT8 \| actin 8 \| chr1:18216539-18217947 FORWARD LENGTH=377;AT3G18780.2 \| Symbols: ACT2, DER1, LSR2, ENL2 \| actin 2 \| chr3:6475535-6476832 FORWARD LENGTH=377;AT3G18780.1 \| Symbols: ACT2, DER1, LSR2, ENL2 \| actin 2 \| chr3:6475535-6476728 FORWARD LENGTH=371 | 0.19884 | 0.192607716 | 0.195723858 | 5.109239163 |
| AT3G58510 | AT3G58510.1 \| Symbols: \| DEA(D/H)-box RNA helicase family protein \| chr3:21640608-21643464 FORWARD LENGTH=612;AT3G58510.2 \| Symbols: \| DEA(D/H)-box RNA helicase family protein \| chr3:21640608-21643464 FORWARD LENGTH=612;AT3G58510.3 \| Symbols: \| DEA(D/H)-box RNA helicase family protein \| chr3:21640608-21643464 FORWARD LENGTH=612;AT3G58570.1 \| Symbols: \| P-loop containing nucleoside triphosphate hydrolases superfamily protein \| chr3:21657099-21660352 FORWARD LENGTH=646;AT2G42520.1 \| Symbols: \| P-loop containing nucleoside triphosphate hydrolases superfamily protein \| chr2:17705382-17708744 FORWARD LENGTH=633 | 0.096631 | 0.293685756 | 0.195158378 | 5.124043403 |

| | | | | | |
|---|---|---|---|---|---|
| AT2G43150 | AT2G43150.1 \| Symbols: \| Proline-rich extensin-like family protein \| chr2:17945893-17946531 FORWARD LENGTH=212 | | 0.19485201 | 0.19485201 | 5.1321 |
| AT3G48870 | AT3G48870.1 \| Symbols: ATCLPC, ATHSP93-III, HSP93-III \| Clp ATPase \| chr3:18122363-18126008 REVERSE LENGTH=952;AT3G48870.2 \| Symbols: HSP93-III \| Clp ATPase \| chr3:18122363-18125915 REVERSE LENGTH=921;AT5G50920.1 \| Symbols: CLPC, ATHSP93-V, HSP93-V, DCA1, CLPC1 \| CLPC homologue 1 \| chr5:20715710-20719800 REVERSE LENGTH=929;AT5G51070.1 \| Symbols: ERD1, CLPD, SAG15 \| Clp ATPase \| chr5:20764479-20768481 FORWARD LENGTH=945 | 0.080857 | 0.3057356 | 0.1932963 | 5.17340477 |
| AT5G15550 | AT5G15550.1 \| Symbols: \| Transducin/WD40 repeat-like superfamily protein \| chr5:5059051-5062003 REVERSE LENGTH=433;AT5G15550.2 \| Symbols: \| Transducin/WD40 repeat-like superfamily protein \| chr5:5059315-5062003 REVERSE LENGTH=402 | | 0.19305392 | 0.19305392 | 5.1799 |
| AT5G63400 | AT5G63400.1 \| Symbols: ADK1 \| adenylate kinase 1 \| chr5:25393274-25394817 REVERSE LENGTH=246;AT5G63400.2 \| Symbols: ADK1 \| adenylate kinase 1 \| chr5:25393502-25394817 REVERSE LENGTH=190;AT5G50370.1 \| Symbols: \| Adenylate kinase family protein \| chr5:20509382-20510631 REVERSE LENGTH=248 | 0.1743 | 0.210939313 | 0.192619656 | 5.191578154 |
| AT1G51060 | AT1G51060.1 \| Symbols: HTA10 \| histone H2A 10 \| chr1:18926948-18927443 FORWARD LENGTH=132;AT4G27230.1 \| Symbols: HTA2 \| histone H2A 2 \| chr4:13637515-13638325 REVERSE LENGTH=131;AT4G27230.2 \| Symbols: HTA2 \| histone H2A 2 \| chr4:13637515-13638325 REVERSE LENGTH=131;AT5G54640.1 \| Symbols: HTA1, RAT5, ATHTA1 \| Histone superfamily protein \| chr5:22196540-22197279 FORWARD LENGTH=130 | 0.2634 | 0.121397528 | 0.192398764 | 5.197538582 |
| AT3G09500 | AT3G09500.1 \| Symbols: \| Ribosomal L29 family protein \| chr3:2917047-2917895 FORWARD LENGTH=123;AT3G55170.1 \| Symbols: \| Ribosomal L29 family protein \| chr3:20453136-20454293 REVERSE LENGTH=123;AT3G55170.2 \| Symbols: \| Ribosomal L29 family protein \| chr3:20453136-20454293 REVERSE LENGTH=123 | 0.20416 | 0.179478436 | 0.191819218 | 5.213241985 |
| AT4G16720 | AT4G16720.1 \| Symbols: \| Ribosomal protein L23/L15e family protein \| chr4:9400156-9401315 REVERSE LENGTH=204;AT4G17390.1 \| Symbols: \| Ribosomal protein L23/L15e family protein \| chr4:9714407-9715545 REVERSE LENGTH=204 | 0.065766 | 0.317833646 | 0.191799823 | 5.213769151 |
| AT5G04830 | AT5G04830.1 \| Symbols: \| Nuclear transport factor 2 (NTF2) family protein \| chr5:1402294-1403671 REVERSE LENGTH=178;AT5G04830.2 \| Symbols: \| Nuclear transport factor 2 (NTF2) family protein \| chr5:1402294-1403671 REVERSE LENGTH=178 | 0.15903 | 0.224487047 | 0.191758524 | 5.21489205 |
| AT4G21710 | AT4G21710.1 \| Symbols: NRPB2, EMB1989, RPB2 \| DNA-directed RNA polymerase family protein \| chr4:11535684-11542200 REVERSE LENGTH=1188 | | 0.190211706 | 0.190211706 | 5.2573 |
| AT1G04170 | AT1G04170.1 \| Symbols: EIF2 GAMMA \| eukaryotic translation initiation factor 2 gamma subunit \| chr1:1097423-1099702 FORWARD LENGTH=465 | | 0.190197235 | 0.190197235 | 5.2577 |
| AT4G16155 | AT4G16155.1 \| Symbols: \| dihydrolipoyl dehydrogenases \| chr4:9153570-9157322 REVERSE LENGTH=630;AT3G16950.2 \| Symbols: LPD1, ptlpd1 \| lipoamide dehydrogenase 1 \| chr3:5786508-5790383 REVERSE LENGTH=623;AT3G16950.1 \| Symbols: LPD1, ptlpd1 \| lipoamide dehydrogenase 1 \| chr3:5786761-5790383 REVERSE LENGTH=570 | 0.048661 | 0.331663958 | 0.190162479 | 5.258660936 |
| AT5G49500 | AT5G49500.1 \| Symbols: \| Signal recognition particle, SRP54 subunit protein \| chr5:20077556-20080206 REVERSE LENGTH=497;AT1G48900.1 \| Symbols: \| Signal recognition particle, SRP54 subunit protein \| chr1:18084972-18087743 REVERSE LENGTH=495;AT1G48900.2 \| Symbols: \| Signal recognition particle, SRP54 subunit protein \| chr1:18085031-18087743 REVERSE LENGTH=431 | | 0.190114068 | 0.190114068 | 5.26 |
| AT4G30800 | AT4G30800.1 \| Symbols: \| Nucleic acid-binding, OB-fold-like protein \| chr4:15001216-15002392 FORWARD LENGTH=159 | | 0.189962387 | 0.189962387 | 5.2642 |
| AT4G10320 | AT4G10320.1 \| Symbols: \| tRNA synthetase class I (I, L, M and V) family protein \| chr4:6397526-6404509 REVERSE LENGTH=1190 | 0.12409 | 0.255128074 | 0.189609037 | 5.274010221 |

| | | | | | |
|---|---|---|---|---|---|
| AT5G26742 | AT5G26742.2 \| Symbols: emb1138 \| DEAD box RNA helicase (RH3) \| chr5:9285540-9288871 REVERSE LENGTH=748;AT5G26742.1 \| Symbols: emb1138 \| DEAD box RNA helicase (RH3) \| chr5:9285540-9288871 REVERSE LENGTH=747;AT5G26742.3 \| Symbols: emb1138 \| DEAD box RNA helicase (RH3) \| chr5:9285540-9288618 REVERSE LENGTH=655 | 0.091432 | 0.287174775 | 0.189303387 | 5.282525655 |
| AT3G52140 | AT3G52140.4 \| Symbols: \| tetratricopeptide repeat (TPR)-containing protein \| chr3:19333232-19341295 FORWARD LENGTH=1419;AT3G52140.2 \| Symbols: \| tetratricopeptide repeat (TPR)-containing protein \| chr3:19333232-19341295 FORWARD LENGTH=1407;AT3G52140.1 \| Symbols: \| tetratricopeptide repeat (TPR)-containing protein \| chr3:19333232-19341295 FORWARD LENGTH=1403;AT3G52140.3 \| Symbols: \| tetratricopeptide repeat (TPR)-containing protein \| chr3:19333232-19341295 FORWARD LENGTH=1396 | | 0.189114566 | 0.189114566 | 5.2878 |
| AT5G57290 | AT5G57290.1 \| Symbols: \| 60S acidic ribosomal protein family \| chr5:23207049-23207835 REVERSE LENGTH=120;AT5G57290.3 \| Symbols: \| 60S acidic ribosomal protein family \| chr5:23207049-23207835 REVERSE LENGTH=119;AT5G57290.2 \| Symbols: \| 60S acidic ribosomal protein family \| chr5:23207089-23207835 REVERSE LENGTH=89 | 0.20825 | 0.169943749 | 0.189096874 | 5.28829471 |
| AT2G21390 | AT2G21390.1 \| Symbols: \| Coatomer, alpha subunit \| chr2:9152428-9156577 FORWARD LENGTH=1218 | | 0.188686366 | 0.188686366 | 5.2998 |
| AT3G07110 | AT3G07110.2 \| Symbols: \| Ribosomal protein L13 family protein \| chr3:2252092-2253332 FORWARD LENGTH=207;AT3G07110.1 \| Symbols: \| Ribosomal protein L13 family protein \| chr3:2252092-2253332 FORWARD LENGTH=206 | 0.10414 | 0.273044998 | 0.188592499 | 5.302437826 |
| AT5G66680 | AT5G66680.1 \| Symbols: DGL1 \| dolichyl-diphosphooligosaccharide-protein glycosyltransferase 48kDa subunit family protein \| chr5:26617840-26620581 REVERSE LENGTH=437 | | 0.188228208 | 0.188228208 | 5.3127 |
| AT3G20050 | AT3G20050.1 \| Symbols: ATTCP-1, TCP-1 \| T-complex protein 1 alpha subunit \| chr3:6998544-7002266 REVERSE LENGTH=545 | | 0.187856928 | 0.187856928 | 5.3232 |
| AT5G40200 | AT5G40200.1 \| Symbols: DegP9 \| DegP protease 9 \| chr5:16070402-16073101 FORWARD LENGTH=592 | | 0.187641904 | 0.187641904 | 5.3293 |
| AT1G06220 | AT1G06220.1 \| Symbols: MEE5, CLO, GFA1 \| Ribosomal protein S5/Elongation factor G/III/V family protein \| chr1:1900524-1904583 FORWARD LENGTH=987;AT1G06220.2 \| Symbols: MEE5 \| Ribosomal protein S5/Elongation factor G/III/V family protein \| chr1:1900524-1904583 FORWARD LENGTH=987;AT5G25230.1 \| Symbols: \| Ribosomal protein S5/Elongation factor G/III/V family protein \| chr5:8739709-8743594 FORWARD LENGTH=973 | 0.16833 | 0.206701255 | 0.187515627 | 5.332888859 |
| AT4G11010 | AT4G11010.1 \| Symbols: NDPK3 \| nucleoside diphosphate kinase 3 \| chr4:6732780-6734298 REVERSE LENGTH=238 | | 0.187483595 | 0.187483595 | 5.3338 |
| AT5G24650 | AT5G24650.1 \| Symbols: \| Mitochondrial import inner membrane translocase subunit Tim17/Tim22/Tim23 family protein \| chr5:8437123-8438859 FORWARD LENGTH=259 | 0.093556 | 0.281072573 | 0.187314286 | 5.338621089 |
| AT4G04020 | AT4G04020.1 \| Symbols: FIB \| fibrillin \| chr4:1932161-1933546 FORWARD LENGTH=318 | | 0.186521926 | 0.186521926 | 5.3613 |
| AT3G06650 | AT3G06650.1 \| Symbols: ACLB-1 \| ATP-citrate lyase B-1 \| chr3:2079247-2082633 REVERSE LENGTH=608;AT5G49460.1 \| Symbols: ACLB-2 \| ATP citrate lyase subunit B 2 \| chr5:20055048-20058195 FORWARD LENGTH=608 | 0.095734 | 0.277031332 | 0.186382666 | 5.365305802 |
| AT5G49890 | AT5G49890.1 \| Symbols: CLC-C, ATCLC-C \| chloride channel C \| chr5:20288489-20292143 REVERSE LENGTH=779 | | 0.185880516 | 0.185880516 | 5.3798 |

| | | | | | |
|---|---|---|---|---|---|
| AT5G14610 | AT5G14610.1 \| Symbols: \| DEAD box RNA helicase family protein \| chr5:4711271-4714713 FORWARD LENGTH=712;AT5G14610.2 \| Symbols: \| DEAD box RNA helicase family protein \| chr5:4711271-4714713 FORWARD LENGTH=645;AT3G01540.2 \| Symbols: DRH1, ATDRH1 \| DEAD box RNA helicase 1 \| chr3:213077-216142 REVERSE LENGTH=619;AT3G01540.3 \| Symbols: DRH1, ATDRH1 \| DEAD box RNA helicase 1 \| chr3:213077-216142 REVERSE LENGTH=619;AT3G01540.4 \| Symbols: DRH1, ATDRH1 \| DEAD box RNA helicase 1 \| chr3:213077-216142 REVERSE LENGTH=619;AT3G01540.1 \| Symbols: DRH1, ATDRH1 \| DEAD box RNA helicase 1 \| chr3:213077-216142 REVERSE LENGTH=618 | | 0.185435867 | 0.185435867 | 5.3927 |
| AT5G16280 | AT5G16280.1 \| Symbols: \| Tetratricopeptide repeat (TPR)-like superfamily protein \| chr5:5323377-5331345 REVERSE LENGTH=1272;AT5G16280.2 \| Symbols: \| Tetratricopeptide repeat (TPR)-like superfamily protein \| chr5:5323377-5331345 REVERSE LENGTH=1244 | | 0.184979652 | 0.184979652 | 5.406 |
| AT1G07070 | AT1G07070.1 \| Symbols: \| Ribosomal protein L35Ae family protein \| chr1:2168652-2169703 FORWARD LENGTH=112;AT1G74270.1 \| Symbols: \| Ribosomal protein L35Ae family protein \| chr1:27928415-27929466 REVERSE LENGTH=112;AT1G41880.1 \| Symbols: \| Ribosomal protein L35Ae family protein \| chr1:15651585-15652427 REVERSE LENGTH=111;AT3G55750.1 \| Symbols: \| Ribosomal protein L35Ae family protein \| chr3:20698641-20699553 FORWARD LENGTH=111 | 0.18444 | | 0.18444 | 5.421817393 |
| AT3G48890 | AT3G48890.1 \| Symbols: ATMP2, ATMAPR3, MSBP2, MAPR3 \| membrane-associated progesterone binding protein 3 \| chr3:18129669-18131353 FORWARD LENGTH=233;AT5G52240.1 \| Symbols: MSBP1, ATMP1, AtMAPR5 \| membrane steroid binding protein 1 \| chr5:21213121-21214557 FORWARD LENGTH=220;AT5G52240.2 \| Symbols: MSBP1 \| membrane steroid binding protein 1 \| chr5:21213121-21214619 FORWARD LENGTH=175 | 0.13152 | 0.235211102 | 0.183365551 | 5.453587081 |
| AT3G43520 | AT3G43520.1 \| Symbols: \| Transmembrane proteins 14C \| chr3:15406088-15407699 FORWARD LENGTH=240 | 0.072528 | 0.293530586 | 0.183029293 | 5.463606311 |
| AT5G13680 | AT5G13680.1 \| Symbols: ELO2, ABO1 \| IKI3 family protein \| chr5:4410522-4415471 REVERSE LENGTH=1319 | | 0.182671757 | 0.182671757 | 5.4743 |
| AT5G16120 | AT5G16120.2 \| Symbols: \| alpha/beta-Hydrolases superfamily protein \| chr5:5265820-5267775 FORWARD LENGTH=369;AT5G16120.1 \| Symbols: \| alpha/beta-Hydrolases superfamily protein \| chr5:5266048-5267775 FORWARD LENGTH=351 | 0.18265 | | 0.18265 | 5.474952094 |
| AT1G04690 | AT1G04690.1 \| Symbols: KAB1, KV-BETA1 \| potassium channel beta subunit 1 \| chr1:1313662-1315420 FORWARD LENGTH=328 | | 0.182548375 | 0.182548375 | 5.478 |
| AT3G46560 | AT3G46560.1 \| Symbols: TIM9, emb2474 \| Tim10/DDP family zinc finger protein \| chr3:17138632-17139301 FORWARD LENGTH=93 | | 0.182488412 | 0.182488412 | 5.4798 |
| AT5G47700 | AT5G47700.1 \| Symbols: \| 60S acidic ribosomal protein family \| chr5:19328019-19328724 REVERSE LENGTH=113;AT5G47700.2 \| Symbols: \| 60S acidic ribosomal protein family \| chr5:19328019-19328724 REVERSE LENGTH=113 | 0.2459 | 0.118985293 | 0.182442647 | 5.481174594 |
| AT1G67120 | AT1G67120.1 \| Symbols: \| ATPases;nucleotide binding;ATP binding;nucleoside-triphosphatases;transcription factor binding \| chr1:25069727-25095526 REVERSE LENGTH=5393 | | 0.182295464 | 0.182295464 | 5.4856 |
| AT5G14220 | AT5G14220.1 \| Symbols: HEMG2, MEE61, PPO2 \| Flavin containing amine oxidoreductase family \| chr5:4583506-4587369 REVERSE LENGTH=508;AT5G14220.2 \| Symbols: HEMG2, MEE61, PPO2 \| Flavin containing amine oxidoreductase family \| chr5:4583506-4587369 REVERSE LENGTH=478 | 0.078966 | 0.285551114 | 0.182258557 | 5.486710843 |
| AT4G18465 | AT4G18465.1 \| Symbols: \| RNA helicase family protein \| chr4:10197056-10201611 FORWARD LENGTH=695 | | 0.181775217 | 0.181775217 | 5.5013 |
| AT1G36180 | AT1G36180.1 \| Symbols: ACC2 \| acetyl-CoA carboxylase 2 \| chr1:13546047-13558339 FORWARD LENGTH=2355 | | 0.180241885 | 0.180241885 | 5.5481 |

| | | | | | |
|---|---|---|---|---|---|
| AT2G07698 | AT2G07698.1 \| Symbols: \| ATPase, F1 complex, alpha subunit protein \| chr2:3361474-3364028 FORWARD LENGTH=777;ATMG01190.1 \| Symbols: ATP1 \| ATP synthase subunit 1 \| chrM:302166-303689 REVERSE LENGTH=507 | 0.17109 | 0.189261313 | 0.180175657 | 5.550139343 |
| AT5G01500 | AT5G01500.1 \| Symbols: TAAC \| thylakoid ATP/ADP carrier \| chr5:199017-201329 FORWARD LENGTH=415 | 0.17978 | | 0.17978 | 5.562353988 |
| AT1G61570 | AT1G61570.1 \| Symbols: TIM13 \| translocase of the inner mitochondrial membrane 13 \| chr1:22718897-22719473 REVERSE LENGTH=87 | | 0.179391504 | 0.179391504 | 5.5744 |
| AT2G34480 | AT2G34480.1 \| Symbols: \| Ribosomal protein L18ae/LX family protein \| chr2:14532916-14534161 REVERSE LENGTH=178;AT1G29970.2 \| Symbols: RPL18AA \| 60S ribosomal protein L18A-1 \| chr1:10498458-10501311 REVERSE LENGTH=312;AT1G29965.1 \| Symbols: \| Ribosomal protein L18ae/LX family protein \| chr1:10498458-10499373 REVERSE LENGTH=178;AT3G14600.1 \| Symbols: \| Ribosomal protein L18ae/LX family protein \| chr3:4910773-4911933 FORWARD LENGTH=178 | 0.14557 | 0.21256244 | 0.17906622 | 5.584526213 |
| AT2G32730 | AT2G32730.1 \| Symbols: \| 26S proteasome regulatory complex, non-ATPase subcomplex, Rpn2/Psmd1 subunit \| chr2:13880189-13885464 FORWARD LENGTH=1004;AT1G04810.1 \| Symbols: \| 26S proteasome regulatory complex, non-ATPase subcomplex, Rpn2/Psmd1 subunit \| chr1:1350304-1355261 FORWARD LENGTH=1001 | 0.11707 | 0.240581244 | 0.178825622 | 5.592039821 |
| AT5G20920 | AT5G20920.1 \| Symbols: EIF2 BETA, EMB1401 \| eukaryotic translation initiation factor 2 beta subunit \| chr5:7094994-7096661 REVERSE LENGTH=268;AT5G20920.3 \| Symbols: EIF2 BETA \| eukaryotic translation initiation factor 2 beta subunit \| chr5:7094994-7096661 REVERSE LENGTH=268;AT5G20920.2 \| Symbols: EIF2 BETA \| eukaryotic translation initiation factor 2 beta subunit \| chr5:7094994-7096661 REVERSE LENGTH=267 | 0.1787 | | 0.1787 | 5.595970901 |
| AT5G65010 | AT5G65010.2 \| Symbols: ASN2 \| asparagine synthetase 2 \| chr5:25969224-25972278 FORWARD LENGTH=579;AT5G10240.1 \| Symbols: ASN3 \| asparagine synthetase 3 \| chr5:3212934-3216418 REVERSE LENGTH=578;AT5G65010.1 \| Symbols: ASN2 \| asparagine synthetase 2 \| chr5:25969224-25972278 FORWARD LENGTH=578;AT5G10240.2 \| Symbols: ASN3 \| asparagine synthetase 3 \| chr5:3212934-3216418 REVERSE LENGTH=577 | | 0.178616083 | 0.178616083 | 5.5986 |
| AT4G03560 | AT4G03560.1 \| Symbols: ATTPC1, TPC1, ATCCH1, FOU2 \| two-pore channel 1 \| chr4:1580253-1585691 FORWARD LENGTH=733 | | 0.177992951 | 0.177992951 | 5.6182 |
| AT1G62390 | AT1G62390.1 \| Symbols: Phox2 \| Octicosapeptide/Phox/Bem1p (PB1) domain-containing protein / tetratricopeptide repeat (TPR)-containing protein \| chr1:23084632-23086887 REVERSE LENGTH=751 | | 0.177973945 | 0.177973945 | 5.6188 |
| AT2G40290 | AT2G40290.1 \| Symbols: \| Eukaryotic translation initiation factor 2 subunit 1 \| chr2:16829030-16830889 REVERSE LENGTH=344;AT2G40290.2 \| Symbols: \| Eukaryotic translation initiation factor 2 subunit 1 \| chr2:16829872-16830889 REVERSE LENGTH=241;AT2G40290.3 \| Symbols: \| Eukaryotic translation initiation factor 2 subunit 1 \| chr2:16829872-16830889 REVERSE LENGTH=241 | 0.14373 | 0.212161073 | 0.177945536 | 5.619697018 |
| AT1G52930 | AT1G52930.1 \| Symbols: \| Ribosomal RNA processing Brix domain protein \| chr1:19711261-19713103 FORWARD LENGTH=320;AT3G15460.1 \| Symbols: \| Ribosomal RNA processing Brix domain protein \| chr3:5214279-5216188 REVERSE LENGTH=315 | | 0.177705116 | 0.177705116 | 5.6273 |

| | | | | | |
|---|---|---|---|---|---|
| AT2G34040 | AT2G34040.1 \| Symbols: \| Apoptosis inhibitory protein 5 (API5) \| chr2:14377957-14382246 REVERSE LENGTH=553;AT2G34040.2 \| Symbols: \| Apoptosis inhibitory protein 5 (API5) \| chr2:14378571-14382246 REVERSE LENGTH=442;REV__AT3G59090.2 \| Symbols: \| CONTAINS InterPro DOMAIN/s: Protein of unknown function DUF1084 (InterPro:IPR009457); BEST Arabidopsis thaliana protein match is: tobamovirus multiplication 1 (TAIR:AT4G21790.1); Has 198 Blast hits to 197 proteins in 29 species: Archae - 0; Bacteria - 0; Metazoa - 0; Fungi - 2; Plants - 190; Viruses - 0; Other Eukaryotes - 6 (source: NCBI BLink). \| chr3:21839334-21842449 FORWARD LENGTH=373;REV__AT3G59090.1 \| Symbols: \| CONTAINS InterPro DOMAIN/s: Protein of unknown function DUF1084 (InterPro:IPR009457); BEST Arabidopsis thaliana protein match is: tobamovirus multiplication 1 (TAIR:AT4G21790.1); Has 196 Blast hits to 195 proteins in 29 species: Archae - 0; Bacteria - 0; Metazoa - 0; Fungi - 2; Plants - 188; Viruses - 0; Other Eukaryotes - 6 (source: NCBI BLink). \| chr3:21839334-21842348 FORWARD LENGTH=367;REV__AT3G59090.3 \| Symbols: \| LOCATED IN: endomembrane system; EXPRESSED IN: 24 plant structures; EXPRESSED DURING: 15 growth stages; CONTAINS InterPro DOMAIN/s: Protein of unknown function DUF1084 (InterPro:IPR009457); BEST Arabidopsis thaliana protein match is: tobamovirus multiplication 1 (TAIR:AT4G21790.1); Has 35333 Blast hits to 34131 proteins in 2444 species: Archae - 798; Bacteria - 22429; Metazoa - 974; Fungi - 991; Plants - 531; Viruses - 0; Other Eukaryotes - 9610 (source: NCBI BLink). \| chr3:21839380-21842348 FORWARD LENGTH=361 | 0.10761 | 0.246998963 | 0.177304481 | 5.640015372 |
| AT1G50200 | AT1G50200.1 \| Symbols: ALATS, ACD \| Alanyl-tRNA synthetase \| chr1:18591429-18598311 REVERSE LENGTH=1003;AT1G50200.2 \| Symbols: ALATS \| Alanyl-tRNA synthetase \| chr1:18591429-18598167 REVERSE LENGTH=963 | | 0.176019151 | 0.176019151 | 5.6812 |
| AT1G02500 | AT1G02500.1 \| Symbols: SAM1, SAM-1, MAT1, AtSAM1 \| S-adenosylmethionine synthetase 1 \| chr1:519037-520218 FORWARD LENGTH=393;AT1G02500.2 \| Symbols: SAM1, SAM-1, MAT1, AtSAM1 \| S-adenosylmethionine synthetase 1 \| chr1:519037-520218 FORWARD LENGTH=393;AT3G17390.1 \| Symbols: MTO3, SAMS3, MAT4 \| S-adenosylmethionine synthetase family protein \| chr3:5952484-5953665 REVERSE LENGTH=393;AT4G01850.1 \| Symbols: SAM-2, MAT2, SAM2, AtSAM2 \| S-adenosylmethionine synthetase 2 \| chr4:796298-797479 REVERSE LENGTH=393;AT4G01850.2 \| Symbols: SAM-2, MAT2, SAM2, AtSAM2 \| S-adenosylmethionine synthetase 2 \| chr4:796298-797479 REVERSE LENGTH=393 | 0.17575 | | 0.17575 | 5.689900427 |
| AT5G40770 | AT5G40770.1 \| Symbols: ATPHB3, PHB3 \| prohibitin 3 \| chr5:16315589-16316621 REVERSE LENGTH=277;AT3G27280.1 \| Symbols: ATPHB4, PHB4 \| prohibitin 4 \| chr3:10076904-10078051 FORWARD LENGTH=279;AT3G27280.2 \| Symbols: ATPHB4, PHB4 \| prohibitin 4 \| chr3:10076904-10078051 FORWARD LENGTH=279 | 0.12063 | 0.228211506 | 0.174420753 | 5.733262709 |
| AT2G44120 | AT2G44120.2 \| Symbols: \| Ribosomal protein L30/L7 family protein \| chr2:18249227-18250417 REVERSE LENGTH=247;AT2G44120.1 \| Symbols: \| Ribosomal protein L30/L7 family protein \| chr2:18249227-18250402 REVERSE LENGTH=242;AT3G13580.1 \| Symbols: \| Ribosomal protein L30/L7 family protein \| chr3:4433809-4435109 FORWARD LENGTH=244;AT3G13580.2 \| Symbols: \| Ribosomal protein L30/L7 family protein \| chr3:4433809-4435109 FORWARD LENGTH=244;AT3G13580.3 \| Symbols: \| Ribosomal protein L30/L7 family protein \| chr3:4433809-4435109 FORWARD LENGTH=244 | 0.082325 | 0.266439305 | 0.174382153 | 5.734531804 |
| AT3G11400 | AT3G11400.2 \| Symbols: EIF3G1, ATEIF3G1 \| eukaryotic translation initiation factor 3G1 \| chr3:3578536-3580366 FORWARD LENGTH=321;AT3G11400.1 \| Symbols: EIF3G1, ATEIF3G1 \| eukaryotic translation initiation factor 3G1 \| chr3:3578536-3580366 FORWARD LENGTH=294 | | 0.173235167 | 0.173235167 | 5.7725 |

| | | | | | |
|---|---|---|---|---|---|
| AT1G36240 | AT1G36240.1 \| Symbols: \| Ribosomal protein L7Ae/L30e/S12e/Gadd45 family protein \| chr1:13614890-13616233 FORWARD LENGTH=112;AT1G77940.1 \| Symbols: \| Ribosomal protein L7Ae/L30e/S12e/Gadd45 family protein \| chr1:29304116-29305288 REVERSE LENGTH=112;AT3G18740.1 \| Symbols: \| Ribosomal protein L7Ae/L30e/S12e/Gadd45 family protein \| chr3:6453437-6453870 FORWARD LENGTH=112 | 0.17225 | | 0.17225 | 5.805515239 |
| AT3G18740 | AT3G18740.1 \| Symbols: \| Ribosomal protein L7Ae/L30e/S12e/Gadd45 family protein \| chr3:6453437-6453870 FORWARD LENGTH=112;AT1G36240.1 \| Symbols: \| Ribosomal protein L7Ae/L30e/S12e/Gadd45 family protein \| chr1:13614890-13616233 FORWARD LENGTH=112 | | 0.17027363 | 0.17027363 | 5.8729 |
| AT1G64090 | AT1G64090.2 \| Symbols: RTNLB3 \| Reticulan like protein B3 \| chr1:23789395-23790669 FORWARD LENGTH=271;AT1G64090.1 \| Symbols: RTNLB3 \| Reticulan like protein B3 \| chr1:23789395-23790669 FORWARD LENGTH=255 | 0.16998 | | 0.16998 | 5.883045064 |
| AT3G11250 | AT3G11250.1 \| Symbols: \| Ribosomal protein L10 family protein \| chr3:3521453-3522826 FORWARD LENGTH=323 | 0.16921 | | 0.16921 | 5.909816205 |
| AT1G80070 | AT1G80070.1 \| Symbols: SUS2, EMB33, EMB177, EMB14 \| Pre-mRNA-processing-splicing factor \| chr1:30118052-30127574 FORWARD LENGTH=2359;AT4G38780.1 \| Symbols: \| Pre-mRNA-processing-splicing factor \| chr4:18101438-18111029 REVERSE LENGTH=2332 | 0.2124 | 0.124296173 | 0.168348086 | 5.940073457 |
| AT3G15190 | AT3G15190.1 \| Symbols: \| chloroplast 30S ribosomal protein S20, putative \| chr3:5116216-5117412 FORWARD LENGTH=202 | | 0.167599638 | 0.167599638 | 5.9666 |
| AT3G02260 | AT3G02260.1 \| Symbols: BIG, DOC1, TIR3, UMB1, ASA1, LPR1, CRM1 \| auxin transport protein (BIG) \| chr3:431152-448489 REVERSE LENGTH=5098 | | 0.16737242 | 0.16737242 | 5.9747 |
| AT1G66070 | AT1G66070.1 \| Symbols: \| Translation initiation factor eIF3 subunit \| chr1:24596194-24597520 REVERSE LENGTH=226;AT1G66070.2 \| Symbols: \| Translation initiation factor eIF3 subunit \| chr1:24596194-24597520 REVERSE LENGTH=226 | | 0.167263239 | 0.167263239 | 5.9786 |
| AT1G26340 | AT1G26340.1 \| Symbols: B5 #6, ATCB5-A, CB5-A \| cytochrome B5 isoform A \| chr1:9113992-9114755 FORWARD LENGTH=135 | 0.16724 | | 0.16724 | 5.979430758 |
| AT4G14910 | AT4G14910.2 \| Symbols: HISN5B \| HISTIDINE BIOSYNTHESIS 5B \| chr4:8528507-8530105 REVERSE LENGTH=308;AT4G14910.3 \| Symbols: HISN5B \| HISTIDINE BIOSYNTHESIS 5B \| chr4:8528441-8530105 REVERSE LENGTH=282;AT4G14910.1 \| Symbols: HISN5B \| HISTIDINE BIOSYNTHESIS 5B \| chr4:8528451-8530105 REVERSE LENGTH=272 | | 0.167062047 | 0.167062047 | 5.9858 |
| AT2G28000 | AT2G28000.1 \| Symbols: CPN60A, CH-CPN60A, SLP \| chaperonin-60alpha \| chr2:11926603-11929184 FORWARD LENGTH=586 | 0.16673 | 0.167383627 | 0.167056813 | 5.985987524 |
| AT5G61170 | AT5G61170.1 \| Symbols: \| Ribosomal protein S19e family protein \| chr5:24611158-24612202 FORWARD LENGTH=143 | | 0.166450281 | 0.166450281 | 6.0078 |
| AT5G26830 | AT5G26830.1 \| Symbols: \| Threonyl-tRNA synthetase \| chr5:9437351-9441568 FORWARD LENGTH=709 | | 0.165920027 | 0.165920027 | 6.027 |
| AT5G27670 | AT5G27670.1 \| Symbols: HTA7 \| histone H2A 7 \| chr5:9792807-9793365 REVERSE LENGTH=150 | 0.16521 | | 0.16521 | 6.052902367 |
| AT2G38770 | AT2G38770.1 \| Symbols: EMB2765 \| P-loop containing nucleoside triphosphate hydrolases superfamily protein \| chr2:16203185-16210253 REVERSE LENGTH=1509 | | 0.164063525 | 0.164063525 | 6.0952 |
| AT1G42960 | AT1G42960.1 \| Symbols: \| expressed protein localized to the inner membrane of the chloroplast. \| chr1:16125863-16127080 FORWARD LENGTH=168 | 0.1459 | 0.180063382 | 0.162981691 | 6.135658508 |
| AT4G19210 | AT4G19210.1 \| Symbols: ATRLI2, RLI2 \| RNAse l inhibitor protein 2 \| chr4:10501906-10504776 FORWARD LENGTH=605 | 0.15423 | 0.171679714 | 0.162954857 | 6.136668875 |
| AT3G18130 | AT3G18130.1 \| Symbols: RACK1C_AT \| receptor for activated C kinase 1C \| chr3:6211109-6212371 REVERSE LENGTH=326;AT1G48630.1 \| Symbols: RACK1B_AT \| receptor for activated C kinase 1B \| chr1:17981977-17983268 REVERSE LENGTH=326 | 0.16285 | | 0.16285 | 6.140620203 |
| AT1G33590 | AT1G33590.1 \| Symbols: \| Leucine-rich repeat (LRR) family protein \| chr1:12177788-12179221 FORWARD LENGTH=477 | 0.081098 | 0.244403167 | 0.162750584 | 6.144371203 |

| | | | | | |
|---|---|---|---|---|---|
| AT1G65820 | AT1G65820.3 \| Symbols: \| microsomal glutathione s-transferase, putative \| chr1:24485213-24486682 FORWARD LENGTH=194;AT1G65820.2 \| Symbols: \| microsomal glutathione s-transferase, putative \| chr1:24485213-24486682 FORWARD LENGTH=162;AT1G65820.1 \| Symbols: \| microsomal glutathione s-transferase, putative \| chr1:24485213-24486682 FORWARD LENGTH=146 | | 0.162261273 | 0.162261273 | 6.1629 |
| AT3G55200 | AT3G55200.1 \| Symbols: \| Cleavage and polyadenylation specificity factor (CPSF) A subunit protein \| chr3:20460533-20464361 FORWARD LENGTH=1214;AT3G55220.1 \| Symbols: \| Cleavage and polyadenylation specificity factor (CPSF) A subunit protein \| chr3:20467116-20470944 REVERSE LENGTH=1214 | 0.17169 | 0.152541339 | 0.162115669 | 6.168435192 |
| AT4G16143 | AT4G16143.1 \| Symbols: IMPA-2 \| importin alpha isoform 2 \| chr4:9134450-9137134 REVERSE LENGTH=535;AT4G16143.2 \| Symbols: IMPA-2 \| importin alpha isoform 2 \| chr4:9134450-9137134 REVERSE LENGTH=535 | 0.054403 | 0.268636669 | 0.161519834 | 6.1911901 |
| AT1G66580 | AT1G66580.1 \| Symbols: SAG24, RPL10C \| senescence associated gene 24 \| chr1:24839208-24840439 FORWARD LENGTH=221 | 0.12436 | 0.198542697 | 0.161451348 | 6.193816345 |
| AT3G07230 | AT3G07230.1 \| Symbols: \| wound-responsive protein-related \| chr3:2299920-2300183 FORWARD LENGTH=46 | | 0.160346348 | 0.160346348 | 6.2365 |
| AT2G42270 | AT2G42270.1 \| Symbols: \| U5 small nuclear ribonucleoprotein helicase \| chr2:17604330-17610848 FORWARD LENGTH=2172;AT1G20960.1 \| Symbols: emb1507 \| U5 small nuclear ribonucleoprotein helicase, putative \| chr1:7302591-7309914 REVERSE LENGTH=2171;AT1G20960.2 \| Symbols: emb1507 \| U5 small nuclear ribonucleoprotein helicase, putative \| chr1:7302591-7309914 REVERSE LENGTH=2171 | 0.16 | | 0.16 | 6.25 |
| AT3G57660 | AT3G57660.1 \| Symbols: NRPA1 \| nuclear RNA polymerase A1 \| chr3:21353746-21362814 FORWARD LENGTH=1670 | | 0.159459115 | 0.159459115 | 6.2712 |
| AT2G21330 | AT2G21330.1 \| Symbols: FBA1 \| fructose-bisphosphate aldolase 1 \| chr2:9128416-9130152 REVERSE LENGTH=399;AT2G21330.3 \| Symbols: FBA1 \| fructose-bisphosphate aldolase 1 \| chr2:9128416-9130152 REVERSE LENGTH=389;AT2G21330.2 \| Symbols: FBA1 \| fructose-bisphosphate aldolase 1 \| chr2:9128762-9130152 REVERSE LENGTH=311 | 0.12389 | 0.194658569 | 0.159274284 | 6.27847743 |
| AT1G60660 | AT1G60660.1 \| Symbols: B5 #5, ATCB5LP, CB5LP \| cytochrome B5-like protein \| chr1:22342589-22342954 REVERSE LENGTH=121 | | 0.159182439 | 0.159182439 | 6.2821 |
| AT3G13930 | AT3G13930.1 \| Symbols: \| Dihydrolipoamide acetyltransferase, long form protein \| chr3:4596240-4600143 FORWARD LENGTH=539 | 0.083792 | 0.234186553 | 0.158989277 | 6.289732377 |
| AT4G27090 | AT4G27090.1 \| Symbols: \| Ribosomal protein L14 \| chr4:13594104-13595187 REVERSE LENGTH=134 | 0.1906 | 0.127124569 | 0.158862285 | 6.294760282 |
| AT4G30190 | AT4G30190.2 \| Symbols: HA2 \| H(+)-ATPase 2 \| chr4:14770820-14775920 REVERSE LENGTH=981;AT4G30190.1 \| Symbols: AHA2, PMA2, HA2 \| H(+)-ATPase 2 \| chr4:14770820-14775920 REVERSE LENGTH=948 | | 0.158657126 | 0.158657126 | 6.3029 |
| AT1G30880 | AT1G30880.1 \| Symbols: \| unknown protein; Has 24 Blast hits to 24 proteins in 8 species: Archae - 0; Bacteria - 0; Metazoa - 2; Fungi - 0; Plants - 22; Viruses - 0; Other Eukaryotes - 0 (source: NCBI BLink). \| chr1:10993379-10994056 REVERSE LENGTH=120 | 0.21451 | 0.1016033 | 0.15805665 | 6.326845468 |
| AT5G22650 | AT5G22650.1 \| Symbols: HD2B, HDT02, HDT2, ATHD2B, HDA4, HD2, ATHD2 \| histone deacetylase 2B \| chr5:7534120-7536054 FORWARD LENGTH=306;AT5G22650.2 \| Symbols: HD2B, HDT02, HDT2, ATHD2B, HDA4, HD2 \| histone deacetylase 2B \| chr5:7534120-7536054 FORWARD LENGTH=305 | | 0.157604413 | 0.157604413 | 6.345 |
| AT4G30220 | AT4G30220.2 \| Symbols: RUXF \| small nuclear ribonucleoprotein F \| chr4:14803100-14804259 REVERSE LENGTH=96;AT4G30220.1 \| Symbols: RUXF \| small nuclear ribonucleoprotein F \| chr4:14803100-14804259 REVERSE LENGTH=88;AT2G14285.1 \| Symbols: \| Small nuclear ribonucleoprotein family protein \| chr2:6052612-6053500 REVERSE LENGTH=61 | | 0.156754554 | 0.156754554 | 6.3794 |

| | | | | | |
|---|---|---|---|---|---|
| AT5G16150 | AT5G16150.1 \| Symbols: GLT1, PGLCT \| plastidic GLC translocator \| chr5:5272904-5275678 FORWARD LENGTH=546;AT5G16150.2 \| Symbols: GLT1, PGLCT \| plastidic GLC translocator \| chr5:5272904-5275678 FORWARD LENGTH=546;AT5G16150.3 \| Symbols: GLT1, PGLCT \| plastidic GLC translocator \| chr5:5272904-5275678 FORWARD LENGTH=546 | | 0.155848204 | 0.155848204 | 6.4165 |
| AT3G22230 | AT3G22230.1 \| Symbols: \| Ribosomal L27e protein family \| chr3:7844136-7844543 REVERSE LENGTH=135;AT2G32220.1 \| Symbols: \| Ribosomal L27e protein family \| chr2:13678945-13679352 FORWARD LENGTH=135 | | 0.155748684 | 0.155748684 | 6.4206 |
| AT3G23990 | AT3G23990.1 \| Symbols: HSP60, HSP60-3B \| heat shock protein 60 \| chr3:8669013-8672278 FORWARD LENGTH=577;AT2G33210.1 \| Symbols: HSP60-2 \| heat shock protein 60-2 \| chr2:14075093-14078568 REVERSE LENGTH=585;AT2G33210.2 \| Symbols: HSP60-2 \| heat shock protein 60-2 \| chr2:14075093-14078568 REVERSE LENGTH=580 | 0.10345 | 0.207623951 | 0.155536976 | 6.429339359 |
| AT5G08180 | AT5G08180.1 \| Symbols: \| Ribosomal protein L7Ae/L30e/S12e/Gadd45 family protein \| chr5:2631843-2633374 REVERSE LENGTH=156;AT5G08180.2 \| Symbols: \| Ribosomal protein L7Ae/L30e/S12e/Gadd45 family protein \| chr5:2631843-2633374 REVERSE LENGTH=156 | | 0.15521202 | 0.15521202 | 6.4428 |
| AT3G09680 | AT3G09680.1 \| Symbols: \| Ribosomal protein S12/S23 family protein \| chr3:2969197-2970291 REVERSE LENGTH=142;AT5G02960.1 \| Symbols: \| Ribosomal protein S12/S23 family protein \| chr5:693280-694396 REVERSE LENGTH=142 | | 0.154547562 | 0.154547562 | 6.4705 |
| AT3G46830 | AT3G46830.1 \| Symbols: ATRAB11A, ATRABA2C, ATRAB-A2C, RAB-A2C, RABA2c \| RAB GTPase homolog A2C \| chr3:17246699-17248362 REVERSE LENGTH=217;AT4G18430.1 \| Symbols: AtRABA1e, RABA1e \| RAB GTPase homolog A1E \| chr4:10183903-10185223 REVERSE LENGTH=217;AT5G59150.1 \| Symbols: ATRABA2D, ATRAB-A2D, RABA2D \| RAB GTPase homolog A2D \| chr5:23876858-23878244 FORWARD LENGTH=217;AT1G07410.1 \| Symbols: ATRABA2B, RAB-A2B, ATRAB-A2B, RABA2b \| RAB GTPase homolog A2B \| chr1:2276270-2277154 FORWARD LENGTH=214;AT1G06400.1 \| Symbols: ARA2, ATRABA1A, ATRAB11E, ARA-2 \| Ras-related small GTP-binding family protein \| chr1:1951089-1952686 REVERSE LENGTH=216;AT3G54840.1 \| Symbols: ARA6, ATRABF1, ARA-6, ATRAB5C \| Ras-related small GTP-binding family protein \| chr3:20318597-20320782 FORWARD LENGTH=202;AT3G54840.2 \| Symbols: ARA6, RABF1 \| Ras-related small GTP-binding family protein \| chr3:20318597-20320737 FORWARD LENGTH=193 | 0.15415 | | 0.15415 | 6.487187804 |
| AT1G09760 | AT1G09760.1 \| Symbols: U2A' \| U2 small nuclear ribonucleoprotein A \| chr1:3159476-3161603 REVERSE LENGTH=249 | 0.10946 | 0.198275007 | 0.153867504 | 6.499098093 |
| AT4G24820 | AT4G24820.1 \| Symbols: \| 26S proteasome, regulatory subunit Rpn7;Proteasome component (PCI) domain \| chr4:12790471-12792599 REVERSE LENGTH=387;AT4G24820.2 \| Symbols: \| 26S proteasome, regulatory subunit Rpn7;Proteasome component (PCI) domain \| chr4:12790471-12792599 REVERSE LENGTH=387 | 0.13121 | 0.176074937 | 0.153642469 | 6.508617104 |
| AT1G11910 | AT1G11910.1 \| Symbols: APA1, ATAPA1 \| aspartic proteinase A1 \| chr1:4017119-4019874 REVERSE LENGTH=506 | 0.19314 | 0.113864092 | 0.153502046 | 6.514571152 |
| AT3G09200 | AT3G09200.1 \| Symbols: \| Ribosomal protein L10 family protein \| chr3:2823364-2825020 REVERSE LENGTH=320;AT3G09200.2 \| Symbols: \| Ribosomal protein L10 family protein \| chr3:2823364-2825020 REVERSE LENGTH=287;AT2G40010.1 \| Symbols: \| Ribosomal protein L10 family protein \| chr2:16708578-16710448 REVERSE LENGTH=317 | 0.065793 | 0.240621767 | 0.153207383 | 6.527100576 |

| | | | | | |
|---|---|---|---|---|---|
| AT1G75630 | AT1G75630.2 \| Symbols: AVA-P4 \| vacuolar H+-pumping ATPase 16 kDa proteolipid subunit 4 \| chr1:28400662-28402032 FORWARD LENGTH=200;AT1G75630.1 \| Symbols: AVA-P4 \| vacuolar H+-pumping ATPase 16 kDa proteolipid subunit 4 \| chr1:28400662-28402032 FORWARD LENGTH=166;AT1G19910.1 \| Symbols: AVA-P2, AVA-2PE, ATVHA-C2 \| ATPase, F0/V0 complex, subunit C protein \| chr1:6913317-6914322 FORWARD LENGTH=165;AT2G16510.1 \| Symbols: \| ATPase, F0/V0 complex, subunit C protein \| chr2:7160007-7160811 REVERSE LENGTH=164;AT4G34720.1 \| Symbols: AVA-P1, VHA-C1, ATVHA-C1 \| ATPase, F0/V0 complex, subunit C protein \| chr4:16568223-16569165 REVERSE LENGTH=164;AT4G38920.1 \| Symbols: AVA-P3, ATVHA-C3, VHA-C3 \| vacuolar-type H(+)-ATPase C3 \| chr4:18147330-18148853 FORWARD LENGTH=164 | | 0.152692736 | 0.152692736 | 6.5491 |
| AT3G46740 | AT3G46740.1 \| Symbols: TOC75-III, MAR1 \| translocon at the outer envelope membrane of chloroplasts 75-III \| chr3:17216104-17219296 REVERSE LENGTH=818 | 0.068497 | 0.235521326 | 0.152009163 | 6.578550784 |
| AT4G00860 | AT4G00860.1 \| Symbols: ATOZI1, AT0ZI1 \| Protein of unknown function (DUF1138) \| chr4:359548-359868 REVERSE LENGTH=80 | | 0.151756582 | 0.151756582 | 6.5895 |
| AT1G64790 | AT1G64790.1 \| Symbols: ILA \| ILITYHIA \| chr1:24065232-24081908 REVERSE LENGTH=2610;AT1G64790.2 \| Symbols: ILA \| ILITYHIA \| chr1:24065232-24083403 REVERSE LENGTH=2696 | | 0.151620827 | 0.151620827 | 6.5954 |
| AT2G16360 | AT2G16360.1 \| Symbols: \| Ribosomal protein S25 family protein \| chr2:7076713-7077366 REVERSE LENGTH=125;AT2G21580.1 \| Symbols: \| Ribosomal protein S25 family protein \| chr2:9236629-9237510 FORWARD LENGTH=108;AT4G34555.1 \| Symbols: \| Ribosomal protein S25 family protein \| chr4:16504381-16505339 REVERSE LENGTH=108;AT4G39200.1 \| Symbols: \| Ribosomal protein S25 family protein \| chr4:18257464-18258464 FORWARD LENGTH=108;AT2G21580.2 \| Symbols: \| Ribosomal protein S25 family protein \| chr2:9236629-9237510 FORWARD LENGTH=107;AT4G39200.2 \| Symbols: \| Ribosomal protein S25 family protein \| chr4:18257464-18258464 FORWARD LENGTH=107 | 0.15136 | | 0.15136 | 6.606765328 |
| AT1G20960 | AT1G20960.1 \| Symbols: emb1507 \| U5 small nuclear ribonucleoprotein helicase, putative \| chr1:7302591-7309914 REVERSE LENGTH=2171;AT1G20960.2 \| Symbols: emb1507 \| U5 small nuclear ribonucleoprotein helicase, putative \| chr1:7302591-7309914 REVERSE LENGTH=2171;AT2G42270.1 \| Symbols: \| U5 small nuclear ribonucleoprotein helicase \| chr2:17604330-17610848 FORWARD LENGTH=2172 | | 0.15006678 | 0.15006678 | 6.6637 |
| AT4G13340 | AT4G13340.1 \| Symbols: \| Leucine-rich repeat (LRR) family protein \| chr4:7758610-7760892 FORWARD LENGTH=760;AT3G24480.1 \| Symbols: \| Leucine-rich repeat (LRR) family protein \| chr3:8901154-8902638 REVERSE LENGTH=494;AT4G18670.1 \| Symbols: \| Leucine-rich repeat (LRR) family protein \| chr4:10275918-10278491 REVERSE LENGTH=857;AT1G62440.1 \| Symbols: LRX2 \| leucine-rich repeat/extensin 2 \| chr1:23111818-23115293 FORWARD LENGTH=826 | | 0.149779076 | 0.149779076 | 6.6765 |
| AT3G57150 | AT3G57150.1 \| Symbols: NAP57, AtNAP57, CBF5, AtCBF5 \| homologue of NAP57 \| chr3:21154255-21155952 REVERSE LENGTH=565 | | 0.149452257 | 0.149452257 | 6.6911 |
| AT4G27585 | AT4G27585.1 \| Symbols: \| SPFH/Band 7/PHB domain-containing membrane-associated protein family \| chr4:13766984-13769832 REVERSE LENGTH=411 | 0.075869 | 0.222588256 | 0.149228628 | 6.701127073 |
| AT4G39200 | AT4G39200.1 \| Symbols: \| Ribosomal protein S25 family protein \| chr4:18257464-18258464 FORWARD LENGTH=108;AT4G39200.2 \| Symbols: \| Ribosomal protein S25 family protein \| chr4:18257464-18258464 FORWARD LENGTH=107 | | 0.14831954 | 0.14831954 | 6.7422 |
| AT5G27540 | AT5G27540.1 \| Symbols: MIRO1, emb2473 \| MIRO-related GTP-ase 1 \| chr5:9722816-9727112 FORWARD LENGTH=648;AT5G27540.2 \| Symbols: MIRO1 \| MIRO-related GTP-ase 1 \| chr5:9722816-9727112 FORWARD LENGTH=648 | | 0.147939936 | 0.147939936 | 6.7595 |

| | | | | | |
|---|---|---|---|---|---|
| AT1G08360 | AT1G08360.1 \| Symbols: \| Ribosomal protein L1p/L10e family \| chr1:2636231-2637694 FORWARD LENGTH=216 | | 0.147570982 | 0.147570982 | 6.7764 |
| AT4G30990 | AT4G30990.2 \| Symbols: \| ARM repeat superfamily protein \| chr4:15084456-15097860 FORWARD LENGTH=2620;AT4G30990.1 \| Symbols: \| ARM repeat superfamily protein \| chr4:15084456-15097860 FORWARD LENGTH=2599 | | 0.147331821 | 0.147331821 | 6.7874 |
| AT5G53560 | AT5G53560.1 \| Symbols: ATB5-A, B5 #2, ATCB5-E, CB5-E \| cytochrome B5 isoform E \| chr5:21759628-21760353 FORWARD LENGTH=134 | 0.18652 | 0.107620616 | 0.147070308 | 6.799469004 |
| AT2G19730 | AT2G19730.1 \| Symbols: \| Ribosomal L28e protein family \| chr2:8511752-8512995 FORWARD LENGTH=143;AT2G19730.2 \| Symbols: \| Ribosomal L28e protein family \| chr2:8511752-8512995 FORWARD LENGTH=143;AT2G19730.3 \| Symbols: \| Ribosomal L28e protein family \| chr2:8511752-8512995 FORWARD LENGTH=143 | | 0.147054498 | 0.147054498 | 6.8002 |
| AT1G18630 | AT1G18630.1 \| Symbols: GR-RBP6 \| glycine-rich RNA-binding protein 6 \| chr1:6415226-6416283 FORWARD LENGTH=155 | | 0.146271538 | 0.146271538 | 6.8366 |
| AT5G19760 | AT5G19760.1 \| Symbols: \| Mitochondrial substrate carrier family protein \| chr5:6679591-6681845 REVERSE LENGTH=298 | | 0.14522219 | 0.14522219 | 6.886 |
| AT1G33810 | AT1G33810.1 \| Symbols: \| unknown protein; FUNCTIONS IN: molecular_function unknown; INVOLVED IN: biological_process unknown; LOCATED IN: chloroplast thylakoid membrane, chloroplast, chloroplast envelope; EXPRESSED IN: 22 plant structures; EXPRESSED DURING: 13 growth stages; Has 39 Blast hits to 39 proteins in 18 species: Archae - 0; Bacteria - 0; Metazoa - 0; Fungi - 0; Plants - 39; Viruses - 0; Other Eukaryotes - 0 (source: NCBI BLink). \| chr1:12265084-12266655 FORWARD LENGTH=138 | 0.17088 | 0.119350257 | 0.145115129 | 6.891080273 |
| AT1G07920 | AT1G07920.1 \| Symbols: \| GTP binding Elongation factor Tu family protein \| chr1:2455559-2457001 FORWARD LENGTH=449;AT1G07930.1 \| Symbols: \| GTP binding Elongation factor Tu family protein \| chr1:2459014-2460458 FORWARD LENGTH=449;AT1G07940.1 \| Symbols: \| GTP binding Elongation factor Tu family protein \| chr1:2463350-2464792 REVERSE LENGTH=449;AT1G07940.2 \| Symbols: \| GTP binding Elongation factor Tu family protein \| chr1:2463350-2464792 REVERSE LENGTH=449;AT5G60390.1 \| Symbols: \| GTP binding Elongation factor Tu family protein \| chr5:24289226-24290675 FORWARD LENGTH=449;AT5G60390.3 \| Symbols: \| GTP binding Elongation factor Tu family protein \| chr5:24289226-24290675 FORWARD LENGTH=449;AT1G07930.2 \| Symbols: \| GTP binding Elongation factor Tu family protein \| chr1:2459014-2460458 FORWARD LENGTH=372;AT5G60390.2 \| Symbols: \| GTP binding Elongation factor Tu family protein \| chr5:24289226-24290577 FORWARD LENGTH=400 | 0.080337 | 0.209815153 | 0.145076076 | 6.892935242 |
| AT3G15430 | AT3G15430.1 \| Symbols: \| Regulator of chromosome condensation (RCC1) family protein \| chr3:5209408-5211562 FORWARD LENGTH=488;AT3G15430.2 \| Symbols: \| Regulator of chromosome condensation (RCC1) family protein \| chr3:5209408-5211562 FORWARD LENGTH=488 | 0.14471 | | 0.14471 | 6.910372469 |
| AT5G25460 | AT5G25460.1 \| Symbols: \| Protein of unknown function, DUF642 \| chr5:8863430-8865394 FORWARD LENGTH=369;AT5G11420.1 \| Symbols: \| Protein of unknown function, DUF642 \| chr5:3644655-3646991 FORWARD LENGTH=366 | | 0.144370976 | 0.144370976 | 6.9266 |

| | | | | | |
|---|---|---|---|---|---|
| AT3G13920 | AT3G13920.2 \| Symbols: EIF4A1, RH4, TIF4A1 \| eukaryotic translation initiation factor 4A1 \| chr3:4592586-4594128 REVERSE LENGTH=415;AT1G54270.1 \| Symbols: EIF4A-2 \| eif4a-2 \| chr1:20260495-20262018 FORWARD LENGTH=412;AT3G13920.1 \| Symbols: EIF4A1, RH4, TIF4A1 \| eukaryotic translation initiation factor 4A1 \| chr3:4592635-4594128 REVERSE LENGTH=412;AT1G54270.2 \| Symbols: EIF4A-2 \| eif4a-2 \| chr1:20260495-20262018 FORWARD LENGTH=407;AT3G13920.4 \| Symbols: EIF4A1 \| eukaryotic translation initiation factor 4A1 \| chr3:4592635-4594128 REVERSE LENGTH=407;AT3G13920.3 \| Symbols: EIF4A1, RH4, TIF4A1 \| eukaryotic translation initiation factor 4A1 \| chr3:4592635-4594094 REVERSE LENGTH=402;AT1G72730.1 \| Symbols: \| DEA(D/H)-box RNA helicase family protein \| chr1:27378040-27379593 REVERSE LENGTH=414 | 0.090498 | 0.198145359 | 0.14432168 | 6.928965918 |
| AT1G55490 | AT1G55490.1 \| Symbols: CPN60B, LEN1 \| chaperonin 60 beta \| chr1:20715717-20718673 REVERSE LENGTH=600;AT1G55490.2 \| Symbols: CPN60B, LEN1 \| chaperonin 60 beta \| chr1:20715717-20718673 REVERSE LENGTH=600;AT5G56500.1 \| Symbols: \| TCP-1/cpn60 chaperonin family protein \| chr5:22874058-22876966 FORWARD LENGTH=597;AT5G56500.2 \| Symbols: \| TCP-1/cpn60 chaperonin family protein \| chr5:22874058-22876966 FORWARD LENGTH=597;AT3G13470.1 \| Symbols: \| TCP-1/cpn60 chaperonin family protein \| chr3:4389685-4392624 FORWARD LENGTH=596 | 0.040259 | 0.245682136 | 0.142970568 | 6.994446566 |
| AT1G56070 | AT1G56070.1 \| Symbols: LOS1 \| Ribosomal protein S5/Elongation factor G/III/V family protein \| chr1:20968245-20971077 REVERSE LENGTH=843 | 0.11344 | 0.170564055 | 0.142002028 | 7.042152964 |
| AT1G23290 | AT1G23290.1 \| Symbols: RPL27A, RPL27AB \| Ribosomal protein L18e/L15 superfamily protein \| chr1:8263007-8263447 FORWARD LENGTH=146 | 0.10088 | 0.181274359 | 0.141077179 | 7.088318638 |
| AT1G70770 | AT1G70770.1 \| Symbols: \| Protein of unknown function DUF2359, transmembrane \| chr1:26688622-26691185 REVERSE LENGTH=610;AT1G70770.2 \| Symbols: \| Protein of unknown function DUF2359, transmembrane \| chr1:26688622-26691185 REVERSE LENGTH=610 | | 0.140827219 | 0.140827219 | 7.1009 |
| AT2G03510 | AT2G03510.1 \| Symbols: \| SPFH/Band 7/PHB domain-containing membrane-associated protein family \| chr2:1066717-1068934 FORWARD LENGTH=356 | 0.05945 | 0.220960293 | 0.140205147 | 7.132405788 |
| AT5G50850 | AT5G50850.1 \| Symbols: MAB1 \| Transketolase family protein \| chr5:20689671-20692976 FORWARD LENGTH=363 | 0.14004 | 0.13981321 | 0.139926605 | 7.146603761 |
| AT1G06530 | AT1G06530.1 \| Symbols: \| Tropomyosin-related \| chr1:2001625-2002596 FORWARD LENGTH=323 | 0.13888 | | 0.13888 | 7.200460829 |
| AT5G42020 | AT5G42020.1 \| Symbols: BIP, BIP2 \| Heat shock protein 70 (Hsp 70) family protein \| chr5:16807697-16810480 REVERSE LENGTH=668;AT5G42020.2 \| Symbols: BIP \| Heat shock protein 70 (Hsp 70) family protein \| chr5:16807697-16810480 REVERSE LENGTH=613 | 0.093546 | 0.1833315 | 0.13843875 | 7.223411075 |
| AT2G47250 | AT2G47250.1 \| Symbols: \| RNA helicase family protein \| chr2:19399923-19402981 REVERSE LENGTH=729;AT3G62310.1 \| Symbols: \| RNA helicase family protein \| chr3:23057516-23060561 REVERSE LENGTH=726 | | 0.138333633 | 0.138333633 | 7.2289 |
| AT3G11510 | AT3G11510.1 \| Symbols: \| Ribosomal protein S11 family protein \| chr3:3623757-3624866 REVERSE LENGTH=150 | 0.14036 | 0.133008792 | 0.136684396 | 7.316124076 |
| AT5G47030 | AT5G47030.1 \| Symbols: \| ATPase, F1 complex, delta/epsilon subunit \| chr5:19090384-19092034 FORWARD LENGTH=203 | 0.1363 | | 0.1363 | 7.336757153 |
| AT2G21640 | AT2G21640.1 \| Symbols: \| Encodes a protein of unknown function that is a marker for oxidative stress response. \| chr2:9254985-9255453 FORWARD LENGTH=104 | 0.1357 | | 0.1357 | 7.369196758 |
| AT1G31970 | AT1G31970.1 \| Symbols: STRS1 \| DEA(D/H)-box RNA helicase family protein \| chr1:11479921-11482707 FORWARD LENGTH=537 | 0.06114 | 0.209661188 | 0.135400594 | 7.385491985 |
| AT1G04510 | AT1G04510.1 \| Symbols: MAC3A \| MOS4-associated complex 3A \| chr1:1226749-1230592 FORWARD LENGTH=523;AT1G04510.2 \| Symbols: MAC3A \| MOS4-associated complex 3A \| chr1:1226749-1230592 FORWARD LENGTH=523 | | 0.134417636 | 0.134417636 | 7.4395 |
| AT5G47930 | AT5G47930.1 \| Symbols: \| Zinc-binding ribosomal protein family protein \| chr5:19406423-19407329 REVERSE LENGTH=84 | | 0.134262429 | 0.134262429 | 7.4481 |

| | | | | | |
|---|---|---|---|---|---|
| AT4G30810 | AT4G30810.1 \| Symbols: scpl29 \| serine carboxypeptidase-like 29 \| chr4:15003474-15006017 FORWARD LENGTH=479 | | 0.133806115 | 0.133806115 | 7.4735 |
| AT2G39460 | AT2G39460.1 \| Symbols: ATRPL23A, RPL23A, RPL23AA \| ribosomal protein L23AA \| chr2:16475049-16475904 FORWARD LENGTH=154;AT2G39460.2 \| Symbols: ATRPL23A, RPL23A, RPL23AA \| ribosomal protein L23AA \| chr2:16475049-16475904 FORWARD LENGTH=154;AT3G55280.1 \| Symbols: RPL23AB \| ribosomal protein L23AB \| chr3:20500667-20501519 FORWARD LENGTH=154;AT3G55280.2 \| Symbols: RPL23AB \| ribosomal protein L23AB \| chr3:20500667-20501519 FORWARD LENGTH=154;AT3G55280.3 \| Symbols: RPL23AB \| ribosomal protein L23AB \| chr3:20500667-20501519 FORWARD LENGTH=148 | 0.13882 | 0.128676944 | 0.133748472 | 7.47672094 |
| AT1G77940 | AT1G77940.1 \| Symbols: \| Ribosomal protein L7Ae/L30e/S12e/Gadd45 family protein \| chr1:29304116-29305288 REVERSE LENGTH=112 | | 0.132613683 | 0.132613683 | 7.5407 |
| AT1G03230 | AT1G03230.1 \| Symbols: \| Eukaryotic aspartyl protease family protein \| chr1:790110-791414 FORWARD LENGTH=434 | 0.019443 | 0.244893961 | 0.13216848 | 7.566100454 |
| AT2G07707 | AT2G07707.1 \| Symbols: \| Plant mitochondrial ATPase, F0 complex, subunit 8 protein \| chr2:3386292-3386768 FORWARD LENGTH=158;ATMG00480.1 \| Symbols: ORFB, ATP8 \| Plant mitochondrial ATPase, F0 complex, subunit 8 protein \| chrM:129909-130385 FORWARD LENGTH=158 | | 0.13191916 | 0.13191916 | 7.5804 |
| AT1G48460 | AT1G48460.1 \| Symbols: \| unknown protein; FUNCTIONS IN: molecular_function unknown; INVOLVED IN: biological_process unknown; LOCATED IN: chloroplast envelope; EXPRESSED IN: 21 plant structures; EXPRESSED DURING: 13 growth stages; BEST Arabidopsis thaliana protein match is: unknown protein (TAIR:AT5G63040.1); Has 60 Blast hits to 60 proteins in 14 species: Archae - 0; Bacteria - 0; Metazoa - 0; Fungi - 0; Plants - 60; Viruses - 0; Other Eukaryotes - 0 (source: NCBI BLink). \| chr1:17911469-17913149 FORWARD LENGTH=340 | | 0.131566829 | 0.131566829 | 7.6007 |
| AT1G07670 | AT1G07670.1 \| Symbols: ATECA4, ECA4 \| endomembrane-type CA-ATPase 4 \| chr1:2370305-2374196 REVERSE LENGTH=1061;AT4G00900.1 \| Symbols: ECA2, ATECA2 \| ER-type Ca2+-ATPase 2 \| chr4:382690-386226 REVERSE LENGTH=1054 | | 0.131428496 | 0.131428496 | 7.6087 |
| AT3G13300 | AT3G13300.1 \| Symbols: VCS \| Transducin/WD40 repeat-like superfamily protein \| chr3:4304085-4309949 FORWARD LENGTH=1344;AT3G13300.2 \| Symbols: VCS \| Transducin/WD40 repeat-like superfamily protein \| chr3:4304085-4309949 FORWARD LENGTH=1309;AT3G13290.1 \| Symbols: VCR \| varicose-related \| chr3:4297529-4303113 FORWARD LENGTH=1340 | | 0.131224985 | 0.131224985 | 7.6205 |
| AT5G48810 | AT5G48810.1 \| Symbols: ATB5-B, B5 #3, ATCB5-D, CB5-D \| cytochrome B5 isoform D \| chr5:19789249-19790180 REVERSE LENGTH=140 | 0.13012 | | 0.13012 | 7.685213649 |
| AT3G06530 | AT3G06530.2 \| Symbols: \| ARM repeat superfamily protein \| chr3:2022596-2033637 FORWARD LENGTH=2199;AT3G06530.1 \| Symbols: \| ARM repeat superfamily protein \| chr3:2022596-2033637 FORWARD LENGTH=2197;AT3G06530.3 \| Symbols: \| ARM repeat superfamily protein \| chr3:2022596-2033637 FORWARD LENGTH=2188 | | 0.1296546 | 0.1296546 | 7.7128 |
| AT3G08580 | AT3G08580.1 \| Symbols: AAC1 \| ADP/ATP carrier 1 \| chr3:2605706-2607030 REVERSE LENGTH=381;AT3G08580.2 \| Symbols: AAC1 \| ADP/ATP carrier 1 \| chr3:2605706-2607030 REVERSE LENGTH=381 | | 0.129587394 | 0.129587394 | 7.7168 |
| AT3G44890 | AT3G44890.1 \| Symbols: RPL9 \| ribosomal protein L9 \| chr3:16386505-16387963 FORWARD LENGTH=197 | | 0.129587394 | 0.129587394 | 7.7168 |
| AT5G43970 | AT5G43970.1 \| Symbols: TOM22-V, TOM9-2, ATTOM22-V \| translocase of outer membrane 22-V \| chr5:17692888-17693187 FORWARD LENGTH=99 | | 0.128617363 | 0.128617363 | 7.775 |
| AT3G47520 | AT3G47520.1 \| Symbols: MDH \| malate dehydrogenase \| chr3:17513657-17514868 FORWARD LENGTH=403 | 0.094469 | 0.160632249 | 0.127550624 | 7.840024349 |
| AT3G54110 | AT3G54110.1 \| Symbols: ATPUMP1, UCP, PUMP1, ATUCP1, UCP1 \| plant uncoupling mitochondrial protein 1 \| chr3:20038890-20040996 FORWARD LENGTH=306 | 0.11719 | 0.135358293 | 0.126274147 | 7.91927743 |

| | | | | | |
|---|---|---|---|---|---|
| AT3G51260 | AT3G51260.1 \| Symbols: PAD1 \| 20S proteasome alpha subunit PAD1 \| chr3:19031086-19032746 FORWARD LENGTH=250;AT3G51260.2 \| Symbols: PAD1 \| 20S proteasome alpha subunit PAD1 \| chr3:19031086-19032746 FORWARD LENGTH=243;AT5G66140.1 \| Symbols: PAD2 \| proteasome alpha subunit D2 \| chr5:26437445-26438677 REVERSE LENGTH=250 | 0.12497 | | 0.12497 | 8.001920461 |
| AT1G15930 | AT1G15930.1 \| Symbols: \| Ribosomal protein L7Ae/L30e/S12e/Gadd45 family protein \| chr1:5471702-5472741 FORWARD LENGTH=144;AT1G15930.2 \| Symbols: \| Ribosomal protein L7Ae/L30e/S12e/Gadd45 family protein \| chr1:5471702-5472741 FORWARD LENGTH=144 | | 0.123881965 | 0.123881965 | 8.0722 |
| AT4G22240 | AT4G22240.1 \| Symbols: \| Plastid-lipid associated protein PAP / fibrillin family protein \| chr4:11766090-11767227 REVERSE LENGTH=310;AT4G04020.1 \| Symbols: FIB \| fibrillin \| chr4:1932161-1933546 FORWARD LENGTH=318 | 0.074416 | 0.171227013 | 0.122821506 | 8.141896557 |
| AT4G31300 | AT4G31300.2 \| Symbols: PBA1 \| N-terminal nucleophile aminohydrolases (Ntn hydrolases) superfamily protein \| chr4:15188927-15190935 FORWARD LENGTH=234;AT4G31300.1 \| Symbols: PBA1 \| N-terminal nucleophile aminohydrolases (Ntn hydrolases) superfamily protein \| chr4:15188927-15190935 FORWARD LENGTH=233;AT4G31300.3 \| Symbols: PBA1 \| N-terminal nucleophile aminohydrolases (Ntn hydrolases) superfamily protein \| chr4:15188927-15190935 FORWARD LENGTH=233 | | 0.1221568 | 0.1221568 | 8.1862 |
| AT5G16660 | AT5G16660.1 \| Symbols: \| unknown protein; FUNCTIONS IN: molecular_function unknown; INVOLVED IN: biological_process unknown; LOCATED IN: chloroplast thylakoid membrane, chloroplast, membrane, chloroplast envelope; EXPRESSED IN: 23 plant structures; EXPRESSED DURING: 14 growth stages; BEST Arabidopsis thaliana protein match is: unknown protein (TAIR:AT3G02900.1); Has 30201 Blast hits to 17322 proteins in 780 species: Archae - 12; Bacteria - 1396; Metazoa - 17338; Fungi - 3422; Plants - 5037; Viruses - 0; Other Eukaryotes - 2996 (source: NCBI BLink). \| chr5:5465699-5467006 REVERSE LENGTH=168;AT5G16660.2 \| Symbols: \| unknown protein; FUNCTIONS IN: molecular_function unknown; INVOLVED IN: biological_process unknown; LOCATED IN: chloroplast, membrane; EXPRESSED IN: 23 plant structures; EXPRESSED DURING: 14 growth stages; BEST Arabidopsis thaliana protein match is: unknown protein (TAIR:AT3G02900.1); Has 106 Blast hits to 106 proteins in 32 species: Archae - 0; Bacteria - 28; Metazoa - 0; Fungi - 0; Plants - 78; Viruses - 0; Other Eukaryotes - 0 (source: NCBI BLink). \| chr5:5465699-5467006 REVERSE LENGTH=166 | 0.11716 | 0.126478214 | 0.121819107 | 8.208892875 |
| AT5G64270 | AT5G64270.1 \| Symbols: \| splicing factor, putative \| chr5:25706909-25710718 FORWARD LENGTH=1269 | | 0.121216529 | 0.121216529 | 8.2497 |
| AT5G61790 | AT5G61790.1 \| Symbols: CNX1, ATCNX1 \| calnexin 1 \| chr5:24827394-24829642 REVERSE LENGTH=530;AT5G07340.2 \| Symbols: \| Calreticulin family protein \| chr5:2317300-2319458 FORWARD LENGTH=540;AT5G07340.1 \| Symbols: \| Calreticulin family protein \| chr5:2317300-2319458 FORWARD LENGTH=532 | 0.061152 | 0.18123165 | 0.121191825 | 8.251381632 |
| AT1G53850 | AT1G53850.1 \| Symbols: PAE1, ATPAE1 \| 20S proteasome alpha subunit E1 \| chr1:20104131-20105792 REVERSE LENGTH=237;AT1G53850.2 \| Symbols: PAE1 \| 20S proteasome alpha subunit E1 \| chr1:20104131-20105792 REVERSE LENGTH=237;AT3G14290.1 \| Symbols: PAE2 \| 20S proteasome alpha subunit E2 \| chr3:4764364-4766381 FORWARD LENGTH=237 | 0.12061 | | 0.12061 | 8.291186469 |
| AT1G55210 | AT1G55210.1 \| Symbols: \| Disease resistance-responsive (dirigent-like protein) family protein \| chr1:20598057-20598620 REVERSE LENGTH=187;AT1G55210.2 \| Symbols: \| Disease resistance-responsive (dirigent-like protein) family protein \| chr1:20598057-20598620 REVERSE LENGTH=187 | 0.12042 | | 0.12042 | 8.304268394 |
| AT5G25100 | AT5G25100.2 \| Symbols: \| Endomembrane protein 70 protein family \| chr5:8648374-8651015 REVERSE LENGTH=651;AT5G25100.1 \| Symbols: \| Endomembrane protein 70 protein family \| chr5:8648374-8651015 REVERSE LENGTH=644 | | 0.120277601 | 0.120277601 | 8.3141 |

| | | | | | |
|---|---|---|---|---|---|
| AT3G02090 | AT3G02090.2 \| Symbols: MPPBETA \| Insulinase (Peptidase family M16) protein \| chr3:365624-368534 FORWARD LENGTH=535;AT3G02090.1 \| Symbols: MPPBETA \| Insulinase (Peptidase family M16) protein \| chr3:365624-368526 FORWARD LENGTH=531 | 0.06102 | 0.179533214 | 0.120276607 | 8.314168702 |
| AT4G31985 | AT4G31985.1 \| Symbols: \| Ribosomal protein L39 family protein \| chr4:15469931-15470366 FORWARD LENGTH=51;AT2G25210.1 \| Symbols: \| Ribosomal protein L39 family protein \| chr2:10740157-10740379 FORWARD LENGTH=44 | | 0.119284769 | 0.119284769 | 8.3833 |
| AT5G64420 | AT5G64420.1 \| Symbols: \| DNA polymerase V family \| chr5:25756416-25761122 FORWARD LENGTH=1306 | 0.11883 | | 0.11883 | 8.415383321 |
| AT5G64140 | AT5G64140.1 \| Symbols: RPS28 \| ribosomal protein S28 \| chr5:25667529-25667723 REVERSE LENGTH=64 | | 0.118090245 | 0.118090245 | 8.4681 |
| AT1G18080 | AT1G18080.1 \| Symbols: ATARCA, RACK1A_AT, RACK1A \| Transducin/WD40 repeat-like superfamily protein \| chr1:6222325-6223901 FORWARD LENGTH=327 | | 0.117702448 | 0.117702448 | 8.496 |
| AT4G00100 | AT4G00100.1 \| Symbols: ATRPS13A, RPS13, PFL2, RPS13A \| ribosomal protein S13A \| chr4:37172-38123 FORWARD LENGTH=151 | | 0.116211505 | 0.116211505 | 8.605 |
| AT2G18740 | AT2G18740.1 \| Symbols: \| Small nuclear ribonucleoprotein family protein \| chr2:8123334-8124710 FORWARD LENGTH=88;AT4G30330.1 \| Symbols: \| Small nuclear ribonucleoprotein family protein \| chr4:14836773-14837919 REVERSE LENGTH=88;AT2G18740.2 \| Symbols: \| Small nuclear ribonucleoprotein family protein \| chr2:8123334-8124581 FORWARD LENGTH=78 | | 0.115928588 | 0.115928588 | 8.626 |
| AT3G10090 | AT3G10090.1 \| Symbols: \| Nucleic acid-binding, OB-fold-like protein \| chr3:3108960-3109154 REVERSE LENGTH=64;AT5G03850.1 \| Symbols: \| Nucleic acid-binding, OB-fold-like protein \| chr5:1028542-1028736 REVERSE LENGTH=64 | 0.16449 | 0.06643194 | 0.11546097 | 8.660935361 |
| AT5G20090 | AT5G20090.1 \| Symbols: \| Uncharacterised protein family (UPF0041) \| chr5:6787246-6788000 REVERSE LENGTH=110;AT5G20090.2 \| Symbols: \| Uncharacterised protein family (UPF0041) \| chr5:6787246-6788000 REVERSE LENGTH=110;AT5G20090.3 \| Symbols: \| Uncharacterised protein family (UPF0041) \| chr5:6787543-6788000 REVERSE LENGTH=97 | 0.071037 | 0.158677266 | 0.114857133 | 8.706468397 |
| AT1G21720 | AT1G21720.1 \| Symbols: PBC1 \| proteasome beta subunit C1 \| chr1:7626394-7628070 FORWARD LENGTH=204 | 0.11481 | | 0.11481 | 8.710042679 |
| AT5G03345 | AT5G03345.1 \| Symbols: \| FUNCTIONS IN: molecular_function unknown; INVOLVED IN: biological_process unknown; LOCATED IN: endomembrane system; EXPRESSED IN: 24 plant structures; EXPRESSED DURING: 15 growth stages; CONTAINS InterPro DOMAIN/s: Magnesium transporter (InterPro:IPR018937); Has 30201 Blast hits to 17322 proteins in 780 species: Archae - 12; Bacteria - 1396; Metazoa - 17338; Fungi - 3422; Plants - 5037; Viruses - 0; Other Eukaryotes - 2996 (source: NCBI BLink). \| chr5:814196-815478 FORWARD LENGTH=104;AT5G03345.2 \| Symbols: \| FUNCTIONS IN: molecular_function unknown; INVOLVED IN: biological_process unknown; LOCATED IN: endomembrane system; EXPRESSED IN: 24 plant structures; EXPRESSED DURING: 15 growth stages; CONTAINS InterPro DOMAIN/s: Magnesium transporter (InterPro:IPR018937); Has 155 Blast hits to 155 proteins in 63 species: Archae - 0; Bacteria - 0; Metazoa - 105; Fungi - 0; Plants - 44; Viruses - 0; Other Eukaryotes - 6 (source: NCBI BLink). \| chr5:814196-815478 FORWARD LENGTH=104 | | 0.114092735 | 0.114092735 | 8.7648 |
| AT3G22370 | AT3G22370.1 \| Symbols: AOX1A, ATAOX1A \| alternative oxidase 1A \| chr3:7906890-7908416 FORWARD LENGTH=354;AT3G27620.1 \| Symbols: AOX1C \| alternative oxidase 1C \| chr3:10229289-10230595 REVERSE LENGTH=329;AT3G22360.1 \| Symbols: AOX1B \| alternative oxidase 1B \| chr3:7904156-7905384 FORWARD LENGTH=325 | 0.10638 | 0.12092922 | 0.11365461 | 8.798587224 |

| | | | | | |
|---|---|---|---|---|---|
| AT1G78570 | AT1G78570.1 \| Symbols: RHM1, ROL1, ATRHM1 \| rhamnose biosynthesis 1 \| chr1:29550110-29552207 FORWARD LENGTH=669;AT1G53500.1 \| Symbols: MUM4, RHM2, ATRHM2, ATMUM4 \| NAD-dependent epimerase/dehydratase family protein \| chr1:19967157-19969239 REVERSE LENGTH=667;AT3G14790.1 \| Symbols: RHM3, ATRHM3 \| rhamnose biosynthesis 3 \| chr3:4964791-4966875 FORWARD LENGTH=664 | 0.1131 | | 0.1131 | 8.84173298 |
| AT1G06950 | AT1G06950.1 \| Symbols: ATTIC110, TIC110 \| translocon at the inner envelope membrane of chloroplasts 110 \| chr1:2130303-2135563 REVERSE LENGTH=1016 | 0.062097 | 0.16336666 | 0.11273183 | 8.870609121 |
| AT2G07440 | AT2G07440.1 \| Symbols: \| two-component responsive regulator-related / response regulator protein-related \| chr2:3093821-3094750 REVERSE LENGTH=136 | | 0.112090031 | 0.112090031 | 8.9214 |
| ATCG00120 | ATCG00120.1 \| Symbols: ATPA \| ATP synthase subunit alpha \| chrC:9938-11461 REVERSE LENGTH=507 | 0.049326 | 0.174532254 | 0.111929127 | 8.934224976 |
| AT1G14320 | AT1G14320.1 \| Symbols: SAC52, RPL10, RPL10A \| Ribosomal protein L16p/L10e family protein \| chr1:4888270-4889408 FORWARD LENGTH=220;AT1G26910.1 \| Symbols: RPL10B \| Ribosomal protein L16p/L10e family protein \| chr1:9321709-9322813 FORWARD LENGTH=221;AT1G14320.2 \| Symbols: SAC52, RPL10A \| Ribosomal protein L16p/L10e family protein \| chr1:4888270-4889408 FORWARD LENGTH=163 | 0.048353 | 0.174788506 | 0.111570753 | 8.962922393 |
| AT2G20490 | AT2G20490.1 \| Symbols: NOP10, EDA27 \| nucleolar RNA-binding Nop10p family protein \| chr2:8831897-8832723 FORWARD LENGTH=64;AT2G20490.2 \| Symbols: NOP10, EDA27 \| nucleolar RNA-binding Nop10p family protein \| chr2:8831897-8832723 FORWARD LENGTH=63 | 0.070716 | 0.151382119 | 0.111049059 | 9.005028999 |
| AT4G02060 | AT4G02060.1 \| Symbols: PRL, MCM7 \| Minichromosome maintenance (MCM2/3/5) family protein \| chr4:901484-905297 FORWARD LENGTH=716;AT4G02060.2 \| Symbols: PRL \| Minichromosome maintenance (MCM2/3/5) family protein \| chr4:901484-905297 FORWARD LENGTH=716 | 0.12414 | 0.097675327 | 0.110907664 | 9.016509477 |
| AT3G25920 | AT3G25920.1 \| Symbols: RPL15 \| ribosomal protein L15 \| chr3:9491268-9492558 REVERSE LENGTH=277 | | 0.10863307 | 0.10863307 | 9.2053 |
| AT4G18100 | AT4G18100.1 \| Symbols: \| Ribosomal protein L32e \| chr4:10035715-10036475 REVERSE LENGTH=133 | 0.083669 | 0.132828585 | 0.108248792 | 9.237978348 |
| AT2G36160 | AT2G36160.1 \| Symbols: \| Ribosomal protein S11 family protein \| chr2:15169925-15171159 FORWARD LENGTH=150 | 0.073112 | 0.14079153 | 0.106951765 | 9.350009325 |
| AT1G08880 | AT1G08880.1 \| Symbols: HTA5, H2AXA, G-H2AX, GAMMA-H2AX \| Histone superfamily protein \| chr1:2847144-2847676 REVERSE LENGTH=142;AT1G54690.1 \| Symbols: HTA3, H2AXB, G-H2AX, GAMMA-H2AX \| gamma histone variant H2AX \| chr1:20414607-20415206 REVERSE LENGTH=142;AT2G38810.1 \| Symbols: HTA8 \| histone H2A 8 \| chr2:16219444-16220679 REVERSE LENGTH=136;AT2G38810.2 \| Symbols: HTA8 \| histone H2A 8 \| chr2:16219444-16220679 REVERSE LENGTH=136;AT2G38810.3 \| Symbols: HTA8 \| histone H2A 8 \| chr2:16219444-16220679 REVERSE LENGTH=136;AT3G54560.1 \| Symbols: HTA11 \| histone H2A 11 \| chr3:20196532-20197466 FORWARD LENGTH=136;AT1G52740.1 \| Symbols: HTA9 \| histone H2A protein 9 \| chr1:19645409-19646221 FORWARD LENGTH=134;AT3G20670.1 \| Symbols: HTA13 \| histone H2A 13 \| chr3:7229472-7229963 FORWARD LENGTH=132 | 0.10608 | | 0.10608 | 9.426847662 |
| AT3G44110 | AT3G44110.1 \| Symbols: ATJ3, ATJ \| DNAJ homologue 3 \| chr3:15869115-15871059 REVERSE LENGTH=420;AT3G44110.2 \| Symbols: ATJ3, ATJ \| DNAJ homologue 3 \| chr3:15869179-15871059 REVERSE LENGTH=343 | 0.10601 | | 0.10601 | 9.433072352 |
| AT3G01280 | AT3G01280.1 \| Symbols: VDAC1, ATVDAC1 \| voltage dependent anion channel 1 \| chr3:85754-87612 FORWARD LENGTH=276 | 0.024965 | 0.182558373 | 0.103761687 | 9.637468641 |

| | | | | | |
|---|---|---|---|---|---|
| AT3G27240 | AT3G27240.1 \| Symbols: \| Cytochrome C1 family \| chr3:10056144-10058370 REVERSE LENGTH=307;AT5G40810.1 \| Symbols: \| Cytochrome C1 family \| chr5:16340200-16342327 FORWARD LENGTH=307;AT5G40810.2 \| Symbols: \| Cytochrome C1 family \| chr5:16340670-16342327 FORWARD LENGTH=260 | | 0.102740078 | 0.102740078 | 9.7333 |
| AT1G70600 | AT1G70600.1 \| Symbols: \| Ribosomal protein L18e/L15 superfamily protein \| chr1:26621168-26621608 REVERSE LENGTH=146 | 0.10263 | | 0.10263 | 9.743739647 |
| AT1G09850 | AT1G09850.1 \| Symbols: XBCP3 \| xylem bark cysteine peptidase 3 \| chr1:3201848-3203875 FORWARD LENGTH=437 | | 0.101274027 | 0.101274027 | 9.8742 |
| AT4G31880 | AT4G31880.1 \| Symbols: \| LOCATED IN: cytosol, chloroplast; EXPRESSED IN: 24 plant structures; EXPRESSED DURING: 14 growth stages; BEST Arabidopsis thaliana protein match is: Tudor/PWWP/MBT superfamily protein (TAIR:AT1G15940.1); Has 137162 Blast hits to 70781 proteins in 2973 species: Archae - 289; Bacteria - 24182; Metazoa - 56725; Fungi - 20130; Plants - 6559; Viruses - 758; Other Eukaryotes - 28519 (source: NCBI BLink). \| chr4:15419435-15423939 REVERSE LENGTH=873;AT4G31880.2 \| Symbols: \| LOCATED IN: cytosol; EXPRESSED IN: 24 plant structures; EXPRESSED DURING: 14 growth stages; BEST Arabidopsis thaliana protein match is: Tudor/PWWP/MBT superfamily protein (TAIR:AT1G15940.1). \| chr4:15419435-15423939 REVERSE LENGTH=872 | | 0.10119409 | 0.10119409 | 9.882 |
| AT2G07734 | AT2G07734.1 \| Symbols: \| Alpha-L RNA-binding motif/Ribosomal protein S4 family protein \| chr2:3472604-3473692 REVERSE LENGTH=362;ATMG00290.1 \| Symbols: RPS4 \| mitochondrial ribosomal protein S4 \| chrM:82028-83116 REVERSE LENGTH=362 | | 0.101143938 | 0.101143938 | 9.8869 |
| AT4G35000 | AT4G35000.1 \| Symbols: APX3 \| ascorbate peroxidase 3 \| chr4:16665007-16667541 REVERSE LENGTH=287;AT4G35970.1 \| Symbols: APX5 \| ascorbate peroxidase 5 \| chr4:17028651-17030205 FORWARD LENGTH=279 | 0.10092 | | 0.10092 | 9.908838684 |
| AT1G03860 | AT1G03860.1 \| Symbols: ATPHB2, PHB2 \| prohibitin 2 \| chr1:979611-981157 REVERSE LENGTH=286;AT1G03860.3 \| Symbols: ATPHB2, PHB2 \| prohibitin 2 \| chr1:979611-981157 REVERSE LENGTH=286 | 0.10859 | 0.092764378 | 0.100677189 | 9.932736577 |
| AT1G28290 | AT1G28290.1 \| Symbols: AGP31 \| arabinogalactan protein 31 \| chr1:9889331-9890843 REVERSE LENGTH=359;AT1G28290.2 \| Symbols: AGP31 \| arabinogalactan protein 31 \| chr1:9889331-9890843 REVERSE LENGTH=315 | | 0.100147216 | 0.100147216 | 9.9853 |
| AT3G26020 | AT3G26020.4 \| Symbols: \| Protein phosphatase 2A regulatory B subunit family protein \| chr3:9514363-9516585 FORWARD LENGTH=529;AT3G26020.3 \| Symbols: \| Protein phosphatase 2A regulatory B subunit family protein \| chr3:9514363-9516585 FORWARD LENGTH=512;AT3G26020.1 \| Symbols: \| Protein phosphatase 2A regulatory B subunit family protein \| chr3:9514363-9516585 FORWARD LENGTH=510;AT3G26020.2 \| Symbols: \| Protein phosphatase 2A regulatory B subunit family protein \| chr3:9514363-9516585 FORWARD LENGTH=510;AT5G25510.1 \| Symbols: \| Protein phosphatase 2A regulatory B subunit family protein \| chr5:8882728-8884325 REVERSE LENGTH=500;AT3G09880.1 \| Symbols: ATB' BETA \| Protein phosphatase 2A regulatory B subunit family protein \| chr3:3029902-3031931 FORWARD LENGTH=499;AT3G54930.1 \| Symbols: \| Protein phosphatase 2A regulatory B subunit family protein \| chr3:20351084-20352659 REVERSE LENGTH=497;AT1G13460.1 \| Symbols: \| Protein phosphatase 2A regulatory B subunit family protein \| chr1:4616504-4618180 FORWARD LENGTH=492;AT1G13460.2 \| Symbols: \| Protein phosphatase 2A regulatory B subunit family protein \| chr1:4616504-4618180 FORWARD LENGTH=492;AT3G26030.1 \| Symbols: ATB' DELTA \| serine/threonine protein phosphatase 2A 55 kDa regulatory subunit B prime delta \| chr3:9517741-9519260 FORWARD LENGTH=477 | 0.097279 | | 0.097279 | 10.27971093 |

| | | | | | |
|---|---|---|---|---|---|
| AT5G13490 | AT5G13490.1 \| Symbols: AAC2 \| ADP/ATP carrier 2 \| chr5:4336034-4337379 FORWARD LENGTH=385;AT5G13490.2 \| Symbols: AAC2 \| ADP/ATP carrier 2 \| chr5:4336034-4337379 FORWARD LENGTH=385;AT3G08580.1 \| Symbols: AAC1 \| ADP/ATP carrier 1 \| chr3:2605706-2607030 REVERSE LENGTH=381;AT3G08580.2 \| Symbols: AAC1 \| ADP/ATP carrier 1 \| chr3:2605706-2607030 REVERSE LENGTH=381;AT4G28390.1 \| Symbols: AAC3, ATAAC3 \| ADP/ATP carrier 3 \| chr4:14041486-14042781 REVERSE LENGTH=379 | 0.025763 | 0.168335999 | 0.097049499 | 10.30402018 |
| AT4G20980 | AT4G20980.1 \| Symbols: \| Eukaryotic translation initiation factor 3 subunit 7 (eIF-3) \| chr4:11216997-11218772 FORWARD LENGTH=591;AT4G20980.2 \| Symbols: \| Eukaryotic translation initiation factor 3 subunit 7 (eIF-3) \| chr4:11216997-11218772 FORWARD LENGTH=591;AT4G20980.3 \| Symbols: \| Eukaryotic translation initiation factor 3 subunit 7 (eIF-3) \| chr4:11216997-11218772 FORWARD LENGTH=591;AT4G20980.4 \| Symbols: \| Eukaryotic translation initiation factor 3 subunit 7 (eIF-3) \| chr4:11216997-11218772 FORWARD LENGTH=591;AT5G44320.1 \| Symbols: \| Eukaryotic translation initiation factor 3 subunit 7 (eIF-3) \| chr5:17854901-17856667 REVERSE LENGTH=588 | 0.096673 | | 0.096673 | 10.34414987 |
| AT1G64330 | AT1G64330.1 \| Symbols: \| myosin heavy chain-related \| chr1:23872172-23873970 FORWARD LENGTH=555 | 0.096303 | | 0.096303 | 10.38389251 |
| AT2G16640 | AT2G16640.1 \| Symbols: ATTOC132, TOC132 \| multimeric translocon complex in the outer envelope membrane 132 \| chr2:7211489-7215109 REVERSE LENGTH=1206;AT3G16620.1 \| Symbols: ATTOC120, TOC120 \| translocon outer complex protein 120 \| chr3:5658469-5661738 REVERSE LENGTH=1089 | | 0.095210892 | 0.095210892 | 10.503 |
| AT3G02900 | AT3G02900.1 \| Symbols: \| unknown protein; FUNCTIONS IN: molecular_function unknown; INVOLVED IN: biological_process unknown; LOCATED IN: chloroplast; EXPRESSED IN: 22 plant structures; EXPRESSED DURING: 13 growth stages; BEST Arabidopsis thaliana protein match is: unknown protein (TAIR:AT5G16660.1); Has 80 Blast hits to 80 proteins in 21 species: Archae - 0; Bacteria - 4; Metazoa - 0; Fungi - 0; Plants - 76; Viruses - 0; Other Eukaryotes - 0 (source: NCBI BLink). \| chr3:645850-647445 FORWARD LENGTH=162;AT3G02900.2 \| Symbols: \| unknown protein; FUNCTIONS IN: molecular_function unknown; INVOLVED IN: biological_process unknown; EXPRESSED IN: 22 plant structures; EXPRESSED DURING: 13 growth stages; BEST Arabidopsis thaliana protein match is: unknown protein (TAIR:AT5G16660.1). \| chr3:645810-647445 FORWARD LENGTH=161 | | 0.095201828 | 0.095201828 | 10.504 |
| AT3G26520 | AT3G26520.1 \| Symbols: TIP2, SITIP, GAMMA-TIP2, TIP1;2 \| tonoplast intrinsic protein 2 \| chr3:9722770-9723703 REVERSE LENGTH=253 | | 0.094750805 | 0.094750805 | 10.554 |
| AT5G15750 | AT5G15750.1 \| Symbols: \| Alpha-L RNA-binding motif/Ribosomal protein S4 family protein \| chr5:5141449-5142701 FORWARD LENGTH=182 | 0.092939 | | 0.092939 | 10.75974564 |
| AT5G11670 | AT5G11670.1 \| Symbols: ATNADP-ME2, NADP-ME2 \| NADP-malic enzyme 2 \| chr5:3754456-3758040 FORWARD LENGTH=588 | 0.091344 | | 0.091344 | 10.94762655 |
| AT5G13450 | AT5G13450.1 \| Symbols: ATP5 \| delta subunit of Mt ATP synthase \| chr5:4310558-4311941 REVERSE LENGTH=238;AT5G13450.2 \| Symbols: ATP5 \| delta subunit of Mt ATP synthase \| chr5:4310558-4311941 REVERSE LENGTH=190 | 0.036886 | 0.143274686 | 0.090080343 | 11.10120105 |
| AT2G09990 | AT2G09990.1 \| Symbols: \| Ribosomal protein S5 domain 2-like superfamily protein \| chr2:3781442-3781882 FORWARD LENGTH=146 | 0.090559 | 0.087919817 | 0.089239409 | 11.2058116 |

| | | | | | |
|---|---|---|---|---|---|
| AT5G15320 | AT5G15320.1 \| Symbols: \| unknown protein; FUNCTIONS IN: molecular_function unknown; INVOLVED IN: biological_process unknown; LOCATED IN: endomembrane system; EXPRESSED IN: 23 plant structures; EXPRESSED DURING: 15 growth stages; BEST Arabidopsis thaliana protein match is: unknown protein (TAIR:AT3G01130.1); Has 64 Blast hits to 64 proteins in 11 species: Archae - 0; Bacteria - 0; Metazoa - 0; Fungi - 0; Plants - 64; Viruses - 0; Other Eukaryotes - 0 (source: NCBI BLink). \| chr5:4977647-4978658 FORWARD LENGTH=53;AT5G15320.2 \| Symbols: \| unknown protein; FUNCTIONS IN: molecular_function unknown; INVOLVED IN: biological_process unknown; LOCATED IN: endomembrane system; EXPRESSED IN: 23 plant structures; EXPRESSED DURING: 15 growth stages; BEST Arabidopsis thaliana protein match is: unknown protein (TAIR:AT3G01130.1); Has 64 Blast hits to 64 proteins in 11 species: Archae - 0; Bacteria - 0; Metazoa - 0; Fungi - 0; Plants - 64; Viruses - 0; Other Eukaryotes - 0 (source: NCBI BLink). \| chr5:4977647-4978658 FORWARD LENGTH=50 | | 0.089190153 | 0.089190153 | 11.212 |
| AT5G39740 | AT5G39740.1 \| Symbols: OLI7, RPL5B \| ribosomal protein L5 B \| chr5:15903365-15905185 FORWARD LENGTH=301;AT5G39740.2 \| Symbols: RPL5B \| ribosomal protein L5 B \| chr5:15903484-15905185 FORWARD LENGTH=301 | | 0.088597502 | 0.088597502 | 11.287 |
| AT5G15090 | AT5G15090.1 \| Symbols: VDAC3, ATVDAC3 \| voltage dependent anion channel 3 \| chr5:4889641-4891389 REVERSE LENGTH=274;AT5G15090.2 \| Symbols: VDAC3 \| voltage dependent anion channel 3 \| chr5:4889641-4891389 REVERSE LENGTH=274 | 0.040832 | 0.130943184 | 0.085887592 | 11.64312537 |
| AT3G47370 | AT3G47370.1 \| Symbols: \| Ribosomal protein S10p/S20e family protein \| chr3:17453671-17454437 REVERSE LENGTH=122;AT3G47370.2 \| Symbols: \| Ribosomal protein S10p/S20e family protein \| chr3:17453671-17454437 REVERSE LENGTH=122;AT3G47370.3 \| Symbols: \| Ribosomal protein S10p/S20e family protein \| chr3:17453671-17454437 REVERSE LENGTH=122 | 0.050134 | 0.121118163 | 0.085626081 | 11.67868462 |
| AT5G13650 | AT5G13650.2 \| Symbols: \| elongation factor family protein \| chr5:4397821-4402364 FORWARD LENGTH=676;AT5G13650.1 \| Symbols: \| elongation factor family protein \| chr5:4397821-4402364 FORWARD LENGTH=675 | | 0.08486803 | 0.08486803 | 11.783 |
| AT2G28720 | AT2G28720.1 \| Symbols: \| Histone superfamily protein \| chr2:12327043-12327498 FORWARD LENGTH=151;AT3G45980.1 \| Symbols: H2B, HTB9 \| Histone superfamily protein \| chr3:16897492-16897944 REVERSE LENGTH=150;AT5G59910.1 \| Symbols: HTB4 \| Histone superfamily protein \| chr5:24127206-24127658 FORWARD LENGTH=150;AT1G07790.1 \| Symbols: HTB1 \| Histone superfamily protein \| chr1:2413049-2413495 FORWARD LENGTH=148;AT3G46030.1 \| Symbols: HTB11 \| Histone superfamily protein \| chr3:16913614-16914051 REVERSE LENGTH=145;AT5G22880.1 \| Symbols: H2B, HTB2 \| histone B2 \| chr5:7652130-7652567 REVERSE LENGTH=145;AT2G37470.1 \| Symbols: \| Histone superfamily protein \| chr2:15736832-15737248 FORWARD LENGTH=138;AT5G02570.1 \| Symbols: \| Histone superfamily protein \| chr5:576742-577140 REVERSE LENGTH=132;AT3G53650.1 \| Symbols: \| Histone superfamily protein \| chr3:19889358-19889774 FORWARD LENGTH=138;AT3G09480.1 \| Symbols: \| Histone superfamily protein \| chr3:2914890-2915270 REVERSE LENGTH=126 | 0.083076 | 0.084552296 | 0.083814148 | 11.93115991 |
| AT2G15400 | AT2G15400.1 \| Symbols: RBP36B, NRPE3B \| DNA-directed RNA polymerase family protein \| chr2:6713022-6714386 FORWARD LENGTH=319;AT2G15430.1 \| Symbols: RBP36A, RPB35.5A, NRPB3, NRPD3, NRPE3A \| DNA-directed RNA polymerase family protein \| chr2:6733661-6735482 FORWARD LENGTH=319 | 0.081674 | | 0.081674 | 12.24379852 |
| AT5G54750 | AT5G54750.2 \| Symbols: \| Transport protein particle (TRAPP) component \| chr5:22242080-22243477 FORWARD LENGTH=199;AT5G54750.1 \| Symbols: \| Transport protein particle (TRAPP) component \| chr5:22242080-22243477 FORWARD LENGTH=186 | 0.079008 | | 0.079008 | 12.65694613 |

| | | | | | |
|---|---|---|---|---|---|
| AT5G17770 | AT5G17770.1 \| Symbols: ATCBR, CBR1, CBR \| NADH:cytochrome B5 reductase 1 \| chr5:5864543-5866495 REVERSE LENGTH=281 | 0.077319 | | 0.077319 | 12.93343163 |
| AT3G55460 | AT3G55460.1 \| Symbols: SCL30, At-SCL30 \| SC35-like splicing factor 30 \| chr3:20561024-20563502 FORWARD LENGTH=262 | 0.076246 | | 0.076246 | 13.11544212 |
| AT5G65750 | AT5G65750.1 \| Symbols: \| 2-oxoglutarate dehydrogenase, E1 component \| chr5:26304212-26307947 FORWARD LENGTH=1025;AT3G55410.1 \| Symbols: \| 2-oxoglutarate dehydrogenase, E1 component \| chr3:20541897-20545728 FORWARD LENGTH=1017 | 0.075931 | | 0.075931 | 13.16985158 |
| AT3G07430 | AT3G07430.1 \| Symbols: emb1990, YLMG1-1, ATYLMG1-1 \| YGGT family protein \| chr3:2379251-2379949 FORWARD LENGTH=232;AT4G27990.1 \| Symbols: YLMG1-2, ATYLMG1-2 \| YGGT family protein \| chr4:13923991-13924647 REVERSE LENGTH=218 | | 0.075289866 | 0.075289866 | 13.282 |
| AT4G25630 | AT4G25630.1 \| Symbols: FIB2, ATFIB2 \| fibrillarin 2 \| chr4:13074239-13076205 FORWARD LENGTH=320;AT5G52470.1 \| Symbols: FIB1, FBR1, ATFIB1, ATFBR1, SKIP7 \| fibrillarin 1 \| chr5:21294290-21296509 FORWARD LENGTH=308;AT5G52470.2 \| Symbols: FIB1, FBR1, ATFIB1, ATFBR1, SKIP7 \| fibrillarin 1 \| chr5:21294290-21296409 FORWARD LENGTH=273;AT5G52490.1 \| Symbols: \| Fibrillarin family protein \| chr5:21298857-21300239 FORWARD LENGTH=292 | 0.074722 | | 0.074722 | 13.38293943 |
| AT2G38550 | AT2G38550.1 \| Symbols: \| Transmembrane proteins 14C \| chr2:16132178-16134229 FORWARD LENGTH=335 | 0.074683 | | 0.074683 | 13.3899281 |
| AT3G59410 | AT3G59410.2 \| Symbols: GCN2 \| protein kinase family protein \| chr3:21950575-21959151 FORWARD LENGTH=1265;AT3G59410.1 \| Symbols: GCN2 \| protein kinase family protein \| chr3:21950575-21959151 FORWARD LENGTH=1241 | | 0.074299725 | 0.074299725 | 13.459 |
| AT5G08680 | AT5G08680.1 \| Symbols: \| ATP synthase alpha/beta family protein \| chr5:2821992-2824683 FORWARD LENGTH=559;AT5G08670.1 \| Symbols: \| ATP synthase alpha/beta family protein \| chr5:2818395-2821149 REVERSE LENGTH=556;AT5G08690.1 \| Symbols: \| ATP synthase alpha/beta family protein \| chr5:2825739-2828352 FORWARD LENGTH=556 | 0.035952 | 0.112381017 | 0.074166508 | 13.48317486 |
| AT5G10840 | AT5G10840.1 \| Symbols: \| Endomembrane protein 70 protein family \| chr5:3424910-3427797 REVERSE LENGTH=648 | | 0.072290899 | 0.072290899 | 13.833 |
| AT1G07280 | AT1G07280.1 \| Symbols: \| Tetratricopeptide repeat (TPR)-like superfamily protein \| chr1:2238505-2240991 FORWARD LENGTH=552;AT1G07280.2 \| Symbols: \| Tetratricopeptide repeat (TPR)-like superfamily protein \| chr1:2238505-2240991 FORWARD LENGTH=552;AT1G07280.3 \| Symbols: \| Tetratricopeptide repeat (TPR)-like superfamily protein \| chr1:2238505-2240991 FORWARD LENGTH=552 | 0.072255 | | 0.072255 | 13.83987267 |
| AT4G28470 | AT4G28470.1 \| Symbols: RPN1B, ATRPN1B \| 26S proteasome regulatory subunit S2 1B \| chr4:14067082-14072357 REVERSE LENGTH=891 | | 0.066462847 | 0.066462847 | 15.046 |
| AT5G66760 | AT5G66760.1 \| Symbols: SDH1-1 \| succinate dehydrogenase 1-1 \| chr5:26653776-26657224 FORWARD LENGTH=634;AT2G18450.1 \| Symbols: SDH1-2 \| succinate dehydrogenase 1-2 \| chr2:7997510-8000801 REVERSE LENGTH=632 | | 0.06532532 | 0.06532532 | 15.308 |
| AT5G42150 | AT5G42150.1 \| Symbols: \| Glutathione S-transferase family protein \| chr5:16846247-16847909 FORWARD LENGTH=315 | 0.064758 | | 0.064758 | 15.44210754 |
| AT4G13170 | AT4G13170.1 \| Symbols: \| Ribosomal protein L13 family protein \| chr4:7655133-7656542 REVERSE LENGTH=206;AT3G24830.1 \| Symbols: \| Ribosomal protein L13 family protein \| chr3:9064613-9065871 FORWARD LENGTH=206;AT5G48760.1 \| Symbols: \| Ribosomal protein L13 family protein \| chr5:19771315-19772686 REVERSE LENGTH=206;AT5G48760.2 \| Symbols: \| Ribosomal protein L13 family protein \| chr5:19771315-19772686 REVERSE LENGTH=206 | 0.064444 | | 0.064444 | 15.5173484 |
| AT2G02955 | AT2G02955.1 \| Symbols: MEE12 \| maternal effect embryo arrest 12 \| chr2:859592-861772 FORWARD LENGTH=666 | | 0.063897764 | 0.063897764 | 15.65 |
| AT5G13780 | AT5G13780.1 \| Symbols: \| Acyl-CoA N-acyltransferases (NAT) superfamily protein \| chr5:4447610-4448603 REVERSE LENGTH=192 | | 0.061564982 | 0.061564982 | 16.243 |

| | | | | | |
|---|---|---|---|---|---|
| AT3G10860 | AT3G10860.1 \| Symbols: \| Cytochrome b-c1 complex, subunit 8 protein \| chr3:3399815-3400514 FORWARD LENGTH=72;AT5G05370.1 \| Symbols: \| Cytochrome b-c1 complex, subunit 8 protein \| chr5:1590977-1591869 REVERSE LENGTH=72 | | 0.056462086 | 0.056462086 | 17.711 |
| AT2G19680 | AT2G19680.1 \| Symbols: \| Mitochondrial ATP synthase subunit G protein \| chr2:8501865-8502622 FORWARD LENGTH=122;AT2G19680.2 \| Symbols: \| Mitochondrial ATP synthase subunit G protein \| chr2:8501865-8502622 FORWARD LENGTH=122 | | 0.054614965 | 0.054614965 | 18.31 |
| AT2G19750 | AT2G19750.1 \| Symbols: \| Ribosomal protein S30 family protein \| chr2:8515037-8515806 FORWARD LENGTH=62;AT4G29390.1 \| Symbols: \| Ribosomal protein S30 family protein \| chr4:14465212-14465902 REVERSE LENGTH=62;AT5G56670.1 \| Symbols: \| Ribosomal protein S30 family protein \| chr5:22935413-22935981 REVERSE LENGTH=62 | 0.008722 | 0.096135359 | 0.052428679 | 19.07353024 |
| AT1G54290 | AT1G54290.1 \| Symbols: \| Translation initiation factor SUI1 family protein \| chr1:20268710-20269390 REVERSE LENGTH=113;AT4G27130.1 \| Symbols: \| Translation initiation factor SUI1 family protein \| chr4:13604814-13605525 REVERSE LENGTH=113;AT5G54760.1 \| Symbols: \| Translation initiation factor SUI1 family protein \| chr5:22244732-22245517 FORWARD LENGTH=113;AT5G54760.2 \| Symbols: \| Translation initiation factor SUI1 family protein \| chr5:22244732-22245517 FORWARD LENGTH=113;AT5G54760.3 \| Symbols: \| Translation initiation factor SUI1 family protein \| chr5:22244732-22245517 FORWARD LENGTH=113 | | 0.048253233 | 0.048253233 | 20.724 |
| AT4G32470 | AT4G32470.1 \| Symbols: \| Cytochrome bd ubiquinol oxidase, 14kDa subunit \| chr4:15669641-15671095 REVERSE LENGTH=122;AT5G25450.1 \| Symbols: \| Cytochrome bd ubiquinol oxidase, 14kDa subunit \| chr5:8857036-8857849 FORWARD LENGTH=122;AT5G25450.2 \| Symbols: \| Cytochrome bd ubiquinol oxidase, 14kDa subunit \| chr5:8857036-8857849 FORWARD LENGTH=106;AT4G32470.2 \| Symbols: \| Cytochrome bd ubiquinol oxidase, 14kDa subunit \| chr4:15669705-15671095 REVERSE LENGTH=101 | 0.045262 | 0.050859526 | 0.048060763 | 20.80699385 |
| AT1G26830 | AT1G26830.1 \| Symbols: CUL3A, ATCUL3A, ATCUL3, CUL3 \| cullin 3 \| chr1:9296063-9298374 FORWARD LENGTH=732;AT1G69670.1 \| Symbols: CUL3B, ATCUL3B \| cullin 3B \| chr1:26202169-26204442 REVERSE LENGTH=732 | 0.045083 | | 0.045083 | 22.18131003 |
| AT3G46430 | AT3G46430.1 \| Symbols: \| unknown protein; FUNCTIONS IN: molecular_function unknown; INVOLVED IN: biological_process unknown; LOCATED IN: mitochondrion, mitochondrial respiratory chain complex III; EXPRESSED IN: 25 plant structures; EXPRESSED DURING: 15 growth stages; BEST Arabidopsis thaliana protein match is: unknown protein (TAIR:AT5G59613.2); Has 51 Blast hits to 51 proteins in 16 species: Archae - 0; Bacteria - 0; Metazoa - 0; Fungi - 0; Plants - 51; Viruses - 0; Other Eukaryotes - 0 (source: NCBI BLink). \| chr3:17087687-17088497 FORWARD LENGTH=55;AT5G59613.1 \| Symbols: \| unknown protein; BEST Arabidopsis thaliana protein match is: unknown protein (TAIR:AT3G46430.1); Has 30201 Blast hits to 17322 proteins in 780 species: Archae - 12; Bacteria - 1396; Metazoa - 17338; Fungi - 3422; Plants - 5037; Viruses - 0; Other Eukaryotes - 2996 (source: NCBI BLink). \| chr5:24016162-24016500 REVERSE LENGTH=55;AT5G59613.2 \| Symbols: \| unknown protein; FUNCTIONS IN: molecular_function unknown; INVOLVED IN: biological_process unknown; LOCATED IN: mitochondrial respiratory chain complex III; BEST Arabidopsis thaliana protein match is: unknown protein (TAIR:AT3G46430.1). \| chr5:24015726-24016500 REVERSE LENGTH=55 | | 0.041758884 | 0.041758884 | 23.947 |
| AT3G18524 | AT3G18524.1 \| Symbols: MSH2, ATMSH2 \| MUTS homolog 2 \| chr3:6368151-6372409 REVERSE LENGTH=937 | 0.04149 | | 0.04149 | 24.1021933 |
| AT2G07727 | AT2G07727.1 \| Symbols: \| Di-haem cytochrome, transmembrane;Cytochrome b/b6, C-terminal \| chr2:3450863-3452044 FORWARD LENGTH=393;ATMG00220.1 \| Symbols: COB \| apocytochrome b \| chrM:60235-61416 FORWARD LENGTH=393 | | 0.041298422 | 0.041298422 | 24.214 |

| | | | | | |
|---|---|---|---|---|---|
| AT4G25730 | AT4G25730.1 \| Symbols: \| FtsJ-like methyltransferase family protein \| chr4:13102906-13106718 REVERSE LENGTH=821 | | 0.035339435 | 0.035339435 | 28.297 |
| AT5G13440 | AT5G13440.1 \| Symbols: \| Ubiquinol-cytochrome C reductase iron-sulfur subunit \| chr5:4308431-4310022 REVERSE LENGTH=274;AT5G13430.1 \| Symbols: \| Ubiquinol-cytochrome C reductase iron-sulfur subunit \| chr5:4305414-4307399 REVERSE LENGTH=272 | 0.034347 | | 0.034347 | 29.11462428 |
| AT3G50140 | AT3G50140.1 \| Symbols: \| Plant protein of unknown function (DUF247) \| chr3:18592472-18594562 REVERSE LENGTH=539;AT3G50190.1 \| Symbols: \| Plant protein of unknown function (DUF247) \| chr3:18609164-18611169 REVERSE LENGTH=493 | 0.033289 | | 0.033289 | 30.03995314 |
| AT3G19970 | AT3G19970.1 \| Symbols: \| alpha/beta-Hydrolases superfamily protein \| chr3:6959644-6961367 FORWARD LENGTH=434 | 0.026416 | | 0.026416 | 37.85584494 |
| AT5G20320 | AT5G20320.1 \| Symbols: DCL4, ATDCL4 \| dicer-like 4 \| chr5:6859571-6869068 REVERSE LENGTH=1702;AT5G20320.2 \| Symbols: DCL4 \| dicer-like 4 \| chr5:6859571-6869068 REVERSE LENGTH=1688 | 0.021956 | | 0.021956 | 45.54563673 |
| AT5G53080 | AT5G53080.1 \| Symbols: \| Tetratricopeptide repeat (TPR)-like superfamily protein \| chr5:21520624-21523158 FORWARD LENGTH=564 | 0.021688 | | 0.021688 | 46.10844707 |

**Supplemental Dataset 4. LC/MS-MS data corresponding to discussed proteins.** Data from CCV MS/MS and dimethyl labeling experiments (Supplemental Tables 1-3) is presented for proteins discussed in the manuscript and categorized by function.

| | | | Total Spectral Counts | | | | | Fold Enrichment (CCV:DFGL) | | | iBAQ Values | | | |
|---|---|---|---|---|---|---|---|---|---|---|---|---|---|---|
| **Accession** | **Label** | **MW** | **Rep 1** | **Rep 2** | **Rep 3** | **Rep 4** | **Sum** | **Rep 1** | **Rep 2** | **Average** | **Rep 1** | **Rep 2** | **Rep 3** | **Sum** |
| **Clathrin** | | | | | | | | | | | | | | |
| AT3G11130 | CHC1 | 193 | 2144 | 3376 | 12637 | 9651 | 27,808 | 7.34 | 4.88 | 6.11 | 6.7E+08 | 6.8E+08 | 2.6E+08 | 1.6E+09 |
| AT3G08530 | CHC2 | 193 | 2101 | 3145 | 12447 | 9390 | 27,083 | 6.82 | 3.67 | 5.25 | 6.7E+08 | 6.6E+08 | 2.4E+08 | 1.6E+09 |
| AT2G20760 | CLC1 | 37 | 144 | 441 | 906 | 648 | 2,139 | 3.57 | 12.77 | 8.17 | 8.9E+08 | 1.6E+09 | 2.5E+08 | 2.7E+09 |
| AT2G40060 | CLC2 | 29 | 143 | 628 | 1011 | 408 | 2,190 | 4.47 | 31.87 | 18.17 | 6.2E+08 | 1.0E+09 | 2.6E+08 | 1.9E+09 |
| AT3G51890 | CLC3 | 29 | 82 | 574 | 582 | 614 | 1,852 | 11.07 | 6.49 | 8.78 | 1.4E+08 | 1.8E+08 | 2.0E+07 | 3.4E+08 |
| **Adaptins** | | | | | | | | | | | | | | |
| AT4G11380 | AP1/2B1 | 99 | 214 | 204 | 1662 | 1692 | 3,772 | 3.53 | 7.65 | 5.59 | 7.1E+07 | 5.9E+07 | 1.8E+07 | 1.5E+08 |
| AT4G23460 | AP1/2B2 | 99 | 201 | 186 | 1394 | 1506 | 3,287 | 4.13 | 4.81 | 4.47 | 1.9E+07 | 1.7E+07 | 1.1E+07 | 4.7E+07 |
| AT1G60070 | AP1G1 | 95 | 159 | 325 | 1935 | 1345 | 3,764 | 5.06 | 6.20 | 5.63 | 9.5E+07 | 1.2E+08 | 2.9E+07 | 2.5E+08 |
| AT1G23900 | AP1G2 | 96 | 201 | 372 | 1819 | 1353 | 3,745 | 4.25 | 5.61 | 4.93 | 3.4E+07 | 4.5E+07 | 2.4E+07 | 1.0E+08 |
| AT1G10730 | AP1M1 | 49 | 26 | 49 | 348 | 406 | 829 | | 8.02 | 8.02 | 3.8E+05 | 5.4E+05 | 6.2E+04 | 9.8E+05 |
| AT1G60780 | AP1M2 | 49 | 96 | 258 | 908 | 793 | 2,055 | 3.07 | 4.50 | 3.78 | 5.0E+07 | 6.6E+07 | 2.2E+07 | 1.4E+08 |
| AT2G17380 | AP1S1 | 19 | 37 | 89 | 353 | 144 | 623 | | 7.01 | 7.01 | 5.0E+07 | 7.9E+07 | 1.0E+07 | 1.4E+08 |
| AT4G35410 | AP1S2 | 19 | 26 | 46 | 291 | 117 | 480 | 10.33 | | 10.33 | 2.8E+06 | 6.9E+06 | 1.3E+06 | 1.1E+07 |
| AT5G22770 | AP2A1 | 112 | 155 | 216 | 823 | 779 | 1,973 | | | | 1.5E+06 | 1.2E+06 | 5.4E+05 | 3.3E+06 |
| AT5G22780 | AP2A2 | 112 | 155 | 220 | 862 | 814 | 2,051 | 1.56 | 3.68 | 2.62 | 3.4E+06 | 2.7E+06 | 3.3E+05 | 6.4E+06 |
| AT5G46630 | AP2M | 49 | 30 | 87 | 429 | 543 | 1,089 | | 2.89 | 2.89 | 2.0E+07 | 2.1E+07 | 3.9E+06 | 4.4E+07 |
| AT1G47830 | AP2S | 17 | 2 | 4 | 30 | 25 | 61 | | 5.71 | 5.71 | 2.8E+07 | 2.7E+07 | 2.9E+06 | 5.7E+07 |
| AT1G48760 | AP3D | 97 | | | 9 | 10 | 19 | | | | | | | |
| AT1G56590 | AP3M | 46 | | | 6 | 5 | 11 | | | | | | | |
| AT5G11490 | AP4B | 93 | 108 | 114 | 465 | 333 | 1,020 | | 6.27 | 6.27 | 4.5E+07 | 6.4E+07 | 9.0E+06 | 1.2E+08 |
| AT1G31730 | AP4E | 104 | 175 | 260 | 779 | 528 | 1,742 | 3.30 | 5.86 | 4.58 | 6.4E+07 | 8.7E+07 | 1.3E+07 | 1.6E+08 |
| AT4G24550 | AP4M | 51 | 103 | 213 | 467 | 562 | 1,345 | 3.47 | 2.88 | 3.17 | 2.9E+07 | 4.1E+07 | 6.6E+06 | 7.7E+07 |
| AT2G19790 | AP4S | 17 | 22 | 40 | 70 | 45 | 177 | | | | 5.6E+07 | 8.7E+07 | 2.6E+06 | 1.5E+08 |
| AT3G19870 | AP5B | 122 | | | 4 | 4 | 8 | | | | | | | |
| AT2G20790 | AP5M | 67 | | | 3 | 4 | 7 | | | | | | | |
| **TPC** | | | | | | | | | | | | | | |
| AT3G01780 | TPLATE | 131 | 1 | 0 | 101 | 107 | 209 | | 0.53 | 0.53 | 6.1E+06 | 5.3E+06 | 1.3E+06 | 1.3E+07 |
| AT1G20760 | AthEH1 | 110 | 1 | 3 | 51 | 49 | 104 | | | | 3.4E+05 | 7.9E+05 | 1.2E+05 | 1.3E+06 |
| AT1G21630 | AtEH2 | 132 | | 2 | 58 | 60 | 120 | | | | 2.7E+06 | 3.0E+06 | 9.3E+05 | 6.6E+06 |
| AT2G07360 | TASH3 | 130 | 1 | 12 | 113 | 93 | 219 | | 1.29 | 1.29 | 6.6E+06 | 8.1E+06 | 2.2E+06 | 1.7E+07 |
| AT5G57460 | TML | 71 | 1 | 2 | 86 | 59 | 148 | | 1.38 | 1.38 | 3.4E+06 | 4.3E+06 | 9.5E+05 | 8.7E+06 |
| AT3G50590 | TWD40-1 | 176 | 1 | 3 | 88 | 91 | 183 | | 0.20 | 0.20 | 4.4E+06 | 4.2E+06 | 7.3E+05 | 9.4E+06 |
| AT5G24710 | TWD40-2 | 148 | 1 | 3 | 126 | 125 | 255 | | 0.25 | 0.25 | 8.2E+06 | 9.0E+06 | 2.7E+06 | 2.0E+07 |
| AT1G15370 | LOLITA | 17 | | | 5 | 1 | 6 | | | | | | | |
| **DRPs** | | | | | | | | | | | | | | |
| AT5G42080 | DRP1A | 68 | | 1 | 35 | 18 | 54 | | | | 7.2E+05 | 7.5E+05 | 0.0E+00 | 1.5E+06 |
| AT1G14830 | DRP1C | 69 | | 1 | 27 | 8 | 36 | | | | | | | |

| | | | | | | | | | | | | | | |
|---|---|---|---|---|---|---|---|---|---|---|---|---|---|---|
| AT3G60190 | DRP1E | 70 | | | 10 | 7 | 17 | | | | | | | |
| AT1G10290 | DRP2A | 99 | | | 18 | 7 | 25 | | | | | | | |
| AT1G59610 | DRP2B | 100 | | 1 | 19 | 7 | 27 | | | | | | | |
| **ENTH/ANTH** | | | | | | | | | | | | | | |
| AT2G01600 | AtECA1 | 64 | 51 | 80 | 383 | 361 | 875 | 2.66 | 6.79 | 4.72 | 1.9E+07 | 2.3E+07 | 2.3E+06 | 4.5E+07 |
| AT1G03050 | AtECA2 | 67 | | | 18 | 10 | 28 | | | | | | | |
| AT2G25430 | AtECA4 | 72 | 100 | 147 | 672 | 668 | 1,587 | 3.78 | 8.61 | 6.19 | 2.0E+07 | 3.0E+07 | 1.5E+06 | 5.2E+07 |
| AT4G32285 | CAP1 | 71 | 157 | 248 | 863 | 705 | 1,973 | 6.04 | 5.78 | 5.91 | 3.3E+07 | 4.9E+07 | 4.4E+06 | 8.6E+07 |
| AT2G38410 | TOL6 | 72 | 35 | 24 | 219 | 279 | 557 | 3.52 | 4.03 | 3.78 | 3.7E+06 | 4.7E+06 | 7.4E+05 | 9.1E+06 |
| AT4G32760 | TOL9 | 73 | 33 | 80 | 263 | 147 | 523 | 8.88 | 3.86 | 6.37 | 1.4E+07 | 1.4E+07 | 8.7E+06 | 3.7E+07 |
| AT3G59290 | EPSIN3 | 110 | 31 | 90 | 206 | 163 | 490 | | | | 6.0E+05 | 9.8E+05 | 3.1E+05 | 1.9E+06 |
| AT5G35200 | PICALM3 | 61 | | 2 | 44 | 28 | 74 | | | | 2.1E+05 | 6.3E+05 | 0.0E+00 | 8.4E+05 |
| AT1G14910 | PICALM1B | 76 | 17 | 13 | 111 | 101 | 242 | | | | 1.3E+06 | 1.4E+06 | 2.9E+05 | 2.9E+06 |
| AT5G63640 | TOL5 | 49 | 1 | 1 | 7 | 8 | 17 | | | | | | | |
| AT5G01760 | TOL7 | 60 | | | 7 | 7 | 14 | | | | | | | |
| AT4G25940 | PICALM2B | 68 | 4 | 2 | 29 | 32 | 67 | | | | | | | |
| AT5G11710 | EpsinR1 | 61 | 140 | 314 | 796 | 565 | 1,815 | 3.31 | 6.66 | 4.98 | 1.1E+08 | 1.7E+08 | 3.3E+07 | 3.2E+08 |
| AT2G43160 | EpsinR2 | 95 | 93 | 270 | 401 | 386 | 1,150 | 4.32 | 4.68 | 4.50 | 3.0E+07 | 4.0E+07 | 1.0E+07 | 8.1E+07 |
| AT3G16270 | MTV1 | 75 | 96 | 244 | 666 | 488 | 1,494 | | 6.74 | 6.74 | 1.7E+07 | 2.6E+07 | 9.6E+06 | 5.3E+07 |
| **SNAREs** | | | | | | | | | | | | | | |
| AT1G77140 | VPS45 | 65 | 14 | 10 | 315 | 213 | 552 | | 5.25 | 5.25 | 5.9E+06 | 3.2E+06 | 2.1E+06 | 1.1E+07 |
| AT1G28490 | SYP61 | 28 | 17 | 95 | 154 | 101 | 367 | 3.65 | 7.54 | 5.60 | 5.3E+06 | 8.8E+06 | 2.5E+06 | 1.7E+07 |
| AT3G09740 | SYP71 | 30 | 34 | 42 | 123 | 123 | 322 | 2.66 | 8.43 | 5.55 | 4.2E+07 | 4.0E+07 | 1.6E+06 | 8.3E+07 |
| AT3G54300 | VAMP727 | 27 | 31 | 103 | 86 | 141 | 361 | 2.54 | 8.06 | 5.30 | 5.4E+07 | 7.4E+07 | 6.3E+06 | 1.3E+08 |
| AT2G33120 | VAMP722 | 26 | 26 | 81 | 132 | 94 | 333 | 2.06 | 7.20 | 4.63 | 2.7E+06 | 2.3E+06 | 3.6E+05 | 5.4E+06 |
| AT5G08080 | SYP132 | 34 | 29 | 51 | 125 | 95 | 300 | | 7.63 | 7.63 | 1.4E+08 | 1.1E+08 | 3.7E+06 | 2.6E+08 |
| AT3G05710 | SYP43 | 37 | 5 | 6 | 70 | 127 | 208 | | 8.72 | 8.72 | | 9.5E+05 | 5.3E+04 | 1.0E+06 |
| AT3G11820 | SYP121 | 38 | 20 | 24 | 52 | 55 | 151 | | 6.86 | 6.86 | 2.4E+06 | 2.2E+06 | 8.1E+04 | 4.8E+06 |
| AT5G11150 | VAMP713 | 25 | 2 | 0 | 53 | 46 | 101 | 0.23 | 0.18 | 0.21 | 1.6E+07 | 5.8E+06 | 2.0E+06 | 2.4E+07 |
| AT4G02195 | SYP42 | 36 | 10 | 8 | 68 | 116 | 202 | | | | 9.3E+05 | 6.1E+05 | 1.4E+05 | 1.7E+06 |
| AT5G46860 | SYP22 | 29 | 6 | 13 | 20 | 56 | 95 | | | | 1.9E+07 | 1.5E+07 | 9.7E+05 | 3.5E+07 |
| AT1G48240 | NPSN12 | 30 | 5 | 3 | 42 | 23 | 73 | | | | 8.4E+06 | 1.2E+07 | 0.0E+00 | 2.0E+07 |
| AT1G08560 | KNOLLE | 35 | 4 | 3 | 33 | 34 | 74 | 1.41 | 11.15 | 6.28 | | | | |
| AT5G26980 | SYP41 | 36 | 6 | 11 | 31 | 76 | 124 | | 12.55 | 12.55 | | 9.6E+05 | | 9.6E+05 |
| AT5G22360 | VAMP714 | 25 | 1 | 3 | 32 | 16 | 52 | | 0.31 | 0.31 | 2.1E+06 | | 2.3E+05 | 2.4E+06 |
| AT4G15780 | VAMP724 | 25 | 5 | 8 | 20 | 19 | 52 | | | | | | | |
| AT1G26670 | VTI12 | 25 | 1 | 12 | 32 | 8 | 53 | 6.59 | | 6.59 | | | | |
| AT5G39510 | VTI11 | 25 | 4 | 6 | 23 | 20 | 53 | 0.39 | 1.60 | 1.00 | 4.0E+06 | 2.8E+06 | 9.9E+05 | 7.7E+06 |
| AT3G52400 | SYP122 | 38 | 4 | 5 | 16 | 19 | 44 | | 5.15 | 5.15 | 2.4E+07 | 1.5E+07 | 2.9E+04 | 3.9E+07 |
| AT1G04750 | VAMP721 | 25 | | | | | | | | | 2.1E+06 | 2.1E+06 | 0.0E+00 | 4.2E+06 |
| AT5G16830 | SYP21 | 31 | 8 | 17 | 14 | 10 | 49 | | 2.36 | 2.36 | | | | |
| AT4G32150 | VAMP711 | 25 | | 1 | 27 | 11 | 39 | | | | | | | |
| AT1G16240 | SYP51 | 26 | | 2 | 13 | 8 | 23 | | | | | | | |
| AT3G17440 | NPSN13 | 30 | 4 | 5 | 42 | 14 | 65 | | 6.05 | 6.05 | 1.2E+07 | 1.4E+07 | 0.0E+00 | 2.6E+07 |
| AT1G79590 | SYP52 | 26 | | | 9 | 8 | 17 | | | | | | | |
| AT5G61210 | SNAP33 | 34 | 5 | 0 | 2 | 9 | 16 | | | | | | | |

| | | | | | | | | | | | | | | |
|---|---|---|---|---|---|---|---|---|---|---|---|---|---|---|
| AT2G35190 | NPSN11 | 30 | | 2 | 14 | 10 | 26 | | | | 1.8E+06 | 7.2E+05 | 0.0E+00 | 2.5E+06 |
| AT1G04760 | VAMP726 | 25 | 15 | 36 | 87 | 35 | 173 | | | | | | | |
| AT1G80500 | TRAPPC2 | 15 | 2 | 0 | 5 | 7 | 14 | | 0.39 | 0.39 | | | | |
| **VSRs** | | | | | | | | | | | | | | |
| AT3G52850 | VSR1 | 69 | 95 | 116 | 431 | 360 | 1,002 | 3.68 | 2.99 | 3.33 | 4.2E+07 | 5.5E+07 | 7.4E+06 | 1.0E+08 |
| AT2G14740 | VSR3 | 70 | 79 | 67 | 429 | 376 | 951 | | | | 8.4E+05 | 8.2E+05 | 8.6E+05 | 2.5E+06 |
| AT2G14720 | VSR4 | 70 | 84 | 75 | 458 | 398 | 1,015 | 1.47 | 4.03 | 2.75 | 6.0E+06 | 8.0E+06 | 2.2E+06 | 1.6E+07 |
| AT1G30900 | VSR6 | 70 | 2 | 4 | 13 | 14 | 33 | | | | | | | |
| AT4G20110 | VSR7 | 70 | | | 19 | 11 | 30 | | | | | | | |
| **LRRK-RLKs** | | | | | | | | | | | | | | |
| AT5G46330 | FLS2 | 129 | 2 | 0 | 7 | 12 | 21 | | | | | | | |
| AT4G33430 | BAK1 | 74 | 1 | 3 | 49 | 21 | 74 | | | | 4.0E+06 | 2.6E+06 | 0.0E+00 | 6.6E+06 |
| AT2G01820 | | 102 | 73 | 61 | 690 | 546 | 1,370 | 2.93 | 3.51 | 3.22 | 9.5E+07 | 7.4E+07 | 1.3E+07 | 1.8E+08 |
| AT3G49490 | | 105 | 24 | 78 | 323 | 212 | 637 | | 5.08 | 5.08 | 2.5E+06 | 4.0E+06 | 5.6E+06 | 1.2E+07 |
| AT3G02880 | KIN7 | 68 | 61 | 24 | 301 | 221 | 607 | 2.49 | 5.94 | 4.22 | 2.5E+07 | 1.9E+07 | 6.2E+05 | 4.5E+07 |
| AT4G08850 | MIK2 | 115 | 16 | 11 | 267 | 197 | 491 | 2.67 | 4.82 | 3.74 | 4.8E+07 | 4.2E+07 | 3.0E+06 | 9.3E+07 |
| AT1G11130 | | 84 | 11 | 6 | 86 | 74 | 177 | | 5.32 | 5.32 | 1.2E+07 | 8.9E+06 | 7.6E+04 | 2.1E+07 |
| AT5G16590 | | 67 | 32 | 6 | 137 | 111 | 286 | | 2.98 | 2.98 | 3.4E+06 | 3.2E+06 | 0.0E+00 | 6.6E+06 |
| AT5G49760 | | 105 | | | 11 | 14 | 25 | | | | 1.9E+05 | 1.3E+05 | 0.0E+00 | 3.3E+05 |
| AT5G58150 | | 87 | | | 7 | 6 | 13 | | | | | | | |
| AT3G23750 | | 100 | 14 | 4 | 91 | 103 | 212 | | | | 7.0E+06 | 3.0E+06 | 4.4E+04 | 1.0E+07 |
| AT1G33590 | | 52 | | 3 | 60 | 98 | 161 | 0.08 | 0.24 | 0.16 | 6.4E+07 | 3.7E+07 | 7.0E+06 | 1.1E+08 |
| AT1G27190 | | 65 | 4 | 8 | 86 | 51 | 149 | | 0.74 | 0.74 | 3.1E+06 | 3.8E+05 | 0.0E+00 | 3.4E+06 |
| AT2G37050 | | 104 | | | 66 | 56 | 122 | | | | 4.6E+06 | 3.7E+06 | 0.0E+00 | 8.3E+06 |
| AT5G05160 | | 71 | | | 51 | 64 | 115 | | | | 2.4E+06 | 1.6E+06 | 0.0E+00 | 3.9E+06 |
| AT1G56140 | | 114 | | 1 | 54 | 50 | 105 | | | | 1.2E+06 | 9.7E+05 | 0.0E+00 | 2.1E+06 |
| AT2G26730 | | 72 | 11 | 0 | 50 | 42 | 103 | | | | 1.2E+06 | 4.6E+05 | 0.0E+00 | 1.6E+06 |
| AT5G10290 | | 69 | | | 49 | 33 | 82 | | | | 6.1E+05 | 4.0E+05 | 0.0E+00 | 1.0E+06 |
| AT3G28450 | | 67 | 1 | 1 | 40 | 31 | 73 | | 2.23 | 2.23 | 9.1E+06 | 4.7E+06 | 0.0E+00 | 1.4E+07 |
| AT1G06840 | | 106 | | 1 | 43 | 26 | 70 | | 1.81 | 1.81 | 8.9E+05 | 3.3E+05 | 2.7E+04 | 1.2E+06 |
| AT5G65700 | | 109 | | | 30 | 24 | 54 | | | | 1.2E+06 | 4.1E+05 | 5.4E+04 | 1.6E+06 |
| AT2G36570 | | 74 | 2 | 2 | 22 | 37 | 63 | | | | 6.0E+06 | 2.7E+06 | 0.0E+00 | 8.7E+06 |
| AT2G16250 | | 100 | | | 27 | 24 | 51 | | | | 1.8E+06 | 1.2E+06 | 0.0E+00 | 3.0E+06 |
| AT5G10020 | | 115 | | | 24 | 26 | 50 | | | | 1.1E+06 | 8.0E+05 | 0.0E+00 | 1.9E+06 |
| AT4G22730 | | 76 | | | 16 | 27 | 43 | | | | 9.7E+05 | 4.0E+05 | 0.0E+00 | 1.4E+06 |
| AT2G41820 | | 97 | | | 23 | 21 | 44 | | | | 2.1E+06 | 1.9E+07 | 0.0E+00 | 2.1E+07 |
| AT1G51850 | | 96 | | | 30 | 12 | 42 | | | | | | | |
| AT5G07910 | | | 1 | 6 | 17 | 11 | 35 | | 6.76 | 6.76 | | | | |
| AT3G14350 | | 78 | 3 | 1 | 22 | 17 | 43 | | | | 1.5E+06 | 1.6E+06 | 0.0E+00 | 3.0E+06 |
| AT3G14840 | | 112 | | 1 | 27 | 35 | 63 | | | | | | | |
| AT4G20940 | | 105 | | | 15 | 19 | 34 | | | | | | | |
| AT5G62710 | | 67 | | 1 | 19 | 15 | 35 | | | | | | | |
| AT4G34220 | | 83 | | | 20 | 13 | 33 | | | | | | | |
| AT5G67200 | | 74 | | | 16 | 17 | 33 | | | | | | | |
| AT3G08680 | | 69 | | | 13 | 12 | 25 | | | | | | | |
| AT3G47570 | | 111 | 1 | 1 | 18 | 13 | 33 | | | | | | | |

| | | | | | | | | | | | | | | |
|---|---|---|---|---|---|---|---|---|---|---|---|---|---|---|
| AT1G63430 | | 74 | | | 17 | 13 | 30 | | | | | | | |
| AT5G65240 | | 68 | 1 | 0 | 36 | 27 | 64 | | | | | | | |
| AT1G53430 | | 114 | | 1 | 22 | 22 | 45 | | | | | | | |
| AT1G31420 | | 65 | | | 14 | 11 | 25 | | | | | | | |
| AT5G21090 | | 24 | 1 | 3 | 7 | 5 | 16 | | | | | | | |
| AT5G14210 | | 86 | | | 6 | 6 | 12 | | | | | | | |
| AT1G67720 | | 104 | | | 9 | 3 | 12 | | | | | | | |
| AT2G24230 | | 94 | | 1 | 20 | 17 | 38 | | | | | | | |
| AT1G68400 | | 74 | | | 5 | 10 | 15 | | | | | | | |
| AT5G01890 | | 104 | | | 8 | 8 | 16 | | | | | | | |
| AT1G09970 | | 107 | | 1 | 7 | 7 | 15 | | | | | | | |
| AT1G10850 | | 73 | | | 8 | 6 | 14 | | | | | | | |
| AT4G23740 | | 71 | | | 6 | 6 | 12 | | | | | | | |
| AT4G36180 | | 124 | | | 5 | 7 | 12 | | | | | | | |
| AT2G25790 | | 106 | | | 4 | 7 | 11 | | | | | | | |
| AT2G31880 | | 71 | | 1 | 15 | 17 | 33 | | | | | | | |
| AT1G56145 | | 112 | | 2 | 24 | 23 | 49 | | | | | | | |
| AT2G35620 | | 64 | | | 14 | 9 | 23 | | | | | | | |
| AT5G58300 | | 72 | | | 5 | 8 | 13 | | | | | | | |
| AT2G27060 | | 110 | 1 | 0 | 2 | 4 | 7 | | | | 3.5E+05 | 1.1E+05 | 0.0E+00 | 4.7E+05 |
| AT5G53320 | | 67 | | | 10 | 1 | 11 | | | | | | | |
| AT1G53440 | | 115 | | 1 | 21 | 19 | 41 | | | | | | | |
| AT1G25320 | | 76 | | | 7 | 6 | 13 | | | | | | | |
| AT1G15740 | | 64 | | | 4 | 2 | 6 | | | | | | | |
| **CDPKs** | | | | | | | | | | | | | | |
| AT4G04720 | CPK21 | 60 | 50 | 34 | 387 | 328 | 799 | 3.90 | 6.80 | 5.35 | 3.6E+07 | 2.1E+07 | 1.8E+06 | 5.9E+07 |
| AT4G23650 | CPK3 | 59 | 38 | 12 | 144 | 148 | 342 | | 2.64 | 2.64 | 2.2E+07 | 8.2E+06 | 1.1E+05 | 3.0E+07 |
| AT3G57530 | CPK32 | 61 | 2 | 4 | 64 | 53 | 123 | | | | 2.7E+06 | 2.0E+06 | 0.0E+00 | 4.6E+06 |
| AT4G04700 | CPK27 | 55 | 3 | 3 | 70 | 36 | 112 | | 3.20 | 3.20 | | | | |
| AT5G24430 | CPK | 67 | | | 51 | 31 | 82 | | 3.13 | 3.13 | | | | |
| AT3G49370 | CPK | 66 | | | 24 | 17 | 41 | | | | | | | |
| AT1G49580 | CPK | 68 | | | 19 | 9 | 28 | | | | | | | |
| AT5G19450 | CDPK19 | 60 | 1 | 7 | 46 | 39 | 93 | | | | 2.4E+06 | 1.8E+06 | 0.0E+00 | 4.2E+06 |
| AT5G04870 | CPK1 | 68 | | | 5 | 2 | 7 | | | | | | | |
| AT3G51850 | CPK13 | 59 | | | 26 | 17 | 43 | | | | 6.1E+06 | 3.4E+06 | 0.0E+00 | 9.5E+06 |
| AT4G21940 | CPK15 | 63 | | 11 | 129 | 134 | 274 | | | | | | | |
| AT4G04740 | CPK23 | 59 | 8 | 12 | 73 | 60 | 153 | | | | | | | |
| AT5G66210 | CPK29 | 59 | | | 22 | 10 | 32 | | | | | | | |
| AT1G76040 | CPK29 | 64 | | | 60 | 77 | 137 | | | | | | | |
| AT1G74740 | CPK30 | 61 | | 2 | 24 | 21 | 47 | | | | | | | |
| AT4G04695 | CPK31 | 55 | | | 25 | 17 | 42 | | | | | | | |
| AT1G50700 | CPK33 | 59 | | 2 | 37 | 39 | 78 | | | | | | | |
| AT2G17290 | CPK6 | 61 | | | 8 | 5 | 13 | | | | | | | |
| AT5G12480 | CPK7 | 60 | | 5 | 35 | 23 | 63 | | | | 2.1E+06 | 1.1E+06 | 0.0E+00 | 3.2E+06 |
| **CrRLK1Ls** | | | | | | | | | | | | | | |
| AT5G24010 | | 92 | 13 | 13 | 186 | 164 | 376 | 7.60 | 4.49 | 6.05 | 2.1E+07 | 1.3E+07 | 2.2E+05 | 3.3E+07 |

| | | | | | | | | | | | | | | |
|---|---|---|---|---|---|---|---|---|---|---|---|---|---|---|
| AT3G51550 | FER | 98 | 9 | 10 | 141 | 97 | 257 | 3.29 | 5.63 | 4.46 | 9.1E+06 | 8.5E+06 | 8.4E+04 | 1.8E+07 |
| AT3G46290 | HERK1 | 91 | 12 | 6 | 81 | 91 | 190 | | 3.16 | 3.16 | 9.4E+06 | 6.1E+06 | 0.0E+00 | 1.5E+07 |
| AT2G39360 | CrRLK1L | 91 | | | 7 | 19 | 26 | | | | | | | |
| AT5G38990 | CrRLK1L | 98 | | | 5 | 8 | 13 | | | | | | | |
| AT5G59700 | CrRLK1L | 92 | | 3 | 29 | 26 | 58 | | | | | | | |
| AT5G54380 | THE1 | 93 | 2 | 3 | 30 | 26 | 61 | | | | 9.3E+05 | 8.8E+05 | 0.0E+00 | 1.8E+06 |
| AT1G30570 | HERK2 | 94 | | 3 | 37 | 32 | 72 | | | | 3.9E+05 | 4.3E+05 | 0.0E+00 | 8.2E+05 |
| **Cargo** | | | | | | | | | | | | | | |
| AT4G32410 | CESA1 | 122 | | | 27 | 18 | 45 | | | | | | | |
| AT5G05170 | CESA3 | 120 | | | 25 | 18 | 43 | | | | | | | |
| AT5G64740 | CESA6 | 123 | | | 14 | 7 | 21 | | | | | | | |
| AT3G53420 | PIP2;1 | 30 | | 1 | 18 | 20 | 39 | | | | 1.5E+06 | 9.2E+05 | 0.0E+00 | 2.4E+06 |
| AT5G47910 | RBOHD | 104 | | | 69 | 32 | 101 | | | | 1.1E+06 | 1.2E+06 | 0.0E+00 | 2.3E+06 |
| AT1G73590 | PIN1 | 67 | 5 | 2 | 34 | 32 | 73 | | | | | | | |
| AT1G70940 | PIN3 | 69 | | | 15 | 27 | 42 | | | | | | | |
| AT1G77110 | PIN6 | 62 | | | 6 | 13 | 19 | | | | | | | |
| AT1G69840 | AtHIR1 | 31 | 43 | 41 | 95 | 79 | 258 | 1.36 | 4.20 | 2.78 | 2.3E+07 | 1.2E+07 | 9.2E+05 | 3.6E+07 |
| AT3G01290 | AtHIR2 | 31 | 26 | 16 | 30 | 48 | 120 | | 5.83 | 5.83 | | | | |
| AT5G51570 | AtHIR3 | 32 | 33 | 54 | 180 | 135 | 402 | 0.47 | 4.42 | 2.45 | 5.9E+06 | 1.9E+06 | 6.2E+06 | 1.4E+07 |
| AT5G62740 | AtHIR4 | 31 | 58 | 112 | 99 | 146 | 415 | 1.02 | 5.27 | 3.15 | 3.5E+07 | 2.8E+07 | 3.4E+06 | 6.6E+07 |
| AT2G18960 | AHA1 | 104 | 106 | 42 | 859 | 506 | 1,513 | 1.61 | 0.44 | 1.03 | 4.5E+07 | 2.8E+07 | 3.9E+06 | 7.6E+07 |
| AT5G57350 | AHA3 | 104 | 37 | 14 | 426 | 280 | 757 | | | | 1.7E+06 | 4.9E+05 | 0.0E+00 | 2.2E+06 |
| AT4G30190 | AHA2 | 104 | 94 | 22 | 743 | 458 | 1,317 | | 0.16 | 0.16 | 2.5E+07 | 2.0E+07 | 1.6E+06 | 4.7E+07 |
| AT3G28710 | V0-C/D | 41 | 28 | 72 | 255 | 191 | 546 | 0.52 | 3.82 | 2.17 | 3.4E+06 | 3.3E+06 | 4.3E+05 | 7.1E+06 |
| AT3G28715 | V0-C/D | 41 | 28 | 71 | 255 | 193 | 547 | | 3.57 | 3.57 | 1.0E+07 | 3.3E+06 | 1.2E+05 | 1.4E+07 |
| AT1G78900 | V1-A | 69 | 77 | 72 | 973 | 693 | 1,815 | 0.12 | 0.46 | 0.29 | 1.8E+08 | 1.1E+08 | 2.0E+07 | 3.2E+08 |
| AT2G28520 | V1-A1 | 93 | 17 | 26 | 378 | 199 | 620 | 3.92 | 1.91 | 2.91 | 7.0E+06 | 8.3E+06 | 4.0E+06 | 1.9E+07 |
| AT2G21410 | V1-A2 | 93 | | 4 | 92 | 85 | 181 | 0.67 | 0.21 | 0.44 | 9.0E+05 | 4.8E+05 | 0.0E+00 | 1.4E+06 |
| AT4G39080 | V1-A3 | 93 | 10 | 5 | 181 | 143 | 339 | 0.39 | 0.21 | 0.30 | 1.8E+07 | 1.0E+07 | 8.9E+06 | 3.7E+07 |
| AT1G76030 | V1-B | 54 | 56 | 97 | 579 | 602 | 1,334 | | | | 3.4E+07 | 2.3E+07 | 3.0E+06 | 6.0E+07 |
| AT1G12840 | V1-C | 43 | 18 | 36 | 173 | 131 | 358 | 0.22 | 2.00 | 1.11 | 5.2E+07 | 2.9E+07 | 4.3E+06 | 8.6E+07 |
| AT3G58730 | V1-D | 29 | 21 | 36 | 91 | 58 | 206 | 0.31 | 2.26 | 1.29 | 3.6E+07 | 2.0E+07 | 3.9E+06 | 6.0E+07 |
| AT4G11150 | V1-E1 | 26 | 26 | 64 | 144 | 134 | 368 | 0.33 | 1.67 | 1.00 | 9.7E+07 | 6.0E+07 | 3.3E+06 | 1.6E+08 |
| AT1G64200 | V1-E3 | 27 | | 34 | 78 | 116 | 228 | | 1.48 | 1.48 | 4.4E+06 | 7.4E+05 | 2.1E+06 | 7.2E+06 |
| AT4G23710 | V1-G2 | 12 | 2 | 0 | 13 | 4 | 19 | 0.37 | 0.69 | 0.53 | 7.7E+07 | 4.3E+07 | 4.1E+06 | 1.2E+08 |
| **Tethering** | | | | | | | | | | | | | | |
| AT5G03540 | EXO70A1 | 72 | 17 | 2 | 47 | 29 | 95 | | | | | | | |
| AT5G58430 | **EXO70B1** | 71 | | | 28 | 28 | 56 | | | | | | | |
| AT1G72470 | **EXO70D1** | 71 | | | 8 | 4 | 12 | | | | | | | |
| AT1G54090 | **EXO70D2** | 71 | | | 8 | 5 | 13 | | | | | | | |
| AT3G14090 | EXO70D3 | 71 | | 1 | 5 | 3 | 9 | | | | | | | |
| AT3G29400 | **EXO70E1** | 75 | | | 2 | 2 | 4 | | | | | | | |
| AT5G50380 | **EXO70F1** | 77 | | | 28 | 16 | 44 | | | | 1.2E+05 | 8.1E+05 | 0.0E+00 | 9.3E+05 |
| AT4G31540 | **EXO70G1** | 78 | | | 4 | 4 | 8 | | | | | | | |
| AT5G59730 | EXO70H7 | 71 | | | 7 | 4 | 11 | | | | | | | |
| AT5G49830 | EXO84B | 90 | 7 | 2 | 38 | 25 | 72 | | | | | | | |

| | | | | | | | | | | | | | | |
|---|---|---|---|---|---|---|---|---|---|---|---|---|---|---|
| AT5G12370 | **SEC10** | 90 | 19 | 3 | 45 | 32 | 99 | | | | 2.9E+05 | 5.7E+05 | 0.0E+00 | 8.6E+05 |
| AT3G56640 | **SEC15A** | 90 | | | 8 | 5 | 13 | | | | | | | |
| AT4G02350 | SEC15B | 88 | 15 | 2 | 45 | 35 | 97 | | | | | | | |
| AT1G47550 | **SEC3A** | 100 | 18 | 1 | 31 | 19 | 69 | | | | 1.9E+05 | 2.1E+05 | 0.0E+00 | 4.0E+05 |
| AT1G76850 | **SEC5A** | 122 | 10 | 1 | 31 | 9 | 51 | | | | | | | |
| AT1G21170 | SEC5B | 123 | 2 | 0 | 17 | 11 | 30 | | | | | | | |
| AT3G10380 | Sec8 | 117 | 10 | 0 | 43 | 41 | 94 | | | | 1.3E+05 | 1.4E+05 | 0.0E+00 | 2.7E+05 |
| AT5G54750 | Bet3 | 21 | | | 3 | 3 | 6 | 0.08 | | 0.08 | | | | |
| AT3G05000 | Trs33 | 20 | | 1 | 4 | 2 | 7 | | | | | | | |
| AT5G16280 | Trs85 | 140 | | | 5 | 11 | 16 | | 0.18 | 0.18 | | | | |
| **Regulators** | | | | | | | | | | | | | | |
| AT3G59770 | SAC9 | 180 | 55 | 117 | 840 | 540 | 1,552 | 5.80 | 5.25 | 5.53 | 6.8E+06 | 8.0E+06 | 4.3E+05 | 1.5E+07 |
| AT1G77140 | VPS45 | 65 | 14 | 10 | 315 | 213 | 552 | | 5.25 | 5.25 | 5.9E+06 | 3.2E+06 | 2.1E+06 | 1.1E+07 |
| AT5G54310 | MTV4 | 53 | 13 | 12 | 97 | 74 | 196 | | | | 2.9E+06 | 2.5E+06 | 4.6E+06 | 9.9E+06 |
| AT3G60860 | BIG2 | 200 | | | 4 | 6 | 10 | | | | | | | |
| AT1G01960 | BIG3 | 195 | | | 6 | 4 | 10 | | | | 4.3E+04 | 4.4E+04 | 0.0E+00 | 8.7E+04 |
| AT5G06970 | PATROL1 | 125 | 21 | 6 | 70 | 44 | 141 | | | | 6.3E+05 | 1.4E+06 | 0.0E+00 | 2.1E+06 |
| AT1G71410 | SCYL2B | 99 | 57 | 137 | 406 | 194 | 794 | | 14.94 | 14.94 | 1.6E+07 | 1.7E+07 | 3.8E+06 | 3.7E+07 |
| AT3G20290 | EHD1 | 61 kD | 92 | 41 | 57 | 27 | 217 | | | | 2.2E+06 | 2.5E+06 | 0.0E+00 | 4.7E+06 |
| AT4G05520 | EHD2 | 61 kD | 117 | 52 | 101 | 81 | 351 | | | | 2.8E+07 | 3.4E+07 | 4.6E+04 | 6.2E+07 |
| AT1G22870 | SCYL2A | 100 | 31 | 49 | 131 | 72 | 283 | | | | 4.9E+05 | 1.3E+05 | 0.0E+00 | 6.2E+05 |
| AT4G34660 | SH3 domain-containing p | 41 kD | 49 | 306 | 304 | 268 | 927 | 7.13 | 9.40 | 8.27 | 1.9E+07 | 2.9E+07 | 8.5E+06 | 5.6E+07 |
| AT5G06140 | SNX1 | 47 | 52 | 73 | 177 | 162 | 464 | | 3.79 | 3.79 | 2.4E+07 | 2.6E+07 | 2.5E+06 | 5.3E+07 |
| AT5G58440 | SNX2A | 66 | | 12 | 86 | 83 | 181 | | | | 1.7E+06 | 2.4E+06 | 4.7E+05 | 4.6E+06 |
| AT5G07120 | SNX2B | 64 | 35 | 62 | 179 | 186 | 462 | | 4.48 | 4.48 | 1.6E+07 | 2.0E+07 | 3.2E+06 | 3.9E+07 |
| AT4G12770 | AUXILIN-LIKE2 | 98 | 29 | 202 | 315 | 264 | 810 | | | | 1.2E+06 | 1.3E+06 | 1.3E+06 | 3.9E+06 |
| AT4G12780 | AUXILIN-LIKE1 | 100 | 36 | 254 | 394 | 310 | 994 | 3.98 | 3.75 | 3.86 | 6.7E+05 | 1.9E+06 | 1.4E+06 | 4.0E+06 |
| AT1G06400 | RabA1a | 24 | 15 | 12 | 25 | 27 | 79 | | | | 4.8E+06 | 1.9E+06 | 0.0E+00 | 6.7E+06 |
| AT1G16920 | RabA1b | 24 | 31 | 39 | 80 | 97 | 247 | | | | 6.4E+06 | 4.3E+06 | 2.1E+05 | 1.1E+07 |
| AT5G45750 | RabA1c | 24 | 38 | 48 | 95 | 104 | 285 | | | | 5.0E+07 | 2.3E+07 | 2.2E+06 | 7.5E+07 |
| AT4G18800 | RabA1d | 24 | 23 | 44 | 81 | 90 | 238 | | 2.86 | 2.86 | 4.4E+06 | 1.7E+06 | 0.0E+00 | 6.0E+06 |
| AT3G15060 | RabA1g | 24 | 22 | 38 | 73 | 83 | 216 | | 2.86 | 2.86 | 3.5E+06 | 1.5E+06 | 0.0E+00 | 5.0E+06 |
| AT1G09630 | RabA2a | 24 | 17 | 25 | 46 | 41 | 129 | | | | | | | |
| AT1G07410 | RabA2b | 24 | 15 | 0 | 38 | 41 | 94 | | | | | | | |
| AT3G46830 | RabA2c | 24 | 26 | 33 | 56 | 45 | 160 | 0.15 | | 0.15 | 5.1E+06 | 5.5E+06 | 0.0E+00 | 1.1E+07 |
| AT5G65270 | RabA4a | 25 | 7 | 16 | 26 | 27 | 76 | | 0.46 | 0.46 | 6.6E+05 | | | 6.6E+05 |
| AT4G39990 | RabA4b | 24 | 5 | 14 | 23 | 20 | 62 | | | | 4.2E+06 | 1.8E+06 | 3.6E+05 | 6.4E+06 |
| AT5G47520 | RabA5a | 24 | | 11 | 34 | 64 | 109 | | | | | | | |
| AT3G07410 | RabA5b | 24 | 10 | 13 | 17 | 19 | 59 | | | | 1.3E+06 | 1.0E+06 | 0.0E+00 | 2.4E+06 |
| AT2G43130 | RabA5c | 24 | | 12 | 16 | 15 | 43 | | | | | | | |
| AT1G73640 | RabA6a | 26 | | 11 | 11 | 9 | 31 | | | | | | | |
| AT4G35860 | RabB1b | 23 | 6 | 8 | 14 | 20 | 48 | | | | 1.3E+06 | 9.4E+05 | 0.0E+00 | 2.2E+06 |
| AT4G17170 | RabB1c | 23 | 8 | 14 | 20 | 19 | 61 | | | | | | | |
| AT1G43890 | RabC1 | 24 | 13 | 18 | 44 | 35 | 110 | 0.15 | 0.66 | 0.41 | 1.1E+07 | 6.8E+06 | 5.5E+05 | 1.9E+07 |
| AT3G11730 | RabD1 | 23 | 12 | 26 | 36 | 30 | 104 | | | | | | | |

| | | | | | | | | | | | | | | |
|---|---|---|---|---|---|---|---|---|---|---|---|---|---|---|
| AT1G02130 | RabD2a | 23 | | 33 | 33 | 35 | 101 | | | | | | | |
| AT5G47200 | RabD2b | 22 | 29 | 103 | 136 | 86 | 354 | | | | 4.8E+07 | 3.0E+07 | 1.8E+06 | 8.0E+07 |
| AT3G53610 | RabE1a | 24 | 39 | 58 | 65 | 49 | 211 | | | | 1.1E+06 | 1.4E+06 | 5.4E+04 | 2.5E+06 |
| AT4G20360 | RabE1b | 52 | 8 | 18 | 66 | 52 | 144 | 0.15 | 0.40 | 0.27 | 2.1E+07 | 1.5E+07 | 1.1E+07 | 4.7E+07 |
| AT3G46060 | RabE1c | 24 | 51 | 91 | 101 | 82 | 325 | 0.58 | | 0.58 | 9.4E+06 | 7.8E+06 | 0.0E+00 | 1.7E+07 |
| AT5G03520 | RabE1d | 24 | 38 | 62 | 76 | 54 | 230 | | | | 3.2E+06 | 2.9E+06 | 0.0E+00 | 6.1E+06 |
| AT3G54840 | RabF1 | 22 | | 3 | 14 | 23 | 40 | | | | | | | |
| AT4G19640 | RabF2b | 22 | | 3 | 13 | 12 | 28 | | | | 1.6E+06 | 1.2E+06 | 1.4E+06 | 4.2E+06 |
| AT4G09720 | RabG3a | 23 | 1 | 9 | 23 | 17 | 50 | | 0.99 | 0.99 | 4.6E+06 | 3.3E+06 | 2.3E+05 | 8.1E+06 |
| AT3G16100 | RabG3c | 23 | | 13 | 18 | 23 | 54 | | | | | | | |
| AT1G49300 | RabG3e | 23 | 3 | 9 | 21 | 21 | 54 | 0.27 | | 0.27 | 4.1E+06 | 1.2E+06 | 0.0E+00 | 5.4E+06 |
| AT3G55020 | RABGAP | 87 | 3 | 0 | 2 | 1 | 6 | | | | | | | |
| AT2G44100 | RabGDI1 | 50 | 7 | 1 | 17 | 28 | 53 | | | | 2.8E+06 | 1.9E+06 | 0.0E+00 | 4.7E+06 |
| AT3G59920 | RabGDI2 | 50 | 19 | 3 | 28 | 50 | 100 | | 35.80 | 35.80 | 9.0E+06 | 5.6E+06 | 0.0E+00 | 1.5E+07 |
| AT1G08770 | PRA1.E | 22 | | | 6 | 5 | 11 | 0.83 | 1.18 | 1.00 | | | | |
| AT2G40380 | PRA1.B2 | 23 | 4 | 8 | 10 | 8 | 30 | | 3.86 | 3.86 | | | | |
| AT3G56110 | PRA1.B1 | 23 | 10 | 3 | 25 | 14 | 52 | | | | 7.9E+06 | 1.1E+07 | 0.0E+00 | 1.9E+07 |
| AT1G49040 | SCD1 | 132 | 2 | 0 | 11 | 6 | 19 | | | | | | | |
| AT3G48860 | SCD2 | 64 | 4 | 1 | 18 | 12 | 35 | | | | 1.0E+06 | 8.3E+05 | 0.0E+00 | 1.8E+06 |

**Supplemental Dataset 5. Predicted protein localization data for proteins more than two-fold depleted between DFGL and CCV fractions.** The consensus subcellular localization predicted by SUBA4 algorithm for the accession numbers corresponding to the 539 proteins which were more than two fold depleted between DFGL and CCV are shown, as well as the corresponding total spectral counts of these proteins in the LC/MS-MS dataset corresponding to CCVs digested in gel and average fold depletion. The annotated localizations collected from TAIR for the same accession numbers are additionally shown.

| | Average Fold Depletion Across Replicates | Total Spectral Count Across Four Replicates | SUBA Consensus Data | TAIR Consensus Data |
|---|---|---|---|---|
| AT1G50480 | 2.0034 | 45 | cytosol | plasma membrane:17317660;plas |
| AT2G44610 | 2.0038 | 141 | golgi | plasma membrane:17151019 |
| AT2G06210 | 2.0123 | 59 | nucleus | |
| AT3G59650 | 2.0125 | 20 | mitochondrion | |
| AT3G19760 | 2.0306 | 66 | nucleus | nucleus:14617066;nucleus:15496 |
| AT1G26880 | 2.0393 | 49 | cytosol,nucleus | plastid:15028209;nucleus:154964 |
| AT1G53750 | 2.0441 | 354 | cytosol,nucleus | plasma membrane:17317660 |
| AT1G07830 | 2.0447 | 21 | mitochondrion | |
| AT2G38040 | 2.0506 | 167 | plastid | plastid:15028209;plastid:186331 |
| AT5G23540 | 2.064125633 | 192 | cytosol,nucleus | |
| AT1G20220 | 2.0677 | 8 | nucleus | |
| AT4G26600 | 2.0734 | 61 | nucleus | nucleus:15496452 |
| AT3G16950 | 2.0816 | 782 | plastid | plastid:15028209;plastid:110562 |
| AT4G38800 | 2.093626999 | 12 | cytosol | plasma membrane:17644812 |
| AT3G02200 | 2.098373084 | 177 | cytosol,nucleus | |
| AT5G15810 | 2.0984 | 41 | mitochondrion | |
| AT3G57560 | 2.108718894 | 25 | plastid | plastid:16377628;plastid:184314 |
| AT5G58290 | 2.109332247 | 397 | cytosol,nucleus | |
| AT2G07696 | 2.117268423 | 28 | mitochondrion | |

| | | | | |
|---|---|---|---|---|
| AT3G58570 | 2.1206 | 205 | nucleus | |
| AT2G36130 | 2.1232 | 16 | cytosol | |
| AT1G27090 | 2.125 | 63 | nucleus | |
| AT3G18190 | 2.1277 | 82 | cytosol | |
| AT5G57110 | 2.1361 | 414 | plasma membrane | plasma membrane:16618929;plas[illegible] |
| AT2G33040 | 2.1602 | 27 | mitochondrion | nucleus:15610358;plastid:18431[illegible] |
| AT5G52280 | 2.1679 | 13 | nucleus | plastid:15028209 |
| AT3G12580 | 2.1703 | 600 | cytosol | mitochondrion:17137349;plasma[illegible] |
| AT5G65270 | 2.1808 | 76 | golgi | plasma membrane:17317660;plas[illegible] |
| AT1G30120 | 2.1808 | 40 | plastid | plastid:12766230;plastid:184314[illegible] |
| AT2G43070 | 2.1876 | 34 | vacuole | plasma membrane:17151019 |
| AT5G28060 | 2.198889934 | 140 | cytosol | cytosol:17934214 |
| AT3G17390 | 2.1994 | 280 | nucleus,plasma membrane | plasma membrane:17644812;nucl[illegible] |
| ATCG00770 | 2.1996 | 11 | plastid | plastid:18431481 |
| AT5G09810 | 2.200098902 | 669 | cytosol | plasma membrane:17317660;mito[illegible] |
| AT4G38510 | 2.203605819 | 1147 | vacuole | vacuole:15539469;plasma membr[illegible] |
| ATCG00820 | 2.2049 | 13 | plastid | plastid:18431481;plastid:16207[illegible] |
| AT3G61820 | 2.2248 | 73 | extracellular | |
| AT5G67630 | 2.2367 | 86 | nucleus | plastid:15028209;nucleus:15496[illegible] |
| AT5G39850 | 2.250174389 | 180 | cytosol | |
| AT2G21410 | 2.272605688 | 181 | vacuole | plastid:15028209;vacuole:16618[illegible] |
| AT1G71880 | 2.286 | 266 | plasma membrane | plasma membrane:17317660;plas[illegible] |
| AT5G05470 | 2.294044746 | 36 | cytosol | |
| AT3G16240 | 2.296494488 | 233 | vacuole | plastid:15028209;vacuole:15539[illegible] |
| AT2G07690 | 2.3006 | 103 | nucleus | nucleus:19357199 |
| AT3G11560 | 2.3012 | 25 | plastid | plastid:18431481;plastid:129389[illegible] |
| AT4G17530 | 2.3104668 | 345 | cytosol | vacuole:15539469;plasma membr[illegible] |
| AT3G22200 | 2.3105 | 16 | mitochondrion | plasma membrane:17151019;mito[illegible] |
| AT1G24280 | 2.3195 | 78 | plastid | |
| AT3G06720 | 2.3212 | 66 | nucleus | cytosol:18433157;nucleus:10428[illegible] |

| | | | | |
|---|---|---|---|---|
| AT3G22320 | 2.3354 | 23 | nucleus | |
| AT3G18420 | 2.3354 | 6 | plastid | plastid:12766230;plastid:184314 |
| AT5G22330 | 2.354127561 | 181 | nucleus | plastid:15028209;nucleus:154694 |
| AT2G42790 | 2.3674 | 51 | peroxisome | peroxisome:17951448 |
| AT2G31610 | 2.372265365 | 501 | cytosol | vacuole:15539469;plasma membr |
| AT5G12190 | 2.375872086 | 17 | nucleus | |
| AT1G49670 | 2.3796 | 92 | peroxisome | mitochondrion:14671022;peroxis |
| AT2G29550 | 2.3831 | 1606 | cytosol | plasma membrane:17317660;vacu |
| AT1G42440 | 2.3913 | 102 | mitochondrion | |
| AT2G01690 | 2.399178963 | 404 | golgi | plasma membrane:17151019 |
| AT1G24180 | 2.410498667 | 423 | mitochondrion | mitochondrion:18385124;mitoch |
| AT4G02930 | 2.424178622 | 247 | mitochondrion | mitochondrion:17137349;mitoch |
| AT5G22100 | 2.4352 | 51 | mitochondrion | |
| AT1G43710 | 2.4366 | 57 | cytosol | |
| AT5G62200 | 2.4384 | 20 | vacuole | plasma membrane:17151019 |
| AT2G34470 | 2.442 | 20 | cytosol | |
| AT5G36230 | 2.4445 | 70 | cytosol | |
| AT1G43890 | 2.448178336 | 110 | golgi | plasma membrane:17644812 |
| AT1G30580 | 2.448892015 | 61 | cytosol | |
| AT3G06610 | 2.4513 | 11 | cytosol | |
| AT3G05530 | 2.464608638 | 539 | cytosol,nucleus | nucleus:11742986 |
| AT5G46430 | 2.4682 | 156 | cytosol | cytosol:17934214 |
| AT2G27100 | 2.500374585 | 91 | nucleus | plastid:15028209;cytosol:184331 |
| AT3G59140 | 2.5081 | 28 | plasma membrane | plasma membrane:17151019 |
| AT5G55920 | 2.5505 | 143 | nucleus | |
| AT5G23740 | 2.5554 | 75 | cytosol | |
| AT1G80500 | 2.567 | 14 | golgi,plasma membrane | |
| AT5G02870 | 2.574295303 | 814 | cytosol | plastid:15028209;plasma membra |
| AT4G00810 | 2.58 | 17 | cytosol | nucleus:18433157;cytosol:18433 |
| AT1G75500 | 2.591083756 | 55 | vacuole | plasma membrane:15060130 |

| | | | | |
|---|---|---|---|---|
| AT1G31280 | 2.5954 | 26 | nucleus | |
| AT2G17510 | 2.6029 | 22 | cytosol | |
| AT5G02450 | 2.605241059 | 91 | cytosol | plasma membrane:17151019;cyto |
| AT1G08130 | 2.6143 | 28 | nucleus,mitochondrion | nucleus:16790030;mitochondrion |
| AT2G24820 | 2.618 | 4 | plastid | plastid:15028209;plastid:127662 |
| AT3G24830 | 2.649 | 264 | cytosol | cytosol:17934214 |
| AT3G25780 | 2.6792 | 15 | plasma membrane | plasma membrane:17151019;plas |
| AT3G62870 | 2.686527448 | 446 | cytosol | plasma membrane:17151019;cyto |
| AT5G02490 | 2.6875 | 754 | cytosol | plasma membrane:17317660 |
| AT1G04270 | 2.690447532 | 212 | cytosol | plasma membrane:17151019;cyto |
| ATCG01120 | 2.707 | 16 | plastid | plastid:18633119;plastid:184314 |
| AT1G29320 | 2.7252 | 52 | nucleus | |
| ATCG00790 | 2.7281 | 16 | plastid | plastid:18431481 |
| AT5G23900 | 2.730558483 | 100 | cytosol | cytosol:17934214 |
| AT2G17930 | 2.7432 | 222 | golgi | |
| AT5G14040 | 2.752148374 | 9 | mitochondrion | plastid:15028209;mitochondrion |
| AT4G29830 | 2.7562 | 20 | nucleus | |
| AT1G44575 | 2.7569 | 18 | plastid | plastid:15028209;plastid:184314 |
| AT2G26780 | 2.758 | 80 | cytosol,nucleus | |
| AT5G18620 | 2.7583 | 74 | nucleus | |
| AT3G15000 | 2.7618 | 20 | mitochondrion | mitochondrion:14671022;mitoch |
| AT2G47110 | 2.7689 | 1969 | cytosol | cytosol:17934214 |
| AT2G39990 | 2.769740872 | 93 | cytosol,nucleus | nucleus:14617066;nucleus:15610 |
| AT3G52090 | 2.7797 | 14 | nucleus | |
| AT3G53430 | 2.7801 | 135 | cytosol | cytosol:17934214 |
| AT5G35590 | 2.790065486 | 15 | cytosol,nucleus | plasma membrane:17151019 |
| AT1G50920 | 2.7938 | 249 | nucleus | |
| AT4G37870 | 2.8062 | 68 | cytosol | cytosol:18433157;nucleus:18433 |
| AT1G74470 | 2.8094 | 66 | plastid | plastid:15028209;plastid:153221 |
| AT1G20200 | 2.812859046 | 313 | cytosol,nucleus | plasma membrane:17317660 |

| | | | | |
|---|---|---|---|---|
| AT1G16350 | 2.8168 | 30 | peroxisome | |
| AT1G15420 | 2.81781863 | 31 | nucleus | |
| AT4G38130 | 2.8359 | 18 | nucleus | |
| AT2G29580 | 2.8413 | 29 | nucleus | |
| AT2G36880 | 2.842626516 | 150 | cytosol | plasma membrane:17317660 |
| AT2G43460 | 2.849252024 | 99 | cytosol | cytosol:17934214 |
| AT5G44340 | 2.854599565 | 1367 | cytosol | |
| AT3G55620 | 2.854948805 | 49 | cytosol | nucleus:15496452 |
| AT4G16630 | 2.8861 | 56 | nucleus | |
| AT5G18380 | 2.890185573 | 152 | cytosol | cytosol:11598216;plastid:184314 |
| AT4G10450 | 2.8929 | 79 | cytosol | cytosol:17934214 |
| AT3G52580 | 2.896078879 | 82 | cytosol | |
| AT1G59870 | 2.9054 | 294 | plasma membrane | mitochondrion:15276431;plasma |
| AT2G42520 | 2.912 | 235 | nucleus | peroxisome:17951448 |
| AT4G33510 | 2.9167 | 44 | plastid | plastid:18431481 |
| AT5G20650 | 2.924181664 | 19 | vacuole | vacuole:15539469;plasma membr |
| AT5G63120 | 2.9336 | 42 | nucleus | |
| AT5G62390 | 2.945 | 543 | endoplasmic reticulum | plasma membrane:17317660;endo |
| AT1G73180 | 2.9579 | 13 | cytosol | plasma membrane:17317660 |
| AT3G42660 | 2.9618 | 104 | nucleus | |
| AT5G27120 | 2.9637 | 82 | nucleus | nucleus:15496452 |
| AT1G35680 | 2.9652 | 9 | plastid | nucleus:14617066;plastid:186331 |
| AT3G15010 | 2.9705 | 25 | nucleus | nucleus:15496452 |
| AT5G15200 | 2.991999013 | 421 | cytosol | plastid:15028209;plasma membra |
| AT1G18540 | 3.001210897 | 343 | cytosol | plastid:15028209;endoplasmic ret |
| AT3G61650 | 3.033089504 | 111 | cytosol | plasma membrane:17317660;cyto |
| AT1G09270 | 3.0343 | 14 | nucleus | |
| AT5G20290 | 3.042126054 | 386 | cytosol,nucleus | plasma membrane:17317660;plast |
| AT1G72370 | 3.044897869 | 273 | cytosol | plasma membrane:17317660;nucl |
| AT4G22690 | 3.046630122 | 121 | endoplasmic reticulum | plasma membrane:17317660;mito |

| | | | | |
|---|---|---|---|---|
| AT1G72440 | 3.0572 | 62 | nucleus | |
| AT4G31580 | 3.076537928 | 39 | nucleus | nucleus:15686520 |
| AT1G56330 | 3.077710759 | 121 | endoplasmic reticulum,cy | plasma membrane:17317660;endo |
| AT5G06160 | 3.0798 | 10 | nucleus | cytosol:18433157 |
| AT4G05410 | 3.0892 | 27 | nucleus | nucleus:15496452 |
| AT5G62690 | 3.090930216 | 1785 | cytosol | plasma membrane:17644812;nucl |
| AT3G14990 | 3.092287442 | 314 | golgi | vacuole:15539469;plasma membr |
| AT2G01250 | 3.099465215 | 394 | cytosol | plastid:15028209;plasma membra |
| AT5G48370 | 3.1021 | 19 | peroxisome | peroxisome:12154131 |
| AT1G15750 | 3.1342 | 7 | cytosol | nucleus:16763149;cytosol:18433 |
| AT5G41520 | 3.144137787 | 122 | cytosol | cytosol:11598216;plastid:184314 |
| AT1G53500 | 3.1543 | 143 | cytosol | |
| AT1G74050 | 3.171076829 | 445 | cytosol | plasma membrane:17317660;cyto |
| AT2G20450 | 3.174577784 | 231 | cytosol | cytosol:17934214;endoplasmic re |
| AT2G22780 | 3.1962 | 5 | peroxisome | peroxisome:12154131;plastid:184 |
| AT3G49240 | 3.1987 | 43 | mitochondrion | mitochondrion:17137349;plastid |
| AT1G77180 | 3.2019 | 56 | nucleus | nucleus:14617066;nucleus:15496 |
| AT3G42050 | 3.226897769 | 345 | golgi,vacuole | vacuole:15539469;plasma membr |
| AT5G22360 | 3.2285 | 52 | golgi | vacuole:14760709 |
| AT3G22660 | 3.232463491 | 19 | nucleus | |
| AT5G11240 | 3.2331 | 87 | nucleus | |
| AT1G02690 | 3.259264413 | 45 | nucleus | |
| AT1G54270 | 3.2595 | 653 | cytosol | plasma membrane:17644812;cyto |
| AT5G39950 | 3.2762 | 16 | cytosol | plasma membrane:17317660 |
| AT4G23850 | 3.286861864 | 66 | plasma membrane | plasma membrane:17317660 |
| AT2G30970 | 3.294922365 | 149 | mitochondrion | mitochondrion:18385124;mitoch |
| AT2G20230 | 3.302 | 31 | vacuole | vacuole:15539469;plasma membr |
| AT3G03920 | 3.30523086 | 19 | nucleus | plastid:15322131;nucleus:15496[illegible] |
| AT1G02780 | 3.31016034 | 190 | cytosol | plasma membrane:17317660;cyto |
| ATCG00780 | 3.3123 | 9 | plastid | plastid:18431481 |

| | | | | |
|---|---|---|---|---|
| AT4G02840 | 3.3169 | 45 | nucleus | plasma membrane:17151019;nucl |
| AT5G49160 | 3.3199 | 6 | nucleus | |
| AT4G21150 | 3.327161071 | 16 | endoplasmic reticulum | mitochondrion:15276431;plasma |
| AT3G22520 | 3.3362 | 112 | nucleus | plastid:18633119;plastid:184314 |
| AT2G46280 | 3.34273682 | 104 | cytosol | |
| AT4G12650 | 3.36 | 464 | golgi | golgi:16618929;plasma membrane |
| AT3G49910 | 3.363667224 | 168 | cytosol | plastid:15028209;nucleus:14617( |
| AT4G39080 | 3.364166538 | 339 | vacuole | vacuole:15539469;plasma membr |
| AT1G07660 | 3.368792052 | 217 | nucleus | plasma membrane:17151019;plas |
| AT3G22330 | 3.3998 | 105 | mitochondrion | mitochondrion:11743115;nucleu: |
| AT4G31700 | 3.400401054 | 271 | cytosol | plasma membrane:17317660;plas |
| AT2G02040 | 3.4016 | 79 | vacuole | plasma membrane:17151019;vacu |
| AT5G27770 | 3.40224735 | 77 | cytosol | plasma membrane:17317660;cyto |
| AT3G04920 | 3.437949032 | 135 | cytosol | plasma membrane:17317660;plas |
| AT1G78900 | 3.443851704 | 1815 | golgi,vacuole | plastid:15028209;vacuole:155394 |
| AT1G60200 | 3.447 | 39 | nucleus | |
| AT5G49930 | 3.4562 | 47 | nucleus | |
| AT1G62020 | 3.4586 | 83 | cytosol | plasma membrane:17317660 |
| AT2G31060 | 3.4815 | 29 | mitochondrion | |
| AT5G46280 | 3.4888 | 127 | nucleus | nucleus:19357199 |
| AT5G58070 | 3.490338462 | 49 | endoplasmic reticulum | vacuole:15539469;mitochondrior |
| AT4G20850 | 3.498840951 | 1449 | plastid | plastid:15028209;vacuole:155394 |
| AT1G63660 | 3.5114 | 105 | cytosol | |
| AT2G37860 | 3.5231 | 22 | plastid | plastid:18431481 |
| AT4G39460 | 3.5311 | 4 | plastid | plastid:16618929;plastid:127662 |
| AT5G60980 | 3.534115672 | 11 | nucleus | |
| AT1G80410 | 3.537539589 | 247 | cytosol | |
| AT1G56110 | 3.5509 | 149 | nucleus | nucleus:15610358;nucleus:15496 |
| ATCG00480 | 3.553235834 | 97 | plastid | plastid:16414959;plastid:169230 |
| AT4G29130 | 3.5623 | 31 | mitochondrion | nucleus:17081979;plasma membr |

| | | | | |
|---|---|---|---|---|
| AT2G34660 | 3.5726 | 139 | vacuole | plasma membrane:17151019;vacu |
| AT2G19740 | 3.5745 | 39 | cytosol | plasma membrane:17317660;plas |
| AT2G21160 | 3.5837 | 5 | endoplasmic reticulum | plasma membrane:17644812;plas |
| AT1G80930 | 3.5888 | 62 | nucleus | nucleus:18433157;cytosol:18433 |
| AT3G52200 | 3.5952 | 296 | mitochondrion | mitochondrion:17137349;mitoch |
| AT4G23630 | 3.609587063 | 36 | plasma membrane | vacuole:15539469;plasma membr |
| AT2G33340 | 3.6197 | 109 | nucleus | plastid:15028209 |
| AT4G35300 | 3.6203 | 78 | vacuole | vacuole:15539469;plasma membr |
| AT1G58380 | 3.646941808 | 371 | cytosol | plasma membrane:17317660;plas |
| AT3G45970 | 3.6561 | 5 | extracellular | |
| AT4G20360 | 3.65964898 | 144 | plastid | plastid:15322131;nucleus:14617 |
| AT5G02500 | 3.667468719 | 951 | cytosol | plasma membrane:17317660;nucl |
| AT1G49300 | 3.670263525 | 54 | plasma membrane | plasma membrane:17151019 |
| AT3G48930 | 3.682707702 | 128 | cytosol | cytosol:17934214 |
| AT2G47800 | 3.685866157 | 134 | plasma membrane | plasma membrane:17151019;plas |
| AT5G52820 | 3.7012 | 42 | nucleus | |
| AT2G18720 | 3.7107 | 59 | cytosol | |
| ATCG00380 | 3.7113 | 13 | plastid | plastid:18633119;plastid:184314 |
| AT1G59900 | 3.728256426 | 161 | mitochondrion | mitochondrion:18385124;nucleus |
| AT3G28500 | 3.744595146 | 80 | cytosol | plasma membrane:17317660;cyto |
| AT3G09440 | 3.7635 | 619 | cytosol | plasma membrane:17317660;vacu |
| AT1G22410 | 3.7719 | 73 | plastid | |
| AT1G04480 | 3.7819 | 113 | cytosol,nucleus | |
| AT5G18520 | 3.787 | 222 | golgi | |
| AT2G32060 | 3.8158 | 20 | cytosol | nucleus:14617066;cytosol:17934 |
| AT5G26710 | 3.822854707 | 175 | cytosol | |
| AT3G02080 | 3.826282679 | 113 | cytosol | cytosol:17934214 |
| AT1G51650 | 3.8306 | 7 | mitochondrion | mitochondrion:14671022;mitoch |
| AT3G13870 | 3.830920191 | 11 | endoplasmic reticulum,go | plasma membrane:17151019;endo |
| AT3G63410 | 3.840246837 | 21 | plastid | plastid:15028209;plastid:127872 |

| | | | | |
|---|---|---|---|---|
| AT4G21110 | 3.8416 | 15 | nucleus | |
| AT1G27400 | 3.848122896 | 165 | cytosol,nucleus,plastid | plasma membrane:17317660;plas |
| AT1G71380 | 3.8618 | 45 | extracellular | plasma membrane:17317660 |
| AT1G80750 | 3.8686215 | 35 | cytosol | cytosol:17934214;nucleus:15496 |
| AT1G53210 | 3.882613783 | 339 | plasma membrane | vacuole:15539469;plasma membr |
| AT1G07810 | 3.887 | 29 | endoplasmic reticulum | plasma membrane:17317660;endo |
| AT3G45030 | 3.894020575 | 45 | cytosol | cytosol:17934214 |
| AT1G09590 | 3.894649043 | 145 | cytosol | plastid:15028209;cytosol:179342 |
| AT4G16566 | 3.903 | 9 | peroxisome | peroxisome:17951448 |
| AT3G06730 | 3.91 | 11 | plastid | plastid:18431481 |
| AT4G22010 | 3.9116 | 53 | extracellular | |
| AT5G61140 | 3.918 | 4 | cytosol,golgi | |
| AT4G36130 | 3.932668597 | 275 | cytosol | plasma membrane:17317660;cyto |
| AT1G34430 | 3.934000049 | 727 | plastid | plasma membrane:17317660;plas |
| AT1G52360 | 3.9386 | 33 | cytosol | |
| AT4G39980 | 3.944 | 35 | plastid | |
| AT5G24710 | 3.9445 | 255 | nucleus,plasma membrane | plasma membrane:17317660 |
| AT2G27710 | 3.944757603 | 218 | cytosol | plasma membrane:17317660;cyto |
| AT4G03430 | 3.945240068 | 26 | nucleus | nucleus:16751345 |
| AT2G22125 | 3.9745 | 159 | plasma membrane | plasma membrane:17644812 |
| AT1G04810 | 3.9903 | 177 | cytosol,nucleus | plasma membrane:17317660 |
| AT4G32520 | 3.993 | 67 | plastid | plastid:18431481 |
| AT2G01750 | 3.9973 | 34 | cytosol | plasma membrane:17317660 |
| AT5G16130 | 3.999709796 | 134 | cytosol | plasma membrane:17317660;cyto |
| AT3G09630 | 4.005702798 | 859 | cytosol | plasma membrane:17317660;plas |
| AT4G34450 | 4.006882633 | 210 | cytosol | plastid:18431481 |
| AT3G56150 | 4.009907853 | 373 | cytosol,nucleus | nucleus:18433157;cytosol:18433 |
| AT5G08540 | 4.0118 | 30 | plastid | plastid:12766230;plastid:184314 |
| AT5G19770 | 4.013244263 | 820 | cytosol | plasma membrane:17317660;cyto |
| AT3G18600 | 4.0141 | 41 | nucleus | nucleus:15496452 |

| | | | | |
|---|---|---|---|---|
| AT1G63810 | 4.017 | 19 | cytosol | |
| AT2G45810 | 4.0246 | 69 | nucleus | |
| AT3G04840 | 4.031883324 | 442 | cytosol | plasma membrane:17317660;plas |
| AT3G60240 | 4.0538 | 13 | No consensus | |
| AT2G30860 | 4.056136935 | 21 | cytosol | vacuole:15539469;plastid:186331 |
| AT3G57330 | 4.0716 | 182 | vacuole | plasma membrane:17151019;plas |
| AT5G27850 | 4.073287264 | 284 | cytosol | cytosol:17934214;vacuole:15215 |
| AT2G18900 | 4.0804 | 44 | nucleus | |
| AT4G29430 | 4.086 | 22 | cytosol | mitochondrion:11971146;cytosol |
| AT4G15000 | 4.108009581 | 143 | cytosol | cytosol:17934214 |
| AT1G04820 | 4.111469961 | 1082 | cytosol | plastid:15028209;plasma membra |
| AT1G07770 | 4.120525703 | 329 | cytosol | plasma membrane:17317660;mito |
| AT5G22060 | 4.131454856 | 155 | cytosol | plasma membrane:17317660 |
| AT2G27720 | 4.144340935 | 74 | cytosol | nucleus:14617066;nucleus:18433 |
| AT2G40360 | 4.1855 | 130 | nucleus | |
| AT3G05560 | 4.192288457 | 80 | cytosol | plasma membrane:17317660;cyto |
| AT1G48830 | 4.200591907 | 153 | cytosol | plasma membrane:17317660;plas |
| AT3G60245 | 4.2223 | 104 | cytosol | cytosol:17934214 |
| AT4G15770 | 4.2279 | 52 | cytosol | |
| AT1G31230 | 4.2383 | 4 | plastid | plastid:18633119;plastid:184314 |
| AT2G42740 | 4.250493661 | 275 | cytosol | plastid:15028209;cytosol:179342 |
| AT2G43770 | 4.2644 | 50 | nucleus | |
| AT5G57870 | 4.286084171 | 62 | cytosol,nucleus | nucleus:15610358 |
| AT2G20580 | 4.291606305 | 547 | cytosol,nucleus | plasma membrane:17317660 |
| AT3G49010 | 4.295250487 | 386 | cytosol | plasma membrane:17317660;plas |
| AT1G10490 | 4.2997 | 96 | nucleus | |
| AT1G43170 | 4.306234924 | 883 | cytosol | plasma membrane:17317660;plas |
| AT3G07050 | 4.3081 | 78 | nucleus | nucleus:15496452 |
| AT1G15500 | 4.324833315 | 41 | plastid | plastid:16618929;plastid:127662 |
| AT5G14520 | 4.3275 | 125 | nucleus | |

| | | | | |
|---|---|---|---|---|
| AT3G52990 | 4.3327 | 347 | cytosol | |
| AT1G15440 | 4.3442 | 24 | nucleus | |
| AT4G00630 | 4.3474 | 6 | plastid | plastid:18431481;plastid:129389 |
| AT5G10360 | 4.34897629 | 274 | cytosol | plasma membrane:17317660;cyto |
| AT5G06360 | 4.3505 | 32 | cytosol,nucleus | |
| AT3G13580 | 4.3585 | 318 | cytosol | plastid:15028209;cytosol:179342 |
| AT5G57490 | 4.3923 | 5 | mitochondrion | mitochondrion:14671022;mitoch |
| AT4G39260 | 4.4008 | 17 | nucleus | plastid:15028209;peroxisome:179 |
| AT3G55410 | 4.4043 | 123 | mitochondrion | mitochondrion:14671022 |
| ATCG00160 | 4.4153 | 12 | plastid | plastid:18633119;plastid:184314 |
| AT2G27530 | 4.415431231 | 227 | cytosol | cytosol:17934214 |
| AT5G20010 | 4.423021254 | 149 | cytosol | plasma membrane:17644812 |
| AT5G52650 | 4.425425103 | 84 | cytosol | cytosol:17934214 |
| AT5G14030 | 4.440162206 | 5 | endoplasmic reticulum | plasma membrane:17151019 |
| AT1G52280 | 4.4518 | 45 | golgi,plasma membrane,va | vacuole:15539469;plasma membr |
| AT4G02510 | 4.4698 | 53 | plastid | plastid:15028209;plastid:106466 |
| AT5G48760 | 4.4711 | 117 | cytosol | cytosol:17934214 |
| AT5G62190 | 4.4749 | 65 | nucleus | nucleus:9121476;nucleus:151331 |
| AT5G16840 | 4.5196 | 4 | cytosol | |
| AT1G30400 | 4.529289202 | 255 | vacuole | vacuole:15539469;plasma membr |
| AT1G09770 | 4.534687274 | 95 | nucleus | plastid:15028209 |
| AT2G05920 | 4.56659464 | 303 | extracellular | |
| AT3G27300 | 4.600443058 | 86 | cytosol | plastid:18431481 |
| AT3G12800 | 4.60635221 | 11 | peroxisome | plasma membrane:17151019;perc |
| AT3G05060 | 4.6338 | 89 | nucleus | nucleus:15496452 |
| AT3G58750 | 4.636 | 70 | peroxisome | peroxisome:15923350 |
| AT1G31660 | 4.6507 | 31 | nucleus | |
| AT4G16130 | 4.6769 | 61 | cytosol | |
| AT1G47530 | 4.6979 | 26 | vacuole | plasma membrane:17151019 |
| AT3G02530 | 4.7133 | 64 | cytosol | |

| | | | | |
|---|---|---|---|---|
| AT1G65030 | 4.7352 | 34 | nucleus | nucleus:15496452 |
| AT4G13930 | 4.735664748 | 261 | cytosol | plasma membrane:17644812 |
| AT1G28110 | 4.739473152 | 21 | extracellular | |
| AT1G48630 | 4.7395 | 109 | cytosol | |
| AT3G23400 | 4.7466 | 5 | plastid | plastid:15028209;plastid:164149 |
| AT1G22780 | 4.750295772 | 222 | cytosol | plasma membrane:17151019;cyto |
| AT3G54540 | 4.7584 | 57 | nucleus | |
| AT2G03270 | 4.788932537 | 148 | nucleus | |
| AT1G64550 | 4.814210263 | 90 | cytosol | |
| AT1G33120 | 4.815369003 | 458 | cytosol | plasma membrane:17317660;plas |
| AT4G11420 | 4.8156 | 346 | cytosol,nucleus | plasma membrane:17317660 |
| AT5G11150 | 4.826622484 | 101 | vacuole | vacuole:14760709 |
| AT1G69620 | 4.843318642 | 69 | cytosol | plasma membrane:17151019;cyto |
| AT2G18020 | 4.865422551 | 349 | cytosol | plastid:15028209;plasma membra |
| AT3G61110 | 4.9101 | 41 | cytosol | cytosol:17934214 |
| AT1G01100 | 4.913800906 | 73 | cytosol | plasma membrane:17317660;nucl |
| AT1G15690 | 4.922632236 | 765 | vacuole | plastid:15028209;vacuole:15539 |
| AT3G62700 | 4.9428 | 140 | vacuole | vacuole:15539469;plasma membr |
| AT5G14120 | 4.9434 | 63 | vacuole | vacuole:15539469;plasma membr |
| AT1G51980 | 4.9468 | 23 | mitochondrion | plastid:15028209;mitochondrion |
| AT5G05000 | 4.970144781 | 22 | plastid | plastid:15028209;plastid:129389 |
| AT3G50590 | 4.9844 | 183 | nucleus,plasma membrane | plasma membrane:17317660 |
| AT1G55150 | 4.988 | 76 | cytosol | |
| AT4G34670 | 4.989344186 | 453 | cytosol | plasma membrane:17317660;cyto |
| AT5G27640 | 4.993011499 | 149 | cytosol | nucleus:18433157;cytosol:18433 |
| AT2G24050 | 5.0039 | 45 | nucleus | |
| AT1G76300 | 5.0144 | 23 | nucleus | nucleus:16839879;nucleus:15496 |
| AT5G52470 | 5.0392 | 99 | nucleus | nucleus:10829025;nucleus:15496 |
| AT2G41840 | 5.071041222 | 382 | cytosol | cytosol:11598216;nucleus:15496 |
| AT5G06110 | 5.0758 | 61 | nucleus | |

| | | | | |
|---|---|---|---|---|
| AT3G25860 | 5.0788418 | 531 | plastid | plastid:10364395;plastid:184314 |
| AT5G28540 | 5.101817854 | 187 | endoplasmic reticulum | plastid:15028209;vacuole:155394 |
| AT4G08950 | 5.106730671 | 15 | extracellular | |
| AT3G58510 | 5.124043403 | 261 | nucleus | plasma membrane:17317660;perc |
| AT3G48870 | 5.17340477 | 98 | plastid | plastid:15028209;plastid:145931 |
| AT5G15550 | 5.1799 | 60 | nucleus | nucleus:15496452 |
| AT5G63400 | 5.191578154 | 6 | mitochondrion | mitochondrion:15276431;plasma |
| AT1G51060 | 5.197538582 | 42 | nucleus | nucleus:15496452 |
| AT3G09500 | 5.213241985 | 133 | cytosol | cytosol:17934214;nucleus:15496 |
| AT4G16720 | 5.213769151 | 161 | cytosol | plasma membrane:17151019;cyto |
| AT5G04830 | 5.21489205 | 52 | cytosol,peroxisome | |
| AT4G21710 | 5.2573 | 107 | nucleus | |
| AT1G04170 | 5.2577 | 110 | cytosol | |
| AT4G16155 | 5.258660936 | 827 | plastid | plastid:11056213;plastid:184314 |
| AT4G30800 | 5.2642 | 42 | cytosol | cytosol:11598216 |
| AT4G10320 | 5.274010221 | 80 | cytosol | |
| AT5G26742 | 5.282525655 | 91 | plastid | plastid:18431481;plastid:162077 |
| AT3G52140 | 5.2878 | 259 | cytosol | plasma membrane:17317660;plas |
| AT5G57290 | 5.28829471 | 72 | cytosol | nucleus:18433157 |
| AT2G21390 | 5.2998 | 76 | cytosol | |
| AT3G07110 | 5.302437826 | 229 | cytosol | plastid:15028209;cytosol:179342 |
| AT5G66680 | 5.3127 | 5 | endoplasmic reticulum | vacuole:15539469;plasma membr |
| AT3G20050 | 5.3232 | 90 | cytosol | |
| AT5G40200 | 5.3293 | 12 | nucleus | plastid:15028209;nucleus:154964 |
| AT1G06220 | 5.332888859 | 307 | nucleus | plasma membrane:17317660;nucl |
| AT4G11010 | 5.3338 | 18 | mitochondrion | mitochondrion:18385124;plastid |
| AT5G24650 | 5.338621089 | 82 | plastid | plasma membrane:17151019;plas |
| AT3G06650 | 5.365305802 | 45 | cytosol | |
| AT5G49890 | 5.3798 | 32 | vacuole | plasma membrane:17151019 |
| AT5G14610 | 5.3927 | 39 | nucleus | |

| | | | | |
|---|---|---|---|---|
| AT5G16280 | 5.406 | 16 | golgi | |
| AT3G48890 | 5.453587081 | 7 | endoplasmic reticulum | nucleus:18433157;plastid:153221 |
| AT3G43520 | 5.463606311 | 11 | plastid | plastid:18431481;plastid:129389 |
| AT5G13680 | 5.4743 | 23 | cytosol | |
| AT5G16120 | 5.474952094 | 72 | cytosol | |
| AT1G04690 | 5.478 | 36 | plasma membrane | plasma membrane:17644812 |
| AT5G47700 | 5.481174594 | 68 | cytosol | plasma membrane:17317660;nucl |
| AT1G67120 | 5.4856 | 26 | nucleus | plastid:12938931 |
| AT5G14220 | 5.486710843 | 42 | plastid,mitochondrion | plastid:12938931;plastid:166189 |
| AT4G18465 | 5.5013 | 62 | nucleus | |
| AT1G36180 | 5.5481 | 49 | plastid | |
| AT2G07698 | 5.550139343 | 146 | plastid,mitochondrion | mitochondrion:15276431;mitoch |
| AT5G01500 | 5.562353988 | 13 | plastid | plastid:17261580;plastid:166189 |
| AT1G61570 | 5.5744 | 5 | mitochondrion | mitochondrion:14671022;mitoch |
| AT2G34480 | 5.584526213 | 384 | cytosol | plasma membrane:17317660;cyto |
| AT2G32730 | 5.592039821 | 183 | cytosol,nucleus | cytosol:18433157 |
| AT5G20920 | 5.595970901 | 52 | cytosol | plasma membrane:17317660 |
| AT5G65010 | 5.5986 | 42 | golgi | |
| AT4G03560 | 5.6182 | 52 | vacuole | plasma membrane:17151019;vacu |
| AT1G62390 | 5.6188 | 41 | nucleus | |
| AT2G40290 | 5.619697018 | 78 | cytosol | nucleus:14617066 |
| AT1G52930 | 5.6273 | 39 | nucleus | |
| AT2G34040 | 5.640015372 | 54 | cytosol | |
| AT1G50200 | 5.6812 | 97 | mitochondrion | plastid:16251277;mitochondrion |
| AT1G02500 | 5.689900427 | 134 | cytosol | plasma membrane:17317660 |
| AT5G40770 | 5.733262709 | 5 | mitochondrion | vacuole:15539469;mitochondrion |
| AT2G44120 | 5.734531804 | 395 | cytosol | plastid:15028209;cytosol:179342 |
| AT3G11400 | 5.7725 | 21 | cytosol | plasma membrane:17317660 |
| AT3G18740 | 5.8729 | 141 | cytosol | cytosol:17934214 |
| AT1G64090 | 5.883045064 | 45 | endoplasmic reticulum | |

| | | | | |
|---|---|---|---|---|
| AT3G11250 | 5.909816205 | 219 | cytosol | |
| AT1G80070 | 5.940073457 | 323 | nucleus | plastid:15028209 |
| AT3G02260 | 5.9747 | 540 | nucleus | |
| AT1G26340 | 5.979430758 | 7 | plastid | plastid:17322552 |
| AT2G28000 | 5.985987524 | 38 | plastid | plastid:15028209;mitochondrion |
| AT5G61170 | 6.0078 | 102 | cytosol | vacuole:15539469;cytosol:17934 |
| AT5G26830 | 6.027 | 50 | plastid,mitochondrion | plasma membrane:17317660;plas |
| AT5G27670 | 6.052902367 | 20 | nucleus | nucleus:15496452 |
| AT2G38770 | 6.0952 | 129 | cytosol | plasma membrane:17317660;plas |
| AT1G42960 | 6.135658508 | 53 | plastid | plastid:15322131;plastid:129389 |
| AT4G19210 | 6.136668875 | 133 | cytosol | |
| AT3G18130 | 6.140620203 | 126 | cytosol | nucleus:15496452 |
| AT1G33590 | 6.144371203 | 161 | extracellular | plastid:15028209 |
| AT3G55200 | 6.168435192 | 103 | nucleus | plastid:15028209 |
| AT4G16143 | 6.1911901 | 61 | nucleus | cytosol:18433157;nucleus:15496 |
| AT1G66580 | 6.193816345 | 239 | cytosol | plasma membrane:17151019;cyto |
| AT3G07230 | 6.2365 | 4 | cytosol | |
| AT3G57660 | 6.2712 | 92 | nucleus | nucleus:14617066 |
| AT3G13930 | 6.289732377 | 827 | mitochondrion | plastid:15028209;mitochondrion |
| AT4G27090 | 6.294760282 | 296 | cytosol | plastid:15028209;endoplasmic ret |
| AT4G30190 | 6.3029 | 1317 | plasma membrane | plasma membrane:14506206;plas |
| AT5G22650 | 6.345 | 10 | nucleus | plasma membrane:17151019;cyto |
| AT3G22230 | 6.4206 | 127 | cytosol | cytosol:17934214 |
| AT3G23990 | 6.429339359 | 23 | mitochondrion | mitochondrion:16407270;mitoch |
| AT5G08180 | 6.4428 | 28 | nucleus | nucleus:15496452 |
| AT3G46830 | 6.487187804 | 160 | cytosol | plasma membrane:17317660;cyto |
| AT1G09760 | 6.499098093 | 35 | nucleus | plastid:15028209;nucleus:14617( |
| AT4G24820 | 6.508617104 | 222 | cytosol,nucleus | plasma membrane:17317660 |
| AT1G11910 | 6.514571152 | 115 | vacuole | vacuole:15215502 |
| AT3G09200 | 6.527100576 | 456 | cytosol | plastid:15028209;plasma membra |

| | | | | |
|---|---|---|---|---|
| AT3G46740 | 6.578550784 | 79 | plastid | mitochondrion:16618929;plastid |
| AT1G64790 | 6.5954 | 282 | golgi | nucleus:18433157;cytosol:18433 |
| AT1G20960 | 6.6637 | 286 | nucleus | nucleus:15496452 |
| AT3G57150 | 6.6911 | 99 | nucleus | cytosol:18433157;nucleus:15496 |
| AT4G27585 | 6.701127073 | 6 | mitochondrion | mitochondrion:18385124;mitoch |
| AT4G39200 | 6.7422 | 91 | cytosol,nucleus | cytosol:17934214 |
| AT5G27540 | 6.7595 | 19 | mitochondrion | mitochondrion:18344283;mitoch |
| AT1G08360 | 6.7764 | 107 | cytosol | plasma membrane:17317660;cyto |
| AT4G30990 | 6.7874 | 67 | nucleus | golgi:16618929 |
| AT5G53560 | 6.799469004 | 12 | endoplasmic reticulum | vacuole:15539469;plasma membr |
| AT2G19730 | 6.8002 | 41 | cytosol | plasma membrane:17317660;plas |
| AT1G07920 | 6.892935242 | 3605 | cytosol | plastid:15028209;mitochondrion |
| AT5G25460 | 6.9266 | 40 | extracellular | |
| AT3G13920 | 6.928965918 | 645 | cytosol | cytosol:18433157;nucleus:15496 |
| AT1G56070 | 7.042152964 | 306 | cytosol | cytosol:18433157;plastid:184314 |
| AT1G70770 | 7.1009 | 10 | endoplasmic reticulum | plasma membrane:17317660;nucl |
| AT2G03510 | 7.132405788 | 47 | endoplasmic reticulum | plasma membrane:17151019;endo |
| AT5G50850 | 7.146603761 | 366 | mitochondrion | mitochondrion:15276431;plasma |
| AT1G06530 | 7.200460829 | 29 | mitochondrion | mitochondrion:14671022;plasma |
| AT5G42020 | 7.223411075 | 191 | endoplasmic reticulum | plasma membrane:17317660;vacu |
| AT2G47250 | 7.2289 | 55 | nucleus | plastid:12938931 |
| AT3G11510 | 7.316124076 | 99 | cytosol | cytosol:11598216;nucleus:15496 |
| AT2G21640 | 7.369196758 | 4 | mitochondrion | mitochondrion:12492832 |
| AT1G31970 | 7.385491985 | 128 | nucleus | |
| AT1G04510 | 7.4395 | 82 | nucleus | nucleus:15496452 |
| AT5G47930 | 7.4481 | 26 | cytosol | plasma membrane:17317660;cyto |
| AT1G77940 | 7.5407 | 112 | cytosol | cytosol:17934214 |
| AT1G03230 | 7.566100454 | 138 | extracellular | |
| AT1G07670 | 7.6087 | 34 | endoplasmic reticulum | plasma membrane:17317660;endo |
| AT3G13300 | 7.6205 | 34 | cytosol | nucleus:18433157;cytosol:18433 |

| | | | | |
|---|---|---|---|---|
| AT5G48810 | 7.685213649 | 8 | endoplasmic reticulum | endoplasmic reticulum:16618929 |
| AT3G06530 | 7.7128 | 75 | plasma membrane | vacuole:15539469;plastid:184314 |
| AT3G08580 | 7.7168 | 23 | mitochondrion | plastid:15028209;mitochondrion |
| AT3G44890 | 7.7168 | 3 | plastid | plastid:15028209;plastid:162077 |
| AT5G43970 | 7.775 | 5 | mitochondrion | plasma membrane:17151019;plas |
| AT3G47520 | 7.840024349 | 35 | plastid | plastid:15028209;plastid:169230 |
| AT3G54110 | 7.91927743 | 4 | mitochondrion | plastid:15028209;mitochondrion |
| AT3G51260 | 8.001920461 | 18 | cytosol,nucleus | plastid:15028209;cytoskeleton:11 |
| AT1G15930 | 8.0722 | 35 | cytosol | cytosol:17934214 |
| AT4G22240 | 8.141896557 | 10 | plastid | plastid:16414959;plastid:164613 |
| AT4G31300 | 8.1862 | 4 | cytosol,nucleus | plasma membrane:17151019;vacu |
| AT5G16660 | 8.208892875 | 17 | plastid | plastid:18431481;plastid:147299 |
| AT5G64270 | 8.2497 | 73 | nucleus | plastid:15028209 |
| AT5G61790 | 8.251381632 | 29 | endoplasmic reticulum | vacuole:15539469;mitochondrion |
| AT1G53850 | 8.291186469 | 7 | cytosol,nucleus | plasma membrane:17151019 |
| AT1G55210 | 8.304268394 | 5 | extracellular | |
| AT5G25100 | 8.3141 | 223 | golgi | plasma membrane:17317660;golg |
| AT3G02090 | 8.314168702 | 104 | mitochondrion | plastid:15028209;mitochondrion |
| AT4G31985 | 8.3833 | 35 | cytosol,nucleus | cytosol:17934214 |
| AT5G64420 | 8.415383321 | 139 | nucleus | |
| AT1G18080 | 8.496 | 61 | cytosol | plastid:18431481 |
| AT4G00100 | 8.605 | 171 | cytosol | cytosol:10849343;nucleus:15496 |
| AT2G18740 | 8.626 | 33 | nucleus | |
| AT3G10090 | 8.660935361 | 10 | cytosol | cytosol:17934214 |
| AT1G21720 | 8.710042679 | 5 | cytosol,nucleus | plasma membrane:17151019 |
| AT3G22370 | 8.798587224 | 15 | mitochondrion | mitochondrion:15276431;plastid |
| AT1G78570 | 8.84173298 | 117 | cytosol | |
| AT1G06950 | 8.870609121 | 51 | plastid | plastid:15028209;plastid:127662 |
| ATCG00120 | 8.934224976 | 63 | plastid | plastid:16923014;plastid:164613 |
| AT1G14320 | 8.962922393 | 247 | cytosol | plastid:15028209;plasma membra |

| | | | | |
|---|---|---|---|---|
| AT2G20490 | 9.005028999 | 13 | nucleus | nucleus:15133128;nucleus:15496 |
| AT4G02060 | 9.016509477 | 63 | nucleus | nucleus:10751170;nucleus:19357 |
| AT4G18100 | 9.237978348 | 175 | cytosol | cytosol:17934214;nucleus:15496 |
| AT2G36160 | 9.350009325 | 102 | cytosol | plastid:15028209;plasma membra |
| AT1G08880 | 9.426847662 | 33 | nucleus | nucleus:17785529 |
| AT3G44110 | 9.433072352 | 184 | nucleus | plasma membrane:17317660;plas |
| AT3G01280 | 9.637468641 | 47 | mitochondrion | plastid:19326079;mitochondrion |
| AT1G70600 | 9.743739647 | 178 | cytosol | cytosol:17934214 |
| AT1G09850 | 9.8742 | 15 | endoplasmic reticulum | vacuole:15539469 |
| AT4G31880 | 9.882 | 18 | nucleus | plastid:15028209;cytosol:184331 |
| AT2G07734 | 9.8869 | 45 | mitochondrion | |
| AT4G35000 | 9.908838684 | 11 | peroxisome | mitochondrion:15276431;vacuole |
| AT1G28290 | 9.9853 | 15 | extracellular | plasma membrane:17885091 |
| AT3G26020 | 10.27971093 | 18 | nucleus | |
| AT4G20980 | 10.34414987 | 27 | cytosol | |
| AT1G64330 | 10.38389251 | 96 | vacuole | vacuole:15539469;plasma membr |
| AT2G16640 | 10.503 | 5 | nucleus | |
| AT3G02900 | 10.504 | 7 | plastid | |
| AT5G15750 | 10.75974564 | 10 | cytosol | cytosol:17934214 |
| AT5G11670 | 10.94762655 | 55 | cytosol | plasma membrane:17317660;plas |
| AT5G13450 | 11.10120105 | 3 | mitochondrion | plasma membrane:17317660;mitc |
| AT2G09990 | 11.2058116 | 148 | cytosol | plastid:15028209;cytosol:179342 |
| AT5G39740 | 11.287 | 742 | cytosol | plasma membrane:17317660;vacu |
| AT5G15090 | 11.64312537 | 264 | mitochondrion | plastid:19326079;mitochondrion |
| AT3G47370 | 11.67868462 | 35 | cytosol | cytosol:17934214;nucleus:15496 |
| AT5G13650 | 11.783 | 27 | plastid | plastid:18633119;plastid:184314 |
| AT5G54750 | 12.65694613 | 6 | golgi | |
| AT5G17770 | 12.93343163 | 4 | endoplasmic reticulum | plasma membrane:17317660;endo |
| AT3G55460 | 13.11544212 | 15 | nucleus | nucleus:15133128 |
| AT5G65750 | 13.16985158 | 77 | mitochondrion | mitochondrion:14671022 |

| | | | | |
|---|---|---|---|---|
| AT4G25630 | 13.38293943 | 111 | nucleus | nucleus:10829025;nucleus:15496 |
| AT2G38550 | 13.3899281 | 29 | plastid | plastid:12938931;plastid:166189[illegible] |
| AT5G10840 | 13.833 | 179 | golgi | golgi:16618929;plasma membran[illegible] |
| AT4G28470 | 15.046 | 274 | cytosol,nucleus | |
| AT5G42150 | 15.44210754 | 22 | mitochondrion | mitochondrion:14671022 |
| AT5G13780 | 16.243 | 15 | cytosol | |
| AT1G54290 | 20.724 | 4 | cytosol | |
| AT4G32470 | 20.80699385 | 4 | mitochondrion | mitochondrion:15276431;plastid |
| AT3G18524 | 24.1021933 | 4 | nucleus | plasma membrane:14506206 |
| AT4G25730 | 28.297 | 40 | nucleus | |

| |
|---|
| tid:18431481 |
| [illegible]452 |
| [illegible]452 |
| 19;plastid:12766230;plastid:12938931 |
| 13;plastid:12938931;nucleus:15496452 |
| 81 |

ma membrane:17644812
481;mitochondrion:15276431;mitochondrion:12681508;nucleus:15496452

membrane:17644812
ma membrane:16618929
81

eus:15496452

ochondrion:14671022;nucleus:15496452
ane:17151019;plastid:12766230;vacuole:15215502
01

452

929;mitochondrion:14671022;vacuole:16461582;vacuole:15215502;plasma membrane:17151019
ma membrane:7920705;vacuole:15215502

469;plastid:12766230;plasma membrane:15060130;vacuole:8624437

31
ane:15060130
ochondrion:12492832

841;nucleus:15496452

81
496;nucleus:15496452

ane:17151019;plastid:18431481;cytosol:17934214

some:17951448
uole:14760709

ondrion:14671022;nucleus:18433157;cytosol:18433157
ondrion:11743115

57;nucleus:18433157;nucleus:17442570;nucleus:15496452

ne:17317660;vacuole:15215502;plasma membrane:17151019;cytosol:17934214;nucleus:15496452
157

sol:17934214
:16790030
30

ma membrane:16618929
sol:17934214

sol:11598216
81

:14671022;plasma membrane:17151019;mitochondrion:15276431;plastid:16618929

81;plastid:14729914;plastid:15322131

ondrion:15276431

358

157;nucleus:15496452
31;plastid:12938931

·81

membrane:17317660;plastid:18431481;plastid:16473969;plastid:12938931;plasma membrane:16618929

ane:17151019

oplasmic reticulum:16618929

19;plastid:18431481;plastid:12938931

ne:17151019;cytosol:11598216;plastid:18431481;nucleus:15496452
ticulum:16618929;cytosol:17934214;plasma membrane:15060130
skeleton:16603653;mitochondrion:14671022

tid:15028209;cytosol:17934214;nucleus:15496452
eus:14617066;cytosol:11598216;nucleus:15610358;plastid:18431481
ochondrion:15276431

oplasmic reticulum:9159953;cytosol:9159953

eus:15496452
ane:17151019;plasma membrane:17644812
ne:17317660;plastid:18431481;vacuole:15215502;plasma membrane:17151019;cytosol:17934214

157
81

sol:17934214
ticulum:16618929;vacuole:15215502
431481;peroxisome:17376163
:18431481
452
ane:17151019;plastid:18431481;vacuole:16618929

sol:18433157

ondrion:14671022
ane:17151019
452
sol:17934214;nucleus:15496452

eus:15496452

membrane:17151019;endoplasmic reticulum:16618929
81;plastid:12938931

e:17151019;vacuole:14760709
066;plasma membrane:17151019;cytosol:17934214;nucleus:15496452
ane:17151019;vacuole:16461582;plastid:18431481;vacuole:16618929
tid:18431481;plasma membrane:17644812
s:15496452
tid:15028209;cytosol:11598216;nucleus:15496452
uole:15215502;vacuole:16618929
osol:17934214;nucleus:15496452
tid:18431481;cytosol:17934214;nucleus:15496452
469;vacuole:14760709;vacuole:16618929;plasma membrane:17317660;plastid:18431481;plastid:12938931;plastid:12766230

n:15276431;plasma membrane:17151019;endoplasmic reticulum:16618929;plasma membrane:19302169
469;plasma membrane:17151019

30;plastid:18431481;plastid:17098813;mitochondrion:16618929

452
14;plastid:15322131;plastid:18431481;plastid:14729914;plastid:16207701
ane:17151019;plastid:16618929;mitochondrion:16920781;vacuole:15539469

ıole:15215502;vacuole:16618929
tid:18431481;cytosol:17934214
ma membrane:17151019;endoplasmic reticulum:16618929;vacuole:15215502;plastid:15322131
157
ondrion:12492832
ane:15060130

ane:17151019
tid:15028209;cytosol:17934214

)66;plastid:16207701;plastid:18431481;plastid:12938931;nucleus:15496452
eus:14617066;plastid:18431481;plasma membrane:17644812;nucleus:15496452

ma membrane:16506311;vacuole:14760709;vacuole:16618929

81
s:18433157;mitochondrion:11743115;cytosol:18433157
ısol:18433157
ıole:14760709;plastid:18431481

214

ondrion:15276431
oplasmic reticulum:16618929;plasma membrane:17644812
52;plastid:12938931

tid:15028209;cytosol:17934214;vacuole:15215502

452
ane:17151019;vacuole:16618929
oplasmic reticulum:9238019

14;nucleus:15496452

osol:17934214;vacuole:15215502
tid:18431481;plastid:12938931

osol:18433157;nucleus:18433157;plastid:18431481;nucleus:15496452

osol:17934214;nucleus:15496452
tid:15028209;cytosol:17934214;vacuole:15215502;nucleus:15496452

157
81;plastid:15322131
osol:18433157

tid:15028209;cytosol:11598216

l19;plastid:18431481;plasma membrane:17644812
tid:18431481;vacuole:15215502
502

:17934214;vacuole:15539469

ne:15060130;cytosol:18433157
ochondrion:11971146;plasma membrane:17151019

157;cytosol:18433157

sol:17934214;nucleus:15496452
tid:15028209;cytosol:17934214

81
14;vacuole:15215502;nucleus:15496452

ma membrane:17151019;cytosol:18433157;cytosol:17934214;nucleus:15496452

ma membrane:17151019;cytosol:2227432;vacuole:15215502;nucleus:15496452

30;plastid:18431481;mitochondrion:16618929

31
sol:11598216

14
ondrion:15276431;plasma membrane:15060130
951448;plasma membrane:17644812;nucleus:15496452

81

ane:17151019
06;mitochondrion:16618929;plastid:12938931;plastid:16618929;cytosol:18433157

28;nucleus:15496452

ane:17151019;vacuole:16618929

xisome:17951448

59;plastid:16461379;plastid:15322131;nucleus:14617066;plasma membrane:16618929
sol:17934214

tid:15028209;vacuole:15215502;nucleus:15496452

sol:17934214
ne:17151019;cytosol:17934214;vacuole:15215502;nucleus:15496452

eus:18433157;cytosol:18433157
69;vacuole:1311852;vacuole:16618929;mitochondrion:14671022;plastid:18431481;plasma membrane:17644812;cytosol:16210544;plasma membrane:171
ane:17151019;vacuole:16618929
ane:17151019
:15276431;mitochondrion:14671022;plastid:16618929;plasma membrane:17151019;mitochondrion:17137349;mitochondrion:18189341
31;plastid:18431481;plastid:16618929;plastid:10373488;mitochondrion:16618929;plastid:12766230

sol:17934214;nucleus:15496452
157

452
452
452

81;plastid:12938931
469;plasma membrane:17151019

oxisome:17951448;nucleus:15496452
20;mitochondrion:14671022;plastid:12938931;plastid:15659100;plastid:10427773;plastid:16207701

membrane:17151019;mitochondrion:11743115;plastid:16618929

452
sol:17934214

81;plastid:12938931

01
tid:15028209

14;nucleus:15496452
ane:17151019;endoplasmic reticulum:16618929;nucleus:15496452

452
eus:18702672
:16618929;mitochondrion:11718729;mitochondrion:11743114
tid:12938931;vacuole:14760709;plastid:18431481

131
31

eus:18433157;cytosol:18433157

29;plastid:18431481;mitochondrion:16618929

ondrion:11743115;plastid:12938931;plastid:18431481;vacuole:15215502;plasma membrane:17151019;nucleus:15496452
29;plastid:12766230;plastid:18431481;mitochondrion:16618929
ondrion:14730085
sol:17934214

uole:18028262;vacuole:14760709;vacuole:15772667

:16251277

n:15276431;plasma membrane:15060130;mitochondrion:11743115;nucleus:15496452
14;plastid:18431481;vacuole:15215502;nucleus:15496452

:18385124;plastid:18633119;mitochondrion:11743115;plastid:12938931
214
tid:16251277;mitochondrion:16251277

tid:15028209
31;plastid:12177442;plastid:14729914;plastid:16618929;mitochondrion:16618929;plastid:18431481

452
sol:17934214;vacuole:15215502

:14671022;mitochondrion:15276431;plastid:12938931
ciculum:16618929;vacuole:15215502;plasma membrane:17151019;cytosol:17934214;nucleus:15496452
ma membrane:8883393
sol:18433157;nucleus:15496452

ondrion:14671022;plasma membrane:17151019;mitochondrion:15276431

sol:18239134
066;nucleus:15133128;nucleus:15496452

ne:17317660;nucleus:14617066;nucleus:18433157;cytosol:18433157;nucleus:15496452

:18431481;plastid:16618929;plasma membrane:17151019;plastid:12938931;plastid:15908591
157

452
ondrion:11743115;plastid:16618929

ondrion:15276431
sol:17934214

ane:17151019;endoplasmic reticulum:16618929;plastid:15322131
tid:18431481;cytosol:17934214
:14671022;plasma membrane:17644812;nucleus:15496452

452
81;plasma membrane:17644812;nucleus:15496452
eus:18433157;endoplasmic reticulum:16618929
oplasmic reticulum:16618929;nucleus:15496452
membrane:17151019;mitochondrion:11743115;nucleus:15496452
membrane:17151019;plastid:16618929
iole:15539469;nucleus:14617066;endoplasmic reticulum:16618929;nucleus:15496452

452

sol:17934214

oplasmic reticulum:16618929
157

481
:15276431;mitochondrion:14671022;plastid:12938931;plastid:18431481;plasma membrane:17644812;vacuole:15215502;nucleus:15496452
01
tid:16618929;mitochondrion:14730085
14;plastid:9774405;mitochondrion:14671022;plasma membrane:17151019;plastid:18633119;plastid:12766230
:14671022;mitochondrion:17148605;plasma membrane:17151019;mitochondrion:15276431
.387208;plasma membrane:17151019;vacuole:15215502;nucleus:11387208

79;plastid:18431481;plastid:14729914;plastid:15322131
ıole:15215502
14;plastid:12938931

ı:15276431;plasma membrane:17317660;mitochondrion:14671022;plastid:18431481;endoplasmic reticulum:16618929

i:16618929
:15276431;plasma membrane:17151019;nucleus:15496452;mitochondrion:18189341;mitochondrion:14730085

452

:16618929;mitochondrion:16618929

30;plastid:15659100
79;plastid:18431481;plastid:15322131
ıne:17151019;cytosol:17934214;vacuole:15215502;nucleus:15496452

452
199
452
ne:17151019;cytosol:11598216

ma membrane:17644812;nucleus:15496452
:15276431;plastid:18431481;vacuole:15215502;plastid:16618929;mitochondrion:19326079;plasma membrane:17151019;plastid:12938931

57

e:14760709;plastid:18431481;plasma membrane:17151019;plastid:12938931;peroxisome:17951448

ane:17151019

tid:18431481
ochondrion:14671022;mitochondrion:15276431;plastid:18431481
14
uole:15539469;cytosol:17934214;nucleus:15496452
:18385124;mitochondrion:11743115;plastid:12938931;plastid:18431481;plasma membrane:17644812;plastid:16618929;vacuole:15215502;nucleus:15496
452
81

oplasmic reticulum:16618929

452
29;plastid:12766230;plastid:18431481;mitochondrion:16618929
e:17151019

:16618929;plasma membrane:17151019;mitochondrion:18189341;mitochondrion:16618929

51019;plastid:12938931;plasma membrane:16210544
